# Helix Design Studio: designing antimicrobial peptide candidates for membrane environments

**DOI:** 10.64898/2026.10.20.760466

**Authors:** Daniel Conde-Torres, Rebeca García-Fandiño, Ángel Piñeiro

## Abstract

The activity of many antimicrobial peptides (AMPs) depends on how their sequences interact with different membrane environments. This creates a design challenge: improving compatibility with a target environment can also favour interactions under conditions where activity is undesirable. Helix Design Studio is an open-source web application that addresses this challenge by analysing and designing peptide sequences on a prescribed ***α***-helical scaffold. A published environmental scoring framework combines intrinsic helix propensity, residue-pair interactions, electrostatics and hydrophobic exposure to evaluate the assumed helical state in idealized solvents and interfacial environments. A shared, sequence-dependent alignment rule links the amphipathic residue pattern to the interfacial geometry during both analysis and optimization. Users can inspect score contributions, generate candidate families and search for sequences that favour a target environment or balance compatibility across environments. Computational demonstrations show that environmental assumptions and helix orientation affect candidate rankings and sequence selection. Agreement with an independent descriptor implementation and exhaustive enumeration of a tractable sequence space support computational consistency. Downloadable sequences, scores and calculation settings provide reproducible starting points for simulation and experimental characterization. The scores assess model compatibility; they do not predict folding equilibria or establish biological selectivity. Helix Design Studio is freely available at https://helix.simbioslab.com/.

**Scientific Contribution:** Helix Design Studio connects sequence analysis and candidate generation with explicit design objectives across solvent and interfacial environments. An interpretable scoring framework supports diverse peptide families, preferences for a target over competing environments and compromises between conditions; downloadable sequences, scores and calculation settings make the design choices reproducible.

## 1 Introduction

Antimicrobial peptides (AMPs) are widespread components of biological defence and have long provided molecular templates for the development of anti-infective agents[1]. Their activities encompass antibacterial, antifungal and antiviral effects, while many host-defence peptides also participate in the regulation of immune responses[2]. This functional breadth is accompanied by considerable structural and mechanistic diversity. Membrane interactions, effects on intracellular targets and modulation of host responses all contribute to the biological repertoire of these molecules, providing several routes through which peptide sequences can be developed into therapeutic candidates[3].

Within this diversity, membrane-active peptides offer a direct connection between molecular organization and biological function. Their interactions with lipid bilayers depend on the combined effects of charge, hydrophobicity, amphipathicity and conformation, making these properties central to understanding and modifying their activity[4]. Amphipathic *α*-helices provide a well-defined geometrical framework for exploring these relationships: the position of residues along the sequence determines their distribution around the helical surface and the segregation of polar and hydrophobic groups[5]. Sequence changes can therefore reshape the molecular surface presented to a membrane, linking residue selection to the physicochemical basis of peptide–membrane nteractions.

That relationship is also shaped by the membrane environment. Differences between microbial and host membranes influence peptide interactions and contribute to selective activity[6]. The relevance of lipid composition extends beyond infection: experiments with model membranes show that cancer-associated lipid changes can alter peptide binding and membrane permeabilization[7]. Lipid alterations in cancer and cellular senescence also motivate membrane-targeted therapeutic strategies, as discussed in the cancer–aging–infection (CAIn) framework[8]. This environmental dependence also extends to peptide conformation: membrane association can promote *α*-helical folding, with the extent of this response depending on the membrane environment[9]. For peptides whose activity involves a membrane-associated helical state, this coupling suggests an attractive design objective: favouring that conformation at target membranes while limiting its formation in non-target environments. Such behaviour could contribute to selectivity alongside differences in binding and membrane perturbation, making environmental responsiveness a property worth considering explicitly during sequence design.

Translating this biological rationale into a design strategy requires a way to compare the consequences of sequence changes across environments. A substitution that improves compatibility with a target condition may also improve compatibility with a condition in which peptide activity is undesirable. Conversely, some applications may benefit from sequences that retain favourable properties across several conditions. These objectives require different comparisons, but both depend on evaluating candidates under a common set of assumptions. Linking sequence generation to an interpretable environmental score provides a way to express the intended preference during the search and to examine which molecular contributions produce it. The same framework can then be used to assess existing sequences, explore alternatives and identify compromises between design objectives.

Here, we present Helix Design Studio, an open-source web application that integrates these tasks for peptide sequences represented on a prescribed *α*-helical scaffold. The application builds on a previously published environmental scoring framework combining intrinsic helix propensity, residue interactions, electrostatics and hydrophobic exposure[10]. It evaluates the compatibility of the prescribed helical state with idealized solvent and interfacial environments and uses that evaluation to generate candidate families, favour a target over competing environments or explore compromises between conditions. Because sequence changes alter the amphipathic pattern, a sequence-dependent alignment based on the hydrophobic moment is applied during both scoring and optimization. Inspectable scoring contributions and exportable calculation settings connect the resulting candidates to the assumptions that produced them, supporting reproducible selection of sequences for subsequent simulation and experimental investigation of folding, membrane interactions and biological selectivity.

Figure 1 summarizes how this framework connects sequence choices, environmental comparisons and candidate selection.

**Fig. 1:**
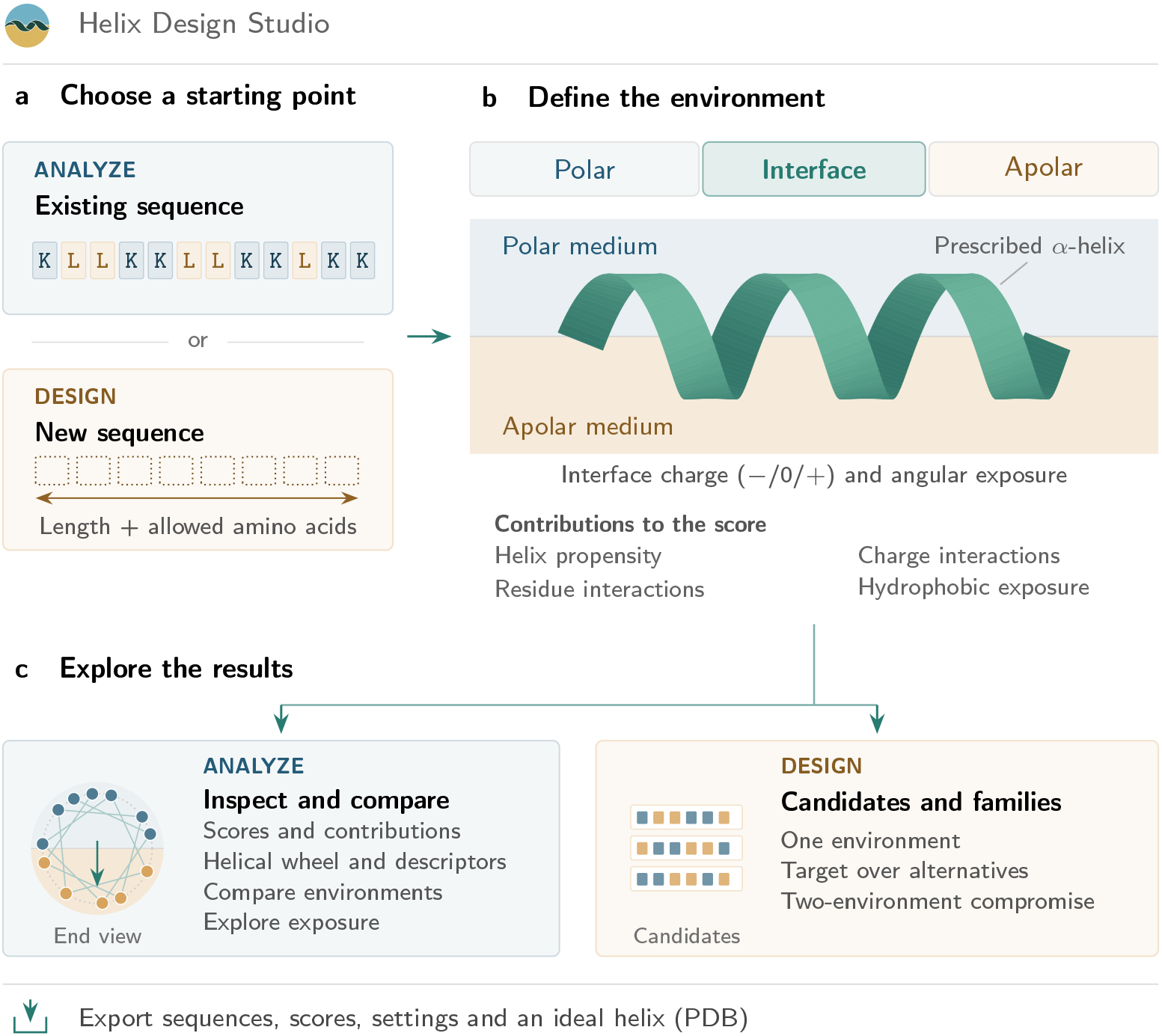
Two routes through Helix Design Studio: sequence analysis and design. (a) Users supply an existing sequence or specify peptide length and allowed amino acids for a design search. (b) A shared scoring model evaluates a prescribed *α*-helical state in a polar medium, an apolar medium or an interface with selected charge and angular exposure. The four contribution groups summarize the model; the interface shown is schematic. (c) Analysis provides scores, their contributions, helical-wheel views and descriptors, environmental comparisons and exposure scans. Design generates individual candidates or families for one environment, target preference or a compromise between environments. The end-on wheel depicts the illustrative input, with charged lysine residues in blue and hydrophobic leucines in gold; its hydrophobic moment points towards the lower, apolar region. Candidate strips illustrate diversity without representing calculated results. Sequences, scores, settings and ideal helical structures can be exported for subsequent studies. These outputs assess model compatibility, not folding or biological activity.

## 2 Implementation

### 2.1 Architecture and distribution

The application has three layers (Fig. 2). A static HTML/CSS/JavaScript interface presents the workflows and renders their outputs. A Python backend validates inputs and coordinates calculations, including progressive delivery of family and specificity results to the browser. In the sequence-analysis workflow, users enter a sequence, choose an alphabet and environment, then inspect summary statistics, individual score contributions and the helical wheel. The Python engine performs sequence encoding, calibration, scoring and classical optimization using NumPy[11]. The scoring core and four numerical parameter tables are distributed with the application. The web workflows run on conventional CPU hardware with the Python runtime and a WebGL-capable browser.

**Fig. 2:**
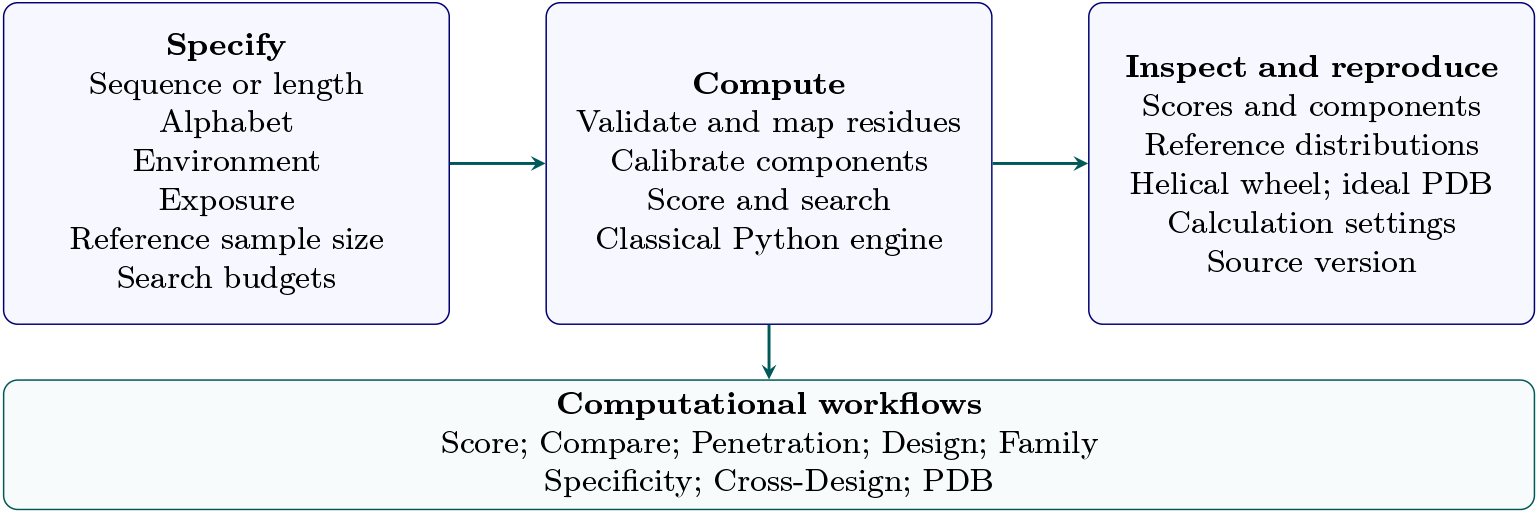
Software architecture and user workflows. Browser inputs, server-side calculations and interactive outputs are connected through a shared scoring and search engine. Calibration, reference statistics, orientation and source version form part of the reproducible calculation. PDB output represents a prescribed ideal helix.

The public repository[12] contains the runtime, pinned dependencies, installation instructions, an optional container definition, tests and parameter-verification tools. Additional file 1 provides the complete source snapshot used in this study. Original project code is released under MIT, while inherited third-party notices remain applicable. The vendored 3Dmol.js 2.4.2 bundle was matched by checksum to its versioned distribution and is accompanied by its licence[13].

The numerical examples, consistency checks and execution-time measurements use Helix Design Studio version 1.0.0rc6, following the protocol in Section 2.7. The application is publicly accessible through its web interface[14]. Downloadable calculation records identify the software version and input settings used to obtain each result.

### 2.2 Browser-based use and result export

The overview offers two entry points: analysis of an existing sequence and design of a new one (Fig. 1). Dedicated panels guide users through each task, while Behind the model explains the model and the interpretation of results. In Check a sequence (the Score workflow), the user enters a sequence, selects an environmental preset and sampling settings, and inspects the score contributions, reference statistics and helical wheel. Compare environments (Compare) evaluates the same sequence in two environments. Design a peptide (Design) and Create a family (Family) generate candidates for a selected environment. Design for selectivity (Specificity) explores target–competitor preferences, while Design for two environments (Cross-Design) seeks compromises between conditions. Explore exposure (Penetration) scans angular exposure, and Build a 3D helix (PDB) displays an idealized helical structure for inspection and export. The workflow identifiers in parentheses are used throughout this article and the evaluation records.

Calculation results can be downloaded as JSON records containing the inputs, scores, calibration and reference settings, and software provenance. Specificity also provides CSV export, and the structural panel allows PDB download. These exports support selection of candidates for subsequent simulation or experimental characterization and preserve the settings needed to reproduce a comparison. The examples in the Results section illustrate these web workflows using a common evaluation protocol.

### 2.3 Inputs, environmental conventions and score interpretation

The web interface accepts a sequence or length, an allowed amino-acid alphabet, an environment and workflow-specific sampling or search settings. Advanced settings expose the calibration sample size, random seed and an optional exact interfacial half-width; the exposure slider provides the primary geometric control. Calculation records preserve these settings and the score weights. The interface constrains calculations to at most 40 residues and bounds sampling and search budgets. The model charge is the sum of fixed side-chain charges: +1 for Lys/Arg, −1 for Asp/Glu and zero for the other standard residues, including His.

Five presets represent homogeneous polar and apolar media and anionic, neutral and cationic interfaces. The interfacial model assigns scaffold positions to two regions using a helical wheel with a 100^◦^ increment per residue. The presets specify idealized scoring conditions through their polarity, charge and exposure parameters, which determine the scoring components and their calibration.

The single-residue helix-propensity contribution averages the individual amino-acid penalties tabulated by Pace and Scholtz[15]. The separate helix-neighbour contribution uses M1 pairing propensities from Nacar[16] together with the M3 and M4 interaction coefficients of Muñoz and Serrano[17] (Section 2.6). The exposure contribution uses the Fauchère–Pliska hydrophobicity scale[18]; hydrophobic residues lower the score in the apolar sector, whereas polar residues lower it in the polar sector. The interfacial charge contribution averages min(*σq_i_,* 0) over polar-facing residues, with division by the total sequence length *L*, where *q_i_* is the side-chain charge of residue *i* and *σ* = −1, 0 or +1 denotes anionic, neutral or cationic interfaces. It therefore rewards opposite-charge complementarity in that sector. The separate residue–residue electrostatic contribution contains the signed product *q_i_q_j_* and distinguishes attraction from repulsion.

Let **s** = (*s*_1_*, . . ., s_L_*) denote the peptide sequence being evaluated, where *L* is its length and *s_i_* is the amino acid at position *i*. Each *s_i_* belongs to the allowed amino-acid alphabet *A*. The symbol *E* denotes the selected solvent or interface parameters. For the complete sequence, the model evaluates six raw contributions *h_a_*(**s**; *ɛ*), with *a* = 1*, . . .,* 6 indexing, in order, hydrophobic exposure, coupling of charge to the interface, local helix propensity, residue-pair interactions, helix-neighbour effects and screened electrostatics[10]. Thus, *a* labels a type of score contribution, and *h_a_* is its value before standardization and weighting.

Each contribution is first calibrated against an ensemble of *N*_cal_ random sequences, denoted 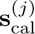 for *j* = 1*, . . . , N*_cal_. These sequences have length *L*, and each residue is drawn independently with equal probability from *A*; sampling is independent of the query composition. For component *a*, *µ_a_* is the mean of the values 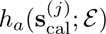 over this ensemble, and *σ_a_* is their population standard deviation, calculated with divisor *N*_cal_. Calibration and query evaluation use the same environment and orientation convention (Section 2.4). The statistics therefore depend on the length, alphabet, environment and orientation convention, with these dependencies implicit in the notation. Their numerical estimates also depend on the calibration sample size and seed.

The score assigned to the query sequence is the weighted sum of its six standardized contributions:

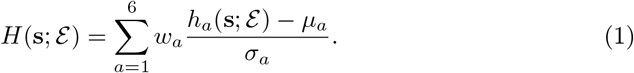

Here *w_a_* is the dimensionless weight specifying the relative contribution of component *a*; the sum runs over components, whereas the index *j* above runs over calibration sequences. The calibration statistics are held fixed when evaluating the query. If a component’s estimated *σ_a_* is at most 10^−12^, its denominator in Eq. 1 is replaced by one to avoid division by zero or numerical amplification. The default weights are 6.0 for exposure, 3.0 for charge coupling, 1.5 for local helix propensity, 0.1 for pair interactions, 1.5 for helix neighbours and 4.0 for electrostatics. The default weights provide a heuristic, physically motivated balance among the standardized contributions, following the original method[10], and yield a dimensionless model score *H*.

A second, independently sampled ensemble contains *N*_ref_ random reference sequences 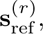 with *r* = 1*, . . ., N*_ref_, drawn using the same length and uniform alphabet rule. Each reference sequence is evaluated with the already calibrated model, using the same weights, environment and orientation convention, to obtain 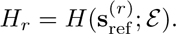 Let *H*_ref_ and *σ_H,_*_ref_ denote the mean and population standard deviation of these total scores. The query’s standardized total score and empirical percentile are then

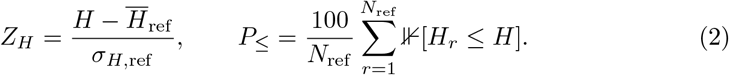

Here *H* abbreviates the query score *H*(**s**; *ɛ*), and ⊮[*H_r_ ≤ H*] equals one when reference sequence *r* has an equal or lower score and zero otherwise. If the reference standard deviation is exactly zero, its denominator is replaced by one. Thus, the component standardization used to construct *H* and the subsequent standardization of *H* to obtain *Z_H_* are distinct operations on separately sampled ensembles. Lower values of *H* and *Z_H_* indicate greater compatibility within the model. The reported percentile includes ties and has a nominal resolution of 100*/N*_ref_ percentage points. A percentile of zero indicates that the query scored below every sequence in the sampled reference ensemble. Returned scores include the alphabet, environment, weights, calibration count/seed, reference count/seed and orientation mode. Changing the alphabet changes the reference problem, even when the same chemical sequence is retained.

The number of random reference sequences, *N*_ref_, is specified for each calculation. Design, Family and Cross-Design use *N*_ref_ = 500, whereas Specificity uses *N*_ref_ = 3000; Score allows the reference count to be selected, with a default of 500. These are differences between named application functions. They are distinct from repeating a calculation with a different random seed. Changing *N*_ref_ can change the estimated reference mean and standard deviation and hence the reported *Z_H_*, even for the same sequence and calibrated score *H*. Changing the seed also changes the sampled calibration and reference sequences. Repeating a calculation with the same software, complete settings and seed reproduces the same samples and score. Exact reproduction therefore requires both ensemble sizes, *N*_cal_ and *N*_ref_, and the seed, together with the remaining recorded settings.

### 2.4 Orientation, exposure scans and structural outputs

Here, orientation refers to rotation of the ideal helix about its long axis relative to the model interface. The helix axis is parallel to the interface. The figures and web interface display the polar medium above and the apolar medium below, with the centre of the apolar sector (0^◦^) pointing downwards in end-on wheel views. The helical-wheel representation defines the transverse hydrophobic-moment vector, ***µ****_H,_*_⊥_ = (*µ_x_, µ_y_*), in the plane perpendicular to that axis. Every interfacial scoring and design workflow uses the same automatic rule: the peptide is rotated about its helix axis until this vector points towards the centre of the apolar sector. The exposure control described below changes the width of the apolar sector, while alignment determines the rotation of the peptide within that geometry.

With the apolar sector centred at 0^◦^, the applied wheel phase is − atan2(*µ_y_, µ_x_*) modulo 360^◦^, where *µ_x_* and *µ_y_* are the components of the transverse hydrophobic moment before rotation in the model’s angular coordinate system. The moment is calculated from the Fauchère–Pliska hydrophobicities at a 100^◦^ helical angle and returned both as a total magnitude and per residue[5, 19]. For a vanishing moment the preset angle is retained. The moment determines the alignment and is also returned as a sequence descriptor. Scores in homogeneous polar and apolar environments are invariant under rotation because all residues occupy the same medium.

During Design, Family, Specificity and Cross-Design searches, the alignment is recalculated for every proposed sequence before its score is used to accept or reject the mutation. For comparisons involving several environments, each proposed sequence is evaluated under the same alignment rule in every interfacial environment. Calibration and random-reference sequences are each aligned by their own moments as well. Their statistics are prepared once for the declared length, alphabet, environment, sample sizes and seeds and reused throughout the search. Thus, the search, reported score, Family ranking, Specificity margins and Cross-Design objectives use the same orientation convention.

This design rule differs from the fixed-orientation sequence searches in the original method article[10]. Sequence-dependent rotation couples the environmental contributions across residue positions, so the classical optimizer reevaluates the complete objective for every proposed mutation. Hydrophobic-moment alignment specifies a geometric convention for placing the sequence relative to the interface. The controlled comparison in the Results section assesses sensitivity to the orientation rule within the model.

The Penetration workflow samples exposure from 0 to 100 in 5% increments. At 0 and 100 it evaluates the homogeneous polar and apolar presets, respectively. Intermediate settings use the linear mapping *α*(*p*) = 40^◦^ + (1.1^◦^)*p*, where *α* is the apolar-sector half-width and *p* is the displayed percentage. This operational mapping defines the range explored by the interfacial control: the sampled settings from 5 to 95 correspond to half-widths of 45.5^◦^ to 144.5^◦^, retaining polar and apolar sectors at each interfacial point. The percentage labels an angular exposure coordinate rather than a calibrated membrane-insertion depth; the corresponding half-width specifies the geometry used in each calculation. All sampled points use the same supplied seed and the same separate random streams for calibration and reference sequences. The minimum reported *Z_H_* identifies the preferred sampled exposure setting under that protocol.

The structural workflow uses PeptideBuilder and Biopython to construct an idealized helix with backbone angles *ϕ* = −57^◦^ and *ψ* = −47^◦^[20, 21]. The resulting PDB is available for inspection in 3Dmol.js and export. The exported PDB represents the prescribed ideal helical geometry, including sidechains.

### 2.5 Search objectives and candidate collections

Single-environment design uses simulated annealing with single-residue substitutions, a linearly decreasing temperature and a floor of 10^−4^. Multiple restarts explore independent trajectories. Family generation sequentially searches for candidates while adding a soft penalty for similarity to previously accepted sequences. The returned collection is characterized by its observed pairwise Hamming distances, which quantify the diversity achieved under this soft penalty.

For target environment *t* and selected off-target environments *k*, *H_t_* and *H_k_* are model scores standardized with the calibration statistics of their respective environments. The specificity objective is

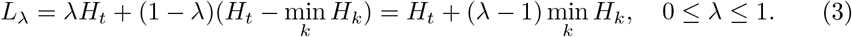

At *λ* = 0, minimization favours separation from the most favourable competitor; at *λ* = 1, it reduces to target-only optimization. A positive margin *m* = min*_k_ H_k_ − H_t_* indicates a lower target score than every selected competitor within the specified model. The penalty used to encourage collection diversity is additional to the unpenalized objective reported with each candidate.

For two environments, the compromise objective is

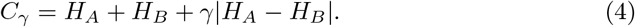

It balances the combined score with their disparity. At *γ* = 1 it equals 2 max(*H_A_, H_B_*). Larger weights can favour reducing imbalance even by worsening the better-scoring environment. Displaying both component scores is therefore essential to interpretation. Here *H_A_* and *H_B_* likewise use the calibration statistics of environments A and B, respectively. Both objectives express preferences relative to the corresponding environmental backgrounds on a dimensionless model scale; their differences are not membrane-transfer free energies.

### 2.6 Reproducibility and parameter provenance

The model uses four 20 *×* 20 parameter matrices for the residue-pair interaction and helix-neighbour contributions to *H*. The Miyazawa–Jernigan (MJ) matrix supplies the residue-pair coefficients *e_ij_* from the upper triangle and diagonal of Table 3 in Miyazawa and Jernigan (1996)[22]; these entries are symmetrized when loaded. For the helix-neighbour contribution, M1 uses the pairing propensities in Figure 3 of Nacar 2020)[16], applied in the model at sequence separation *k* = 1. M3 and M4 use the side-chain interaction coefficients in Table 2 of Muñoz and Serrano (1994)[17] for separations *k* = 3 and *k* = 4, respectively. Each matrix entry specifies a coefficient for a pair of amino-acid identities.

**Fig. 3:**
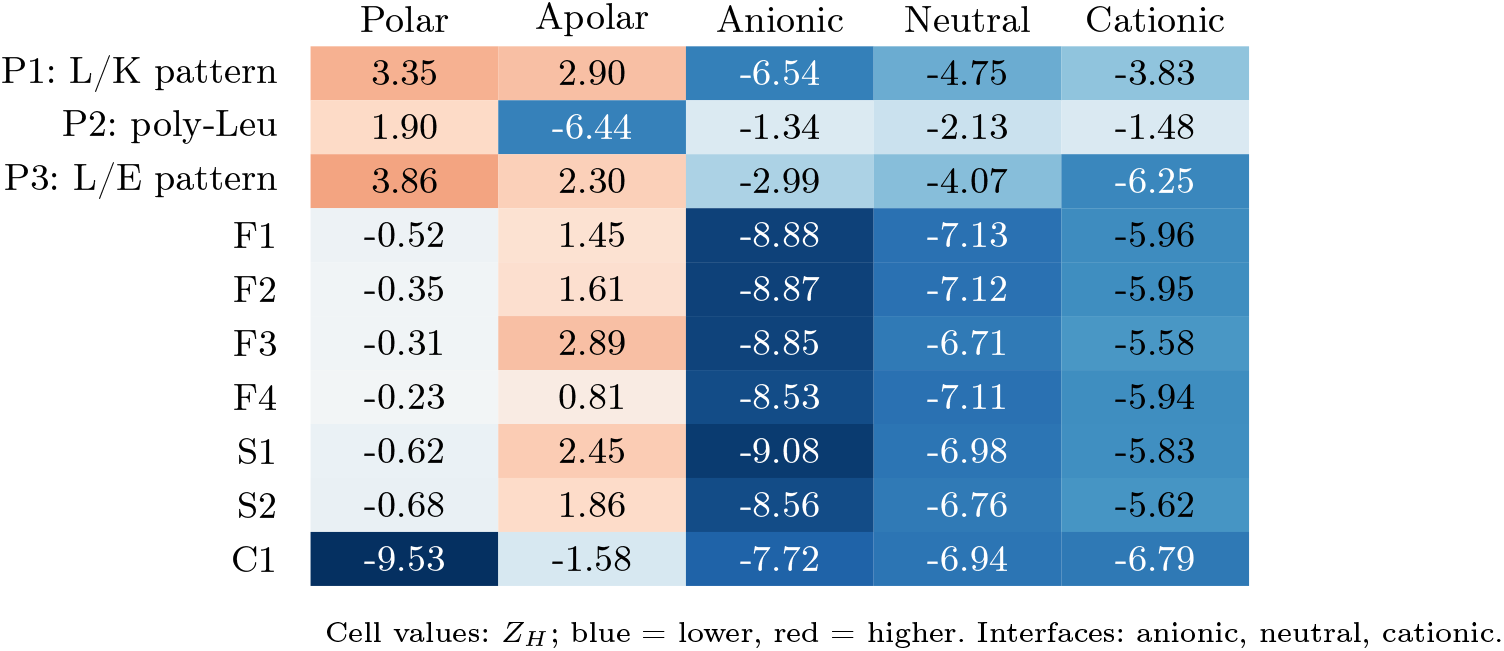
Environmental profiles under a common evaluation protocol. P1–P3 are predefined model sequences; F1–F4, S1–S2 and C1 are candidates from Create a family, Design for selectivity and Design for two environments, respectively (Table 2). All sequences use the same 20-residue alphabet, 2,000 calibration sequences, 1,000 reference sequences and seed 42. Each interfacial query, calibration sequence and reference sequence is aligned to its own hydrophobic moment. All scoring and search evaluations use the same sequence-dependent alignment rule; this common analysis can use a different reference sample count from the native workflow.

The rows and columns of each matrix are labelled by amino-acid identity. For example, an alanine–lysine pair uses the coefficient in the alanine row and lysine column, identified by their one-letter labels A and K. When loading a table, the software matches these labels to the selected amino-acid alphabet and reorders or selects rows and columns accordingly. This preserves the coefficient assigned to each amino-acid pair even if the alphabet is reordered or restricted to a subset. The loaders also check matrix dimensions, unique amino-acid labels and finite numerical values. Calibration caches are versioned and include the parameter-table digest and scoring convention, so statistics are associated with the numerical model used to obtain them. Tests compare raw scores across alphabet permutations and subsets. These checks distinguish raw-score invariance from the sampling variation of standardized scores; changing the alphabet order can change the sampled sequences.

For the helix-neighbour matrices, rows identify the residue at position *i*, and columns identify the residue at position *i* + *k*. The raw neighbour contribution is

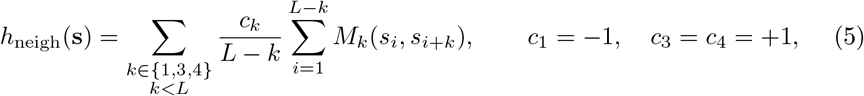

where *k* is the separation in sequence positions and only separations smaller than the peptide length *L* contribute. Higher Nacar pairing propensities thereby lower the score, while the energetic signs of M3 and M4 retain their source interpretation. The three averaged contributions have fixed relative coefficients −1, +1, +1 and are calibrated jointly as one empirical descriptor. This is a dimensionless comparative scoring construction; the internal combination of propensity ratios and tabulated energy coefficients defines its empirical scale. Miyazawa–Jernigan coefficients are summed over sequence separations one through four and calibrated as a separate component.

Random sampling is controlled by a common integer seed, recorded with the calculation and available in Advanced settings; its default is 42. Component calibration uses this seed directly. Reference sampling and sequence searches use separate pseudorandom-number generators initialized reproducibly through NumPy SeedSequence. Reference initialization combines the seed with task identifier 1; a single search combines it with task identifier 2. Family and Specificity additionally include a search-attempt index, starting at zero, so successive attempts use distinct random streams. The exact initialization protocol is preserved in the downloaded calculation metadata.

Search and score calculations share their declared calibration and reference settings. Reconstructing Eqs. 3 and 4 uses the returned aligned model scores for the relevant environments. The reproducibility metadata record the complete input settings, software version, parameter-set checksum, seed protocol and runtime versions. Interrupted Family and Specificity calculations retain fully evaluated candidates in downloadable JSON, with their provenance and an explicit partial-result marker.

The Fauchère–Pliska values for Ser and Thr are −0.04 and 0.26, respectively, in both the exposure score and the hydrophobic-moment descriptor, matching the primary scale and its HELIQUEST transcription[18, 19]. The numerical examples use the dentified software revision and the orientation and sampling protocols described here. M1 reproduces the helical pairing odds ratios in Figure 3 of Nacar (2020)[16]. The published matrix describes hydrogen-bonded pairs at positions *n, n* + 4, with the earlier carbonyl acceptor in the columns and the later amide donor in the rows. The software transposes that table to preserve ordered sequence pairs. All 400 stored coefficients agree with the published values at three-decimal precision. Their residue order is A, R, N, D, C, Q, E, G, H, I, L, K, M, F, P, S, T, W, Y, V. The score subtracts their nearest-neighbour average (Eq. 5), so that greater enrichment of a pair in the source helices gives a more favourable contribution. The published zeros yield zero in this linear contribution. Reusing globular-protein hydrogen-bonded *n, n* + 4 pairing statistics at *i, i* + 1 remains an empirical model assumption; these ratios describe enrichment relative to the source background.

M3 and M4 reproduce the *i, i* + 3 and *i, i* + 4 side-chain interaction coefficients, respectively, in Table 2 of Muñoz and Serrano (1994)[17]. The source reports contributions in kcal mol^−1^ for 18 amino acids. Its row and column orders differ for phenylalanine and tyrosine; both axes are mapped independently by residue identity. All 324 published entries of each matrix agree with the distributed files. The software extends these tables to 20 *×* 20 by assigning zero to pairs involving cysteine or proline (76 entries per matrix). The zero coefficients are model defaults for residues absent from the source tables. M3 and M4 use the residue order A, G, S, T, N, D, Q, E, H, K, R, Y, F, W, V, I, M, L, C, P.

Standardization against the calibration ensemble expresses the combined empirical neighbour contribution on a dimensionless comparative scale. The source distribution includes independently labelled primary-source transcriptions and an executable comparison. Additional file 2 supplies the complete runtime tables, SHA-256 checksums and source mappings. These records identify the numerical dataset used in every example.

### 2.7 Evaluation protocol

The workflow examples were computed with the Helix Design Studio calculation engine, using the scoring and design settings described below. The evaluated software revision is preserved in Additional file 1. Complete input settings and calculation outputs, sequence identifiers, source hashes, package versions, calibration statistics and plotting inputs are included in the evaluation package. The predefined model sequences P1–P3 (Table 2) were selected before calculating the reported examples to represent contrasting charge and hydrophobicity patterns.

Common rescoring uses the same 20-residue alphabet, 2,000 calibration sequences, 1,000 reference sequences and seed 42. The Family example specifies 4 candidates for the anionic interface, length 20, 3,000 steps per restart and 2 restarts per candidate search, with at most 16 candidate searches. The Specificity example generates 2 candidates for the anionic interface against the neutral interface and homogeneous polar medium, with *λ* = 0.5 and 3,000 steps. Its total budget of 4 restarts is shared across attempts, including unsuccessful attempts. Compromise design uses the anionic interface and homogeneous polar medium with *γ* = 0.5, 3,000 steps, 2 restarts and seed 42. Complete input records also preserve the remaining calculation settings and defaults.

All three design examples use 2,000 calibration sequences, seed 42 and the ordered alphabet ARNDCQEGHILKMFPSTWYV. Every interfacial environment has an apolar-sector half-width of 90^◦^. To reproduce these settings in the browser, set Allowed residues to ARNDCQEGHILKMFPSTWYV for each calculation, set Calibration sample size to 2,000 and Random seed to 42 in Advanced settings, and set Exact apolar-sector half-width to 90^◦^ for each interface. Specificity uses 2,000 calibration sequences in this experiment; its application default is 10,000. Homogeneous environments retain their preset definitions. Table 1 distinguishes the search settings from the subsequent common evaluation.

**Table 1:** Effective sampling and search settings for the illustrated design examples. All sequences have 20 residues. Calibration and final reference sequences are independently sampled.

| Calculation | Calibration sequences | Reference sequences | Steps per restart | Restarts | Candidates requested |
| --- | --- | --- | --- | --- | --- |
| Family | 2,000 | 500 | 3,000 | 2 | 4 |
| Specificity | 2,000 | 3,000 | 3,000 | 4 | 2 |
| Compromise | 2,000 | 500 | 3,000 | 2 | 1 |
| Common evaluation | 2,000 | 1,000 | — | — | — |

Family uses the stated restart budget for each attempted search, up to a maximum of 16 candidate searches, stopping when the requested family is complete. Specificity shares four total restarts across attempts. Each generated sequence is evaluated in all five environments under the common protocol. The model score $H$ uses the same calibration in the native result and the common evaluation; $Z_H$ is referenced to the ensemble size specified in each row.

The exact-search check enumerates 4,096 sequences of length 6 over A, E, K and L. Enumeration and 5 annealing runs use the same sequence-aligned anionic-interface Hamiltonian, calibrated with 2,000 sequences and seed 42. Search seeds are 42, 43, 44, 45 and 46, with 1,000 steps and 2 restarts. A recovered minimum requires an absolute score gap of at most 1 *×* 10^−9^. The enumeration provides an exact reference for this finite optimizer–objective problem.

Sampling sensitivity uses calibration/reference-count pairs 500/500, 500/2000, 2000/500 and 2000/2000, with seeds 42, 43 and 44. The exposure scan contains 21 sampled settings, all using seed 42. Runtime measurements use lengths 8, 16 and 24 and seeds 42, 43 and 44 with sequential Score and Design calculations. Each operation first calculates the component-calibration means and standard deviations from 2,000 random sequences, saves these statistics and is then repeated immediately with the same settings using the saved calibration. Both measurements include evaluation of the random reference ensemble and, for Design, the sequence search. Design uses 3,000 steps per restart and 2 restarts. Elapsed time covers input handling and computation after application initialization. Minimum-to-maximum ranges summarize the observed variation across the three seeds.

### 2.8 Use of AI-assisted tools

ChatGPT/Codex (OpenAI) and Claude (Anthropic) assisted with manuscript drafting and editing, software and analysis-code development, computational verification, and preparation of figures and reproducibility materials. The authors reviewed and edited the resulting content and retain full responsibility for the accuracy, integrity and interpretation of the work.

## 3 Results and discussion

### 3.1 Execution, consistency and tractable optimization

Consistency checks covered input validation, calibrated scores, reconstruction of the design objectives and preservation of calculation settings in downloaded results. The evaluated examples returned finite numerical values. All 18 paired calculations with identical settings produced identical computational results with empty and populated calibration caches.

Exhaustive enumeration of 4,096 sequences in the 4-residue-alphabet, length-6 problem found a minimum of *H* = −24.555206. The unique minimizing sequence was `LLKKLL`. Simulated annealing recovered the minimum in 5 of 5 trials. The largest observed absolute optimality gap was 0. Every enumerated score and search trial is retained in the evaluation package. The enumerated minimum provides an exact reference for this finite optimizer–objective problem.

### 3.2 Shared descriptor calculation

The hydrophobic moment per residue returned by Helix Design Studio, 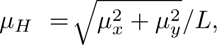 was compared with modlAMP 4.3.2 for the same 6 sequences. Both calculations used the Fauchère–Pliska scale, the full 20-residue sequence and a 100^◦^ helical angle. The comparison included P1, P2 and P3 from Table 2 and three additional sequences: 20 serines, 20 threonines and 10 alternating Ser–Thr pairs. These additional sequences check the assignment of the serine and threonine hydrophobicity values. The hydrophobic moments per residue for P1, P2 and P3 were 0.861239, 0.109274 and 0.747505, respectively. Across all 6 sequences, the largest absolute difference in hydrophobic moment per residue was 1.11 *×* 10^−16^. A separate check compared the mean of the hydrophobicity values used by the scoring model with the mean calculated by modlAMP; its maximum absolute difference was 1.11 *×* 10^−16^. The values agree within floating-point rounding precision; complete sequences, descriptor values and calculation settings are provided in Additional file 2. HELIQUEST documentation supplies the reference descriptor and charge conventions. modlAMP reports fractional, pH-dependent charges that include terminal-group ionization; Helix Design Studio reports the integer side-chain charge defined in Section 2.3. Agreement assesses mplementation consistency under the shared descriptor definitions.

**Table 2:**
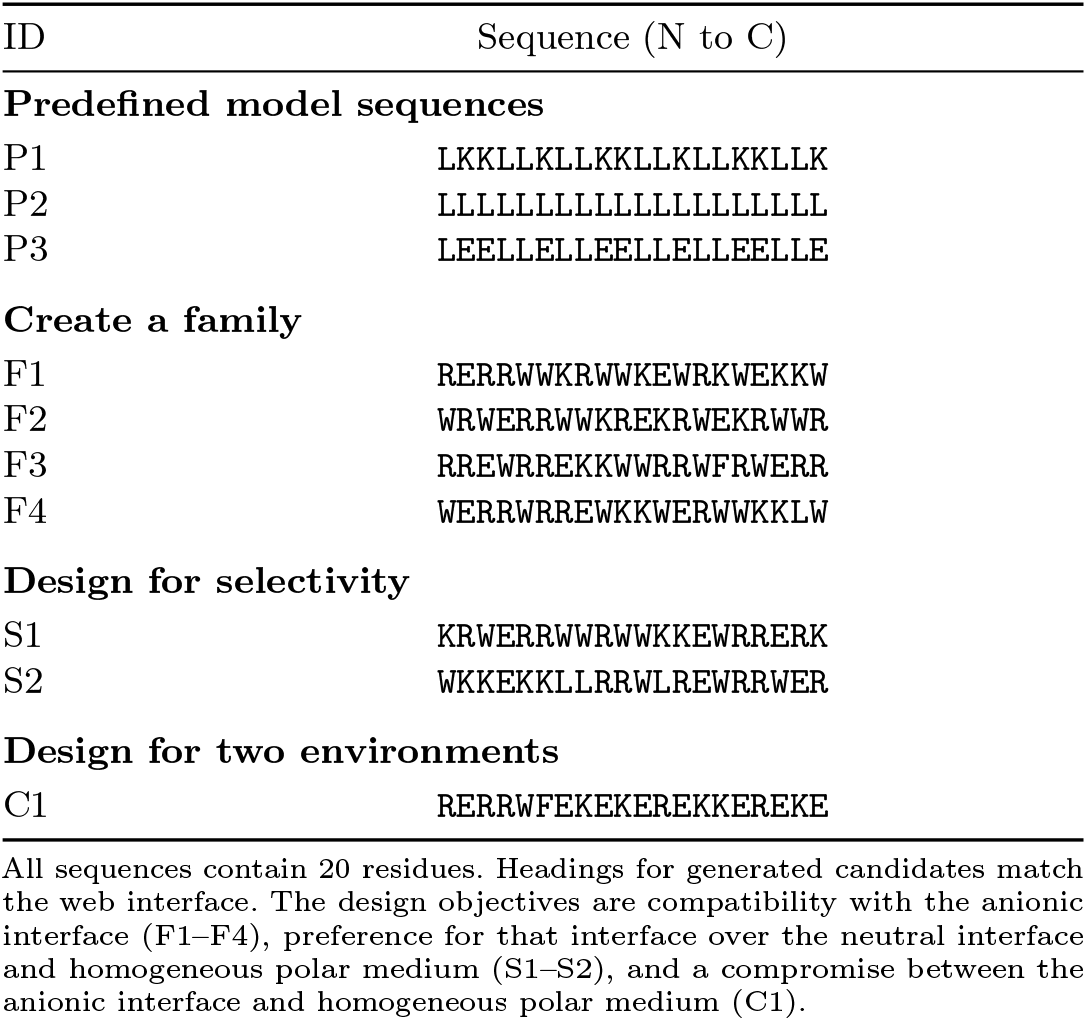
Predefined model sequences and candidates generated by the illustrated application workflows.

### 3.3 Sequences and environmental profiles

The examples comprise three predefined model sequences and seven computationally generated candidates. P1 and P3 combine leucine with positively charged lysine or negatively charged glutamate, respectively, whereas P2 consists entirely of leucine. These contrasting inputs illustrate the response of the environmental scoring model. The remaining sequences demonstrate three candidate-generation workflows, identified in Table 2 by their interface labels.

Generated examples are included regardless of their environmental ranking. Figure 3 applies the same post-search protocol to all sequences, allowing target and alternative environments to be inspected together. P1 scored *Z_H_* = −6.536 at the anionic interface, −4.752 at the neutral interface and −3.829 at the cationic interface. The corresponding anionic and cationic scores of P3 were −2.986 and −6.251. Poly-leucine attained its lowest score in the homogeneous apolar preset (*Z_H_* = −6.443). These observations describe the model’s response to the specified sequences and environments.

### 3.4 Candidate diversity and specificity

The family workflow returned 4 distinct candidates. Their 6 pairwise Hamming distances ranged from 10 to 20 substitutions, with median 18 (Fig. 4a). The observed distances quantify diversity achieved under the soft similarity penalty. Aligned family scores ranged from −78.453 to −75.347.

**Fig. 4:**
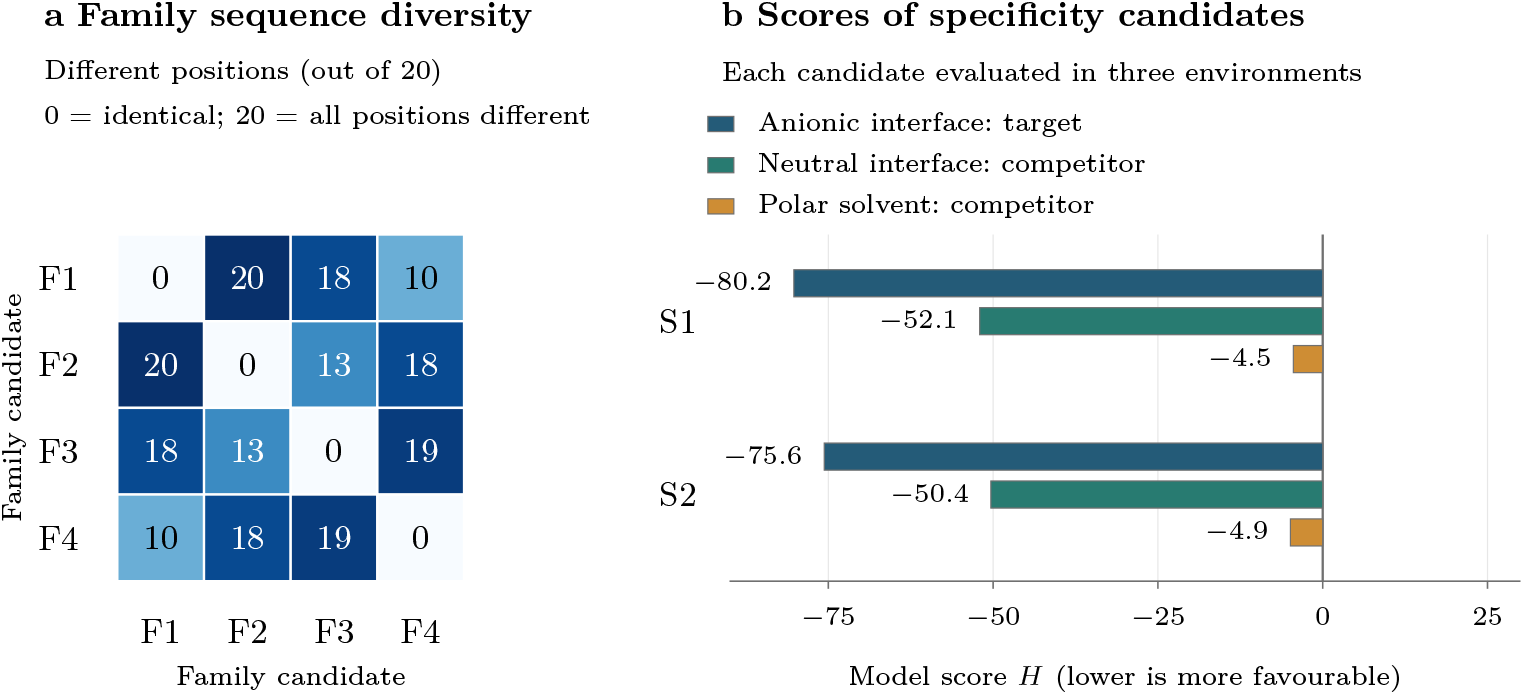
Sequence diversity and environmental scores. (a) Each cell gives the number of amino-acid positions that differ between two 20-residue family members, F1–F4; darker cells indicate more differences. For example, F1 and F2 differ at 20 of 20 positions. (b) Horizontal bars show the dimensionless model score *H* of each specificity candidate in the anionic target interface, neutral competing interface and homogeneous polar competitor. Values are printed at the bar ends. Lower *H* indicates greater compatibility with the specified model environment. Candidate identifiers refer to Table 2; specificity searches use *λ* = 0.5 (Eq. 3).

The specificity calculation returned 2 distinct candidates. Figure 4b compares the aligned model score *H* of each candidate across the target and competing environments. These scores are used in the specificity search objective. In candidate order, target-versus-lowest-competitor score margins were 28.191 and 25.275, and objectives were −54.217 and −50.453. The largest absolute discrepancy between reconstructed and recorded objectives was 0, within the declared 10^−8^ check. Score margins and differences of *Z_H_* values are distinct quantities because the environments have separate background distributions. Search and reported scores use the same per-sequence alignment, calibration and reference convention. Common rescoring in Fig. 3 additionally uses a shared reference-ensemble size; it is a separate analysis of the generated sequences.

The compromise example had aligned model scores *H_A_* = −68.224 and *H_B_* = −68.062 and an objective of −136.205. Reconstruction of Eq. 4 gave an absolute discrepancy of 0. The combined model score sums these same aligned scores; the compromise objective additionally includes their weighted absolute difference. The common profile is included as C1. C1 illustrates compromise design for the specified environment pair and weight.

### 3.5 Exposure control and sampling dependence

For P1, a minimum sampled exposure-scan score occurred at 55%, with *Z_H_* = −7.148 Fig. 5a). This interfacial setting has an angular half-width of 100.5^◦^. The full 21-point profile is retained. Every sampled point uses the same supplied seed. Repeated mask assignments, geometry-dependent calibration and the special homogeneous endpoints affect interpretation. The minimum identifies the preferred sampled angular-mask setting under this protocol.

**Fig. 5:**
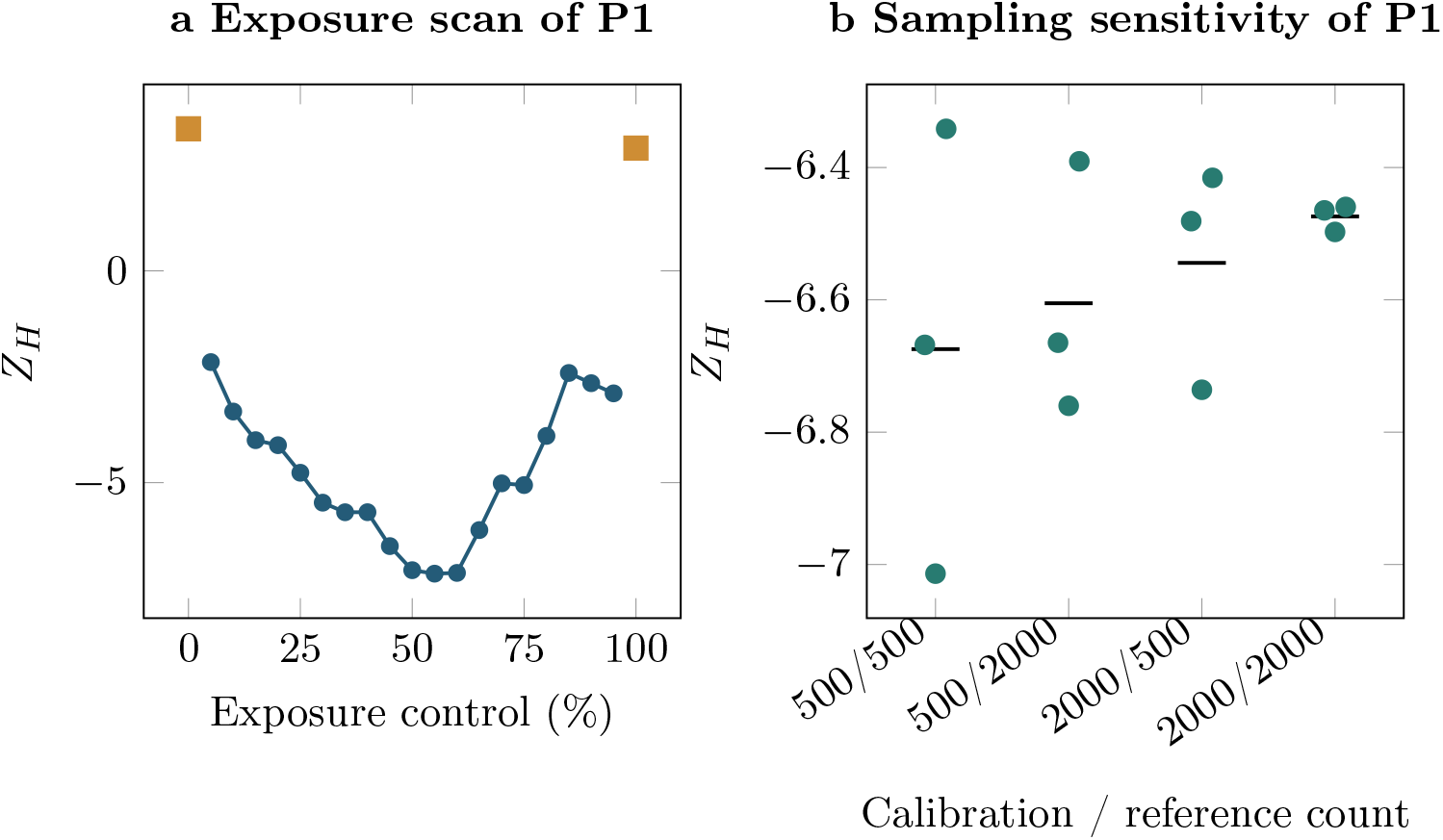
Exposure and finite-sampling sensitivity. (a) The 21-point exposure scan for P1. Gold squares are homogeneous endpoints; intermediate points are interfacial angular masks. Every sampled point uses the same supplied seed and the same separate calibration/reference random streams. (b) Scores of P1 at the default anionic interface for 3 seeds at each sample-count pair. Points show individual runs; horizontal segments show their arithmetic means.

Across 12 sampling-sensitivity evaluations, P1 at the default anionic interface produced *Z_H_* values between -7.014 and -6.342 (Fig. 5b). The replicate seeds vary component calibration and the final reference distribution while keeping the sequence fixed. The observed range characterizes finite-sampling variation for this input.

### 3.6 Measured computational cost

Figure 6 shows the computation time required to evaluate one sequence or design one candidate at lengths 8, 16 and 24 residues. Each operation was measured with the component calibration calculated as part of the operation and then with the same calibration reused. The saved quantities are the means and standard deviations of the six raw score components (Eq. 1); they can be reused when the calibration settings match. In both conditions, the random reference ensemble is evaluated and Design performs the sequence search. At 24 residues, evaluation took 114.7 ms with calibration computed and 46.0 ms with calibration reused; the corresponding design times were 372.4 and 293.8 ms. Both conditions produced identical scientific outputs. The measurements cover input handling and computation after application initialization. The recorded hardware was Apple M3 Max with 16 logical CPUs and 48 GiB RAM (macOS-26.6.2; Python 3.12.8), and calculations ran sequentially. Individual timings and package versions are preserved in the evaluation data.

**Fig. 6:**
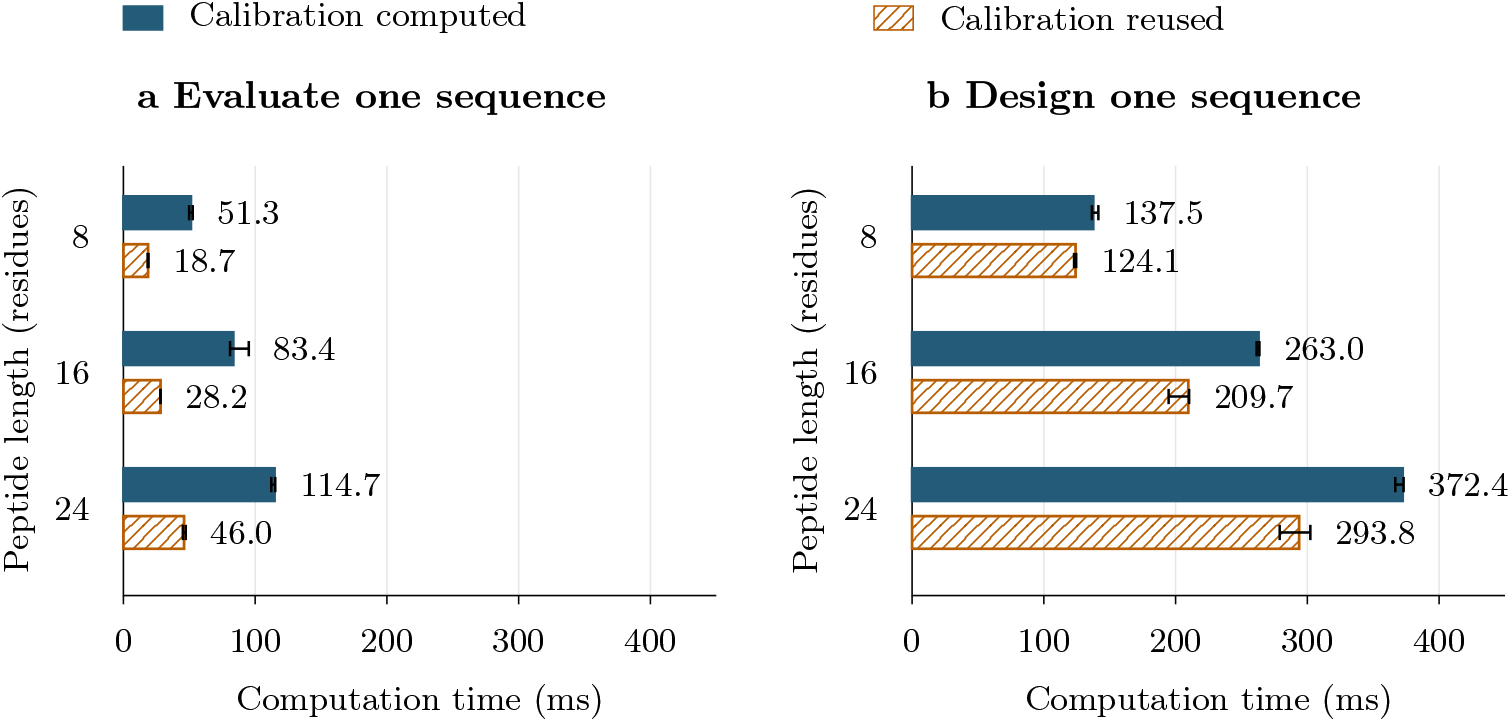
Computation times for sequence evaluation and design. (a) Evaluation of one supplied sequence. (b) Search for one candidate sequence. Blue filled bars include calculation of the component-calibration means and standard deviations; orange hatched bars reuse these statistics with identical calculation settings. Both conditions evaluate the random reference ensemble and, in Design, perform the sequence search. Bar lengths and printed values are medians in milliseconds; whiskers span the observed minimum and maximum across 3 runs seeds 42, 43 and 44). Both panels use the same linear time scale. Elapsed time covers input handling and computation after application initialization. The study used Apple M3 Max with 16 logical CPUs and 48 GiB RAM (macOS-26.6.2; Python 3.12.8); Design used 3,000 steps per restart and 2 restarts.

### 3.7 Updated interaction parameters and their effects

The present software updates the helix-neighbour tables relative to those in the initial implementation associated with our published method[10]. Checking the transcribed tables against their primary sources identified discrepancies in M1 and two interchanged values in M4. The updated M1 reproduces the helical pairing propensities in Figure 3 of Nacar (2020)[16], with its axes mapped to the software’s ordered residue pairs. M3 and M4 use the interaction coefficients in Table 2 of Muñoz and Serrano (1994)[17]; M3 is unchanged, and M4 corrects the F–H and Y–H coefficients to −0.800 and −0.850 kcal mol^−1^, respectively. The source mappings and the retained cysteine/proline zero defaults are described in Section 2.6.

To measure the effect of these updates, we compared the initial and source-verified tables using the same current calculation engine, orientation rule, weights and random streams. Only the M1 and M4 tables were varied, with the negative M1 coefficient in Eq. 5 held fixed in every case. Each parameter set was recalibrated with the same sampled sequences. Twelve reference sequences were evaluated in all five environments, giving 60 paired *Z_H_* values, with the full amino-acid alphabet, 1,000 calibration sequences, 500 reference sequences and seed 42. Updating M1 alone, M4 alone and both tables separated their contributions to the observed differences.

Table 3 shows that M1 accounts for most of the change in standardized scores. With both tables updated, the median absolute change in *Z_H_* was 0.1084 and the maximum was 0.2448. The Spearman correlation across the 60 paired sequence–environment scores was 0.9953, indicating that their overall ordering was largely preserved in this benchmark.

**Table 3:**
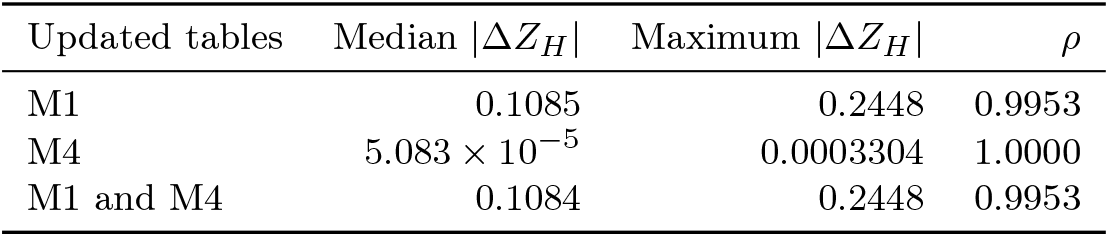
Effect of updating the helix-neighbour tables on standardized scores. Absolute differences are measured relative to the initial tables. Each row summarizes the same 60 paired sequence–environment scores; *ρ* is their Spearman rank correlation.

| Updated tables | Median $\vert \Delta Z_H\vert $ | Maximum $\vert \Delta Z_H\vert $ | $\rho$ |
| --- | --- | --- | --- |
| M1 | 0.1085 | 0.2448 | 0.9953 |
| M4 | $5.083 \times 10^{-5}$ | 0.0003304 | 1.0000 |
| M1 and M4 | 0.1084 | 0.2448 | 0.9953 |

Sequence design was evaluated in six paired searches of length 18, using the anionic interface and homogeneous polar medium, seeds 1–3, 4,000 steps and two restarts. Updating both tables changed 6 of the 6 returned sequences, with differences at 60 of 108 residue positions (55.6%). These results show that the parameter update can substantially change the candidate sequences selected by optimization, even when the benchmark scores remain highly correlated.

The M1 coefficient was also changed from +1 to −1 (Eq. 5), so that higher helical pairing propensities lower the minimized score. A matched comparison varying only this coefficient, with both conditions recalibrated using the same sampled sequences, gave a median absolute change in *Z_H_* of 0.3628 (maximum 3.4388) across the 60 scores and changed all six designs (76 of 108 residue positions). Additional file 2 provides the paired scores, sequences and complete protocols for both comparisons. All results reported here use the source-verified tables and the corrected scoring convention.

### 3.8 Effect of helix orientation on the designed sequences

For interfacial scoring, the peptide is rotated about its helical axis to orient the transverse component of its hydrophobic moment towards the hydrophobic phase Fig. 7). This component lies in the plane perpendicular to the helix axis. We examined how applying this rotation during sequence design affects the candidates selected. Two search strategies were compared. The fixed-orientation strategy evaluates every proposed sequence at the same rotation about the helix axis, following the search convention of the original method[10]. The aligned strategy recalculates the rotation for each proposed sequence to orient its transverse hydrophobic moment towards the apolar sector, as used by the present application. Applying this orientation rule throughout the search can change which mutations are accepted. Both strategies use the same current engine, source-verified parameter tables, weights and search budgets.

**Fig. 7:**
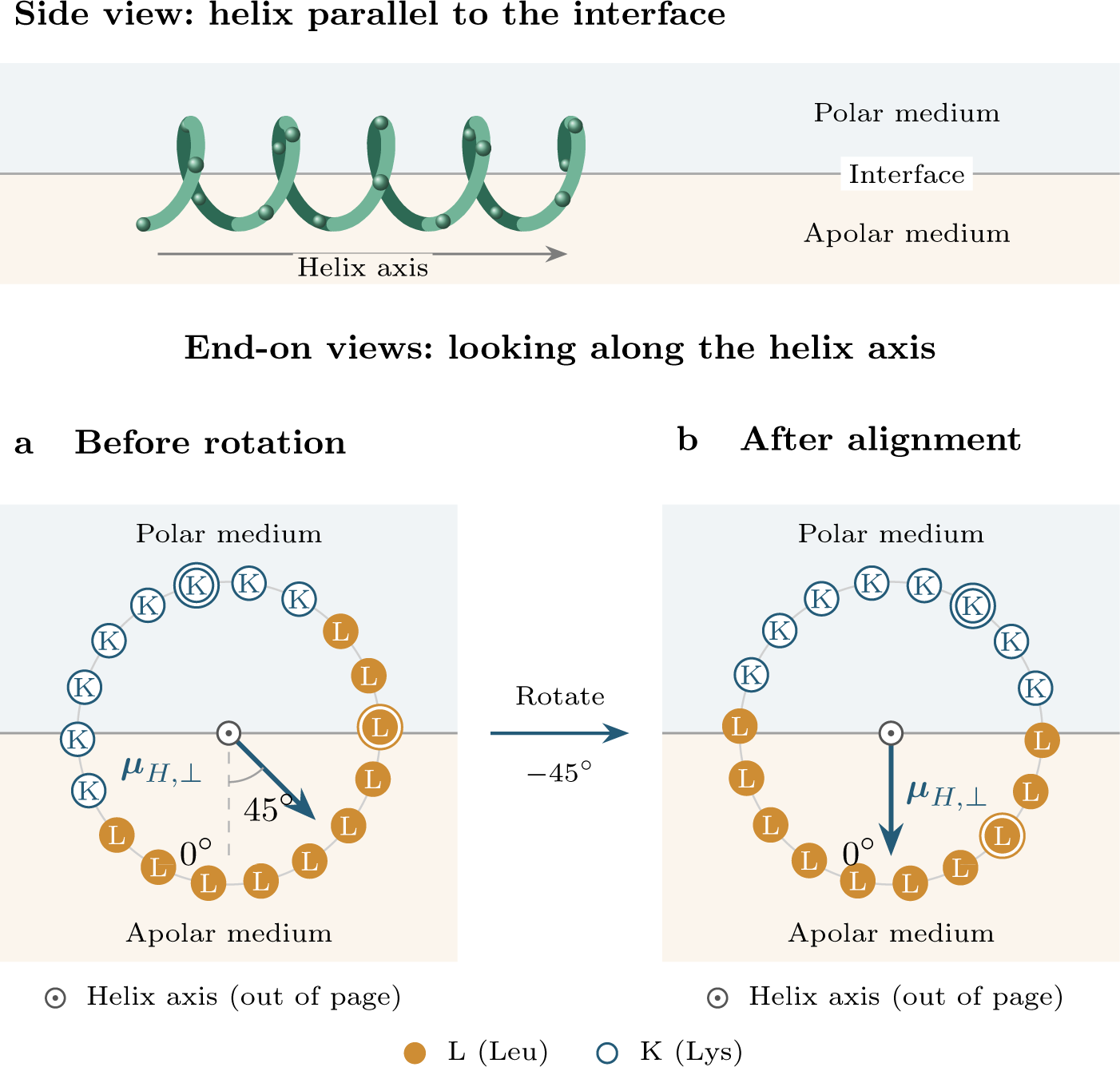
Helix alignment at a horizontal polar–apolar interface. The upper schematic shows the prescribed helix from the side, with its long axis parallel to the interface. Below, endon helical-wheel views show the same P1 sequence (Table 2) before (a) and after (b) rotation about that axis, which points out of the drawing plane. The polar medium is above and the apolar medium below in every view. The transverse hydrophobic moment, ***µ****_H,_*_⊥_, lies in the wheel plane and rotates with the residues. After rotation it points downwards, towards the centre of the apolar sector, displayed as 0^◦^ to match the web application. The initial 45^◦^ offset is illustrative. A 180^◦^ apolar sector, helix centre and exposure setting are held fixed. Residues are projected at a 100^◦^ helical angle; concentric circles mark coincident projections of pairs 1/19 and 2/20. During optimization, alignment is recalculated for every proposed sequence.

Each of the four workflows was run under both orientation rules with matched seeds 42, 43 and 44, giving 12 paired comparisons. All sequences have 20 residues and use the full amino-acid alphabet. Each strategy calibrates the model with 2,000 sequences under its own orientation rule. Design, Family and Cross-Design use 3,000 steps and two restarts per search; Family returns four candidates. Specificity requests two candidates and shares four total restarts across its searches.

The two strategies returned different sets of sequences in every paired comparison. No identical amino-acid sequence was shared between the fixed and aligned searches, either within a matched seed or across the three seeds. Thus, changing the orientation rule during search changes the identities of the designed candidates in these tests.

To compare the resulting candidates on a common basis, all returned sequences were evaluated again with hydrophobic-moment alignment, 2,000 calibration sequences, 1,000 reference sequences and seed 42. Lower scores indicate more favourable values of the relevant model objective: *H* for Design and Family, *L_λ_* with *λ* = 0.5 for Specificity, and *C_γ_* with *γ* = 0.5 for Cross-Design. For Family and Specificity, the comparison selects the lowest-scoring member of each returned collection after this common evaluation. The best candidates from matched searches differed at 6–20 of their 20 residue positions.

Table 4 shows that aligned search produced the more favourable best candidate for all three seeds in each of the four workflows. Changing the orientation rule therefore changed the designed sequences in all tested cases, while its effect on the best model score depended on the workflow and search seed. The application consistently uses hydrophobic-moment alignment during search and result presentation. Complete candidate sequences, scores, settings and source hashes are preserved in Additional file 2.

**Table 4:** Sequence identity and model scores after searches with fixed or aligned helix orientation. Each workflow has three paired searches with matched seeds.

| Workflow | Candidates per search | Shared sequences | Aligned search lower score | Fixed search lower score |
| --- | --- | --- | --- | --- |
| Design | 1 | 0 | 3/3 | 0/3 |
| Family | 4 | 0 | 3/3 | 0/3 |
| Specificity | 2 | 0 | 3/3 | 0/3 |
| Cross-Design | 1 | 0 | 3/3 | 0/3 |

Shared sequences are exact matches between strategies in the paired searches; none occurred. The last two columns count the paired searches in which each strategy produced the lower-scoring best candidate under the common aligned evaluation. Each workflow uses its own model objective.

### 3.9 Relationship to existing tools

Table 5 summarizes documented functionality; the executed descriptor comparison is reported separately. HELIQUEST[19, 23] offers sequence design against selected physicochemical properties; modlAMP[24] provides programmatic generation and analysis of peptide libraries; helixvis[25] addresses programmable visualization; SuPepMem[26] makes peptide–membrane simulation data accessible. Helix Design Studio[12] provides an accessible, inspectable implementation of the environmental Hamiltonian[10] and its associated decision workflows. The executed descriptor comparison assesses implementation consistency under shared calculation conventions.

**Table 5:** Scope of related software resources.

| Resource | Access mode | Documented scope relevant to this study |
| --- | --- | --- |
| HELIQUEST[19, 23] | Browser | Helical descriptors, composition, screening, mutation and genetic-algorithm design. |
| modlAMP[24] | Python | Descriptors, peptide/library generation, analysis and visualization. |
| helixvis[25] | R | Helical-wheel and wenxiang visualization with programmable customization. |
| SuPepMem[26] | Browser/database | Molecular-simulation data and descriptors for innate immune-system peptides interacting with model membranes. |
| Helix Design Studio[12] | Browser | Decomposed environmental objective, linked scoring/design tasks, target-competitor analysis, exposure scan and idealized PDB export. |

Capabilities were assessed from the cited primary papers and project documentation, with the literature review updated on 15 October 2026. The table summarizes the documented capabilities of the selected resources.

### 3.10 Interpretation and limitations

The software supports inspection of score contributions, environmental preferences, candidate diversity and sampling sensitivity within an explicit model. The evaluation characterizes computational behaviour. The generated candidates are intended for subsequent simulation and experimental characterization. Favourable scores alone do not establish antimicrobial potency, host-cell safety or membrane selectivity.

The geometric and chemical scope is a prescribed *α*-helical scaffold, the 20 standard amino acids, fixed integer side-chain charges and simplified solvent or interface parameters. These assumptions describe a specified helical state rather than an equilibrium folding ensemble. Hydrophobic-moment alignment selects a prescribed wheel phase, and exposure scans vary an angular mask with homogeneous endpoints. These geometric operations do not determine equilibrium orientations, physical insertion depths or membrane-insertion free energies. Evaluation of terminal modifications, noncanonical residues and pH-dependent charge states requires extending the chemical model. The empirical transfer of M1 propensities and the Cys/Pro defaults are specified with the parameter sources in Section 2.6. The interfacial charge component retains favourable opposite-charge coupling; same-sign and neutral residues receive the same zero raw contribution in that component. Residue–residue electrostatics is treated separately. The electrostatic factors and sequence-distance law are effective model choices; uniform multiplicative factors cancel upon component standardization.

Calibration/reference ensembles, residue alphabet, phase and weights define each result. Optimizing a sequence favours its own model score. Independent validation requires evidence beyond the optimized scoring model, such as experimental measurements. Comparisons across calculation protocols require matching these settings. Search outcomes reflect the specified stochastic budgets, and the exact benchmark establishes optimality within its enumerated sequence space. Computational timings characterize sequential calculations on the specified hardware configuration; deployment capacity and concurrent performance require dedicated measurements. Synthesis feasibility, toxicity and aggregation also require assessment during candidate selection. The distributed parameter tables and explicit calculation protocols make the numerical model reproducible. Further evaluation should use experimentally characterized peptide systems and examine deployment performance, usability and additional chemical constraints.

## 4 Conclusions

Helix Design Studio makes environmental preference an explicit objective in the design of peptide sequences on a prescribed *α*-helical scaffold. By connecting sequence analysis, candidate generation and comparisons between environments within a shared scoring framework, it allows users to examine why a sequence is favoured under one set of conditions and what compromises arise under another. Decomposed score contributions, sequence-dependent alignment and exportable calculation settings make these choices inspectable and reproducible through a browser-based workflow.

The computational evaluation supports implementation consistency and shows that numerical parameters affect environmental scores, while orientation conventions can change the sequences selected by optimization. These findings underline the value of recording model assumptions alongside each design. The resulting scores describe the compatibility of an assumed helical state; environment-dependent folding, membrane selectivity and biological activity remain questions for independent investigation. With its open-source implementation and reproducible candidate records, Helix Design Studio provides a practical starting point for turning membrane-interaction hypotheses into peptide sequences for simulation and experimental testing.

## Supporting information

Additional file 1. Evaluated software source and dependencies

Additional file 2. Computational evaluation, inputs and results

## Availability and requirements

**Project name:** Helix Design Studio.

**Project home page:** Web application[14]; source repository[12].

**Archived version:** v1.0.0rc6, commit d6ccec1920f2; Additional file 1.

**Operating systems:** Browser-based access; Linux or macOS for server deployment.

**Programming languages:** Python, JavaScript, HTML and CSS.

**Other requirements:** WebGL for the 3D viewer; Python 3.12 and packaged dependencies for server installation.

**License:** MIT; third-party licence notices accompany the source.

**Restrictions for non-academic use:** None.

## Declarations

### Availability of data and materials

Source code is publicly available in the project repository[12]. Additional file 1 provides the exact source snapshot used for the evaluation. Additional file 2 contains the executable study, machine-readable calculation inputs and outputs, summary tables, calibration records, numerical parameter tables, source hashes and figure-generation script. Additional file 3 contains the figure assets. The evaluated version is identified by ts public tag, commit and archive checksums. The main workflow inputs are predefined model sequences and computationally generated candidates; the parameter-correction comparison additionally uses publicly documented reference sequences. No private usage data or human-subject records were used.

### Ethics approval and consent to participate

Not applicable.

### Consent for publication

Not applicable.

### Authors’ contributions

D.C.-T. developed the software, carried out the computational evaluation, prepared the figures and wrote the original manuscript. R.G.-F. and Á.P. conceived and directed the work, supervised the software development and evaluation, and revised the manuscript.

## Acknowledgements

We thank CESGA for providing access to classical and quantum computing resources and for its technical support.

## Additional files

*Additional file 1: software source*

File: `additional_file_1_source_1.0.0rc6.zip`. Format: ZIP (`.zip`). The complete evaluated source revision, including licences, dependency specifications, tests and run nstructions.

*Additional file 2: reproducible evaluation*

File: `additional_file_2_evaluation.zip`. Format: ZIP (`.zip`). Study and plotting scripts, complete input settings and calculation outputs, machine-readable results, complete neighbour parameter tables and provenance metadata. Predefined model sequences are distinguished from computationally generated candidates; native search and common rescoring outputs are stored separately.

*Additional file 3: figures*

File: `additional_file_3_figures.zip`. Format: ZIP (`.zip`). Vector PDF and raster PNG exports of the seven main figures, with their legends. The graphical abstract is supplied separately.

