## Additional file 1. Evaluated software source and dependencies for "Helix Design Studio: designing antimicrobial peptide candidates for membrane environments": engine.py

```
from __future__ import annotations

import math
import os
import statistics
import sys
import tempfile
import threading
import time
from contextlib import contextmanager
from contextvars import ContextVar
from dataclasses import dataclass
from pathlib import Path
from typing import Any, Callable, Dict, Generator, List, Optional

import numpy as np

def _resolve_code_dir() -> Path:
    # QFOLD_CODE_DIR se mantiene como nombre heredado.
    env_override = os.environ.get("HELIX_CODE_DIR") or os.environ.get("QFOLD_CODE_DIR")
    candidate = (Path(env_override).expanduser().resolve() if env_override
                 else Path(__file__).resolve().parents[1] / "Code")
    if not (candidate / "core" / "hamiltonian_builder.py").is_file():
        raise FileNotFoundError(f"Could not locate the Helix Design Studio runtime at {candidate}.")
    return candidate


CODE_DIR = _resolve_code_dir()
if str(CODE_DIR) not in sys.path:
    sys.path.insert(0, str(CODE_DIR))

from core.hamiltonian_builder import (  # noqa: E402
    FAUCHERE_PLISKA_HYDROPHOBICITY,
    HamiltonianBuilder,
    hydrophobic_moment,
)

DEFAULT_ALPHABET_16 = ["A", "P", "D", "E", "Q", "L", "M", "F", "K", "S", "T", "W", "Y", "V", "H", "G"]
DEFAULT_WEIGHTS = {
    "lambda_env": 6.0,
    "lambda_charge": 3.0,
    "lambda_local": 1.5,
    "lambda_pairwise": 0.1,
    "lambda_helix_pairs": 1.5,
    "lambda_electrostatic": 4.0,
}


def default_cache_dir() -> str:
    """Se lee en cada llamada para respetar HELIX_CACHE_DIR aunque cambie tras importar."""
    return os.environ.get("HELIX_CACHE_DIR") or str(Path(tempfile.gettempdir()) / "helix-design-studio-cache")


ENVIRONMENT_PRESETS: Dict[str, Dict[str, Any]] = {
    "homogeneous_polar": {
        "label": "Homogeneous polar medium",
        "wheel_phase_deg": 90.0,
        "wheel_halfwidth_deg": -1.0,
        "membrane_charge": "neu",
    },
    "homogeneous_apolar": {
        "label": "Homogeneous apolar medium",
        "wheel_phase_deg": 90.0,
        "wheel_halfwidth_deg": 180.0,
        "membrane_charge": "neu",
    },
    "interfacial_neg": {
        "label": "Interfacial, anionic membrane",
        "wheel_phase_deg": 90.0,
        "wheel_halfwidth_deg": 90.0,
        "membrane_charge": "neg",
    },
    "interfacial_neu": {
        "label": "Interfacial, neutral membrane",
        "wheel_phase_deg": 90.0,
        "wheel_halfwidth_deg": 90.0,
        "membrane_charge": "neu",
    },
    "interfacial_pos": {
        "label": "Interfacial, cationic membrane",
        "wheel_phase_deg": 90.0,
        "wheel_halfwidth_deg": 90.0,
        "membrane_charge": "pos",
    },
}

HYDROPHOBICITY_FP = FAUCHERE_PLISKA_HYDROPHOBICITY

RESIDUE_COLORS = {
    "hydrophobic": "#b05a1b",
    "polar": "#23796a",
    "positive": "#e03131",
    "negative": "#265dcb",
    "special": "#6c4bf0",
}

HYDROPHOBIC = {"A", "V", "L", "I", "M", "F", "W", "Y", "C", "P"}
POLAR = {"Q", "N", "S", "T", "G"}
POSITIVE = {"K", "R"}
NEGATIVE = {"D", "E"}

THREE_LETTER_CODES = {
    "A": "ALA",
    "R": "ARG",
    "N": "ASN",
    "D": "ASP",
    "C": "CYS",
    "Q": "GLN",
    "E": "GLU",
    "G": "GLY",
    "H": "HIS",
    "I": "ILE",
    "L": "LEU",
    "K": "LYS",
    "M": "MET",
    "F": "PHE",
    "P": "PRO",
    "S": "SER",
    "T": "THR",
    "W": "TRP",
    "Y": "TYR",
    "V": "VAL",
}


def normalize_residues(residues: Optional[List[str] | str]) -> List[str]:
    if residues is None:
        return list(DEFAULT_ALPHABET_16)
    if isinstance(residues, str):
        if "," in residues:
            toks = [t.strip().upper() for t in residues.split(",") if t.strip()]
        else:
            toks = [c.strip().upper() for c in residues if c.strip()]
    else:
        toks = [str(t).strip().upper() for t in residues if str(t).strip()]
    seen: List[str] = []
    for aa in toks:
        if aa not in seen:
            seen.append(aa)
    return seen or list(DEFAULT_ALPHABET_16)


def normalize_weights(weights: Optional[Dict[str, float]]) -> Dict[str, float]:
    merged = dict(DEFAULT_WEIGHTS)
    if weights:
        merged.update({k: float(v) for k, v in weights.items() if v is not None})
    return merged


def resolve_environment(environment: Optional[Dict[str, Any]]) -> Dict[str, Any]:
    environment = environment or {}
    preset = environment.get("preset", "interfacial_neg")
    base = dict(ENVIRONMENT_PRESETS.get(preset, ENVIRONMENT_PRESETS["interfacial_neg"]))
    for key in ("wheel_phase_deg", "wheel_halfwidth_deg", "membrane_charge"):
        if environment.get(key) is not None:
            base[key] = environment[key]
    base["preset"] = preset
    return base


def is_interfacial(environment: Dict[str, Any]) -> bool:
    return str(environment.get("preset", "")).startswith("interfacial_")


def physical_environment_key(environment: Dict[str, Any]) -> tuple[Any, ...]:
    """Configuración física y regla de orientación del Hamiltoniano.

    En un sector parcial, el preset determina si cada secuencia se alinea por su
    momento hidrofóbico. La fase base sigue importando cuando ese momento es nulo.
    """
    env = resolve_environment(environment)
    half_width = float(env["wheel_halfwidth_deg"])
    if half_width >= 180.0:
        # Todo el contorno es apolar: el término de carga, que solo actúa en agua, es nulo.
        return ("all_apolar",)
    charge = str(env["membrane_charge"])
    if half_width < 0.0:
        return ("all_polar", charge)
    return ("sector", half_width, float(env["wheel_phase_deg"]), charge, is_interfacial(env))


def effective_environment(builder: HamiltonianBuilder, codes: List[int], environment: Dict[str, Any]) -> Dict[str, Any]:
    """Entorno con la fase de rueda realmente usada para puntuar esta secuencia."""
    env = dict(environment)
    env["wheel_phase_deg"] = builder.phase_for(codes)
    if builder.align_phase:
        descriptor = hydrophobic_moment_descriptor(codes_to_seq(codes, builder.amino_acids))
        env["phase_aligned_to_hydrophobic_moment"] = descriptor["magnitude"] >= 1e-10
        env["hydrophobic_moment_phase_deg"] = descriptor["phase_deg"]
    return env


def hydrophobic_moment_descriptor(sequence: str) -> Dict[str, float]:
    cleaned = sequence.strip().upper()
    mx, my = hydrophobic_moment(cleaned)
    magnitude = math.hypot(mx, my)
    phase_deg = math.degrees(math.atan2(my, mx)) if magnitude >= 1e-10 else 0.0
    return {
        "x": float(mx),
        "y": float(my),
        "magnitude": float(magnitude),
        "magnitude_per_residue": float(magnitude / len(cleaned)) if cleaned else 0.0,
        "phase_deg": float(phase_deg),
    }


def penetration_percent_to_halfwidth(percent: float) -> float:
    clamped = max(0.0, min(100.0, float(percent)))
    return 40.0 + (clamped / 100.0) * 110.0


def penetration_percent_to_environment(percent: int, interface_preset: str) -> Dict[str, Any]:
    clamped = max(0, min(100, int(percent)))
    if clamped <= 0:
        env = dict(ENVIRONMENT_PRESETS["homogeneous_polar"])
        env["preset"] = "homogeneous_polar"
        env["penetration_regime"] = "polar_endpoint"
        return env
    if clamped >= 100:
        env = dict(ENVIRONMENT_PRESETS["homogeneous_apolar"])
        env["preset"] = "homogeneous_apolar"
        env["penetration_regime"] = "apolar_endpoint"
        return env
    env = dict(ENVIRONMENT_PRESETS.get(interface_preset, ENVIRONMENT_PRESETS["interfacial_neg"]))
    env["preset"] = interface_preset
    env["wheel_halfwidth_deg"] = penetration_percent_to_halfwidth(clamped)
    env["penetration_regime"] = "interfacial"
    return env


MJ_MAX_DISTANCE = 4  # contactos MJ entre i e i+1..i+4
DESIGN_N_RANDOM = 500


def build_builder(
    length: int,
    amino_acids: List[str],
    environment: Dict[str, Any],
    n_decoys: int = 2000,
    zscore_seed: int = 42,
    stats_cache_dir: Optional[str] = None,
    oriented: bool = False,
) -> HamiltonianBuilder:
    """En interfaces, oriented=True alinea cada secuencia por su propio momento
    hidrofóbico: consulta, propuestas, calibración y referencias comparten esta regla.
    oriented=False permite comparaciones explícitas con una fase fija."""
    return HamiltonianBuilder(
        L=length,
        amino_acids=amino_acids,
        n_decoys=n_decoys,
        zscore_seed=zscore_seed,
        stats_cache_dir=stats_cache_dir or default_cache_dir(),
        stats_cache_max_files=int(os.environ.get("HELIX_CACHE_MAX_FILES") or 2000),
        wheel_phase_deg=float(environment["wheel_phase_deg"]),
        wheel_halfwidth_deg=float(environment["wheel_halfwidth_deg"]),
        membrane_charge=str(environment["membrane_charge"]),
        align_phase=oriented and is_interfacial(environment),
        max_interaction_dist=MJ_MAX_DISTANCE,
        interrupt_check=_check_limits,
    )


def seq_to_codes(sequence: str, amino_acids: List[str]) -> List[int]:
    aa_to_idx = {aa: i for i, aa in enumerate(amino_acids)}
    try:
        return [aa_to_idx[aa] for aa in sequence]
    except KeyError as exc:
        raise ValueError(f"Residue {exc.args[0]} is not present in the selected alphabet") from exc


def codes_to_seq(codes: List[int], amino_acids: List[str]) -> str:
    return "".join(amino_acids[idx] for idx in codes)


class CalculationTimeout(RuntimeError):
    """El cálculo superó el tiempo máximo asignado a una petición."""


class CalculationCancelled(RuntimeError):
    """El cliente abandonó la petición y el cálculo se detuvo."""


@dataclass(frozen=True)
class CalculationLimits:
    deadline: Optional[float] = None  # instante límite en time.monotonic()
    cancelled: Optional[threading.Event] = None


# Un ContextVar se copia al hilo del pool que ejecuta el cálculo.
_calculation_limits: ContextVar[Optional[CalculationLimits]] = ContextVar("calculation_limits", default=None)


@contextmanager
def calculation_limits(limits: Optional[CalculationLimits]) -> Generator[None, None, None]:
    token = _calculation_limits.set(limits)
    try:
        yield
    finally:
        _calculation_limits.reset(token)


def _check_limits() -> None:
    limits = _calculation_limits.get()
    if limits is None:
        return
    if limits.cancelled is not None and limits.cancelled.is_set():
        raise CalculationCancelled("The calculation was cancelled.")
    if limits.deadline is not None and time.monotonic() > limits.deadline:
        raise CalculationTimeout("The calculation exceeded the time limit of the web server.")


def compute_breakdown(builder: HamiltonianBuilder, codes: List[int], weights: Dict[str, float]) -> Dict[str, float]:
    _check_limits()
    raw = builder.raw_terms(codes)
    z = {
        "local": builder._apply_z_score("helix_local", raw["helix_local"]),
        "env": builder._apply_z_score("env_pol", raw["env_pol"]),
        "charge": builder._apply_z_score("env_chg", raw["env_chg"]),
        "pairwise": builder._apply_z_score("pw_int", raw["pw_int"]),
        "electrostatic": builder._apply_z_score("electrostatic", raw["electrostatic"]),
        "neigh": builder._apply_z_score("helix_neigh", raw["helix_neigh"]),
    }
    weighted = {
        "local": weights["lambda_local"] * z["local"],
        "env": weights["lambda_env"] * z["env"],
        "charge": weights["lambda_charge"] * z["charge"],
        "pairwise": weights["lambda_pairwise"] * z["pairwise"],
        "electrostatic": weights["lambda_electrostatic"] * z["electrostatic"],
        "neigh": weights["lambda_helix_pairs"] * z["neigh"],
    }
    weighted["total"] = float(sum(weighted.values()))
    return weighted


def residue_family(aa: str) -> str:
    if aa in POSITIVE:
        return "positive"
    if aa in NEGATIVE:
        return "negative"
    if aa in POLAR:
        return "polar"
    if aa in HYDROPHOBIC:
        return "hydrophobic"
    return "special"


def wheel_data(sequence: str, environment: Dict[str, Any]) -> List[Dict[str, Any]]:
    phase = float(environment["wheel_phase_deg"])
    half_width = float(environment["wheel_halfwidth_deg"])
    data = []
    for idx, aa in enumerate(sequence, start=1):
        theta = ((idx - 1) * 100.0 + phase) % 360.0
        signed = theta if theta <= 180.0 else theta - 360.0
        env = "membrane" if abs(signed) <= half_width else "water"
        rad = math.radians(theta - 90.0)
        data.append(
            {
                "index": idx,
                "residue": aa,
                "angle_deg": theta,
                "signed_angle_deg": signed,
                "x": round(math.cos(rad), 6),
                "y": round(math.sin(rad), 6),
                "environment": env,
                "family": residue_family(aa),
                "color": RESIDUE_COLORS[residue_family(aa)],
            }
        )
    return data


def model_net_charge(sequence: str) -> int:
    charge_map = {"R": 1, "K": 1, "D": -1, "E": -1}
    return int(sum(charge_map.get(aa, 0) for aa in sequence))


# Flujos aleatorios independientes derivados de la semilla del usuario (SeedSequence): la
# calibración usa la semilla tal cual; referencias y búsquedas usan subflujos distintos.
REFERENCE_STREAM = 1
SEARCH_STREAM = 2


SEED_PROTOCOL = (
    "numpy.random.default_rng with SeedSequence entropy: calibration decoys [seed]; "
    "random reference [seed, 1]; search [seed, 2] for single searches and [seed, 2, i] "
    "for the i-th run of family and specificity searches"
)


def rng_seed(seed: int, stream: int, *index: int) -> List[int]:
    return [int(seed), stream, *index]


def random_reference(
    builder: HamiltonianBuilder,
    amino_acids: List[str],
    weights: Dict[str, float],
    n_random: int,
    seed: int | List[int],
) -> Dict[str, float]:
    rng = np.random.default_rng(seed)
    energies = []
    for _ in range(n_random):
        codes = rng.integers(0, len(amino_acids), size=builder.L).tolist()
        energies.append(compute_breakdown(builder, codes, weights)["total"])
    mean = statistics.fmean(energies)
    std = statistics.pstdev(energies) or 1.0
    return {"mean": float(mean), "std": float(std), "energies": energies, "seed_entropy": seed}


def score_sequence_with_builder(
    sequence: str,
    builder: HamiltonianBuilder,
    environment: Dict[str, Any],
    amino_acids: List[str],
    weights: Dict[str, float],
    ref: Dict[str, float],
    n_random: int,
) -> Dict[str, Any]:
    sequence = sequence.strip().upper()
    codes = seq_to_codes(sequence, amino_acids)
    environment = effective_environment(builder, codes, environment)
    breakdown = compute_breakdown(builder, codes, weights)
    mu_descriptor = hydrophobic_moment_descriptor(sequence)
    energy = breakdown["total"]
    z_score = (energy - ref["mean"]) / ref["std"]
    percentile = 100.0 * sum(e <= energy for e in ref["energies"]) / len(ref["energies"])
    return {
        "sequence": sequence,
        "length": len(sequence),
        "environment": environment,
        "weights": weights,
        "energy": energy,
        "z_score": float(z_score),
        "percentile_lower_is_better": float(percentile),
        "random_reference": {
            "mean": ref["mean"], "std": ref["std"], "n": n_random, "seed_entropy": ref.get("seed_entropy"),
        },
        "calibration": {
            "n_decoys": int(builder.kwargs.get("n_decoys", 0)),
            "seed_entropy": int(builder.kwargs.get("zscore_seed", 0)),
            "orientation": "own hydrophobic moment" if builder.align_phase else "fixed wheel phase",
            "terms": {term: {"mu": mu, "sigma": sigma} for term, (mu, sigma) in builder.stats.items()},
        },
        "breakdown": breakdown,
        "observables": {
            "abs_mu_h": mu_descriptor["magnitude"],
            "abs_mu_h_per_residue": mu_descriptor["magnitude_per_residue"],
            "mu_h_phase_deg": mu_descriptor["phase_deg"],
        },
        "wheel": wheel_data(sequence, environment),
        "model_net_charge": model_net_charge(sequence),
        "alphabet": amino_acids,
    }


@dataclass
class ScoringModel:
    """Builder calibrado y referencia aleatoria para un entorno y una regla de orientación."""
    builder: HamiltonianBuilder
    environment: Dict[str, Any]
    reference: Dict[str, Any]
    n_random: int


def scoring_model(
    length: int,
    amino_acids: List[str],
    environment: Dict[str, Any],
    weights: Dict[str, float],
    *,
    n_decoys: int,
    seed: int,
    n_random: int,
    oriented: bool,
) -> ScoringModel:
    builder = build_builder(length, amino_acids, environment, n_decoys=n_decoys, zscore_seed=seed, oriented=oriented)
    reference = random_reference(builder, amino_acids, weights, n_random=n_random, seed=rng_seed(seed, REFERENCE_STREAM))
    return ScoringModel(builder, environment, reference, n_random)


def report_models(
    length: int,
    amino_acids: List[str],
    environment: Dict[str, Any],
    weights: Dict[str, float],
    *,
    n_decoys: int,
    seed: int,
    n_random: int,
) -> tuple[ScoringModel, ScoringModel]:
    """Un único modelo para la búsqueda y el resultado; se conserva la tupla por compatibilidad."""
    model = scoring_model(length, amino_acids, environment, weights,
                          n_decoys=n_decoys, seed=seed, n_random=n_random, oriented=True)
    return model, model


def score_with_model(sequence: str, model: ScoringModel, amino_acids: List[str], weights: Dict[str, float]) -> Dict[str, Any]:
    return score_sequence_with_builder(
        sequence, model.builder, model.environment, amino_acids, weights, model.reference, model.n_random
    )


def report_sequence(
    sequence: str,
    design: ScoringModel,
    scoring: ScoringModel,
    amino_acids: List[str],
    weights: Dict[str, float],
) -> Dict[str, Any]:
    """Puntuación optimizada, con el mismo protocolo que Score.

    design_geometry conserva el esquema de exportación anterior, pero ahora
    reproduce la misma puntuación y orientación que el resultado principal.
    """
    report = score_with_model(sequence, scoring, amino_acids, weights)
    in_design = report if design is scoring else score_with_model(sequence, design, amino_acids, weights)
    orientation = in_design["calibration"]["orientation"]
    if design.builder.align_phase and not in_design["environment"]["phase_aligned_to_hydrophobic_moment"]:
        orientation = "preset wheel phase (vanishing hydrophobic moment)"
    report["design_geometry"] = {
        "energy": in_design["energy"],
        "z_score": in_design["z_score"],
        "percentile_lower_is_better": in_design["percentile_lower_is_better"],
        "breakdown": in_design["breakdown"],
        "wheel_phase_deg": in_design["environment"]["wheel_phase_deg"],
        "orientation": orientation,
    }
    return report


def hamming_distance_codes(a: List[int], b: List[int]) -> int:
    return sum(x != y for x, y in zip(a, b))


def diversity_penalty(
    codes: List[int],
    accepted_codes: List[List[int]],
    min_distance: int,
    penalty_weight: float,
) -> float:
    if not accepted_codes:
        return 0.0
    penalty = 0.0
    for prior in accepted_codes:
        distance = hamming_distance_codes(codes, prior)
        deficit = max(0, min_distance - distance)
        if deficit:
            penalty += penalty_weight * deficit
    return penalty


def score_sequence(
    sequence: str,
    environment: Optional[Dict[str, Any]] = None,
    residues: Optional[List[str] | str] = None,
    weights: Optional[Dict[str, float]] = None,
    n_decoys: int = 2000,
    n_random: int = 500,
    seed: int = 42,
) -> Dict[str, Any]:
    sequence = sequence.strip().upper()
    amino_acids = normalize_residues(residues)
    seq_to_codes(sequence, amino_acids)  # valida antes de calibrar
    env = resolve_environment(environment)
    w = normalize_weights(weights)
    model = scoring_model(len(sequence), amino_acids, env, w, n_decoys=n_decoys, seed=seed, n_random=n_random, oriented=True)
    return score_with_model(sequence, model, amino_acids, w)


def simulated_annealing(
    builder: HamiltonianBuilder,
    amino_acids: List[str],
    weights: Dict[str, float],
    steps: int,
    restarts: int,
    seed: int | List[int],
    penalty_fn: Optional[Callable[[List[int]], float]] = None,
) -> Dict[str, Any]:
    rng = np.random.default_rng(seed)
    n_aa = len(amino_acids)
    best_codes = None
    best_objective = float("inf")
    best_energy = float("inf")

    for _ in range(restarts):
        current = rng.integers(0, n_aa, size=builder.L).tolist()
        current_energy = compute_breakdown(builder, current, weights)["total"]
        current_objective = current_energy + penalty_fn(current) if penalty_fn else current_energy
        local_best = list(current)
        local_best_energy = current_energy
        local_best_objective = current_objective

        for step in range(steps):
            temperature = max(1e-4, 2.0 * (1.0 - step / steps))
            proposal = list(current)
            pos = int(rng.integers(0, builder.L))
            old_code = proposal[pos]
            new_code = int(rng.integers(0, n_aa - 1))
            if new_code >= old_code:
                new_code += 1
            proposal[pos] = new_code

            proposal_energy = compute_breakdown(builder, proposal, weights)["total"]
            proposal_objective = proposal_energy + penalty_fn(proposal) if penalty_fn else proposal_energy
            delta = proposal_objective - current_objective
            if delta <= 0 or rng.random() < math.exp(-delta / temperature):
                current = proposal
                current_energy = proposal_energy
                current_objective = proposal_objective
                if current_objective < local_best_objective:
                    local_best = list(current)
                    local_best_energy = current_energy
                    local_best_objective = current_objective

        if local_best_objective < best_objective:
            best_codes = list(local_best)
            best_energy = local_best_energy
            best_objective = local_best_objective

    assert best_codes is not None
    sequence = codes_to_seq(best_codes, amino_acids)
    return {
        "sequence": sequence,
        "energy": best_energy,
        "objective": best_objective,
        "breakdown": compute_breakdown(builder, best_codes, weights),
        "codes": list(best_codes),
    }


def _iter_specificity_search(
    target_builder: HamiltonianBuilder,
    off_builders: List[HamiltonianBuilder],
    amino_acids: List[str],
    weights: Dict[str, float],
    lambda_balance: float,
    steps: int,
    restarts: int,
    seed: int | List[int],
    accepted_codes: Optional[List[List[int]]] = None,
    min_distance: int = 0,
    penalty_weight: float = 0.0,
    progress_callback: Optional[Callable[[Dict[str, Any]], None]] = None,
) -> Dict[str, Any]:
    rng = np.random.default_rng(seed)
    n_aa = len(amino_acids)
    accepted_codes = accepted_codes or []
    best_codes = None
    best_objective = float("inf")
    best_penalized_objective = float("inf")
    best_target_energy = float("inf")
    best_off_energies: List[float] = []
    checkpoint = max(250, min(2000, steps // 30))

    for restart_idx in range(restarts):
        current = rng.integers(0, n_aa, size=target_builder.L).tolist()
        current_target = compute_breakdown(target_builder, current, weights)["total"]
        current_off = [compute_breakdown(builder, current, weights)["total"] for builder in off_builders]
        current_raw_objective = current_target + (lambda_balance - 1.0) * min(current_off)
        current_objective = current_raw_objective + diversity_penalty(
            current,
            accepted_codes,
            min_distance=min_distance,
            penalty_weight=penalty_weight,
        )
        local_best = list(current)
        local_best_target = current_target
        local_best_off = list(current_off)
        local_best_objective = current_raw_objective
        local_best_penalized_objective = current_objective

        progress_payload = {
            "phase": "annealing",
            "restart": restart_idx + 1,
            "restarts": restarts,
            "step": 0,
            "steps": steps,
            "progress_fraction": restart_idx / max(1, restarts),
            "latest_sequence": codes_to_seq(current, amino_acids),
            "best_sequence": codes_to_seq(local_best, amino_acids),
            "best_objective": local_best_objective,
        }
        if progress_callback:
            progress_callback(progress_payload)
        yield {"type": "progress", "payload": progress_payload}

        for step_idx in range(steps):
            temperature = max(1e-4, 2.0 * (1.0 - step_idx / steps))
            proposal = list(current)
            pos = int(rng.integers(0, target_builder.L))
            old_code = proposal[pos]
            new_code = int(rng.integers(0, n_aa - 1))
            if new_code >= old_code:
                new_code += 1
            proposal[pos] = new_code

            proposal_target = compute_breakdown(target_builder, proposal, weights)["total"]
            proposal_off = [compute_breakdown(builder, proposal, weights)["total"] for builder in off_builders]
            proposal_raw_objective = proposal_target + (lambda_balance - 1.0) * min(proposal_off)
            proposal_objective = proposal_raw_objective + diversity_penalty(
                proposal,
                accepted_codes,
                min_distance=min_distance,
                penalty_weight=penalty_weight,
            )
            delta = proposal_objective - current_objective
            if delta <= 0 or rng.random() < math.exp(-delta / temperature):
                current = proposal
                current_target = proposal_target
                current_off = proposal_off
                current_objective = proposal_objective
                current_raw_objective = proposal_raw_objective
                if current_objective < local_best_penalized_objective:
                    local_best = list(current)
                    local_best_target = current_target
                    local_best_off = list(current_off)
                    local_best_objective = current_raw_objective
                    local_best_penalized_objective = current_objective

            if (step_idx + 1) % checkpoint == 0 or step_idx + 1 == steps:
                overall = (restart_idx + (step_idx + 1) / max(1, steps)) / max(1, restarts)
                use_previous_best = (
                    best_codes is not None and best_penalized_objective <= local_best_penalized_objective
                )
                best_sequence_codes = best_codes if use_previous_best else local_best
                best_sequence = codes_to_seq(best_sequence_codes, amino_acids)
                best_obj = best_objective if use_previous_best else local_best_objective
                progress_payload = {
                    "phase": "annealing",
                    "restart": restart_idx + 1,
                    "restarts": restarts,
                    "step": step_idx + 1,
                    "steps": steps,
                    "progress_fraction": overall,
                    "latest_sequence": codes_to_seq(current, amino_acids),
                    "best_sequence": best_sequence,
                    "best_objective": best_obj,
                }
                if progress_callback:
                    progress_callback(progress_payload)
                yield {"type": "progress", "payload": progress_payload}

        if local_best_penalized_objective < best_penalized_objective:
            best_codes = list(local_best)
            best_objective = local_best_objective
            best_penalized_objective = local_best_penalized_objective
            best_target_energy = local_best_target
            best_off_energies = list(local_best_off)

    assert best_codes is not None
    yield {
        "type": "result",
        "payload": {
            "sequence": codes_to_seq(best_codes, amino_acids),
            "codes": list(best_codes),
            "objective": best_objective,
            "penalized_objective": best_penalized_objective,
            "best_target_energy": best_target_energy,
            "best_off_energies": best_off_energies,
        },
    }


SPECIFICITY_N_RANDOM = 3000


def _build_specificity_candidate_payload(
    sequence: str,
    *,
    length: int,
    amino_acids: List[str],
    target_env: Dict[str, Any],
    off_target_envs: List[Dict[str, Any]],
    target_models: tuple[ScoringModel, ScoringModel],
    off_models: List[tuple[ScoringModel, ScoringModel]],
    weights: Dict[str, float],
    lambda_balance: float,
    steps: int,
    restarts: int,
    seed: int,
    attempt: int,
    objective: float,
    best_target_energy: float,
    best_off_energies: List[float],
) -> Dict[str, Any]:
    # Objetivo y márgenes usan las mismas puntuaciones mostradas en cada entorno.
    target_score = report_sequence(sequence, *target_models, amino_acids, weights)
    off_target_scores = [report_sequence(sequence, *models, amino_acids, weights) for models in off_models]
    target_design = target_score["design_geometry"]
    off_design = [item["design_geometry"] for item in off_target_scores]
    worst_index = min(range(len(off_design)), key=lambda idx: off_design[idx]["energy"])
    worst_off_target = off_target_scores[worst_index]
    specificity_design = {
        "objective": objective,
        "objective_formula": "H_target + (lambda - 1) * min(H_offtargets)",
        "objective_formula_expanded": "lambda * H_target + (1 - lambda) * (H_target - min(H_offtargets))",
        "lambda_balance": lambda_balance,
        "target_preset": target_env["preset"],
        "off_target_presets": [env["preset"] for env in off_target_envs],
        "worst_off_target_preset": worst_off_target["environment"]["preset"],
        "energy_margin": off_design[worst_index]["energy"] - target_design["energy"],
        "z_score_margin": min(item["z_score"] for item in off_design) - target_design["z_score"],
        "margins_geometry": "same orientation rule as Score in each environment",
        "steps": steps,
        "restarts": restarts,
        "seed": seed,
        "attempt": attempt,
        "best_target_energy": best_target_energy,
        "best_off_target_energy": min(best_off_energies) if best_off_energies else None,
    }
    return {
        "sequence": sequence,
        "length": length,
        "alphabet": amino_acids,
        "target_environment": target_score,
        "off_target_environments": off_target_scores,
        "specificity_design": specificity_design,
    }


def design_sequence(
    length: int,
    environment: Optional[Dict[str, Any]] = None,
    residues: Optional[List[str] | str] = None,
    weights: Optional[Dict[str, float]] = None,
    n_decoys: int = 2000,
    steps: int = 12000,
    restarts: int = 8,
    seed: int = 42,
) -> Dict[str, Any]:
    amino_acids = normalize_residues(residues)
    env = resolve_environment(environment)
    w = normalize_weights(weights)
    design, scoring = report_models(length, amino_acids, env, w, n_decoys=n_decoys, seed=seed, n_random=DESIGN_N_RANDOM)
    result = simulated_annealing(
        design.builder, amino_acids, w, steps=steps, restarts=restarts, seed=rng_seed(seed, SEARCH_STREAM)
    )
    score = report_sequence(result["sequence"], design, scoring, amino_acids, w)
    score["design_settings"] = {"steps": steps, "restarts": restarts, "seed": seed}
    return score


def design_family(
    length: int,
    environment: Optional[Dict[str, Any]] = None,
    residues: Optional[List[str] | str] = None,
    weights: Optional[Dict[str, float]] = None,
    n_decoys: int = 2000,
    steps: int = 10000,
    restarts: int = 6,
    family_size: int = 8,
    oversample_factor: int = 3,
    seed: int = 42,
) -> Dict[str, Any]:
    final_update = None
    for update in design_family_stream(
        length=length,
        environment=environment,
        residues=residues,
        weights=weights,
        n_decoys=n_decoys,
        steps=steps,
        restarts=restarts,
        family_size=family_size,
        oversample_factor=oversample_factor,
        seed=seed,
    ):
        if update["type"] == "final":
            final_update = update["payload"]
    if final_update is None:
        raise RuntimeError("Family generation did not produce a final result")
    return final_update


def design_family_stream(
    length: int,
    environment: Optional[Dict[str, Any]] = None,
    residues: Optional[List[str] | str] = None,
    weights: Optional[Dict[str, float]] = None,
    n_decoys: int = 2000,
    steps: int = 10000,
    restarts: int = 6,
    family_size: int = 8,
    oversample_factor: int = 3,
    seed: int = 42,
) -> Generator[Dict[str, Any], None, None]:
    amino_acids = normalize_residues(residues)
    env = resolve_environment(environment)
    w = normalize_weights(weights)
    design, scoring = report_models(length, amino_acids, env, w, n_decoys=n_decoys, seed=seed, n_random=DESIGN_N_RANDOM)
    builder = design.builder

    unique_results: Dict[str, Dict[str, Any]] = {}
    accepted_codes: List[List[int]] = []
    total_runs = max(family_size * max(oversample_factor, 4), family_size + 6)
    min_distance = max(2, math.ceil(length * 0.15))
    penalty_weight = 6.0

    yield {
        "type": "meta",
        "payload": {
            "target": family_size,
            "total_runs_budget": total_runs,
            "min_distance": min_distance,
            "environment": env,
        },
    }

    for run_idx in range(total_runs):
        penalty_fn = None
        if accepted_codes:
            penalty_fn = lambda codes, prior=list(accepted_codes): diversity_penalty(
                codes, prior, min_distance=min_distance, penalty_weight=penalty_weight
            )
        candidate = simulated_annealing(
            builder,
            amino_acids,
            w,
            steps=steps,
            restarts=restarts,
            seed=rng_seed(seed, SEARCH_STREAM, run_idx),
            penalty_fn=penalty_fn,
        )
        sequence = candidate["sequence"]
        yield {
            "type": "progress",
            "payload": {
                "attempt": run_idx + 1,
                "accepted": len(unique_results),
                "target": family_size,
                "latest_sequence": sequence,
            },
        }
        if sequence in unique_results:
            continue
        scored = report_sequence(sequence, design, scoring, amino_acids, w)
        min_hamming = None
        if accepted_codes:
            min_hamming = min(hamming_distance_codes(candidate["codes"], prior) for prior in accepted_codes)
        unique_results[sequence] = scored
        accepted_codes.append(candidate["codes"])
        yield {
            "type": "candidate",
            "payload": {
                "attempt": run_idx + 1,
                "accepted": len(unique_results),
                "target": family_size,
                "min_hamming_to_previous": min_hamming,
                "member": scored,
            },
        }
        if len(unique_results) >= family_size:
            break

    # Orden por la misma energía que se optimizó y se muestra en Score.
    ranked = sorted(unique_results.values(), key=lambda item: item["design_geometry"]["energy"])[:family_size]
    payload = {
        "environment": env,
        "weights": w,
        "alphabet": amino_acids,
        "family_size_requested": family_size,
        "family_size_returned": len(ranked),
        "design_settings": {
            "steps": steps,
            "restarts": restarts,
            "seed": seed,
            "oversample_factor": oversample_factor,
            "min_distance": min_distance,
            "penalty_weight": penalty_weight,
        },
        "members": ranked,
        "status_message": (
            f"Generated {len(ranked)} unique candidates"
            if len(ranked) >= family_size
            else f"Generated {len(ranked)} unique candidates out of the requested {family_size}"
        ),
    }
    yield {"type": "final", "payload": payload}


def compare_environments(
    sequence: str,
    environment_a: Dict[str, Any],
    environment_b: Dict[str, Any],
    residues: Optional[List[str] | str] = None,
    weights: Optional[Dict[str, float]] = None,
    n_decoys: int = 2000,
    seed: int = 42,
) -> Dict[str, Any]:
    amino_acids = normalize_residues(residues)
    w = normalize_weights(weights)
    env_a = resolve_environment(environment_a)
    env_b = resolve_environment(environment_b)
    scored_a = score_sequence(sequence, env_a, amino_acids, w, n_decoys=n_decoys, seed=seed)
    scored_b = score_sequence(sequence, env_b, amino_acids, w, n_decoys=n_decoys, seed=seed)
    return {
        "sequence": sequence.upper(),
        "environment_a": scored_a,
        "environment_b": scored_b,
        "energy_gap": abs(scored_a["energy"] - scored_b["energy"]),
        "z_score_gap": abs(scored_a["z_score"] - scored_b["z_score"]),
        "joint_energy": scored_a["energy"] + scored_b["energy"],
    }


def optimize_penetration(
    sequence: str,
    environment: Dict[str, Any],
    residues: Optional[List[str] | str] = None,
    weights: Optional[Dict[str, float]] = None,
    n_decoys: int = 2000,
    n_random: int = 500,
    percent_min: int = 0,
    percent_max: int = 100,
    percent_step: int = 5,
    seed: int = 42,
) -> Dict[str, Any]:
    amino_acids = normalize_residues(residues)
    w = normalize_weights(weights)
    env_base = resolve_environment(environment)
    if not str(env_base["preset"]).startswith("interfacial_"):
        raise ValueError("Optimal penetration is only defined for interfacial presets.")
    if percent_step <= 0:
        raise ValueError("percent_step must be positive.")
    if percent_min > percent_max:
        raise ValueError("percent_min must be smaller than or equal to percent_max.")

    profile: List[Dict[str, Any]] = []
    best_result: Optional[Dict[str, Any]] = None
    best_percent: Optional[int] = None

    for percent in range(percent_min, percent_max + 1, percent_step):
        env = penetration_percent_to_environment(percent, str(env_base["preset"]))
        scored = score_sequence(
            sequence=sequence,
            environment=env,
            residues=amino_acids,
            weights=w,
            n_decoys=n_decoys,
            n_random=n_random,
            seed=seed,
        )
        profile.append(
            {
                "penetration_percent": percent,
                "wheel_halfwidth_deg": env.get("wheel_halfwidth_deg"),
                "environment_preset": env["preset"],
                "penetration_regime": env.get("penetration_regime", "interfacial"),
                "energy": scored["energy"],
                "z_score": scored["z_score"],
                "percentile_lower_is_better": scored["percentile_lower_is_better"],
            }
        )
        if best_result is None or scored["z_score"] < best_result["z_score"]:
            best_result = scored
            best_percent = percent

    assert best_result is not None and best_percent is not None
    return {
        "sequence": sequence.strip().upper(),
        "interface_preset": env_base["preset"],
        "optimization_target": "minimum_z_score",
        "best_penetration_percent": best_percent,
        "best_score": best_result,
        "profile": profile,
        "scan_settings": {
            "percent_min": percent_min,
            "percent_max": percent_max,
            "percent_step": percent_step,
            "n_random": n_random,
            "n_decoys": n_decoys,
            "seed": seed,
        },
    }


def cross_environment_design(
    length: int,
    environment_a: Dict[str, Any],
    environment_b: Dict[str, Any],
    lambda_gap: float,
    residues: Optional[List[str] | str] = None,
    weights: Optional[Dict[str, float]] = None,
    n_decoys: int = 2000,
    steps: int = 16000,
    restarts: int = 10,
    seed: int = 42,
) -> Dict[str, Any]:
    amino_acids = normalize_residues(residues)
    w = normalize_weights(weights)
    env_a = resolve_environment(environment_a)
    env_b = resolve_environment(environment_b)
    design_a, scoring_a = report_models(length, amino_acids, env_a, w, n_decoys=n_decoys, seed=seed, n_random=DESIGN_N_RANDOM)
    design_b, scoring_b = report_models(length, amino_acids, env_b, w, n_decoys=n_decoys, seed=seed, n_random=DESIGN_N_RANDOM)
    builder_a, builder_b = design_a.builder, design_b.builder
    rng = np.random.default_rng(rng_seed(seed, SEARCH_STREAM))
    n_aa = len(amino_acids)
    best_codes = None
    best_obj = float("inf")
    best_a = 0.0
    best_b = 0.0

    for _ in range(restarts):
        current = rng.integers(0, n_aa, size=length).tolist()
        current_a = compute_breakdown(builder_a, current, w)["total"]
        current_b = compute_breakdown(builder_b, current, w)["total"]
        current_obj = current_a + current_b + lambda_gap * abs(current_a - current_b)
        local_best = list(current)
        local_obj = current_obj
        local_a = current_a
        local_b = current_b

        for step in range(steps):
            temperature = max(1e-4, 2.0 * (1.0 - step / steps))
            proposal = list(current)
            pos = int(rng.integers(0, length))
            old_code = proposal[pos]
            new_code = int(rng.integers(0, n_aa - 1))
            if new_code >= old_code:
                new_code += 1
            proposal[pos] = new_code
            prop_a = compute_breakdown(builder_a, proposal, w)["total"]
            prop_b = compute_breakdown(builder_b, proposal, w)["total"]
            prop_obj = prop_a + prop_b + lambda_gap * abs(prop_a - prop_b)
            delta = prop_obj - current_obj
            if delta <= 0 or rng.random() < math.exp(-delta / temperature):
                current = proposal
                current_obj = prop_obj
                current_a = prop_a
                current_b = prop_b
                if current_obj < local_obj:
                    local_best = list(current)
                    local_obj = current_obj
                    local_a = current_a
                    local_b = current_b

        if local_obj < best_obj:
            best_codes = list(local_best)
            best_obj = local_obj
            best_a = local_a
            best_b = local_b

    assert best_codes is not None
    sequence = codes_to_seq(best_codes, amino_acids)
    # La búsqueda y Compare usan la misma regla de orientación en cada entorno.
    scored_a = report_sequence(sequence, design_a, scoring_a, amino_acids, w)
    scored_b = report_sequence(sequence, design_b, scoring_b, amino_acids, w)
    comparison = {
        "sequence": sequence,
        "environment_a": scored_a,
        "environment_b": scored_b,
        "energy_gap": abs(scored_a["energy"] - scored_b["energy"]),
        "z_score_gap": abs(scored_a["z_score"] - scored_b["z_score"]),
        "joint_energy": scored_a["energy"] + scored_b["energy"],
    }
    comparison["cross_design"] = {
        "lambda_gap": lambda_gap,
        "objective": best_obj,
        "energy_a": best_a,
        "energy_b": best_b,
        "steps": steps,
        "restarts": restarts,
        "seed": seed,
    }
    return comparison


def design_specificity(
    length: int,
    target_environment: Dict[str, Any],
    off_target_environments: List[Dict[str, Any]],
    residues: Optional[List[str] | str] = None,
    lambda_balance: float = 0.0,
    weights: Optional[Dict[str, float]] = None,
    n_decoys: int = 10000,
    steps: int = 60000,
    restarts: int = 20,
    num_sequences: int = 1,
    seed: int = 42,
) -> Dict[str, Any]:
    final_update = None
    for update in design_specificity_stream(
        length=length,
        target_environment=target_environment,
        off_target_environments=off_target_environments,
        residues=residues,
        lambda_balance=lambda_balance,
        weights=weights,
        n_decoys=n_decoys,
        steps=steps,
        restarts=restarts,
        num_sequences=num_sequences,
        seed=seed,
    ):
        if update["type"] == "final":
            final_update = update["payload"]
    if final_update is None:
        raise RuntimeError("Specificity design did not produce a final result")
    return final_update


def design_specificity_stream(
    length: int,
    target_environment: Dict[str, Any],
    off_target_environments: List[Dict[str, Any]],
    residues: Optional[List[str] | str] = None,
    lambda_balance: float = 0.0,
    weights: Optional[Dict[str, float]] = None,
    n_decoys: int = 10000,
    steps: int = 60000,
    restarts: int = 20,
    num_sequences: int = 1,
    seed: int = 42,
) -> Generator[Dict[str, Any], None, None]:
    amino_acids = normalize_residues(residues)
    w = normalize_weights(weights)
    target_env = resolve_environment(target_environment)
    off_target_envs = [resolve_environment(env) for env in off_target_environments]
    # Se descartan off-targets físicamente idénticos entre sí o al objetivo (mismo Hamiltoniano).
    seen: set[tuple[Any, ...]] = {physical_environment_key(target_env)}
    unique_off_targets: List[Dict[str, Any]] = []
    for env in off_target_envs:
        key = physical_environment_key(env)
        if key not in seen:
            seen.add(key)
            unique_off_targets.append(env)
    off_target_envs = unique_off_targets
    if not off_target_envs:
        raise ValueError("Please provide at least one off-target environment different from the target.")

    lambda_balance = float(lambda_balance)
    if not (0.0 <= lambda_balance <= 1.0):
        raise ValueError("lambda_balance must lie between 0 and 1.")
    num_sequences = max(1, int(num_sequences))
    total_restart_budget = max(1, int(restarts))
    if num_sequences > total_restart_budget:
        raise ValueError("num_sequences cannot exceed restarts: each candidate needs at least one restart.")
    restarts_per_attempt = max(1, total_restart_budget // num_sequences)
    total_attempts = max(1, total_restart_budget // restarts_per_attempt)

    yield {
        "type": "meta",
        "payload": {
            "target_environment": target_env,
            "off_target_environments": off_target_envs,
            "alphabet_size": len(amino_acids),
            "steps": steps,
            "restarts": restarts_per_attempt,
            "total_restart_budget": total_restart_budget,
            "restarts_per_attempt": restarts_per_attempt,
            "num_sequences": num_sequences,
            "lambda_balance": lambda_balance,
            "phase": "initializing",
            "progress_fraction": 0.0,
        },
    }

    builder_specs = [("target", target_env)] + [(f"off_target_{idx + 1}", env) for idx, env in enumerate(off_target_envs)]
    models: Dict[str, tuple[ScoringModel, ScoringModel]] = {}
    built: Dict[str, HamiltonianBuilder] = {}
    total_builders = len(builder_specs)
    for idx, (label, env) in enumerate(builder_specs, start=1):
        models[label] = report_models(
            length, amino_acids, env, w, n_decoys=n_decoys, seed=seed, n_random=SPECIFICITY_N_RANDOM
        )
        built[label] = models[label][0].builder
        yield {
            "type": "phase",
            "payload": {
                "phase": "calibrating",
                "label": label,
                "completed": idx,
                "total": total_builders,
                "progress_fraction": 0.1 * (idx / max(1, total_builders)),
            },
        }

    target_builder = built["target"]
    off_builders = [built[f"off_target_{idx + 1}"] for idx in range(len(off_target_envs))]
    unique_results: Dict[str, Dict[str, Any]] = {}
    accepted_codes: List[List[int]] = []
    min_distance = max(2, math.ceil(length * 0.15))
    penalty_weight = 6.0

    for attempt_idx in range(total_attempts):
        yield {
            "type": "phase",
            "payload": {
                "phase": "collecting",
                "completed": len(unique_results),
                "total": num_sequences,
                "attempt": attempt_idx + 1,
                "attempts_total": total_attempts,
                "progress_fraction": 0.1 + 0.55 * (attempt_idx / max(1, total_attempts)),
            },
        }

        result: Optional[Dict[str, Any]] = None
        for update in _iter_specificity_search(
            target_builder=target_builder,
            off_builders=off_builders,
            amino_acids=amino_acids,
            weights=w,
            lambda_balance=lambda_balance,
            steps=steps,
            restarts=restarts_per_attempt,
            seed=rng_seed(seed, SEARCH_STREAM, attempt_idx),
            accepted_codes=accepted_codes,
            min_distance=min_distance,
            penalty_weight=penalty_weight,
        ):
            if update["type"] == "progress":
                inner = update["payload"]
                inner_fraction = float(inner.get("progress_fraction", 0.0))
                overall = 0.1 + 0.55 * ((attempt_idx + inner_fraction) / max(1, total_attempts))
                yield {
                    "type": "progress",
                    "payload": {
                        **inner,
                        "phase": "collecting",
                        "accepted": len(unique_results),
                        "target_count": num_sequences,
                        "attempt": attempt_idx + 1,
                        "attempts_total": total_attempts,
                        "progress_fraction": overall,
                    },
                }
            elif update["type"] == "result":
                result = update["payload"]

        if result is None:
            raise RuntimeError("Specificity search attempt did not return a result")

        sequence = result["sequence"]
        if sequence in unique_results:
            continue

        candidate_payload = _build_specificity_candidate_payload(
            sequence,
            length=length,
            amino_acids=amino_acids,
            target_env=target_env,
            off_target_envs=off_target_envs,
            target_models=models["target"],
            off_models=[models[f"off_target_{idx + 1}"] for idx in range(len(off_target_envs))],
            weights=w,
            lambda_balance=lambda_balance,
            steps=steps,
            restarts=restarts_per_attempt,
            seed=seed,
            attempt=attempt_idx + 1,
            objective=result["objective"],
            best_target_energy=result["best_target_energy"],
            best_off_energies=result["best_off_energies"],
        )
        unique_results[sequence] = candidate_payload
        accepted_codes.append(result["codes"])
        yield {
            "type": "candidate",
            "payload": {
                "accepted": len(unique_results),
                "target": num_sequences,
                "attempt": attempt_idx + 1,
                "sequence": sequence,
                "objective": candidate_payload["specificity_design"]["objective"],
                "z_score_margin": candidate_payload["specificity_design"]["z_score_margin"],
                "member": candidate_payload,
            },
        }
        if len(unique_results) >= num_sequences:
            break

    ranked_candidates = sorted(
        unique_results.values(),
        key=lambda item: (
            round(float(item["specificity_design"]["objective"]), 8),
            -float(item["specificity_design"]["z_score_margin"]),
            float(item["target_environment"]["design_geometry"]["z_score"]),
            item["sequence"],
        ),
    )
    if not ranked_candidates:
        raise RuntimeError("Specificity design did not recover any candidate sequence.")

    for idx, candidate in enumerate(ranked_candidates, start=1):
        candidate["specificity_design"]["rank"] = idx

    top_candidate = ranked_candidates[0]
    payload = {
        **top_candidate,
        "candidates": ranked_candidates,
        "specificity_collection": {
            "requested": num_sequences,
            "returned": len(ranked_candidates),
            "attempts_budget": total_attempts,
            "total_restart_budget": total_restart_budget,
            "restarts_per_attempt": restarts_per_attempt,
            "lambda_balance": lambda_balance,
            "target_preset": target_env["preset"],
            "off_target_presets": [env["preset"] for env in off_target_envs],
        },
    }
    yield {"type": "final", "payload": payload}
def generate_helix_pdb(sequence: str) -> str:
    sequence = sequence.strip().upper()
    if not sequence:
        raise RuntimeError("Sequence cannot be empty")
    for aa in sequence:
        if aa not in THREE_LETTER_CODES:
            raise RuntimeError(f"Unsupported residue for PDB export: {aa}")

    try:
        import PeptideBuilder
        from Bio.PDB import PDBIO
    except Exception as exc:
        raise RuntimeError(
            "PDB export requires PeptideBuilder and Biopython in the application environment."
        ) from exc

    phi = [-57.0] * max(0, len(sequence) - 1)
    psi = [-47.0] * max(0, len(sequence) - 1)
    structure = PeptideBuilder.make_structure(sequence, phi, psi)

    with tempfile.TemporaryDirectory(prefix="helix_pdb_") as tmpdir:
        out_path = Path(tmpdir) / "helix.pdb"
        io = PDBIO()
        io.set_structure(structure)
        io.save(str(out_path))
        pdb_text = out_path.read_text()
    if pdb_text.strip():
        return pdb_text if pdb_text.endswith("\n") else pdb_text + "\n"
    raise RuntimeError("Failed to generate PDB for the requested sequence.")
```
