## Additional file 1. Evaluated software source and dependencies for "Helix Design Studio: designing antimicrobial peptide candidates for membrane environments": main.py

```
from __future__ import annotations

import asyncio
import hashlib
import hmac
import json
import logging
import os
import platform
import secrets
import threading
import tempfile
import time
from collections import deque
from datetime import datetime, timedelta, timezone
from pathlib import Path
from urllib.parse import urlsplit
from typing import Any, AsyncIterator, Callable, Deque, Dict, Iterator, List, Optional

from fastapi import FastAPI, HTTPException, Request
from fastapi.exceptions import RequestValidationError
from fastapi.middleware.cors import CORSMiddleware
from fastapi.responses import FileResponse, JSONResponse, StreamingResponse
from fastapi.staticfiles import StaticFiles
from starlette.concurrency import run_in_threadpool
from pydantic import BaseModel, ConfigDict, Field, field_validator, model_validator

from .engine import (
    CalculationCancelled,
    CalculationLimits,
    CalculationTimeout,
    DEFAULT_ALPHABET_16,
    DEFAULT_WEIGHTS,
    ENVIRONMENT_PRESETS,
    SEED_PROTOCOL,
    calculation_limits,
    compare_environments,
    cross_environment_design,
    design_specificity,
    design_specificity_stream,
    design_family,
    design_family_stream,
    design_sequence,
    generate_helix_pdb,
    optimize_penetration,
    physical_environment_key,
    score_sequence,
)
from core.hamiltonian_builder import parameter_set_sha256  # noqa: E402  (ruta añadida por .engine)


APP_DIR = Path(__file__).resolve().parent
STATIC_DIR = APP_DIR / "static"
USAGE_LOGGING_ENABLED = os.environ.get("HELIX_USAGE_LOGGING", "0") == "1"
VAR_DIR = Path(
    os.environ.get("HELIX_VAR_DIR")
    or Path(tempfile.gettempdir()) / "helix-design-studio-usage"
).expanduser()
USAGE_LOG_PATH = VAR_DIR / "usage_events.jsonl"

CORS_ORIGINS = [
    origin.strip()
    for origin in os.environ.get("HELIX_CORS_ORIGINS", "https://simbios.usc.es").split(",")
    if origin.strip()
]
MAX_CONCURRENT_JOBS = max(1, int(os.environ.get("HELIX_MAX_CONCURRENT_JOBS", "4")))
MAX_JOB_SECONDS = float(os.environ.get("HELIX_MAX_JOB_SECONDS", "1800"))
MAX_REQUEST_BYTES = 64 * 1024
DISCONNECT_POLL_SECONDS = 1.0
MAX_LOGGED_TEXT = 200
CONTENT_SECURITY_POLICY = (
    "default-src 'self'; script-src 'self'; style-src 'self' 'unsafe-inline'; "
    "img-src 'self' data: blob:; connect-src 'self'; font-src 'self' data:; worker-src 'self' blob:; "
    "object-src 'none'; base-uri 'self'; form-action 'self'; frame-ancestors 'none'"
)
# Clave para los hashes del registro de uso: sin ella, los hashes de IPs o secuencias cortas
# serian reversibles por fuerza bruta. Sin HELIX_LOG_HASH_KEY se genera una por proceso.
LOG_HASH_KEY = (os.environ.get("HELIX_LOG_HASH_KEY") or secrets.token_hex(32)).encode("utf-8")

logger = logging.getLogger("helix_design_studio")


class RequestLimitsMiddleware:
    """Rechaza cuerpos grandes antes de parsearlos y añade cabeceras de seguridad."""

    def __init__(self, app: Any) -> None:
        self.app = app

    async def __call__(self, scope: Dict[str, Any], receive: Any, send: Any) -> None:
        if scope["type"] != "http":
            await self.app(scope, receive, send)
            return
        length = dict(scope.get("headers") or []).get(b"content-length")
        if length is not None and (not length.isdigit() or int(length) > MAX_REQUEST_BYTES):
            await JSONResponse({"detail": "Request body too large."}, status_code=413)(scope, receive, send)
            return
        received = 0

        async def limited_receive() -> Dict[str, Any]:
            nonlocal received
            message = await receive()
            if message["type"] == "http.request":
                received += len(message.get("body", b""))
                if received > MAX_REQUEST_BYTES:
                    # HTTPException: FastAPI convierte otras excepciones del cuerpo en un 400 genérico.
                    raise HTTPException(status_code=413, detail="Request body too large.")
            return message

        async def secured_send(message: Dict[str, Any]) -> None:
            if message["type"] == "http.response.start":
                headers = list(message.get("headers", []))
                headers += [
                    (b"x-content-type-options", b"nosniff"),
                    (b"referrer-policy", b"strict-origin-when-cross-origin"),
                    (b"x-frame-options", b"DENY"),
                ]
                if not scope["path"].startswith(("/docs", "/redoc")):
                    headers.append((b"content-security-policy", CONTENT_SECURITY_POLICY.encode("ascii")))
                message = {**message, "headers": headers}
            await send(message)

        await self.app(scope, limited_receive, secured_send)


APP_VERSION = "1.0.0rc6"
PARAMETERS_SHA256 = parameter_set_sha256()

app = FastAPI(title="Helix Design Studio", version=APP_VERSION)
app.add_middleware(RequestLimitsMiddleware)

if CORS_ORIGINS:
    app.add_middleware(
        CORSMiddleware,
        allow_origins=CORS_ORIGINS,
        allow_methods=["GET", "POST", "OPTIONS"],
        allow_headers=["*"],
    )

app.mount("/static", StaticFiles(directory=str(STATIC_DIR)), name="static")


@app.exception_handler(RequestValidationError)
async def validation_error_handler(request: Request, exc: RequestValidationError) -> JSONResponse:
    errors = [
        {"loc": list(error.get("loc", ())), "msg": str(error.get("msg", "Invalid value")), "type": error.get("type")}
        for error in exc.errors()
    ]
    return JSONResponse(status_code=422, content={"detail": errors})


MAX_SEQUENCE_LENGTH = 40
MAX_ALLOWED_RESIDUES = 20
MIN_ALLOWED_RESIDUES = 2
MAX_RANDOM_REFERENCE = 2000
MAX_N_DECOYS = 10000
MAX_DESIGN_STEPS = 50000
MAX_FAMILY_STEPS = 20000
MAX_CROSS_STEPS = 60000
MAX_RESTARTS = 20
MAX_FAMILY_SIZE = 12
MAX_SPECIFICITY_CANDIDATES = 10
MAX_OVERSAMPLE_FACTOR = 5
MAX_PENETRATION_SCAN_POINTS = 21
MAX_OFF_TARGET_ENVIRONMENTS = len(ENVIRONMENT_PRESETS) - 1
MAX_WEIGHT = 100.0
MAX_SEED = 2**32 - 1
MEMBRANE_CHARGES = ("neg", "neu", "pos")
SIMBIOS_CONTACT_MSG = "For heavier runs or custom large-scale calculations, please contact SIMBIOS via https://simbios.usc.es/."
FULL_ALPHABET_20 = ["A", "R", "N", "D", "C", "Q", "E", "G", "H", "I", "L", "K", "M", "F", "P", "S", "T", "W", "Y", "V"]
STANDARD_RESIDUES = frozenset(FULL_ALPHABET_20)
BUSY_MSG = "The server is busy with other calculations. Please try again in a few minutes."

_job_slots = threading.BoundedSemaphore(MAX_CONCURRENT_JOBS)


def _clean_sequence(value: str) -> str:
    seq = str(value or "").strip().upper()
    if not seq:
        raise ValueError("Sequence cannot be empty.")
    if len(seq) > MAX_SEQUENCE_LENGTH:
        raise ValueError(
            f"Sequence length exceeds the current web limit of {MAX_SEQUENCE_LENGTH} residues. {SIMBIOS_CONTACT_MSG}"
        )
    invalid = sorted(set(seq) - STANDARD_RESIDUES)
    if invalid:
        raise ValueError(
            f"Sequence contains unsupported residue codes ({', '.join(invalid)}). Use the 20 standard one-letter codes."
        )
    return seq


def _validate_residues(value: List[str]) -> List[str]:
    residues = [str(aa).strip().upper() for aa in value if str(aa).strip()]
    invalid = sorted(set(residues) - STANDARD_RESIDUES)
    if invalid:
        raise ValueError(
            f"Allowed residues must be standard one-letter codes; unsupported: {', '.join(invalid)}."
        )
    unique: List[str] = []
    for aa in residues:
        if aa not in unique:
            unique.append(aa)
    if len(unique) < MIN_ALLOWED_RESIDUES:
        raise ValueError("Please provide at least two distinct allowed residue types.")
    if len(unique) > MAX_ALLOWED_RESIDUES:
        raise ValueError(
            f"Allowed-residue sets are limited to {MAX_ALLOWED_RESIDUES} residue types in the web app. {SIMBIOS_CONTACT_MSG}"
        )
    return unique


def _hash_value(value: str) -> str:
    return hmac.new(LOG_HASH_KEY, value.encode("utf-8"), hashlib.sha256).hexdigest()[:16]


def _classify_error(exc: Exception) -> tuple[int, str]:
    """ValueError describes an invalid request; anything else is an internal failure."""
    if isinstance(exc, ValueError):
        return 400, str(exc)[:MAX_LOGGED_TEXT * 2]
    if isinstance(exc, CalculationTimeout):
        return 400, f"{exc} Please reduce steps, restarts or requested sequences. {SIMBIOS_CONTACT_MSG}"
    logger.error("Calculation failed", exc_info=exc)
    return 500, "Internal error while running the calculation."


def _acquire_job_slot() -> None:
    if not _job_slots.acquire(blocking=False):
        raise HTTPException(status_code=503, detail=BUSY_MSG)


def _sequence_summary(sequence: Optional[str]) -> Dict[str, Any]:
    if not sequence:
        return {}
    seq = str(sequence).strip().upper()
    if not seq:
        return {}
    return {
        "sequence_length": len(seq),
        "sequence_hash": _hash_value(seq),
    }


def _environment_summary(environment: Optional[Dict[str, Any]], prefix: str = "environment") -> Dict[str, Any]:
    if not environment:
        return {}
    return {
        f"{prefix}_preset": environment.get("preset"),
        f"{prefix}_membrane_charge": environment.get("membrane_charge"),
        f"{prefix}_wheel_halfwidth_deg": environment.get("wheel_halfwidth_deg"),
    }


def _client_ip_from_request(request: Request) -> str:
    # X-Forwarded-For solo es fiable si lo reescribe un proxy de confianza; uvicorn lo aplica
    # a request.client con --proxy-headers/--forwarded-allow-ips.
    if request.client and request.client.host:
        return request.client.host
    return "unknown"


def _url_origin(url: str) -> Optional[str]:
    """Solo esquema y host: la ruta o la query de un referer pueden contener datos privados."""
    try:
        parts = urlsplit(url[:2048])
    except ValueError:  # p. ej. "https://[invalid"
        return None
    return f"{parts.scheme}://{parts.netloc}"[:MAX_LOGGED_TEXT] if parts.scheme and parts.netloc else None


def _usage_context(request: Request) -> Dict[str, Any]:
    anonymous_id = request.headers.get("x-anonymous-id", "").strip()[:64] or None
    user_agent = request.headers.get("user-agent", "").strip() or "unknown"
    client_ip = _client_ip_from_request(request)
    return {
        "anonymous_user_id": anonymous_id,
        "user_agent_hash": _hash_value(user_agent),
        "client_ip_hash": _hash_value(client_ip),
        "origin": request.headers.get("origin", "")[:MAX_LOGGED_TEXT] or None,
        "referer": _url_origin(request.headers.get("referer", "")),
    }


def _request_summary(payload: Any) -> Dict[str, Any]:
    data = payload.model_dump() if hasattr(payload, "model_dump") else dict(payload)
    summary: Dict[str, Any] = {}
    summary.update(_sequence_summary(data.get("sequence")))
    if "length" in data:
        summary["length"] = data["length"]
    if "residues" in data:
        summary["alphabet_size"] = len(data["residues"])
    if "steps" in data:
        summary["steps"] = data["steps"]
    if "restarts" in data:
        summary["restarts"] = data["restarts"]
    if "family_size" in data:
        summary["family_size"] = data["family_size"]
    if "num_sequences" in data:
        summary["num_sequences"] = data["num_sequences"]
    if "n_random" in data:
        summary["n_random"] = data["n_random"]
    if "lambda_gap" in data:
        summary["lambda_gap"] = data["lambda_gap"]
    if "lambda_balance" in data:
        summary["lambda_balance"] = data["lambda_balance"]
    if "percent_min" in data:
        summary["percent_min"] = data["percent_min"]
    if "percent_max" in data:
        summary["percent_max"] = data["percent_max"]
    if "percent_step" in data:
        summary["percent_step"] = data["percent_step"]
    if "environment" in data:
        summary.update(_environment_summary(data.get("environment"), "environment"))
    if "environment_a" in data:
        summary.update(_environment_summary(data.get("environment_a"), "environment_a"))
    if "environment_b" in data:
        summary.update(_environment_summary(data.get("environment_b"), "environment_b"))
    return summary


def _directory_is_private(path: Path) -> bool:
    """Propio del usuario del servicio y no escribible por otros (el valor por defecto está en /tmp)."""
    info = path.stat()
    if hasattr(os, "getuid") and info.st_uid != os.getuid():
        return False
    return not info.st_mode & 0o022


def _append_usage_event(event: Dict[str, Any]) -> None:
    if not USAGE_LOGGING_ENABLED:
        return
    # Un fallo del registro de uso no debe hacer fallar la peticion del usuario.
    try:
        VAR_DIR.mkdir(mode=0o700, parents=True, exist_ok=True)
        if not _directory_is_private(VAR_DIR):
            logger.warning("Usage log directory is not private to this user; event not written: %s", VAR_DIR)
            return
        flags = os.O_WRONLY | os.O_CREAT | os.O_APPEND | getattr(os, "O_NOFOLLOW", 0)
        descriptor = os.open(USAGE_LOG_PATH, flags, 0o600)
        with os.fdopen(descriptor, "a", encoding="utf-8") as handle:
            handle.write(json.dumps(event, ensure_ascii=True) + "\n")
    except OSError as exc:
        logger.warning("Could not write usage event: %s", exc)


DEFAULT_HEALTH_WINDOW_MINUTES = 60
MAX_HEALTH_WINDOW_MINUTES = 60 * 24 * 7


class UsageLogIndex:
    """Agregados del registro de uso que se actualizan leyendo solo las líneas nuevas.

    /api/visit, /api/usage-metrics y /api/health se consultan en cada carga de página; releer
    el fichero completo en cada petición crecería linealmente con el historial.
    """

    def __init__(self, path: Path) -> None:
        self.path = path
        self.lock = threading.Lock()
        self._reset(None)

    def _reset(self, file_id: Optional[tuple[int, int]]) -> None:
        self.file_id = file_id
        self.offset = 0
        self.total_visits = 0
        self.unique_visitors: set[str] = set()
        self.total_runs = 0
        # (instante, endpoint, fallo del servidor) de la ventana máxima de /api/health.
        self.recent: Deque[tuple[datetime, str, bool]] = deque()

    def _ingest(self, event: Dict[str, Any]) -> None:
        if event.get("status") == "ok":
            if event.get("endpoint") == "visit":
                self.total_visits += 1
                if event.get("anonymous_user_id"):
                    self.unique_visitors.add(str(event["anonymous_user_id"]))
            else:
                self.total_runs += 1
        try:
            stamp = datetime.fromisoformat(str(event["timestamp_utc"]))
        except (KeyError, ValueError):
            return
        if stamp.tzinfo is None:
            stamp = stamp.replace(tzinfo=timezone.utc)
        # Los errores de validacion del cliente no indican un fallo del servicio.
        try:
            http_status = int(event.get("http_status", 500))
        except (TypeError, ValueError):
            http_status = 500
        server_error = event.get("status") == "error" and http_status >= 500
        self.recent.append((stamp, str(event.get("endpoint", "unknown")), server_error))

    def refresh(self) -> None:
        try:
            info = self.path.stat()
        except OSError:
            self._reset(None)
            return
        file_id = (info.st_dev, info.st_ino)
        if file_id != self.file_id or info.st_size < self.offset:
            self._reset(file_id)  # fichero nuevo, rotado o truncado
        try:
            with self.path.open("rb") as handle:
                handle.seek(self.offset)
                for raw in handle:
                    if not raw.endswith(b"\n"):
                        break  # línea a medio escribir: se leerá en la próxima consulta
                    self.offset += len(raw)
                    try:
                        event = json.loads(raw)
                    except ValueError:
                        continue
                    if isinstance(event, dict):
                        self._ingest(event)
        except OSError as exc:
            logger.warning("Could not read usage log: %s", exc)
        cutoff = datetime.now(timezone.utc) - timedelta(minutes=MAX_HEALTH_WINDOW_MINUTES)
        while self.recent and self.recent[0][0] < cutoff:
            self.recent.popleft()

    def metrics(self) -> Dict[str, int]:
        with self.lock:
            self.refresh()
            return {
                "total_visits": self.total_visits,
                "unique_visitors": len(self.unique_visitors),
                "total_runs": self.total_runs,
            }

    def recent_summary(self, window_minutes: int) -> Dict[str, Any]:
        with self.lock:
            self.refresh()
            cutoff = datetime.now(timezone.utc) - timedelta(minutes=window_minutes)
            events = [item for item in self.recent if item[0] >= cutoff]
        failing: Dict[str, int] = {}
        last_error: Optional[Dict[str, Any]] = None
        for stamp, endpoint, server_error in events:
            if server_error:
                failing[endpoint] = failing.get(endpoint, 0) + 1
                last_error = {"timestamp_utc": stamp.isoformat(), "endpoint": endpoint}
        return {
            "window_minutes": window_minutes,
            "events": len(events),
            "errors": sum(failing.values()),
            "failing_endpoints": dict(sorted(failing.items(), key=lambda kv: (-kv[1], kv[0]))),
            "last_error": last_error,
        }


_usage_index = UsageLogIndex(USAGE_LOG_PATH)


def _usage_metrics() -> Dict[str, int]:
    if not USAGE_LOGGING_ENABLED:
        return {"total_visits": 0, "unique_visitors": 0, "total_runs": 0}
    return _usage_index.metrics()


def _recent_usage_summary(window_minutes: int) -> Dict[str, Any]:
    """Resume los eventos de uso de la ventana reciente, agrupados por endpoint.

    Se usa en /api/health: los endpoints de streaming capturan sus excepciones y
    las emiten dentro del stream, por lo que HTTP siempre responde 200 y un
    monitor externo basado en codigos de estado no puede detectar los fallos.
    """
    if not USAGE_LOGGING_ENABLED:
        return {"window_minutes": window_minutes, "events": 0, "errors": 0, "failing_endpoints": {}, "last_error": None}
    return _usage_index.recent_summary(window_minutes)


def _log_usage_event(
    request: Request,
    endpoint: str,
    payload: Any,
    status: str,
    duration_s: float,
    extra: Optional[Dict[str, Any]] = None,
) -> None:
    if not USAGE_LOGGING_ENABLED:
        return
    # El registro de uso nunca debe invalidar una petición (ni un cálculo ya terminado).
    try:
        event = {
            "timestamp_utc": datetime.now(timezone.utc).isoformat(),
            "endpoint": endpoint,
            "status": status,
            "duration_s": round(duration_s, 4),
            **_usage_context(request),
            **_request_summary(payload),
        }
        if extra:
            event.update(extra)
        _append_usage_event(event)
    except Exception:
        logger.warning("Could not record usage event", exc_info=True)


class RequestModel(BaseModel):
    # JSON acepta NaN/Infinity en Python; se rechazan para que no lleguen al modelo. Los campos
    # desconocidos (p. ej. erratas) se rechazan en lugar de ignorarse en silencio.
    model_config = ConfigDict(allow_inf_nan=False, extra="forbid")


def _weight_field(name: str) -> Any:
    return Field(default=DEFAULT_WEIGHTS[name], ge=0.0, le=MAX_WEIGHT)


def _seed_field() -> Any:
    return Field(default=42, ge=0, le=MAX_SEED)


class EnvironmentConfig(RequestModel):
    preset: str = "interfacial_neg"
    wheel_halfwidth_deg: Optional[float] = Field(default=None, ge=-1.0, le=180.0)
    membrane_charge: Optional[str] = None

    @field_validator("preset")
    @classmethod
    def validate_preset(cls, value: str) -> str:
        if value not in ENVIRONMENT_PRESETS:
            raise ValueError(f"Unknown environment preset. Choose one of: {', '.join(ENVIRONMENT_PRESETS)}.")
        return value

    @field_validator("membrane_charge")
    @classmethod
    def validate_membrane_charge(cls, value: Optional[str]) -> Optional[str]:
        if value is not None and value not in MEMBRANE_CHARGES:
            raise ValueError(f"membrane_charge must be one of: {', '.join(MEMBRANE_CHARGES)}.")
        return value


class WeightsConfig(RequestModel):
    lambda_env: float = _weight_field("lambda_env")
    lambda_charge: float = _weight_field("lambda_charge")
    lambda_local: float = _weight_field("lambda_local")
    lambda_pairwise: float = _weight_field("lambda_pairwise")
    lambda_helix_pairs: float = _weight_field("lambda_helix_pairs")
    lambda_electrostatic: float = _weight_field("lambda_electrostatic")


def _check_sequence_in_alphabet(sequence: str, residues: List[str]) -> None:
    missing = sorted(set(sequence) - set(residues))
    if missing:
        raise ValueError(f"Residues {', '.join(missing)} of the sequence are not in the allowed residues.")


def _environment_key(env: EnvironmentConfig) -> tuple[Any, ...]:
    return physical_environment_key(env.model_dump())


class ScoreRequest(RequestModel):
    sequence: str = Field(max_length=4 * MAX_SEQUENCE_LENGTH)
    residues: List[str] = Field(default_factory=lambda: list(FULL_ALPHABET_20), max_length=2 * MAX_ALLOWED_RESIDUES)
    environment: EnvironmentConfig = Field(default_factory=EnvironmentConfig)
    weights: WeightsConfig = Field(default_factory=WeightsConfig)
    n_decoys: int = Field(default=2000, ge=100, le=MAX_N_DECOYS)
    n_random: int = Field(default=500, ge=100, le=MAX_RANDOM_REFERENCE)
    seed: int = _seed_field()

    @field_validator("sequence")
    @classmethod
    def validate_sequence(cls, value: str) -> str:
        return _clean_sequence(value)

    @field_validator("residues")
    @classmethod
    def validate_residues(cls, value: List[str]) -> List[str]:
        return _validate_residues(value)

    @model_validator(mode="after")
    def validate_sequence_alphabet(self) -> "ScoreRequest":
        _check_sequence_in_alphabet(self.sequence, self.residues)
        return self


class DesignRequest(RequestModel):
    length: int = Field(default=20, ge=4, le=MAX_SEQUENCE_LENGTH)
    residues: List[str] = Field(default_factory=lambda: list(FULL_ALPHABET_20), max_length=2 * MAX_ALLOWED_RESIDUES)
    environment: EnvironmentConfig = Field(default_factory=EnvironmentConfig)
    weights: WeightsConfig = Field(default_factory=WeightsConfig)
    n_decoys: int = Field(default=2000, ge=100, le=MAX_N_DECOYS)
    steps: int = Field(default=12000, ge=1000, le=MAX_DESIGN_STEPS)
    restarts: int = Field(default=8, ge=1, le=MAX_RESTARTS)
    seed: int = _seed_field()

    @field_validator("residues")
    @classmethod
    def validate_residues(cls, value: List[str]) -> List[str]:
        return _validate_residues(value)


class CompareRequest(RequestModel):
    sequence: str = Field(max_length=4 * MAX_SEQUENCE_LENGTH)
    residues: List[str] = Field(default_factory=lambda: list(FULL_ALPHABET_20), max_length=2 * MAX_ALLOWED_RESIDUES)
    environment_a: EnvironmentConfig
    environment_b: EnvironmentConfig
    weights: WeightsConfig = Field(default_factory=WeightsConfig)
    n_decoys: int = Field(default=2000, ge=100, le=MAX_N_DECOYS)
    seed: int = _seed_field()

    @field_validator("sequence")
    @classmethod
    def validate_sequence(cls, value: str) -> str:
        return _clean_sequence(value)

    @field_validator("residues")
    @classmethod
    def validate_residues(cls, value: List[str]) -> List[str]:
        return _validate_residues(value)

    @model_validator(mode="after")
    def validate_sequence_alphabet(self) -> "CompareRequest":
        _check_sequence_in_alphabet(self.sequence, self.residues)
        return self


class FamilyDesignRequest(RequestModel):
    length: int = Field(default=20, ge=4, le=MAX_SEQUENCE_LENGTH)
    residues: List[str] = Field(default_factory=lambda: list(DEFAULT_ALPHABET_16), max_length=2 * MAX_ALLOWED_RESIDUES)
    environment: EnvironmentConfig = Field(default_factory=EnvironmentConfig)
    weights: WeightsConfig = Field(default_factory=WeightsConfig)
    n_decoys: int = Field(default=2000, ge=100, le=MAX_N_DECOYS)
    steps: int = Field(default=10000, ge=1000, le=MAX_FAMILY_STEPS)
    restarts: int = Field(default=6, ge=1, le=MAX_RESTARTS)
    family_size: int = Field(default=8, ge=2, le=MAX_FAMILY_SIZE)
    oversample_factor: int = Field(default=3, ge=1, le=MAX_OVERSAMPLE_FACTOR)
    seed: int = _seed_field()

    @field_validator("residues")
    @classmethod
    def validate_residues(cls, value: List[str]) -> List[str]:
        return _validate_residues(value)


class CrossDesignRequest(RequestModel):
    length: int = Field(default=20, ge=4, le=MAX_SEQUENCE_LENGTH)
    residues: List[str] = Field(default_factory=lambda: list(DEFAULT_ALPHABET_16), max_length=2 * MAX_ALLOWED_RESIDUES)
    environment_a: EnvironmentConfig
    environment_b: EnvironmentConfig
    lambda_gap: float = Field(default=0.5, ge=0.0, le=10.0)
    weights: WeightsConfig = Field(default_factory=WeightsConfig)
    n_decoys: int = Field(default=2000, ge=100, le=MAX_N_DECOYS)
    steps: int = Field(default=16000, ge=1000, le=MAX_CROSS_STEPS)
    restarts: int = Field(default=10, ge=1, le=MAX_RESTARTS)
    seed: int = _seed_field()

    @field_validator("residues")
    @classmethod
    def validate_residues(cls, value: List[str]) -> List[str]:
        return _validate_residues(value)


class SpecificityDesignRequest(RequestModel):
    length: int = Field(default=20, ge=4, le=MAX_SEQUENCE_LENGTH)
    residues: List[str] = Field(default_factory=lambda: list(FULL_ALPHABET_20), max_length=2 * MAX_ALLOWED_RESIDUES)
    target_environment: EnvironmentConfig
    off_target_environments: List[EnvironmentConfig] = Field(max_length=MAX_OFF_TARGET_ENVIRONMENTS)
    num_sequences: int = Field(default=10, ge=1, le=MAX_SPECIFICITY_CANDIDATES)
    lambda_balance: float = Field(default=0.0, ge=0.0, le=1.0)
    weights: WeightsConfig = Field(default_factory=WeightsConfig)
    n_decoys: int = Field(default=10000, ge=100, le=MAX_N_DECOYS)
    steps: int = Field(default=60000, ge=1000, le=MAX_CROSS_STEPS)
    restarts: int = Field(default=20, ge=1, le=MAX_RESTARTS)
    seed: int = _seed_field()

    @field_validator("residues")
    @classmethod
    def validate_residues(cls, value: List[str]) -> List[str]:
        return _validate_residues(value)

    @field_validator("off_target_environments")
    @classmethod
    def validate_off_targets(cls, value: List[EnvironmentConfig]) -> List[EnvironmentConfig]:
        if not value:
            raise ValueError("Please provide at least one off-target environment.")
        return value

    @model_validator(mode="after")
    def validate_target_not_in_off_targets(self) -> "SpecificityDesignRequest":
        target_key = _environment_key(self.target_environment)
        if any(_environment_key(env) == target_key for env in self.off_target_environments):
            raise ValueError(
                "An off-target environment is physically identical to the target (same sector and membrane charge)."
            )
        if self.num_sequences > self.restarts:
            raise ValueError("num_sequences cannot exceed restarts: each candidate needs at least one restart.")
        return self


class PenetrationOptimizationRequest(RequestModel):
    sequence: str = Field(max_length=4 * MAX_SEQUENCE_LENGTH)
    residues: List[str] = Field(default_factory=lambda: list(FULL_ALPHABET_20), max_length=2 * MAX_ALLOWED_RESIDUES)
    environment: EnvironmentConfig
    weights: WeightsConfig = Field(default_factory=WeightsConfig)
    n_decoys: int = Field(default=2000, ge=100, le=MAX_N_DECOYS)
    n_random: int = Field(default=500, ge=100, le=MAX_RANDOM_REFERENCE)
    percent_min: int = Field(default=0, ge=0, le=100)
    percent_max: int = Field(default=100, ge=0, le=100)
    percent_step: int = Field(default=5, ge=1, le=20)
    seed: int = _seed_field()

    @field_validator("sequence")
    @classmethod
    def validate_sequence(cls, value: str) -> str:
        return _clean_sequence(value)

    @field_validator("residues")
    @classmethod
    def validate_residues(cls, value: List[str]) -> List[str]:
        return _validate_residues(value)

    @field_validator("environment")
    @classmethod
    def validate_environment(cls, value: EnvironmentConfig) -> EnvironmentConfig:
        if not str(value.preset).startswith("interfacial_"):
            raise ValueError("Optimal penetration can only be scanned for interfacial presets.")
        return value

    @model_validator(mode="after")
    def validate_scan_geometry(self) -> "PenetrationOptimizationRequest":
        if self.percent_min != 0 or self.percent_max != 100 or self.percent_step != 5:
            raise ValueError("Web penetration scans are fixed to 0-100% in 5% increments.")
        n_points = ((self.percent_max - self.percent_min) // self.percent_step) + 1
        if n_points > MAX_PENETRATION_SCAN_POINTS:
            raise ValueError(
                f"Penetration scans are limited to {MAX_PENETRATION_SCAN_POINTS} sampled depths in the web app. {SIMBIOS_CONTACT_MSG}"
            )
        return self

    @model_validator(mode="after")
    def validate_sequence_alphabet(self) -> "PenetrationOptimizationRequest":
        _check_sequence_in_alphabet(self.sequence, self.residues)
        return self


class PdbRequest(RequestModel):
    sequence: str = Field(max_length=4 * MAX_SEQUENCE_LENGTH)

    @field_validator("sequence")
    @classmethod
    def validate_sequence(cls, value: str) -> str:
        return _clean_sequence(value)


@app.get("/")
def index() -> FileResponse:
    return FileResponse(STATIC_DIR / "index.html")


@app.get("/api/health")
def health(window_minutes: int = DEFAULT_HEALTH_WINDOW_MINUTES) -> JSONResponse:
    window_minutes = max(1, min(int(window_minutes), MAX_HEALTH_WINDOW_MINUTES))
    recent = _recent_usage_summary(window_minutes)
    degraded = recent["errors"] > 0

    body: Dict[str, Any] = {
        "status": "degraded" if degraded else "ok",
        "service": "Helix Design Studio",
        "version": APP_VERSION,
        "usage_logging": USAGE_LOGGING_ENABLED,
        "recent": recent,
        "alphabet_default": DEFAULT_ALPHABET_16,
        "environment_presets": ENVIRONMENT_PRESETS,
        "default_weights": DEFAULT_WEIGHTS,
    }
    # 503 para que un monitor externo basado en codigos HTTP vea el fallo: los
    # endpoints de streaming responden 200 aunque la ejecucion falle.
    return JSONResponse(content=body, status_code=503 if degraded else 200)


@app.post("/api/visit")
def register_visit(request: Request) -> Dict[str, Any]:
    _log_usage_event(request, "visit", {}, "ok", 0.0)
    return _usage_metrics()


@app.get("/api/usage-metrics")
def usage_metrics() -> Dict[str, Any]:
    return _usage_metrics()


def _provenance(payload: Any) -> Dict[str, Any]:
    """Todo lo necesario para repetir el cálculo: petición completa, versión, parámetros y semillas."""
    import numpy

    return {
        "software": "Helix Design Studio",
        "version": APP_VERSION,
        "parameters_sha256": PARAMETERS_SHA256,
        "request": payload.model_dump(mode="json"),
        "seed_protocol": SEED_PROTOCOL,
        "python": platform.python_version(),
        "numpy": numpy.__version__,
    }


def _consume_result(task: "asyncio.Future[Any]") -> None:
    # Evita avisos por excepciones de cálculos abandonados (p. ej. cancelados).
    if not task.cancelled():
        task.exception()


async def _await_calculation(request: Request, func: Callable[[], Any], cancelled: threading.Event) -> Any:
    """Ejecuta func en el pool de hilos; si el cliente se desconecta, pide al cálculo que se detenga."""
    task = asyncio.ensure_future(run_in_threadpool(func))
    task.add_done_callback(_consume_result)
    try:
        while not task.done():
            await asyncio.wait({task}, timeout=DISCONNECT_POLL_SECONDS)
            if not task.done() and await request.is_disconnected():
                cancelled.set()
        return task.result()
    finally:
        if not task.done():
            cancelled.set()


async def _run_calculation(
    request: Request,
    endpoint: str,
    payload: Any,
    calculation: Callable[[], Dict[str, Any]],
    summary: Optional[Callable[[Dict[str, Any]], Dict[str, Any]]] = None,
    use_job_slot: bool = True,
) -> Dict[str, Any]:
    started = time.perf_counter()
    if use_job_slot:
        _acquire_job_slot()
    cancelled = threading.Event()
    limits = CalculationLimits(deadline=time.monotonic() + MAX_JOB_SECONDS, cancelled=cancelled)

    def run() -> Dict[str, Any]:
        with calculation_limits(limits):
            return calculation()

    try:
        # Si el cliente se va (pestaña cerrada, Cancelar), el cálculo se detiene y libera su plaza.
        result = await _await_calculation(request, run, cancelled)
    except CalculationCancelled as exc:
        _log_usage_event(request, endpoint, payload, "cancelled", time.perf_counter() - started)
        raise HTTPException(status_code=400, detail=str(exc)) from exc
    except Exception as exc:
        status_code, detail = _classify_error(exc)
        _log_usage_event(
            request, endpoint, payload, "error", time.perf_counter() - started,
            {"error": detail, "http_status": status_code},
        )
        raise HTTPException(status_code=status_code, detail=detail) from exc
    finally:
        cancelled.set()
        if use_job_slot:
            _job_slots.release()
    _log_usage_event(
        request, endpoint, payload, "ok", time.perf_counter() - started, summary(result) if summary else None
    )
    return {**result, "provenance": _provenance(payload)}


def _stream_calculation(
    request: Request,
    endpoint: str,
    payload: Any,
    updates: Callable[[], Iterator[Dict[str, Any]]],
    summary: Callable[[Dict[str, Any]], Dict[str, Any]],
) -> StreamingResponse:
    async def generate() -> AsyncIterator[str]:
        started = time.perf_counter()
        if not _job_slots.acquire(blocking=False):
            yield json.dumps({"type": "error", "detail": BUSY_MSG}) + "\n"
            return
        cancelled = threading.Event()
        limits = CalculationLimits(deadline=time.monotonic() + MAX_JOB_SECONDS, cancelled=cancelled)
        iterator = updates()

        def step() -> Optional[Dict[str, Any]]:
            # Cada paso se ejecuta en un hilo del pool con los límites de esta petición.
            with calculation_limits(limits):
                return next(iterator, None)

        final_payload: Dict[str, Any] = {}
        try:
            provenance = _provenance(payload)
            while True:
                # Cada paso puede tardar (una búsqueda completa): se vigila la desconexión mientras dura.
                update = await _await_calculation(request, step, cancelled)
                if update is None:
                    break
                update_type = update.get("type")
                if update_type in ("meta", "final"):
                    event_payload = update.get("payload", {})
                    if update_type == "final":
                        final_payload = event_payload
                    update = {**update, "payload": {**event_payload, "provenance": provenance}}
                elif update_type == "candidate":
                    event_payload = update.get("payload", {})
                    update = {**update, "payload": {
                        **event_payload,
                        "member": {**event_payload["member"], "provenance": provenance},
                    }}
                yield json.dumps(update) + "\n"
        except CalculationCancelled:
            _log_usage_event(request, endpoint, payload, "cancelled", time.perf_counter() - started)
            return
        except Exception as exc:
            status_code, detail = _classify_error(exc)
            _log_usage_event(
                request, endpoint, payload, "error", time.perf_counter() - started,
                {"error": detail, "http_status": status_code},
            )
            yield json.dumps({"type": "error", "detail": detail}) + "\n"
            return
        finally:
            # También al cerrarse el stream por desconexión: el hilo se detiene en la siguiente evaluación.
            cancelled.set()
            _job_slots.release()
        _log_usage_event(request, endpoint, payload, "ok", time.perf_counter() - started, summary(final_payload))

    return StreamingResponse(generate(), media_type="application/x-ndjson")


def _family_summary(result: Dict[str, Any]) -> Dict[str, Any]:
    return {"family_size_returned": result.get("family_size_returned")}


def _specificity_summary(result: Dict[str, Any]) -> Dict[str, Any]:
    return {"final_sequence_hash": _sequence_summary(result.get("sequence")).get("sequence_hash")}


@app.post("/api/score")
async def score(payload: ScoreRequest, request: Request) -> Dict[str, Any]:
    return await _run_calculation(request, "score", payload, lambda: score_sequence(
        sequence=payload.sequence,
        residues=payload.residues,
        environment=payload.environment.model_dump(),
        weights=payload.weights.model_dump(),
        n_decoys=payload.n_decoys,
        n_random=payload.n_random,
        seed=payload.seed,
    ))


@app.post("/api/design")
async def design(payload: DesignRequest, request: Request) -> Dict[str, Any]:
    return await _run_calculation(request, "design", payload, lambda: design_sequence(
        length=payload.length,
        residues=payload.residues,
        environment=payload.environment.model_dump(),
        weights=payload.weights.model_dump(),
        n_decoys=payload.n_decoys,
        steps=payload.steps,
        restarts=payload.restarts,
        seed=payload.seed,
    ))


@app.post("/api/compare")
async def compare(payload: CompareRequest, request: Request) -> Dict[str, Any]:
    return await _run_calculation(request, "compare", payload, lambda: compare_environments(
        sequence=payload.sequence,
        residues=payload.residues,
        environment_a=payload.environment_a.model_dump(),
        environment_b=payload.environment_b.model_dump(),
        weights=payload.weights.model_dump(),
        n_decoys=payload.n_decoys,
        seed=payload.seed,
    ))


def _family_arguments(payload: FamilyDesignRequest) -> Dict[str, Any]:
    return {
        "length": payload.length,
        "residues": payload.residues,
        "environment": payload.environment.model_dump(),
        "weights": payload.weights.model_dump(),
        "n_decoys": payload.n_decoys,
        "steps": payload.steps,
        "restarts": payload.restarts,
        "family_size": payload.family_size,
        "oversample_factor": payload.oversample_factor,
        "seed": payload.seed,
    }


@app.post("/api/design-family")
async def family_design(payload: FamilyDesignRequest, request: Request) -> Dict[str, Any]:
    return await _run_calculation(
        request, "design-family", payload, lambda: design_family(**_family_arguments(payload)), _family_summary
    )


@app.post("/api/design-family-stream")
def family_design_stream_route(payload: FamilyDesignRequest, request: Request) -> StreamingResponse:
    return _stream_calculation(
        request, "design-family-stream", payload,
        lambda: design_family_stream(**_family_arguments(payload)), _family_summary,
    )


@app.post("/api/cross-design")
async def cross_design(payload: CrossDesignRequest, request: Request) -> Dict[str, Any]:
    return await _run_calculation(request, "cross-design", payload, lambda: cross_environment_design(
        length=payload.length,
        residues=payload.residues,
        environment_a=payload.environment_a.model_dump(),
        environment_b=payload.environment_b.model_dump(),
        lambda_gap=payload.lambda_gap,
        weights=payload.weights.model_dump(),
        n_decoys=payload.n_decoys,
        steps=payload.steps,
        restarts=payload.restarts,
        seed=payload.seed,
    ))


def _specificity_arguments(payload: SpecificityDesignRequest) -> Dict[str, Any]:
    return {
        "length": payload.length,
        "residues": payload.residues,
        "target_environment": payload.target_environment.model_dump(),
        "off_target_environments": [env.model_dump() for env in payload.off_target_environments],
        "num_sequences": payload.num_sequences,
        "lambda_balance": payload.lambda_balance,
        "weights": payload.weights.model_dump(),
        "n_decoys": payload.n_decoys,
        "steps": payload.steps,
        "restarts": payload.restarts,
        "seed": payload.seed,
    }


@app.post("/api/specificity-design")
async def specificity_design(payload: SpecificityDesignRequest, request: Request) -> Dict[str, Any]:
    return await _run_calculation(
        request, "specificity-design", payload,
        lambda: design_specificity(**_specificity_arguments(payload)), _specificity_summary,
    )


@app.post("/api/specificity-design-stream")
def specificity_design_stream_route(payload: SpecificityDesignRequest, request: Request) -> StreamingResponse:
    return _stream_calculation(
        request, "specificity-design-stream", payload,
        lambda: design_specificity_stream(**_specificity_arguments(payload)), _specificity_summary,
    )


@app.post("/api/optimal-penetration")
async def optimal_penetration(payload: PenetrationOptimizationRequest, request: Request) -> Dict[str, Any]:
    return await _run_calculation(request, "optimal-penetration", payload, lambda: optimize_penetration(
        sequence=payload.sequence,
        residues=payload.residues,
        environment=payload.environment.model_dump(),
        weights=payload.weights.model_dump(),
        n_decoys=payload.n_decoys,
        n_random=payload.n_random,
        percent_min=payload.percent_min,
        percent_max=payload.percent_max,
        percent_step=payload.percent_step,
        seed=payload.seed,
    ))


@app.post("/api/pdb")
async def pdb(payload: PdbRequest, request: Request) -> Dict[str, Any]:
    return await _run_calculation(
        request, "pdb", payload,
        lambda: {"sequence": payload.sequence, "pdb": generate_helix_pdb(payload.sequence)},
        use_job_slot=False,
    )
```
