## Additional file 1. Evaluated software source and dependencies for "Helix Design Studio: designing antimicrobial peptide candidates for membrane environments": 3Dmol-min.js

```
/*! For license information please see 3Dmol-min.js.LICENSE.txt */
!function(t,e){"object"==typeof exports&&"object"==typeof module?module.exports=e():"function"==typeof define&&define.amd?define([],e):"object"==typeof exports?exports["3Dmol"]=e():t["3Dmol"]=e()}(this,(()=>(()=>{var __webpack_modules__={620:function(){"use strict";!function(t){if(t.TextEncoder&&t.TextDecoder)return!1;function e(t="utf-8"){if("utf-8"!==t)throw new RangeError(`Failed to construct 'TextEncoder': The encoding label provided ('${t}') is invalid.`)}function i(t="utf-8",e={fatal:!1}){if("utf-8"!==t)throw new RangeError(`Failed to construct 'TextDecoder': The encoding label provided ('${t}') is invalid.`);if(e.fatal)throw new Error("Failed to construct 'TextDecoder': the 'fatal' option is unsupported.")}Object.defineProperty(e.prototype,"encoding",{value:"utf-8"}),e.prototype.encode=function(t,e={stream:!1}){if(e.stream)throw new Error("Failed to encode: the 'stream' option is unsupported.");let i=0;const r=t.length;let s=0,n=Math.max(32,r+(r>>1)+7),a=new Uint8Array(n>>3<<3);for(;i<r;){let e=t.charCodeAt(i++);if(e>=55296&&e<=56319){if(i<r){const r=t.charCodeAt(i);56320==(64512&r)&&(++i,e=((1023&e)<<10)+(1023&r)+65536)}if(e>=55296&&e<=56319)continue}if(s+4>a.length){n+=8,n*=1+i/t.length*2,n=n>>3<<3;const e=new Uint8Array(n);e.set(a),a=e}if(4294967168&e){if(4294965248&e)if(4294901760&e){if(4292870144&e)continue;a[s++]=e>>18&7|240,a[s++]=e>>12&63|128,a[s++]=e>>6&63|128}else a[s++]=e>>12&15|224,a[s++]=e>>6&63|128;else a[s++]=e>>6&31|192;a[s++]=63&e|128}else a[s++]=e}return a.slice(0,s)},Object.defineProperty(i.prototype,"encoding",{value:"utf-8"}),Object.defineProperty(i.prototype,"fatal",{value:!1}),Object.defineProperty(i.prototype,"ignoreBOM",{value:!1}),i.prototype.decode=function(t,e={stream:!1}){if(e.stream)throw new Error("Failed to decode: the 'stream' option is unsupported.");const i=new Uint8Array(t);let r=0;const s=i.length,n=[];for(;r<s;){const t=i[r++];if(0===t)break;if(128&t){if(192==(224&t)){const e=63&i[r++];n.push((31&t)<<6|e)}else if(224==(240&t)){const e=63&i[r++],s=63&i[r++];n.push((31&t)<<12|e<<6|s)}else if(240==(248&t)){let e=(7&t)<<18|(63&i[r++])<<12|(63&i[r++])<<6|63&i[r++];e>65535&&(e-=65536,n.push(e>>>10&1023|55296),e=56320|1023&e),n.push(e)}}else n.push(t)}return String.fromCharCode.apply(null,n)},t.TextEncoder=e,t.TextDecoder=i}("undefined"!=typeof window?window:"undefined"!=typeof self?self:this)},546:(t,e,i)=>{"use strict";i.r(e),i.d(e,{CustomLinear:()=>CustomLinear,Gradient:()=>Gradient,GradientType:()=>GradientType,ROYGB:()=>ROYGB,RWB:()=>RWB,Sinebow:()=>Sinebow,builtinGradients:()=>a,getGradient:()=>n,normalizeValue:()=>s});var r=i(222);class GradientType{}function s(t,e,i){return e>=t?(i<t&&(i=t),i>e&&(i=e),{lo:t,hi:e,val:i}):(i>t&&(i=t),i<e&&(i=e),{lo:e,hi:t,val:i=t-i+e})}function n(t){if(t instanceof GradientType)return t;if(void 0!==t.gradient&&a[t.gradient]){let e=void 0===t.min?-1:t.min,i=void 0===t.max?1:t.max;return void 0===t.mid?void 0===t.colors?new a[t.gradient](e,i):new a[t.gradient](e,i,t.colors):new a[t.gradient](e,i,t.mid)}if("string"==typeof t.gradient&&t.gradient.startsWith("linear_")){let e=t.gradient.split("_");e.shift();let i=void 0===t.min?-1:t.min,r=void 0===t.max?1:t.max;return new CustomLinear(i,r,e)}return t}class RWB extends GradientType{constructor(t,e,i){super(),this.gradient="RWB",this.mult=1,this.mid=i,this.min=t,this.max=e,void 0===e&&Array.isArray(t)&&t.length>=2?(this.max=t[1],this.min=t[0]):t&&e&&!Array.isArray(t)&&(this.min=t,this.max=e)}range(){return void 0!==this.min&&void 0!==this.max?[this.min,this.max]:null}valueToHex(t,e){var i,r;if(t=this.mult*t,e?(i=e[0],r=e[1]):(i=this.min,r=this.max),void 0===t)return 16777215;var n=s(i,r,t);i=n.lo;var a,o=((r=n.hi)+i)/2;return(t=n.val)<(o=e&&void 0!==e[2]?e[2]:void 0!==this.mid?this.mid:(i+r)/2)?16711680+256*(a=Math.floor(255*Math.sqrt((t-i)/(o-i))))+a:t>o?65536*(a=Math.floor(255*Math.sqrt(1-(t-o)/(r-o))))+256*a+255:16777215}}class ROYGB extends GradientType{constructor(t,e){super(),this.gradient="ROYGB",this.mult=1,this.min=t,this.max=e,void 0===e&&Array.isArray(t)&&t.length>=2?(this.max=t[1],this.min=t[0]):t&&e&&!Array.isArray(t)&&(this.min=t,this.max=e)}valueToHex(t,e){var i,r;if(t=this.mult*t,e?(i=e[0],r=e[1]):(i=this.min,r=this.max),void 0===t)return 16777215;var n=s(i,r,t),a=((i=n.lo)+(r=n.hi))/2,o=(i+a)/2,l=(a+r)/2;return(t=n.val)<o?16711680+256*Math.floor(255*Math.sqrt((t-i)/(o-i)))+0:t<a?65536*Math.floor(255*Math.sqrt(1-(t-o)/(a-o)))+65280+0:t<l?65280+1*Math.floor(255*Math.sqrt((t-a)/(l-a))):0+256*Math.floor(255*Math.sqrt(1-(t-l)/(r-l)))+255}range(){return void 0!==this.min&&void 0!==this.max?[this.min,this.max]:null}}class Sinebow extends GradientType{constructor(t,e){super(),this.gradient="Sinebow",this.mult=1,this.min=t,this.max=e,void 0===e&&Array.isArray(t)&&t.length>=2&&(this.max=t[1],this.min=t[0]),e<t&&(this.mult=-1,this.min*=-1,this.max*=-1)}valueToHex(t,e){var i,r;if(t=this.mult*t,e?(i=e[0],r=e[1]):(i=this.min,r=this.max),void 0===t)return 16777215;var s=Gradient.normalizeValue(i,r,t);i=s.lo,r=s.hi;var n=5*(((t=s.val)-i)/(r-i))/6+.5,a=Math.sin(Math.PI*n);a*=255*a;var o=Math.sin(Math.PI*(n+1/3));o*=255*o;var l=Math.sin(Math.PI*(n+2/3));return l*=255*l,65536*Math.floor(a)+256*Math.floor(l)+1*Math.floor(o)}range(){return void 0!==this.min&&void 0!==this.max?[this.min,this.max]:null}}class CustomLinear extends GradientType{constructor(t,e,i){var s;if(super(),this.gradient="linear",this.colors=new Array,Array.isArray(t)&&t.length>=2?(this.max=t[1],this.min=t[0],s=e):(this.min=t,this.max=e,s=i),s)for(let t of s)this.colors.push(r.CC.color(t));else this.colors.push(r.CC.color(0))}range(){return void 0!==this.min&&void 0!==this.max?[this.min,this.max]:null}valueToHex(t,e){var i,n;if(e?(i=e[0],n=e[1]):(i=this.min,n=this.max),void 0===t)return 16777215;var a=s(i,n,t);i=a.lo,n=a.hi,t=a.val;let o=this.colors.length,l=(n-i)/o,h=Math.min(Math.floor((t-i)/l),o-1),c=Math.min(h+1,o-1),d=(t-i-h*l)/l,u=this.colors[h],f=this.colors[c];return new r.Color(u.r+d*(f.r-u.r),u.g+d*(f.g-u.g),u.b+d*(f.b-u.b)).getHex()}}const a={rwb:RWB,RWB,roygb:ROYGB,ROYGB,sinebow:Sinebow,linear:CustomLinear};class Gradient extends GradientType{valueToHex(t,e){return 0}range(){return null}}Gradient.RWB=RWB,Gradient.ROYGB=ROYGB,Gradient.Sinebow=Sinebow,Gradient.CustomLinear=CustomLinear,Gradient.builtinGradients=a,Gradient.normalizeValue=s,Gradient.getGradient=n},848:(t,e,i)=>{"use strict";i.r(e),i.d(e,{VolumeData:()=>VolumeData});var r=i(864),s=i(529),n=i(797),a=i(865),o=i(75);class VolumeData{constructor(t,e,i){if(this.unit={x:1,y:1,z:1},this.origin={x:0,y:0,z:0},this.size={x:0,y:0,z:0},this.data=new Float32Array([]),this.matrix=null,this.inversematrix=null,this.isbinary=new Set(["ccp4","CCP4"]),this.getCoordinates=function(t){var e=t/(this.size.y*this.size.z),i=t%(this.size.y*this.size.z),r=t%this.size.z;return e*=this.unit.x,i*=this.unit.y,r*=this.unit.z,{x:e+=this.origin.x,y:i+=this.origin.y,z:r+=this.origin.z}},this.vasp=function(t){var e=t.replace(/^\s+/,"").split(/[\n\r]/),i=(0,n.VASP)(t)[0].length;if(0==i)return console.warn("No good formating of CHG or CHGCAR file, not atomic information provided in the file."),void(this.data=[]);var r,a=1.889725992,o=parseFloat(e[1]);r=e[2].replace(/^\s+/,"").split(/\s+/);var l=new s.Vector3(parseFloat(r[0]),parseFloat(r[1]),parseFloat(r[2])).multiplyScalar(o*a);r=e[3].replace(/^\s+/,"").split(/\s+/);var h=new s.Vector3(parseFloat(r[0]),parseFloat(r[1]),parseFloat(r[2])).multiplyScalar(o*a);r=e[4].replace(/^\s+/,"").split(/\s+/);var c=new s.Vector3(parseFloat(r[0]),parseFloat(r[1]),parseFloat(r[2])).multiplyScalar(o*a),d=l.x*(h.y*c.z-c.y*h.z)-h.x*(l.y*c.z-c.y*l.z)+c.x*(l.y*h.z-h.y*l.z),u=1/(d=Math.abs(d)/Math.pow(a,3));e.splice(0,8+i+1);var f=e[0].replace(/^\s+/,"").replace(/\s+/g," ").split(" "),p=Math.abs(parseFloat(f[0])),g=Math.abs(parseFloat(f[1])),m=Math.abs(parseFloat(f[2])),v=this.origin=new s.Vector3(0,0,0);this.size={x:p,y:g,z:m},this.unit=new s.Vector3(l.x,h.y,c.z),l=l.multiplyScalar(1/(a*p)),h=h.multiplyScalar(1/(a*g)),c=c.multiplyScalar(1/(a*m)),0==l.y&&0==l.z&&0==h.x&&0==h.z&&0==c.x&&0==c.y||(this.matrix=new s.Matrix4(l.x,h.x,c.x,0,l.y,h.y,c.y,0,l.z,h.z,c.z,0,0,0,0,1),this.matrix=this.matrix.multiplyMatrices(this.matrix,(new s.Matrix4).makeTranslation(v.x,v.y,v.z)),this.origin=new s.Vector3(0,0,0),this.unit=new s.Vector3(1,1,1)),e.splice(0,1);var _=e.join(" "),y=(_=_.replace(/^\s+/,"")).split(/[\s\r]+/);y.splice(p*g*m+1);for(var b=Float32Array.from(y,parseFloat),x=0;x<b.length;x++)b[x]=b[x]*u*.036749309;this.data=b},this.dx=function(t){var e,i=t.split(/[\n\r]+/),r=/gridpositions\s+counts\s+(\d+)\s+(\d+)\s+(\d+)/,n=/^origin\s+(\S+)\s+(\S+)\s+(\S+)/,a=/^delta\s+(\S+)\s+(\S+)\s+(\S+)/,o=/data follows/,l=0;for(l=0;l<i.length;l++){var h=i[l];if(e=r.exec(h)){var c=parseInt(e[1]),d=parseInt(e[2]),u=parseInt(e[3]);this.size={x:c,y:d,z:u}}else if(e=a.exec(h)){var f=parseFloat(e[1]);if(0==parseFloat(e[2])&&0==parseFloat(e[3])||console.warn("Non-orthogonal delta matrix not currently supported in dx format"),h=i[l+=1],null==(e=a.exec(h)))return void console.error("Parse error in dx delta matrix");var p=parseFloat(e[2]);if(0==parseFloat(e[1])&&0==parseFloat(e[3])||console.warn("Non-orthogonal delta matrix not currently supported in dx format"),h=i[l+=1],null==(e=a.exec(h)))return void console.error("Parse error in dx delta matrix");var g=parseFloat(e[3]);0==parseFloat(e[1])&&0==parseFloat(e[2])||console.warn("Non-orthogonal delta matrix not currently supported in dx format"),this.unit=new s.Vector3(f,p,g)}else if(e=n.exec(h)){var m=parseFloat(e[1]),v=parseFloat(e[2]),_=parseFloat(e[3]);this.origin=new s.Vector3(m,v,_)}else if(e=o.exec(h))break}if(l+=1,this.size&&this.origin&&this.unit&&this.size){var y=i.splice(l).join(" ").split(/[\s\r]+/);this.data=Float32Array.from(y,parseFloat)}else console.error("Error parsing dx format")},e=e.toLowerCase(),/\.gz$/.test(e)){e=e.replace(/\.gz$/,"");try{this[e]&&this.isbinary.has(e)?("string"==typeof t&&(t=(0,r.base64ToArray)(t)),t=(0,o.inflate)(t)):t=new TextDecoder("utf-8").decode((0,o.inflate)(t))}catch(t){console.error(t)}}if(this[e]&&(this.isbinary.has(e)&&"string"==typeof t&&(t=(0,r.base64ToArray)(t)),this[e](t)),i){if(i.negate)for(let t=0,e=this.data.length;t<e;t++)this.data[t]=-this.data[t];if(i.normalize){var a=0;for(let t=0,e=this.data.length;t<e;t++)a+=this.data[t];var l=a/this.data.length;a=0;for(let t=0,e=this.data.length;t<e;t++){var h=this.data[t]-l;a+=h*h}var c=a/this.data.length;for(let t=0,e=this.data.length;t<e;t++)this.data[t]=(this.data[t]-l)/c}}}getIndex(t,e,i){if(this.matrix){null==this.inversematrix&&(this.inversematrix=(new s.Matrix4).getInverse(this.matrix));var r=new s.Vector3(t,e,i);t=(r=r.applyMatrix4(this.inversematrix)).x,e=r.y,i=r.z}else t-=this.origin.x,e-=this.origin.y,i-=this.origin.z,t/=this.unit.x,e/=this.unit.y,i/=this.unit.z;return t=Math.round(t),e=Math.round(e),i=Math.round(i),t<0||t>=this.size.x||e<0||e>=this.size.y||i<0||i>=this.size.z?-1:t*this.size.y*this.size.z+e*this.size.z+i}getVal(t,e,i){let r=this.getIndex(t,e,i);return r<0?0:this.data[r]}cube(t){var e=t.split(/\r?\n/);if(!(e.length<6)){var i=(0,a.CUBE)(t,{}).modelData[0].cryst,r=e[2].replace(/^\s+/,"").replace(/\s+/g," ").split(" "),s=parseFloat(r[0]),n=Math.abs(s);this.origin=i.origin,this.size=i.size,this.unit=i.unit,this.matrix=i.matrix4;var o=6;s<0&&o++;var l=e.splice(n+o).join(" "),h=(l=l.replace(/^\s+/,"")).split(/[\s\r]+/);this.data=Float32Array.from(h,parseFloat)}}ccp4(t){var e={};t=new Int8Array(t);var i=new Int32Array(t.buffer,0,56),r=new Float32Array(t.buffer,0,56),n=new DataView(t.buffer);if(e.MAP=String.fromCharCode(n.getUint8(208),n.getUint8(209),n.getUint8(210),n.getUint8(211)),e.MACHST=[n.getUint8(212),n.getUint8(213)],17===e.MACHST[0]&&17===e.MACHST[1])for(var a=t.byteLength,o=0;o<a;o+=4)n.setFloat32(o,n.getFloat32(o),!0);e.NX=i[0],e.NY=i[1],e.NZ=i[2],e.MODE=i[3],e.NXSTART=i[4],e.NYSTART=i[5],e.NZSTART=i[6],e.MX=i[7],e.MY=i[8],e.MZ=i[9],e.xlen=r[10],e.ylen=r[11],e.zlen=r[12],e.alpha=r[13],e.beta=r[14],e.gamma=r[15],e.MAPC=i[16],e.MAPR=i[17],e.MAPS=i[18],e.DMIN=r[19],e.DMAX=r[20],e.DMEAN=r[21],e.ISPG=i[22],e.NSYMBT=i[23],e.LSKFLG=i[24],e.originX=r[49],e.originY=r[50],e.originZ=r[51],e.ARMS=r[54];var l=e,h=[l.xlen,0,0],c=[l.ylen*Math.cos(Math.PI/180*l.gamma),l.ylen*Math.sin(Math.PI/180*l.gamma),0],d=[l.zlen*Math.cos(Math.PI/180*l.beta),l.zlen*(Math.cos(Math.PI/180*l.alpha)-Math.cos(Math.PI/180*l.gamma)*Math.cos(Math.PI/180*l.beta))/Math.sin(Math.PI/180*l.gamma),0];d[2]=Math.sqrt(l.zlen*l.zlen*Math.sin(Math.PI/180*l.beta)*Math.sin(Math.PI/180*l.beta)-d[1]*d[1]);var u=[0,h,c,d],f=[0,l.MX,l.MY,l.MZ],p=[0,l.MAPC,l.MAPR,l.MAPS];this.matrix=new s.Matrix4,this.matrix.set(u[p[1]][0]/f[p[1]],u[p[2]][0]/f[p[2]],u[p[3]][0]/f[p[3]],0,u[p[1]][1]/f[p[1]],u[p[2]][1]/f[p[2]],u[p[3]][1]/f[p[3]],0,u[p[1]][2]/f[p[1]],u[p[2]][2]/f[p[2]],u[p[3]][2]/f[p[3]],0,0,0,0,1),this.matrix=this.matrix.multiplyMatrices(this.matrix,(new s.Matrix4).makeTranslation(l.NXSTART+l.originX,l.NYSTART+l.originY,l.NZSTART+l.originZ)),this.origin=new s.Vector3(0,0,0),this.unit=new s.Vector3(1,1,1),this.size={x:e.NX,y:e.NY,z:e.NZ},this.dimensionorder=[e.MAPC,e.MAPR,e.MAPS];var g=new Float32Array(t.buffer,1024+e.NSYMBT),m=e.NX,v=e.NY,_=e.NZ;this.data=new Float32Array(m*v*_);for(let t=0;t<m;t++)for(let e=0;e<v;e++)for(let i=0;i<_;i++)this.data[(t*v+e)*_+i]=g[(i*v+e)*m+t]}}},638:(t,e,i)=>{"use strict";var r;i.r(e),i.d(e,{BackSide:()=>n,Camera:()=>Camera,ClampToEdgeWrapping:()=>c,Coloring:()=>r,Cylinder:()=>M.Cylinder,DoubleSide:()=>a,EventDispatcher:()=>EventDispatcher,FloatType:()=>g,Fog:()=>Fog,FrontSide:()=>s,Geometry:()=>Geometry,GeometryGroup:()=>GeometryGroup,GeometryIDCount:()=>A,ImposterMaterial:()=>ImposterMaterial,InstancedMaterial:()=>InstancedMaterial,Light:()=>Light,Line:()=>Line,LineBasicMaterial:()=>LineBasicMaterial,LineStyle:()=>N,LinearFilter:()=>d,LinearMipMapLinearFilter:()=>f,Material:()=>Material,MaterialIdCount:()=>x,Matrix3:()=>l.Matrix3,Matrix4:()=>l.Matrix4,Mesh:()=>Mesh,MeshDoubleLambertMaterial:()=>MeshDoubleLambertMaterial,MeshLambertMaterial:()=>MeshLambertMaterial,MeshOutlineMaterial:()=>MeshOutlineMaterial,NearestFilter:()=>u,Object3D:()=>Object3D,Object3DIDCount:()=>C,Projector:()=>Projector,Quaternion:()=>l.Quaternion,R32Format:()=>_,RFormat:()=>v,RGBAFormat:()=>m,Ray:()=>l.Ray,Raycaster:()=>Raycaster,Renderer:()=>Renderer,Scene:()=>Scene,ShaderLib:()=>ct,ShaderUtils:()=>ut,Shading:()=>o,Sphere:()=>M.Sphere,SphereImposterMaterial:()=>SphereImposterMaterial,SphereImposterOutlineMaterial:()=>SphereImposterOutlineMaterial,Sprite:()=>Sprite,SpriteAlignment:()=>h,SpriteMaterial:()=>SpriteMaterial,SpritePlugin:()=>SpritePlugin,StickImposterMaterial:()=>StickImposterMaterial,StickImposterOutlineMaterial:()=>StickImposterOutlineMaterial,Texture:()=>Texture,TextureIdCount:()=>B,TextureOperations:()=>y,Triangle:()=>M.Triangle,UVMapping:()=>UVMapping,UnsignedByteType:()=>p,Vector2:()=>l.Vector2,Vector3:()=>l.Vector3,VolumetricMaterial:()=>VolumetricMaterial,basic:()=>V,clamp:()=>l.clamp,clone:()=>dt,conversionMatrix3:()=>l.conversionMatrix3,degToRad:()=>l.degToRad,instanced:()=>H,intersectObject:()=>U,lambert:()=>q,lambertdouble:()=>Z,outline:()=>K,screen:()=>Q,screenaa:()=>$,sphereimposter:()=>tt,sphereimposteroutline:()=>it,sprite:()=>rt,stickimposter:()=>at,stickimposteroutline:()=>ot,volumetric:()=>ht}),function(t){t[t.NoColors=0]="NoColors",t[t.FaceColors=1]="FaceColors",t[t.VertexColors=2]="VertexColors"}(r||(r={}));const s=0,n=1,a=2;var o;!function(t){t[t.NoShading=0]="NoShading",t[t.FlatShading=1]="FlatShading",t[t.SmoothShading=2]="SmoothShading"}(o||(o={}));var l=i(529);const h={topLeft:new l.Vector2(1,-1),topCenter:new l.Vector2(0,-1),topRight:new l.Vector2(-1,-1),centerLeft:new l.Vector2(1,0),center:new l.Vector2(0,0),centerRight:new l.Vector2(-1,0),bottomLeft:new l.Vector2(1,1),bottomCenter:new l.Vector2(0,1),bottomRight:new l.Vector2(-1,1)},c=1001,d=1006,u=1007,f=1008,p=1009,g=1010,m=1021,v=1022,_=1023;var y;!function(t){t[t.MultiplyOperation=0]="MultiplyOperation",t[t.MixOperation=1]="MixOperation",t[t.AddOperation=2]="AddOperation"}(y||(y={}));class EventDispatcher{constructor(){this.listeners={}}dispatchEvent(t){var e=this.listeners[t.type];if(void 0!==e){t.target=this;for(var i=0,r=e.length;i<r;i++)e[i].call(this,t)}}removeEventListener(t,e){if(e){var i=this.listeners[t].indexOf(e);-1!==i&&this.listeners[t].splice(i,1)}else this.listeners[t]=[]}addEventListener(t,e){void 0===this.listeners[t]&&(this.listeners[t]=[]),-1===this.listeners[t].indexOf(e)&&this.listeners[t].push(e)}}var b=i(222);class Material extends EventDispatcher{constructor(){super(...arguments),this.id=x++,this.name="",this.side=s,this.opacity=1,this.transparent=!1,this.depthTest=!0,this.depthWrite=!0,this.polygonOffset=!1,this.polygonOffsetFactor=0,this.polygonOffsetUnits=0,this.alphaTest=0,this.visible=!0,this.needsUpdate=!0,this.outline=!1,this.wireframe=!1,this.shaded=!1}setValues(t={}){if(void 0!==t)for(var e in t){var i=t[e];if(void 0!==i){if(e in this){var r=this[e];r instanceof b.Color&&i instanceof b.Color?r.copy(i):r instanceof b.Color?r.set(i):r instanceof l.Vector3&&i instanceof l.Vector3?r.copy(i):this[e]=i}}else console.warn("$3Dmol.Material: '"+e+"' parameter is undefined.")}}clone(t=new Material){return t.name=this.name,t.side=this.side,t.opacity=this.opacity,t.transparent=this.transparent,t.depthTest=this.depthTest,t.depthWrite=this.depthWrite,t.polygonOffset=this.polygonOffset,t.polygonOffsetFactor=this.polygonOffsetFactor,t.polygonOffsetUnits=this.polygonOffsetUnits,t.alphaTest=this.alphaTest,t.overdraw=this.overdraw,t.visible=this.visible,t}makeShaded(t){this.shaded=!0,this.uniforms&&(this.uniforms.shading={type:"i",value:t},this.uniforms.vWidth={type:"f",value:1},this.uniforms.vHeight={type:"f",value:1})}dispose(){this.dispatchEvent({type:"dispose"})}}let x=0;class LineBasicMaterial extends Material{constructor(t){super(),this.color=new b.Color(16777215),this.linewidth=1,this.linecap="round",this.linejoin="round",this.vertexColors=!1,this.fog=!0,this.shaderID="basic",this.setValues(t)}clone(t=new LineBasicMaterial){return super.clone.call(this,t),t.color.copy(this.color),t}}const w=65535;class GeometryGroup{constructor(t=0){this.vertexArray=null,this.colorArray=null,this.normalArray=null,this.radiusArray=null,this.faceArray=null,this.lineArray=null,this.atomArray=Array(),this.vertices=0,this.faceidx=0,this.lineidx=0,this.__inittedArrays=!1,this.id=t}setColor(t){var e=this.vertexArray,i=this.colorArray;if(!e)throw new Error("vertex array not initialized");if(!i)throw new Error("color array not initialized");let r=b.CC.color(t);for(var s=0;s<e.length;s+=3)i[s]=r.r,i[s+1]=r.g,i[s+2]=r.b}setColors(t){var e=this.vertexArray,i=this.colorArray;if(!e)throw new Error("vertex array not initialized");if(!i)throw new Error("color array not initialized");if(e.length==i.length)for(var r=0;r<e.length;r+=3){var s=t(e[r],e[r+1],e[r+2]);s instanceof b.Color||(s=b.CC.color(s)),i[r]=s.r,i[r+1]=s.g,i[r+2]=s.b}else console.log("Cannot re-color geometry group due to mismatched lengths.")}getNumVertices(){return this.vertices}getVertices(){return this.vertexArray}getCentroid(){if(!this.vertexArray)throw new Error("vertex array not initialized");for(var t,e,i,r,s=new l.Vector3,n=0;n<this.vertices;++n)t=3*n,e=this.vertexArray[t],i=this.vertexArray[t+1],r=this.vertexArray[t+2],s.x+=e,s.y+=i,s.z+=r;return s.divideScalar(this.vertices),s}setNormals(){var t=this.faceArray,e=this.vertexArray,i=this.normalArray;if(this.vertices&&this.faceidx){if(!t)throw new Error("face array not initialized");if(!e)throw new Error("vertex array not initialized");if(!i)throw new Error("normal array not initialized");for(var r,s,n,a,o,h,c,d=0;d<t.length/3;++d)r=3*t[3*d],s=3*t[3*d+1],n=3*t[3*d+2],a=new l.Vector3(e[r],e[r+1],e[r+2]),o=new l.Vector3(e[s],e[s+1],e[s+2]),h=new l.Vector3(e[n],e[n+1],e[n+2]),a.subVectors(a,o),h.subVectors(h,o),h.cross(a),(c=h).normalize(),i[r]+=c.x,i[s]+=c.x,i[n]+=c.x,i[r+1]+=c.y,i[s+1]+=c.y,i[n+1]+=c.y,i[r+2]+=c.z,i[s+2]+=c.z,i[n+2]+=c.z}}setLineIndices(){if(this.faceidx&&(!this.lineArray||this.lineArray.length!=2*this.faceidx||this.lineidx!=2*this.faceidx)){var t=this.faceArray,e=this.lineArray=new Uint16Array(2*this.faceidx);if(this.lineidx=2*this.faceidx,!t)throw new Error("face array not initialized");for(var i=0;i<this.faceidx/3;++i){var r=3*i,s=2*r,n=t[r],a=t[r+1],o=t[r+2];e[s]=n,e[s+1]=a,e[s+2]=n,e[s+3]=o,e[s+4]=a,e[s+5]=o}}}vrml(t,e){var i,r,s,n,a,o,l,h,c,d,u,f,p,g,m,v,_,y,b="";if(b+=t+"Shape {\n"+t+" appearance Appearance {\n"+t+"  material Material {\n"+t+"   diffuseColor "+(null===(i=null==e?void 0:e.color)||void 0===i?void 0:i.r)+" "+(null===(r=null==e?void 0:e.color)||void 0===r?void 0:r.g)+" "+(null===(s=null==e?void 0:e.color)||void 0===s?void 0:s.b)+"\n",e.wireframe&&this.colorArray){let e=this.colorArray;b+=t+"    emissiveColor "+e[0]+" "+e[1]+" "+e[2]+"\n"}(null==e?void 0:e.transparent)&&(b+=t+"   transparency "+(1-e.opacity)+"\n"),b+=t+"  }\n",b+=t+" }\n";var x=t;if(t+=" ",e instanceof LineBasicMaterial||e.wireframe){let i,r,s;b+=t+"geometry IndexedLineSet {\n"+t+" colorPerVertex TRUE\n"+t+" coord Coordinate {\n"+t+"  point [\n";for(let e=0;e<this.vertices;++e){let l=3*e;i=null===(n=this.vertexArray)||void 0===n?void 0:n[l],r=null===(a=this.vertexArray)||void 0===a?void 0:a[l+1],s=null===(o=this.vertexArray)||void 0===o?void 0:o[l+2],b+=t+"   "+i+" "+r+" "+s+",\n"}if(b+=t+"  ]\n",b+=t+" }\n",this.colorArray&&!e.wireframe){b+=t+" color Color {\n"+t+"  color [\n";for(let e=0;e<this.vertices;++e){let n=3*e;i=this.colorArray[n],r=this.colorArray[n+1],s=this.colorArray[n+2],b+=t+"   "+i+" "+r+" "+s+",\n"}b+=t+"  ]\n",b+=t+" }\n"}if(b+=t+" coordIndex [\n",e.wireframe&&this.faceArray)for(let e=0;e<this.faceidx;e+=3)i=null===(l=this.faceArray)||void 0===l?void 0:l[e],r=null===(h=this.faceArray)||void 0===h?void 0:h[e+1],s=null===(c=this.faceArray)||void 0===c?void 0:c[e+2],b+=t+"  "+i+", "+r+", "+s+", -1,\n";else for(let e=0;e<this.vertices-1;e+=2)b+=t+"  "+e+", "+(e+1)+", -1,\n";b+=t+" ]\n",b+=t+"}\n"}else{let e,i,r;b+=t+"geometry IndexedFaceSet {\n"+t+" colorPerVertex TRUE\n"+t+" normalPerVertex TRUE\n"+t+" solid FALSE\n",b+=t+" coord Coordinate {\n"+t+"  point [\n";for(let s=0;s<this.vertices;++s){let n=3*s;e=null===(d=this.vertexArray)||void 0===d?void 0:d[n],i=null===(u=this.vertexArray)||void 0===u?void 0:u[n+1],r=null===(f=this.vertexArray)||void 0===f?void 0:f[n+2],b+=t+"   "+e+" "+i+" "+r+",\n"}b+=t+"  ]\n",b+=t+" }\n",b+=t+" normal Normal {\n"+t+"  vector [\n";for(let s=0;s<this.vertices;++s){let n=3*s;e=null===(p=this.normalArray)||void 0===p?void 0:p[n],i=null===(g=this.normalArray)||void 0===g?void 0:g[n+1],r=null===(m=this.normalArray)||void 0===m?void 0:m[n+2],b+=t+"   "+e+" "+i+" "+r+",\n"}if(b+=t+"  ]\n",b+=t+" }\n",this.colorArray){b+=t+" color Color {\n"+t+"  color [\n";for(let s=0;s<this.vertices;++s){let n=3*s;e=this.colorArray[n],i=this.colorArray[n+1],r=this.colorArray[n+2],b+=t+"   "+e+" "+i+" "+r+",\n"}b+=t+"  ]\n",b+=t+" }\n"}b+=t+" coordIndex [\n";for(let s=0;s<this.faceidx;s+=3)e=null===(v=this.faceArray)||void 0===v?void 0:v[s],i=null===(_=this.faceArray)||void 0===_?void 0:_[s+1],r=null===(y=this.faceArray)||void 0===y?void 0:y[s+2],b+=t+"  "+e+", "+i+", "+r+", -1,\n";b+=t+" ]\n",b+=t+"}\n"}return b+=x+"}"}truncateArrayBuffers(t=!0,e=!1){var i=this.vertexArray,r=this.colorArray,s=this.normalArray,n=this.faceArray,a=this.lineArray,o=this.radiusArray;this.vertexArray=(null==i?void 0:i.subarray(0,3*this.vertices))||null,this.colorArray=(null==r?void 0:r.subarray(0,3*this.vertices))||null,t?(this.normalArray=(null==s?void 0:s.subarray(0,3*this.vertices))||null,this.faceArray=(null==n?void 0:n.subarray(0,this.faceidx))||null,this.lineidx>0?this.lineArray=(null==a?void 0:a.subarray(0,this.lineidx))||null:this.lineArray=new Uint16Array(0)):(this.normalArray=new Float32Array(0),this.faceArray=new Uint16Array(0),this.lineArray=new Uint16Array(0)),o&&(this.radiusArray=o.subarray(0,this.vertices)),e&&(this.normalArray&&(this.normalArray=new Float32Array(this.normalArray)),this.faceArray&&(this.faceArray=new Uint16Array(this.faceArray)),this.lineArray&&(this.lineArray=new Uint16Array(this.lineArray)),this.vertexArray&&(this.vertexArray=new Float32Array(this.vertexArray)),this.colorArray&&(this.colorArray=new Float32Array(this.colorArray)),this.radiusArray&&(this.radiusArray=new Float32Array(this.radiusArray))),this.__inittedArrays=!0}}class Geometry extends EventDispatcher{constructor(t=!1,e=!1,i=!1){super(),this.name="",this.hasTangents=!1,this.dynamic=!0,this.verticesNeedUpdate=!1,this.elementsNeedUpdate=!1,this.normalsNeedUpdate=!1,this.colorsNeedUpdate=!1,this.buffersNeedUpdate=!1,this.imposter=!1,this.instanced=!1,this.geometryGroups=[],this.groups=0,this.id=A++,this.mesh=t,this.radii=e,this.offset=i}updateGeoGroup(t=0){var e,i=this.groups>0?this.geometryGroups[this.groups-1]:null;return(!i||i.vertices+t>((null===(e=null==i?void 0:i.vertexArray)||void 0===e?void 0:e.length)||0)/3)&&(i=this.addGeoGroup()),i}vrml(t,e){for(var i="",r=this.geometryGroups.length,s=0;s<r;s++){i+=this.geometryGroups[s].vrml(t,e)+",\n"}return i}addGeoGroup(){var t=new GeometryGroup(this.geometryGroups.length);return this.geometryGroups.push(t),this.groups=this.geometryGroups.length,t.vertexArray=new Float32Array(196605),t.colorArray=new Float32Array(196605),this.mesh&&(t.normalArray=new Float32Array(196605),t.faceArray=new Uint16Array(393210),t.lineArray=new Uint16Array(393210)),this.radii&&(t.radiusArray=new Float32Array(w)),t.useOffset=this.offset,t}setUpNormals(...t){for(var e=0;e<this.groups;e++){this.geometryGroups[e].setNormals(...t)}}setColors(...t){for(var e=this.geometryGroups.length,i=0;i<e;i++){this.geometryGroups[i].setColors(...t)}}setColor(...t){for(var e=this.geometryGroups.length,i=0;i<e;i++){this.geometryGroups[i].setColor(...t)}}setUpWireframe(...t){for(var e=0;e<this.groups;e++){this.geometryGroups[e].setLineIndices(...t)}}initTypedArrays(){for(var t=0;t<this.groups;t++){var e=this.geometryGroups[t];!0!==e.__inittedArrays&&e.truncateArrayBuffers(this.mesh,!1)}}dispose(){this.dispatchEvent({type:"dispose"})}get vertices(){for(var t=0,e=0;e<this.groups;e++)t+=this.geometryGroups[e].vertices;return t}}let A=0,C=0;class Object3D{constructor(){this.id=C++,this.name="",this.children=[],this.position=new l.Vector3,this.rotation=new l.Vector3,this.matrix=new l.Matrix4,this.matrixWorld=new l.Matrix4,this.quaternion=new l.Quaternion,this.eulerOrder="XYZ",this.up=new l.Vector3(0,1,0),this.scale=new l.Vector3(1,1,1),this.matrixAutoUpdate=!0,this.matrixWorldNeedsUpdate=!0,this.rotationAutoUpdate=!0,this.useQuaternion=!1,this.visible=!0}lookAt(t){this.matrix.lookAt(t,this.position,this.up),this.rotationAutoUpdate&&(!0===this.useQuaternion?console.error("Unimplemented math operation."):this.rotation instanceof l.Vector3&&this.rotation.setEulerFromRotationMatrix(this.matrix,this.eulerOrder))}add(t){if(t!==this){t.parent=this,this.children.push(t);for(var e=this;void 0!==e.parent;)e=e.parent;void 0!==e&&e instanceof Scene&&e.__addObject(t)}else console.error("Can't add $3Dmol.Object3D to itself")}remove(t){var e=this.children.indexOf(t);if(-1!==e){t.parent=void 0,this.children.splice(e,1);for(var i=this;void 0!==i.parent;)i=i.parent;void 0!==i&&i instanceof Scene&&i.__removeObject(t)}}vrml(t){t||(t=" ");var e=2*Math.atan2(this.quaternion.lengthxyz(),this.quaternion.w),i=0,r=0,s=0;if(0!=e){let t=Math.sin(e/2);i=this.quaternion.x/t,r=this.quaternion.y/t,s=this.quaternion.z/t}var n=t+"Transform {\n"+t+" center "+this.position.x+" "+this.position.y+" "+this.position.z+"\n"+t+" rotation "+i+" "+r+" "+s+" "+e+"\n"+t+" children [\n";this.geometry&&(n+=this.geometry.vrml(t,this.material));for(var a=0;a<this.children.length;a++)n+=this.children[a].vrml(t+" ")+",\n";return n+=" ]\n",n+="}"}updateMatrix(){this.matrix.setPosition(this.position),!1===this.useQuaternion&&this.rotation instanceof l.Vector3?this.matrix.setRotationFromEuler(this.rotation,this.eulerOrder):this.matrix.setRotationFromQuaternion(this.quaternion),1===this.scale.x&&1===this.scale.y&&1===this.scale.z||this.matrix.scale(this.scale),this.matrixWorldNeedsUpdate=!0}updateMatrixWorld(t){!0===this.matrixAutoUpdate&&this.updateMatrix(),!0!==this.matrixWorldNeedsUpdate&&!0!==t||(void 0===this.parent?this.matrixWorld.copy(this.matrix):this.matrixWorld.multiplyMatrices(this.parent.matrixWorld,this.matrix)),this.matrixWorldNeedsUpdate=!1;for(var e=0;e<this.children.length;e++)this.children[e].updateMatrixWorld(!0)}clone(t){void 0===t&&(t=new Object3D),t.name=this.name,t.up.copy(this.up),t.position.copy(this.position),t.rotation instanceof l.Vector3&&this.rotation instanceof l.Vector3?t.rotation.copy(this.rotation):t.rotation=this.rotation,t.eulerOrder=this.eulerOrder,t.scale.copy(this.scale),t.rotationAutoUpdate=this.rotationAutoUpdate,t.matrix.copy(this.matrix),t.matrixWorld.copy(this.matrixWorld),t.quaternion.copy(this.quaternion),t.matrixAutoUpdate=this.matrixAutoUpdate,t.matrixWorldNeedsUpdate=this.matrixWorldNeedsUpdate,t.useQuaternion=this.useQuaternion,t.visible=this.visible;for(var e=0;e<this.children.length;e++){var i=this.children[e];t.add(i.clone())}return t}setVisible(t){this.visible=t;for(var e=0;e<this.children.length;e++){this.children[e].setVisible(t)}}}class Scene extends Object3D{constructor(){super(...arguments),this.fog=null,this.overrideMaterial=null,this.matrixAutoUpdate=!1,this.__objects=[],this.__lights=[],this.__objectsAdded=[],this.__objectsRemoved=[]}__addObject(t){if(t instanceof Light)-1===this.__lights.indexOf(t)&&this.__lights.push(t),t.target&&void 0===t.target.parent&&this.add(t.target);else if(-1===this.__objects.indexOf(t)){this.__objects.push(t),this.__objectsAdded.push(t);var e=this.__objectsRemoved.indexOf(t);-1!==e&&this.__objectsRemoved.splice(e,1)}for(var i=0;i<t.children.length;i++)this.__addObject(t.children[i])}__removeObject(t){var e;t instanceof Light?-1!==(e=this.__lights.indexOf(t))&&this.__lights.splice(e,1):-1!==(e=this.__objects.indexOf(t))&&(this.__objects.splice(e,1),this.__objectsRemoved.push(t),-1!==this.__objectsAdded.indexOf(t)&&this.__objectsAdded.splice(e,1));for(var i=0;i<t.children.length;i++)this.__removeObject(t.children[i])}}class Light extends Object3D{constructor(t,e=1){super(),this.position=new l.Vector3(0,1,0),this.target=new Object3D,this.castShadow=!1,this.onlyShadow=!1,this.color=new b.Color(t),this.intensity=e}}const S=new l.Matrix4;class Projector{static unprojectVector(t,e){return e.projectionMatrixInverse.getInverse(e.projectionMatrix),S.multiplyMatrices(e.matrixWorld,e.projectionMatrixInverse),t.applyProjection(S)}static projectVector(t,e){return e.matrixWorldInverse.getInverse(e.matrixWorld),S.multiplyMatrices(e.projectionMatrix,e.matrixWorldInverse),t.applyProjection(S)}projectVector(t,e){return Projector.projectVector(t,e)}unprojectVector(t,e){return Projector.unprojectVector(t,e)}}var M=i(99);const z=(t,e)=>t.distance-e.distance,T=new l.Matrix4;class Raycaster{constructor(t,e,i,r){this.precision=1e-4,this.linePrecision=.2,this.ray=new l.Ray(t,e),this.ray.direction.lengthSq()>0&&this.ray.direction.normalize(),this.near=r||0,this.far=i||1/0}set(t,e){this.ray.set(t,e)}setFromCamera(t,e){e.ortho?(this.ray.origin.set(t.x,t.y,(e.near+e.far)/(e.near-e.far)).unproject(e),this.ray.direction.set(0,0,-1).transformDirection(e.matrixWorld)):(this.ray.origin.setFromMatrixPosition(e.matrixWorld),this.ray.direction.set(t.x,t.y,t.z),e.projectionMatrixInverse.getInverse(e.projectionMatrix),T.multiplyMatrices(e.matrixWorld,e.projectionMatrixInverse),this.ray.direction.applyProjection(T),this.ray.direction.sub(this.ray.origin).normalize())}intersectObjects(t,e){for(var i=[],r=0,s=e.length;r<s;r++)U(t,e[r],this,i);return i.sort(z),i}}const E=t=>Math.min(Math.max(t,-1),1);var L=new M.Sphere,F=new M.Cylinder,I=new M.Triangle,O=new l.Vector3,D=new l.Vector3,k=new l.Vector3,R=new l.Vector3,P=new l.Vector3;function U(t,e,i,r){if(P.getPositionFromMatrix(t.matrixWorld),void 0===e.intersectionShape)return r;var s,n,a,o,l,h,c,d,u,f,p,g,m,v,_=e.intersectionShape,y=i.linePrecision,b=(y*=t.matrixWorld.getMaxScaleOnAxis())*y;if(void 0!==e.boundingSphere&&e.boundingSphere instanceof M.Sphere&&(L.copy(e.boundingSphere),L.applyMatrix4(t.matrixWorld),!i.ray.isIntersectionSphere(L)))return r;for(s=0,n=_.triangle.length;s<n;s++)if(_.triangle[s]instanceof M.Triangle){if(I.copy(_.triangle[s]),I.applyMatrix4(t.matrixWorld),a=I.getNormal(),(o=i.ray.direction.dot(a))>=0)continue;if(O.subVectors(I.a,i.ray.origin),(c=a.dot(O)/o)<0)continue;D.copy(i.ray.direction).multiplyScalar(c).add(i.ray.origin),D.sub(I.a),k.copy(I.b).sub(I.a),R.copy(I.c).sub(I.a);var x=k.dot(R),w=k.lengthSq(),A=R.lengthSq();if((g=(w*D.dot(R)-x*D.dot(k))/(w*A-x*x))<0||g>1)continue;if((p=(D.dot(k)-g*x)/w)<0||p>1||p+g>1)continue;r.push({clickable:e,distance:c})}for(s=0,n=_.cylinder.length;s<n;s++)if(_.cylinder[s]instanceof M.Cylinder){if(F.copy(_.cylinder[s]),F.applyMatrix4(t.matrixWorld),O.subVectors(F.c1,i.ray.origin),l=O.dot(F.direction),h=O.dot(i.ray.direction),0===(u=1-(o=E(i.ray.direction.dot(F.direction)))*o))continue;m=(o*h-l)/u,v=(h-o*l)/u,D.copy(F.direction).multiplyScalar(m).add(F.c1),k.copy(i.ray.direction).multiplyScalar(v).add(i.ray.origin),d=R.subVectors(D,k).lengthSq();var C=F.radius*F.radius;if(d<=C){if((p=o*(g=c=(f=(o*l-h)*(o*l-h)-u*(O.lengthSq()-l*l-C))<=0?Math.sqrt(d):(h-o*l-Math.sqrt(f))/u)-l)<0||p*p>F.lengthSq()||g<0)continue;r.push({clickable:e,distance:c})}}for(s=0,n=_.line.length;s<n;s+=2){D.copy(_.line[s]),D.applyMatrix4(t.matrixWorld),k.copy(_.line[s+1]),k.applyMatrix4(t.matrixWorld),R.subVectors(k,D);var S=R.lengthSq();R.normalize(),O.subVectors(D,i.ray.origin);var z=O.dot(R);h=O.dot(i.ray.direction),0!==(u=1-(o=E(i.ray.direction.dot(R)))*o)&&(m=(o*h-z)/u,v=(h-o*z)/u,D.add(R.multiplyScalar(m)),k.copy(i.ray.direction).multiplyScalar(v).add(i.ray.origin),(d=R.subVectors(k,D).lengthSq())<b&&m*m<S&&r.push({clickable:e,distance:v}))}for(s=0,n=_.sphere.length;s<n;s++)if(_.sphere[s]instanceof M.Sphere&&(L.copy(_.sphere[s]),L.applyMatrix4(t.matrixWorld),i.ray.isIntersectionSphere(L))){D.subVectors(L.center,i.ray.origin);var T=D.dot(i.ray.direction);if(f=T*T-(D.lengthSq()-L.radius*L.radius),T<0)return r;c=f<=0?T:T-Math.sqrt(f),r.push({clickable:e,distance:c})}return r}class UVMapping{}class Texture extends EventDispatcher{constructor(t,e){super(),this.id=B++,this.name="",this.image=t,this.mapping=new UVMapping,this.wrapS=c,this.wrapT=c,this.anisotropy=1,e?(this.format=v,this.type=g,this.premultiplyAlpha=!1,this.flipY=!1,this.unpackAlignment=1,this.magFilter=u,this.minFilter=u):(this.format=m,this.type=p,this.offset=new l.Vector2(0,0),this.repeat=new l.Vector2(1,1),this.premultiplyAlpha=!1,this.flipY=!0,this.unpackAlignment=4,this.magFilter=d,this.minFilter=f),this.needsUpdate=!1,this.onUpdate=null}clone(t=new Texture){return t.image=this.image,t.mapping=this.mapping,t.wrapS=this.wrapS,t.wrapT=this.wrapT,t.magFilter=this.magFilter,t.minFilter=this.minFilter,t.anisotropy=this.anisotropy,t.format=this.format,t.type=this.type,t.offset.copy(this.offset),t.repeat.copy(this.repeat),t.premultiplyAlpha=this.premultiplyAlpha,t.flipY=this.flipY,t.unpackAlignment=this.unpackAlignment,t}dispose(){this.dispatchEvent({type:"dispose"})}}let B=0;class ImposterMaterial extends Material{constructor(t){super(),this.color=new b.Color(16777215),this.ambient=new b.Color(1048575),this.emissive=new b.Color(0),this.imposter=!0,this.wrapAround=!1,this.wrapRGB=new l.Vector3(1,1,1),this.map=void 0,this.lightMap=null,this.specularMap=null,this.envMap=null,this.reflectivity=1,this.refractionRatio=.98,this.fog=!0,this.wireframe=!1,this.wireframeLinewidth=1,this.wireframeLinecap="round",this.wireframeLinejoin="round",this.shading=o.SmoothShading,this.shaderID=null,this.vertexColors=r.NoColors,this.skinning=!1,this.setValues(t)}clone(t=new ImposterMaterial){return super.clone.call(this,t),t.color.copy(this.color),t.ambient.copy(this.ambient),t.emissive.copy(this.emissive),t.wrapAround=this.wrapAround,t.wrapRGB.copy(this.wrapRGB),t.map=this.map,t.lightMap=this.lightMap,t.specularMap=this.specularMap,t.envMap=this.envMap,t.combine=this.combine,t.reflectivity=this.reflectivity,t.refractionRatio=this.refractionRatio,t.fog=this.fog,t.shading=this.shading,t.shaderID=this.shaderID,t.vertexColors=this.vertexColors,t.skinning=this.skinning,t.morphTargets=this.morphTargets,t.morphNormals=this.morphNormals,t}}class InstancedMaterial extends Material{constructor(t){super(),this.color=new b.Color(16777215),this.ambient=new b.Color(1048575),this.emissive=new b.Color(0),this.wrapAround=!1,this.wrapRGB=new l.Vector3(1,1,1),this.map=void 0,this.lightMap=null,this.specularMap=null,this.envMap=null,this.reflectivity=1,this.refractionRatio=.98,this.fog=!0,this.wireframe=!1,this.wireframeLinewidth=1,this.wireframeLinecap="round",this.wireframeLinejoin="round",this.shading=o.SmoothShading,this.shaderID="instanced",this.vertexColors=r.NoColors,this.skinning=!1,this.sphere=null,this.setValues(t)}clone(t=new InstancedMaterial){return super.clone.call(this,t),t.color.copy(this.color),t.ambient.copy(this.ambient),t.emissive.copy(this.emissive),t.wrapAround=this.wrapAround,t.wrapRGB.copy(this.wrapRGB),t.map=this.map,t.lightMap=this.lightMap,t.specularMap=this.specularMap,t.envMap=this.envMap,t.combine=this.combine,t.reflectivity=this.reflectivity,t.refractionRatio=this.refractionRatio,t.fog=this.fog,t.shading=this.shading,t.shaderID=this.shaderID,t.vertexColors=this.vertexColors,t.skinning=this.skinning,t.morphTargets=this.morphTargets,t.morphNormals=this.morphNormals,t.sphere=this.sphere,t}}class MeshLambertMaterial extends Material{constructor(t){super(),this.color=new b.Color(16777215),this.ambient=new b.Color(1048575),this.emissive=new b.Color(0),this.wrapAround=!1,this.wrapRGB=new l.Vector3(1,1,1),this.map=void 0,this.lightMap=null,this.specularMap=null,this.envMap=null,this.reflectivity=1,this.refractionRatio=.98,this.fog=!0,this.wireframe=!1,this.wireframeLinewidth=1,this.wireframeLinecap="round",this.wireframeLinejoin="round",this.shading=o.SmoothShading,this.shaderID="lambert",this.vertexColors=r.NoColors,this.skinning=!1,this.setValues(t)}clone(t=new MeshLambertMaterial){return super.clone.call(this,t),t.color.copy(this.color),t.ambient.copy(this.ambient),t.emissive.copy(this.emissive),t.wrapAround=this.wrapAround,t.wrapRGB.copy(this.wrapRGB),t.map=this.map,t.lightMap=this.lightMap,t.specularMap=this.specularMap,t.envMap=this.envMap,t.combine=this.combine,t.reflectivity=this.reflectivity,t.refractionRatio=this.refractionRatio,t.fog=this.fog,t.shading=this.shading,t.shaderID=this.shaderID,t.vertexColors=this.vertexColors,t.skinning=this.skinning,t.morphTargets=this.morphTargets,t.morphNormals=this.morphNormals,t}}class MeshDoubleLambertMaterial extends MeshLambertMaterial{constructor(t){super(t),this.shaderID="lambertdouble",this.side=a,this.outline=!1}clone(t=new MeshDoubleLambertMaterial){return super.clone.call(this,t),t}}class MeshOutlineMaterial extends Material{constructor(t){super(),t=t||{},this.fog=!0,this.shaderID="outline",this.wireframe=!1,this.outlineColor=b.CC.color(t.color||new b.Color(0,0,0)),this.outlineWidth=t.width||.1,this.outlinePushback=t.pushback||1,this.outlineMaxPixels=t.maxpixels||0}clone(t=new MeshOutlineMaterial){return super.clone.call(this,t),t.fog=this.fog,t.shaderID=this.shaderID,t.wireframe=this.wireframe,t.outlineColor=this.outlineColor,t.outlineWidth=this.outlineWidth,t.outlinePushback=this.outlinePushback,t.outlineMaxPixels=this.outlineMaxPixels,t}}class SphereImposterMaterial extends ImposterMaterial{constructor(t){super(t),this.shaderID="sphereimposter",this.setValues(t)}clone(t=new SphereImposterMaterial){return super.clone.call(this,t),t}}class SphereImposterOutlineMaterial extends ImposterMaterial{constructor(t){super(t),t=t||{},this.shaderID="sphereimposteroutline",this.outlineColor=b.CC.color(t.color||new b.Color(0,0,0)),this.outlineWidth=t.width||.1,this.outlinePushback=t.pushback||1,this.outlineMaxPixels=t.maxpixels||0,this.setValues(t)}clone(t=new SphereImposterOutlineMaterial){return super.clone.call(this,t),t.outlineColor=this.outlineColor,t.outlineWidth=this.outlineWidth,t.outlinePushback=this.outlinePushback,t.outlineMaxPixels=this.outlineMaxPixels,t}}class SpriteMaterial extends Material{constructor(t){super(),this.color=new b.Color(16777215),this.map=new Texture,this.useScreenCoordinates=!0,this.fog=!1,this.uvOffset=new l.Vector2(0,0),this.uvScale=new l.Vector2(1,1),this.depthTest=!this.useScreenCoordinates,this.sizeAttenuation=!this.useScreenCoordinates,this.screenOffset=this.screenOffset,this.scaleByViewPort=!this.sizeAttenuation,this.alignment=h.center.clone(),this.setValues(t),void 0===(t=t||{}).depthTest&&(this.depthTest=!this.useScreenCoordinates),void 0===t.sizeAttenuation&&(this.sizeAttenuation=!this.useScreenCoordinates),void 0===t.scaleByViewPort&&(this.scaleByViewPort=!this.sizeAttenuation)}clone(t=new SpriteMaterial){return super.clone.call(this,t),t.color.copy(this.color),t.map=this.map,t.useScreenCoordinates=this.useScreenCoordinates,t.screenOffset=this.screenOffset,t.sizeAttenuation=this.sizeAttenuation,t.scaleByViewport=this.scaleByViewPort,t.alignment.copy(this.alignment),t.uvOffset.copy(this.uvOffset),t}}class StickImposterMaterial extends ImposterMaterial{constructor(t){super(t),this.shaderID="stickimposter",this.setValues(t)}clone(t=new StickImposterMaterial){return super.clone.call(this,t),t}}class StickImposterOutlineMaterial extends ImposterMaterial{constructor(t={}){super(t),this.shaderID="stickimposteroutline",this.outlineColor=new b.Color(0,0,0),this.outlineWidth=.1,this.outlinePushback=1,this.outlineMaxPixels=0,t.color&&(this.outlineColor=b.CC.color(t.color)),t.width&&(this.outlineWidth=t.width),t.pushback&&(this.outlinePushback=t.pushback),t.maxpixels&&(this.outlineMaxPixels=t.maxpixels),this.setValues(t)}clone(t=new StickImposterOutlineMaterial){return super.clone.call(this,t),t.outlineColor=this.outlineColor,t.outlineWidth=this.outlineWidth,t.outlinePushback=this.outlinePushback,t.outlineMaxPixels=this.outlineMaxPixels,t}}class VolumetricMaterial extends Material{constructor(t){super(),this.transparent=!1,this.volumetric=!0,this.color=new b.Color(16777215),this.transferfn=null,this.map=void 0,this.extent=[],this.maxdepth=100,this.unit=0,this.texmatrix=null,this.transfermin=-1,this.transfermax=1,this.subsamples=5,this.shaderID="volumetric",this.side=s,this.setValues(t)}clone(t=new VolumetricMaterial){return super.clone.call(this,t),t.transparent=this.transparent,t.volumetric=this.volumetric,t.color=this.color,t.transferfn=this.transferfn,t.map=this.map,t.extent=this.extent,t.maxdepth=this.maxdepth,t.unit=this.unit,t.texmatrix=this.texmatrix,t.transfermin=this.transfermin,t.transfermax=this.transfermax,t.subsamples=this.subsamples,t.shaderID=this.shaderID,t.side=this.side,t}}var N;!function(t){t[t.LineStrip=0]="LineStrip",t[t.LinePieces=1]="LinePieces"}(N||(N={}));class Line extends Object3D{constructor(t,e=new LineBasicMaterial({color:16777215*Math.random()}),i=N.LineStrip){super(),this.geometry=t,this.material=e,this.type=i}clone(t=new Line(this.geometry,this.material,this.type)){return super.clone.call(this,t),t}}class Mesh extends Object3D{constructor(t,e){super(),this.geometry=t,this.material=e}clone(t){return void 0===t&&(t=new Mesh(this.geometry,this.material)),super.clone.call(this,t),t}}class Sprite extends Object3D{constructor(t=new SpriteMaterial){super(),this.material=t,this.rotation3d=this.rotation,this.rotation=0}updateMatrix(){this.matrix.setPosition(this.position),this.rotation3d.set(0,0,this.rotation),this.matrix.setRotationFromEuler(this.rotation3d),1===this.scale.x&&1===this.scale.y||this.matrix.scale(this.scale),this.matrixWorldNeedsUpdate=!0}clone(t=new Sprite(this.material)){return Object3D.prototype.clone.call(this,t),t}}const G={opacity:{type:"f",value:1},fogColor:{type:"c",value:new b.Color(1,1,1)},fogNear:{type:"f",value:1},fogFar:{type:"f",value:2e3}},V={vertexShader:"uniform mat4 modelViewMatrix;\nuniform mat4 projectionMatrix;\nuniform mat4 viewMatrix;\nuniform mat3 normalMatrix;\n\nattribute vec3 position;\nattribute vec3 color;\n\nvarying vec3 vColor;\n\nvoid main() {\n\n    vColor = color;\n    vec4 mvPosition = modelViewMatrix * vec4( position, 1.0 );\n    gl_Position = projectionMatrix * mvPosition;\n\n}".replace("#define GLSLIFY 1",""),fragmentShader:"uniform mat4 viewMatrix;\nuniform float opacity;\nuniform vec3 fogColor;\nuniform float fogNear;\nuniform float fogFar;\nvarying vec3 vColor;\n//DEFINEFRAGCOLOR\nvoid main() {\n    gl_FragColor = vec4( vColor, opacity );\n    float depth = gl_FragCoord.z / gl_FragCoord.w;\n    float fogFactor = smoothstep( fogNear, fogFar, depth );\n    gl_FragColor = mix( gl_FragColor, vec4( fogColor, gl_FragColor.w ), fogFactor );\n}".replace("#define GLSLIFY 1",""),uniforms:G},j={opacity:{type:"f",value:1},fogColor:{type:"c",value:new b.Color(1,1,1)},fogNear:{type:"f",value:1},fogFar:{type:"f",value:2e3},directionalLightColor:{type:"fv",value:[]},directionalLightDirection:{type:"fv",value:[]}},H={fragmentShader:"uniform mat4 viewMatrix;\nuniform float opacity;\n\nuniform vec3 fogColor;\nuniform float fogNear;\nuniform float fogFar;\n#ifdef SHADED\nuniform highp sampler2D shading;\n#endif\nvarying vec3 vLightFront;\nvarying vec3 vColor;\n//DEFINEFRAGCOLOR\n\nvoid main() {\n\n    gl_FragColor = vec4( vec3 ( 1.0 ), opacity );\n\n    #ifndef WIREFRAME\n    gl_FragColor.xyz *= vLightFront;\n    #endif\n\n#ifdef SHADED\n    ivec2 dim = textureSize(shading,0);\n    float shadowFactor = texture2D(shading,vec2(gl_FragCoord.x/float(dim.x),gl_FragCoord.y/float(dim.y))).r;\n    vColor *= shadowFactor;\n#endif\n    gl_FragColor = gl_FragColor * vec4( vColor, opacity );\n    float depth = gl_FragCoord.z / gl_FragCoord.w;\n\n    float fogFactor = smoothstep( fogNear, fogFar, depth );\n\n    gl_FragColor = mix( gl_FragColor, vec4( fogColor, gl_FragColor.w ), fogFactor );\n\n}\n\n\n".replace("#define GLSLIFY 1",""),vertexShader:"\n\nuniform mat4 modelViewMatrix;\nuniform mat4 projectionMatrix;\nuniform mat4 viewMatrix;\nuniform mat3 normalMatrix;\nuniform vec3 directionalLightColor[ 1 ];\nuniform vec3 directionalLightDirection[ 1 ];\n\nattribute vec3 offset;\nattribute vec3 position;\nattribute vec3 normal;\nattribute vec3 color;\nattribute float radius;\n\nvarying vec3 vColor;\nvarying vec3 vLightFront;\n\nvoid main() {\n\n    vColor = color;\n\n    vec3 objectNormal = normal;\n    vec3 transformedNormal = normalMatrix * objectNormal;\n    vec4 mvPosition = modelViewMatrix * vec4( position * radius + offset, 1.0 );\n\n    vLightFront = vec3( 0.0 );\n\n    transformedNormal = normalize( transformedNormal );\n\n    vec4 lDirection = viewMatrix * vec4( directionalLightDirection[ 0 ], 0.0 );\n    vec3 dirVector = normalize( lDirection.xyz );\n    float dotProduct = dot( transformedNormal, dirVector );\n    vec3 directionalLightWeighting = vec3( max( dotProduct, 0.0 ) );\n\n    vLightFront += directionalLightColor[ 0 ] * directionalLightWeighting;\n\n    gl_Position = projectionMatrix * mvPosition;\n}\n\n".replace("#define GLSLIFY 1",""),uniforms:j},W={opacity:{type:"f",value:1},fogColor:{type:"c",value:new b.Color(1,1,1)},fogNear:{type:"f",value:1},fogFar:{type:"f",value:2e3},directionalLightColor:{type:"fv",value:[]},directionalLightDirection:{type:"fv",value:[]}},q={fragmentShader:"uniform mat4 viewMatrix;\nuniform float opacity;\n\nuniform vec3 fogColor;\nuniform float fogNear;\nuniform float fogFar;\n#ifdef SHADED\nuniform highp sampler2D shading;\n#endif\nvarying vec3 vLightFront;\nvarying vec3 vColor;\n//DEFINEFRAGCOLOR\n\nvoid main() {\n\n    gl_FragColor = vec4( vec3 ( 1.0 ), opacity );\n\n    #ifndef WIREFRAME\n    gl_FragColor.xyz *= vLightFront;\n    #endif\n#ifdef SHADED\n    ivec2 dim = textureSize(shading,0);\n    float shadowFactor = texture2D(shading,vec2(gl_FragCoord.x/float(dim.x),gl_FragCoord.y/float(dim.y))).r;\n    vColor *= shadowFactor;\n#endif\n    gl_FragColor = gl_FragColor * vec4( vColor, opacity );\n    float depth = gl_FragCoord.z / gl_FragCoord.w;\n\n    float fogFactor = smoothstep( fogNear, fogFar, depth );\n\n    gl_FragColor = mix( gl_FragColor, vec4( fogColor, gl_FragColor.w ), fogFactor );\n\n}".replace("#define GLSLIFY 1",""),vertexShader:"\nuniform mat4 modelViewMatrix;\nuniform mat4 projectionMatrix;\nuniform mat4 viewMatrix;\nuniform mat3 normalMatrix;\nuniform vec3 directionalLightColor[ 1 ];\nuniform vec3 directionalLightDirection[ 1 ];\n\nattribute vec3 position;\nattribute vec3 normal;\nattribute vec3 color;\n\nvarying vec3 vColor;\nvarying vec3 vLightFront;\n\nvoid main() {\n\n    vColor = color;\n\n    vec3 objectNormal = normal;\n    vec3 transformedNormal = normalMatrix * objectNormal;\n    vec4 mvPosition = modelViewMatrix * vec4( position, 1.0 );\n\n    vLightFront = vec3( 0.0 );\n\n    transformedNormal = normalize( transformedNormal );\n\n    vec4 lDirection = viewMatrix * vec4( directionalLightDirection[ 0 ], 0.0 );\n    vec3 dirVector = normalize( lDirection.xyz );\n    float dotProduct = dot( transformedNormal, dirVector );\n    vec3 directionalLightWeighting = vec3( max( dotProduct, 0.0 ) );\n\n    vLightFront += directionalLightColor[ 0 ] * directionalLightWeighting;\n\n    gl_Position = projectionMatrix * mvPosition;\n}".replace("#define GLSLIFY 1",""),uniforms:W},Y={opacity:{type:"f",value:1},fogColor:{type:"c",value:new b.Color(1,1,1)},fogNear:{type:"f",value:1},fogFar:{type:"f",value:2e3},directionalLightColor:{type:"fv",value:[]},directionalLightDirection:{type:"fv",value:[]}},Z={fragmentShader:"\n#ifdef SHADED\nuniform highp sampler2D shading;\n#endif\n\nuniform mat4 viewMatrix;\nuniform float opacity;\n\nuniform vec3 fogColor;\nuniform float fogNear;\nuniform float fogFar;\n\nvarying vec3 vLightFront;\nvarying vec3 vLightBack;\n\nvarying vec3 vColor;\n\n\n//DEFINEFRAGCOLOR\n\nvoid main() {\n\n    gl_FragColor = vec4( vec3 ( 1.0 ), opacity );\n\n    #ifndef WIREFRAME\n    if ( gl_FrontFacing )\n       gl_FragColor.xyz *= vLightFront;\n    else\n       gl_FragColor.xyz *= vLightBack;\n    #endif\n\n    vec3 color = vColor;\n#ifdef SHADED\n    ivec2 dim = textureSize(shading,0);\n    float shadowFactor = texture2D(shading,vec2(gl_FragCoord.x/float(dim.x),gl_FragCoord.y/float(dim.y))).r;\n    color *= shadowFactor;\n#endif\n    gl_FragColor = gl_FragColor * vec4( color, opacity );\n    float depth = gl_FragCoord.z / gl_FragCoord.w;\n\n    float fogFactor = smoothstep( fogNear, fogFar, depth );\n\n    gl_FragColor = mix( gl_FragColor, vec4( fogColor, gl_FragColor.w ), fogFactor );\n\n}\n\n\n".replace("#define GLSLIFY 1",""),vertexShader:"\n\nuniform mat4 modelViewMatrix;\nuniform mat4 projectionMatrix;\nuniform mat4 viewMatrix;\nuniform mat3 normalMatrix;\nuniform vec3 directionalLightColor[ 1 ];\nuniform vec3 directionalLightDirection[ 1 ];\n\nattribute vec3 position;\nattribute vec3 normal;\nattribute vec3 color;\n\nvarying vec3 vColor;\nvarying vec3 vLightFront;\nvarying vec3 vLightBack;\n\nvoid main() {\n\n    vColor = color;\n\n    vec3 objectNormal = normal;\n    vec3 transformedNormal = normalMatrix * objectNormal;\n    vec4 mvPosition = modelViewMatrix * vec4( position, 1.0 );\n\n    vLightFront = vec3( 0.0 );\n    vLightBack = vec3( 0.0 );\n\n    transformedNormal = normalize( transformedNormal );\n\n    vec4 lDirection = viewMatrix * vec4( directionalLightDirection[ 0 ], 0.0 );\n    vec3 dirVector = normalize( lDirection.xyz );\n    float dotProduct = dot( transformedNormal, dirVector );\n    vec3 directionalLightWeighting = vec3( max( dotProduct, 0.0 ) );\n    vec3 directionalLightWeightingBack = vec3( max( -dotProduct, 0.0 ) );\n\n    vLightFront += directionalLightColor[ 0 ] * directionalLightWeighting;\n    vLightBack += directionalLightColor[ 0 ] * directionalLightWeightingBack;\n\n    gl_Position = projectionMatrix * mvPosition;\n}\n\n".replace("#define GLSLIFY 1",""),uniforms:Y},X={opacity:{type:"f",value:1},outlineColor:{type:"c",value:new b.Color(0,0,0)},fogColor:{type:"c",value:new b.Color(1,1,1)},fogNear:{type:"f",value:1},fogFar:{type:"f",value:2e3},outlineWidth:{type:"f",value:.1},outlinePushback:{type:"f",value:1},outlineMaxPixels:{type:"f",value:0}},K={fragmentShader:"\n\nuniform float opacity;\nuniform vec3 outlineColor;\nuniform vec3 fogColor;\nuniform float fogNear;\nuniform float fogFar;\n//DEFINEFRAGCOLOR\n\nvoid main() {\n    gl_FragColor = vec4( outlineColor, 1 );\n\n    float depth = gl_FragCoord.z / gl_FragCoord.w;\n    float fogFactor = smoothstep( fogNear, fogFar, depth );\n    gl_FragColor = mix( gl_FragColor, vec4( fogColor, gl_FragColor.w ), fogFactor );    \n}\n\n\n".replace("#define GLSLIFY 1",""),vertexShader:"\n\nuniform mat4 modelViewMatrix;\nuniform mat4 projectionMatrix;\nuniform float outlineWidth;\nuniform float outlinePushback;\nuniform float vWidth;\nuniform float vHeight;\nuniform float outlineMaxPixels;\n\nattribute vec3 position;\nattribute vec3 normal;\nattribute vec3 color;\n\nvoid main() {\n\n    vec4 norm = modelViewMatrix*vec4(normalize(normal),0.0);\n    vec4 mvPosition = modelViewMatrix * vec4( position, 1.0 );\n    mvPosition.xy += norm.xy*outlineWidth;\n    vec4 outpos = projectionMatrix * mvPosition;\n\n    if(outlineMaxPixels > 0.0) {\n        vec4 unadjusted = projectionMatrix*modelViewMatrix * vec4( position, 1.0 );\n        float w = outpos.w;\n        //normalize homogeneous coords\n        unadjusted /= unadjusted.w;\n        outpos /= outpos.w;\n        vec2 diff = outpos.xy-unadjusted.xy;\n        //put into pixels\n        diff.x *= vWidth;\n        diff.y *= vHeight;\n        if ( length(diff) > outlineMaxPixels) {\n            vec2 ndiff = normalize(diff)*outlineMaxPixels;\n            ndiff.x /= vWidth;\n            ndiff.y /= vHeight;\n            outpos.xy = unadjusted.xy;\n            outpos.xy += ndiff;\n        }\n        outpos *= w; //if I don't do this things blow up\n    }\n    gl_Position = outpos;\n    mvPosition.z -= outlinePushback; //go backwards in model space\n    vec4 pushpos = projectionMatrix*mvPosition; //project to get z in projection space, I'm probably missing some simple math to do the same thing..\n    gl_Position.z = gl_Position.w*pushpos.z/pushpos.w;\n}\n\n".replace("#define GLSLIFY 1",""),uniforms:X},Q={fragmentShader:"uniform sampler2D colormap;\nvarying highp vec2 vTexCoords;\nuniform vec2 dimensions;\n//DEFINEFRAGCOLOR\nvoid main (void) {\n   gl_FragColor = texture2D(colormap, vTexCoords);\n\n   //gl_FragColor.g = gl_FragColor.b =  gl_FragColor.r; //debug shading \n}\n        ".replace("#define GLSLIFY 1",""),vertexShader:"attribute vec2 vertexPosition;\nvarying highp vec2 vTexCoords;\nconst vec2 scale = vec2(0.5, 0.5);\n\nvoid main() {\n   vTexCoords  = vertexPosition * scale + scale; // scale vertex attribute to [0,1] range\n   gl_Position = vec4(vertexPosition, 0.0, 1.0);\n}\n        ".replace("#define GLSLIFY 1",""),uniforms:{}},$={fragmentShader:"uniform highp sampler2D colormap;\nvarying highp vec2 vTexCoords;\n\n\n// Basic FXAA implementation based on the code on geeks3d.com \n#define FXAA_REDUCE_MIN (1.0/ 128.0)\n#define FXAA_REDUCE_MUL (1.0 / 8.0)\n#define FXAA_SPAN_MAX 8.0\n\n\nvec4 applyFXAA(vec2 fragCoord, highp sampler2D tex)\n{\n    vec4 color;\n    ivec2 dim = textureSize(tex,0);\n    vec2 dimensions = vec2(float(dim.x),float(dim.y));\n    vec2 inverseVP = vec2(1.0 / dimensions.x, 1.0 / dimensions.y);\n    vec4 rgbNW = texture2D(tex, fragCoord + vec2(-1.0, -1.0) * inverseVP);\n    vec4 rgbNE = texture2D(tex, fragCoord + vec2(1.0, -1.0) * inverseVP);\n    vec4 rgbSW = texture2D(tex, fragCoord + vec2(-1.0, 1.0) * inverseVP);\n    vec4 rgbSE = texture2D(tex, fragCoord + vec2(1.0, 1.0) * inverseVP);\n    vec4 rgbM  = texture2D(tex, fragCoord );\n    vec3 luma = vec3(0.299, 0.587, 0.114);\n    float lumaNW = dot(rgbNW.xyz, luma);\n    float lumaNE = dot(rgbNE.xyz, luma);\n    float lumaSW = dot(rgbSW.xyz, luma);\n    float lumaSE = dot(rgbSE.xyz, luma);\n    float lumaM  = dot(rgbM.xyz,  luma);\n    float lumaMin = min(lumaM, min(min(lumaNW, lumaNE), min(lumaSW, lumaSE)));\n    float lumaMax = max(lumaM, max(max(lumaNW, lumaNE), max(lumaSW, lumaSE)));\n\n    vec2 dir;\n    dir.x = -((lumaNW + lumaNE) - (lumaSW + lumaSE));\n    dir.y =  ((lumaNW + lumaSW) - (lumaNE + lumaSE));\n\n    float dirReduce = max((lumaNW + lumaNE + lumaSW + lumaSE) *\n                        (0.25 * FXAA_REDUCE_MUL), FXAA_REDUCE_MIN);\n\n    float rcpDirMin = 1.0 / (min(abs(dir.x), abs(dir.y)) + dirReduce);\n    dir = min(vec2(FXAA_SPAN_MAX, FXAA_SPAN_MAX),\n            max(vec2(-FXAA_SPAN_MAX, -FXAA_SPAN_MAX),\n            dir * rcpDirMin)) * inverseVP;\n\n    vec4 rgbA = 0.5 * (\n        texture2D(tex, fragCoord + dir * (1.0 / 3.0 - 0.5)) +\n        texture2D(tex, fragCoord  + dir * (2.0 / 3.0 - 0.5)));\n    vec4 rgbB = rgbA * 0.5 + 0.25 * (\n        texture2D(tex, fragCoord  + dir * -0.5) +\n        texture2D(tex, fragCoord  + dir * 0.5));\n\n    float lumaB = dot(rgbB.xyz, luma);\n    if ((lumaB < lumaMin) || (lumaB > lumaMax))\n        color = rgbA;\n    else\n        color = rgbB;\n\n    return color;\n}\n\n\n//DEFINEFRAGCOLOR\nvoid main (void) {\n    ivec2 dim = textureSize(colormap,0);\n\n  gl_FragColor = applyFXAA(vTexCoords, colormap);\n}\n        ".replace("#define GLSLIFY 1",""),vertexShader:"attribute vec2 vertexPosition;\nvarying highp vec2 vTexCoords;\nconst vec2 scale = vec2(0.5, 0.5);\n\nvoid main() {\n   vTexCoords  = vertexPosition * scale + scale; // scale vertex attribute to [0,1] range\n   gl_Position = vec4(vertexPosition, 0.0, 1.0);\n}\n        ".replace("#define GLSLIFY 1",""),uniforms:{}},J={opacity:{type:"f",value:1},fogColor:{type:"c",value:new b.Color(1,1,1)},fogNear:{type:"f",value:1},fogFar:{type:"f",value:2e3},directionalLightColor:{type:"fv",value:[]},directionalLightDirection:{type:"fv",value:[]}},tt={vertexShader:"uniform mat4 modelViewMatrix;\nuniform mat4 projectionMatrix;\nuniform mat4 viewMatrix;\nuniform mat3 normalMatrix;\nuniform vec3 directionalLightColor[ 1 ];\nuniform vec3 directionalLightDirection[ 1 ];\n\nattribute vec3 position;\nattribute vec3 normal;\nattribute vec3 color;\n\nvarying vec2 mapping;\nvarying vec3 vColor;\nvarying float rval;\nvarying vec3 vLight;\nvarying vec3 center;\n\nvoid main() {\n\n    vColor = color;\n    vec4 mvPosition = modelViewMatrix * vec4( position, 1.0 );\n    center = mvPosition.xyz;\n    vec4 projPosition = projectionMatrix * mvPosition;\n    vec4 adjust = projectionMatrix* vec4(normal,0.0); adjust.z = 0.0; adjust.w = 0.0;\n    vec4 lDirection = viewMatrix * vec4( directionalLightDirection[ 0 ], 0.0 );\n    vLight = normalize( lDirection.xyz );\n    mapping = normal.xy;\n    rval = abs(normal.x);\n    gl_Position = projPosition+adjust;\n\n}\n".replace("#define GLSLIFY 1",""),fragmentShader:"\nuniform mat4 viewMatrix;\nuniform float opacity;\nuniform mat4 projectionMatrix;\n\nuniform vec3 fogColor;\nuniform float fogNear;\nuniform float fogFar;\nuniform float uDepth;\nuniform vec3 directionalLightColor[ 1 ];\n\nvarying vec3 vColor;\nvarying vec2 mapping;\nvarying float rval;\nvarying vec3 vLight;\nvarying vec3 center;\n\n#ifdef SHADED\nuniform highp sampler2D shading;\n#endif\n\n//DEFINEFRAGCOLOR\n\nvoid main() {\n    float lensqr = dot(mapping,mapping);\n    float rsqr = rval*rval;\n    if(lensqr > rsqr)\n       discard;\n    float z = sqrt(rsqr-lensqr);\n    vec3 cameraPos = center+ vec3(mapping.x,mapping.y,z);\n    vec4 clipPos = projectionMatrix * vec4(cameraPos, 1.0);\n    float ndcDepth = clipPos.z / clipPos.w;\n    gl_FragDepthEXT = ((gl_DepthRange.diff * ndcDepth) + gl_DepthRange.near + gl_DepthRange.far) / 2.0;\n    vec3 norm = normalize(vec3(mapping.x,mapping.y,z));\n    float dotProduct = dot( norm, vLight );\n    vec3 directionalLightWeighting = vec3( max( dotProduct, 0.0 ) );\n    vec3 vLight = directionalLightColor[ 0 ] * directionalLightWeighting;\n    vec3 color = vLight*vColor;\n#ifdef SHADED\n    ivec2 dim = textureSize(shading,0);\n    float shadowFactor = texture2D(shading,vec2(gl_FragCoord.x/float(dim.x),gl_FragCoord.y/float(dim.y))).r;\n    color *= shadowFactor;\n#endif    \n    gl_FragColor = vec4(color, opacity*opacity );\n    float fogFactor = smoothstep( fogNear, fogFar, gl_FragDepthEXT/gl_FragCoord.w );\n    gl_FragColor = mix( gl_FragColor, vec4( fogColor, gl_FragColor.w ), fogFactor );\n}\n\n".replace("#define GLSLIFY 1",""),uniforms:J},et={opacity:{type:"f",value:1},outlineColor:{type:"c",value:new b.Color(0,0,0)},fogColor:{type:"c",value:new b.Color(1,1,1)},fogNear:{type:"f",value:1},fogFar:{type:"f",value:2e3},outlineWidth:{type:"f",value:.1},outlinePushback:{type:"f",value:1},outlineMaxPixels:{type:"f",value:0}},it={fragmentShader:"\n\nuniform float opacity;\nuniform vec3 outlineColor;\nuniform vec3 fogColor;\nuniform float fogNear;\nuniform float fogFar;\nuniform mat4 projectionMatrix;\nvarying vec2 mapping;\nvarying float rval;\nvarying vec3 center;\n\nuniform float outlinePushback;\n\n//DEFINEFRAGCOLOR\n\nvoid main() {\n    float lensqr = dot(mapping,mapping);\n    float rsqr = rval*rval;\n    if(lensqr > rsqr)\n       discard;\n    float z = sqrt(rsqr-lensqr);\n    vec3 cameraPos = center+ vec3(mapping.x,mapping.y,z-outlinePushback);\n    vec4 clipPos = projectionMatrix * vec4(cameraPos, 1.0);\n    float ndcDepth = clipPos.z / clipPos.w;\n    gl_FragDepthEXT = ((gl_DepthRange.diff * ndcDepth) + gl_DepthRange.near + gl_DepthRange.far) / 2.0;\n    gl_FragColor = vec4(outlineColor, 1 );\n}\n\n\n".replace("#define GLSLIFY 1",""),vertexShader:"\n\nuniform mat4 modelViewMatrix;\nuniform mat4 projectionMatrix;\nuniform float outlineWidth;\nuniform float outlinePushback;\nuniform float outlineMaxPixels;\nuniform float vWidth;\nuniform float vHeight;\n\nattribute vec3 position;\nattribute vec3 normal;\nattribute vec3 color;\n\nvarying vec2 mapping;\nvarying float rval;\nvarying vec3 center;\n\nvoid main() {\n\n    vec4 mvPosition = modelViewMatrix * vec4( position, 1.0 );\n    center = mvPosition.xyz;\n    vec4 projPosition = projectionMatrix * mvPosition;\n    vec2 norm = normal.xy + vec2(sign(normal.x)*outlineWidth,sign(normal.y)*outlineWidth);\n\n    vec4 adjust = projectionMatrix* vec4(norm,normal.z,1.0); \n    mapping = norm.xy;\n    rval = abs(norm.x);\n    gl_Position = projPosition+vec4(adjust.xy,0.0,0.0);\n\n    if(outlineMaxPixels > 0.0) {\n        vec4 unadjusted = projectionMatrix*vec4(center.x+normal.x, center.y,center.z,1.0); \n        vec4 ccoord = projectionMatrix*vec4(center.xyz,1.0);\n        adjust = projectionMatrix* vec4(center.x+norm.x,center.y,center.z,1.0); \n        //subtract center \n        unadjusted.xyz -= ccoord.xyz;\n        adjust.xyz -= ccoord.xyz;\n        unadjusted /= unadjusted.w;\n        adjust /= adjust.w;\n        float diff = abs(adjust.x-unadjusted.x);\n        diff *= vWidth;\n        if(diff > outlineMaxPixels) {\n            \n            float fixlen = abs(unadjusted.x) + outlineMaxPixels/vWidth;\n            //adjsut reval by ratio of lengths\n            rval *= fixlen/abs(adjust.x);\n        }\n\n    }\n}\n\n".replace("#define GLSLIFY 1",""),uniforms:et},rt={fragmentShader:"\n\nuniform vec3 color;\nuniform sampler2D map;\nuniform float opacity;\n\nuniform int fogType;\nuniform vec3 fogColor;\nuniform float fogDensity;\nuniform float fogNear;\nuniform float fogFar;\nuniform float alphaTest;\n\nvarying vec2 vUV;\n//DEFINEFRAGCOLOR\n\nvoid main() {\n\n    vec4 texture = texture2D(map, vUV);\n\n    if (texture.a <= alphaTest) discard;\n\n    gl_FragColor = vec4(color * texture.xyz, texture.a * opacity);\n\n    if (fogType > 0) {\n\n        float depth = gl_FragCoord.z / gl_FragCoord.w;\n        float fogFactor = 0.0;\n\n        if (fogType == 1) {\n            fogFactor = smoothstep(fogNear, fogFar, depth);\n        }\n\n        else {\n            const float LOG2 = 1.442695;\n            float fogFactor = exp2(- fogDensity * fogDensity * depth * depth * LOG2);\n            fogFactor = 1.0 - clamp(fogFactor, 0.0, 1.0);\n        }\n\n        gl_FragColor = mix(gl_FragColor, vec4(fogColor, gl_FragColor.w), fogFactor);\n\n    }\n}\n\n".replace("#define GLSLIFY 1",""),vertexShader:"\n\nuniform int useScreenCoordinates;\nuniform vec3 screenPosition;\nuniform mat4 modelViewMatrix;\nuniform mat4 projectionMatrix;\nuniform float rotation;\nuniform vec2 scale;\nuniform vec2 alignment;\nuniform vec2 uvOffset;\nuniform vec2 uvScale;\n\nattribute vec2 position;\nattribute vec2 uv;\n\nvarying vec2 vUV;\n\nvoid main() {\n\n    vUV = uvOffset + uv * uvScale;\n\n    vec2 alignedPosition = position + alignment;\n\n    vec2 rotatedPosition;\n    rotatedPosition.x = ( cos(rotation) * alignedPosition.x - sin(rotation) * alignedPosition.y ) * scale.x;\n    rotatedPosition.y = ( sin(rotation) * alignedPosition.x + cos(rotation) * alignedPosition.y ) * scale.y;\n\n    vec4 finalPosition;\n\n    if(useScreenCoordinates != 0) {\n        finalPosition = vec4(screenPosition.xy + rotatedPosition, screenPosition.z, 1.0);\n    }\n\n    else {\n        finalPosition = projectionMatrix * modelViewMatrix * vec4(0.0, 0.0, 0.0, 1.0); finalPosition /= finalPosition.w;\n        finalPosition.xy += rotatedPosition; \n    }\n\n    gl_Position = finalPosition;\n\n}\n\n".replace("#define GLSLIFY 1",""),uniforms:{}},st={opacity:{type:"f",value:1},fogColor:{type:"c",value:new b.Color(1,1,1)},fogNear:{type:"f",value:1},fogFar:{type:"f",value:2e3},directionalLightColor:{type:"fv",value:[]},directionalLightDirection:{type:"fv",value:[]}},nt="uniform float opacity;\nuniform mat4 projectionMatrix;\n\nuniform vec3 fogColor;\nuniform float fogNear;\nuniform float fogFar;\n\nvarying vec3 vLight;\nvarying vec3 vColor;\nvarying vec3 cposition;\nvarying vec3 p1;\nvarying vec3 p2;\nvarying float r;\n\n#ifdef SHADED\nuniform highp sampler2D shading;\n#endif\n\n//DEFINEFRAGCOLOR\n\n//cylinder-ray intersection testing taken from http://mrl.nyu.edu/~dzorin/cg05/lecture12.pdf\n//also useful: http://stackoverflow.com/questions/9595300/cylinder-impostor-in-glsl\n//with a bit more care (caps) this could be a general cylinder imposter (see also outline)\nvoid main() {\n    vec3 color = abs(vColor);\n    vec3 pos = cposition;\n    vec3 p = pos; //ray point\n    vec3 v = vec3(0.0,0.0,-1.0); //ray normal - orthographic\n    if(projectionMatrix[3][3] == 0.0) v = normalize(pos); //ray normal - perspective\n    vec3 pa = p1; //cyl start\n    vec3 va = normalize(p2-p1); //cyl norm\n    vec3 tmp1 = v-(dot(v,va)*va);\n    vec3 deltap = p-pa;\n    float A = dot(tmp1,tmp1);\n    if(A == 0.0) discard;\n    vec3 tmp2 = deltap-(dot(deltap,va)*va);\n    float B = 2.0*dot(tmp1, tmp2);\n    float C = dot(tmp2,tmp2)-r*r;\n//quadratic equation!\n    float det = (B*B) - (4.0*A*C);\n    if(det < 0.0) discard;\n    float sqrtDet = sqrt(det);\n    float posT = (-B+sqrtDet)/(2.0*A);\n    float negT = (-B-sqrtDet)/(2.0*A);\n    float intersectionT = min(posT,negT);\n    vec3 qi = p+v*intersectionT;\n    float dotp1 = dot(va,qi-p1);\n    float dotp2 = dot(va,qi-p2);\n    vec3 norm;\n    if( dotp1 < 0.0 || dotp2 > 0.0) { //(p-c)^2 + 2(p-c)vt +v^2+t^2 - r^2 = 0\n       vec3 cp;\n       if( dotp1 < 0.0) {  \n//        if(vColor.x < 0.0 ) discard; //color sign bit indicates if we should cap or not\n        cp = p1;\n       } else {\n//          if(vColor.y < 0.0 ) discard;\n          cp = p2;\n       }\n       vec3 diff = p-cp;\n       A = dot(v,v);\n       B = dot(diff,v)*2.0;\n       C = dot(diff,diff)-r*r;\n       det = (B*B) - (4.0*C);\n       if(det < 0.0) discard;\n       sqrtDet = sqrt(det);\n       posT = (-B+sqrtDet)/(2.0);\n       negT = (-B-sqrtDet)/(2.0);\n       float t = min(posT,negT);\n       qi = p+v*t; \n       norm = normalize(qi-cp); \n    } else {\n       norm = normalize(qi-(dotp1*va + p1));\n    }\n    vec4 clipPos = projectionMatrix * vec4(qi, 1.0);\n    float ndcDepth = clipPos.z / clipPos.w;\n    float depth = ((gl_DepthRange.diff * ndcDepth) + gl_DepthRange.near + gl_DepthRange.far) / 2.0;\n    gl_FragDepthEXT = depth;",at={fragmentShader:[nt,"    float dotProduct = dot( norm, vLight );\n    vec3 light = vec3( max( dotProduct, 0.0 ) );\n    color *= light;\n#ifdef SHADED\n    ivec2 dim = textureSize(shading,0);\n    float shadowFactor = texture2D(shading,vec2(gl_FragCoord.x/float(dim.x),gl_FragCoord.y/float(dim.y))).r;\n    color *= shadowFactor;\n#endif    \n    gl_FragColor = vec4(color, opacity*opacity );\n    float fogFactor = smoothstep( fogNear, fogFar, depth );\n    gl_FragColor = mix( gl_FragColor, vec4( fogColor, gl_FragColor.w ), fogFactor );\n}"].join("\n"),vertexShader:"\n\nuniform mat4 modelViewMatrix;\nuniform mat4 projectionMatrix;\nuniform mat4 viewMatrix;\nuniform mat3 normalMatrix;\nuniform vec3 directionalLightColor[ 1 ];\nuniform vec3 directionalLightDirection[ 1 ];\n\nattribute vec3 position;\nattribute vec3 normal;\nattribute vec3 color;\nattribute float radius;\n\nvarying vec3 vColor;\nvarying vec3 vLight;\nvarying vec3 cposition;\nvarying vec3 p1;\nvarying vec3 p2;\nvarying float r;\n\nvoid main() {\n\n    vColor = color; vColor.z = abs(vColor.z); //z indicates which vertex and so would vary\n    r = abs(radius);\n    vec4 to = modelViewMatrix*vec4(normal, 1.0); //normal is other point of cylinder\n    vec4 pt = modelViewMatrix*vec4(position, 1.0);\n    vec4 mvPosition = pt;\n    p1 = pt.xyz; p2 = to.xyz;\n    vec3 norm = to.xyz-pt.xyz;\n    float mult = 1.1; //slop to account for perspective of sphere\n    if(length(p1) > length(p2)) { //billboard at level of closest point\n       mvPosition = to;\n    }\n    vec3 n = normalize(mvPosition.xyz);\n    bool isperspective = (projectionMatrix[3][3] == 0.0);\n//intersect with the plane defined by the camera looking at the billboard point\n    if(color.z >= 0.0) { //p1\n       if(isperspective) { //perspective\n         vec3 pnorm = normalize(p1);\n         float t = dot(mvPosition.xyz-p1,n)/dot(pnorm,n);\n         mvPosition.xyz = p1+t*pnorm; \n       } else { //orthographic\n         mvPosition.xyz = p1;\n       }\n    } else {\n      if(isperspective) { //perspective\n         vec3 pnorm = normalize(p2);\n         float t = dot(mvPosition.xyz-p2,n)/dot(pnorm,n);\n         mvPosition.xyz = p2+t*pnorm;\n       } else { //orthographic\n         mvPosition.xyz = p2;\n       } \n       mult *= -1.0;\n    }\n\n    if(isperspective) { //perspective\n      vec3 cr = normalize(cross(mvPosition.xyz,norm))*radius;\n      vec3 doublecr = normalize(cross(mvPosition.xyz,cr))*radius;\n      mvPosition.xyz +=  mult*(cr + doublecr).xyz;\n    } else {\n      vec3 cr = normalize(cross(vec3(0.0,0.0,-1.0),norm))*radius;\n      vec3 doublecr = normalize(cross(vec3(0.0,0.0,-1.0),cr))*radius;\n      mvPosition.xyz +=  mult*(cr + doublecr).xyz;     \n    }\n\n    cposition = mvPosition.xyz;\n    gl_Position = projectionMatrix * mvPosition;\n    vec4 lDirection = viewMatrix * vec4( directionalLightDirection[ 0 ], 0.0 );\n    vLight = normalize( lDirection.xyz )*directionalLightColor[0]; //not really sure this is right, but color is always white so..\n}\n\n",uniforms:st},ot={fragmentShader:nt+"gl_FragColor = vec4(color,1.0);}",vertexShader:"\n\nuniform mat4 modelViewMatrix;\nuniform mat4 projectionMatrix;\nuniform mat4 viewMatrix;\nuniform mat3 normalMatrix;\nuniform vec3 directionalLightColor[ 1 ];\nuniform vec3 directionalLightDirection[ 1 ];\nuniform vec3 outlineColor;\nuniform float outlineWidth;\nuniform float outlinePushback;\nuniform float outlineMaxPixels;\nuniform float vWidth;\nuniform mat4 projinv;\n\n\nattribute vec3 position;\nattribute vec3 normal;\nattribute vec3 color;\nattribute float radius;\n\nvarying vec3 vColor;\nvarying vec3 vLight;\nvarying vec3 cposition;\nvarying vec3 p1;\nvarying vec3 p2;\nvarying float r;\n\nvoid main() {\n\n    vColor = outlineColor;\n    float rad = radius+sign(radius)*outlineWidth;\n    r = abs(rad);\n\n    vec4 to = modelViewMatrix*vec4(normal, 1.0); //normal is other point of cylinder\n    vec4 pt = modelViewMatrix*vec4(position, 1.0);\n//pushback\n    float scale = 1.0;\n    if(projectionMatrix[3][3] != 0.0) { //orthographic\n        to.z -= outlinePushback;\n        pt.z -= outlinePushback;\n    } else { //perspective\n        vec4 midbefore = pt;\n        if(length(to.xyz) < length(pt)) {\n            midbefore = to;\n        }\n        vec4 midafter = midbefore;\n        midafter.xyz += normalize(midbefore.xyz)*outlinePushback;\n\n        to.xyz += normalize(to.xyz)*outlinePushback;\n        pt.xyz += normalize(pt.xyz)*outlinePushback;\n\n        //figure out a scaling factor for radius to account for perspective setback\n        vec4 midbeforer = vec4(midbefore.x+rad,midbefore.y, midbefore.z, midbefore.w);\n        vec4 midafterr = vec4(midafter.x+rad,midafter.y, midafter.z, midafter.w);\n\n        vec4 mb = projectionMatrix*midbefore;\n        vec4 mbr = projectionMatrix*midbeforer;\n        vec4 ma = projectionMatrix*midafter;\n        vec4 mar = projectionMatrix*midafterr;\n        mb /= mb.w;\n        mbr /= mbr.w;\n        ma /= ma.w;\n        mar /= mar.w;\n        scale = abs((mbr.x-mb.x)/(mar.x-ma.x));\n        rad *= scale;\n        r = abs(rad);\n    }\n    vec4 mvPosition = pt;\n    p1 = pt.xyz; p2 = to.xyz;\n    vec3 norm = to.xyz-pt.xyz;\n    float mult = 1.1; //slop to account for perspective of sphere\n    if(length(p1) > length(p2)) { //billboard at level of closest point\n       mvPosition = to;\n    }\n\n    vec3 n = normalize(mvPosition.xyz);\n//intersect with the plane defined by the camera looking at the billboard point\n    if(color.z >= 0.0) { //p1\n       vec3 pnorm = normalize(p1);\n       float t = dot(mvPosition.xyz-p1,n)/dot(pnorm,n);\n       mvPosition.xyz = p1+t*pnorm;\n    } else {\n       vec3 pnorm = normalize(p2);\n       float t = dot(mvPosition.xyz-p2,n)/dot(pnorm,n);\n       mvPosition.xyz = p2+t*pnorm;\n       mult *= -1.0;\n    }\n\n    if(outlineMaxPixels > 0.0) {\n        vec4 cpos = mvPosition;\n        vec4 unadjusted = projectionMatrix*vec4(cpos.x+abs(scale*radius), cpos.y,cpos.z,cpos.w); \n        vec4 ccoord = projectionMatrix*cpos;\n        vec4 adjust = projectionMatrix*vec4(cpos.x+r,cpos.y,cpos.z,cpos.w); \n        unadjusted /= unadjusted.w;\n        adjust /= adjust.w;\n        unadjusted.xyz -= ccoord.xyz/ccoord.w;\n        adjust.xyz -= ccoord.xyz/ccoord.w;\n        float diff = abs(adjust.x-unadjusted.x);\n        diff *= vWidth; //this should now be in pixels\n        if(diff > outlineMaxPixels) {\n            float fixlen = abs(unadjusted.x) + outlineMaxPixels/vWidth; \n            vec4 pcoord = ccoord;\n            pcoord.x += fixlen*pcoord.w;\n            vec4 altc = projinv*pcoord;\n            r= abs(altc.x-cpos.x);\n        }\n    }\n\n    vec3 cr = normalize(cross(mvPosition.xyz,norm))*rad;\n    vec3 doublecr = normalize(cross(mvPosition.xyz,cr))*rad;\n    mvPosition.xy +=  mult*(cr + doublecr).xy;\n    cposition = mvPosition.xyz;\n    gl_Position = projectionMatrix * mvPosition;\n    vLight = vec3(1.0,1.0,1.0);\n}\n\n",uniforms:{opacity:{type:"f",value:1},fogColor:{type:"c",value:new b.Color(1,1,1)},fogNear:{type:"f",value:1},fogFar:{type:"f",value:2e3},outlineColor:{type:"c",value:new b.Color(0,0,0)},outlineWidth:{type:"f",value:.1},outlinePushback:{type:"f",value:1},outlineMaxPixels:{type:"f",value:0},projinv:{type:"mat4",value:[]}}},lt={opacity:{type:"f",value:1},fogColor:{type:"c",value:new b.Color(1,1,1)},fogNear:{type:"f",value:1},fogFar:{type:"f",value:2e3},data:{type:"i",value:3},colormap:{type:"i",value:4},depthmap:{type:"i",value:5},step:{type:"f",value:1},maxdepth:{type:"f",value:100},subsamples:{type:"f",value:5},textmat:{type:"mat4",value:[]},projinv:{type:"mat4",value:[]},transfermin:{type:"f",value:-.2},transfermax:{type:"f",value:.2}},ht={fragmentShader:"\nuniform highp sampler3D data;\nuniform highp sampler2D colormap;\nuniform highp sampler2D depthmap;\n\n\nuniform mat4 textmat;\nuniform mat4 projinv;\nuniform mat4 projectionMatrix;\n\nuniform float step;\nuniform float subsamples;\nuniform float maxdepth;\nuniform float transfermin;\nuniform float transfermax;\nin  vec4 mvPosition;\nout vec4 color;\nvoid main(void) {\n\n   vec4 pos = mvPosition;\n   bool seengood = false;\n   float i = 0.0;\n   color = vec4(1,1,1,0);\n   float increment = 1.0/subsamples;\n   float maxsteps = (maxdepth*subsamples/step);\n//there's probably a better way to do this..\n//calculate farthest possible point in model coordinates\n   vec4 maxpos = vec4(pos.x,pos.y,pos.z-maxdepth,1.0);\n// convert to projection\n   maxpos = projectionMatrix*maxpos;\n   vec4 startp = projectionMatrix*pos;\n// homogonize\n   maxpos /= maxpos.w;\n   startp /= startp.w;\n//take x,y from start and z from max\n   maxpos = vec4(startp.x,startp.y,maxpos.z,1.0);\n//convert back to model space\n   maxpos = projinv*maxpos;\n   maxpos /= maxpos.w;\n   float incr = step/subsamples;\n//get depth from depthmap\n//startp is apparently [-1,1]\n   vec2 tpos = startp.xy/2.0+0.5;\n   float depth = texture(depthmap, tpos).r;\n//compute vector between start and end\n   vec4 direction = maxpos-pos;\n   for( i = 0.0; i <= maxsteps; i++) {\n      vec4 pt = (pos+(i/maxsteps)*direction);\n      vec4 ppt = projectionMatrix*pt;\n      float ptdepth = ppt.z/ppt.w;\n      ptdepth = ((gl_DepthRange.diff * ptdepth) + gl_DepthRange.near + gl_DepthRange.far) / 2.0;\n      if(ptdepth > depth) break;\n      pt = textmat*pt;\n//       pt /= pt.w;\n      if(pt.x >= -0.01 && pt.y >= -0.01 && pt.z >= -0.01 && pt.x <= 1.01 && pt.y <= 1.01 && pt.z <= 1.01) {\n         seengood = true;\n      } else if(seengood) {\n         break;\n      }\n      if( pt.x < -0.01 || pt.x > 1.01 || pt.y < -0.01 || pt.y > 1.01 || pt.z < -0.01 || pt.z > 1.01  ){\n          color.a = 0.0;\n          continue;\n      }\n      else {\n         float val = texture(data, pt.zyx).r;\n         if(isinf(val)) continue; //masked out\n         float cval = (val-transfermin)/(transfermax-transfermin); //scale to texture 0-1 range\n         vec4 val_color = texture(colormap, vec2(cval,0.5));\n         color.rgb = color.rgb*color.a + (1.0-color.a)*val_color.a*val_color.rgb;\n         color.a += (1.0 - color.a) * val_color.a; \n         if(color.a > 0.0) color.rgb /= color.a;\n//          color = vec4(pt.x, pt.y, pt.z, 1.0);\n      }\n//       color = vec4(pt.x, pt.y, pt.z, 0.0)\n    }\n}\n\n        ".replace("#define GLSLIFY 1",""),vertexShader:"uniform mat4 modelViewMatrix;\nuniform mat4 projectionMatrix;\nuniform mat4 viewMatrix;\n\nin vec3 position;\nout vec4 mvPosition;\nvoid main() {\n\n    mvPosition = modelViewMatrix * vec4( position, 1.0 );\n    gl_Position = projectionMatrix*mvPosition;\n}\n".replace("#define GLSLIFY 1",""),uniforms:lt},ct={basic:V,blur:{fragmentShader:"const float INV_TOTAL_SAMPLES_FACTOR = 1.0 / 9.0;\nuniform highp sampler2D inTex;\nvarying highp vec2 vTexCoords;\n    \nvoid main() {\n \n vec2 texelSize = 1.0 / vec2(textureSize(inTex,0));\n float blurred_visibility_factor = 0.0f;\n for (int t = -1; t <= 1; ++t) {\n  for (int s = -1; s <= 1; ++s) {\n   vec2 offset = vec2(float(s), float(t)) * texelSize;\n   blurred_visibility_factor += texture2D(inTex, vTexCoords + offset).r;\n  }\n }\n    \n gl_FragDepthEXT = blurred_visibility_factor * INV_TOTAL_SAMPLES_FACTOR;\n \n}",vertexShader:"attribute vec2 vertexPosition;\nvarying highp vec2 vTexCoords;\nconst vec2 scale = vec2(0.5, 0.5);\n\nvoid main() {\n   vTexCoords  = vertexPosition * scale + scale; // scale vertex attribute to [0,1] range\n   gl_Position = vec4(vertexPosition, 0.0, 1.0);\n}\n        ",uniforms:{}},instanced:H,lambert:q,lambertdouble:Z,outline:K,screen:Q,screenaa:$,ssao:{fragmentShader:"uniform sampler2D depthmap;\nvarying highp vec2 vTexCoords;\nuniform float vWidth;\nuniform float vHeight;\nuniform float total_strength;\nuniform float radius;\nuniform mat4 projinv;\nuniform mat4 projectionMatrix;\n\n//code for pseudorandom vector from chatgpt\nfloat hash(vec3 p) {\n    p = fract(p * vec3(0.1031, 0.1030, 0.0973));\n    p += dot(p, p.yzx + 33.33);\n    return fract((p.x + p.y) * p.z);\n}\n\n// Generate a pseudorandom vec3 from a seed vec3\nvec3 pseudorandomVec3(vec3 seed) {\n    vec3 randomVec;\n    randomVec.x = hash(seed);\n    randomVec.y = hash(seed + vec3(1.0, 0.0, 17.1));\n    randomVec.z = hash(seed + vec3(0.0, 13.23, 0.0));\n    return randomVec;\n}\n\nvoid main(void)\n{   \n   const float base = 0.2;\n   const float area = 0.75;\n   const int cycles = 1;\n\n   const int samples = 64;\n   vec3 sample_sphere[64] = vec3[](\n      vec3(0.091258,-0.510164,0.000000),\n      vec3(-0.204347,-0.872967,0.187199),\n      vec3(0.009690,-0.263696,-0.110414),\n      vec3(0.175208,-0.563987,0.228527),\n      vec3(-0.001824,-0.003113,-0.000323),\n      vec3(0.411134,-0.719869,-0.261530),\n      vec3(-0.074272,-0.377368,0.276290),\n      vec3(-0.147773,-0.381587,-0.284529),\n      vec3(0.173317,-0.199635,0.063295),\n      vec3(-0.186452,-0.199460,0.076965),\n      vec3(0.143985,-0.308160,-0.307687),\n      vec3(0.053194,-0.148286,0.169589),\n      vec3(-0.547656,-0.486476,-0.317378),\n      vec3(0.020804,-0.015092,-0.004574),\n      vec3(-0.038006,-0.043165,0.054059),\n      vec3(-0.094795,-0.443908,-0.731525),\n      vec3(0.547552,-0.396466,0.461477),\n      vec3(-0.176886,-0.089989,0.007315),\n      vec3(0.074401,-0.048840,-0.074039),\n      vec3(-0.008240,-0.075697,0.178197),\n      vec3(-0.307880,-0.185053,-0.368943),\n      vec3(0.309520,-0.108483,0.041646),\n      vec3(-0.773478,-0.292946,0.538166),\n      vec3(0.184487,-0.231594,-0.820065),\n      vec3(0.207318,-0.100531,0.361797),\n      vec3(-0.173306,-0.037737,-0.055289),\n      vec3(0.548102,-0.105342,-0.253237),\n      vec3(-0.119342,-0.043907,0.285162),\n      vec3(-0.270247,-0.087861,-0.751357),\n      vec3(0.449312,-0.039777,0.236146),\n      vec3(-0.743773,-0.036095,0.196056),\n      vec3(0.148819,-0.004300,-0.231448),\n      vec3(0.008773,0.000809,0.051047),\n      vec3(-0.461467,0.027390,-0.357386),\n      vec3(0.169626,0.013338,-0.014053),\n      vec3(-0.043786,0.007095,0.047331),\n      vec3(0.004821,0.140371,-0.988260),\n      vec3(0.092402,0.023994,0.101860),\n      vec3(-0.295335,0.061530,-0.027372),\n      vec3(0.024903,0.007537,-0.018901),\n      vec3(-0.081463,0.125402,0.447794),\n      vec3(-0.397119,0.231805,-0.631062),\n      vec3(0.163853,0.059014,0.044905),\n      vec3(-0.495220,0.214357,0.254139),\n      vec3(0.306123,0.373687,-0.825715),\n      vec3(0.021665,0.026737,0.053220),\n      vec3(-0.208231,0.117129,-0.098685),\n      vec3(0.139749,0.080968,-0.043086),\n      vec3(-0.153599,0.182814,0.262090),\n      vec3(-0.159673,0.496777,-0.743568),\n      vec3(0.134797,0.117152,0.095753),\n      vec3(-0.155626,0.120533,0.019395),\n      vec3(0.042311,0.054462,-0.049709),\n      vec3(0.001257,0.031288,0.034468),\n      vec3(-0.002271,0.003199,-0.002304),\n      vec3(0.662104,0.717307,0.033854),\n      vec3(-0.373100,0.576021,0.308274),\n      vec3(0.024233,0.231316,-0.173688),\n      vec3(0.161311,0.420217,0.234273),\n      vec3(-0.045248,0.078031,-0.010411),\n      vec3(0.167453,0.376942,-0.094872),\n      vec3(-0.056194,0.433247,0.173218),\n      vec3(-0.016224,0.123149,-0.035569),\n      vec3(0.067127,0.407641,0.028479)\n   );\n\n   float depth = texture2D(depthmap, vTexCoords).r;\n   if(depth == 1.0) {\n      discard;\n   }\n\n   vec4 screen_position = vec4(vTexCoords, depth,1.0);\n   vec4 pos = projinv*screen_position;\n   pos /= pos.w;\n   vec3 position = pos.xyz;\n\n   //approximate the normal from the depth map; I find this simpler\n   //than trying to recompute the exact normal within every possible object shader\n\n   //pixel offset positions in screen space\n   ivec2 dim = textureSize(depthmap,0);\n   vec2 offset1 = vec2(0.0,1.0/float(dim.y));\n   vec2 offset2 = vec2(1.0/float(dim.x),0.0);\n   float depth1 = texture2D(depthmap, vTexCoords + offset1).r;\n   float depth2 = texture2D(depthmap, vTexCoords + offset2).r;\n   \n   vec3 p1 = vec3(screen_position.xy+offset1, depth1 - depth);\n   vec3 p2 = vec3(screen_position.xy+offset2, depth2 - depth);\n\n   //convert to model space\n   vec4 pos1 = projinv*vec4(p1,1);\n   pos1 /= pos1.w;\n   vec4 pos2 = projinv*vec4(p2,1);\n   pos2 /= pos2.w;\n\n   vec3 normal = normalize(cross(pos1.xyz-position, pos2.xyz-position)); //model normal, important we normalize in model space\n\n   //pseudo random number\n\n   float occlusion = 0.0;\n   for(int c = 0; c < cycles; c++) {\n   vec3 random = normalize(pseudorandomVec3(position+float(c)));\n   for(int i=0; i < samples; i++) {\n\n      vec3 ray = radius * reflect(sample_sphere[i],random);\n      vec3 hemi_ray = position + sign(dot(ray,normal)) * ray; //model space\n      vec4 hemi_screen = projectionMatrix*vec4(hemi_ray,1.0);\n      hemi_screen /= hemi_screen.w;\n      \n      float occ_depth = texture2D(depthmap, clamp(hemi_screen.xy,0.0,1.0)).r;\n      float difference = hemi_screen.z - occ_depth;\n      \n      occlusion += step(0.0, difference) * (1.0-smoothstep(0.0, area, difference));\n   }\n   }\n   float ao = 1.0 - total_strength * occlusion * (1.0 / float(cycles*samples));\n   gl_FragDepthEXT = clamp(ao+base,0.0,1.0);\n\n}",vertexShader:"attribute vec2 vertexPosition;\nvarying highp vec2 vTexCoords;\nconst vec2 scale = vec2(0.5, 0.5);\n\nvoid main() {\n   vTexCoords  = vertexPosition * scale + scale; // scale vertex attribute to [0,1] range\n   gl_Position = vec4(vertexPosition, 0.0, 1.0);\n}\n        ",uniforms:{total_strength:{type:"f",value:1},radius:{type:"f",value:5},projinv:{type:"mat4",value:[]}}},sphereimposter:tt,sphereimposteroutline:it,sprite:rt,stickimposter:at,stickimposteroutline:ot,volumetric:ht};function dt(t){let e={};for(const r in t){e[r]={},e[r].type=t[r].type;var i=t[r].value;i instanceof b.Color?e[r].value=i.clone():"number"==typeof i?e[r].value=i:i instanceof Array?e[r].value=[]:console.error("Error copying shader uniforms from ShaderLib: unknown type for uniform")}return e}const ut={clone:dt};class Camera extends Object3D{constructor(t=50,e=1,i=.1,r=2e3,s=!1){super(),this.projectionMatrix=new l.Matrix4,this.projectionMatrixInverse=new l.Matrix4,this.matrixWorldInverse=new l.Matrix4,this.fov=t,this.aspect=e,this.near=i,this.far=r;var n=this.position.z;this.right=n*Math.tan(Math.PI/180*t),this.left=-this.right,this.top=this.right/this.aspect,this.bottom=-this.top,this.ortho=!!s,this.updateProjectionMatrix()}lookAt(t){this.matrix.lookAt(this.position,t,this.up),this.rotationAutoUpdate&&(!1===this.useQuaternion&&this.rotation instanceof l.Vector3?this.rotation.setEulerFromRotationMatrix(this.matrix,this.eulerOrder):console.error("Unimplemented math operation."))}updateProjectionMatrix(){this.ortho?this.projectionMatrix.makeOrthographic(this.left,this.right,this.top,this.bottom,this.near,this.far):this.projectionMatrix.makePerspective(this.fov,this.aspect,this.near,this.far),this.projectionMatrixInverse.getInverse(this.projectionMatrix)}}class Fog{constructor(t,e=1,i=1e3){this.name="",this.color=new b.Color(t),this.near=e,this.far=i}clone(){return new Fog(this.color.getHex(),this.near,this.far)}}class SpritePlugin{constructor(){this.sprite={vertices:null,faces:null,vertexBuffer:null,elementBuffer:null,program:null,attributes:{},uniforms:null}}init(t){this.gl=t.context,this.renderer=t,this.precision=t.getPrecision(),this.sprite.vertices=new Float32Array(16),this.sprite.faces=new Uint16Array(6);var e=0;this.sprite.vertices[e++]=-1,this.sprite.vertices[e++]=-1,this.sprite.vertices[e++]=0,this.sprite.vertices[e++]=0,this.sprite.vertices[e++]=1,this.sprite.vertices[e++]=-1,this.sprite.vertices[e++]=1,this.sprite.vertices[e++]=0,this.sprite.vertices[e++]=1,this.sprite.vertices[e++]=1,this.sprite.vertices[e++]=1,this.sprite.vertices[e++]=1,this.sprite.vertices[e++]=-1,this.sprite.vertices[e++]=1,this.sprite.vertices[e++]=0,this.sprite.vertices[e++]=1,e=0,this.sprite.faces[e++]=0,this.sprite.faces[e++]=1,this.sprite.faces[e++]=2,this.sprite.faces[e++]=0,this.sprite.faces[e++]=2,this.sprite.faces[e++]=3,this.sprite.vertexBuffer=this.gl.createBuffer(),this.sprite.elementBuffer=this.gl.createBuffer(),this.gl.bindBuffer(this.gl.ARRAY_BUFFER,this.sprite.vertexBuffer),this.gl.bufferData(this.gl.ARRAY_BUFFER,this.sprite.vertices,this.gl.STATIC_DRAW),this.gl.bindBuffer(this.gl.ELEMENT_ARRAY_BUFFER,this.sprite.elementBuffer),this.gl.bufferData(this.gl.ELEMENT_ARRAY_BUFFER,this.sprite.faces,this.gl.STATIC_DRAW),this.sprite.program=this.createProgram(ct.sprite,this.precision||1),this.sprite.attributes={};const i={};this.sprite.attributes.position=this.gl.getAttribLocation(this.sprite.program,"position"),this.sprite.attributes.uv=this.gl.getAttribLocation(this.sprite.program,"uv"),i.uvOffset=this.gl.getUniformLocation(this.sprite.program,"uvOffset"),i.uvScale=this.gl.getUniformLocation(this.sprite.program,"uvScale"),i.rotation=this.gl.getUniformLocation(this.sprite.program,"rotation"),i.scale=this.gl.getUniformLocation(this.sprite.program,"scale"),i.alignment=this.gl.getUniformLocation(this.sprite.program,"alignment"),i.color=this.gl.getUniformLocation(this.sprite.program,"color"),i.map=this.gl.getUniformLocation(this.sprite.program,"map"),i.opacity=this.gl.getUniformLocation(this.sprite.program,"opacity"),i.useScreenCoordinates=this.gl.getUniformLocation(this.sprite.program,"useScreenCoordinates"),i.screenPosition=this.gl.getUniformLocation(this.sprite.program,"screenPosition"),i.modelViewMatrix=this.gl.getUniformLocation(this.sprite.program,"modelViewMatrix"),i.projectionMatrix=this.gl.getUniformLocation(this.sprite.program,"projectionMatrix"),i.fogType=this.gl.getUniformLocation(this.sprite.program,"fogType"),i.fogDensity=this.gl.getUniformLocation(this.sprite.program,"fogDensity"),i.fogNear=this.gl.getUniformLocation(this.sprite.program,"fogNear"),i.fogFar=this.gl.getUniformLocation(this.sprite.program,"fogFar"),i.fogColor=this.gl.getUniformLocation(this.sprite.program,"fogColor"),i.alphaTest=this.gl.getUniformLocation(this.sprite.program,"alphaTest"),this.sprite.uniforms=i}render(t,e,i,r,s){var n,a,o,l,h,c,d,u,f,p;if(!this.gl)throw new Error("WebGLRenderer not initialized");let g=[];null===(n=null==t?void 0:t.__webglSprites)||void 0===n||n.forEach((t=>{(s&&0==t.material.depthTest||!s&&t.material.depthTest)&&g.push(t)}));let m=g.length;if(!m)return;const v=this.sprite.attributes,_=this.sprite.uniforms;if(!_)throw new Error("Uniforms not defined");var y=.5*i,b=.5*r;this.gl.useProgram(this.sprite.program),this.gl.enableVertexAttribArray(v.position),this.gl.enableVertexAttribArray(v.uv),this.gl.disable(this.gl.CULL_FACE),this.gl.enable(this.gl.BLEND),this.gl.bindBuffer(this.gl.ARRAY_BUFFER,this.sprite.vertexBuffer),this.gl.vertexAttribPointer(v.position,2,this.gl.FLOAT,!1,16,0),this.gl.vertexAttribPointer(v.uv,2,this.gl.FLOAT,!1,16,8),this.gl.bindBuffer(this.gl.ELEMENT_ARRAY_BUFFER,this.sprite.elementBuffer),this.gl.uniformMatrix4fv(_.projectionMatrix,!1,e.projectionMatrix.elements),this.gl.activeTexture(this.gl.TEXTURE0),this.gl.uniform1i(_.map,0);var x,w=0,A=0,C=t.fog;let S,M,z,T;C?(this.gl.uniform3f(_.fogColor,C.color.r,C.color.g,C.color.b),this.gl.uniform1f(_.fogNear,C.near),this.gl.uniform1f(_.fogFar,C.far),this.gl.uniform1i(_.fogType,1),w=1,A=1):(this.gl.uniform1i(_.fogType,0),w=0,A=0);let E=[];for(x=0;x<m;x++)S=g[x],M=S.material,M&&(0!=M.depthTest||s)&&S.visible&&0!==M.opacity&&(M.useScreenCoordinates?S.z=-S.position.z:(S._modelViewMatrix.multiplyMatrices(e.matrixWorldInverse,S.matrixWorld),S.z=-S._modelViewMatrix.elements[14]));for(g.sort(ft),x=0;x<m;x++)if(S=g[x],M=S.material,M&&S.visible&&0!==M.opacity&&M.map&&M.map.image&&M.map.image.width){this.gl.uniform1f((null==_?void 0:_.alphaTest)||null,M.alphaTest);var L=M.map.image.width,F=M.map.image.height;E[0]=L*this.renderer.devicePixelRatio/i,E[1]=F*this.renderer.devicePixelRatio/r,!0===M.useScreenCoordinates?(this.gl.uniform1i(_.useScreenCoordinates,1),this.gl.uniform3f(_.screenPosition,(S.position.x*this.renderer.devicePixelRatio-y)/y,(b-S.position.y*this.renderer.devicePixelRatio)/b,Math.max(0,Math.min(1,S.position.z)))):(this.gl.uniform1i(_.useScreenCoordinates,0),this.gl.uniformMatrix4fv(_.modelViewMatrix,!1,S._modelViewMatrix.elements)),T=t.fog&&M.fog?A:0,w!==T&&(this.gl.uniform1i(_.fogType,T),w=T),z=1/(M.scaleByViewport?r:1),E[0]*=z*S.scale.x,E[1]*=z*S.scale.y;let e=null===(a=null==M?void 0:M.alignment)||void 0===a?void 0:a.x,s=null===(o=null==M?void 0:M.alignment)||void 0===o?void 0:o.y;M.screenOffset&&(e=(e||0)+2*M.screenOffset.x/L,s=(s||0)+2*M.screenOffset.y/F),this.gl.uniform2f(_.uvScale,(null===(l=null==M?void 0:M.uvScale)||void 0===l?void 0:l.x)||1,(null===(h=null==M?void 0:M.uvScale)||void 0===h?void 0:h.y)||1),this.gl.uniform2f(_.uvOffset,(null===(c=null==M?void 0:M.uvOffset)||void 0===c?void 0:c.x)||0,(null===(d=null==M?void 0:M.uvOffset)||void 0===d?void 0:d.y)||0),this.gl.uniform2f(_.alignment,e||0,s||0),this.gl.uniform1f(_.opacity,M.opacity),this.gl.uniform3f(_.color,(null===(u=null==M?void 0:M.color)||void 0===u?void 0:u.r)||0,(null===(f=null==M?void 0:M.color)||void 0===f?void 0:f.g)||0,(null===(p=null==M?void 0:M.color)||void 0===p?void 0:p.b)||0),this.gl.uniform1f(_.rotation,S.rotation),this.gl.uniform2fv(_.scale,E),this.renderer.setDepthTest(M.depthTest),this.renderer.setDepthWrite(M.depthWrite),this.renderer.setTexture(M.map,0),this.gl.drawElements(this.gl.TRIANGLES,6,this.gl.UNSIGNED_SHORT,0)}this.gl.enable(this.gl.CULL_FACE)}createProgram(t,e){if(!this.gl)throw new Error("WebGL Rendering context not found");var i=this.gl.createProgram();if(!i)throw new Error("Error creating webgl program");var r=this.gl.createShader(this.gl.FRAGMENT_SHADER),s=this.gl.createShader(this.gl.VERTEX_SHADER);if(!r)throw new Error("Unable to create fragment shader SpritePlugin.createProgram");if(!s)throw new Error("Unable to create vertex shader SpritePlugin.createProgram");var n="precision "+e+" float;\n";if(this.gl.shaderSource(r,n+t.fragmentShader),this.gl.shaderSource(s,n+t.vertexShader),this.gl.compileShader(r),this.gl.compileShader(s),!this.gl.getShaderParameter(r,this.gl.COMPILE_STATUS)||!this.gl.getShaderParameter(s,this.gl.COMPILE_STATUS))throw new Error(`Error compiling shader: \n      ${this.gl.getShaderInfoLog(r)} \n      ${this.gl.getShaderInfoLog(s)}`);return this.gl.attachShader(i,r),this.gl.attachShader(i,s),this.gl.linkProgram(i),this.gl.getProgramParameter(i,this.gl.LINK_STATUS)||console.error("Could not initialize shader"),i}}function ft(t,e){return t.z!==e.z?e.z-t.z:e.id-t.id}var pt=null,gt=null;class Renderer{constructor(t){this.context=null,this.devicePixelRatio=1,this.sortObjects=!0,this.autoUpdateObjects=!0,this.autoUpdateScene=!0,this.info={memory:{programs:0,geometries:0,textures:0},render:{calls:0,vertices:0,faces:0,points:0}},this._offscreen=null,this._bitmap=null,this._programs=[],this._programs_counter=0,this._webglversion=1,this._currentProgram=null,this._currentMaterialId=-1,this._currentGeometryGroupHash=null,this._currentCamera=null,this._geometryGroupCounter=0,this._oldDoubleSided=-1,this._oldFlipSided=-1,this._oldDepthTest=-1,this._oldDepthWrite=-1,this._oldPolygonOffset=null,this._oldLineWidth=null,this._viewportWidth=0,this._viewportHeight=0,this._currentWidth=0,this._currentHeight=0,this._enabledAttributes={},this._vector3=new l.Vector3,this._worldInverse=new l.Matrix4,this._projInverse=new l.Matrix4,this._textureMatrix=new l.Matrix4,this._fullProjModelMatrix=new l.Matrix4,this._fullProjModelMatrixInv=new l.Matrix4,this._direction=new l.Vector3,this._lightsNeedUpdate=!0,this._lights={ambient:[0,0,0],directional:{length:0,colors:[],positions:[]},point:{length:0,colors:[],positions:[],distances:[]},spot:{length:0,colors:[],positions:[],distances:[],directions:[],anglesCos:[],exponents:[]},hemi:{length:0,skyColors:[],groundColors:[],positions:[]}},this.sprites=new SpritePlugin,this._screenshader=null,this._AOshader=null,this._blurshader=null,this._vertexattribpos=null,this._aovertexattribpos=null,this._blurvertexattribpos=null,this._screenQuadVBO=null,this._fb=null,this._targetTexture=null,this._depthTexture=null,this._shadingTexture=null,this._scratchTexture=null,this._AOstrength=1,this._AOradius=5,this.SHADE_TEXTURE=3,t=t||{},this.row=t.row,this.col=t.col,this.rows=t.rows,this.cols=t.cols,this._canvas=void 0!==t.canvas?t.canvas:document.createElement("canvas"),this._precision=void 0!==t.precision?t.precision:"highp",this._alpha=void 0===t.alpha||t.alpha,this._premultipliedAlpha=void 0===t.premultipliedAlpha||t.premultipliedAlpha,this._antialias=void 0!==t.antialias&&t.antialias,this._preserveDrawingBuffer=void 0!==t.preserveDrawingBuffer&&t.preserveDrawingBuffer,this._clearColor=void 0!==t.clearColor?new b.Color(t.clearColor):new b.Color(0),this._clearAlpha=void 0!==t.clearAlpha?t.clearAlpha:0,this._outlineMaterial=new MeshOutlineMaterial(t.outline),this._outlineSphereImposterMaterial=new SphereImposterOutlineMaterial(t.outline),this._outlineStickImposterMaterial=new StickImposterOutlineMaterial(t.outline),this._outlineEnabled=!!t.outline,this._AOEnabled=!!t.ambientOcclusion,t.ambientOcclusion&&void 0!==t.ambientOcclusion.strength&&(this._AOstrength=parseFloat(t.ambientOcclusion.strength)),0==this._AOstrength&&(this._AOEnabled=!1),t.ambientOcclusion&&void 0!==t.ambientOcclusion.radius&&(this._AOradius=parseFloat(t.ambientOcclusion.radius)),this.domElement=this._canvas,this._canvas.id=t.id,0!=t.containerWidth&&0!=t.containerHeight&&(this.initGL(),this.setDefaultGLState(),this.context=this._gl,this.isWebGL1()?this._extInstanced=this._gl.getExtension("ANGLE_instanced_arrays"):this._extInstanced={vertexAttribDivisorANGLE:this._gl.vertexAttribDivisor.bind(this._gl),drawElementsInstancedANGLE:this._gl.drawElementsInstanced.bind(this._gl)},this._extFragDepth=this._gl.getExtension("EXT_frag_depth"),this._extFloatLinear=this._gl.getExtension("OES_texture_float_linear"),this._extColorBufferFloat=this._gl.getExtension("EXT_color_buffer_float"),this.sprites.init(this))}supportedExtensions(){return{supportsAIA:Boolean(this._extInstanced),supportsImposters:Boolean(this._extFragDepth)||!this.isWebGL1(),regen:!1}}getContext(){return this._gl}getCanvas(){return this._canvas}isLost(){return null==this._gl||this._gl.isContextLost()}getPrecision(){return this._precision}setClearColorHex(t,e){this._clearColor.setHex(t),this._clearAlpha=e,this.isLost()||this._gl.clearColor(this._clearColor.r,this._clearColor.g,this._clearColor.b,this._clearAlpha)}enableOutline(t){this._outlineMaterial=new MeshOutlineMaterial(t),this._outlineSphereImposterMaterial=new SphereImposterOutlineMaterial(t),this._outlineStickImposterMaterial=new StickImposterOutlineMaterial(t),this._outlineEnabled=!0}disableOutline(){this._outlineEnabled=!1}enableAmbientOcclusion(t){t&&(t.strength&&(this._AOstrength=t.strength),t.scale&&(this._AOradius=t.scale)),this._AOEnabled=this._AOstrength>0}disableAmbientOcclusion(){this._AOEnabled=!1}setViewport(){if(this._offscreen&&(this._offscreen.width=this._canvas.width,this._offscreen.height=this._canvas.height),null!=this.rows&&null!=this.cols&&null!=this.row&&null!=this.col){var t=this._canvas.width/this.cols,e=this._canvas.height/this.rows;this._viewportWidth=t,this._viewportHeight=e,this.isLost()||(this._gl.enable(this._gl.SCISSOR_TEST),this._gl.scissor(t*this.col,e*this.row,t,e),this._gl.viewport(t*this.col,e*this.row,t,e))}}setSize(t,e){if(this.devicePixelRatio=void 0!==window.devicePixelRatio?window.devicePixelRatio:1,this._antialias&&this.devicePixelRatio<2&&(this.devicePixelRatio*=2),null!=this.rows&&null!=this.cols&&null!=this.row&&null!=this.col){var i=t/this.cols,r=e/this.rows;this._canvas.width=t*this.devicePixelRatio,this._canvas.height=e*this.devicePixelRatio,this._viewportWidth=i*this.devicePixelRatio,this._viewportHeight=r*this.devicePixelRatio,this._canvas.style.width=t+"px",this._canvas.style.height=e+"px",this.setViewport()}else this._viewportWidth=this._canvas.width=t*this.devicePixelRatio,this._viewportHeight=this._canvas.height=e*this.devicePixelRatio,this._canvas.style.width=t+"px",this._canvas.style.height=e+"px",this.isLost()||this._gl.viewport(0,0,this._gl.drawingBufferWidth,this._gl.drawingBufferHeight);this.initFrameBuffer()}clear(t,e,i){var r=0;(void 0===t||t)&&(r|=this._gl.COLOR_BUFFER_BIT),(void 0===e||e)&&(r|=this._gl.DEPTH_BUFFER_BIT),(void 0===i||i)&&(r|=this._gl.STENCIL_BUFFER_BIT),this._gl.clear(r)}setMaterialFaces(t,e){var i=t.side===a,r=t.side===n;t.imposter||(r=e?!r:r),this._oldDoubleSided!==i&&(i?this._gl.disable(this._gl.CULL_FACE):this._gl.enable(this._gl.CULL_FACE),this._oldDoubleSided=i),this._oldFlipSided!==r&&(r?this._gl.frontFace(this._gl.CW):this._gl.frontFace(this._gl.CCW),this._oldFlipSided=r),this._gl.cullFace(this._gl.BACK)}setDepthTest(t){this._oldDepthTest!==t&&(t?this._gl.enable(this._gl.DEPTH_TEST):this._gl.disable(this._gl.DEPTH_TEST),this._oldDepthTest=t)}setDepthWrite(t){this._oldDepthWrite!==t&&(this._gl.depthMask(t),this._oldDepthWrite=t)}setBlending(t){t?(this._gl.enable(this._gl.BLEND),this._gl.blendEquationSeparate(this._gl.FUNC_ADD,this._gl.FUNC_ADD),this._gl.blendFuncSeparate(this._gl.SRC_ALPHA,this._gl.ONE_MINUS_SRC_ALPHA,this._gl.ONE,this._gl.ONE_MINUS_SRC_ALPHA)):this._gl.disable(this._gl.BLEND)}initMaterial(t,e,i,r){var s,n;if(t.addEventListener("dispose",this.onMaterialDispose.bind(this)),n=t.shaderID){var a=ct[n];t.vertexShader=a.vertexShader,t.fragmentShader=a.fragmentShader,t.uniforms=ut.clone(a.uniforms),t.shaded&&t.makeShaded(this.SHADE_TEXTURE)}s={wireframe:t.wireframe,fragdepth:t.imposter,volumetric:t.volumetric,shaded:t.shaded},t.program=this.buildProgram(t.fragmentShader,t.vertexShader,t.uniforms,s)}renderBuffer(t,e,i,r,s,n){var a,o;if(r.visible&&(a=this.setProgram(t,e,i,r,n,this))){o=a.attributes;var l,h,c=!1,d=r.wireframe?1:0,u=16777215*s.id+2*a.id+d;if(u!==this._currentGeometryGroupHash&&(this._currentGeometryGroupHash=u,c=!0),c&&(this.disableAttributes(),o.position>=0&&(this._gl.bindBuffer(this._gl.ARRAY_BUFFER,s.__webglVertexBuffer),this.enableAttribute(o.position),this._gl.vertexAttribPointer(o.position,3,this._gl.FLOAT,!1,0,0)),o.color>=0&&(this._gl.bindBuffer(this._gl.ARRAY_BUFFER,s.__webglColorBuffer),this.enableAttribute(o.color),this._gl.vertexAttribPointer(o.color,3,this._gl.FLOAT,!1,0,0)),o.normal>=0&&(this._gl.bindBuffer(this._gl.ARRAY_BUFFER,s.__webglNormalBuffer),this.enableAttribute(o.normal),this._gl.vertexAttribPointer(o.normal,3,this._gl.FLOAT,!1,0,0)),o.offset>=0&&(this._gl.bindBuffer(this._gl.ARRAY_BUFFER,s.__webglOffsetBuffer),this.enableAttribute(o.offset),this._gl.vertexAttribPointer(o.offset,3,this._gl.FLOAT,!1,0,0)),o.radius>=0&&(this._gl.bindBuffer(this._gl.ARRAY_BUFFER,s.__webglRadiusBuffer),this.enableAttribute(o.radius),this._gl.vertexAttribPointer(o.radius,1,this._gl.FLOAT,!1,0,0))),n instanceof Mesh){if("instanced"===r.shaderID){var f=r.sphere.geometryGroups[0];c&&(this._gl.bindBuffer(this._gl.ARRAY_BUFFER,s.__webglVertexBuffer),this._gl.bufferData(this._gl.ARRAY_BUFFER,f.vertexArray,this._gl.STATIC_DRAW),this._gl.bindBuffer(this._gl.ARRAY_BUFFER,s.__webglNormalBuffer),this._gl.bufferData(this._gl.ARRAY_BUFFER,f.normalArray,this._gl.STATIC_DRAW),this._gl.bindBuffer(this._gl.ELEMENT_ARRAY_BUFFER,s.__webglFaceBuffer),this._gl.bufferData(this._gl.ELEMENT_ARRAY_BUFFER,f.faceArray,this._gl.STATIC_DRAW)),l=f.faceidx,this._extInstanced.vertexAttribDivisorANGLE(o.offset,1),this._extInstanced.vertexAttribDivisorANGLE(o.radius,1),this._extInstanced.vertexAttribDivisorANGLE(o.color,1),this._extInstanced.drawElementsInstancedANGLE(this._gl.TRIANGLES,l,this._gl.UNSIGNED_SHORT,0,s.radiusArray.length),this._extInstanced.vertexAttribDivisorANGLE(o.offset,0),this._extInstanced.vertexAttribDivisorANGLE(o.radius,0),this._extInstanced.vertexAttribDivisorANGLE(o.color,0)}else r.wireframe?(h=s.lineidx,this.setLineWidth(r.wireframeLinewidth),c&&this._gl.bindBuffer(this._gl.ELEMENT_ARRAY_BUFFER,s.__webglLineBuffer),this._gl.drawElements(this._gl.LINES,h,this._gl.UNSIGNED_SHORT,0)):(l=s.faceidx,c&&this._gl.bindBuffer(this._gl.ELEMENT_ARRAY_BUFFER,s.__webglFaceBuffer),this._gl.drawElements(this._gl.TRIANGLES,l,this._gl.UNSIGNED_SHORT,0));this.info.render.calls++,this.info.render.vertices+=l,this.info.render.faces+=l/3}else n instanceof Line&&(h=s.vertices,this.setLineWidth(r.linewidth),this._gl.drawArrays(this._gl.LINES,0,h),this.info.render.calls++)}}clearShading(){this._gl.framebufferTexture2D(this._gl.FRAMEBUFFER,this._gl.DEPTH_ATTACHMENT,this._gl.TEXTURE_2D,this._shadingTexture,0),this.clear(!1,!0,!1),this._gl.framebufferTexture2D(this._gl.FRAMEBUFFER,this._gl.DEPTH_ATTACHMENT,this._gl.TEXTURE_2D,this._depthTexture,0)}setShading(t,e,i){let r=t.__lights,s=t.fog,n=[];for(let e=0,r=t.__webglObjects.length;e<r;e++){let r=t.__webglObjects[e];r.render&&r[i]&&n.push(r)}if(0==n.length)return;this._gl.framebufferTexture2D(this._gl.FRAMEBUFFER,this._gl.DEPTH_ATTACHMENT,this._gl.TEXTURE_2D,this._shadingTexture,0),this._gl.framebufferTexture2D(this._gl.FRAMEBUFFER,this._gl.COLOR_ATTACHMENT0,this._gl.TEXTURE_2D,null,0),this.renderObjects(t.__webglObjects,!0,i+"Depth",e,r,s,!1),this._gl.framebufferTexture2D(this._gl.FRAMEBUFFER,this._gl.DEPTH_ATTACHMENT,this._gl.TEXTURE_2D,this._scratchTexture,0),this.clear(!1,!0,!1),this._gl.useProgram(this._AOshader),this._currentProgram=this._AOshader,this.setDepthTest(-1),this.setDepthWrite(-1);let a=this._AOshader.uniforms;this._gl.uniform1f(a.total_strength,this._AOstrength),this._gl.uniform1f(a.radius,this._AOradius),this._fullProjModelMatrix=new l.Matrix4,this._fullProjModelMatrixInv=new l.Matrix4;let o=n[0].object;this._fullProjModelMatrix.multiplyMatrices(e.projectionMatrix,o._modelViewMatrix),this._fullProjModelMatrixInv.getInverse(this._fullProjModelMatrix),this._gl.uniformMatrix4fv(a.projectionMatrix,!1,this._fullProjModelMatrix.elements),this._gl.uniformMatrix4fv(a.projinv,!1,this._fullProjModelMatrixInv.elements),this._gl.bindBuffer(this._gl.ARRAY_BUFFER,this._screenQuadVBO),this._gl.enableVertexAttribArray(this._aovertexattribpos),this._gl.vertexAttribPointer(this._aovertexattribpos,2,this._gl.FLOAT,!1,0,0),this._gl.activeTexture(this._gl.TEXTURE0),this._gl.bindTexture(this._gl.TEXTURE_2D,this._shadingTexture),this._gl.drawArrays(this._gl.TRIANGLES,0,6),this._gl.framebufferTexture2D(this._gl.FRAMEBUFFER,this._gl.DEPTH_ATTACHMENT,this._gl.TEXTURE_2D,this._shadingTexture,0),this.clear(!1,!0,!1),this._gl.useProgram(this._blurshader),this._currentProgram=this._blurshader,this.setDepthTest(-1),this.setDepthWrite(-1),this._gl.bindBuffer(this._gl.ARRAY_BUFFER,this._screenQuadVBO),this._gl.enableVertexAttribArray(this._blurvertexattribpos),this._gl.vertexAttribPointer(this._blurvertexattribpos,2,this._gl.FLOAT,!1,0,0),this._gl.activeTexture(this._gl.TEXTURE0),this._gl.bindTexture(this._gl.TEXTURE_2D,this._scratchTexture),this._gl.drawArrays(this._gl.TRIANGLES,0,6),this._gl.framebufferTexture2D(this._gl.FRAMEBUFFER,this._gl.COLOR_ATTACHMENT0,this._gl.TEXTURE_2D,this._targetTexture,0),this._gl.framebufferTexture2D(this._gl.FRAMEBUFFER,this._gl.DEPTH_ATTACHMENT,this._gl.TEXTURE_2D,this._depthTexture,0)}render(t,e){if(e instanceof Camera==!1)return void console.error("Renderer.render: camera is not an instance of Camera.");var i,r,s,n,a,o=t.__lights,l=t.fog;if(this._currentMaterialId=-1,this._lightsNeedUpdate=!0,this.autoUpdateScene&&t.updateMatrixWorld(),void 0===e.parent&&e.updateMatrixWorld(),e.matrixWorldInverse.getInverse(e.matrixWorld),this.isLost())return;this.autoUpdateObjects&&this.initWebGLObjects(t),this.info.render.calls=0,this.info.render.vertices=0,this.info.render.faces=0,this.info.render.points=0,this._currentWidth=this._viewportWidth,this._currentHeight=this._viewportHeight,this.setViewport(),this.setFrameBuffer(),this._gl.clearColor(this._clearColor.r,this._clearColor.g,this._clearColor.b,this._clearAlpha),this.clear(!0,!0,!0),a=t.__webglObjects;let h=!1,c=this._AOEnabled;for(i=0,r=a.length;i<r;i++)n=(s=a[i]).object,s.render=!1,n.visible&&(this.setupMatrices(n,e),this.unrollBufferMaterial(s),s.render=!0,s.volumetric&&(h=!0),s.hasAO&&(c=!0));if(this.setBlending(!1),c&&this.setShading(t,e,"opaque"),this.renderObjects(t.__webglObjects,!0,"opaque",e,o,l,!1),c&&this.clearShading(),this.renderSprites(t,e,!1),this.renderObjects(t.__webglObjects,!0,"transparentDepth",e,o,l,!0),this.renderObjects(t.__webglObjects,!1,"transparent",e,o,l,!0),h&&this._fb&&(this._gl.framebufferTexture2D(this._gl.FRAMEBUFFER,this._gl.DEPTH_ATTACHMENT,this._gl.TEXTURE_2D,null,0),this.renderObjects(t.__webglObjects,!1,"volumetric",e,o,l,!0)),this.renderFrameBuffertoScreen(),this.setDepthTest(!0),this.setDepthWrite(!0),this.renderSprites(t,e,!0),this._bitmap){const t=this._offscreen.transferToImageBitmap();this._bitmap.transferFromImageBitmap(t),t.close()}}setFrameBuffer(){if(this.isWebGL1()||!this._fb)return;let t=this._viewportWidth,e=this._viewportHeight;this._gl.enable(this._gl.SCISSOR_TEST),this._gl.scissor(0,0,t,e),this._gl.viewport(0,0,t,e),this._gl.bindTexture(this._gl.TEXTURE_2D,this._targetTexture),this._gl.texImage2D(this._gl.TEXTURE_2D,0,this._gl.RGBA,t,e,0,this._gl.RGBA,this._gl.UNSIGNED_BYTE,null),this._gl.texParameteri(this._gl.TEXTURE_2D,this._gl.TEXTURE_MIN_FILTER,this._gl.LINEAR),this._gl.texParameteri(this._gl.TEXTURE_2D,this._gl.TEXTURE_MAG_FILTER,this._gl.LINEAR),this._gl.texParameteri(this._gl.TEXTURE_2D,this._gl.TEXTURE_WRAP_S,this._gl.CLAMP_TO_EDGE),this._gl.texParameteri(this._gl.TEXTURE_2D,this._gl.TEXTURE_WRAP_T,this._gl.CLAMP_TO_EDGE),this._gl.bindTexture(this._gl.TEXTURE_2D,this._depthTexture),this._gl.texImage2D(this._gl.TEXTURE_2D,0,this._gl.DEPTH_COMPONENT32F,t,e,0,this._gl.DEPTH_COMPONENT,this._gl.FLOAT,null),this._gl.texParameteri(this._gl.TEXTURE_2D,this._gl.TEXTURE_MIN_FILTER,this._gl.NEAREST),this._gl.texParameteri(this._gl.TEXTURE_2D,this._gl.TEXTURE_MAG_FILTER,this._gl.NEAREST),this._gl.texParameteri(this._gl.TEXTURE_2D,this._gl.TEXTURE_WRAP_S,this._gl.CLAMP_TO_EDGE),this._gl.texParameteri(this._gl.TEXTURE_2D,this._gl.TEXTURE_WRAP_T,this._gl.CLAMP_TO_EDGE),this._shadingTexture&&(this._gl.deleteTexture(this._shadingTexture),this._shadingTexture=this._gl.createTexture(),this._gl.bindTexture(this._gl.TEXTURE_2D,this._shadingTexture),this._gl.texImage2D(this._gl.TEXTURE_2D,0,this._gl.DEPTH_COMPONENT32F,t,e,0,this._gl.DEPTH_COMPONENT,this._gl.FLOAT,null),this._gl.texParameteri(this._gl.TEXTURE_2D,this._gl.TEXTURE_MIN_FILTER,this._gl.NEAREST),this._gl.texParameteri(this._gl.TEXTURE_2D,this._gl.TEXTURE_MAG_FILTER,this._gl.NEAREST),this._gl.texParameteri(this._gl.TEXTURE_2D,this._gl.TEXTURE_WRAP_S,this._gl.CLAMP_TO_EDGE),this._gl.texParameteri(this._gl.TEXTURE_2D,this._gl.TEXTURE_WRAP_T,this._gl.CLAMP_TO_EDGE),this._gl.deleteTexture(this._scratchTexture),this._scratchTexture=this._gl.createTexture(),this._gl.bindTexture(this._gl.TEXTURE_2D,this._scratchTexture),this._gl.texImage2D(this._gl.TEXTURE_2D,0,this._gl.DEPTH_COMPONENT32F,t,e,0,this._gl.DEPTH_COMPONENT,this._gl.FLOAT,null),this._gl.texParameteri(this._gl.TEXTURE_2D,this._gl.TEXTURE_MIN_FILTER,this._gl.NEAREST),this._gl.texParameteri(this._gl.TEXTURE_2D,this._gl.TEXTURE_MAG_FILTER,this._gl.NEAREST),this._gl.texParameteri(this._gl.TEXTURE_2D,this._gl.TEXTURE_WRAP_S,this._gl.CLAMP_TO_EDGE),this._gl.texParameteri(this._gl.TEXTURE_2D,this._gl.TEXTURE_WRAP_T,this._gl.CLAMP_TO_EDGE)),this._gl.bindFramebuffer(this._gl.FRAMEBUFFER,this._fb),this._gl.framebufferTexture2D(this._gl.FRAMEBUFFER,this._gl.COLOR_ATTACHMENT0,this._gl.TEXTURE_2D,this._targetTexture,0),this._gl.framebufferTexture2D(this._gl.FRAMEBUFFER,this._gl.DEPTH_ATTACHMENT,this._gl.TEXTURE_2D,this._shadingTexture,0),this._gl.clearDepth(1),this._gl.clear(this._gl.DEPTH_BUFFER_BIT),this._gl.framebufferTexture2D(this._gl.FRAMEBUFFER,this._gl.DEPTH_ATTACHMENT,this._gl.TEXTURE_2D,this._depthTexture,0)}initFrameBuffer(){if(this.isWebGL1())return;let t=this._viewportWidth,e=this._viewportHeight;this._gl.enable(this._gl.SCISSOR_TEST),this._gl.scissor(0,0,t,e),this._gl.viewport(0,0,t,e),this._targetTexture=this._gl.createTexture(),this._depthTexture=this._gl.createTexture(),this._shadingTexture=this._gl.createTexture(),this._scratchTexture=this._gl.createTexture(),this._fb=this._gl.createFramebuffer();var i=this._antialias?ct.screenaa:ct.screen;this._screenshader=this.buildProgram(i.fragmentShader,i.vertexShader,i.uniforms,{}),this._vertexattribpos=this._gl.getAttribLocation(this._screenshader,"vertexPosition");this._screenQuadVBO=this._gl.createBuffer(),this._gl.bindBuffer(this._gl.ARRAY_BUFFER,this._screenQuadVBO),this._gl.bufferData(this._gl.ARRAY_BUFFER,new Float32Array([1,1,-1,1,-1,-1,-1,-1,1,-1,1,1]),this._gl.STATIC_DRAW);let r=ct.ssao;this._AOshader=this.buildProgram(r.fragmentShader,r.vertexShader,r.uniforms,{}),this._aovertexattribpos=this._gl.getAttribLocation(this._AOshader,"vertexPosition");let s=ct.blur;this._blurshader=this.buildProgram(s.fragmentShader,s.vertexShader,s.uniforms,{}),this._blurvertexattribpos=this._gl.getAttribLocation(this._blurshader,"vertexPosition")}renderFrameBuffertoScreen(){this.isWebGL1()||null===this._fb||(this.setViewport(),this._gl.bindFramebuffer(this._gl.FRAMEBUFFER,null),this._gl.clear(this._gl.COLOR_BUFFER_BIT|this._gl.DEPTH_BUFFER_BIT),this._gl.frontFace(this._gl.CCW),this._gl.cullFace(this._gl.BACK),this._gl.useProgram(this._screenshader),this._currentProgram=this._screenshader,this.setDepthTest(-1),this.setDepthWrite(-1),this._gl.bindBuffer(this._gl.ARRAY_BUFFER,this._screenQuadVBO),this._gl.enableVertexAttribArray(this._vertexattribpos),this._gl.vertexAttribPointer(this._vertexattribpos,2,this._gl.FLOAT,!1,0,0),this._gl.activeTexture(this._gl.TEXTURE0),this._gl.bindTexture(this._gl.TEXTURE_2D,this._targetTexture),this._gl.drawArrays(this._gl.TRIANGLES,0,6))}initWebGLObjects(t){if(t.__webglObjects||(t.__webglObjects=[],t.__webglObjectsImmediate=[],t.__webglSprites=[],t.__webglFlares=[]),t.__objectsAdded.length){for(;t.__objectsAdded.length;)this.addObject(t.__objectsAdded[0],t),t.__objectsAdded.splice(0,1);this._currentGeometryGroupHash=-1}for(;t.__objectsRemoved.length;)this.removeObject(t.__objectsRemoved[0],t),t.__objectsRemoved.splice(0,1);for(var e=0,i=t.__webglObjects.length;e<i;e++)this.updateObject(t.__webglObjects[e].object)}getYRatio(){return void 0!==this.rows&&void 0!==this.row?this.rows:1}getXRatio(){return void 0!==this.cols&&void 0!==this.col?this.cols:1}getAspect(t,e){null!=t&&null!=e||(t=this._canvas.width,e=this._canvas.height);var i=t/e;null!=this.rows&&null!=this.cols&&null!=this.row&&null!=this.col&&(i=t/this.cols/(e/this.rows));return i}setTexture(t,e,i){if(t.needsUpdate){t.__webglInit||(t.__webglInit=!0,t.addEventListener("dispose",this.onTextureDispose.bind(this)),t.__webglTexture=this._gl.createTexture(),this.info.memory.textures++),this._gl.activeTexture(this._gl.TEXTURE0+e);var r=i?this._gl.TEXTURE_3D:this._gl.TEXTURE_2D;this._gl.bindTexture(r,t.__webglTexture),this._gl.pixelStorei(this._gl.UNPACK_FLIP_Y_WEBGL,t.flipY),this._gl.pixelStorei(this._gl.UNPACK_PREMULTIPLY_ALPHA_WEBGL,t.premultiplyAlpha),this._gl.pixelStorei(this._gl.UNPACK_ALIGNMENT,t.unpackAlignment),this._gl.pixelStorei(this._gl.PACK_ALIGNMENT,t.unpackAlignment);var s=this.paramToGL(t.format),n=this.paramToGL(t.type);if(i)this.setTextureParameters(this._gl.TEXTURE_3D,t),this._gl.texImage3D(this._gl.TEXTURE_3D,0,this._gl.R32F,t.image.size.z,t.image.size.y,t.image.size.x,0,this._gl.RED,this._gl.FLOAT,t.image.data);else{var a=t.image,o=a.width,l=a.height;void 0===o&&(o=a.length,s==this._gl.RGBA&&(o/=4),l=1),this.setTextureParameters(this._gl.TEXTURE_2D,t),this.isWebGL1()?this._gl.texImage2D(this._gl.TEXTURE_2D,0,s,s,n,t.image):this._gl.texImage2D(this._gl.TEXTURE_2D,0,s,o,l,0,s,n,t.image)}t.needsUpdate=!1,t.onUpdate&&t.onUpdate()}else this._gl.activeTexture(this._gl.TEXTURE0+e),i?this._gl.bindTexture(this._gl.TEXTURE_3D,t.__webglTexture):this._gl.bindTexture(this._gl.TEXTURE_2D,t.__webglTexture)}supportsVolumetric(){return!this.isWebGL1()}enableAttribute(t){this._enabledAttributes[t]||(this._gl.enableVertexAttribArray(t),this._enabledAttributes[t]=!0)}disableAttributes(){for(let t in this._enabledAttributes)this._enabledAttributes[t]&&(this._gl.disableVertexAttribArray(t),this._enabledAttributes[t]=!1)}setPolygonOffset(t,e,i){this._oldPolygonOffset!==t&&(t?this._gl.enable(this._gl.POLYGON_OFFSET_FILL):this._gl.disable(this._gl.POLYGON_OFFSET_FILL))}setLineWidth(t){t!==this._oldLineWidth&&(this._gl.lineWidth(t),this._oldLineWidth=t)}deallocateGeometry(t){if(t.__webglInit=void 0,void 0!==t.__webglVertexBuffer&&this._gl.deleteBuffer(t.__webglVertexBuffer),void 0!==t.__webglColorBuffer&&this._gl.deleteBuffer(t.__webglColorBuffer),void 0!==t.geometryGroups)for(var e=0,i=t.groups;e<i;e++){var r=t.geometryGroups[e];void 0!==r.__webglVertexBuffer&&this._gl.deleteBuffer(r.__webglVertexBuffer),void 0!==r.__webglColorBuffer&&this._gl.deleteBuffer(r.__webglColorBuffer),void 0!==r.__webglNormalBuffer&&this._gl.deleteBuffer(r.__webglNormalBuffer),void 0!==r.__webglFaceBuffer&&this._gl.deleteBuffer(r.__webglFaceBuffer),void 0!==r.__webglLineBuffer&&this._gl.deleteBuffer(r.__webglLineBuffer)}}deallocateMaterial(t){var e=t.program;if(void 0!==e){var i,r,s;t.program=void 0;var n=!1;for(i=0,r=this._programs.length;i<r;i++)if((s=this._programs[i]).program===e){s.usedTimes--,0===s.usedTimes&&(n=!0);break}if(!0===n){var a=[];for(i=0,r=this._programs.length;i<r;i++)(s=this._programs[i]).program!==e&&a.push(s);this._programs=a,this._gl.deleteProgram(e),this.info.memory.programs--}}}deallocateTexture(t){if(t.image&&t.image.__webglTextureCube)this._gl.deleteTexture(t.image.__webglTextureCube);else{if(!t.__webglInit)return;t.__webglInit=!1,this._gl.deleteTexture(t.__webglTexture)}}onGeometryDispose(t){var e=t.target;e.removeEventListener("dispose",this.onGeometryDispose),this.deallocateGeometry(e),this.info.memory.geometries--}onTextureDispose(t){var e=t.target;e.removeEventListener("dispose",this.onTextureDispose),this.deallocateTexture(e),this.info.memory.textures--}onMaterialDispose(t){var e=t.target;e.removeEventListener("dispose",this.onMaterialDispose),this.deallocateMaterial(e)}getShader(t,e){var i;return this.isWebGL1()||e.startsWith("#version")||(e=e.replace(/gl_FragDepthEXT/g,"gl_FragDepth"),e="#version 300 es\n"+(e=(e=(e=(e=(e="fragment"==t?e.replace(/varying/g,"in"):e.replace(/varying/g,"out")).replace(/attribute/g,"in")).replace(/texture2D/g,"texture")).replace(/\/\/DEFINEFRAGCOLOR/g,"out vec4 glFragColor;")).replace(/gl_FragColor/g,"glFragColor"))),"fragment"===t?i=this._gl.createShader(this._gl.FRAGMENT_SHADER):"vertex"===t&&(i=this._gl.createShader(this._gl.VERTEX_SHADER)),null==i?null:(this._gl.shaderSource(i,e),this._gl.compileShader(i),this._gl.getShaderParameter(i,this._gl.COMPILE_STATUS)?i:(console.error(this._gl.getShaderInfoLog(i)),console.error("could not initialize shader"),null))}buildProgram(t,e,i,r){var s,n,a,o,l=[];for(s in l.push(t),l.push(e),r)l.push(s),l.push(r[s]);for(o=l.join(),s=0,n=this._programs.length;s<n;s++){var h=this._programs[s];if(h.code===o)return h.usedTimes++,h.program}if(this.isWebGL1()&&r.volumetric)throw new Error("Volumetric rendering requires webgl2 which is not supported by your hardware.");if(null==(a=this._gl.createProgram()))return null;var c,d,u,f="precision "+this._precision+" float;",p=[r.volumetric?"#version 300 es":"",f].join("\n"),g=[r.volumetric?"#version 300 es":"",r.fragdepth&&this.isWebGL1()?"#extension GL_EXT_frag_depth: enable":"",r.shaded?"#define SHADED 1":"",r.wireframe?"#define WIREFRAME 1":"",f].join("\n"),m=this.getShader("fragment",g+t),v=this.getShader("vertex",p+e);for(d in null!=v&&this._gl.attachShader(a,v),null!=m&&this._gl.attachShader(a,m),this._gl.linkProgram(a),this._gl.getProgramParameter(a,this._gl.LINK_STATUS)||console.error("Could not initialize shader"),a.uniforms={},a.attributes={},c=["viewMatrix","modelViewMatrix","projectionMatrix","normalMatrix","vWidth","vHeight"],i)c.push(d);for(u=0;u<c.length;u++){var _=c[u];a.uniforms[_]=this._gl.getUniformLocation(a,_)}for(c=["position","normal","color","lineDistance","offset","radius"],u=0;u<c.length;u++){var y=c[u];a.attributes[y]=this._gl.getAttribLocation(a,y)}return a.id=this._programs_counter++,this._programs.push({program:a,code:o,usedTimes:1}),this.info.memory.programs=this._programs.length,a}setProgram(t,e,i,r,s,n){if(r.needsUpdate&&(r.program&&this.deallocateMaterial(r),this.initMaterial(r,e,i,s),r.needsUpdate=!1),null==r.program)return null;var a=!1,o=r.program,l=o.uniforms,h=r.uniforms;if(o!=this._currentProgram&&(this._gl.useProgram(o),this._currentProgram=o,a=!0),r.id!=this._currentMaterialId&&(this._currentMaterialId=r.id,a=!0),t!=this._currentCamera&&(this._currentCamera=t,a=!0),l.projectionMatrix&&this._gl.uniformMatrix4fv(l.projectionMatrix,!1,t.projectionMatrix.elements),l.modelViewMatrix&&this._gl.uniformMatrix4fv(l.modelViewMatrix,!1,s._modelViewMatrix.elements),l.normalMatrix&&this._gl.uniformMatrix3fv(l.normalMatrix,!1,s._normalMatrix.elements),l.projinv&&(this._projInverse.getInverse(t.projectionMatrix),this._gl.uniformMatrix4fv(l.projinv,!1,this._projInverse.elements)),l.viewMatrix&&this._gl.uniformMatrix4fv(l.viewMatrix,!1,t.matrixWorldInverse.elements),l.vWidth&&this._gl.uniform1f(l.vWidth,this._viewportWidth),l.vHeight&&this._gl.uniform1f(l.vHeight,this._viewportHeight),a){if(h.fogColor.value=i.color,h.fogNear.value=i.near,h.fogFar.value=i.far,r.shaderID.startsWith("lambert")||"instanced"===r.shaderID||r.shaderID.endsWith("imposter"))this._lightsNeedUpdate&&(this.setupLights(o,e),this._lightsNeedUpdate=!1),h.directionalLightColor.value=this._lights.directional.colors,h.directionalLightDirection.value=this._lights.directional.positions;else if(r.shaderID.endsWith("outline"))h.outlineColor.value=r.outlineColor,h.outlineWidth.value=r.outlineWidth,h.outlinePushback.value=r.outlinePushback,h.outlineMaxPixels.value=r.outlineMaxPixels*this.devicePixelRatio;else if("volumetric"===r.shaderID){s._modelViewMatrix.getScale(this._direction),this._worldInverse.getInverse(s._modelViewMatrix),this._projInverse.getInverse(t.projectionMatrix),this._textureMatrix.multiplyMatrices(s.material.texmatrix,this._worldInverse),this._gl.uniformMatrix4fv(l.textmat,!1,this._textureMatrix.elements),this._gl.uniformMatrix4fv(l.projinv,!1,this._projInverse.elements);let e=Math.min(Math.min(this._direction.x,this._direction.y),this._direction.z);h.step.value=s.material.unit*e,h.maxdepth.value=s.material.maxdepth*e,h.transfermax.value=s.material.transfermax,h.transfermin.value=s.material.transfermin,h.subsamples.value=s.material.subsamples,n.setTexture(s.material.transferfn,4,!1),n.setTexture(s.material.map,3,!0),this._gl.activeTexture(this._gl.TEXTURE5),this._gl.bindTexture(this._gl.TEXTURE_2D,this._depthTexture)}h.opacity.value=r.opacity,this.loadMaterialUniforms(l,h)}return h.shading&&(3==h.shading.value?(this._gl.activeTexture(this._gl.TEXTURE0+this.SHADE_TEXTURE),this._gl.bindTexture(this._gl.TEXTURE_2D,this._shadingTexture)):console.error("Invalid shading textures.")),o}loadMaterialUniforms(t,e){var i,r,s,n;for(i in e)t[i]&&(r=e[i].type,s=e[i].value,n=t[i],"f"===r?this._gl.uniform1f(n,s):"i"===r?this._gl.uniform1i(n,s):"fv"===r?this._gl.uniform3fv(n,s):"c"===r?this._gl.uniform3f(n,s.r,s.g,s.b):"f4"===r&&this._gl.uniform4f(n,s[0],s[1],s[2],s[3]))}addObject(t,e){var i,r,s,n;if(!t.__webglInit&&(t.__webglInit=!0,t._modelViewMatrix=new l.Matrix4,t._normalMatrix=new l.Matrix3,void 0!==t.geometry&&void 0===t.geometry.__webglInit&&(t.geometry.__webglInit=!0,t.geometry.addEventListener("dispose",this.onGeometryDispose.bind(this))),t instanceof Mesh||t instanceof Line))for(i=0,r=(s=t.geometry).geometryGroups.length;i<r;i++)(n=s.geometryGroups[i]).id=this._geometryGroupCounter++,n.__webglVertexBuffer||(t instanceof Mesh?(this.createMeshBuffers(n),s.elementsNeedUpdate=!0,s.normalsNeedUpdate=!0):t instanceof Line&&this.createLineBuffers(n),s.verticesNeedUpdate=!0,s.colorsNeedUpdate=!0);if(!t.__webglActive){if(t instanceof Mesh||t instanceof Line)for(i=0,r=(s=t.geometry).geometryGroups.length;i<r;i++)n=s.geometryGroups[i],this.addBuffer(e.__webglObjects,n,t);else t instanceof Sprite&&e.__webglSprites.push(t);t.__webglActive=!0}}updateObject(t){var e,i=t.geometry;if(t instanceof Mesh||t instanceof Line){for(var r=0,s=i.geometryGroups.length;r<s;r++)e=i.geometryGroups[r],(i.verticesNeedUpdate||i.elementsNeedUpdate||i.colorsNeedUpdate||i.normalsNeedUpdate)&&this.setBuffers(e,this._gl.STATIC_DRAW);i.verticesNeedUpdate=!1,i.elementsNeedUpdate=!1,i.normalsNeedUpdate=!1,i.colorsNeedUpdate=!1,i.buffersNeedUpdate=!1}}removeObject(t,e){t instanceof Mesh||t instanceof Line?this.removeInstances(e.__webglObjects,t):t instanceof Sprite&&this.removeInstancesDirect(e.__webglSprites,t),t.__webglActive=!1}removeInstances(t,e){for(var i=t.length-1;i>=0;--i)t[i].object===e&&t.splice(i,1)}removeInstancesDirect(t,e){for(var i=t.length-1;i>=0;--i)t[i]===e&&t.splice(i,1)}unrollBufferMaterial(t){var e=t.object.material;if(e.volumetric)t.opaque=null,t.transparent=null,t.volumetric=e;else if(e.transparent){if(t.opaque=null,t.volumetric=null,t.transparent=e,!e.wireframe){let i=e.clone();i.opacity=0,t.transparentDepth=i}}else{if(t.opaque=e,t.transparent=null,t.volumetric=null,!e.wireframe){let i=e.clone();i.opacity=0,t.opaqueDepth=i}e.hasAO&&(t.hasAO=!0),(this._AOEnabled||t.hasAO)&&(t.opaqueShaded=e.clone(),t.opaqueShaded.shaded=!0)}}setBuffers(t,e){var i=t.vertexArray,r=t.colorArray;if(void 0!==t.__webglOffsetBuffer?(this._gl.bindBuffer(this._gl.ARRAY_BUFFER,t.__webglOffsetBuffer),this._gl.bufferData(this._gl.ARRAY_BUFFER,i,e)):(this._gl.bindBuffer(this._gl.ARRAY_BUFFER,t.__webglVertexBuffer),this._gl.bufferData(this._gl.ARRAY_BUFFER,i,e)),this._gl.bindBuffer(this._gl.ARRAY_BUFFER,t.__webglColorBuffer),this._gl.bufferData(this._gl.ARRAY_BUFFER,r,e),t.normalArray&&void 0!==t.__webglNormalBuffer){var s=t.normalArray;this._gl.bindBuffer(this._gl.ARRAY_BUFFER,t.__webglNormalBuffer),this._gl.bufferData(this._gl.ARRAY_BUFFER,s,e)}if(t.radiusArray&&void 0!==t.__webglRadiusBuffer&&(this._gl.bindBuffer(this._gl.ARRAY_BUFFER,t.__webglRadiusBuffer),this._gl.bufferData(this._gl.ARRAY_BUFFER,t.radiusArray,e)),t.faceArray&&void 0!==t.__webglFaceBuffer){var n=t.faceArray;this._gl.bindBuffer(this._gl.ELEMENT_ARRAY_BUFFER,t.__webglFaceBuffer),this._gl.bufferData(this._gl.ELEMENT_ARRAY_BUFFER,n,e)}if(t.lineArray&&void 0!==t.__webglLineBuffer){var a=t.lineArray;this._gl.bindBuffer(this._gl.ELEMENT_ARRAY_BUFFER,t.__webglLineBuffer),this._gl.bufferData(this._gl.ELEMENT_ARRAY_BUFFER,a,e)}}createMeshBuffers(t){t.radiusArray&&(t.__webglRadiusBuffer=this._gl.createBuffer()),t.useOffset&&(t.__webglOffsetBuffer=this._gl.createBuffer()),t.__webglVertexBuffer=this._gl.createBuffer(),t.__webglNormalBuffer=this._gl.createBuffer(),t.__webglColorBuffer=this._gl.createBuffer(),t.__webglFaceBuffer=this._gl.createBuffer(),t.__webglLineBuffer=this._gl.createBuffer(),this.info.memory.geometries++}createLineBuffers(t){t.__webglVertexBuffer=this._gl.createBuffer(),t.__webglColorBuffer=this._gl.createBuffer(),this.info.memory.geometries++}addBuffer(t,e,i){t.push({buffer:e,object:i,opaque:null,transparent:null})}setupMatrices(t,e){t._modelViewMatrix.multiplyMatrices(e.matrixWorldInverse,t.matrixWorld),t._normalMatrix.getInverse(t._modelViewMatrix),t._normalMatrix.transpose()}filterFallback(t){return this._gl.LINEAR}setTextureParameters(t,e){t==this._gl.TEXTURE_2D?(this._gl.texParameteri(t,this._gl.TEXTURE_WRAP_S,this._gl.CLAMP_TO_EDGE),this._gl.texParameteri(t,this._gl.TEXTURE_WRAP_T,this._gl.CLAMP_TO_EDGE),this._gl.texParameteri(t,this._gl.TEXTURE_MAG_FILTER,this.filterFallback(e.magFilter)),this._gl.texParameteri(t,this._gl.TEXTURE_MIN_FILTER,this.filterFallback(e.minFilter))):(this._gl.texParameteri(t,this._gl.TEXTURE_WRAP_S,this._gl.CLAMP_TO_EDGE),this._gl.texParameteri(t,this._gl.TEXTURE_WRAP_T,this._gl.CLAMP_TO_EDGE),this._gl.texParameteri(t,this._gl.TEXTURE_WRAP_R,this._gl.CLAMP_TO_EDGE),this._extColorBufferFloat&&this._extFloatLinear?(this._gl.texParameteri(t,this._gl.TEXTURE_MAG_FILTER,this._gl.LINEAR),this._gl.texParameteri(t,this._gl.TEXTURE_MIN_FILTER,this._gl.LINEAR)):(this._gl.texParameteri(t,this._gl.TEXTURE_MAG_FILTER,this._gl.NEAREST),this._gl.texParameteri(t,this._gl.TEXTURE_MIN_FILTER,this._gl.NEAREST)))}paramToGL(t){return t===p?this._gl.UNSIGNED_BYTE:t===m?this._gl.RGBA:t===u?this._gl.NEAREST:0}setupLights(t,e){var i,r,s,n,a,o=this._lights,l=o.directional.colors,h=o.directional.positions,c=0,d=0;for(i=0,r=e.length;i<r;i++)if(n=(s=e[i]).color,a=s.intensity,s instanceof Light){if(this._direction.getPositionFromMatrix(s.matrixWorld),this._vector3.getPositionFromMatrix(s.target.matrixWorld),this._direction.sub(this._vector3),this._direction.normalize(),0===this._direction.x&&0===this._direction.y&&0===this._direction.z)continue;h[d]=this._direction.x,h[d+1]=this._direction.y,h[d+2]=this._direction.z,l[d]=n.r*a,l[d+1]=n.g*a,l[d+2]=n.b*a,d+=3,c++}o.ambient[0]=0,o.ambient[1]=0,o.ambient[2]=0,o.directional.length=c}initGL(){try{if(!OffscreenCanvas||null!=this.rows&&null!=this.cols&&null!=this.row&&null!=this.col){if(!(this._gl=this._canvas.getContext("webgl2",{alpha:this._alpha,premultipliedAlpha:this._premultipliedAlpha,antialias:this._antialias,preserveDrawingBuffer:this._preserveDrawingBuffer}))&&!(this._gl=this._canvas.getContext("experimental-webgl",{alpha:this._alpha,premultipliedAlpha:this._premultipliedAlpha,antialias:this._antialias,preserveDrawingBuffer:this._preserveDrawingBuffer}))&&!(this._gl=this._canvas.getContext("webgl",{alpha:this._alpha,premultipliedAlpha:this._premultipliedAlpha,antialias:this._antialias,preserveDrawingBuffer:this._preserveDrawingBuffer})))throw"Error creating WebGL context."}else(null==gt||gt.isContextLost())&&(pt=new OffscreenCanvas(this._canvas.width,this._canvas.height),gt=pt.getContext("webgl2",{alpha:!0,premultipliedAlpha:this._premultipliedAlpha,antialias:this._antialias,preserveDrawingBuffer:this._preserveDrawingBuffer})),this._offscreen=pt,this._gl=gt,this._bitmap=this._canvas.getContext("bitmaprenderer",{alpha:!0});var t=this._gl.getParameter(this._gl.VERSION);this._webglversion=parseInt(t[6])}catch(t){console.error(t)}}isWebGL1(){return 1==this._webglversion}setDefaultGLState(){this._gl.clearDepth(1),this._gl.clearStencil(0),this._gl.enable(this._gl.DEPTH_TEST),this._gl.depthFunc(this._gl.LEQUAL),this._gl.frontFace(this._gl.CCW),this._gl.cullFace(this._gl.BACK),this._gl.enable(this._gl.CULL_FACE),this._gl.enable(this._gl.BLEND),this._gl.blendEquation(this._gl.FUNC_ADD),this._gl.blendFunc(this._gl.SRC_ALPHA,this._gl.ONE_MINUS_SRC_ALPHA),this._gl.clearColor(this._clearColor.r,this._clearColor.g,this._clearColor.b,this._clearAlpha)}renderObjects(t,e,i,r,s,n,a){var o,l,h,c,d,u,f;e?(d=t.length-1,u=-1,f=-1):(d=0,u=t.length,f=1);for(var p=d;p!==u;p+=f)if((o=t[p]).render){if(l=o.object,h=o.buffer,c=o[i],(o.hasAO||this._AOEnabled)&&o[i+"Shaded"]&&(c=o[i+"Shaded"]),!c)continue;this.setBlending(a),this.setDepthTest(c.depthTest),this.setDepthWrite(c.depthWrite),this.setPolygonOffset(c.polygonOffset,c.polygonOffsetFactor,c.polygonOffsetUnits);var g=l._modelViewMatrix.isReflected();if(this.setMaterialFaces(c,g),this.renderBuffer(r,s,n,c,h,l),(this._outlineEnabled||c.outline)&&!c.wireframe&&"basic"!==c.shaderID&&0!==c.opacity){let t=this._outlineMaterial;"sphereimposter"==c.shaderID?t=this._outlineSphereImposterMaterial:"stickimposter"==c.shaderID&&(t=this._outlineStickImposterMaterial),this.renderBuffer(r,s,n,t,h,l)}}}renderSprites(t,e,i){this._currentGeometryGroupHash=-1,this._currentProgram=null,this._currentCamera=null,this._oldDepthWrite=-1,this._oldDepthTest=-1,this._oldDoubleSided=-1,this._currentMaterialId=-1,this._oldFlipSided=-1,this._lightsNeedUpdate=!0,this.sprites.render(t,e,this._currentWidth,this._currentHeight,i),this._currentGeometryGroupHash=-1,this._currentProgram=null,this._currentCamera=null,this._oldDepthWrite=-1,this._oldDepthTest=-1,this._oldDoubleSided=-1,this._currentMaterialId=-1,this._oldFlipSided=-1}}},529:(t,e,i)=>{"use strict";i.r(e),i.d(e,{Matrix3:()=>Matrix3,Matrix4:()=>Matrix4,Quaternion:()=>Quaternion,Ray:()=>Ray,Vector2:()=>Vector2,Vector3:()=>Vector3,clamp:()=>r,conversionMatrix3:()=>d,degToRad:()=>n});class Quaternion{constructor(t,e,i,r){this.x=t||0,this.y=e||0,this.z=i||0,this.w=void 0!==r?r:1}set(t,e,i,r){return this.x=t,this.y=e,this.z=i,this.w=r,this}copy(t){return this.x=t.x,this.y=t.y,this.z=t.z,this.w=t.w,this}conjugate(){return this.x*=-1,this.y*=-1,this.z*=-1,this}inverse(){return this.conjugate().normalize()}length(){return Math.sqrt(this.x*this.x+this.y*this.y+this.z*this.z+this.w*this.w)}lengthxyz(){return Math.sqrt(this.x*this.x+this.y*this.y+this.z*this.z)}normalize(){let t=this.length();return 0===t?(this.x=0,this.y=0,this.z=0,this.w=1):(t=1/t,this.x*=t,this.y*=t,this.z*=t,this.w*=t),this}multiply(t){return this.multiplyQuaternions(this,t)}multiplyScalar(t){return this.x*=t,this.y*=t,this.z*=t,this.w*=t,this}multiplyQuaternions(t,e){const i=t.x,r=t.y,s=t.z,n=t.w,a=e.x,o=e.y,l=e.z,h=e.w;return this.x=i*h+n*a+r*l-s*o,this.y=r*h+n*o+s*a-i*l,this.z=s*h+n*l+i*o-r*a,this.w=n*h-i*a-r*o-s*l,this}sub(t){return this.x-=t.x,this.y-=t.y,this.z-=t.z,this.w-=t.w,this}clone(){return new Quaternion(this.x,this.y,this.z,this.w)}setFromEuler(t){const e=Math.cos(t.x/2),i=Math.cos(t.y/2),r=Math.cos(t.z/2),s=Math.sin(t.x/2),n=Math.sin(t.y/2),a=Math.sin(t.z/2);return this.x=s*i*r+e*n*a,this.y=e*n*r-s*i*a,this.z=e*i*a+s*n*r,this.w=e*i*r-s*n*a,this}}class Vector2{constructor(t,e){this.x=t||0,this.y=e||0}set(t,e){return this.x=t,this.y=e,this}subVectors(t,e){return this.x=t.x-e.x,this.y=t.y-e.y,this}copy(t){return this.x=t.x,this.y=t.y,this}clone(){return new Vector2(this.x,this.y)}}function r(t,e,i){return Math.min(Math.max(t,e),i)}const s=Math.PI/180;function n(t){return t*s}var a,o,l,h,c;class Matrix4{constructor(t=1,e=0,i=0,r=0,s=0,n=1,a=0,o=0,l=0,h=0,c=1,d=0,u=0,f=0,p=0,g=1){void 0!==t&&"number"!=typeof t?this.elements=new Float32Array(t):(this.elements=new Float32Array(16),this.elements[0]=t,this.elements[4]=e,this.elements[8]=i,this.elements[12]=r,this.elements[1]=s,this.elements[5]=n,this.elements[9]=a,this.elements[13]=o,this.elements[2]=l,this.elements[6]=h,this.elements[10]=c,this.elements[14]=d,this.elements[3]=u,this.elements[7]=f,this.elements[11]=p,this.elements[15]=g)}makeScale(t,e,i){throw new Error("Method not implemented.")}set(t,e,i,r,s,n,a,o,l,h,c,d,u,f,p,g){const m=this.elements;return m[0]=t,m[4]=e,m[8]=i,m[12]=r,m[1]=s,m[5]=n,m[9]=a,m[13]=o,m[2]=l,m[6]=h,m[10]=c,m[14]=d,m[3]=u,m[7]=f,m[11]=p,m[15]=g,this}identity(){return this.set(1,0,0,0,0,1,0,0,0,0,1,0,0,0,0,1),this}copy(t){const e=t.elements;return this.set(e[0],e[4],e[8],e[12],e[1],e[5],e[9],e[13],e[2],e[6],e[10],e[14],e[3],e[7],e[11],e[15]),this}matrix3FromTopLeft(){const t=this.elements;return new Matrix3(t[0],t[4],t[8],t[1],t[5],t[9],t[2],t[6],t[10])}setRotationFromEuler(t,e){const i=this.elements,{x:r,y:s,z:n}=t,a=Math.cos(r),o=Math.sin(r),l=Math.cos(s),h=Math.sin(s),c=Math.cos(n),d=Math.sin(n);if(void 0===e||"XYZ"===e){const t=a*c,e=a*d,r=o*c,s=o*d;i[0]=l*c,i[4]=-l*d,i[8]=h,i[1]=e+r*h,i[5]=t-s*h,i[9]=-o*l,i[2]=s-t*h,i[6]=r+e*h,i[10]=a*l}else console.error(`Error with matrix4 setRotationFromEuler. Order: ${e}`);return this}setRotationFromQuaternion(t){const e=this.elements,{x:i,y:r,z:s,w:n}=t,a=i+i,o=r+r,l=s+s,h=i*a,c=i*o,d=i*l,u=r*o,f=r*l,p=s*l,g=n*a,m=n*o,v=n*l;return e[0]=1-(u+p),e[4]=c-v,e[8]=d+m,e[1]=c+v,e[5]=1-(h+p),e[9]=f-g,e[2]=d-m,e[6]=f+g,e[10]=1-(h+u),this}multiplyMatrices(t,e){const i=t.elements,r=e.elements,s=this.elements,n=i[0],a=i[4],o=i[8],l=i[12],h=i[1],c=i[5],d=i[9],u=i[13],f=i[2],p=i[6],g=i[10],m=i[14],v=i[3],_=i[7],y=i[11],b=i[15],x=r[0],w=r[4],A=r[8],C=r[12],S=r[1],M=r[5],z=r[9],T=r[13],E=r[2],L=r[6],F=r[10],I=r[14],O=r[3],D=r[7],k=r[11],R=r[15];return s[0]=n*x+a*S+o*E+l*O,s[4]=n*w+a*M+o*L+l*D,s[8]=n*A+a*z+o*F+l*k,s[12]=n*C+a*T+o*I+l*R,s[1]=h*x+c*S+d*E+u*O,s[5]=h*w+c*M+d*L+u*D,s[9]=h*A+c*z+d*F+u*k,s[13]=h*C+c*T+d*I+u*R,s[2]=f*x+p*S+g*E+m*O,s[6]=f*w+p*M+g*L+m*D,s[10]=f*A+p*z+g*F+m*k,s[14]=f*C+p*T+g*I+m*R,s[3]=v*x+_*S+y*E+b*O,s[7]=v*w+_*M+y*L+b*D,s[11]=v*A+_*z+y*F+b*k,s[15]=v*C+_*T+y*I+b*R,this}multiplyScalar(t){const e=this.elements;return e[0]*=t,e[4]*=t,e[8]*=t,e[12]*=t,e[1]*=t,e[5]*=t,e[9]*=t,e[13]*=t,e[2]*=t,e[6]*=t,e[10]*=t,e[14]*=t,e[3]*=t,e[7]*=t,e[11]*=t,e[15]*=t,this}makeTranslation(t,e,i){return this.set(1,0,0,t,0,1,0,e,0,0,1,i,0,0,0,1),this}snap(t){t||(t=4);const e=Math.pow(10,4),i=this.elements;for(let t=0;t<16;t++){const r=Math.round(i[t]);r===Math.round(i[t]*e)/e&&(i[t]=r)}return this}transpose(){const t=this.elements;let e;return e=t[1],t[1]=t[4],t[4]=e,e=t[2],t[2]=t[8],t[8]=e,e=t[6],t[6]=t[9],t[9]=e,e=t[3],t[3]=t[12],t[12]=e,e=t[7],t[7]=t[13],t[13]=e,e=t[11],t[11]=t[14],t[14]=e,this}setPosition(t){const e=this.elements;return e[12]=t.x,e[13]=t.y,e[14]=t.z,this}translate(t){const e=this.elements;return e[12]+=t.x,e[13]+=t.y,e[14]+=t.z,this}getInverse(t,e){const i=this.elements,r=t.elements,s=r[0],n=r[4],a=r[8],o=r[12],l=r[1],h=r[5],c=r[9],d=r[13],u=r[2],f=r[6],p=r[10],g=r[14],m=r[3],v=r[7],_=r[11],y=r[15];i[0]=c*g*v-d*p*v+d*f*_-h*g*_-c*f*y+h*p*y,i[4]=o*p*v-a*g*v-o*f*_+n*g*_+a*f*y-n*p*y,i[8]=a*d*v-o*c*v+o*h*_-n*d*_-a*h*y+n*c*y,i[12]=o*c*f-a*d*f-o*h*p+n*d*p+a*h*g-n*c*g,i[1]=d*p*m-c*g*m-d*u*_+l*g*_+c*u*y-l*p*y,i[5]=a*g*m-o*p*m+o*u*_-s*g*_-a*u*y+s*p*y,i[9]=o*c*m-a*d*m-o*l*_+s*d*_+a*l*y-s*c*y,i[13]=a*d*u-o*c*u+o*l*p-s*d*p-a*l*g+s*c*g,i[2]=h*g*m-d*f*m+d*u*v-l*g*v-h*u*y+l*f*y,i[6]=o*f*m-n*g*m-o*u*v+s*g*v+n*u*y-s*f*y,i[10]=n*d*m-o*h*m+o*l*v-s*d*v-n*l*y+s*h*y,i[14]=o*h*u-n*d*u-o*l*f+s*d*f+n*l*g-s*h*g,i[3]=c*f*m-h*p*m-c*u*v+l*p*v+h*u*_-l*f*_,i[7]=n*p*m-a*f*m+a*u*v-s*p*v-n*u*_+s*f*_,i[11]=a*h*m-n*c*m-a*l*v+s*c*v+n*l*_-s*h*_,i[15]=n*c*u-a*h*u+a*l*f-s*c*f-n*l*p+s*h*p;const b=s*i[0]+l*i[4]+u*i[8]+m*i[12];if(0===b){const t="Matrix4.getInverse(): can't invert matrix, determinant is 0";if(e)throw new Error(t);return console.warn(t),this.identity(),this}return this.multiplyScalar(1/b),this}isReflected(){const t=this.elements,e=t[0],i=t[4],r=t[8],s=t[1],n=t[5],a=t[9],o=t[2],l=t[6],h=t[10];return e*n*h+s*l*r+o*i*a-o*n*r-s*i*h-e*l*a<0}scale(t){const e=this.elements,{x:i}=t,{y:r}=t,{z:s}=t;return e[0]*=i,e[4]*=r,e[8]*=s,e[1]*=i,e[5]*=r,e[9]*=s,e[2]*=i,e[6]*=r,e[10]*=s,e[3]*=i,e[7]*=r,e[11]*=s,this}getMaxScaleOnAxis(){const t=this.elements,e=t[0]*t[0]+t[1]*t[1]+t[2]*t[2],i=t[4]*t[4]+t[5]*t[5]+t[6]*t[6],r=t[8]*t[8]+t[9]*t[9]+t[10]*t[10];return Math.sqrt(Math.max(e,Math.max(i,r)))}makeFrustum(t,e,i,r,s,n){const a=this.elements,o=2*s/(e-t),l=2*s/(r-i),h=(e+t)/(e-t),c=(r+i)/(r-i),d=-(n+s)/(n-s),u=-2*n*s/(n-s);return a[0]=o,a[4]=0,a[8]=h,a[12]=0,a[1]=0,a[5]=l,a[9]=c,a[13]=0,a[2]=0,a[6]=0,a[10]=d,a[14]=u,a[3]=0,a[7]=0,a[11]=-1,a[15]=0,this}makePerspective(t,e,i,r){const s=i*Math.tan(n(.5*t)),a=-s,o=a*e,l=s*e;return this.makeFrustum(o,l,a,s,i,r)}makeOrthographic(t,e,i,r,s,n){const a=this.elements,o=1/(e-t),l=1/(i-r),h=1/(n-s),c=(e+t)*o,d=(i+r)*l,u=(n+s)*h;return a[0]=2*o,a[4]=0,a[8]=0,a[12]=-c,a[1]=0,a[5]=2*l,a[9]=0,a[13]=-d,a[2]=0,a[6]=0,a[10]=-2*h,a[14]=-u,a[3]=0,a[7]=0,a[11]=0,a[15]=1,this}isEqual(t){const e=t.elements,i=this.elements;return i[0]===e[0]&&i[4]===e[4]&&i[8]===e[8]&&i[12]===e[12]&&i[1]===e[1]&&i[5]===e[5]&&i[9]===e[9]&&i[13]===e[13]&&i[2]===e[2]&&i[6]===e[6]&&i[10]===e[10]&&i[14]===e[14]&&i[3]===e[3]&&i[7]===e[7]&&i[11]===e[11]&&i[15]===e[15]}clone(){const t=this.elements;return new Matrix4(t[0],t[4],t[8],t[12],t[1],t[5],t[9],t[13],t[2],t[6],t[10],t[14],t[3],t[7],t[11],t[15])}isIdentity(){const t=this.elements;return 1===t[0]&&0===t[4]&&0===t[8]&&0===t[12]&&0===t[1]&&1===t[5]&&0===t[9]&&0===t[13]&&0===t[2]&&0===t[6]&&1===t[10]&&0===t[14]&&0===t[3]&&0===t[7]&&0===t[11]&&1===t[15]}isNearlyIdentity(t){return this.clone().snap(t).isIdentity()}getScale(t){const e=this.elements;return t=t||new Vector3,l.set(e[0],e[1],e[2]),h.set(e[4],e[5],e[6]),c.set(e[8],e[9],e[10]),t.x=l.length(),t.y=h.length(),t.z=c.length(),t}lookAt(t,e,i){const r=this.elements;return c.subVectors(t,e).normalize(),0===c.length()&&(c.z=1),l.crossVectors(i,c).normalize(),0===l.length()&&(c.x+=1e-4,l.crossVectors(i,c).normalize()),h.crossVectors(c,l),r[0]=l.x,r[4]=h.x,r[8]=c.x,r[1]=l.y,r[5]=h.y,r[9]=c.y,r[2]=l.z,r[6]=h.z,r[10]=c.z,this}compose(t,e,i){const r=this.elements;return a.identity(),a.setRotationFromQuaternion(e),o.makeScale(i.x,i.y,i.z),this.multiplyMatrices(a,o),r[12]=t.x,r[13]=t.y,r[14]=t.z,this}}a=new Matrix4,o=new Matrix4;class Vector3{constructor(t,e,i){this.x=t||0,this.y=e||0,this.z=i||0,this.atomid=void 0}set(t,e,i){return this.x=t,this.y=e,this.z=i,this}copy(t){return this.x=t.x,this.y=t.y,this.z=t.z,this}add(t){return this.x+=t.x,this.y+=t.y,this.z+=t.z,this}addVectors(t,e){return this.x=t.x+e.x,this.y=t.y+e.y,this.z=t.z+e.z,this}multiplyVectors(t,e){return this.x=t.x*e.x,this.y=t.y*e.y,this.z=t.z*e.z,this}sub(t){return this.x-=t.x,this.y-=t.y,this.z-=t.z,this}subVectors(t,e){return this.x=t.x-e.x,this.y=t.y-e.y,this.z=t.z-e.z,this}multiplyScalar(t){return this.x*=t,this.y*=t,this.z*=t,this}divideScalar(t){return 0!==t?(this.x/=t,this.y/=t,this.z/=t):(this.x=0,this.y=0,this.z=0),this}max(t){return this.x=Math.max(this.x,t.x),this.y=Math.max(this.y,t.y),this.z=Math.max(this.z,t.z),this}min(t){return this.x=Math.min(this.x,t.x),this.y=Math.min(this.y,t.y),this.z=Math.min(this.z,t.z),this}distanceTo(t){return Math.sqrt(this.distanceToSquared(t))}distanceToSquared(t){const e=this.x-t.x,i=this.y-t.y,r=this.z-t.z;return e*e+i*i+r*r}applyMatrix3(t){const{x:e}=this,{y:i}=this,{z:r}=this,s=t.elements;return this.x=s[0]*e+s[3]*i+s[6]*r,this.y=s[1]*e+s[4]*i+s[7]*r,this.z=s[2]*e+s[5]*i+s[8]*r,this}applyMatrix4(t){const{x:e}=this,{y:i}=this,{z:r}=this,s=t.elements;return this.x=s[0]*e+s[4]*i+s[8]*r+s[12],this.y=s[1]*e+s[5]*i+s[9]*r+s[13],this.z=s[2]*e+s[6]*i+s[10]*r+s[14],this}applyProjection(t){const{x:e}=this,{y:i}=this,{z:r}=this,s=t.elements,n=s[3]*e+s[7]*i+s[11]*r+s[15];return this.x=(s[0]*e+s[4]*i+s[8]*r+s[12])/n,this.y=(s[1]*e+s[5]*i+s[9]*r+s[13])/n,this.z=(s[2]*e+s[6]*i+s[10]*r+s[14])/n,this}applyQuaternion(t){const{x:e}=this,{y:i}=this,{z:r}=this,s=t.x,n=t.y,a=t.z,o=t.w,l={};l.x=2*(i*a-r*n),l.y=2*(r*s-e*a),l.z=2*(e*n-i*s);const h={};return h.x=l.y*a-l.z*n,h.y=l.z*s-l.x*a,h.z=l.x*n-l.y*s,this.x=e+o*l.x+h.x,this.y=i+o*l.y+h.y,this.z=r+o*l.z+h.z,this}negate(){return this.multiplyScalar(-1)}dot(t){return this.x*t.x+this.y*t.y+this.z*t.z}length(){return Math.sqrt(this.x*this.x+this.y*this.y+this.z*this.z)}lengthSq(){return this.x*this.x+this.y*this.y+this.z*this.z}normalize(){return this.divideScalar(this.length())}cross(t){const{x:e}=this,{y:i}=this,{z:r}=this;return this.x=i*t.z-r*t.y,this.y=r*t.x-e*t.z,this.z=e*t.y-i*t.x,this}crossVectors(t,e){return this.x=t.y*e.z-t.z*e.y,this.y=t.z*e.x-t.x*e.z,this.z=t.x*e.y-t.y*e.x,this}equals(t){return this.x==t.x&&this.y==t.y&&this.z==t.z}getPositionFromMatrix(t){return this.x=t.elements[12],this.y=t.elements[13],this.z=t.elements[14],this}setEulerFromRotationMatrix(t,e){const i=t.elements,s=i[0],n=i[4],a=i[8],o=i[5],l=i[9],h=i[6],c=i[10];return void 0===e||"XYZ"===e?(this.y=Math.asin(r(a,-1,1)),Math.abs(a)<.99999?(this.x=Math.atan2(-l,c),this.z=Math.atan2(-n,s)):(this.x=Math.atan2(h,o),this.z=0)):console.error(`Error with vector's setEulerFromRotationMatrix: Unknown order: ${e}`),this}rotateAboutVector(t,e){t.normalize();const i=Math.cos(e),r=Math.sin(e),s=this.clone().multiplyScalar(i),n=t.clone().cross(this).multiplyScalar(r),a=t.clone().multiplyScalar(t.clone().dot(this)).multiplyScalar(1-i),o=s.add(n).add(a);return this.x=o.x,this.y=o.y,this.z=o.z,this}setFromMatrixPosition(t){const e=t.elements;return this.x=e[12],this.y=e[13],this.z=e[14],this}transformDirection(t){const{x:e}=this,{y:i}=this,{z:r}=this,s=t.elements;return this.x=s[0]*e+s[4]*i+s[8]*r,this.y=s[1]*e+s[5]*i+s[9]*r,this.z=s[2]*e+s[6]*i+s[10]*r,this.normalize()}clone(){return new Vector3(this.x,this.y,this.z)}unproject(t){const e=a;return e.multiplyMatrices(t.matrixWorld,e.getInverse(t.projectionMatrix)),this.applyMatrix4(e)}}l=new Vector3,h=new Vector3,c=new Vector3;class Matrix3{constructor(t=1,e=0,i=0,r=0,s=1,n=0,a=0,o=0,l=1){this.elements=new Float32Array(9),this.set(t,e,i,r,s,n,a,o,l)}set(t,e,i,r,s,n,a,o,l){const h=this.elements;return h[0]=t,h[3]=e,h[6]=i,h[1]=r,h[4]=s,h[7]=n,h[2]=a,h[5]=o,h[8]=l,this}identity(){return this.set(1,0,0,0,1,0,0,0,1),this}copy(t){const e=t.elements;this.set(e[0],e[3],e[6],e[1],e[4],e[7],e[2],e[5],e[8])}multiplyScalar(t){const e=this.elements;return e[0]*=t,e[3]*=t,e[6]*=t,e[1]*=t,e[4]*=t,e[7]*=t,e[2]*=t,e[5]*=t,e[8]*=t,this}getInverse3(t){const e=t.elements,i=this.elements;i[0]=e[4]*e[8]-e[5]*e[7],i[3]=e[6]*e[5]-e[3]*e[8],i[6]=e[3]*e[7]-e[6]*e[4],i[1]=e[7]*e[2]-e[1]*e[8],i[4]=e[0]*e[8]-e[6]*e[2],i[7]=e[1]*e[6]-e[0]*e[7],i[2]=e[1]*e[5]-e[2]*e[4],i[5]=e[2]*e[3]-e[0]*e[5],i[8]=e[0]*e[4]-e[1]*e[3];const r=e[0]*i[0]+e[3]*i[1]+e[6]*i[2];return this.multiplyScalar(1/r),this}getInverse(t,e){const i=t.elements,r=this.elements;r[0]=i[10]*i[5]-i[6]*i[9],r[1]=-i[10]*i[1]+i[2]*i[9],r[2]=i[6]*i[1]-i[2]*i[5],r[3]=-i[10]*i[4]+i[6]*i[8],r[4]=i[10]*i[0]-i[2]*i[8],r[5]=-i[6]*i[0]+i[2]*i[4],r[6]=i[9]*i[4]-i[5]*i[8],r[7]=-i[9]*i[0]+i[1]*i[8],r[8]=i[5]*i[0]-i[1]*i[4];const s=i[0]*r[0]+i[1]*r[3]+i[2]*r[6];if(0===s){const t="Matrix3.getInverse(): can't invert matrix, determinant is 0";if(e)throw new Error(t);return console.warn(t),this.identity(),this}return this.multiplyScalar(1/s),this}getDeterminant(){const t=this.elements;return t[0]*t[4]*t[8]+t[1]*t[5]*t[6]+t[2]*t[3]*t[7]-t[2]*t[4]*t[6]-t[1]*t[3]*t[8]-t[0]*t[5]*t[7]}transpose(){let t;const e=this.elements;return t=e[1],e[1]=e[3],e[3]=t,t=e[2],e[2]=e[6],e[6]=t,t=e[5],e[5]=e[7],e[7]=t,this}clone(){const t=this.elements;return new Matrix3(t[0],t[3],t[6],t[1],t[4],t[7],t[2],t[5],t[8])}getMatrix4(){const t=this.elements;return new Matrix4(t[0],t[3],t[6],0,t[1],t[4],t[7],0,t[2],t[5],t[8],0)}}class Ray{constructor(t,e){this.origin=void 0!==t?t:new Vector3,this.direction=void 0!==e?e:new Vector3}set(t,e){return this.origin.copy(t),this.direction.copy(e),this}copy(t){return this.origin.copy(t.origin),this.direction.copy(t.direction),this}at(t,e){return(e||new Vector3).copy(this.direction).multiplyScalar(t).add(this.origin)}recast(t){const e=l;return this.origin.copy(this.at(t,e)),this}closestPointToPoint(t,e){const i=e||new Vector3;i.subVectors(t,this.origin);const r=i.dot(this.direction);return i.copy(this.direction).multiplyScalar(r).add(this.origin)}distanceToPoint(t){const e=l,i=e.subVectors(t,this.origin).dot(this.direction);return e.copy(this.direction).multiplyScalar(i).add(this.origin),e.distanceTo(t)}isIntersectionCylinder(){}isIntersectionSphere(t){return this.distanceToPoint(t.center)<=t.radius}isIntersectionPlane(t){return 0!==t.normal.dot(this.direction)||0===t.distanceToPoint(this.origin)}distanceToPlane(t){const e=t.normal.dot(this.direction);if(0===e)return 0===t.distanceToPoint(this.origin)?0:void 0;return-(this.origin.dot(t.normal)+t.constant)/e}intersectPlane(t,e){const i=this.distanceToPlane(t);if(void 0!==i)return this.at(i,e)}applyMatrix4(t){return this.direction.add(this.origin).applyMatrix4(t),this.origin.applyMatrix4(t),this.direction.sub(this.origin),this}clone(){return(new Ray).copy(this)}}function d(t,e,i,r,s,n){r=r*Math.PI/180,s=s*Math.PI/180,n=n*Math.PI/180;const a=t=>t*t,o=Math.cos(r),l=Math.cos(s),h=Math.cos(n),c=Math.sin(n);return new Matrix3(t,e*h,i*l,0,e*c,i*(o-l*h)/c,0,0,i*Math.sqrt(1-a(o)-a(l)-a(h)+2*o*l*h)/c)}},99:(t,e,i)=>{"use strict";i.r(e),i.d(e,{Cylinder:()=>Cylinder,Sphere:()=>Sphere,Triangle:()=>Triangle});var r=i(529);class Sphere{constructor(t={x:0,y:0,z:0},e=0){this.center=new r.Vector3(t.x,t.y,t.z),this.radius=e}set(t,e){return this.center.copy(t),this.radius=e,this}copy(t){return this.center.copy(t.center),this.radius=t.radius,this}applyMatrix4(t){return this.center.applyMatrix4(t),this.radius=this.radius*t.getMaxScaleOnAxis(),this}translate(t){return this.center.add(t),this}equals(t){return t.center.equals(this.center)&&t.radius===this.radius}clone(){return(new Sphere).copy(this)}}let s=new r.Vector3;class Cylinder{constructor(t=new r.Vector3,e=new r.Vector3,i=0){this.c1=t,this.c2=e,this.radius=i,this.direction=(new r.Vector3).subVectors(this.c2,this.c1).normalize()}copy(t){return this.c1.copy(t.c1),this.c2.copy(t.c2),this.direction.copy(t.direction),this.radius=t.radius,this}lengthSq(){return s.subVectors(this.c2,this.c1).lengthSq()}applyMatrix4(t){return this.direction.add(this.c1).applyMatrix4(t),this.c1.applyMatrix4(t),this.c2.applyMatrix4(t),this.direction.sub(this.c1).normalize(),this.radius=this.radius*t.getMaxScaleOnAxis(),this}}const n=new r.Vector3;class Triangle{constructor(t=new r.Vector3,e=new r.Vector3,i=new r.Vector3){this.a=t,this.b=e,this.c=i}copy(t){return this.a.copy(t.a),this.b.copy(t.b),this.c.copy(t.c),this}applyMatrix4(t){return this.a.applyMatrix4(t),this.b.applyMatrix4(t),this.c.applyMatrix4(t),this}getNormal(){var t=this.a.clone();return t.sub(this.b),n.subVectors(this.c,this.b),t.cross(n),t.normalize(),t}}},222:(t,e,i)=>{"use strict";i.r(e),i.d(e,{CC:()=>CC,Color:()=>Color,builtinColorSchemes:()=>h,chains:()=>l,elementColors:()=>a,htmlColors:()=>r,residues:()=>o,ssColors:()=>s});class Color{constructor(t,e,i){return this.r=0,this.g=0,this.b=0,arguments.length>1&&"number"==typeof t?(this.r=t||0,this.g=e||0,this.b=i||0,this):this.set(t||0)}set(t){return t instanceof Color?t.clone():("number"==typeof t?this.setHex(t):"object"==typeof t&&(this.r=(null==t?void 0:t.r)||0,this.g=(null==t?void 0:t.g)||0,this.b=(null==t?void 0:t.b)||0),this)}setHex(t){return t=Math.floor(t),this.r=(t>>16&255)/255,this.g=(t>>8&255)/255,this.b=(255&t)/255,this}getHex(){return Math.round(255*this.r)<<16|Math.round(255*this.g)<<8|Math.round(255*this.b)}clone(){return new Color(this.r,this.g,this.b)}copy(t){return this.r=t.r,this.g=t.g,this.b=t.b,this}scaled(){var t={};return t.r=Math.round(255*this.r),t.g=Math.round(255*this.g),t.b=Math.round(255*this.b),t.a=1,t}}class CC{static color(t){if(!t)return CC.cache[0];if(t instanceof Color)return t;if("number"==typeof t&&void 0!==CC.cache[t])return CC.cache[t];if(t&&Array.isArray(t))return t.map(CC.color);let e=CC.getHex(t),i=new Color(e);return CC.cache[e]=i,i}static getHex(t){var e;if(Array.isArray(t))return t.map(CC.getHex);if("string"==typeof t){let i=t;if(!isNaN(parseInt(i)))return parseInt(i);if(i=i.trim(),4==i.length&&"#"==i[0]&&(i="#"+i[1]+i[1]+i[2]+i[2]+i[3]+i[3]),7==i.length&&"#"==i[0])return parseInt(i.substring(1),16);let r=CC.rgbRegEx.exec(i);if(r){""!=r[1]&&console.log("WARNING: Opacity value in rgba ignored.  Specify separately as opacity attribute.");let t=0;for(let e=2;e<5;e++){t*=256;let i=r[e].endsWith("%")?255*parseFloat(r[e])/100:parseFloat(r[e]);t+=Math.round(i)}return t}return(null===(e=null===window||void 0===window?void 0:window.$3Dmol)||void 0===e?void 0:e.htmlColors[t.toLowerCase()])||0}return t}}CC.rgbRegEx=/rgb(a?)\(\s*([^ ,\)\t]+)\s*,\s*([^ ,\)\t]+)\s*,\s*([^ ,\)\t]+)/i,CC.cache={0:new Color(0)};const r={aliceblue:15792383,antiquewhite:16444375,aqua:65535,aquamarine:8388564,azure:15794175,beige:16119260,bisque:16770244,black:0,blanchedalmond:16772045,blue:255,blueviolet:9055202,brown:10824234,burlywood:14596231,cadetblue:6266528,chartreuse:8388352,chocolate:13789470,coral:16744272,cornflowerblue:6591981,cornsilk:16775388,crimson:14423100,cyan:65535,darkblue:139,darkcyan:35723,darkgoldenrod:12092939,darkgray:11119017,darkgrey:11119017,darkgreen:25600,darkkhaki:12433259,darkmagenta:9109643,darkolivegreen:5597999,darkorange:16747520,darkorchid:10040012,darkred:9109504,darksalmon:15308410,darkseagreen:9419919,darkslateblue:4734347,darkslategray:3100495,darkslategrey:3100495,darkturquoise:52945,darkviolet:9699539,deeppink:16716947,deepskyblue:49151,dimgray:6908265,dimgrey:6908265,dodgerblue:2003199,firebrick:11674146,floralwhite:16775920,forestgreen:2263842,fuchsia:16711935,gainsboro:14474460,ghostwhite:16316671,gold:16766720,goldenrod:14329120,gray:8421504,grey:8421504,green:32768,greenyellow:11403055,honeydew:15794160,hotpink:16738740,indianred:13458524,indigo:4915330,ivory:16777200,khaki:15787660,lavender:15132410,lavenderblush:16773365,lawngreen:8190976,lemonchiffon:16775885,lightblue:11393254,lightcoral:15761536,lightcyan:14745599,lightgoldenrodyellow:16448210,lightgray:13882323,lightgrey:13882323,lightgreen:9498256,lightpink:16758465,lightsalmon:16752762,lightseagreen:2142890,lightskyblue:8900346,lightslategray:7833753,lightslategrey:7833753,lightsteelblue:11584734,lightyellow:16777184,lime:65280,limegreen:3329330,linen:16445670,magenta:16711935,maroon:8388608,mediumaquamarine:6737322,mediumblue:205,mediumorchid:12211667,mediumpurple:9662683,mediumseagreen:3978097,mediumslateblue:8087790,mediumspringgreen:64154,mediumturquoise:4772300,mediumvioletred:13047173,midnightblue:1644912,mintcream:16121850,mistyrose:16770273,moccasin:16770229,navajowhite:16768685,navy:128,oldlace:16643558,olive:8421376,olivedrab:7048739,orange:16753920,orangered:16729344,orchid:14315734,palegoldenrod:15657130,palegreen:10025880,paleturquoise:11529966,palevioletred:14381203,papayawhip:16773077,peachpuff:16767673,peru:13468991,pink:16761035,plum:14524637,powderblue:11591910,purple:8388736,rebeccapurple:6697881,red:16711680,rosybrown:12357519,royalblue:4286945,saddlebrown:9127187,salmon:16416882,sandybrown:16032864,seagreen:3050327,seashell:16774638,sienna:10506797,silver:12632256,skyblue:8900331,slateblue:6970061,slategray:7372944,slategrey:7372944,snow:16775930,springgreen:65407,steelblue:4620980,tan:13808780,teal:32896,thistle:14204888,tomato:16737095,turquoise:4251856,violet:15631086,wheat:16113331,white:16777215,whitesmoke:16119285,yellow:16776960,yellowgreen:10145074},s={pyMol:{h:16711680,s:16776960,c:65280},Jmol:{h:16711808,s:16762880,c:16777215}},n={H:16777215,He:16761035,HE:16761035,Li:11674146,LI:11674146,B:65280,C:13158600,N:9408511,O:15728640,F:14329120,Na:255,NA:255,Mg:2263842,MG:2263842,Al:8421520,AL:8421520,Si:14329120,SI:14329120,P:16753920,S:16762930,Cl:65280,CL:65280,Ca:8421520,CA:8421520,Ti:8421520,TI:8421520,Cr:8421520,CR:8421520,Mn:8421520,MN:8421520,Fe:16753920,FE:16753920,Ni:10824234,NI:10824234,Cu:10824234,CU:10824234,Zn:10824234,ZN:10824234,Br:10824234,BR:10824234,Ag:8421520,AG:8421520,I:10494192,Ba:16753920,BA:16753920,Au:14329120,AU:14329120},a={defaultColor:16716947,Jmol:{H:16777215,He:14286847,HE:14286847,Li:13402367,LI:13402367,Be:12779264,BE:12779264,B:16758197,C:9474192,N:3166456,O:16715021,F:9494608,Ne:11789301,NE:11789301,Na:11230450,NA:11230450,Mg:9109248,MG:9109248,Al:12560038,AL:12560038,Si:1578e4,SI:1578e4,P:16744448,S:16777008,Cl:2093087,CL:2093087,Ar:8442339,AR:8442339,K:9388244,Ca:4062976,CA:4062976,Sc:15132390,SC:15132390,Ti:12567239,TI:12567239,V:10921643,Cr:9083335,CR:9083335,Mn:10255047,MN:10255047,Fe:14706227,FE:14706227,Co:15765664,CO:15765664,Ni:5296208,NI:5296208,Cu:13140019,CU:13140019,Zn:8224944,ZN:8224944,Ga:12750735,GA:12750735,Ge:6721423,GE:6721423,As:12419299,AS:12419299,Se:16752896,SE:16752896,Br:10889513,BR:10889513,Kr:6076625,KR:6076625,Rb:7351984,RB:7351984,Sr:65280,SR:65280,Y:9764863,Zr:9756896,ZR:9756896,Nb:7586505,NB:7586505,Mo:5551541,MO:5551541,Tc:3907230,TC:3907230,Ru:2396047,RU:2396047,Rh:687500,RH:687500,Pd:27013,PD:27013,Ag:12632256,AG:12632256,Cd:16767375,CD:16767375,In:10909043,IN:10909043,Sn:6717568,SN:6717568,Sb:10380213,SB:10380213,Te:13924864,TE:13924864,I:9699476,Xe:4366e3,XE:4366e3,Cs:5707663,CS:5707663,Ba:51456,BA:51456,La:7394559,LA:7394559,Ce:16777159,CE:16777159,Pr:14286791,PR:14286791,Nd:13107143,ND:13107143,Pm:10747847,PM:10747847,Sm:9437127,SM:9437127,Eu:6422471,EU:6422471,Gd:4587463,GD:4587463,Tb:3211207,TB:3211207,Dy:2097095,DY:2097095,Ho:65436,HO:65436,Er:58997,ER:58997,Tm:54354,TM:54354,Yb:48952,YB:48952,Lu:43812,LU:43812,Hf:5096191,HF:5096191,Ta:5089023,TA:5089023,W:2200790,Re:2522539,RE:2522539,Os:2516630,OS:2516630,Ir:1528967,IR:1528967,Pt:13684960,PT:13684960,Au:16765219,AU:16765219,Hg:12105936,HG:12105936,Tl:10900557,TL:10900557,Pb:5724513,PB:5724513,Bi:10375093,BI:10375093,Po:11230208,PO:11230208,At:7688005,AT:7688005,Rn:4358806,RN:4358806,Fr:4325478,FR:4325478,Ra:32e3,RA:32e3,Ac:7384058,AC:7384058,Th:47871,TH:47871,Pa:41471,PA:41471,U:36863,Np:33023,NP:33023,Pu:27647,PU:27647,Am:5528818,AM:5528818,Cm:7888099,CM:7888099,Bk:9064419,BK:9064419,Cf:10565332,CF:10565332,Es:11739092,ES:11739092,Fm:11739066,FM:11739066,Md:11734438,MD:11734438,No:12389767,NO:12389767,Lr:13041766,LR:13041766,Rf:13369433,RF:13369433,Db:13697103,DB:13697103,Sg:14221381,SG:14221381,Bh:14680120,BH:14680120,Hs:15073326,HS:15073326,Mt:15400998,MT:15400998},rasmol:n,defaultColors:Object.assign({},n),greenCarbon:Object.assign(Object.assign({},n),{C:65280}),cyanCarbon:Object.assign(Object.assign({},n),{C:65535}),magentaCarbon:Object.assign(Object.assign({},n),{C:16711935}),yellowCarbon:Object.assign(Object.assign({},n),{C:16776960}),whiteCarbon:Object.assign(Object.assign({},n),{C:16777215}),orangeCarbon:Object.assign(Object.assign({},n),{C:16753920}),purpleCarbon:Object.assign(Object.assign({},n),{C:8388736}),blueCarbon:Object.assign(Object.assign({},n),{C:255})},o={amino:{ALA:13158600,ARG:1334015,ASN:56540,ASP:15075850,CYS:15132160,GLN:56540,GLU:15075850,GLY:15461355,HIS:8553170,ILE:1016335,LEU:1016335,LYS:1334015,MET:15132160,PHE:3289770,PRO:14456450,SER:16422400,THR:16422400,TRP:11819700,TYR:3289770,VAL:1016335,ASX:16738740,GLX:16738740},shapely:{ALA:9240460,ARG:124,ASN:16743536,ASP:10485826,CYS:16777072,GLN:16731212,GLU:6684672,GLY:16777215,HIS:7368959,ILE:19456,LEU:4546117,LYS:4671416,MET:12099650,PHE:5459026,PRO:5395026,SER:16740418,THR:12078080,TRP:5195264,TYR:9203788,VAL:16747775,ASX:16711935,GLX:16711935},nucleic:{A:10526975,G:16740464,I:8454143,C:16747595,T:10551200,U:16744576}},l={atom:{A:12636415,B:11599792,C:16761032,D:16777088,E:16761087,F:11596016,G:16765040,H:15761536,I:16113331,J:49151,K:13458524,L:6737322,M:10145074,N:15631086,O:52945,P:65407,Q:3978097,R:139,S:12433259,T:25600,U:8388608,V:8421376,W:8388736,X:32896,Y:12092939,Z:11674146},hetatm:{A:9478351,B:8441752,C:13602992,D:13619056,E:13603023,F:8437952,G:13607008,H:12603504,I:12955267,J:42959,K:11881548,L:5682578,M:9090346,N:12481214,O:46753,P:53103,Q:3447649,R:187,S:10854235,T:37888,U:11534336,V:11579392,W:11534512,X:45232,Y:15250963,Z:12726834}},h={ssPyMol:{prop:"ss",map:s.pyMol},ssJmol:{prop:"ss",map:s.Jmol},Jmol:{prop:"elem",map:a.Jmol},amino:{prop:"resn",map:o.amino},shapely:{prop:"resn",map:o.shapely},nucleic:{prop:"resn",map:o.nucleic},chain:{prop:"chain",map:l.atom},rasmol:{prop:"elem",map:a.rasmol},default:{prop:"elem",map:a.defaultColors},greenCarbon:{prop:"elem",map:a.greenCarbon},chainHetatm:{prop:"chain",map:l.hetatm},cyanCarbon:{prop:"elem",map:a.cyanCarbon},magentaCarbon:{prop:"elem",map:a.magentaCarbon},purpleCarbon:{prop:"elem",map:a.purpleCarbon},whiteCarbon:{prop:"elem",map:a.whiteCarbon},orangeCarbon:{prop:"elem",map:a.orangeCarbon},yellowCarbon:{prop:"elem",map:a.yellowCarbon},blueCarbon:{prop:"elem",map:a.blueCarbon}}},421:(t,e,i)=>{"use strict";i.r(e),i.d(e,{CAP:()=>it,CC:()=>s.CC,CONTEXTS_PER_VIEWPORT:()=>Wt,Color:()=>s.Color,CustomLinear:()=>r.CustomLinear,Cylinder:()=>J.Cylinder,GLDraw:()=>rt,GLModel:()=>GLModel,GLShape:()=>GLShape,GLViewer:()=>GLViewer,GLVolumetricRender:()=>GLVolumetricRender,Gradient:()=>r.Gradient,GradientType:()=>r.GradientType,Label:()=>Label,LabelCount:()=>a,MarchingCube:()=>MarchingCube,MarchingCubeInitializer:()=>MarchingCubeInitializer,Matrix3:()=>$.Matrix3,Matrix4:()=>$.Matrix4,Parsers:()=>K,PausableTimer:()=>L.PausableTimer,PointGrid:()=>PointGrid,ProteinSurface:()=>ProteinSurface,Quaternion:()=>$.Quaternion,ROYGB:()=>r.ROYGB,RWB:()=>r.RWB,Ray:()=>$.Ray,Sinebow:()=>r.Sinebow,Sphere:()=>J.Sphere,SurfaceType:()=>Q,Triangle:()=>J.Triangle,Vector2:()=>$.Vector2,Vector3:()=>$.Vector3,VolumeData:()=>st.VolumeData,adjustVolumeStyle:()=>L.adjustVolumeStyle,applyPartialCharges:()=>h,autoinit:()=>Xt,autoload:()=>$t,base64ToArray:()=>L.base64ToArray,bondLength:()=>C.bondLength,builtinColorSchemes:()=>s.builtinColorSchemes,builtinGradients:()=>r.builtinGradients,chains:()=>s.chains,clamp:()=>$.clamp,conversionMatrix3:()=>$.conversionMatrix3,createStereoViewer:()=>Zt,createViewer:()=>qt,createViewerGrid:()=>Yt,deepCopy:()=>L.deepCopy,degToRad:()=>$.degToRad,download:()=>L.download,drawCartoon:()=>xt,elementColors:()=>s.elementColors,extend:()=>L.extend,get:()=>L.get,getAtomProperty:()=>L.getAtomProperty,getColorFromStyle:()=>L.getColorFromStyle,getElement:()=>L.getElement,getExtent:()=>L.getExtent,getGradient:()=>r.getGradient,getPropertyRange:()=>L.getPropertyRange,getbin:()=>L.getbin,htmlColors:()=>s.htmlColors,inflateString:()=>L.inflateString,isEmptyObject:()=>L.isEmptyObject,isNumeric:()=>L.isNumeric,makeFunction:()=>L.makeFunction,mergeGeos:()=>L.mergeGeos,normalizeValue:()=>r.normalizeValue,partialCharges:()=>l,processing_autoinit:()=>Kt,residues:()=>s.residues,setBondLength:()=>C.setBondLength,setSyncSurface:()=>et,specStringToObject:()=>L.specStringToObject,splitMesh:()=>wt,ssColors:()=>s.ssColors,subdivide_spline:()=>nt,syncSurface:()=>tt,viewers:()=>Qt});var r=i(546),s=i(222),n=i(638);let a=0;function o(t,e,i){var r=i;return void 0!==t&&(t instanceof s.Color?r=t.scaled():void 0!==(r=s.CC.color(t)).scaled&&(r=r.scaled())),void 0!==e&&(r.a=parseFloat(e)),r}class Label{constructor(t,e){this.id=a++,this.stylespec=e||{},this.canvas=document.createElement("canvas"),this.canvas.width=134,this.canvas.height=35,this.context=this.canvas.getContext("2d"),this.sprite=new n.Sprite,this.text=t,this.frame=this.stylespec.frame}getStyle(){return this.stylespec}hide(){this.sprite&&(this.sprite.visible=!1)}show(){this.sprite&&(this.sprite.visible=!0)}setContext(){var t=this.stylespec,e=void 0!==t.useScreen&&t.useScreen,i=t.showBackground;"0"!==i&&"false"!==i||(i=!1),void 0===i&&(i=!0);var s=t.font?t.font:"sans-serif",a=parseInt(t.fontSize)?parseInt(t.fontSize):18,l=o(t.fontColor,t.fontOpacity,{r:255,g:255,b:255,a:1}),h=t.padding?t.padding:4,c=t.borderThickness?t.borderThickness:0,d=o(t.backgroundColor,t.backgroundOpacity,{r:0,g:0,b:0,a:1}),u=o(t.borderColor,t.borderOpacity,d),f=t.position?t.position:{x:-10,y:1,z:1},p=void 0===t.inFront||t.inFront;"false"!==p&&"0"!==p||(p=!1);var g=t.alignment||n.SpriteAlignment.topLeft;"string"==typeof g&&g in n.SpriteAlignment&&(g=n.SpriteAlignment[g]);var m="";t.bold&&(m="bold "),this.context.font=m+a+"px  "+s;var v=this.context.measureText(this.text).width;i||(c=0);var _=v+2.5*c+2*h,y=1.25*a+2*c+2*h;if(t.backgroundImage){var b=t.backgroundImage,x=t.backgroundWidth?t.backgroundWidth:b.width,w=t.backgroundHeight?t.backgroundHeight:b.height;x>_&&(_=x),w>y&&(y=w)}if(this.canvas.width=_,this.canvas.height=y,this.context.clearRect(0,0,this.canvas.width,this.canvas.height),m="",t.bold&&(m="bold "),this.context.font=m+a+"px  "+s,this.context.fillStyle="rgba("+d.r+","+d.g+","+d.b+","+d.a+")",this.context.strokeStyle="rgba("+u.r+","+u.g+","+u.b+","+u.a+")",t.backgroundGradient){let e=this.context.createLinearGradient(0,y/2,_,y/2),i=r.Gradient.getGradient(t.backgroundGradient),s=i.range(),n=-1,a=1;s&&(n=s[0],a=s[1]);let l=a-n;for(let t=0;t<1.01;t+=.1){let r=o(i.valueToHex(n+l*t)),s="rgba("+r.r+","+r.g+","+r.b+","+r.a+")";e.addColorStop(t,s)}this.context.fillStyle=e}this.context.lineWidth=c,i&&function(t,e,i,r,s,n,a){t.beginPath(),t.moveTo(e+n,i),t.lineTo(e+r-n,i),t.quadraticCurveTo(e+r,i,e+r,i+n),t.lineTo(e+r,i+s-n),t.quadraticCurveTo(e+r,i+s,e+r-n,i+s),t.lineTo(e+n,i+s),t.quadraticCurveTo(e,i+s,e,i+s-n),t.lineTo(e,i+n),t.quadraticCurveTo(e,i,e+n,i),t.closePath(),t.fill(),a&&t.stroke()}(this.context,c,c,_-2*c,y-2*c,6,c>0),t.backgroundImage&&this.context.drawImage(b,0,0,_,y),this.context.fillStyle="rgba("+l.r+","+l.g+","+l.b+","+l.a+")",this.context.fillText(this.text,c+h,a+c+h,v);var A=new n.Texture(this.canvas);A.needsUpdate=!0,this.sprite.material=new n.SpriteMaterial({map:A,useScreenCoordinates:e,alignment:g,depthTest:!p,screenOffset:t.screenOffset||null}),this.sprite.scale.set(1,1,1),this.sprite.position.set(f.x,f.y,f.z)}dispose(){void 0!==this.sprite.material.map&&this.sprite.material.map.dispose(),void 0!==this.sprite.material&&this.sprite.material.dispose()}}const l={"ALA:N":-.15,"ALA:CA":.1,"ALA:CB":0,"ALA:C":.6,"ALA:O":-.55,"ARG:N":-.15,"ARG:CA":.1,"ARG:CB":0,"ARG:CG":0,"ARG:CD":.1,"ARG:NE":-.1,"ARG:CZ":.5,"ARG:NH1":.25,"ARG:NH2":.25,"ARG:C":.6,"ARG:O":-.55,"ASN:N":-.15,"ASN:CA":.1,"ASN:CB":0,"ASN:CG":.55,"ASN:OD1":-.55,"ASN:ND2":0,"ASN:C":.6,"ASN:O":-.55,"ASP:N":-.15,"ASP:CA":.1,"ASP:CB":0,"ASP:CG":.14,"ASP:OD1":-.57,"ASP:OD2":-.57,"ASP:C":.6,"ASP:O":-.55,"CYS:N":-.15,"CYS:CA":.1,"CYS:CB":.19,"CYS:SG":-.19,"CYS:C":.6,"CYS:O":-.55,"GLN:N":-.15,"GLN:CA":.1,"GLN:CB":0,"GLN:CG":0,"GLN:CD":.55,"GLN:OE1":-.55,"GLN:NE2":0,"GLN:C":.6,"GLN:O":-.55,"GLU:N":-.15,"GLU:CA":.1,"GLU:CB":0,"GLU:CG":0,"GLU:CD":.14,"GLU:OE1":-.57,"GLU:OE2":-.57,"GLU:C":.6,"GLU:O":-.55,"GLY:N":-.15,"GLY:CA":.1,"GLY:C":.6,"GLY:O":-.55,"HIS:N":-.15,"HIS:CA":.1,"HIS:CB":0,"HIS:CG":.1,"HIS:ND1":-.1,"HIS:CD2":.1,"HIS:NE2":-.4,"HIS:CE1":.3,"HIS:C":.6,"HIS:O":-.55,"ILE:N":-.15,"ILE:CA":.1,"ILE:CB":0,"ILE:CG2":0,"ILE:CG1":0,"ILE:CD":0,"ILE:C":.6,"ILE:O":-.55,"LEU:N":-.15,"LEU:CA":.1,"LEU:CB":0,"LEU:CG":0,"LEU:CD1":0,"LEU:CD2":0,"LEU:C":.6,"LEU:O":-.55,"LYS:N":-.15,"LYS:CA":.1,"LYS:CB":0,"LYS:CG":0,"LYS:CD":0,"LYS:CE":.25,"LYS:NZ":.75,"LYS:C":.6,"LYS:O":-.55,"MET:N":-.15,"MET:CA":.1,"MET:CB":0,"MET:CG":.06,"MET:SD":-.12,"MET:CE":.06,"MET:C":.6,"MET:O":-.55,"PHE:N":-.15,"PHE:CA":.1,"PHE:CB":0,"PHE:CG":0,"PHE:CD1":0,"PHE:CD2":0,"PHE:CE1":0,"PHE:CE2":0,"PHE:CZ":0,"PHE:C":.6,"PHE:O":-.55,"PRO:N":-.25,"PRO:CD":.1,"PRO:CA":.1,"PRO:CB":0,"PRO:CG":0,"PRO:C":.6,"PRO:O":-.55,"SER:N":-.15,"SER:CA":.1,"SER:CB":.25,"SER:OG":-.25,"SER:C":.6,"SER:O":-.55,"THR:N":-.15,"THR:CA":.1,"THR:CB":.25,"THR:OG1":-.25,"THR:CG2":0,"THR:C":.6,"THR:O":-.55,"TRP:N":-.15,"TRP:CA":.1,"TRP:CB":0,"TRP:CG":-.03,"TRP:CD2":.1,"TRP:CE2":-.04,"TRP:CE3":-.03,"TRP:CD1":.06,"TRP:NE1":-.06,"TRP:CZ2":0,"TRP:CZ3":0,"TRP:CH2":0,"TRP:C":.6,"TRP:O":-.55,"TYR:N":-.15,"TYR:CA":.1,"TYR:CB":0,"TYR:CG":0,"TYR:CD1":0,"TYR:CE1":0,"TYR:CD2":0,"TYR:CE2":0,"TYR:CZ":.25,"TYR:OH":-.25,"TYR:C":.6,"TYR:O":-.55,"VAL:N":-.15,"VAL:CA":.1,"VAL:CB":0,"VAL:CG1":0,"VAL:CG2":0,"VAL:C":.6,"VAL:O":-.55};function h(t,e){if((!e||void 0===t.partialCharge)&&t.resn&&t.atom){var i=t.resn+":"+t.atom;t.properties.partialCharge=l[i]}}var c=i(797),d=i(865),u=i(392);function f(t,e){for(var i=[[]],r=void 0===(e=e||{}).assignBonds||e.assignBonds,s=t.trimStart().split(/\r?\n|\r/);s.length>0&&!(s.length<3);){var a=parseInt(s[0]);if(isNaN(a)||a<=0)break;if(s.length<a+2)break;var o=/Lattice\s*=\s*["\{\}]([^"\{\}]+)["\{\}]\s*/gi.exec(s[1]);if(null!=o&&o.length>1){var l=new Float32Array(o[1].split(/\s+/)),h=new n.Matrix3(l[0],l[3],l[6],l[1],l[4],l[7],l[2],l[5],l[8]);i.modelData=[{cryst:{matrix:h}}]}for(var c=2,d=i[i.length-1].length,f=d+a,p=d;p<f;p++){var g=s[c++].trim().split(/\s+/),m={};m.serial=p;var v=g[0];m.atom=m.elem=v[0].toUpperCase()+v.substring(1,2).toLowerCase(),m.x=parseFloat(g[1]),m.y=parseFloat(g[2]),m.z=parseFloat(g[3]),m.hetflag=!0,m.bonds=[],m.bondOrder=[],m.properties={},i[i.length-1][p]=m,g.length>=7&&(m.dx=parseFloat(g[4]),m.dy=parseFloat(g[5]),m.dz=parseFloat(g[6]))}if(!e.multimodel)break;i.push([]),s.splice(0,c)}if(r)for(let t=0;t<i.length;t++)(0,u.assignBonds)(i[t],e);if(e.onemol){var _=i;(i=[]).push(_[0]);for(let t=1;t<_.length;t++){let e=i[0].length;for(let r=0;r<_[t].length;r++){let s=_[t][r];for(let t=0;t<s.bonds.length;t++)s.bonds[t]=s.bonds[t]+e;s.index=i[0].length,s.serial=i[0].length,i[0].push(s)}}}return i}function p(t,e){var i="V2000",r=t.split(/\r?\n|\r/);return r.length>3&&r[3].length>38&&(i=r[3].substring(34,39)),"V2000"===i?function(t,e){var i=[[]],r=!1;for(void 0!==e.keepH&&(r=!e.keepH);t.length>0&&!(t.length<4);){var s=parseInt(t[3].substring(0,3));if(isNaN(s)||s<=0)break;var n=parseInt(t[3].substring(3,6)),a=4;if(t.length<4+s+n)break;var o,l,h=[],c=i[i.length-1].length,d=c+s;for(o=c;o<d;o++,a++){var u={},f=(l=t[a]).substring(31,34).replace(/ /g,"");u.atom=u.elem=f[0].toUpperCase()+f.substring(1).toLowerCase(),"H"===u.elem&&r||(u.serial=o,h[o]=i[i.length-1].length,u.x=parseFloat(l.substring(0,10)),u.y=parseFloat(l.substring(10,20)),u.z=parseFloat(l.substring(20,30)),u.hetflag=!0,u.bonds=[],u.bondOrder=[],u.properties={},u.index=i[i.length-1].length,i[i.length-1].push(u))}for(o=0;o<n;o++,a++){l=t[a];var p=h[parseInt(l.substring(0,3))-1+c],g=h[parseInt(l.substring(3,6))-1+c],m=parseFloat(l.substring(6));void 0!==p&&void 0!==g&&(i[i.length-1][p].bonds.push(g),i[i.length-1][p].bondOrder.push(m),i[i.length-1][g].bonds.push(p),i[i.length-1][g].bondOrder.push(m))}if(!e.multimodel)break;for(e.onemol||i.push([]);"$$$$"!==t[a]&&a<t.length;)a++;t.splice(0,++a)}return i}(r,e):"V3000"===i?function(t,e){var i=[[]],r=!1;for(void 0!==e.keepH&&(r=!e.keepH);t.length>0&&!(t.length<8)&&t[4].startsWith("M  V30 BEGIN CTAB")&&t[5].startsWith("M  V30 COUNTS")&&!(t[5].length<14);){var s=t[5].substring(13).match(/\S+/g);if(s.length<2)break;var n=parseInt(s[0]);if(isNaN(n)||n<=0)break;var a=parseInt(s[1]),o=7;if(t.length<8+n+a)break;var l,h=[],c=i[i.length-1].length,d=c+n;for(l=c;l<d;l++,o++){var u=t[o].substring(6).match(/\S+/g);if(u.length>4){var f={},p=u[1].replace(/ /g,"");f.atom=f.elem=p[0].toUpperCase()+p.substring(1).toLowerCase(),"H"===f.elem&&r||(f.serial=l,h[l]=i[i.length-1].length,f.x=parseFloat(u[2]),f.y=parseFloat(u[3]),f.z=parseFloat(u[4]),f.hetflag=!0,f.bonds=[],f.bondOrder=[],f.properties={},f.index=i[i.length-1].length,i[i.length-1].push(f))}}if("M  V30 END ATOM"!==t[o])break;if(o++,0===a||"M  V30 BEGIN BOND"!==t[o])break;for(o++,l=0;l<a;l++,o++){var g=t[o].substring(6).match(/\S+/g);if(g.length>3){var m=h[parseInt(g[2])-1+c],v=h[parseInt(g[3])-1+c],_=parseFloat(g[1]);void 0!==m&&void 0!==v&&(i[i.length-1][m].bonds.push(v),i[i.length-1][m].bondOrder.push(_),i[i.length-1][v].bonds.push(m),i[i.length-1][v].bondOrder.push(_))}}if(!e.multimodel)break;for(e.onemol||i.push([]);"$$$$"!==t[o]&&o<t.length;)o++;t.splice(0,++o)}return i}(r,e):[[""]]}function g(t,e){var i=[[]];"string"==typeof t&&(t=JSON.parse(t));for(var r=t.m,s=r[0].a,n=r[0].b,a=r[0].s,o=void 0!==e&&void 0!==e.parseStyle?e.parseStyle:void 0!==a,l=i[i.length-1].length,h=0;h<s.length;h++){var c=s[h],d={};d.id=c.i,d.x=c.x,d.y=c.y,d.z=c.z||0,d.bonds=[],d.bondOrder=[];var u=c.l||"C";d.elem=u[0].toUpperCase()+u.substring(1).toLowerCase(),d.serial=i[i.length-1].length,o&&(d.style=a[c.s||0]),i[i.length-1].push(d)}for(let t=0;t<n.length;t++){let e=n[t],r=e.b+l,s=e.e+l,a=e.o||1,o=i[i.length-1][r],h=i[i.length-1][s];o.bonds.push(s),o.bondOrder.push(a),h.bonds.push(r),h.bondOrder.push(a)}return i}function m(t,e){!function(t,e){const i=e||3.2,r=i*i,s=[];for(let e=0,i=t.length;e<i;e++){t[e].index=e;const i=t[e];i.hetflag||"N"!==i.atom&&"O"!==i.atom||(s.push(i),i.hbondOther=null,i.hbondDistanceSq=Number.POSITIVE_INFINITY)}s.sort((function(t,e){return t.z-e.z}));for(let t=0,e=s.length;t<e;t++){const n=s[t];for(let a=t+1;a<e;a++){const t=s[a],e=t.z-n.z;if(e>i)break;if(t.atom==n.atom)continue;const o=Math.abs(t.y-n.y);if(o>i)continue;const l=Math.abs(t.x-n.x);if(l>i)continue;const h=l*l+o*o+e*e;h>r||t.chain==n.chain&&Math.abs(t.resi-n.resi)<4||(h<n.hbondDistanceSq&&(n.hbondOther=t,n.hbondDistanceSq=h),h<t.hbondDistanceSq&&(t.hbondOther=n,t.hbondDistanceSq=h))}}}(t,e);const i={};let r,s,n,a,o,l;for(r=0,s=t.length;r<s;r++)if(o=t[r],void 0===i[o.chain]&&(i[o.chain]=[]),isFinite(o.hbondDistanceSq)){const t=o.hbondOther;void 0===i[t.chain]&&(i[t.chain]=[]),4===Math.abs(t.resi-o.resi)&&(i[o.chain][o.resi]="h")}for(n in i)for(a=1;a<i[n].length-1;a++){const t=i[n][a-1],e=i[n][a+1];l=i[n][a],"h"==t&&t==e&&l!=t&&(i[n][a]=t)}for(r=0,s=t.length;r<s;r++)o=t[r],isFinite(o.hbondDistanceSq)&&"h"!=i[o.chain][o.resi]&&"h"!==o.ss&&(i[o.chain][o.resi]="maybesheet");for(let e=0,r=t.length;e<r;e++)if(o=t[e],isFinite(o.hbondDistanceSq)&&"maybesheet"==i[o.chain][o.resi]){let t=o.hbondOther,e=i[t.chain][t.resi];"maybesheet"!=e&&"s"!=e||(i[o.chain][o.resi]="s",i[t.chain][t.resi]="s")}for(let t in i){for(let e=1;e<i[t].length-1;e++){const r=i[t][e-1],s=i[t][e+1];l=i[t][e],"s"==r&&r==s&&l!=r&&(i[t][e]=r)}for(let e=0;e<i[t].length;e++){const r=i[t][e];"h"!=r&&"s"!=r||i[t][e-1]!=r&&i[t][e+1]!=r&&delete i[t][e]}}for(r=0,s=t.length;r<s;r++)o=t[r],l=i[o.chain][o.resi],delete o.hbondOther,delete o.hbondDistanceSq,void 0!==l&&"maybesheet"!==l&&(o.ss=l,i[o.chain][o.resi-1]!=l&&(o.ssbegin=!0),i[o.chain][o.resi+1]!=l&&(o.ssend=!0))}function v(t,e,i,r){const s=!i.duplicateAssemblyAtoms,a=e.length;let o=a,l=-1,h=null,c=null;(i.normalizeAssembly||i.wrapAtoms)&&r&&(h=(0,n.conversionMatrix3)(r.a,r.b,r.c,r.alpha,r.beta,r.gamma),c=new n.Matrix3,c.getInverse3(h));let d=function(t){let e=t.clone().applyMatrix3(c);const i=[e.x,e.y,e.z],r=[0,0,0];for(let t=0;t<3;t++){for(;i[t]<-.001;)i[t]+=1,r[t]+=1;for(;i[t]>1.001;)i[t]-=1,r[t]-=1}const s=new n.Vector3(r[0],r[1],r[2]);return s.applyMatrix3(h),s};if(i.normalizeAssembly&&r)for(let i=0;i<t.length;i++){const r=new n.Vector3(0,0,0);for(let s=0;s<a;s++){const a=new n.Vector3(e[s].x,e[s].y,e[s].z);a.applyMatrix4(t[i]),r.add(a)}r.divideScalar(a);const s=d(r);t[i].isNearlyIdentity()&&s.lengthSq()>.001&&(l=i),t[i].translate(s)}if(s){if(t.length>1)for(let i=0;i<e.length;i++){var u=[];for(let r=0;r<t.length;r++)if(!t[r].isNearlyIdentity()){var f=new n.Vector3;f.set(e[i].x,e[i].y,e[i].z),f.applyMatrix4(t[r]),u.push(f)}e[i].symmetries=u}}else{for(let t=0;t<a;t++)e[t].sym=-1;for(let s=0;s<t.length;s++)if(t[s].isNearlyIdentity()||l==s)for(let t=0;t<a;t++)e[t].sym=s;else{let l=new n.Vector3;for(let n=0;n<a;n++){const a=[];for(let t=0;t<e[n].bonds.length;t++)a.push(e[n].bonds[t]+o);if(l.set(e[n].x,e[n].y,e[n].z),l.applyMatrix4(t[s]),i.wrapAtoms&&r){let t=d(l);l.add(t)}const h={};for(const t in e[n])h[t]=e[n][t];h.x=l.x,h.y=l.y,h.z=l.z,h.bonds=a,h.sym=s,h.index=e.length,e.push(h)}o=e.length}if(i.wrapAtoms&&r){let t=new n.Vector3;for(let i=0;i<a;i++){t.set(e[i].x,e[i].y,e[i].z);let r=d(t);t.add(r),e[i].x=t.x,e[i].y=t.y,e[i].z=t.z}}if(l>=0){const i=new n.Vector3;for(let r=0;r<a;r++)i.set(e[r].x,e[r].y,e[r].z),i.applyMatrix4(t[l]),e[r].x=i.x,e[r].y=i.y,e[r].z=i.z}t.length=0}}var _=i(408);const y=new Set(["ABU","ACD","ALA","ALB","ALI","ARG","AR0","ASN","ASP","ASX","BAS","CYS","CYH","CYX","CSS","CSH","GLN","GLU","GLX","GLY","HIS","HIE","HID","HIP","HYP","ILE","ILU","LEU","LYS","MET","PCA","PGA","PHE","PR0","PRO","PRZ","SER","THR","TRP","TYR","VAL","A","1MA","C","5MC","OMC","G","1MG","2MG","M2G","7MG","OMG","YG","I","T","U","+U","H2U","5MU","PSU","ACE","F0R","H2O","HOH","WAT"]);function b(t,e){const i=[],r=[];for(let e=0,s=t.length;e<s;e++){const s=t[e];s.index=e,s.hetflag||!y.has(s.resn)?r.push(s):i.push(s)}(0,u.assignBonds)(r,e),i.sort((function(t,e){return t.chain!==e.chain?t.chain<e.chain?-1:1:t.resi-e.resi}));let s,n=-1,a=-1;for(let t=0,r=i.length;t<r;t++){const r=i[t];r.resi!==n&&(n=r.resi,s||a++,s=!1),r.reschain=a;for(let n=t+1;n<i.length;n++){const t=i[n];if(t.chain!==r.chain||t.resi-r.resi>1)break;(0,_.areConnected)(r,t,e)&&(-1===r.bonds.indexOf(t.index)&&(r.bonds.push(t.index),r.bondOrder.push(1),t.bonds.push(r.index),t.bondOrder.push(1)),r.resi!==t.resi&&(s=!0))}}}function x(t,e={}){const i=[],r=!e.doAssembly,s=i.modelData=[],a=void 0===e.assignBonds||e.assignBonds;function o(t,e){const i=[];let r=0,s=0;for(;s<t.length;){for(;t.substring(s,s+e.length)!==e&&s<t.length;)if("'"===t[s]){for(s++;s<t.length&&"'"!==t[s];)s++;for(;t.substring(s,s+e.length)!==e&&s<t.length;)s++}else if('"'===t[s]){for(s++;s<t.length&&'"'!==t[s];)s++;s++}else s++;i.push(t.substring(r,s)),r=s+=e.length}return i}const l=t.split(/\r?\n|\r/),h=[];let c=!1;for(let t=0;t<l.length;t++){var d=l[t].split("#")[0];if(c?";"===d[0]&&(c=!1):";"===d[0]&&(c=!0),c||""!==d){if(!c&&"_"===(d=d.trim())[0]){const t=d.split(/\s/)[0].indexOf(".");if(t>-1){let e=d.split("");e[t]="_",d=(d=e.join("")).substring(0,t)+"_"+d.substring(t+1)}}h.push(d)}}let u=0;for(;u<h.length;){for(;!h[u].startsWith("data_")||"data_global"===h[u];)u++;u++;const t={};for(;u<h.length&&!h[u].startsWith("data_");)if(void 0===h[u][0])u++;else if("_"===h[u][0]){const e=h[u].split(/\s/)[0].toLowerCase(),i=t[e]=t[e]||[],r=h[u].substring(h[u].indexOf(e)+e.length);if(""===r)if(u++,";"===h[u][0]){let t=h[u].substring(1);for(u++;";"!==h[u];)t=t+"\n"+h[u],u++;i.push(t)}else i.push(h[u]);else i.push(r.trim());u++}else if("loop_"===h[u].substring(0,5)){u++;const e=[];for(;""===h[u]||"_"===h[u][0];){if(""!==h[u]){let i=h[u].split(/\s/)[0].toLowerCase(),r=t[i]=t[i]||[];e.push(r)}u++}let i=0;for(;u<h.length&&"_"!==h[u][0]&&!h[u].startsWith("loop_")&&!h[u].startsWith("data_");){let t=o(h[u]," ");for(let r=0;r<t.length;r++)""!==t[r]&&(e[i].push(t[r]),i=(i+1)%e.length);u++}}else u++;s.push({symmetries:[]}),i.push([]);const e=void 0!==t._atom_site_id?t._atom_site_id.length:t._atom_site_label.length;let a;if(void 0!==t._cell_length_a){const e=parseFloat(t._cell_length_a),i=parseFloat(t._cell_length_b),r=parseFloat(t._cell_length_c),o=parseFloat(t._cell_angle_alpha)||90,l=parseFloat(t._cell_angle_beta)||90,h=parseFloat(t._cell_angle_gamma)||90;a=(0,n.conversionMatrix3)(e,i,r,o,l,h),s[s.length-1].cryst={a:e,b:i,c:r,alpha:o,beta:l,gamma:h}}for(let r=0;r<e;r++){if(void 0!==t._atom_site_group_pdb&&"TER"===t._atom_site_group_pdb[r])continue;const e={};if(void 0!==t._atom_site_cartn_x)e.x=parseFloat(t._atom_site_cartn_x[r]),e.y=parseFloat(t._atom_site_cartn_y[r]),e.z=parseFloat(t._atom_site_cartn_z[r]);else{const i=(f=a,p=parseFloat(t._atom_site_fract_x[r]),g=parseFloat(t._atom_site_fract_y[r]),_=parseFloat(t._atom_site_fract_z[r]),new n.Vector3(p,g,_).applyMatrix3(f));e.x=i.x,e.y=i.y,e.z=i.z}e.chain=t._atom_site_auth_asym_id?t._atom_site_auth_asym_id[r]:t._atom_site_label_asym_id?t._atom_site_label_asym_id[r]:void 0,e.resi=t._atom_site_auth_seq_id?parseInt(t._atom_site_auth_seq_id[r]):t._atom_site_label_seq_id?t._atom_site_label_seq_id[r]:void 0,e.resn=t._atom_site_auth_comp_id?t._atom_site_auth_comp_id[r].trim():t._atom_site_label_comp_id?t._atom_site_label_comp_id[r].trim():void 0,e.atom=t._atom_site_auth_atom_id?t._atom_site_auth_atom_id[r].replace(/"/gm,""):t._atom_site_label_atom_id?t._atom_site_label_atom_id[r].replace(/"/gm,""):void 0,e.hetflag=!t._atom_site_group_pdb||"HETA"===t._atom_site_group_pdb[r]||"HETATM"===t._atom_site_group_pdb[r],t._atom_site_b_iso_or_equiv&&(e.b=parseFloat(t._atom_site_b_iso_or_equiv[r]));let s="X";t._atom_site_type_symbol?s=t._atom_site_type_symbol[r].replace(/\(?\+?\d+.*/,""):t._atom_site_label&&(s=t._atom_site_label[r].split("_")[0].replace(/\(?\d+.*/,"")),e.elem=s[0].toUpperCase()+s.substring(1,2).toLowerCase(),e.bonds=[],e.ss="c",e.serial=r,e.bondOrder=[],e.properties={},i[i.length-1].push(e)}if(void 0!==t._pdbx_struct_oper_list_id&&!r)for(let e=0;e<t._pdbx_struct_oper_list_id.length;e++){const i=parseFloat(t["_pdbx_struct_oper_list_matrix[1][1]"][e]),r=parseFloat(t["_pdbx_struct_oper_list_matrix[1][2]"][e]),a=parseFloat(t["_pdbx_struct_oper_list_matrix[1][3]"][e]),o=parseFloat(t["_pdbx_struct_oper_list_vector[1]"][e]),l=parseFloat(t["_pdbx_struct_oper_list_matrix[2][1]"][e]),h=parseFloat(t["_pdbx_struct_oper_list_matrix[2][2]"][e]),c=parseFloat(t["_pdbx_struct_oper_list_matrix[2][3]"][e]),d=parseFloat(t["_pdbx_struct_oper_list_vector[2]"][e]),u=parseFloat(t["_pdbx_struct_oper_list_matrix[3][1]"][e]),f=parseFloat(t["_pdbx_struct_oper_list_matrix[3][2]"][e]),p=parseFloat(t["_pdbx_struct_oper_list_matrix[3][3]"][e]),g=parseFloat(t["_pdbx_struct_oper_list_vector[3]"][e]),m=new n.Matrix4(i,r,a,o,l,h,c,d,u,f,p,g);s[s.length-1].symmetries.push(m)}const l=function(t){const e=t.match("-"),i=(t=t.replace(/[-xyz]/g,"")).split("/");let r,s;return s=void 0===i[1]?1:parseInt(i[1]),r=""===i[0]?1:parseInt(i[0]),r/s*(e?-1:1)};if(void 0!==t._symmetry_equiv_pos_as_xyz&&!r)for(let e=0;e<t._symmetry_equiv_pos_as_xyz.length;e++){const i=t._symmetry_equiv_pos_as_xyz[e].replace(/["' ]/g,"").split(",").map((function(t){return t.replace(/-/g,"+-")}));let r=new n.Matrix4(0,0,0,0,0,0,0,0,0,0,0,0,0,0,0,1);for(let t=0;t<3;t++){const e=i[t].split("+");for(let i=0;i<e.length;i++){const s=e[i];if(""===s)continue;const n=l(s);s.match("x")?r.elements[t+0]=n:s.match("y")?r.elements[t+4]=n:s.match("z")?r.elements[t+8]=n:r.elements[t+12]=n}}const o=a.getMatrix4(),h=(new n.Matrix4).getInverse(o,!0);r=(new n.Matrix4).multiplyMatrices(r,h),r=(new n.Matrix4).multiplyMatrices(o,r),s[s.length-1].symmetries.push(r)}}var f,p,g,_;for(let t=0;t<i.length;t++)!a||e.duplicateAssemblyAtoms&&!e.dontConnectDuplicatedAtoms||b(i[t],e),m(i[t],e.hbondCutoff),v(s[t].symmetries,i[t],e,s[t].cryst),e.duplicateAssemblyAtoms&&!e.dontConnectDuplicatedAtoms&&a&&b(i[t],e);return i}let w={"C.1":"C",C1:"C","C.2":"C",C2:"C","C.3":"C",C3:"C","C.ar":"C",Car:"C","C.cat":"C",Ccat:"C","H.spc":"H",Hspc:"H","H.t3p":"H",Ht3p:"H","N.1":"N",N1:"N","N.2":"N",N2:"N","N.3":"N",N3:"N","N.4":"N",N4:"N","N.am":"N",Nam:"N","N.ar":"N",Nar:"N","N.p13":"N",Np13:"N","O.2":"O",O2:"O","O.3":"O",O3:"O","O.co2":"O",Oco2:"O","O.spc":"O",Ospc:"O","O.t3p":"O",Ot3p:"O","P.3":"P",P3:"P","S.2":"S",S2:"S","S.3":"S",S3:"S","S.o":"S",So:"S","S.o2":"S",So2:"S"};function A(t,e){var i=[[]],r=!1;void 0!==e.keepH&&(r=!e.keepH);var s=t.search(/@<TRIPOS>MOLECULE/),n=t.search(/@<TRIPOS>ATOM/);if(-1==s||-1==n)return i;for(var a=t.substring(s).split(/\r?\n|\r/);a.length>0;){var o=[],l=a[2].replace(/^\s+/,"").replace(/\s+/g," ").split(" "),h=parseInt(l[0]),c=0;l.length>1&&(c=parseInt(l[1]));var d,u=4;for(d=3;d<a.length;d++)if("@<TRIPOS>ATOM"==a[d]){u=d+1;break}var f=i[i.length-1].length,p=f+h;for(d=f;d<p;d++){var g={},m=(l=a[u++].replace(/^\s+/,"").replace(/\s+/g," ").split(" "))[5];if(m=void 0!==w[m]?w[m]:(m=m.split(".")[0])[0].toUpperCase()+m.substring(1).toLowerCase(),g.atom=l[1],g.elem=m,"H"==g.elem&&r);else{var v=i[i.length-1].length,_=parseInt(l[0]);g.serial=_,g.x=parseFloat(l[2]),g.y=parseFloat(l[3]),g.z=parseFloat(l[4]),g.atom=l[5];var y=parseFloat(l[8]);g.index=v,g.bonds=[],g.bondOrder=[],g.properties={charge:y,partialCharge:y},o[_]=v,i[i.length-1].push(g)}}for(var b=!1;u<a.length;)if("@<TRIPOS>BOND"==a[u++]){b=!0;break}if(b&&c)for(d=0;d<c;d++){l=a[u++].replace(/^\s+/,"").replace(/\s+/g," ").split(" ");var x=parseInt(l[1]),A=i[i.length-1][o[x]],C=parseInt(l[2]),S=i[i.length-1][o[C]],M=parseInt(l[3]);isNaN(M)&&(M=1),void 0!==A&&void 0!==S&&(A.bonds.push(o[C]),A.bondOrder.push(M),S.bonds.push(o[x]),S.bondOrder.push(M))}if(!e.multimodel)break;e.onemol||i.push([]),a.splice(0,u),t=a.join("\n")}return i}var C=i(40);function S(t,e){let i=t.replace(/ /g,"");return i.length>0&&"H"===i[0]&&"Hg"!==i&&"He"!==i&&"Hf"!==i&&"Hs"!==i&&"Ho"!==i&&(i="H"),i.length>1&&(i=i[0].toUpperCase()+i.substring(1).toLowerCase(),void 0===C.bondTable[i]?i=i[0]:e&&("Ca"===i||"Cd"===i)&&(i="C")),i}function M(t){for(const e in t)return!1;return!0}function z(t,e,i){const r=[],s=void 0===e.assignBonds||e.assignBonds,a=!e.keepH,o=!!e.noSecondaryStructure,l=!e.noComputeSecondaryStructure,h=!e.doAssembly,c=e.altLoc?e.altLoc:"A",d={symmetries:[],cryst:void 0};let u,f=[];const p=[];let g;const _={};for(let e=0;e<t.length;e++){g=t[e].replace(/^\s*/,"");const s=g.substring(0,6);let o,l,m;if(0===s.indexOf("END")){if(f=t.slice(e+1),"END"===s)for(const t in i)i.hasOwnProperty(t)&&delete i[t];break}if("ATOM  "===s||"HETATM"===s){let t,e,i,n,o,l,h,d,f,m,v,_;if(v=g.substring(16,17)," "!==v&&v!==c&&"*"!==c)continue;if(m=parseInt(g.substring(6,11)),u=g.substring(12,16).replace(/ /g,""),t=g.substring(17,20).replace(/ /g,""),e=g.substring(21,22),i=parseInt(g.substring(22,26)),n=g.substring(26,27),o=parseFloat(g.substring(30,38)),l=parseFloat(g.substring(38,46)),h=parseFloat(g.substring(46,54)),_=parseFloat(g.substring(60,68)),f=g.substring(76,78).replace(/ /g,""),f=""===f||void 0===C.bondTable[f]?S(g.substring(12,14),"A"==g[0]):f[0].toUpperCase()+f.substring(1).toLowerCase(),"H"===f&&a)continue;d="H"==s[0],p[m]=r.length,r.push({resn:t,x:o,y:l,z:h,elem:f,hetflag:d,altLoc:v,chain:e,resi:i,icode:n,rescode:i+(" "!==n?"^"+n:""),serial:m,atom:u,bonds:[],ss:"c",bondOrder:[],properties:{},b:_,pdbline:g})}else if("SHEET "===s){o=g.substring(21,22),l=parseInt(g.substring(22,26)),m=parseInt(g.substring(33,37)),o in i||(i[o]={}),i[o][l]="s1";for(let t=l+1;t<m;t++)i[o][t]="s";i[o][m]="s2"}else if("CONECT"===s){const t=p[parseInt(g.substring(6,11))],e=r[t],i=[11,16,21,26];for(let s=0;s<4;s++){const n=p[parseInt(g.substring(i[s],i[s]+5))];let a=t+":"+n;const o=r[n];if(void 0!==e&&void 0!==o)if(_[a]){_[a]+=1;for(let t=0;t<e.bonds.length;t++)if(e.bonds[t]==n){const i=_[a];e.bondOrder[t]=i>=4?1:i}}else _[a]=1,0!=e.bonds.length&&e.bonds[e.bonds.length-1]===n||(e.bonds.push(n),e.bondOrder.push(1))}}else if("HELIX "===s){o=g.substring(19,20),l=parseInt(g.substring(21,25)),m=parseInt(g.substring(33,37)),o in i||(i[o]={}),i[o][l]="h1";for(let t=l+1;t<m;t++)i[o][t]="h";i[o][m]="h2"}else if(h||"REMARK"!==s||"BIOMT"!==g.substring(13,18)){if("CRYST1"===s){let t,e,i,r,s,n;t=parseFloat(g.substring(7,15)),e=parseFloat(g.substring(16,24)),i=parseFloat(g.substring(25,33)),r=parseFloat(g.substring(34,40)),s=parseFloat(g.substring(41,47)),n=parseFloat(g.substring(48,54)),d.cryst={a:t,b:e,c:i,alpha:r,beta:s,gamma:n}}else if("ANISOU"===s){const t=r[p[parseInt(g.substring(6,11))]];if(t){const e=g.substring(30).trim().split(/\s+/),i={u11:parseInt(e[0]),u22:parseInt(e[1]),u33:parseInt(e[2]),u12:parseInt(e[3]),u13:parseInt(e[4]),u23:parseInt(e[5])};t.uMat=i}}}else{let i,r=new n.Matrix4;for(i=1;i<=3;i++)if(g=t[e].replace(/^\s*/,""),parseInt(g.substring(18,19))==i)r.elements[i-1]=parseFloat(g.substring(23,33)),r.elements[i-1+4]=parseFloat(g.substring(33,43)),r.elements[i-1+8]=parseFloat(g.substring(43,53)),r.elements[i-1+12]=parseFloat(g.substring(53)),e++;else for(;"BIOMT"===g.substring(13,18);)e++,g=t[e].replace(/^\s*/,"");r.elements[3]=0,r.elements[7]=0,r.elements[11]=0,r.elements[15]=1,d.symmetries.push(r),e--}}if(function(t,e){for(let i=0,r=t.length;i<r;i++){const r=t[i];for(let i=0;i<r.bonds.length;i++){const s=t[r.bonds[i]],n=e[r.serial];s&&n&&s.bonds.indexOf(n)<0&&(s.bonds.push(n),s.bondOrder.push(r.bondOrder[i]))}}}(r,p),s&&b(r,e),h||v(d.symmetries,r,e,d.cryst),l&&!o&&m(r,e.hbondCutoff),!M(i))for(let t=0;t<r.length;t++){const e=r[t];if(void 0!==e&&(e.chain in i&&e.resi in i[e.chain])){const t=i[e.chain][e.resi];e.ss=t[0],t.length>1&&("1"==t[1]?e.ssbegin=!0:"2"==t[1]&&(e.ssend=!0))}}return[r,d,f]}function T(t,e){e=e||{};var i=[],r={};i.modelData=[];for(var s=t.split(/\r?\n|\r/);s.length>0;){var n=z(s,e,r),a=n[0],o=n[1];if(s=n[2],0!=a.length){if(e.multimodel&&e.onemol&&i.length>0)for(var l=i[0].length,h=0;h<a.length;h++){var c=a[h];c.index=h;for(var d=0;d<c.bonds.length;d++)c.bonds[d]+=l;i[0].push(c)}else i.modelData.push(o),i.push(a);if(!e.multimodel)break}}return i}function E(t,e){var i=[[]],r=!e.noSecondaryStructure;i.modelData=[{symmetries:[]}];var s,n=[],a=t.split(/\r?\n|\r/);for(let t=0;t<a.length;t++){var o=(s=a[t].replace(/^\s*/,"")).substring(0,6);if(0==o.indexOf("END")){if(e.multimodel){e.onemol||i.push([]);continue}break}if("ATOM  "==o||"HETATM"==o){var l;let t=parseInt(s.substring(6,11)),e=s.substring(12,16).replace(/ /g,""),r=s.substring(17,20).trim(),a=s.substring(21,22),o=parseInt(s.substring(22,26));var h=s.substring(30).trim().split(/\s+/),c=parseFloat(h[0]),d=parseFloat(h[1]),u=parseFloat(h[2]),f=parseFloat(h[3]),p=parseFloat(h[4]),g=e[0];e.length>1&&e[1].toUpperCase()!=e[1]&&(g=e.substring(0,2)),l="H"==s[0],n[t]=i[i.length-1].length,i[i.length-1].push({resn:r,x:c,y:d,z:u,elem:g,hetflag:l,chain:a,resi:o,serial:t,atom:e,bonds:[],ss:"c",bondOrder:[],properties:{charge:f,partialCharge:f,radius:p},pdbline:s})}else if("CONECT"==o){var v=parseInt(s.substring(6,11)),_=i[i.length-1][n[v]];for(let t=0;t<4;t++){var y=parseInt(s.substring([11,16,21,26][t],[11,16,21,26][t]+5)),x=i[i.length-1][n[y]];void 0!==_&&void 0!==x&&(_.bonds.push(n[y]),_.bondOrder.push(1))}}}for(let t=0;t<i.length;t++)b(i[t],e),r&&m(i[t],e.hbondCutoff);return i}var L=i(864),F=i(471),I=function(t){return String.fromCharCode.apply(null,t).replace(/\0/g,"")},O=function(t){return 0==t||2==t||4==t?"h":3==t?"s":"c"};let D=new Set(["D-SACCHARIDE","D-SACCHARIDE 1,4 AND 1,4 LINKING","D-SACCHARIDE 1,4 AND 1,6 LINKING","L-SACCHARIDE","L-SACCHARIDE 1,4 AND 1,4 LINKING","L-SACCHARIDE 1,4 AND 1,6 LINKING","NON-POLYMER","OTHER","PEPTIDE-LIKE","SACCHARIDE"]);function k(t,e){var i=!e.keepH,r=e.altLoc?e.altLoc:"A",s=!!e.noSecondaryStructure,a=!e.noComputeSecondaryStructure,o=!e.doAssembly,l=e.assemblyIndex?e.assemblyIndex:0;t="string"==typeof t?(0,L.base64ToArray)(t):new Uint8Array(t);var h,c,d,u,f,p,g=F.decode(t),_=[[]],y=_.modelData=[],b=0,x=0,w=0,A=g.secStructList,C=g.bFactorList,S=g.altLocList,M=g.occupancyList,z=g.bondAtomList,T=g.bondOrderList,E=g.numModels;if(0==E)return _;e.multimodel||(E=1);var k=[];if(!o&&g.bioAssemblyList&&g.bioAssemblyList.length>0){var R=g.bioAssemblyList[l].transformList;for(h=0,p=R.length;h<p;h++){var P=new n.Matrix4(R[h].matrix);P.transpose(),k.push(P)}}var U=null;if(g.unitCell){var B=g.unitCell;U={a:B[0],b:B[1],c:B[2],alpha:B[3],beta:B[4],gamma:B[5]}}let N=[];g.entityList.forEach((t=>{t.chainIndexList.forEach((e=>{N[e]="polymer"==t.type}))}));var G=0;for(f=0;f<E;f++){var V=g.chainsPerModel[f],j=_[_.length-1],H=[];for(y.push({symmetries:k,cryst:U}),h=0;h<V;++h){var W=g.groupsPerChain[b],q=I(g.chainIdList.subarray(4*b,4*b+4));g.chainNameList&&(q=I(g.chainNameList.subarray(4*b,4*b+4)));var Y=x,Z="";for(c=0;c<W;++c){var X=g.groupList[g.groupTypeList[x]],K=X.atomNameList.length,Q=0,$=!1,J=!1;if(A){Q=A[x];var tt=O(Q);0!=x&&tt==Z||($=!0),Z=tt;var et=x+1;(et>=A.length||O(A[et]!=tt))&&(J=!0)}var it=g.groupIdList[x],rt=X.groupName;let t=X.chemCompType;var st=w;let e=D.has(t)||!N[b];for(d=0;d<K;++d){var nt=X.elementList[d];if(i&&"H"==nt)w+=1;else{var at="";C&&(at=C[w]);var ot="";S&&S[w]&&(ot=String.fromCharCode(S[w]));var lt="";if(M&&(lt=M[w]),""==ot||ot==r||"*"==r){var ht=g.atomIdList[w],ct=X.atomNameList[d],dt=0;X.atomChargeList&&(dt=X.atomChargeList[d]);var ut=g.xCoordList[w],ft=g.yCoordList[w],pt=g.zCoordList[w];H[w]=j.length,j.push({resn:rt,x:ut,y:ft,z:pt,elem:nt,hetflag:e,chain:q,resi:it,icode:ot,rescode:it+(" "!=ot?"^"+ot:""),serial:ht,altLoc:ot,index:w,atom:ct,bonds:[],ss:O(Q),ssbegin:$,ssend:J,bondOrder:[],properties:{charge:dt,occupancy:lt},b:at}),w+=1}else w+=1}}var gt=X.bondAtomList;for(d=0,u=X.bondOrderList.length;d<u;++d){var mt=st+gt[2*d],vt=st+gt[2*d+1],_t=X.bondOrderList[d],yt=H[mt],bt=H[vt],xt=j[yt],wt=j[bt];xt&&wt&&(xt.bonds.push(bt),xt.bondOrder.push(_t),wt.bonds.push(yt),wt.bondOrder.push(_t))}x+=1}for(x=Y,c=0;c<W;++c)x+=1;b+=1}if(z)for(let t=G,e=z.length;t<e;t+=2){let e=z[t],i=z[t+1],r=T?T[t/2]:1;if(e>=w){G=t;break}let s=H[e],n=H[i],a=j[s],o=j[n];a&&o&&(a.bonds.push(n),a.bondOrder.push(r),o.bonds.push(s),o.bondOrder.push(r))}e.multimodel&&(e.onemol||_.push([]))}if(!o)for(let t=0;t<_.length;t++)v(y[t].symmetries,_[t],e,y[t].cryst);return a&&!s&&m(_,e.hbondCutoff),_}function R(t){var e,i=[],r=0,s=t.split(/\r?\n|\r/);if(!(s.length>0&&s[0].includes("VERSION")))return[];var n=s.filter((function(t){return t.includes("POINTERS")||t.includes("ATOM_NAME")||t.includes("CHARGE")||t.includes("RADII")||t.includes("BONDS_INC_HYDROGEN")||t.includes("BONDS_WITHOUT_HYDROGEN")})),a=c("POINTERS");if(-1==a)return[];var o=d(a),l=parseInt(s[a+1].slice(0,o[1]));if(isNaN(l)||l<=0)return[];if(-1==(a=c("ATOM_NAME")))return[];var h=(o=d(a))[0];for(let t=0;t<l/o[0];t++){t==parseInt(l/o[0])&&(h=l%o[0]);for(let t=0;t<h;t++){let e={},n={charge:"",radii:""};e.serial=r,e.x=0,e.y=0,e.z=0,e.atom=s[a+1].slice(o[1]*t,o[1]*(t+1)).trim(),e.elem=s[a+1].slice(o[1]*t,o[1]*t+1).trim(),e.properties=n,e.bonds=[],e.bondOrder=[],i.push(e),r++}a++}if(-1!=(a=c("CHARGE"))){r=0,h=(o=d(a))[0];for(let t=0;t<l/o[0];t++){t==parseInt(l/o[0])&&(h=l%o[0]);for(let t=0;t<h;t++)i[r].properties.charge=parseFloat(s[a+1].slice(o[1]*t,o[1]*(t+1))),r++;a++}}if(-1!=(a=c("RADII"))){r=0,h=(o=d(a))[0];for(let t=0;t<l/o[0];t++){t==parseInt(l/o[0])&&(h=l%o[0]);for(let t=0;t<h;t++)i[r].properties.radii=parseFloat(s[a+1].slice(o[1]*t,o[1]*(t+1))),r++;a++}}if(-1!=(a=c("BONDS_WITHOUT_HYDROGEN")))for(r=0,h=(o=d(a))[0],a+=1;!s[a].match(/^%FLAG/);){s[a+1].match(/^%FLAG/)&&(h=l%o[0]);for(let t=0;t<h;t++)r%3==0?e=parseInt(s[a].slice(o[1]*t,o[1]*(t+1))/3):r%3==1&&i[e].bonds.push(parseInt(s[a].slice(o[1]*t,o[1]*(t+1))/3)),r++;a++}if(-1!=(a=c("BONDS_INC_HYDROGEN")))for(r=0,h=(o=d(a))[0],a+=1;!s[a].match(/^%FLAG/);){s[a+1].match(/^%FLAG/)&&(h=l%o[0]);for(let t=0;t<h;t++)r%3==0?e=parseInt(s[a].slice(o[1]*t,o[1]*(t+1))/3):r%3==1&&i[e].bonds.push(parseInt(s[a].slice(o[1]*t,o[1]*(t+1))/3)),r++;a++}function c(t){var e=s.indexOf(n.filter((function(e){return e.includes(t)}))[0]);if(Number.isInteger(e)&&e>0){for(;!s[e].includes("FORMAT");)e++;return e}return-1}function d(t){var e=s[t].match(/\((\d*)\S*/),i=s[t].match(/[a-zA-Z](\d*)\)\s*/);return null==i&&(i=s[t].match(/[a-zA-Z](\d*)\.\d*\)\s*/)),[e[1],i[1]]}return[i]}function P(t,e){const i=[],r=t.split(/\r?\n|\r/);for(;r.length>0;){const t=parseInt(r[1]);if(r.length<3||isNaN(t)||t<=0||r.length<t+3)break;const e=[];i.push(e);let s=2;const n=e.length,a=n+t;for(let t=n;t<a;t++){const i=r[s++],n={};n.serial=t,n.atom=i.slice(10,15).trim(),n.elem=S(n.atom,!0),n.x=10*parseFloat(i.slice(20,28)),n.y=10*parseFloat(i.slice(28,36)),n.z=10*parseFloat(i.slice(36,44)),n.resi=parseInt(i.slice(0,5)),n.resn=i.slice(5,10).trim(),n.bonds=[],n.bondOrder=[],n.properties={},i.length>44&&(n.dx=10*parseFloat(i.slice(44,52)),n.dy=10*parseFloat(i.slice(52,60)),n.dz=10*parseFloat(i.slice(60,68))),e[t]=n}if(r.length<=s+3){const t=r[s++].trim().split(/\s+/);if(3===t.length){for(let e=0;e<3;e++)t[e]=(10*parseFloat(t[e])).toString();i.box=t}}r.splice(0,++s)}for(let t=0;t<i.length;t++)b(i[t],e);return i}const U={id:"serial",type:"atom",element:"elem",q:"charge",radius:"radius",x:"x",xu:"x",xs:"x",xsu:"x",y:"y",yu:"y",ys:"y",ysu:"y",z:"z",zu:"z",zs:"z",zsu:"z"};function B(t,e){const i=[],r=t.split(/\r?\n|\r/);let s=0,n=0,a=0;for(;a<r.length-9;){for(var o=a;o<r.length;o++)if(r[o].match(/ITEM: NUMBER OF ATOMS/)&&(n=parseInt(r[o+1])),r[o].match(/ITEM: ATOMS/)){s=o+1;break}const t=r[s-1].replace("ITEM: ATOMS ","").split(" ");i.push([]);for(let e=s;e<s+n;e++){const n={},a={},o=r[e].split(" ");for(let e=0;e<o.length;e++){const i=U[t[e]];void 0!==i&&("serial"===i?n[i]=parseInt(o[e]):"x"===i||"y"===i||"z"===i?n[i]=parseFloat(o[e]):"charge"===i||"radius"===i?a[i]=parseFloat(o[e]):n[i]=o[e]),n.properties=a,n.bonds=[],n.bondOrder=[]}i[i.length-1][e-s]=n}a=s+n-1}if(e.assignBonds)for(let t=0;t<i.length;t++)(0,u.assignBonds)(i[t],e);return i}var N;!function(t){let e,i;!function(t){t[t.Int8=1]="Int8",t[t.Int16=2]="Int16",t[t.Int32=3]="Int32",t[t.Uint8=4]="Uint8",t[t.Uint16=5]="Uint16",t[t.Uint32=6]="Uint32"}(e=t.IntDataType||(t.IntDataType={})),function(t){t[t.Float32=32]="Float32",t[t.Float64=33]="Float64"}(i=t.FloatDataType||(t.FloatDataType={})),t.getDataType=function(e){let i;return i=e instanceof Int8Array?t.IntDataType.Int8:e instanceof Int16Array?t.IntDataType.Int16:e instanceof Int32Array?t.IntDataType.Int32:e instanceof Uint8Array?t.IntDataType.Uint8:e instanceof Uint16Array?t.IntDataType.Uint16:e instanceof Uint32Array?t.IntDataType.Uint32:e instanceof Float32Array?t.FloatDataType.Float32:e instanceof Float64Array?t.FloatDataType.Float64:t.IntDataType.Int32,i},t.isSignedIntegerDataType=function(t){if(t instanceof Int8Array||t instanceof Int16Array||t instanceof Int32Array)return!0;for(let e=0,i=t.length;e<i;e++)if(e<0)return!1;return!0}}(N||(N={}));const G=13330===new Uint16Array(new Uint8Array([18,52]).buffer)[0];function V(t){let e=t.data;for(let i=t.encoding.length-1;i>=0;i--)e=j(e,t.encoding[i]);return e}function j(t,e){switch(e.kind){case"ByteArray":switch(e.type){case N.IntDataType.Uint8:return t;case N.IntDataType.Int8:return function(t){return new Int8Array(t.buffer,t.byteOffset)}(t);case N.IntDataType.Int16:return function(t){return q(t,2,Int16Array)}(t);case N.IntDataType.Uint16:return function(t){return q(t,2,Uint16Array)}(t);case N.IntDataType.Int32:return function(t){return q(t,4,Int32Array)}(t);case N.IntDataType.Uint32:return function(t){return q(t,4,Uint32Array)}(t);case N.FloatDataType.Float32:return function(t){return q(t,4,Float32Array)}(t);case N.FloatDataType.Float64:return function(t){return q(t,8,Float64Array)}(t);default:throw new Error("unreachable")}case"FixedPoint":return function(t,e){const i=t.length,r=W(e.srcType,i),s=1/e.factor;for(let e=0;e<i;e++)r[e]=s*t[e];return r}(t,e);case"IntervalQuantization":return function(t,e){const i=t.length,r=W(e.srcType,i),s=(e.max-e.min)/(e.numSteps-1),n=e.min;for(let e=0;e<i;e++)r[e]=n+s*t[e];return r}(t,e);case"RunLength":return function(t,e){const i=H(e.srcType,e.srcSize);let r=0;for(let e=0,s=t.length;e<s;e+=2){const s=t[e],n=t[e+1];for(let t=0;t<n;++t)i[r++]=s}return i}(t,e);case"Delta":return function(t,e){const i=t.length,r=H(e.srcType,i);if(!i)return t;r[0]=t[0]+(0|e.origin);for(let e=1;e<i;++e)r[e]=t[e]+r[e-1];return r}(t,e);case"IntegerPacking":return function(t,e){return t.length===e.srcSize?t:e.isUnsigned?function(t,e){const i=1===e.byteCount?255:65535,r=t.length,s=new Int32Array(e.srcSize);let n=0,a=0;for(;n<r;){let e=0,r=t[n];for(;r===i;)e+=r,n++,r=t[n];e+=r,s[a]=e,n++,a++}return s}(t,e):function(t,e){const i=1===e.byteCount?127:32767,r=-i-1,s=t.length,n=new Int32Array(e.srcSize);let a=0,o=0;for(;a<s;){let e=0,s=t[a];for(;s===i||s===r;)e+=s,a++,s=t[a];e+=s,n[o]=e,a++,o++}return n}(t,e)}(t,e);case"StringArray":return function(t,e){const i=V({encoding:e.offsetEncoding,data:e.offsets}),r=V({encoding:e.dataEncoding,data:t}),s=e.stringData,n=new Array(i.length);n[0]="";for(let t=1,e=i.length;t<e;t++)n[t]=s.substring(i[t-1],i[t]);let a=0;const o=new Array(r.length);for(let t=0,e=r.length;t<e;t++)o[a++]=n[r[t]+1];return o}(t,e)}}function H(t,e){switch(t){case N.IntDataType.Int8:return new Int8Array(e);case N.IntDataType.Int16:return new Int16Array(e);case N.IntDataType.Int32:return new Int32Array(e);case N.IntDataType.Uint8:return new Uint8Array(e);case N.IntDataType.Uint16:return new Uint16Array(e);case N.IntDataType.Uint32:return new Uint32Array(e);default:return new Int32Array(e)}}function W(t,e){switch(t){case N.FloatDataType.Float32:return new Float32Array(e);case N.FloatDataType.Float64:default:return new Float64Array(e)}}function q(t,e,i){return 0==t.byteOffset&&t.byteLength==t.buffer.byteLength||(t=new Uint8Array(t)),new i(G?t.buffer:function(t,e){const i=new ArrayBuffer(t.length),r=new Uint8Array(i);for(let i=0,s=t.length;i<s;i+=e)for(let s=0;s<e;s++)r[i+e-s-1]=t[i+s];return i}(t,e))}function Y(t){const e=Object.create(null),i=Object.create(null);for(const i of t.columns)e[i.name]=i;return{rowCount:t.rowCount,name:t.name.substring(1),fieldNames:t.columns.map((t=>t.name)),getField(t){const r=e[t];if(r)return i[t]||(i[t]=V(r.data)),i[t]}}}var Z=i(471);class Connectivity{constructor(t){if(this.C={},t){let e=t.getField("comp_id"),i=t.getField("atom_id_1"),r=t.getField("atom_id_2"),s=t.getField("value_order");for(let t=0;t<e.length;t++){let n=e[t],a=i[t],o=r[t],l=s[t],h=1;"doub"==l?h=2:"trip"==l&&(h=3),null==this.C[n]&&(this.C[n]={}),null==this.C[n][a]&&(this.C[n][a]={}),null==this.C[n][o]&&(this.C[n][o]={}),this.C[n][a][o]=h,this.C[n][o][a]=h}}}order(t,e,i){return void 0!==this.C[t]&&void 0!==this.C[t][e]&&void 0!==this.C[t][e][i]?this.C[t][e][i]:0}}class StructConn{constructor(t){if(this.C=[],t){let e=t.getField("conn_type_id"),i=t.getField("ptnr1_label_asym_id"),r=t.getField("ptnr1_label_seq_id"),s=t.getField("ptnr1_label_comp_id"),n=t.getField("ptnr1_label_atom_id"),a=t.getField("ptnr2_label_asym_id"),o=t.getField("ptnr2_label_seq_id"),l=t.getField("ptnr2_label_comp_id"),h=t.getField("ptnr2_label_atom_id"),c=t.getField("pdbx_value_order");for(let t=0;t<e.length;t++)if("disulf"==e[t]||"covale"==e[t]){let e=c?""==c[t]?1:parseInt(c[t]):1;this.C.push([[i[t],r[t],s[t],n[t]],[a[t],o[t],l[t],h[t]],e])}}}}class Residues{constructor(){this.R={}}add(t){null==this.R[t.lchain]&&(this.R[t.lchain]={}),null==this.R[t.lchain][t.lresi]&&(this.R[t.lchain][t.lresi]={}),null==this.R[t.lchain][t.lresi][t.lresn]&&(this.R[t.lchain][t.lresi][t.lresn]=[]),this.R[t.lchain][t.lresi][t.lresn].push(t),this.R[t.lchain][t.lresi][t.lresn][t.atom]=t}geta([t,e,i,r]){if(void 0!==this.R[t]&&void 0!==this.R[t][e]&&void 0!==this.R[t][e][i])return this.R[t][e][i][r]}setBonds(t,e){for(let e in this.R)for(let i in this.R[e])for(let r in this.R[e][i]){let s=this.R[e][i][r];for(let e=0;e<s.length;e++)for(let i=e+1;i<s.length;i++){let n=s[e],a=s[i],o=t.order(r,n.atom,a.atom);n.altLoc!=a.altLoc&&""!=n.altLoc&&""!=a.altLoc&&(o=0),o>0&&(n.bonds.push(a.index),a.bonds.push(n.index),n.bondOrder.push(o),a.bondOrder.push(o))}}for(let t of e.C){let e=t[0],i=t[1],r=t[2],s=this.geta(e),n=this.geta(i);null!=s&&null!=n&&(s.bonds.push(n.index),n.bonds.push(s.index),s.bondOrder.push(r),n.bondOrder.push(r))}}}function X(t,e){var i=!e.keepH,r=e.altLoc?e.altLoc:"A",s=!e.noComputeSecondaryStructure;const a=!e.doAssembly,o=void 0===e.assignBonds||e.assignBonds;if("string"==typeof t)try{t=(0,L.base64ToArray)(t)}catch(e){t=(new TextEncoder).encode(t)}else t=new Uint8Array(t);var l=Z.decodeMsgpack(t);31==l&&(t=(0,L.inflateString)(t,!1),l=Z.decodeMsgpack(t));var h=[],c=h.modelData=[],d=l.dataBlocks.length;if(0==d)return h;e.multimodel||(d=1);for(let t=0;t<d;t++){let s=h.length;const o=[];c.push({symmetries:[]}),h.push([]);const u=l.dataBlocks[t],f=Object.create(null);for(const t of u.categories)f[t.name.substr(1)]=Y(t);let p={},g=f.struct_conf;if(g){let t=g.getField("conf_type_id"),e=g.getField("beg_label_asym_id"),i=g.getField("beg_label_seq_id"),r=g.getField("end_label_seq_id");for(let s=0;s<t.length;s++)if(t[s].startsWith("H")){let t=e[s],n=i[s],a=r[s];t in p||(p[t]={}),p[t][n]="h1";for(let e=n+1;e<a;e++)p[t][e]="h";p[t][a]="h2"}}let m=f.struct_sheet_range;if(m){let t=m.getField("id"),e=m.getField("beg_label_asym_id"),i=m.getField("beg_label_seq_id"),r=m.getField("end_label_seq_id");for(let s=0;s<t.length;s++){let t=e[s],n=i[s],a=r[s];t in p||(p[t]={}),p[t][n]="s1";for(let e=n+1;e<a;e++)p[t][e]="s";p[t][a]="s2"}}let v=f.pdbx_struct_oper_list,_=v.getField("id");if(_&&!a){let t=v.getField("matrix[1][1]"),e=v.getField("matrix[1][2]"),i=v.getField("matrix[1][3]"),r=v.getField("matrix[2][1]"),s=v.getField("matrix[2][2]"),a=v.getField("matrix[2][3]"),o=v.getField("matrix[3][1]"),l=v.getField("matrix[3][2]"),h=v.getField("matrix[3][3]"),d=v.getField("vector[1]"),u=v.getField("vector[2]"),f=v.getField("vector[3]");for(let p=0;p<_.length;p++){const g=new n.Matrix4(t[p],e[p],i[p],d[p],r[p],s[p],a[p],u[p],o[p],l[p],h[p],f[p]);c[c.length-1].symmetries.push(g)}}let y=new Connectivity(f.chem_comp_bond),b=new Residues,x=new StructConn(f.struct_conn),w=f.atom_site,A=w.rowCount,C=w.getField("group_PDB"),S=w.getField("Cartn_x"),z=w.getField("Cartn_y"),T=w.getField("Cartn_z"),E=w.getField("auth_asym_id"),L=w.getField("label_asym_id"),F=w.getField("auth_seq_id"),I=w.getField("label_seq_id"),O=w.getField("auth_comp_id"),D=w.getField("label_comp_id"),k=w.getField("auth_atom_id"),R=w.getField("label_atom_id"),P=w.getField("type_symbol"),U=w.getField("B_iso_or_equiv"),B=w.getField("id"),N=w.getField("label_alt_id"),G=w.getField("pdbx_PDB_model_num"),V=G?G[0]:0;for(let t=0;t<A;t++){if(void 0!==C&&"TER"===C[t])continue;if(G&&G[t]!=V){if(V=G[t],!e.multimodel)break;e.onemol||(h.push([]),c.push(c[c.length-1]),V=G[t],b.setBonds(y,x),b=new Residues)}const s={};s.x=S[t],s.y=z[t],s.z=T[t],s.chain=E?E[t]:L?L[t]:void 0,s.lchain=L?L[t]:void 0,s.resi=F?F[t]:I?I[t]:void 0,s.lresi=I?I[t]:void 0,s.resn=O?O[t].trim():D?D[t].trim():void 0,s.lresn=D?D[t].trim():void 0,s.atom=k?k[t].replace(/"/gm,""):R?R[t].replace(/"/gm,""):void 0,s.icode=N?N[t]:void 0,s.altLoc=s.icode,s.hetflag=!C||"HETA"===C[t]||"HETATM"===C[t];let n="X";P&&(n=P[t].replace(/\(?\+?\d+.*/,"")),s.elem=n[0].toUpperCase()+n.substring(1,2).toLowerCase(),U&&(s.b=U[t]),i&&"H"==s.elem||(""!=s.altLoc&&s.altLoc!=r&&"*"!=r||(s.bonds=[],s.ss="c",s.serial=B[t],s.model=V,s.bondOrder=[],s.properties={},s.index=h[h.length-1].length,o[s.serial]=[h.length,s.index],h[h.length-1].push(s),b.add(s)))}if(b.setBonds(y,x),!M(p))for(let t=s;t<h.length;t++){let e=h[t];for(let t=0;t<e.length;t++){const i=e[t];if(void 0!==i&&(i.lchain in p&&i.lresi in p[i.lchain])){const t=p[i.lchain][i.lresi];i.ss=t[0],t.length>1&&("1"==t[1]?i.ssbegin=!0:"2"==t[1]&&(i.ssend=!0))}}}e.multimodel&&t<d-1&&(e.onemol||(h.push([]),c.push({symmetries:[]})))}for(let t=0;t<h.length;t++)!o||e.duplicateAssemblyAtoms&&!e.dontConnectDuplicatedAtoms||b(h[t],e),s&&m(h[t],e.hbondCutoff),v(c[t].symmetries,h[t],e,c[t].cryst),e.duplicateAssemblyAtoms&&!e.dontConnectDuplicatedAtoms&&o&&b(h[t],e);return h}const K={vasp:c.VASP,VASP:c.VASP,cube:d.CUBE,CUBE:d.CUBE,xyz:f,XYZ:f,sdf:p,SDF:p,json:g,cdjson:g,CDJSON:g,mcif:x,cif:x,CIF:x,mol2:A,MOL2:A,pdb:T,PDB:T,pdbqt:T,PDBQT:T,pqr:E,PQR:E,mmtf:k,MMTF:k,prmtop:R,PRMTOP:R,gro:P,GRO:P,lammpstrj:B,LAMMPSTRJ:B,bcif:X,BCIF:X};var Q,$=i(529),J=i(99);!function(t){t[t.VDW=1]="VDW",t[t.MS=2]="MS",t[t.SAS=3]="SAS",t[t.SES=4]="SES"}(Q||(Q={}));var tt=!1;function et(t){tt=t}(window.navigator.userAgent.indexOf("MSIE ")>=0||window.navigator.userAgent.indexOf("Trident/")>=0)&&(tt=!0);class MarchingCubeInitializer{constructor(){this.ISDONE=2,this.edgeTable=new Uint32Array([0,0,0,0,0,0,0,2816,0,0,0,1792,0,3328,3584,3840,0,0,0,138,0,21,0,134,0,0,0,652,0,2067,3865,3600,0,0,0,42,0,0,0,294,0,0,21,28,0,3875,1049,3360,0,168,162,170,0,645,2475,2210,0,687,293,172,4010,3747,3497,3232,0,0,0,0,0,69,0,900,0,0,0,1792,138,131,1608,1920,0,81,0,2074,84,85,84,86,0,81,0,3676,330,1105,1881,1616,0,0,0,42,0,69,0,502,0,0,21,3580,138,2035,1273,1520,2816,104,2337,106,840,581,367,102,2816,3695,3429,3180,1898,1635,1385,1120,0,0,0,0,0,0,0,3910,0,0,69,588,42,2083,41,2880,0,0,0,1722,0,2293,4095,3830,0,255,757,764,2538,2291,3065,2800,0,0,81,338,0,3925,1119,3414,84,855,85,340,2130,2899,89,2384,1792,712,194,1162,4036,3781,3535,3270,708,719,197,204,3018,2755,2505,2240,0,0,0,0,168,420,168,1958,162,162,676,2988,170,163,680,928,3328,3096,3328,3642,52,53,1855,1590,2340,2111,2869,2620,298,51,825,560,3584,3584,3090,3482,1668,1941,1183,1430,146,2975,2069,2460,154,915,153,400,3840,3592,3329,3082,1796,1541,1295,1030,2818,2575,2309,2060,778,515,265,0]),this.triTable=[[],[],[],[],[],[],[],[11,9,8],[],[],[],[8,10,9],[],[10,8,11],[9,11,10],[8,10,9,8,11,10],[],[],[],[1,7,3],[],[4,2,0],[],[2,1,7],[],[],[],[2,7,3,2,9,7],[],[1,4,11,1,0,4],[3,8,0,11,9,4,11,10,9],[4,11,9,11,10,9],[],[],[],[5,3,1],[],[],[],[2,5,8,2,1,5],[],[],[2,4,0],[3,2,4],[],[0,9,1,8,10,5,8,11,10],[3,4,0,3,10,4],[5,8,10,8,11,10],[],[3,5,7],[7,1,5],[1,7,3,1,5,7],[],[9,2,0,9,7,2],[0,3,8,1,7,11,1,5,7],[11,1,7,1,5,7],[],[9,1,0,5,3,2,5,7,3],[8,2,5,8,0,2],[2,5,3,5,7,3],[3,9,1,3,8,9,7,11,10,7,10,5],[9,1,0,10,7,11,10,5,7],[3,8,0,7,10,5,7,11,10],[11,5,7,11,10,5],[],[],[],[],[],[0,6,2],[],[7,2,9,7,9,8],[],[],[],[8,10,9],[7,1,3],[7,1,0],[6,9,3,6,10,9],[7,10,8,10,9,8],[],[6,0,4],[],[11,1,4,11,3,1],[2,4,6],[2,0,4,2,4,6],[2,4,6],[1,4,2,4,6,2],[],[6,0,4],[],[2,11,3,6,9,4,6,10,9],[8,6,1,8,1,3],[10,0,6,0,4,6],[8,0,3,9,6,10,9,4,6],[10,4,6,10,9,4],[],[],[],[5,3,1],[],[0,6,2],[],[7,4,8,5,2,1,5,6,2],[],[],[2,4,0],[7,4,8,2,11,3,10,5,6],[7,1,3],[5,6,10,0,9,1,8,7,4],[5,6,10,7,0,3,7,4,0],[10,5,6,4,8,7],[9,11,8],[3,5,6],[0,5,11,0,11,8],[6,3,5,3,1,5],[3,9,6,3,8,9],[9,6,0,6,2,0],[0,3,8,2,5,6,2,1,5],[1,6,2,1,5,6],[9,11,8],[1,0,9,6,10,5,11,3,2],[6,10,5,2,8,0,2,11,8],[3,2,11,10,5,6],[10,5,6,9,3,8,9,1,3],[0,9,1,5,6,10],[8,0,3,10,5,6],[10,5,6],[],[],[],[],[],[],[],[1,10,2,9,11,6,9,8,11],[],[],[6,0,2],[3,6,9,3,2,6],[3,5,1],[0,5,1,0,11,5],[0,3,5],[6,9,11,9,8,11],[],[],[],[4,5,9,7,1,10,7,3,1],[],[11,6,7,2,4,5,2,0,4],[11,6,7,8,0,3,1,10,2,9,4,5],[6,7,11,1,10,2,9,4,5],[],[4,1,0,4,5,1,6,7,3,6,3,2],[9,4,5,0,6,7,0,2,6],[4,5,9,6,3,2,6,7,3],[6,7,11,5,3,8,5,1,3],[6,7,11,4,1,0,4,5,1],[4,5,9,3,8,0,11,6,7],[9,4,5,7,11,6],[],[],[0,6,4],[8,6,4,8,1,6],[],[0,10,2,0,9,10,4,8,11,4,11,6],[10,2,1,6,0,3,6,4,0],[10,2,1,11,4,8,11,6,4],[4,2,6],[1,0,9,2,4,8,2,6,4],[2,4,0,2,6,4],[8,2,4,2,6,4],[11,4,1,11,6,4],[0,9,1,4,11,6,4,8,11],[3,6,0,6,4,0],[8,6,4,8,11,6],[10,8,9],[6,3,9,6,7,3],[6,7,1],[10,7,1,7,3,1],[7,11,6,8,10,2,8,9,10],[11,6,7,10,0,9,10,2,0],[2,1,10,7,11,6,8,0,3],[1,10,2,6,7,11],[7,2,6,7,9,2],[1,0,9,3,6,7,3,2,6],[7,0,6,0,2,6],[2,7,3,2,6,7],[7,11,6,3,9,1,3,8,9],[9,1,0,11,6,7],[0,3,8,11,6,7],[11,6,7],[],[],[],[],[5,3,7],[8,5,2,8,7,5],[5,3,7],[1,10,2,5,8,7,5,9,8],[1,7,5],[1,7,5],[9,2,7,9,7,5],[11,3,2,8,5,9,8,7,5],[1,3,7,1,7,5],[0,7,1,7,5,1],[9,3,5,3,7,5],[9,7,5,9,8,7],[8,10,11],[3,4,10,3,10,11],[8,10,11],[5,9,4,1,11,3,1,10,11],[2,4,5],[5,2,4,2,0,4],[0,3,8,5,9,4,10,2,1],[2,1,10,9,4,5],[2,8,5,2,11,8],[3,2,11,1,4,5,1,0,4],[9,4,5,8,2,11,8,0,2],[11,3,2,9,4,5],[8,5,3,5,1,3],[5,0,4,5,1,0],[3,8,0,4,5,9],[9,4,5],[11,9,10],[11,9,10],[1,11,4,1,10,11],[8,7,4,11,1,10,11,3,1],[2,7,9,2,9,10],[4,8,7,0,10,2,0,9,10],[2,1,10,0,7,4,0,3,7],[10,2,1,8,7,4],[1,7,4],[3,2,11,4,8,7,9,1,0],[11,4,2,4,0,2],[2,11,3,7,4,8],[4,1,7,1,3,7],[1,0,9,8,7,4],[3,4,0,3,7,4],[8,7,4],[8,9,10,8,10,11],[3,9,11,9,10,11],[0,10,8,10,11,8],[10,3,1,10,11,3],[2,8,10,8,9,10],[9,2,0,9,10,2],[8,0,3,1,10,2],[10,2,1],[1,11,9,11,8,9],[11,3,2,0,9,1],[11,0,2,11,8,0],[11,3,2],[8,1,3,8,9,1],[9,1,0],[8,0,3],[]],this.edgeTable2=[0,265,515,778,2060,2309,2575,2822,1030,1295,1541,1804,3082,3331,3593,3840,400,153,915,666,2460,2197,2975,2710,1430,1183,1941,1692,3482,3219,3993,3728,560,825,51,314,2620,2869,2111,2358,1590,1855,1077,1340,3642,3891,3129,3376,928,681,419,170,2988,2725,2479,2214,1958,1711,1445,1196,4010,3747,3497,3232,2240,2505,2755,3018,204,453,719,966,3270,3535,3781,4044,1226,1475,1737,1984,2384,2137,2899,2650,348,85,863,598,3414,3167,3925,3676,1370,1107,1881,1616,2800,3065,2291,2554,764,1013,255,502,3830,4095,3317,3580,1786,2035,1273,1520,2912,2665,2403,2154,876,613,367,102,3942,3695,3429,3180,1898,1635,1385,1120,1120,1385,1635,1898,3180,3429,3695,3942,102,367,613,876,2154,2403,2665,2912,1520,1273,2035,1786,3580,3317,4095,3830,502,255,1013,764,2554,2291,3065,2800,1616,1881,1107,1370,3676,3925,3167,3414,598,863,85,348,2650,2899,2137,2384,1984,1737,1475,1226,4044,3781,3535,3270,966,719,453,204,3018,2755,2505,2240,3232,3497,3747,4010,1196,1445,1711,1958,2214,2479,2725,2988,170,419,681,928,3376,3129,3891,3642,1340,1077,1855,1590,2358,2111,2869,2620,314,51,825,560,3728,3993,3219,3482,1692,1941,1183,1430,2710,2975,2197,2460,666,915,153,400,3840,3593,3331,3082,1804,1541,1295,1030,2822,2575,2309,2060,778,515,265,0],this.triTable2=[[],[8,3,0],[9,0,1],[8,3,1,8,1,9],[11,2,3],[11,2,0,11,0,8],[11,2,3,0,1,9],[2,1,11,1,9,11,11,9,8],[10,1,2],[8,3,0,1,2,10],[9,0,2,9,2,10],[3,2,8,2,10,8,8,10,9],[10,1,3,10,3,11],[1,0,10,0,8,10,10,8,11],[0,3,9,3,11,9,9,11,10],[8,10,9,8,11,10],[8,4,7],[3,0,4,3,4,7],[1,9,0,8,4,7],[9,4,1,4,7,1,1,7,3],[2,3,11,7,8,4],[7,11,4,11,2,4,4,2,0],[3,11,2,4,7,8,9,0,1],[2,7,11,2,1,7,1,4,7,1,9,4],[10,1,2,8,4,7],[2,10,1,0,4,7,0,7,3],[4,7,8,0,2,10,0,10,9],[2,7,3,2,9,7,7,9,4,2,10,9],[8,4,7,11,10,1,11,1,3],[11,4,7,1,4,11,1,11,10,1,0,4],[3,8,0,7,11,4,11,9,4,11,10,9],[7,11,4,4,11,9,11,10,9],[9,5,4],[3,0,8,4,9,5],[5,4,0,5,0,1],[4,8,5,8,3,5,5,3,1],[11,2,3,9,5,4],[9,5,4,8,11,2,8,2,0],[3,11,2,1,5,4,1,4,0],[8,5,4,2,5,8,2,8,11,2,1,5],[2,10,1,9,5,4],[0,8,3,5,4,9,10,1,2],[10,5,2,5,4,2,2,4,0],[3,4,8,3,2,4,2,5,4,2,10,5],[5,4,9,1,3,11,1,11,10],[0,9,1,4,8,5,8,10,5,8,11,10],[3,4,0,3,10,4,4,10,5,3,11,10],[4,8,5,5,8,10,8,11,10],[9,5,7,9,7,8],[0,9,3,9,5,3,3,5,7],[8,0,7,0,1,7,7,1,5],[1,7,3,1,5,7],[11,2,3,8,9,5,8,5,7],[9,2,0,9,7,2,2,7,11,9,5,7],[0,3,8,2,1,11,1,7,11,1,5,7],[2,1,11,11,1,7,1,5,7],[1,2,10,5,7,8,5,8,9],[9,1,0,10,5,2,5,3,2,5,7,3],[5,2,10,8,2,5,8,5,7,8,0,2],[10,5,2,2,5,3,5,7,3],[3,9,1,3,8,9,7,11,10,7,10,5],[9,1,0,10,7,11,10,5,7],[3,8,0,7,10,5,7,11,10],[11,5,7,11,10,5],[11,7,6],[0,8,3,11,7,6],[9,0,1,11,7,6],[7,6,11,3,1,9,3,9,8],[2,3,7,2,7,6],[8,7,0,7,6,0,0,6,2],[1,9,0,3,7,6,3,6,2],[7,6,2,7,2,9,2,1,9,7,9,8],[1,2,10,6,11,7],[2,10,1,7,6,11,8,3,0],[11,7,6,10,9,0,10,0,2],[7,6,11,3,2,8,8,2,10,8,10,9],[6,10,7,10,1,7,7,1,3],[6,10,1,6,1,7,7,1,0,7,0,8],[9,0,3,6,9,3,6,10,9,6,3,7],[6,10,7,7,10,8,10,9,8],[8,4,6,8,6,11],[11,3,6,3,0,6,6,0,4],[0,1,9,4,6,11,4,11,8],[1,9,4,11,1,4,11,3,1,11,4,6],[3,8,2,8,4,2,2,4,6],[2,0,4,2,4,6],[1,9,0,3,8,2,2,8,4,2,4,6],[9,4,1,1,4,2,4,6,2],[10,1,2,11,8,4,11,4,6],[10,1,2,11,3,6,6,3,0,6,0,4],[0,2,10,0,10,9,4,11,8,4,6,11],[2,11,3,6,9,4,6,10,9],[8,4,6,8,6,1,6,10,1,8,1,3],[1,0,10,10,0,6,0,4,6],[8,0,3,9,6,10,9,4,6],[10,4,6,10,9,4],[9,5,4,7,6,11],[4,9,5,3,0,8,11,7,6],[6,11,7,4,0,1,4,1,5],[6,11,7,4,8,5,5,8,3,5,3,1],[4,9,5,6,2,3,6,3,7],[9,5,4,8,7,0,0,7,6,0,6,2],[4,0,1,4,1,5,6,3,7,6,2,3],[7,4,8,5,2,1,5,6,2],[6,11,7,1,2,10,9,5,4],[11,7,6,8,3,0,1,2,10,9,5,4],[11,7,6,10,5,2,2,5,4,2,4,0],[7,4,8,2,11,3,10,5,6],[4,9,5,6,10,7,7,10,1,7,1,3],[5,6,10,0,9,1,8,7,4],[5,6,10,7,0,3,7,4,0],[10,5,6,4,8,7],[5,6,9,6,11,9,9,11,8],[0,9,5,0,5,3,3,5,6,3,6,11],[0,1,5,0,5,11,5,6,11,0,11,8],[11,3,6,6,3,5,3,1,5],[9,5,6,3,9,6,3,8,9,3,6,2],[5,6,9,9,6,0,6,2,0],[0,3,8,2,5,6,2,1,5],[1,6,2,1,5,6],[1,2,10,5,6,9,9,6,11,9,11,8],[1,0,9,6,10,5,11,3,2],[6,10,5,2,8,0,2,11,8],[3,2,11,10,5,6],[10,5,6,9,3,8,9,1,3],[0,9,1,5,6,10],[8,0,3,10,5,6],[10,5,6],[10,6,5],[8,3,0,10,6,5],[0,1,9,5,10,6],[10,6,5,9,8,3,9,3,1],[3,11,2,10,6,5],[6,5,10,2,0,8,2,8,11],[1,9,0,6,5,10,11,2,3],[1,10,2,5,9,6,9,11,6,9,8,11],[1,2,6,1,6,5],[0,8,3,2,6,5,2,5,1],[5,9,6,9,0,6,6,0,2],[9,6,5,3,6,9,3,9,8,3,2,6],[11,6,3,6,5,3,3,5,1],[0,5,1,0,11,5,5,11,6,0,8,11],[0,5,9,0,3,5,3,6,5,3,11,6],[5,9,6,6,9,11,9,8,11],[10,6,5,4,7,8],[5,10,6,7,3,0,7,0,4],[5,10,6,0,1,9,8,4,7],[4,5,9,6,7,10,7,1,10,7,3,1],[7,8,4,2,3,11,10,6,5],[11,6,7,10,2,5,2,4,5,2,0,4],[11,6,7,8,0,3,1,10,2,9,4,5],[6,7,11,1,10,2,9,4,5],[7,8,4,5,1,2,5,2,6],[4,1,0,4,5,1,6,7,3,6,3,2],[9,4,5,8,0,7,0,6,7,0,2,6],[4,5,9,6,3,2,6,7,3],[6,7,11,4,5,8,5,3,8,5,1,3],[6,7,11,4,1,0,4,5,1],[4,5,9,3,8,0,11,6,7],[9,4,5,7,11,6],[10,6,4,10,4,9],[8,3,0,9,10,6,9,6,4],[1,10,0,10,6,0,0,6,4],[8,6,4,8,1,6,6,1,10,8,3,1],[2,3,11,6,4,9,6,9,10],[0,10,2,0,9,10,4,8,11,4,11,6],[10,2,1,11,6,3,6,0,3,6,4,0],[10,2,1,11,4,8,11,6,4],[9,1,4,1,2,4,4,2,6],[1,0,9,3,2,8,2,4,8,2,6,4],[2,4,0,2,6,4],[3,2,8,8,2,4,2,6,4],[1,4,9,11,4,1,11,1,3,11,6,4],[0,9,1,4,11,6,4,8,11],[11,6,3,3,6,0,6,4,0],[8,6,4,8,11,6],[6,7,10,7,8,10,10,8,9],[9,3,0,6,3,9,6,9,10,6,7,3],[6,1,10,6,7,1,7,0,1,7,8,0],[6,7,10,10,7,1,7,3,1],[7,11,6,3,8,2,8,10,2,8,9,10],[11,6,7,10,0,9,10,2,0],[2,1,10,7,11,6,8,0,3],[1,10,2,6,7,11],[7,2,6,7,9,2,2,9,1,7,8,9],[1,0,9,3,6,7,3,2,6],[8,0,7,7,0,6,0,2,6],[2,7,3,2,6,7],[7,11,6,3,9,1,3,8,9],[9,1,0,11,6,7],[0,3,8,11,6,7],[11,6,7],[11,7,5,11,5,10],[3,0,8,7,5,10,7,10,11],[9,0,1,10,11,7,10,7,5],[3,1,9,3,9,8,7,10,11,7,5,10],[10,2,5,2,3,5,5,3,7],[5,10,2,8,5,2,8,7,5,8,2,0],[9,0,1,10,2,5,5,2,3,5,3,7],[1,10,2,5,8,7,5,9,8],[2,11,1,11,7,1,1,7,5],[0,8,3,2,11,1,1,11,7,1,7,5],[9,0,2,9,2,7,2,11,7,9,7,5],[11,3,2,8,5,9,8,7,5],[1,3,7,1,7,5],[8,7,0,0,7,1,7,5,1],[0,3,9,9,3,5,3,7,5],[9,7,5,9,8,7],[4,5,8,5,10,8,8,10,11],[3,0,4,3,4,10,4,5,10,3,10,11],[0,1,9,4,5,8,8,5,10,8,10,11],[5,9,4,1,11,3,1,10,11],[3,8,4,3,4,2,2,4,5,2,5,10],[10,2,5,5,2,4,2,0,4],[0,3,8,5,9,4,10,2,1],[2,1,10,9,4,5],[8,4,5,2,8,5,2,11,8,2,5,1],[3,2,11,1,4,5,1,0,4],[9,4,5,8,2,11,8,0,2],[11,3,2,9,4,5],[4,5,8,8,5,3,5,1,3],[5,0,4,5,1,0],[3,8,0,4,5,9],[9,4,5],[7,4,11,4,9,11,11,9,10],[3,0,8,7,4,11,11,4,9,11,9,10],[11,7,4,1,11,4,1,10,11,1,4,0],[8,7,4,11,1,10,11,3,1],[2,3,7,2,7,9,7,4,9,2,9,10],[4,8,7,0,10,2,0,9,10],[2,1,10,0,7,4,0,3,7],[10,2,1,8,7,4],[2,11,7,2,7,1,1,7,4,1,4,9],[3,2,11,4,8,7,9,1,0],[7,4,11,11,4,2,4,0,2],[2,11,3,7,4,8],[9,1,4,4,1,7,1,3,7],[1,0,9,8,7,4],[3,4,0,3,7,4],[8,7,4],[8,9,10,8,10,11],[0,9,3,3,9,11,9,10,11],[1,10,0,0,10,8,10,11,8],[10,3,1,10,11,3],[3,8,2,2,8,10,8,9,10],[9,2,0,9,10,2],[8,0,3,1,10,2],[10,2,1],[2,11,1,1,11,9,11,8,9],[11,3,2,0,9,1],[11,0,2,11,8,0],[11,3,2],[8,1,3,8,9,1],[9,1,0],[8,0,3],[]]}march(t,e,i,r){let s=!!r.fulltable,n=r.hasOwnProperty("origin")&&r.origin.hasOwnProperty("x")?r.origin:{x:0,y:0,z:0},a=!!r.voxel,o=r.matrix,l=r.nX||0,h=r.nY||0,c=r.nZ||0,d=r.scale||1,u=null;u=r.unitCube?r.unitCube:{x:d,y:d,z:d};let f,p,g=new Int32Array(l*h*c);for(f=0,p=g.length;f<p;++f)g[f]=-1;let m=function(t,i,r,s,l,d){let f={x:0,y:0,z:0},p=l;if(!!!(s&1<<l)&&!!(s&1<<d)&&(p=d),1&p&&r++,2&p&&i++,4&p&&t++,o){let e=new $.Vector3(t,i,r);e=e.applyMatrix4(o),f={x:e.x,y:e.y,z:e.z}}else f.x=n.x+u.x*t,f.y=n.y+u.y*i,f.z=n.z+u.z*r;let m=(h*t+i)*c+r;return a?(e.push(f),e.length-1):(g[m]<0&&(g[m]=e.length,e.push(f)),g[m])},v=new Int32Array(12),_=s?this.edgeTable2:this.edgeTable,y=s?this.triTable2:this.triTable;for(f=0;f<l-1;++f)for(let r=0;r<h-1;++r)for(let s=0;s<c-1;++s){let n=0;for(let e=0;e<8;++e){n|=!!(t[(h*(f+((4&e)>>2))+r+((2&e)>>1))*c+s+(1&e)]&this.ISDONE)<<e}if(0===n||255===n)continue;let o=_[n];if(0===o)continue;let l=y[n];1&o&&(v[0]=m(f,r,s,n,0,1)),2&o&&(v[1]=m(f,r,s,n,1,3)),4&o&&(v[2]=m(f,r,s,n,3,2)),8&o&&(v[3]=m(f,r,s,n,2,0)),16&o&&(v[4]=m(f,r,s,n,4,5)),32&o&&(v[5]=m(f,r,s,n,5,7)),64&o&&(v[6]=m(f,r,s,n,7,6)),128&o&&(v[7]=m(f,r,s,n,6,4)),256&o&&(v[8]=m(f,r,s,n,0,4)),512&o&&(v[9]=m(f,r,s,n,1,5)),1024&o&&(v[10]=m(f,r,s,n,3,7)),2048&o&&(v[11]=m(f,r,s,n,2,6));for(let t=0;t<l.length;t+=3){let r=v[l[t]],s=v[l[t+1]],n=v[l[t+2]];a&&t>=3&&(e.push(e[r]),r=e.length-1,e.push(e[s]),s=e.length-1,e.push(e[n]),n=e.length-1),i.push(r),i.push(s),i.push(n)}}}laplacianSmooth(t,e,i){let r,s,n,a,o,l=new Array(e.length);for(r=0,s=e.length;r<s;r++)l[r]={x:0,y:0,z:0};let h,c=new Array(20);for(r=0;r<20;r++)c[r]=new Array(e.length);for(r=0,s=e.length;r<s;r++)c[0][r]=0;for(r=0,s=i.length/3;r<s;r++){let t=3*r,e=3*r+1,s=3*r+2;for(h=!0,n=0,a=c[0][i[t]];n<a;n++)if(i[e]==c[n+1][i[t]]){h=!1;break}for(h&&(c[0][i[t]]++,c[c[0][i[t]]][i[t]]=i[e]),h=!0,n=0,a=c[0][i[t]];n<a;n++)if(i[s]==c[n+1][i[t]]){h=!1;break}for(h&&(c[0][i[t]]++,c[c[0][i[t]]][i[t]]=i[s]),h=!0,n=0,a=c[0][i[e]];n<a;n++)if(i[t]==c[n+1][i[e]]){h=!1;break}for(h&&(c[0][i[e]]++,c[c[0][i[e]]][i[e]]=i[t]),h=!0,n=0,a=c[0][i[e]];n<a;n++)if(i[s]==c[n+1][i[e]]){h=!1;break}for(h&&(c[0][i[e]]++,c[c[0][i[e]]][i[e]]=i[s]),h=!0,n=0;n<c[0][i[s]];n++)if(i[t]==c[n+1][i[s]]){h=!1;break}for(h&&(c[0][i[s]]++,c[c[0][i[s]]][i[s]]=i[t]),h=!0,n=0,a=c[0][i[s]];n<a;n++)if(i[e]==c[n+1][i[s]]){h=!1;break}h&&(c[0][i[s]]++,c[c[0][i[s]]][i[s]]=i[e])}let d=.5;for(o=0;o<t;o++){for(r=0,s=e.length;r<s;r++)if(c[0][r]<3)l[r].x=e[r].x,l[r].y=e[r].y,l[r].z=e[r].z;else if(3==c[0][r]||4==c[0][r]){for(l[r].x=0,l[r].y=0,l[r].z=0,n=0,a=c[0][r];n<a;n++)l[r].x+=e[c[n+1][r]].x,l[r].y+=e[c[n+1][r]].y,l[r].z+=e[c[n+1][r]].z;l[r].x+=d*e[r].x,l[r].y+=d*e[r].y,l[r].z+=d*e[r].z,l[r].x/=d+c[0][r],l[r].y/=d+c[0][r],l[r].z/=d+c[0][r]}else{for(l[r].x=0,l[r].y=0,l[r].z=0,n=0,a=c[0][r];n<a;n++)l[r].x+=e[c[n+1][r]].x,l[r].y+=e[c[n+1][r]].y,l[r].z+=e[c[n+1][r]].z;l[r].x+=1*e[r].x,l[r].y+=1*e[r].y,l[r].z+=1*e[r].z,l[r].x/=1+c[0][r],l[r].y/=1+c[0][r],l[r].z/=1+c[0][r]}for(r=0,s=e.length;r<s;r++)e[r].x=l[r].x,e[r].y=l[r].y,e[r].z=l[r].z}}}let MarchingCube=new MarchingCubeInitializer;class PointGrid{constructor(t,e,i){this.data=new Int32Array(t*e*i*3),this.width=e,this.height=i}set(t,e,i,r){let s=3*((t*this.width+e)*this.height+i);this.data[s]=r.ix,this.data[s+1]=r.iy,this.data[s+2]=r.iz}get(t,e,i){let r=3*((t*this.width+e)*this.height+i);return{ix:this.data[r],iy:this.data[r+1],iz:this.data[r+2]}}}class ProteinSurface{constructor(){this.INOUT=1,this.ISDONE=2,this.ISBOUND=4,this.ptranx=0,this.ptrany=0,this.ptranz=0,this.probeRadius=1.4,this.defaultScaleFactor=2,this.scaleFactor=this.defaultScaleFactor,this.pHeight=0,this.pWidth=0,this.pLength=0,this.cutRadius=0,this.vpBits=null,this.vpDistance=null,this.vpAtomID=null,this.pminx=0,this.pminy=0,this.pminz=0,this.pmaxx=0,this.pmaxy=0,this.pmaxz=0,this.depty={},this.widxz={},this.faces=[],this.verts=[],this.vdwRadii={H:1.2,Li:1.82,Na:2.27,K:2.75,C:1.7,N:1.55,O:1.52,F:1.47,P:1.8,S:1.8,CL:1.75,BR:1.85,SE:1.9,ZN:1.39,CU:1.4,NI:1.63,X:2},this.nb=[new Int32Array([1,0,0]),new Int32Array([-1,0,0]),new Int32Array([0,1,0]),new Int32Array([0,-1,0]),new Int32Array([0,0,1]),new Int32Array([0,0,-1]),new Int32Array([1,1,0]),new Int32Array([1,-1,0]),new Int32Array([-1,1,0]),new Int32Array([-1,-1,0]),new Int32Array([1,0,1]),new Int32Array([1,0,-1]),new Int32Array([-1,0,1]),new Int32Array([-1,0,-1]),new Int32Array([0,1,1]),new Int32Array([0,1,-1]),new Int32Array([0,-1,1]),new Int32Array([0,-1,-1]),new Int32Array([1,1,1]),new Int32Array([1,1,-1]),new Int32Array([1,-1,1]),new Int32Array([-1,1,1]),new Int32Array([1,-1,-1]),new Int32Array([-1,-1,1]),new Int32Array([-1,1,-1]),new Int32Array([-1,-1,-1])],ProteinSurface.MarchingCube||(ProteinSurface.MarchingCube=new MarchingCubeInitializer)}getVDWIndex(t){return t.elem&&void 0!==this.vdwRadii[t.elem]?t.elem:"X"}getFacesAndVertices(t){let e={};for(let i=0,r=t.length;i<r;i++)e[t[i]]=!0;let i=this.verts;for(let t=0,e=i.length;t<e;t++)i[t].x=i[t].x/this.scaleFactor-this.ptranx,i[t].y=i[t].y/this.scaleFactor-this.ptrany,i[t].z=i[t].z/this.scaleFactor-this.ptranz;let r=[];for(let t=0,s=this.faces.length;t<s;t+=3){let s=this.faces[t],n=this.faces[t+1],a=this.faces[t+2],o=i[s].atomid,l=i[n].atomid,h=i[a].atomid,c=o;l<c&&(c=l),h<c&&(c=h),e[c]&&(s!==n&&n!==a&&s!==a&&(r.push(s),r.push(n),r.push(a)))}return this.vpBits=null,this.vpDistance=null,this.vpAtomID=null,{vertices:i,faces:r}}initparm(t,e,i){i>1e6&&(this.scaleFactor=this.defaultScaleFactor/2);let r=1/this.scaleFactor*5.5;this.pminx=t[0][0],this.pmaxx=t[1][0],this.pminy=t[0][1],this.pmaxy=t[1][1],this.pminz=t[0][2],this.pmaxz=t[1][2],e?(this.pminx-=this.probeRadius+r,this.pminy-=this.probeRadius+r,this.pminz-=this.probeRadius+r,this.pmaxx+=this.probeRadius+r,this.pmaxy+=this.probeRadius+r,this.pmaxz+=this.probeRadius+r):(this.pminx-=r,this.pminy-=r,this.pminz-=r,this.pmaxx+=r,this.pmaxy+=r,this.pmaxz+=r),this.pminx=Math.floor(this.pminx*this.scaleFactor)/this.scaleFactor,this.pminy=Math.floor(this.pminy*this.scaleFactor)/this.scaleFactor,this.pminz=Math.floor(this.pminz*this.scaleFactor)/this.scaleFactor,this.pmaxx=Math.ceil(this.pmaxx*this.scaleFactor)/this.scaleFactor,this.pmaxy=Math.ceil(this.pmaxy*this.scaleFactor)/this.scaleFactor,this.pmaxz=Math.ceil(this.pmaxz*this.scaleFactor)/this.scaleFactor,this.ptranx=-this.pminx,this.ptrany=-this.pminy,this.ptranz=-this.pminz,this.pLength=Math.ceil(this.scaleFactor*(this.pmaxx-this.pminx))+1,this.pWidth=Math.ceil(this.scaleFactor*(this.pmaxy-this.pminy))+1,this.pHeight=Math.ceil(this.scaleFactor*(this.pmaxz-this.pminz))+1,this.boundingatom(e),this.cutRadius=this.probeRadius*this.scaleFactor,this.vpBits=new Uint8Array(this.pLength*this.pWidth*this.pHeight),this.vpDistance=new Float64Array(this.pLength*this.pWidth*this.pHeight),this.vpAtomID=new Int32Array(this.pLength*this.pWidth*this.pHeight)}boundingatom(t){let e={};for(const i in this.vdwRadii){let r=this.vdwRadii[i];e[i]=t?(r+this.probeRadius)*this.scaleFactor+.5:r*this.scaleFactor+.5;let s=e[i]*e[i];this.widxz[i]=Math.floor(e[i])+1,this.depty[i]=new Int32Array(this.widxz[i]*this.widxz[i]);let n=0;for(let t=0;t<this.widxz[i];t++)for(let e=0;e<this.widxz[i];e++){let r=t*t+e*e;if(r>s)this.depty[i][n]=-1;else{let t=Math.sqrt(s-r);this.depty[i][n]=Math.floor(t)}n++}}}fillvoxels(t,e){for(let t=0,e=this.vpBits.length;t<e;t++)this.vpBits[t]=0,this.vpDistance[t]=-1,this.vpAtomID[t]=-1;for(let i in e){let r=t[e[i]];void 0!==r&&this.fillAtom(r,t)}for(let t=0,e=this.vpBits.length;t<e;t++)this.vpBits[t]&this.INOUT&&(this.vpBits[t]|=this.ISDONE)}fillAtom(t,e){let i=Math.floor(.5+this.scaleFactor*(t.x+this.ptranx)),r=Math.floor(.5+this.scaleFactor*(t.y+this.ptrany)),s=Math.floor(.5+this.scaleFactor*(t.z+this.ptranz)),n=this.getVDWIndex(t),a=0,o=this.pWidth*this.pHeight;for(let l=0,h=this.widxz[n];l<h;l++)for(let c=0;c<h;c++){if(-1!=this.depty[n][a])for(let h=-1;h<2;h++)for(let d=-1;d<2;d++)for(let u=-1;u<2;u++)if(0!==h&&0!==d&&0!==u){let f=h*l,p=u*c;for(let l=0;l<=this.depty[n][a];l++){let n=l*d,a=i+f,h=r+n,c=s+p;if(a<0||h<0||c<0||a>=this.pLength||h>=this.pWidth||c>=this.pHeight)continue;let u=a*o+h*this.pHeight+c;if(this.vpBits[u]&this.INOUT){let a=e[this.vpAtomID[u]];if(a.serial!=t.serial){let e=i+f-Math.floor(.5+this.scaleFactor*(a.x+this.ptranx)),o=r+n-Math.floor(.5+this.scaleFactor*(a.y+this.ptrany)),l=s+p-Math.floor(.5+this.scaleFactor*(a.z+this.ptranz));f*f+n*n+p*p<e*e+o*o+l*l&&(this.vpAtomID[u]=t.serial)}}else this.vpBits[u]|=this.INOUT,this.vpAtomID[u]=t.serial}}a++}}fillvoxelswaals(t,e){for(let t=0,e=this.vpBits.length;t<e;t++)this.vpBits[t]&=~this.ISDONE;for(let i in e){let r=t[e[i]];void 0!==r&&this.fillAtomWaals(r,t)}}fillAtomWaals(t,e){let i=0,r=Math.floor(.5+this.scaleFactor*(t.x+this.ptranx)),s=Math.floor(.5+this.scaleFactor*(t.y+this.ptrany)),n=Math.floor(.5+this.scaleFactor*(t.z+this.ptranz)),a=this.getVDWIndex(t),o=this.pWidth*this.pHeight;for(let l=0,h=this.widxz[a];l<h;l++)for(let c=0;c<h;c++){if(-1!=this.depty[a][i])for(let h=-1;h<2;h++)for(let d=-1;d<2;d++)for(let u=-1;u<2;u++)if(0!==h&&0!==d&&0!==u){let f=h*l,p=u*c;for(let l=0;l<=this.depty[a][i];l++){let i=l*d,a=r+f,h=s+i,c=n+p;if(a<0||h<0||c<0||a>=this.pLength||h>=this.pWidth||c>=this.pHeight)continue;let u=a*o+h*this.pHeight+c;if(this.vpBits[u]&this.ISDONE){let a=e[this.vpAtomID[u]];if(a.serial!=t.serial){let e=r+f-Math.floor(.5+this.scaleFactor*(a.x+this.ptranx)),o=s+i-Math.floor(.5+this.scaleFactor*(a.y+this.ptrany)),l=n+p-Math.floor(.5+this.scaleFactor*(a.z+this.ptranz));f*f+i*i+p*p<e*e+o*o+l*l&&(this.vpAtomID[u]=t.serial)}}else this.vpBits[u]|=this.ISDONE,this.vpAtomID[u]=t.serial}}i++}}buildboundary(){let t=this.pWidth*this.pHeight;for(let e=0;e<this.pLength;e++)for(let i=0;i<this.pHeight;i++)for(let r=0;r<this.pWidth;r++){let s=e*t+r*this.pHeight+i;if(this.vpBits[s]&this.INOUT){let n=0;for(;n<26;){let a=e+this.nb[n][0],o=i+this.nb[n][2],l=r+this.nb[n][1];if(a>-1&&a<this.pLength&&l>-1&&l<this.pWidth&&o>-1&&o<this.pHeight&&!(this.vpBits[a*t+l*this.pHeight+o]&this.INOUT)){this.vpBits[s]|=this.ISBOUND;break}n++}}}}fastdistancemap(){let t,e=new PointGrid(this.pLength,this.pWidth,this.pHeight),i=this.pWidth*this.pHeight,r=this.cutRadius*this.cutRadius,s=[],n=[];for(let r=0;r<this.pLength;r++)for(let n=0;n<this.pWidth;n++)for(let a=0;a<this.pHeight;a++)if(t=r*i+n*this.pHeight+a,this.vpBits[t]&=~this.ISDONE,this.vpBits[t]&this.INOUT&&this.vpBits[t]&this.ISBOUND){let i={ix:r,iy:n,iz:a};e.set(r,n,a,i),s.push(i),this.vpDistance[t]=0,this.vpBits[t]|=this.ISDONE,this.vpBits[t]&=~this.ISBOUND}do{n=this.fastoneshell(s,e),s=[];for(let e=0,a=n.length;e<a;e++)t=i*n[e].ix+this.pHeight*n[e].iy+n[e].iz,this.vpBits[t]&=~this.ISBOUND,this.vpDistance[t]<=1.0404*r&&s.push({ix:n[e].ix,iy:n[e].iy,iz:n[e].iz})}while(0!==s.length);s=[],n=[],e=null;let a=this.scaleFactor-.5;a<0&&(a=0);let o=r-.5/(.1+a);for(let e=0;e<this.pLength;e++)for(let r=0;r<this.pWidth;r++)for(let s=0;s<this.pHeight;s++)t=e*i+r*this.pHeight+s,this.vpBits[t]&=~this.ISBOUND,this.vpBits[t]&this.INOUT&&(this.vpBits[t]&this.ISDONE&&!(this.vpBits[t]&this.ISDONE&&this.vpDistance[t]>=o)||(this.vpBits[t]|=this.ISBOUND))}fastoneshell(t,e){let i,r,s,n,a,o,l,h,c,d=[];if(0===t.length)return d;let u={ix:-1,iy:-1,iz:-1},f=this.pWidth*this.pHeight;for(let p=0,g=t.length;p<g;p++){i=t[p].ix,r=t[p].iy,s=t[p].iz,h=e.get(i,r,s);for(let t=0;t<6;t++)u.ix=i+this.nb[t][0],u.iy=r+this.nb[t][1],u.iz=s+this.nb[t][2],u.ix<this.pLength&&u.ix>-1&&u.iy<this.pWidth&&u.iy>-1&&u.iz<this.pHeight&&u.iz>-1&&(c=u.ix*f+this.pHeight*u.iy+u.iz,this.vpBits[c]&this.INOUT&&!(this.vpBits[c]&this.ISDONE)?(e.set(u.ix,u.iy,s+this.nb[t][2],h),n=u.ix-h.ix,a=u.iy-h.iy,o=u.iz-h.iz,l=n*n+a*a+o*o,this.vpDistance[c]=l,this.vpBits[c]|=this.ISDONE,this.vpBits[c]|=this.ISBOUND,d.push({ix:u.ix,iy:u.iy,iz:u.iz})):this.vpBits[c]&this.INOUT&&this.vpBits[c]&this.ISDONE&&(n=u.ix-h.ix,a=u.iy-h.iy,o=u.iz-h.iz,l=n*n+a*a+o*o,l<this.vpDistance[c]&&(e.set(u.ix,u.iy,u.iz,h),this.vpDistance[c]=l,this.vpBits[c]&this.ISBOUND||(this.vpBits[c]|=this.ISBOUND,d.push({ix:u.ix,iy:u.iy,iz:u.iz})))))}for(let p=0,g=t.length;p<g;p++){i=t[p].ix,r=t[p].iy,s=t[p].iz,h=e.get(i,r,s);for(let t=6;t<18;t++)u.ix=i+this.nb[t][0],u.iy=r+this.nb[t][1],u.iz=s+this.nb[t][2],u.ix<this.pLength&&u.ix>-1&&u.iy<this.pWidth&&u.iy>-1&&u.iz<this.pHeight&&u.iz>-1&&(c=u.ix*f+this.pHeight*u.iy+u.iz,this.vpBits[c]&this.INOUT&&!(this.vpBits[c]&this.ISDONE)?(e.set(u.ix,u.iy,s+this.nb[t][2],h),n=u.ix-h.ix,a=u.iy-h.iy,o=u.iz-h.iz,l=n*n+a*a+o*o,this.vpDistance[c]=l,this.vpBits[c]|=this.ISDONE,this.vpBits[c]|=this.ISBOUND,d.push({ix:u.ix,iy:u.iy,iz:u.iz})):this.vpBits[c]&this.INOUT&&this.vpBits[c]&this.ISDONE&&(n=u.ix-h.ix,a=u.iy-h.iy,o=u.iz-h.iz,l=n*n+a*a+o*o,l<this.vpDistance[c]&&(e.set(u.ix,u.iy,u.iz,h),this.vpDistance[c]=l,this.vpBits[c]&this.ISBOUND||(this.vpBits[c]|=this.ISBOUND,d.push({ix:u.ix,iy:u.iy,iz:u.iz})))))}for(let p=0,g=t.length;p<g;p++){i=t[p].ix,r=t[p].iy,s=t[p].iz,h=e.get(i,r,s);for(let t=18;t<26;t++)u.ix=i+this.nb[t][0],u.iy=r+this.nb[t][1],u.iz=s+this.nb[t][2],u.ix<this.pLength&&u.ix>-1&&u.iy<this.pWidth&&u.iy>-1&&u.iz<this.pHeight&&u.iz>-1&&(c=u.ix*f+this.pHeight*u.iy+u.iz,this.vpBits[c]&this.INOUT&&!(this.vpBits[c]&this.ISDONE)?(e.set(u.ix,u.iy,s+this.nb[t][2],h),n=u.ix-h.ix,a=u.iy-h.iy,o=u.iz-h.iz,l=n*n+a*a+o*o,this.vpDistance[c]=l,this.vpBits[c]|=this.ISDONE,this.vpBits[c]|=this.ISBOUND,d.push({ix:u.ix,iy:u.iy,iz:u.iz})):this.vpBits[c]&this.INOUT&&this.vpBits[c]&this.ISDONE&&(n=u.ix-h.ix,a=u.iy-h.iy,o=u.iz-h.iz,l=n*n+a*a+o*o,l<this.vpDistance[c]&&(e.set(u.ix,u.iy,u.iz,h),this.vpDistance[c]=l,this.vpBits[c]&this.ISBOUND||(this.vpBits[c]|=this.ISBOUND,d.push({ix:u.ix,iy:u.iy,iz:u.iz})))))}return d}marchingcubeinit(t){for(let e=0,i=this.vpBits.length;e<i;e++)1==t?this.vpBits[e]&=~this.ISBOUND:4==t?(this.vpBits[e]&=~this.ISDONE,this.vpBits[e]&this.ISBOUND&&(this.vpBits[e]|=this.ISDONE),this.vpBits[e]&=~this.ISBOUND):2==t?this.vpBits[e]&this.ISBOUND&&this.vpBits[e]&this.ISDONE?this.vpBits[e]&=~this.ISBOUND:this.vpBits[e]&this.ISBOUND&&!(this.vpBits[e]&this.ISDONE)&&(this.vpBits[e]|=this.ISDONE):3==t&&(this.vpBits[e]&=~this.ISBOUND)}marchingcube(t){this.marchingcubeinit(t),this.verts=[],this.faces=[],ProteinSurface.MarchingCube.march(this.vpBits,this.verts,this.faces,{smooth:1,nX:this.pLength,nY:this.pWidth,nZ:this.pHeight});let e=this.pWidth*this.pHeight;for(let t=0,i=this.verts.length;t<i;t++)this.verts[t].atomid=this.vpAtomID[this.verts[t].x*e+this.pHeight*this.verts[t].y+this.verts[t].z];ProteinSurface.MarchingCube.laplacianSmooth(1,this.verts,this.faces)}}ProteinSurface.MarchingCube=new MarchingCubeInitializer;var it,rt,st=i(848);function nt(t,e){var i,r,s,n,a,o,l,h,c,d=[],u=t;for((u=[]).push(t[0]),i=1,r=t.length-1;i<r;i++)if(a=t[i],o=t[i+1],a.smoothen){var f=new $.Vector3((a.x+o.x)/2,(a.y+o.y)/2,(a.z+o.z)/2);f.atom=a.atom,u.push(f)}else u.push(a);for(u.push(t[t.length-1]),i=-1,s=u.length;i<=s-3;i++)if(n=u[-1===i?0:i],a=u[i+1],o=u[i+2],l=u[i===s-3?s-1:i+3],h=(new $.Vector3).subVectors(o,n).multiplyScalar(.5),c=(new $.Vector3).subVectors(l,a).multiplyScalar(.5),!o.skip)for(var p=0;p<e;p++){var g=1/e*p,m=a.x+g*h.x+g*g*(-3*a.x+3*o.x-2*h.x-c.x)+g*g*g*(2*a.x-2*o.x+h.x+c.x),v=a.y+g*h.y+g*g*(-3*a.y+3*o.y-2*h.y-c.y)+g*g*g*(2*a.y-2*o.y+h.y+c.y),_=a.z+g*h.z+g*g*(-3*a.z+3*o.z-2*h.z-c.z)+g*g*g*(2*a.z-2*o.z+h.z+c.z),y=new $.Vector3(m,v,_);y.atom=p<e/2?a.atom:o.atom,d.push(y)}return d.push(u[u.length-1]),d}!function(t){t[t.NONE=0]="NONE",t[t.FLAT=1]="FLAT",t[t.ROUND=2]="ROUND"}(it||(it={})),function(t){function e(t,e,i){var r,s,n,a,o,l=Math.hypot(t,e);l<1e-4?(s=0,n=1):(s=-t/l,n=e/l),e=-s*t+n*e,(r=Math.hypot(e,i))<1e-4?(a=0,o=1):(a=i/r,o=e/r);var h=new Float32Array(9);return h[0]=n,h[1]=s,h[2]=0,h[3]=-s*o,h[4]=n*o,h[5]=a,h[6]=s*a,h[7]=-n*a,h[8]=o,h}var i=new class CylVertexCache{constructor(){this.cache={};let t,e=[],i=Math.pow(2,4),r=2,s=Math.pow(2,r),n=i/s;for(e[0]=new $.Vector3(-1,0,0),e[n]=new $.Vector3(0,0,1),e[2*n]=new $.Vector3(1,0,0),e[3*n]=new $.Vector3(0,0,-1),r=3;r<=4;r++){for(s=Math.pow(2,r-1),n=i/s,t=0;t<s-1;t++)e[n/2+t*n]=e[t*n].clone().add(e[(t+1)*n]).normalize();t=s-1,e[n/2+t*n]=e[t*n].clone().add(e[0]).normalize()}this.basisVectors=e}getVerticesForRadius(t,e,i){if(void 0!==this.cache&&void 0!==this.cache[t]&&void 0!==this.cache[t][e+i])return this.cache[t][e+i];for(var r,s=this.basisVectors.length,n=[],a=[],o=0;o<s;o++)n.push(this.basisVectors[o].clone().multiplyScalar(t)),n.push(this.basisVectors[o].clone().multiplyScalar(t)),r=this.basisVectors[o].clone().normalize(),a.push(r),a.push(r);var l=[],h=10,c=s;var d,u,f=2*Math.PI,p=Math.PI,g=!1,m=!1;for(u=0;u<=h;u++){g=0===u||u===h,m=5===u;var v=[],_=[];for(d=0;d<=c;d++)if(m){var y=d<c?2*d:0;_.push(y+1),v.push(y)}else{var b=d/c,x=u/h;if(g&&0!==d)g&&v.push(n.length-1);else if(d<c){var w=new $.Vector3;w.x=-t*Math.cos(0+b*f)*Math.sin(0+x*p),w.y=1==e?0:t*Math.cos(0+x*p),w.z=t*Math.sin(0+b*f)*Math.sin(0+x*p),Math.abs(w.x)<1e-5&&(w.x=0),Math.abs(w.y)<1e-5&&(w.y=0),Math.abs(w.z)<1e-5&&(w.z=0),e==it.FLAT?(r=new $.Vector3(0,Math.cos(0+x*p),0)).normalize():(r=new $.Vector3(w.x,w.y,w.z)).normalize(),n.push(w),a.push(r),v.push(n.length-1)}else v.push(n.length-c)}m&&l.push(_),l.push(v)}var A={vertices:n,normals:a,verticesRows:l,w:c,h};return t in this.cache||(this.cache[t]={}),this.cache[t][e+i]=A,A}};t.drawCylinder=function(t,r,s,n,a,o=0,l=0){if(!r||!s)return;let h=function(t){if("string"==typeof t){let e=t;return"flat"==e.toLowerCase()?it.FLAT:"round"==e.toLowerCase()?it.ROUND:it.NONE}return t};o=h(o);var c=(l=h(l))||o;a=a||{r:0,g:0,b:0};var d,u,f,p,g,m,v=e(s.x-r.x,s.y-r.y,s.z-r.z),_=i.getVerticesForRadius(n,l,"to"),y=_.w,b=_.h,x=c?b*y+2:2*y,w=t.updateGeoGroup(x),A=_.vertices,C=_.normals,S=_.verticesRows,M=S[b/2],z=S[b/2+1],T=w.vertices,E=w.vertexArray,L=w.normalArray,F=w.colorArray,I=w.faceArray;for(f=0;f<y;++f){var O=2*f;p=v[0]*A[O].x+v[3]*A[O].y+v[6]*A[O].z,g=v[1]*A[O].x+v[4]*A[O].y+v[7]*A[O].z,m=v[5]*A[O].y+v[8]*A[O].z,d=3*(T+O),u=w.faceidx,E[d]=p+r.x,E[d+1]=g+r.y,E[d+2]=m+r.z,E[d+3]=p+s.x,E[d+4]=g+s.y,E[d+5]=m+s.z,L[d]=p,L[d+3]=p,L[d+1]=g,L[d+4]=g,L[d+2]=m,L[d+5]=m,F[d]=a.r,F[d+3]=a.r,F[d+1]=a.g,F[d+4]=a.g,F[d+2]=a.b,F[d+5]=a.b,I[u]=z[f]+T,I[u+1]=z[f+1]+T,I[u+2]=M[f]+T,I[u+3]=M[f]+T,I[u+4]=z[f+1]+T,I[u+5]=M[f+1]+T,w.faceidx+=6}if(c){var D,k,R,P,U,B,N,G,V,j,H,W,q,Y,Z,X,K,Q,$,J,tt,et,rt,st,nt,at,ot,lt,ht,ct,dt,ut,ft=o?b+1:b/2+1;for(g=l?0:b/2;g<ft;g++)if(g!==b/2){var pt=g<=b/2?s:r,gt=i.getVerticesForRadius(n,l,"to"),mt=i.getVerticesForRadius(n,o,"from");for(pt===s?(A=gt.vertices,C=gt.normals,S=gt.verticesRows):pt==r&&(A=mt.vertices,C=mt.normals,S=mt.verticesRows),p=0;p<y;p++)u=w.faceidx,ht=3*((D=S[g][p+1])+T),ct=3*((k=S[g][p])+T),dt=3*((R=S[g+1][p])+T),ut=3*((P=S[g+1][p+1])+T),U=v[0]*A[D].x+v[3]*A[D].y+v[6]*A[D].z,B=v[0]*A[k].x+v[3]*A[k].y+v[6]*A[k].z,N=v[0]*A[R].x+v[3]*A[R].y+v[6]*A[R].z,G=v[0]*A[P].x+v[3]*A[P].y+v[6]*A[P].z,V=v[1]*A[D].x+v[4]*A[D].y+v[7]*A[D].z,j=v[1]*A[k].x+v[4]*A[k].y+v[7]*A[k].z,H=v[1]*A[R].x+v[4]*A[R].y+v[7]*A[R].z,W=v[1]*A[P].x+v[4]*A[P].y+v[7]*A[P].z,q=v[5]*A[D].y+v[8]*A[D].z,Y=v[5]*A[k].y+v[8]*A[k].z,Z=v[5]*A[R].y+v[8]*A[R].z,X=v[5]*A[P].y+v[8]*A[P].z,E[ht]=U+pt.x,E[ct]=B+pt.x,E[dt]=N+pt.x,E[ut]=G+pt.x,E[ht+1]=V+pt.y,E[ct+1]=j+pt.y,E[dt+1]=H+pt.y,E[ut+1]=W+pt.y,E[ht+2]=q+pt.z,E[ct+2]=Y+pt.z,E[dt+2]=Z+pt.z,E[ut+2]=X+pt.z,F[ht]=a.r,F[ct]=a.r,F[dt]=a.r,F[ut]=a.r,F[ht+1]=a.g,F[ct+1]=a.g,F[dt+1]=a.g,F[ut+1]=a.g,F[ht+2]=a.b,F[ct+2]=a.b,F[dt+2]=a.b,F[ut+2]=a.b,K=v[0]*C[D].x+v[3]*C[D].y+v[6]*C[D].z,Q=v[0]*C[k].x+v[3]*C[k].y+v[6]*C[k].z,$=v[0]*C[R].x+v[3]*C[R].y+v[6]*C[R].z,J=v[0]*C[P].x+v[3]*C[P].y+v[6]*C[P].z,tt=v[1]*C[D].x+v[4]*C[D].y+v[7]*C[D].z,et=v[1]*C[k].x+v[4]*C[k].y+v[7]*C[k].z,rt=v[1]*C[R].x+v[4]*C[R].y+v[7]*C[R].z,st=v[1]*C[P].x+v[4]*C[P].y+v[7]*C[P].z,nt=v[5]*C[D].y+v[8]*C[D].z,at=v[5]*C[k].y+v[8]*C[k].z,ot=v[5]*C[R].y+v[8]*C[R].z,lt=v[5]*C[P].y+v[8]*C[P].z,0===g?(L[ht]=K,L[dt]=$,L[ut]=J,L[ht+1]=tt,L[dt+1]=rt,L[ut+1]=st,L[ht+2]=nt,L[dt+2]=ot,L[ut+2]=lt,I[u]=D+T,I[u+1]=R+T,I[u+2]=P+T,w.faceidx+=3):g===ft-1?(L[ht]=K,L[ct]=Q,L[dt]=$,L[ht+1]=tt,L[ct+1]=et,L[dt+1]=rt,L[ht+2]=nt,L[ct+2]=at,L[dt+2]=ot,I[u]=D+T,I[u+1]=k+T,I[u+2]=R+T,w.faceidx+=3):(L[ht]=K,L[ct]=Q,L[ut]=J,L[ht+1]=tt,L[ct+1]=et,L[ut+1]=st,L[ht+2]=nt,L[ct+2]=at,L[ut+2]=lt,L[ct]=Q,L[dt]=$,L[ut]=J,L[ct+1]=et,L[dt+1]=rt,L[ut+1]=st,L[ct+2]=at,L[dt+2]=ot,L[ut+2]=lt,I[u]=D+T,I[u+1]=k+T,I[u+2]=P+T,I[u+3]=k+T,I[u+4]=R+T,I[u+5]=P+T,w.faceidx+=6)}}w.vertices+=x},t.drawCone=function(t,r,s,n,a){if(!r||!s)return;a=a||{r:0,g:0,b:0};let o=new $.Vector3(s.x-r.x,s.y-r.y,s.z-r.z);var l=e(o.x,o.y,o.z);o=o.normalize();var h,c,d,u,f,p,g=i.basisVectors.length,m=i.basisVectors,v=g+2,_=t.updateGeoGroup(v),y=_.vertices,b=_.vertexArray,x=_.normalArray,w=_.colorArray,A=_.faceArray;for(b[h=3*y]=r.x,b[h+1]=r.y,b[h+2]=r.z,x[h]=-o.x,x[h+1]=-o.y,x[h+2]=-o.z,w[h]=a.r,w[h+1]=a.g,w[h+2]=a.b,b[h+3]=s.x,b[h+4]=s.y,b[h+5]=s.z,x[h+3]=o.x,x[h+4]=o.y,x[h+5]=o.z,w[h+3]=a.r,w[h+4]=a.g,w[h+5]=a.b,h+=6,d=0;d<g;++d){var C=m[d].clone();C.multiplyScalar(n),u=l[0]*C.x+l[3]*C.y+l[6]*C.z,f=l[1]*C.x+l[4]*C.y+l[7]*C.z,p=l[5]*C.y+l[8]*C.z,b[h]=u+r.x,b[h+1]=f+r.y,b[h+2]=p+r.z,x[h]=u,x[h+1]=f,x[h+2]=p,w[h]=a.r,w[h+1]=a.g,w[h+2]=a.b,h+=3}for(_.vertices+=g+2,c=_.faceidx,d=0;d<g;d++){var S=y+2+d,M=y+2+(d+1)%g;A[c]=S,A[c+1]=M,A[c+2]=y,A[c+=3]=S,A[c+1]=M,A[c+2]=y+1,c+=3}_.faceidx+=6*g};var r=new class SphereVertexCache{constructor(){this.cache=new Map}getVerticesForRadius(t,e){e=e||2,this.cache.has(e)||this.cache.set(e,new Map);let i=this.cache.get(e);if(i.has(t))return i.get(t);var r={vertices:[],verticesRows:[],normals:[]},s=16*e,n=10*e;t<1&&(s=10*e,n=8*e);var a,o,l=2*Math.PI,h=Math.PI;for(o=0;o<=n;o++){let e=[];for(a=0;a<=s;a++){let i=a/s,d=o/n,u=-t*Math.cos(0+i*l)*Math.sin(0+d*h),f=t*Math.cos(0+d*h),p=t*Math.sin(0+i*l)*Math.sin(0+d*h);var c=new $.Vector3(u,f,p);c.normalize(),r.vertices.push({x:u,y:f,z:p}),r.normals.push(c),e.push(r.vertices.length-1)}r.verticesRows.push(e)}return i.set(t,r),r}};t.drawSphere=function(t,e,i,s,n){var a=r.getVerticesForRadius(i,n),o=a.vertices,l=a.normals,h=t.updateGeoGroup(o.length),c=h.vertices,d=h.vertexArray,u=h.colorArray,f=h.faceArray,p=h.lineArray,g=h.normalArray;for(let t=0,i=o.length;t<i;++t){let i=3*(c+t),r=o[t];d[i]=r.x+e.x,d[i+1]=r.y+e.y,d[i+2]=r.z+e.z,u[i]=s.r,u[i+1]=s.g,u[i+2]=s.b}h.vertices+=o.length;let m=a.verticesRows,v=m.length-1;for(let t=0;t<v;t++){let e=m[t].length-1;for(let r=0;r<e;r++){let e=h.faceidx,s=h.lineidx,n=m[t][r+1]+c,a=3*n,d=m[t][r]+c,u=3*d,v=m[t+1][r]+c,_=3*v,y=m[t+1][r+1]+c,b=3*y,x=l[n-c],w=l[d-c],A=l[v-c],C=l[y-c];Math.abs(o[n-c].y)===i?(g[a]=x.x,g[_]=A.x,g[b]=C.x,g[a+1]=x.y,g[_+1]=A.y,g[b+1]=C.y,g[a+2]=x.z,g[_+2]=A.z,g[b+2]=C.z,f[e]=n,f[e+1]=v,f[e+2]=y,p[s]=n,p[s+1]=v,p[s+2]=n,p[s+3]=y,p[s+4]=v,p[s+5]=y,h.faceidx+=3,h.lineidx+=6):Math.abs(o[v-c].y)===i?(g[a]=x.x,g[u]=w.x,g[_]=A.x,g[a+1]=x.y,g[u+1]=w.y,g[_+1]=A.y,g[a+2]=x.z,g[u+2]=w.z,g[_+2]=A.z,f[e]=n,f[e+1]=d,f[e+2]=v,p[s]=n,p[s+1]=d,p[s+2]=n,p[s+3]=v,p[s+4]=d,p[s+5]=v,h.faceidx+=3,h.lineidx+=6):(g[a]=x.x,g[u]=w.x,g[b]=C.x,g[a+1]=x.y,g[u+1]=w.y,g[b+1]=C.y,g[a+2]=x.z,g[u+2]=w.z,g[b+2]=C.z,g[u]=w.x,g[_]=A.x,g[b]=C.x,g[u+1]=w.y,g[_+1]=A.y,g[b+1]=C.y,g[u+2]=w.z,g[_+2]=A.z,g[b+2]=C.z,f[e]=n,f[e+1]=d,f[e+2]=y,f[e+3]=d,f[e+4]=v,f[e+5]=y,p[s]=n,p[s+1]=d,p[s+2]=n,p[s+3]=y,p[s+4]=d,p[s+5]=v,p[s+6]=v,p[s+7]=y,h.faceidx+=6,h.lineidx+=8)}}}}(rt||(rt={}));const at=.5,ot=1.3,lt=.8,ht=.4,ct=.4;function dt(t,e,i,r){for(var n,a,o,l,h=0,c=e.length;h<c;h++){l=Math.round(h*(r.length-1)/c),o=s.CC.color(r[l]);var d=t.updateGeoGroup(2),u=d.vertexArray,f=d.colorArray,p=d.faceArray;u[a=3*(n=d.vertices)]=e[h].x,u[a+1]=e[h].y,u[a+2]=e[h].z,u[a+3]=i[h].x,u[a+4]=i[h].y,u[a+5]=i[h].z;for(var g=0;g<6;++g)f[a+3*g]=o.r,f[a+1+3*g]=o.g,f[a+2+3*g]=o.b;if(h>0){var m=[n,n+1,n-1,n-2],v=d.faceidx;p[v]=m[0],p[v+1]=m[1],p[v+2]=m[3],p[v+3]=m[1],p[v+4]=m[2],p[v+5]=m[3],d.faceidx+=6}d.vertices+=2}}function ut(t,e,i,r,n,a,o){o&&"default"!==o||(o="rectangle"),"edged"===o?function(t,e,i,r,n){if(!(e.length<2)){var a,o;if(a=e[0],o=e[e.length-1],a=nt(a,r),o=nt(o,r),!n)return dt(t,a,o,i);var l,h,c,d,u,f,p,g,m,v,_,y,b,x,w,A,C,S,M,z,T,E,L=[],F=[[0,2,-6,-8],[-4,-2,6,4],[7,-1,-5,3],[-3,5,1,-7]];for(b=0,x=a.length;b<x;b++){if(v=Math.round(b*(i.length-1)/x),m=s.CC.color(i[v]),L.push(h=a[b]),L.push(h),L.push(c=o[b]),L.push(c),b<x-1){var I=a[b+1].clone().sub(a[b]);l=o[b].clone().sub(a[b]).cross(I).normalize().multiplyScalar(n)}for(L.push(d=a[b].clone().add(l)),L.push(d),L.push(u=o[b].clone().add(l)),L.push(u),void 0!==h.atom&&(_=h.atom),z=(M=t.updateGeoGroup(8)).vertexArray,T=M.colorArray,E=M.faceArray,z[p=3*(f=M.vertices)]=h.x,z[p+1]=h.y,z[p+2]=h.z,z[p+3]=h.x,z[p+4]=h.y,z[p+5]=h.z,z[p+6]=c.x,z[p+7]=c.y,z[p+8]=c.z,z[p+9]=c.x,z[p+10]=c.y,z[p+11]=c.z,z[p+12]=d.x,z[p+13]=d.y,z[p+14]=d.z,z[p+15]=d.x,z[p+16]=d.y,z[p+17]=d.z,z[p+18]=u.x,z[p+19]=u.y,z[p+20]=u.z,z[p+21]=u.x,z[p+22]=u.y,z[p+23]=u.z,w=0;w<8;++w)T[p+3*w]=m.r,T[p+1+3*w]=m.g,T[p+2+3*w]=m.b;if(b>0){var O=void 0!==y&&void 0!==_&&y.serial!==_.serial;for(w=0;w<4;w++){var D=[f+F[w][0],f+F[w][1],f+F[w][2],f+F[w][3]];if(E[g=M.faceidx]=D[0],E[g+1]=D[1],E[g+2]=D[3],E[g+3]=D[1],E[g+4]=D[2],E[g+5]=D[3],M.faceidx+=6,_.clickable||y.clickable||_.hoverable||y.hoverable){var k=L[D[3]].clone(),R=L[D[0]].clone(),P=L[D[2]].clone(),U=L[D[1]].clone();if(k.atom=L[D[3]].atom||null,P.atom=L[D[2]].atom||null,R.atom=L[D[0]].atom||null,U.atom=L[D[1]].atom||null,O){var B=k.clone().add(R).multiplyScalar(.5),N=P.clone().add(U).multiplyScalar(.5),G=k.clone().add(U).multiplyScalar(.5);w%2==0?((y.clickable||y.hoverable)&&(A=new J.Triangle(B,G,k),C=new J.Triangle(N,P,G),S=new J.Triangle(G,P,k),y.intersectionShape.triangle.push(A),y.intersectionShape.triangle.push(C),y.intersectionShape.triangle.push(S)),(_.clickable||_.hoverable)&&(A=new J.Triangle(R,U,G),C=new J.Triangle(U,N,G),S=new J.Triangle(R,G,B),_.intersectionShape.triangle.push(A),_.intersectionShape.triangle.push(C),_.intersectionShape.triangle.push(S))):((_.clickable||_.hoverable)&&(A=new J.Triangle(B,G,k),C=new J.Triangle(N,P,G),S=new J.Triangle(G,P,k),_.intersectionShape.triangle.push(A),_.intersectionShape.triangle.push(C),_.intersectionShape.triangle.push(S)),(y.clickable||y.hoverable)&&(A=new J.Triangle(R,U,G),C=new J.Triangle(U,N,G),S=new J.Triangle(R,G,B),y.intersectionShape.triangle.push(A),y.intersectionShape.triangle.push(C),y.intersectionShape.triangle.push(S)))}else(_.clickable||_.hoverable)&&(A=new J.Triangle(R,U,k),C=new J.Triangle(U,P,k),_.intersectionShape.triangle.push(A),_.intersectionShape.triangle.push(C))}}}M.vertices+=8,y=_}var V=L.length-8;for(z=(M=t.updateGeoGroup(8)).vertexArray,T=M.colorArray,E=M.faceArray,p=3*(f=M.vertices),g=M.faceidx,b=0;b<4;b++){L.push(L[2*b]),L.push(L[V+2*b]);var j=L[2*b],H=L[V+2*b];z[p+6*b]=j.x,z[p+1+6*b]=j.y,z[p+2+6*b]=j.z,z[p+3+6*b]=H.x,z[p+4+6*b]=H.y,z[p+5+6*b]=H.z,T[p+6*b]=m.r,T[p+1+6*b]=m.g,T[p+2+6*b]=m.b,T[p+3+6*b]=m.r,T[p+4+6*b]=m.g,T[p+5+6*b]=m.b}V+=8,A=[f,f+2,f+6,f+4],C=[f+1,f+5,f+7,f+3],E[g]=A[0],E[g+1]=A[1],E[g+2]=A[3],E[g+3]=A[1],E[g+4]=A[2],E[g+5]=A[3],E[g+6]=C[0],E[g+7]=C[1],E[g+8]=C[3],E[g+9]=C[1],E[g+10]=C[2],E[g+11]=C[3],M.faceidx+=12,M.vertices+=8}}(t,e,i,r,n):"rectangle"!==o&&"oval"!==o&&"parabola"!==o||function(t,e,i,r,n,a,o){var l,h,c,d,u,f,p,g,m,v;if((c=e.length)<2||e[0].length<2)return;for(l=0;l<c;l++)e[l]=nt(e[l],r);if(d=e[0].length,!n)return dt(t,e[0],e[c-1],i);var _=[],y=[],b=[];for(h=0;h<c;h++)_.push(.25+1.5*Math.sqrt((c-1)*h-Math.pow(h,2))/(c-1)),y.push(.5),b.push(2*(Math.pow(h/c,2)-h/c)+.6);var x,w,A,C,S,M,z,T,E,L,F=[];for(h=0;h<2*c-1;h++)F[h]=[h,h+1,h+1-2*c,h-2*c];F[2*c-1]=[h,h+1-2*c,h+1-4*c,h-2*c];let I=t.updateGeoGroup();for(l=0;l<d;l++){let r=t.groups,a=!1;var O,D;for(I=t.updateGeoGroup(2*c),r!=t.groups&&l>0&&(l-=1,a=!0),M=Math.round(l*(i.length-1)/d),S=s.CC.color(i[M]),m=p,v=g,p=[],g=[],u=[],void 0!==e[0][l].atom&&(C=e[0][l].atom,"oval"===o?f=_:"rectangle"===o?f=y:"parabola"===o&&(f=b)),f||(f=y),h=0;h<c;h++)O=l<d-1?e[h][l+1].clone().sub(e[h][l]):e[h][l-1].clone().sub(e[h][l]).negate(),D=h<c-1?e[h+1][l].clone().sub(e[h][l]):e[h-1][l].clone().sub(e[h][l]).negate(),u[h]=D.cross(O).normalize().multiplyScalar(n*f[h]);for(h=0;h<c;h++)p[h]=e[h][l].clone().add(u[h].clone().negate());for(h=0;h<c;h++)g[h]=e[h][l].clone().add(u[h]);for(z=I.vertexArray,T=I.colorArray,E=I.faceArray,w=3*(x=I.vertices),h=0;h<c;h++)z[w+3*h+0]=p[h].x,z[w+3*h+1]=p[h].y,z[w+3*h+2]=p[h].z;for(h=0;h<c;h++)z[w+3*h+0+3*c]=g[c-1-h].x,z[w+3*h+1+3*c]=g[c-1-h].y,z[w+3*h+2+3*c]=g[c-1-h].z;for(h=0;h<2*c;++h)T[w+3*h+0]=S.r,T[w+3*h+1]=S.g,T[w+3*h+2]=S.b;if(l>0&&!a){for(h=0;h<2*c;h++)L=[x+F[h][0],x+F[h][1],x+F[h][2],x+F[h][3]],E[A=I.faceidx]=L[0],E[A+1]=L[1],E[A+2]=L[3],E[A+3]=L[1],E[A+4]=L[2],E[A+5]=L[3],I.faceidx+=6;if(C.clickable||C.hoverable){var k=[];for(h in k.push(new J.Triangle(m[0],p[0],p[c-1])),k.push(new J.Triangle(m[0],p[c-1],m[c-1])),k.push(new J.Triangle(m[c-1],p[c-1],g[c-1])),k.push(new J.Triangle(m[c-1],g[c-1],v[c-1])),k.push(new J.Triangle(g[0],v[0],v[c-1])),k.push(new J.Triangle(g[c-1],g[0],v[c-1])),k.push(new J.Triangle(p[0],m[0],v[0])),k.push(new J.Triangle(g[0],p[0],v[0])),k)C.intersectionShape.triangle.push(k[h])}}I.vertices+=2*c}for(z=I.vertexArray,T=I.colorArray,E=I.faceArray,w=3*(x=I.vertices),A=I.faceidx,l=0;l<c-1;l++)L=[l,l+1,2*c-2-l,2*c-1-l],E[A=I.faceidx]=L[0],E[A+1]=L[1],E[A+2]=L[3],E[A+3]=L[1],E[A+4]=L[2],E[A+5]=L[3],I.faceidx+=6;for(l=0;l<c-1;l++)L=[x-1-l,x-2-l,x-2*c+l+1,x-2*c+l],E[A=I.faceidx]=L[0],E[A+1]=L[1],E[A+2]=L[3],E[A+3]=L[1],E[A+4]=L[2],E[A+5]=L[3],I.faceidx+=6}(t,e,i,r,n,0,o)}function ft(t){return t&&"C"===t.elem&&"CA"===t.atom}function pt(t,e){if(t&&e&&t.chain===e.chain){if(!(t.hetflag||e.hetflag||t.reschain!==e.reschain||t.resi!==e.resi&&t.resi!==e.resi-1))return!0;if(t.resi<e.resi){var i=t.x-e.x,r=t.y-e.y,s=t.z-e.z,n=i*i+r*r+s*s;if("CA"==t.atom&&"CA"==e.atom&&n<16)return!0;if(("P"==t.atom||"P"==e.atom)&&n<64)return!0}}return!1}function gt(t,e,i,r,s){if(null!=e&&0!=e.vertices){s&&(e.initTypedArrays(),e.setUpNormals());var a=new n.MeshDoubleLambertMaterial;a.vertexColors=n.Coloring.FaceColors,"number"==typeof i&&i>=0&&i<1&&(a.transparent=!0,a.opacity=i),a.outline=r;var o=new n.Mesh(e,a);t.add(o)}}function mt(t,e,i,r,s,n,a,o,l){var h,c,d,u,f,p;if(r&&s&&a){var g=s.sub(r);g.normalize();var m=o[l];for(c=l+1;c<o.length&&(m=o[c]).atom!=a.atom;c++);if((m=m?new $.Vector3(m.x,m.y,m.z):new $.Vector3(0,0,0)).sub(r),"arrow start"===a.ss){var v=m.clone().multiplyScalar(.3).cross(s);r.add(v);var _=m.clone().cross(g).normalize();g.rotateAboutVector(_,.43)}for(a.style.cartoon.ribbon?h=a.style.cartoon.thickness||ht:a.style.cartoon.width?h=a.style.cartoon.width:"c"===a.ss?h="P"===a.atom?lt:at:"arrow start"===a.ss?(h=ot,f=!0):h="arrow end"===a.ss||"h"===a.ss&&a.style.cartoon.tubes||"tube start"===a.ss?at:ot,null!=n&&g.dot(n)<0&&g.negate(),g.multiplyScalar(h),c=0;c<e;c++)d=2*c/(e-1)-1,(u=new $.Vector3(r.x+d*g.x,r.y+d*g.y,r.z+d*g.z)).atom=a,i&&"s"===a.ss&&(u.smoothen=!0),t[c].push(u);if(f)for(g.multiplyScalar(2),c=0;c<e;c++)d=2*c/(e-1)-1,(u=new $.Vector3(r.x+d*g.x,r.y+d*g.y,r.z+d*g.z)).atom=a,u.smoothen=!1,u.skip=!0,t[c].push(u);return p=a.style.cartoon.style||"default",t.style?t.style!=p&&(console.log("Warning: a cartoon chain's strand-style is ambiguous"),t.style="default"):t.style=p,"arrow start"!==a.ss&&"arrow end"!==a.ss||(a.ss="s"),f}}const vt={C:!0,CA:!0,O:!0,P:!0,OP2:!0,O2P:!0,"O5'":!0,"O3'":!0,"C5'":!0,"C2'":!0,"O5*":!0,"O3*":!0,"C5*":!0,"C2*":!0,N1:!0,N3:!0},_t={DA:!0,DG:!0,A:!0,G:!0},yt={DT:!0,DC:!0,U:!0,C:!0,T:!0},bt={DA:!0,DG:!0,A:!0,G:!0,DT:!0,DC:!0,U:!0,C:!0,T:!0};function xt(t,e,i,a=10){let o=a,l=a;var h,c,d,u,f,p,g,m,v,_,y,b,x,w,A,C,S,M,z=new n.Geometry(!0),T=new n.Geometry(!0),E=[],F=[],I=1,O=!1,D={};for(var k in r.Gradient.builtinGradients)r.Gradient.builtinGradients.hasOwnProperty(k)&&(D[k]=new r.Gradient.builtinGradients[k](i[1],i[0]));var R=function(t,e){return i&&"spectrum"===e.color?e.colorscheme in D?D[e.colorscheme].valueToHex(t.resi):D.sinebow.valueToHex(t.resi):(0,L.getColorFromStyle)(t,e).getHex()};for(m=0;m<o;m++)F[m]=[];var P=!1,U=!1,B=[];for(m in e){if("C"===(u=e[m]).elem&&"CA"===u.atom){var N=pt(d,u);N&&"s"===u.ss?P=!0:P&&(d&&c&&d.style.cartoon.arrows&&c.style.cartoon.arrows&&(d.ss="arrow end",c.ss="arrow start"),P=!1),N&&("h"===d.ss||"tube start"==d.ss)&&d.style.cartoon.tubes?!U&&"tube start"!=d.ss&&u.style.cartoon.tubes&&(u.ss="tube start",U=!0):U&&("tube start"===d.ss?d.ss="tube end":c&&c.style.cartoon.tubes&&(c.ss="tube end"),U=!1),c=d,d=u}u&&u.atom in vt&&B.push(u)}U&&d.style.cartoon.tubes&&(d.ss="tube end",U=!1);var G=function(e){F[0].length>0&&ut(T,F,E,l,g,0,F.style);var i=[],r=null;if(e){for(m=0;m<o;m++)i[m]=F[m][F[m].length-1];r=E[E.length-1]}for(F=[],m=0;m<o;m++)F[m]=[];if(E=[],e){for(m=0;m<o;m++)F[m].push(i[m]);E.push(r)}gt(t,T,I,O,!0),gt(t,z,I,O,!1),T=new n.Geometry(!0),z=new n.Geometry(!0)};d=void 0;for(var V=0;V<B.length;V++){var j=(u=B[V]).resn.trim(),H=j in bt;if(I=1,h=u.style.cartoon,d&&d.style.cartoon&&(I=d.style.cartoon.opacity),d&&d.style.cartoon&&d.style.cartoon.outline&&(O=d.style.cartoon.outline),!d||!d.style.cartoon||u.style.cartoon&&d.style.cartoon.opacity==u.style.cartoon.opacity||G(d.chain==u.chain),"trace"===h.style){if(u.hetflag);else if("C"===u.elem&&"CA"===u.atom||H&&"P"===u.atom||"BB"===u.atom){if(p=R(u,h),g=(0,L.isNumeric)(h.thickness)?h.thickness:ht,pt(d,u))if(p==f){var W=s.CC.color(p);rt.drawCylinder(z,d,u,g,W,2,2)}else{var q=(new $.Vector3).addVectors(d,u).multiplyScalar(.5),Y=s.CC.color(f),Z=s.CC.color(p);rt.drawCylinder(z,d,q,g,Y,2,0),rt.drawCylinder(z,q,u,g,Z,0,2)}if((!0===u.clickable||u.hoverable)&&void 0!==u.intersectionShape){var X=new $.Vector3(u.x,u.y,u.z);u.intersectionShape.sphere.push(new J.Sphere(X,g))}d=u,f=p}}else{if(ft(u)||H&&("P"===u.atom||0==u.atom.indexOf("O5"))){if(M)if("tube end"===u.ss)M=!1,S=new $.Vector3(u.x,u.y,u.z),rt.drawCylinder(z,C,S,2,s.CC.color(f),1,1),u.ss="h";else{if(d.chain==u.chain&&"tube end"!==d.ss)continue;M=!1,d.ss="h",S=new $.Vector3(d.x,d.y,d.z),rt.drawCylinder(z,C,S,2,s.CC.color(f),1,1)}if(d&&(!pt(d,u)||"tube start"===d.ss)){for("tube start"===d.ss&&(M=!0,C=new $.Vector3(d.x,d.y,d.z),d.ss="h"),A&&(w=b?(new $.Vector3).addVectors(d,b).multiplyScalar(.5):new $.Vector3(d.x,d.y,d.z),rt.drawCylinder(z,w,A,ct,s.CC.color(A.color),0,2),mt(F,o,!0,b,x,y,d,B,V),E.push(p),w=null,A=null),F[0].length>0&&ut(T,F,E,l,g,0,F.style),F=[],m=0;m<o;m++)F[m]=[];E=[]}if(void 0===d||d.rescode!=u.rescode||d.resi!=u.resi){if(A&&null!=d){var K=(w=(new $.Vector3).addVectors(d,u).multiplyScalar(.5)).clone().sub(A).multiplyScalar(.02);w.add(K),rt.drawCylinder(z,w,A,ct,s.CC.color(A.color),0,2),w=null,A=null}p=R(u,h),E.push(p),g=(0,L.isNumeric)(h.thickness)?h.thickness:ht,d=u,(v=new $.Vector3(d.x,d.y,d.z)).resi=d.resi,f=p}!0!==u.clickable&&!0!==u.hoverable||void 0!==u.intersectionShape&&void 0!==u.intersectionShape.triangle||(u.intersectionShape={sphere:null,cylinder:[],line:[],triangle:[]})}else null!=d&&(ft(d)&&"O"===u.atom||H&&"P"===d.atom&&("OP2"===u.atom||"O2P"===u.atom)||H&&0==d.atom.indexOf("O5")&&0==u.atom.indexOf("C5"))?((_=new $.Vector3(u.x,u.y,u.z)).resi=u.resi,"OP2"!==u.atom&&"O2P"!==u.atom||(x=new $.Vector3(u.x,u.y,u.z))):H&&0==u.atom.indexOf("O3")?b=new $.Vector3(u.x,u.y,u.z):("N1"===u.atom&&j in _t||"N3"===u.atom&&j in yt)&&((A=new $.Vector3(u.x,u.y,u.z)).color=(0,L.getColorFromStyle)(u,h).getHex());_&&v&&_.resi===v.resi&&(mt(F,o,!0,v,_,y,d,B,V),y=_,v=null,_=null,E.push(p))}}A&&(w=b?(new $.Vector3).addVectors(d,b).multiplyScalar(.5):new $.Vector3(d.x,d.y,d.z),rt.drawCylinder(z,w,A,ct,s.CC.color(A.color),0,2),mt(F,o,!0,b,x,y,d,B,V),E.push(p)),G(!1)}class GLShape{static finalizeGeo(t){var e=t.updateGeoGroup(0);e.vertices>0&&e.truncateArrayBuffers(!0,!0)}static updateColor(t,e){var i,r,n;e=e||s.CC.color(e),t.colorsNeedUpdate=!0,e.constructor!==Array&&(i=e.r,r=e.g,n=e.b);for(let s in t.geometryGroups){let a=t.geometryGroups[s],o=a.colorArray;for(let t=0,s=a.vertices;t<s;++t){if(e.constructor===Array){let s=e[t];i=s.r,r=s.g,n=s.b}o[3*t]=i,o[3*t+1]=r,o[3*t+2]=n}}}static drawArrow(t,e,i){var r=i.start,s=i.end,n=i.radius,a=i.radiusRatio,o=i.mid,l=i.midpos;if(!r||!s)return;var h=e.updateGeoGroup(51),c=new $.Vector3(s.x,s.y,s.z).sub(r);if(l){let t=c.length();o=l>0?l/t:(t+l)/t}c.multiplyScalar(o);var d=new $.Vector3(r.x,r.y,r.z).add(c),u=c.clone().negate();let f=new $.Vector3(r.x,r.y,r.z);t.intersectionShape.cylinder.push(new J.Cylinder(f,d.clone(),n)),t.intersectionShape.sphere.push(new J.Sphere(f,n));var p=[];p[0]=c.clone(),Math.abs(p[0].x)>1e-4?p[0].y+=1:p[0].x+=1,p[0].cross(c),p[0].normalize(),p[4]=p[0].clone(),p[4].crossVectors(p[0],c),p[4].normalize(),p[8]=p[0].clone().negate(),p[12]=p[4].clone().negate(),p[2]=p[0].clone().add(p[4]).normalize(),p[6]=p[4].clone().add(p[8]).normalize(),p[10]=p[8].clone().add(p[12]).normalize(),p[14]=p[12].clone().add(p[0]).normalize(),p[1]=p[0].clone().add(p[2]).normalize(),p[3]=p[2].clone().add(p[4]).normalize(),p[5]=p[4].clone().add(p[6]).normalize(),p[7]=p[6].clone().add(p[8]).normalize(),p[9]=p[8].clone().add(p[10]).normalize(),p[11]=p[10].clone().add(p[12]).normalize(),p[13]=p[12].clone().add(p[14]).normalize(),p[15]=p[14].clone().add(p[0]).normalize();var g,m,v,_,y,b,x,w,A,C,S,M,z,T,E,L,F,I,O,D,k,R,P=h.vertices,U=h.vertexArray,B=h.faceArray,N=h.normalArray,G=h.lineArray;for(m=0,v=p.length;m<v;++m){g=3*(P+3*m);var V=p[m].clone().multiplyScalar(n).add(r),j=p[m].clone().multiplyScalar(n).add(d),H=p[m].clone().multiplyScalar(n*a).add(d);if(U[g]=V.x,U[g+1]=V.y,U[g+2]=V.z,U[g+3]=j.x,U[g+4]=j.y,U[g+5]=j.z,U[g+6]=H.x,U[g+7]=H.y,U[g+8]=H.z,m>0){var W=U[g-3],q=U[g-2],Y=U[g-1],Z=new $.Vector3(W,q,Y),X=new $.Vector3(s.x,s.y,s.z),K=d.clone(),Q=new $.Vector3(H.x,H.y,H.z);t.intersectionShape.triangle.push(new J.Triangle(Q,X,Z)),t.intersectionShape.triangle.push(new J.Triangle(Z.clone(),K,Q.clone()))}}h.vertices+=48,U[g=3*h.vertices]=r.x,U[g+1]=r.y,U[g+2]=r.z,U[g+3]=d.x,U[g+4]=d.y,U[g+5]=d.z,U[g+6]=s.x,U[g+7]=s.y,U[g+8]=s.z,h.vertices+=3;var tt=h.vertices-3,et=h.vertices-2,it=h.vertices-1,rt=3*tt,st=3*et,nt=3*it;for(m=0,v=p.length-1;m<v;++m){var at=P+3*m;g=3*at,y=h.faceidx,b=h.lineidx,z=3*(x=at),T=3*(w=at+1),E=3*(A=at+2),L=3*(C=at+4),F=3*(S=at+5),I=3*(M=at+3),O=D=p[m],k=R=p[m+1],N[z]=O.x,N[T]=D.x,N[I]=R.x,N[z+1]=O.y,N[T+1]=D.y,N[I+1]=R.y,N[z+2]=O.z,N[T+2]=D.z,N[I+2]=R.z,N[T]=D.x,N[L]=k.x,N[I]=R.x,N[T+1]=D.y,N[L+1]=k.y,N[I+1]=R.y,N[T+2]=D.z,N[L+2]=k.z,N[I+2]=R.z,N[E]=D.x,N[F]=k.x,N[E+1]=D.y,N[F+1]=k.y,N[E+2]=D.z,N[F+2]=k.z,B[y]=x,B[y+1]=w,B[y+2]=M,B[y+3]=w,B[y+4]=C,B[y+5]=M,B[y+6]=x,B[y+7]=M,B[y+8]=tt,B[y+9]=A,B[y+10]=et,B[y+11]=S,B[y+12]=A,B[y+13]=it,B[y+14]=S,G[b]=x,G[b+1]=w,G[b+2]=x,G[b+3]=M,G[b+4]=C,G[b+5]=M,G[b+6]=x,G[b+7]=M,G[b+8]=A,G[b+9]=w,G[b+10]=A,G[b+11]=S,G[b+12]=C,G[b+13]=S,G[b+14]=A,G[b+15]=it,G[b+16]=A,G[b+17]=S,G[b+18]=it,G[b+19]=S,h.faceidx+=15,h.lineidx+=20}_=[P+45,P+46,P+1,P,P+47,P+2],y=h.faceidx,b=h.lineidx,z=3*(x=_[0]),T=3*(w=_[1]),E=3*(A=_[4]),L=3*(C=_[2]),F=3*(S=_[5]),I=3*(M=_[3]),O=D=p[15],k=R=p[0],N[z]=O.x,N[T]=D.x,N[I]=R.x,N[z+1]=O.y,N[T+1]=D.y,N[I+1]=R.y,N[z+2]=O.z,N[T+2]=D.z,N[I+2]=R.z,N[T]=D.x,N[L]=k.x,N[I]=R.x,N[T+1]=D.y,N[L+1]=k.y,N[I+1]=R.y,N[T+2]=D.z,N[L+2]=k.z,N[I+2]=R.z,N[E]=D.x,N[F]=k.x,N[E+1]=D.y,N[F+1]=k.y,N[E+2]=D.z,N[F+2]=k.z,c.normalize(),u.normalize(),N[rt]=u.x,N[st]=N[nt]=c.x,N[rt+1]=u.y,N[st+1]=N[nt+1]=c.y,N[rt+2]=u.z,N[st+2]=N[nt+2]=c.z,B[y]=x,B[y+1]=w,B[y+2]=M,B[y+3]=w,B[y+4]=C,B[y+5]=M,B[y+6]=x,B[y+7]=M,B[y+8]=tt,B[y+9]=A,B[y+10]=et,B[y+11]=S,B[y+12]=A,B[y+13]=it,B[y+14]=S,G[b]=x,G[b+1]=w,G[b+2]=x,G[b+3]=M,G[b+4]=C,G[b+5]=M,G[b+6]=x,G[b+7]=M,G[b+8]=A,G[b+9]=w,G[b+10]=A,G[b+11]=S,G[b+12]=C,G[b+13]=S,G[b+14]=A,G[b+15]=it,G[b+16]=A,G[b+17]=S,G[b+18]=it,G[b+19]=S,h.faceidx+=15,h.lineidx+=20}static updateBoundingFromPoints(t,e,i,r){t.center.set(0,0,0);let s=1/0,n=1/0,a=1/0,o=-1/0,l=-1/0,h=-1/0;t.box&&(s=t.box.min.x,o=t.box.max.x,n=t.box.min.y,l=t.box.max.y,a=t.box.min.z,h=t.box.max.z);for(let t=0,e=r;t<e;t++){var c=i[3*t],d=i[3*t+1],u=i[3*t+2];c<s&&(s=c),d<n&&(n=d),u<a&&(a=u),c>o&&(o=c),d>l&&(l=d),u>h&&(h=u)}t.center.set((o+s)/2,(l+n)/2,(h+a)/2),t.radius=t.center.distanceTo({x:o,y:l,z:h}),t.box={min:{x:s,y:n,z:a},max:{x:o,y:l,z:h}}}static addCustomGeo(t,e,i,r,s){var n,a,o,l,h,c,d,u,f,p=e.addGeoGroup(),g=i.vertexArr,m=i.normalArr,v=i.faceArr;p.vertices=g.length,p.faceidx=v.length;var _=p.vertexArray,y=p.colorArray;for(r.constructor!==Array&&(u=r.r,f=r.g,l=r.b),c=0,d=p.vertices;c<d;++c)n=3*c,a=g[c],_[n]=a.x,_[n+1]=a.y,_[n+2]=a.z,r.constructor===Array&&(u=(h=r[c]).r,f=h.g,l=h.b),y[n]=u,y[n+1]=f,y[n+2]=l;if(s)for(c=0,d=p.faceidx/3;c<d;++c){o=v[n=3*c],l=v[n+1],h=v[n+2];var b=new $.Vector3,x=new $.Vector3,w=new $.Vector3;t.intersectionShape.triangle.push(new J.Triangle(b.copy(g[o]),x.copy(g[l]),w.copy(g[h])))}if(s){var A=new $.Vector3(0,0,0),C=0;for(let t=0;t<e.geometryGroups.length;t++)A.add(e.geometryGroups[t].getCentroid()),C++;A.divideScalar(C),GLShape.updateBoundingFromPoints(t.boundingSphere,{centroid:A},_,p.vertices)}if(p.faceArray=new Uint16Array(v),p.truncateArrayBuffers(!0,!0),m.length<p.vertices)p.setNormals();else{var S,M=p.normalArray=new Float32Array(3*p.vertices);for(c=0,d=p.vertices;c<d;++c)n=3*c,S=m[c],M[n]=S.x,M[n+1]=S.y,M[n+2]=S.z}p.setLineIndices(),p.lineidx=p.lineArray.length}static updateFromStyle(t,e){void 0!==e.color?(t.color=e.color||new s.Color,e.color instanceof s.Color||(t.color=s.CC.color(e.color))):t.color=s.CC.color(0),t.wireframe=!!e.wireframe,t.opacity=e.alpha?(0,$.clamp)(e.alpha,0,1):1,void 0!==e.opacity&&(t.opacity=(0,$.clamp)(e.opacity,0,1)),t.side=void 0!==e.side?e.side:n.DoubleSide,t.linewidth=void 0===e.linewidth?1:e.linewidth,t.clickable=!!e.clickable,t.callback=(0,L.makeFunction)(e.callback),t.hoverable=!!e.hoverable,t.hover_callback=(0,L.makeFunction)(e.hover_callback),t.unhover_callback=(0,L.makeFunction)(e.unhover_callback),t.contextMenuEnabled=!!e.contextMenuEnabled,t.hidden=e.hidden,t.frame=e.frame}constructor(t){this.color=16777215,this.hidden=!1,this.wireframe=!1,this.opacity=1,this.linewidth=1,this.clickable=!1,this.hoverable=!1,this.contextMenuEnabled=!1,this.side=n.DoubleSide,this.stylespec=t||{},this.boundingSphere=new J.Sphere,this.intersectionShape={sphere:[],cylinder:[],line:[],triangle:[]},GLShape.updateFromStyle(this,this.stylespec),this.components=[],this.shapeObj=null,this.renderedShapeObj=null,this.geo=new n.Geometry(!0),this.linegeo=new n.Geometry(!0)}updateStyle(t){for(var e in t)this.stylespec[e]=t[e];if(GLShape.updateFromStyle(this,this.stylespec),t.voldata&&t.volscheme){(0,L.adjustVolumeStyle)(t);const e=t.volscheme,i=t.voldata,r=s.CC,n=e.range()||[-1,1];this.geo.setColors((function(t,s,a){let o=i.getVal(t,s,a);return r.color(e.valueToHex(o,n))})),delete this.color}}addCustom(t){t.vertexArr=t.vertexArr||[],t.faceArr=t.faceArr||[],t.normalArr=t.normalArr||[],GLShape.drawCustom(this,this.geo,t)}addSphere(t){t.center||(t.center=new $.Vector3(0,0,0)),t.radius=t.radius?(0,$.clamp)(t.radius,0,1/0):1.5,t.color=s.CC.color(t.color),this.intersectionShape.sphere.push(new J.Sphere(t.center,t.radius)),rt.drawSphere(this.geo,t.center,t.radius,t.color,t.quality),this.components.push({centroid:new $.Vector3(t.center.x,t.center.y,t.center.z)});var e=this.geo.updateGeoGroup(0);GLShape.updateBoundingFromPoints(this.boundingSphere,this.components,e.vertexArray,e.vertices)}addBox(t){var e,i,r,s=t.dimensions||{w:1,h:1,d:1};e="number"==typeof s.w?{x:s.w,y:0,z:0}:s.w,i="number"==typeof s.h?{x:0,y:s.h,z:0}:s.h,r="number"==typeof s.d?{x:0,y:0,z:s.d}:s.d;var n=t.corner;null==n&&(n=void 0!==t.center?{x:t.center.x-.5*(e.x+i.x+r.x),y:t.center.y-.5*(e.y+i.y+r.y),z:t.center.z-.5*(e.z+i.z+r.z)}:{x:0,y:0,z:0});var a=[{x:n.x,y:n.y,z:n.z},{x:n.x+e.x,y:n.y+e.y,z:n.z+e.z},{x:n.x+i.x,y:n.y+i.y,z:n.z+i.z},{x:n.x+e.x+i.x,y:n.y+e.y+i.y,z:n.z+e.z+i.z},{x:n.x+r.x,y:n.y+r.y,z:n.z+r.z},{x:n.x+e.x+r.x,y:n.y+e.y+r.y,z:n.z+e.z+r.z},{x:n.x+i.x+r.x,y:n.y+i.y+r.y,z:n.z+i.z+r.z},{x:n.x+e.x+i.x+r.x,y:n.y+e.y+i.y+r.y,z:n.z+e.z+i.z+r.z}],o=[],l=[];o.splice(o.length,0,a[0],a[1],a[2],a[3]),l.splice(l.length,0,0,2,1,1,2,3);var h=4;o.splice(o.length,0,a[2],a[3],a[6],a[7]),l.splice(l.length,0,h+0,h+2,h+1,h+1,h+2,h+3),h+=4,o.splice(o.length,0,a[4],a[5],a[0],a[1]),l.splice(l.length,0,h+0,h+2,h+1,h+1,h+2,h+3),h+=4,o.splice(o.length,0,a[6],a[7],a[4],a[5]),l.splice(l.length,0,h+0,h+2,h+1,h+1,h+2,h+3),h+=4,o.splice(o.length,0,a[3],a[1],a[7],a[5]),l.splice(l.length,0,h+0,h+2,h+1,h+1,h+2,h+3),h+=4,o.splice(o.length,0,a[2],a[6],a[0],a[4]),l.splice(l.length,0,h+0,h+2,h+1,h+1,h+2,h+3),h+=4;var c=(0,L.extend)({},t);c.vertexArr=o,c.faceArr=l,c.normalArr=[],GLShape.drawCustom(this,this.geo,c);var d=new $.Vector3;this.components.push({centroid:d.addVectors(a[0],a[7]).multiplyScalar(.5)});var u=this.geo.updateGeoGroup(0);GLShape.updateBoundingFromPoints(this.boundingSphere,this.components,u.vertexArray,u.vertices)}addCylinder(t){var e,i;e=t.start?new $.Vector3(t.start.x||0,t.start.y||0,t.start.z||0):new $.Vector3(0,0,0),t.end?void 0===(i=new $.Vector3(t.end.x,t.end.y||0,t.end.z||0)).x&&(i.x=3):i=new $.Vector3(0,0,0);var r=t.radius||.1,n=s.CC.color(t.color);this.intersectionShape.cylinder.push(new J.Cylinder(e,i,r)),rt.drawCylinder(this.geo,e,i,r,n,t.fromCap,t.toCap);var a=new $.Vector3;this.components.push({centroid:a.addVectors(e,i).multiplyScalar(.5)});var o=this.geo.updateGeoGroup(0);GLShape.updateBoundingFromPoints(this.boundingSphere,this.components,o.vertexArray,o.vertices)}addDashedCylinder(t){var e,i;t.dashLength=t.dashLength||.25,t.gapLength=t.gapLength||.25,e=t.start?new $.Vector3(t.start.x||0,t.start.y||0,t.start.z||0):new $.Vector3(0,0,0),t.end?void 0===(i=new $.Vector3(t.end.x,t.end.y||0,t.end.z||0)).x&&(i.x=3):i=new $.Vector3(3,0,0);for(var r=t.radius||.1,n=s.CC.color(t.color),a=Math.sqrt(Math.pow(e.x-i.x,2)+Math.pow(e.y-i.y,2)+Math.pow(e.z-i.z,2)),o=a/(t.gapLength+t.dashLength),l=new $.Vector3(t.start.x||0,t.start.y||0,t.start.z||0),h=new $.Vector3(t.end.x,t.end.y||0,t.end.z||0),c=new $.Vector3((i.x-e.x)/(a/t.gapLength),(i.y-e.y)/(a/t.gapLength),(i.z-e.z)/(a/t.gapLength)),d=new $.Vector3((i.x-e.x)/(a/t.dashLength),(i.y-e.y)/(a/t.dashLength),(i.z-e.z)/(a/t.dashLength)),u=0;u<o;u++)h=new $.Vector3(l.x+d.x,l.y+d.y,l.z+d.z),this.intersectionShape.cylinder.push(new J.Cylinder(l,h,r)),rt.drawCylinder(this.geo,l,h,r,n,t.fromCap,t.toCap),l=new $.Vector3(h.x+c.x,h.y+c.y,h.z+c.z);var f=new $.Vector3;this.components.push({centroid:f.addVectors(e,i).multiplyScalar(.5)});var p=this.geo.updateGeoGroup(0);GLShape.updateBoundingFromPoints(this.boundingSphere,this.components,p.vertexArray,p.vertices)}addCurve(t){t.points=t.points||[],t.smooth=t.smooth||10,void 0===t.fromCap&&(t.fromCap=2),void 0===t.toCap&&(t.toCap=2);var e=nt(t.points,t.smooth);if(e.length<3)console.log("Too few points in addCurve");else{var i=t.radius||.1,r=s.CC.color(t.color),n=0,a=e.length-1,o=e[0].distanceTo(e[1]),l=Math.ceil(2*i/o);if(t.toArrow){let t={start:e[a-=l],end:e[e.length-1],radius:i,color:r,mid:1e-4};this.addArrow(t)}if(t.fromArrow){let t={start:e[n+=l],end:e[0],radius:i,color:r,mid:1e-4};this.addArrow(t)}for(var h=Math.ceil(e.length/2),c={radius:i,color:r,fromCap:2,toCap:2},d=n;d<a;d++)c.start=e[d],c.end=e[d+1],c.fromCap=2,c.toCap=2,d<h?(c.fromCap=2,c.toCap=0):d>h?(c.fromCap=0,c.toCap=2):(c.fromCap=2,c.toCap=2),this.addCylinder(c)}}addLine(t){var e,i;e=t.start?new $.Vector3(t.start.x||0,t.start.y||0,t.start.z||0):new $.Vector3(0,0,0),t.end?void 0===(i=new $.Vector3(t.end.x,t.end.y||0,t.end.z||0)).x&&(i.x=3):i=new $.Vector3(3,0,0);var r=this.geo.updateGeoGroup(2),s=r.vertices,n=3*s,a=r.vertexArray;a[n]=e.x,a[n+1]=e.y,a[n+2]=e.z,a[n+3]=i.x,a[n+4]=i.y,a[n+5]=i.z,r.vertices+=2;var o=r.lineArray,l=r.lineidx;o[l]=s,o[l+1]=s+1,r.lineidx+=2;var h=new $.Vector3;this.components.push({centroid:h.addVectors(e,i).multiplyScalar(.5)}),r=this.geo.updateGeoGroup(0),GLShape.updateBoundingFromPoints(this.boundingSphere,this.components,r.vertexArray,r.vertices)}addArrow(t){if(t.start?t.start=new $.Vector3(t.start.x||0,t.start.y||0,t.start.z||0):t.start=new $.Vector3(0,0,0),t.dir instanceof $.Vector3&&"number"==typeof t.length){var e=t.dir.clone().multiplyScalar(t.length).add(t.start);t.end=e}else t.end?(t.end=new $.Vector3(t.end.x,t.end.y||0,t.end.z||0),void 0===t.end.x&&(t.end.x=3)):t.end=new $.Vector3(3,0,0);t.radius=t.radius||.1,t.radiusRatio=t.radiusRatio||1.618034,t.mid=0<t.mid&&t.mid<1?t.mid:.618034,GLShape.drawArrow(this,this.geo,t);var i=new $.Vector3;this.components.push({centroid:i.addVectors(t.start,t.end).multiplyScalar(.5)});var r=this.geo.updateGeoGroup(0);GLShape.updateBoundingFromPoints(this.boundingSphere,this.components,r.vertexArray,r.vertices)}static distance_from(t,e){return Math.sqrt(Math.pow(t.x-e.x,2)+Math.pow(t.y-e.y,2)+Math.pow(t.z-e.z,2))}static inSelectedRegion(t,e,i){for(var r=0;r<e.length;r++)if(GLShape.distance_from(e[r],t)<=i)return!0;return!1}addIsosurface(t,e,i,r){var s,n,a=void 0!==e.isoval&&"number"==typeof e.isoval?e.isoval:0,o=!!e.voxel,l=void 0===e.smoothness?1:e.smoothness,h=t.size.x,c=t.size.y,d=t.size.z,u=new Int16Array(h*c*d),f=t.data;for(s=0,n=u.length;s<n;++s)u[s]=-1;var p=new Uint8Array(h*c*d);for(s=0,n=f.length;s<n;++s){(a>=0?f[s]-a:a-f[s])>0&&(p[s]|=GLShape.ISDONE)}var g=[],m=[];MarchingCube.march(p,g,m,{fulltable:!0,voxel:o,unitCube:t.unit,origin:t.origin,matrix:t.matrix,nX:h,nY:c,nZ:d}),!o&&l>0&&MarchingCube.laplacianSmooth(l,g,m);var v=[],_=[],y=[];if(e.selectedRegion&&void 0===e.coords&&(e.coords=e.selectedRegion),void 0===e.coords&&void 0!==e.selection&&(r?e.coords=r.selectedAtoms(e.selection):console.log("addIsosurface needs viewer is selection provided.")),void 0!==e.coords){var b=e.coords[0].x,x=e.coords[0].y,w=e.coords[0].z,A=e.coords[0].x,C=e.coords[0].y,S=e.coords[0].z;for(let t=0;t<e.coords.length;t++)e.coords[t].x>b?b=e.coords[t].x:e.coords[t].x<A&&(A=e.coords[t].x),e.coords[t].y>x?x=e.coords[t].y:e.coords[t].y<C&&(C=e.coords[t].y),e.coords[t].z>w?w=e.coords[t].z:e.coords[t].z<S&&(S=e.coords[t].z);var M=2;void 0!==e.radius&&(M=e.radius),void 0!==e.selectedOffset&&(M=e.selectedOffset),void 0!==e.seldist&&(M=e.seldist),A-=M,b+=M,C-=M,x+=M,S-=M,w+=M;for(let t=0;t<g.length;t++)g[t].x>A&&g[t].x<b&&g[t].y>C&&g[t].y<x&&g[t].z>S&&g[t].z<w&&GLShape.inSelectedRegion(g[t],e.coords,M)?(v.push(_.length),_.push(g[t])):v.push(-1);for(let t=0;t+2<m.length;t+=3)-1!==v[m[t]]&&-1!==v[m[t+1]]&&-1!==v[m[t+2]]&&(y.push(m[t]-(m[t]-v[m[t]])),y.push(m[t+1]-(m[t+1]-v[m[t+1]])),y.push(m[t+2]-(m[t+2]-v[m[t+2]])));g=_,m=y}GLShape.drawCustom(this,this.geo,{vertexArr:g,faceArr:m,normalArr:[],clickable:e.clickable,hoverable:e.hoverable}),this.updateStyle(e);var z=new $.Vector3(t.origin.x,t.origin.y,t.origin.z),T=new $.Vector3(t.size.x*t.unit.x,t.size.y*t.unit.y,t.size.z*t.unit.z),E=new $.Vector3(0,0,0),L=z.clone(),F=z.clone().add(T);for(let t=0;t<g.length;t++)E.add(g[t]),L.max(g[t]),F.min(g[t]);E.divideScalar(g.length);var I=E.distanceTo(F),O=E.distanceTo(L);this.boundingSphere.center=E,this.boundingSphere.radius=Math.max(I,O),"function"==typeof i&&i()}addVolumetricData(t,e,i){t=new st.VolumeData(t,e),this.addIsosurface(t,i)}finalize(){return GLShape.finalizeGeo(this.geo),this.geo.initTypedArrays(),this.geo}globj(t){if(this.renderedShapeObj&&(t.remove(this.renderedShapeObj),this.renderedShapeObj=null),!this.hidden){GLShape.finalizeGeo(this.geo),this.geo.initTypedArrays(),this.wireframe&&this.geo.setUpWireframe(),void 0!==this.color&&GLShape.updateColor(this.geo,this.color),this.shapeObj=new n.Object3D;var e=null;e=this.side==n.DoubleSide?new n.MeshDoubleLambertMaterial({wireframe:this.wireframe,side:this.side,transparent:this.opacity<1,opacity:this.opacity,wireframeLinewidth:this.linewidth,vertexColors:n.Coloring.VertexColors}):new n.MeshLambertMaterial({wireframe:this.wireframe,side:this.side,transparent:this.opacity<1,opacity:this.opacity,wireframeLinewidth:this.linewidth,vertexColors:n.Coloring.VertexColors});var i=new n.Mesh(this.geo,e);if(this.shapeObj.add(i),this.linegeo&&this.linegeo.vertices>0){var r=new n.LineBasicMaterial({linewidth:this.linewidth,color:this.color}),s=new n.Line(this.linegeo,r,n.LineStyle.LinePieces);this.shapeObj.add(s)}this.renderedShapeObj=this.shapeObj.clone(),t.add(this.renderedShapeObj)}}removegl(t){this.renderedShapeObj&&(void 0!==this.renderedShapeObj.geometry&&this.renderedShapeObj.geometry.dispose(),void 0!==this.renderedShapeObj.material&&this.renderedShapeObj.material.dispose(),t.remove(this.renderedShapeObj),this.renderedShapeObj=null),this.shapeObj=null}get position(){return this.boundingSphere.center}get x(){return this.boundingSphere.center.x}get y(){return this.boundingSphere.center.y}get z(){return this.boundingSphere.center.z}}function wt(t){if(t.vertexArr.length<64e3)return[t];var e=[{vertexArr:[],normalArr:[],faceArr:[]}];t.colorArr&&(e.colorArr=[]);var i=[],r=[],s=0,n=t.faceArr;for(let o=0,l=n.length;o<l;o+=3){let l=e[s];for(let e=0;e<3;e++){var a=n[o+e];i[a]!==s&&(i[a]=s,r[a]=l.vertexArr.length,l.vertexArr.push(t.vertexArr[a]),t.normalArr&&t.normalArr[a]&&l.normalArr.push(t.normalArr[a]),t.colorArr&&t.colorArr[a]&&l.colorArr.push(t.colorArr[a])),l.faceArr.push(r[a])}l.vertexArr.length>=64e3&&(e.push({vertexArr:[],normalArr:[],faceArr:[]}),t.colorArr&&(e.colorArr=[]),s++)}return e}GLShape.ISDONE=2,GLShape.drawCustom=function(t,e,i){var r=i,n=r.vertexArr,a=r.faceArr;0!==n.length&&0!==a.length||console.warn("Error adding custom shape component: No vertices and/or face indices supplied!");var o=i.color;void 0===o&&(o=t.color),o=s.CC.color(o);for(var l=wt(r),h=0,c=l.length;h<c;h++)GLShape.addCustomGeo(t,e,l[h],l[h].colorArr?l[h].colorArr:o,i.clickable)};class GLVolumetricRender{static interpolateArray(t,e){function i(t,e,i){return t+(e-t)*i}var r=[],s=(t.length-1)/(e-1);r[0]=t[0];for(var n=1;n<e-1;n++){var a=n*s,o=Math.floor(a),l=Math.ceil(a),h=a-o;r[n]=i(t[o],t[l],h)}return r[e-1]=t[t.length-1],r}constructor(t,e,i){this.hidden=!1,this.boundingSphere=new J.Sphere,this.renderedShapeObj=null,this.shapeObj=null,this.subsamples=5,this.data=null,this.transferfunctionbuffer=[],this.min=0,this.max=0,e=e||{};var r=Object.assign([],e.transferfn);this.subsamples=e.subsamples||5;var n,a,o,l,h,c,d,u,f,p;r.forEach((function(t){t.value=parseFloat(t.value)})),r.sort((function(t,e){return t.value-e.value})),this.min=r[0].value,0==r.length&&r.push(r[0]),this.max=r[r.length-1].value;for(let t=0;t<r.length-1;t++)if(o=s.CC.color(r[t].color),l=s.CC.color(r[t+1].color),f=r[t].opacity,p=r[t+1].opacity,(n=Math.floor(256*(r[t].value-this.min)/(this.max-this.min)))!=(a=Math.floor(256*(r[t+1].value-this.min)/(this.max-this.min)))){h=GLVolumetricRender.interpolateArray([255*o.r,255*l.r],a-n),c=GLVolumetricRender.interpolateArray([255*o.g,255*l.g],a-n),d=GLVolumetricRender.interpolateArray([255*o.b,255*l.b],a-n),u=GLVolumetricRender.interpolateArray([255*f,255*p],a-n);for(let t=0;t<h.length;t++)this.transferfunctionbuffer.push(h[t]),this.transferfunctionbuffer.push(c[t]),this.transferfunctionbuffer.push(d[t]),this.transferfunctionbuffer.push(u[t])}if(this.transferfunctionbuffer=new Uint8ClampedArray(this.transferfunctionbuffer),t.matrix){let e=new $.Vector3(0,0,0),i=new $.Vector3(t.size.x,t.size.y,t.size.z),r=new $.Vector3(1,1,1);e.applyMatrix4(t.matrix),i.applyMatrix4(t.matrix),r.applyMatrix4(t.matrix).sub(e),this.extent=[[e.x,e.y,e.z],[i.x,i.y,i.z]];for(let e=1;e<7;e++)i.x=1&e?t.size.x:0,i.y=2&e?t.size.y:0,i.z=4&e?t.size.z:0,i.applyMatrix4(t.matrix),this.extent[0][0]=Math.min(this.extent[0][0],i.x),this.extent[0][1]=Math.min(this.extent[0][1],i.y),this.extent[0][2]=Math.min(this.extent[0][2],i.z),this.extent[1][0]=Math.max(this.extent[1][0],i.x),this.extent[1][1]=Math.max(this.extent[1][1],i.y),this.extent[1][2]=Math.max(this.extent[1][2],i.z);let s=i.x-e.x,n=i.y-e.y,a=i.z-e.z;this.maxdepth=Math.sqrt(s*s+n*n+a*a),this.minunit=Math.min(Math.min(r.x,r.y),r.z),this.texmatrix=(new $.Matrix4).identity().scale({x:t.size.x,y:t.size.y,z:t.size.z}),this.texmatrix=this.texmatrix.multiplyMatrices(t.matrix,this.texmatrix),this.texmatrix=this.texmatrix.getInverse(this.texmatrix)}else{this.texmatrix=(new $.Matrix4).identity();let e=t.unit.x*t.size.x,i=t.unit.y*t.size.y,r=t.unit.z*t.size.z;this.texmatrix.makeTranslation(-t.origin.x/e,-t.origin.y/i,-t.origin.z/r),this.texmatrix.scale({x:1/e,y:1/i,z:1/r}),this.minunit=Math.min(Math.min(t.unit.x,t.unit.y),t.unit.z),this.extent=[[t.origin.x,t.origin.y,t.origin.z],[t.origin.x+e,t.origin.y+i,t.origin.z+r]],this.maxdepth=Math.sqrt(e*e+i*i+r*r)}var g=new GLShape({});if(g.addBox({corner:{x:this.extent[0][0],y:this.extent[0][1],z:this.extent[0][2]},dimensions:{w:this.extent[1][0]-this.extent[0][0],h:this.extent[1][1]-this.extent[0][1],d:this.extent[1][2]-this.extent[0][2]}}),this.geo=g.finalize(),this.boundingSphere.center=new $.Vector3((this.extent[0][0]+this.extent[1][0])/2,(this.extent[0][1]+this.extent[1][1])/2,(this.extent[0][2]+this.extent[1][2])/2),this.boundingSphere.radius=this.maxdepth/2,void 0===e.coords&&void 0!==e.selection&&(i?e.coords=i.selectedAtoms(e.selection):console.log("Need to provide viewer to volumetric renderer if selection specified.")),void 0!==e.coords&&void 0!==e.seldist){let i=new Uint8Array(t.data.length),r=e.seldist,s=r*r;for(let n=0,a=e.coords.length;n<a;n++){let a=e.coords[n],o=a.x-r,l=a.y-r,h=a.z-r,c=a.x+r,d=a.y+r,u=a.z+r;if(t.getIndex(o,l,h)>=0||t.getIndex(c,d,u)>=0)for(let e=o;e<c;e+=this.minunit)for(let r=l;r<d;r+=this.minunit)for(let n=h;n<u;n+=this.minunit){let o=t.getIndex(e,r,n);if(o>=0&&!i[o]){(e-a.x)*(e-a.x)+(r-a.y)*(r-a.y)+(n-a.z)*(n-a.z)<s&&(i[o]=1)}}}for(let e=0,r=t.data.length;e<r;e++)0==i[e]&&(t.data[e]=1/0)}this.data=t}globj(t){if(this.renderedShapeObj&&(t.remove(this.renderedShapeObj),this.renderedShapeObj=null),!this.hidden){this.shapeObj=new n.Object3D;var e,i=new n.Texture(this.data,!0),r=new n.Texture(this.transferfunctionbuffer,!1);i.needsUpdate=!0,r.needsUpdate=!0,r.flipY=!1,e=new n.VolumetricMaterial({transferfn:r,transfermin:this.min,transfermax:this.max,map:i,extent:this.extent,maxdepth:this.maxdepth,texmatrix:this.texmatrix,unit:this.minunit,subsamples:this.subsamples});var s=new n.Mesh(this.geo,e);this.shapeObj.add(s),this.renderedShapeObj=this.shapeObj.clone(),t.add(this.renderedShapeObj)}}removegl(t){this.renderedShapeObj&&(void 0!==this.renderedShapeObj.geometry&&this.renderedShapeObj.geometry.dispose(),void 0!==this.renderedShapeObj.material&&this.renderedShapeObj.material.dispose(),t.remove(this.renderedShapeObj),this.renderedShapeObj=null),this.shapeObj=null}get position(){return this.boundingSphere.center}get x(){return this.boundingSphere.center.x}get y(){return this.boundingSphere.center.y}get z(){return this.boundingSphere.center.z}}i(620);function At(t,e="utf8"){return new TextDecoder(e).decode(t)}const Ct=new TextEncoder;const St=(()=>{const t=new Uint8Array(4);return!((new Uint32Array(t.buffer)[0]=1)&t[0])})(),Mt={int8:globalThis.Int8Array,uint8:globalThis.Uint8Array,int16:globalThis.Int16Array,uint16:globalThis.Uint16Array,int32:globalThis.Int32Array,uint32:globalThis.Uint32Array,uint64:globalThis.BigUint64Array,int64:globalThis.BigInt64Array,float32:globalThis.Float32Array,float64:globalThis.Float64Array};class IOBuffer{constructor(t=8192,e={}){let i=!1;"number"==typeof t?t=new ArrayBuffer(t):(i=!0,this.lastWrittenByte=t.byteLength);const r=e.offset?e.offset>>>0:0,s=t.byteLength-r;let n=r;(ArrayBuffer.isView(t)||t instanceof IOBuffer)&&(t.byteLength!==t.buffer.byteLength&&(n=t.byteOffset+r),t=t.buffer),this.lastWrittenByte=i?s:0,this.buffer=t,this.length=s,this.byteLength=s,this.byteOffset=n,this.offset=0,this.littleEndian=!0,this._data=new DataView(this.buffer,n,s),this._mark=0,this._marks=[]}available(t=1){return this.offset+t<=this.length}isLittleEndian(){return this.littleEndian}setLittleEndian(){return this.littleEndian=!0,this}isBigEndian(){return!this.littleEndian}setBigEndian(){return this.littleEndian=!1,this}skip(t=1){return this.offset+=t,this}back(t=1){return this.offset-=t,this}seek(t){return this.offset=t,this}mark(){return this._mark=this.offset,this}reset(){return this.offset=this._mark,this}pushMark(){return this._marks.push(this.offset),this}popMark(){const t=this._marks.pop();if(void 0===t)throw new Error("Mark stack empty");return this.seek(t),this}rewind(){return this.offset=0,this}ensureAvailable(t=1){if(!this.available(t)){const e=2*(this.offset+t),i=new Uint8Array(e);i.set(new Uint8Array(this.buffer)),this.buffer=i.buffer,this.length=this.byteLength=e,this._data=new DataView(this.buffer)}return this}readBoolean(){return 0!==this.readUint8()}readInt8(){return this._data.getInt8(this.offset++)}readUint8(){return this._data.getUint8(this.offset++)}readByte(){return this.readUint8()}readBytes(t=1){return this.readArray(t,"uint8")}readArray(t,e){const i=Mt[e].BYTES_PER_ELEMENT*t,r=this.byteOffset+this.offset,s=this.buffer.slice(r,r+i);if(this.littleEndian===St&&"uint8"!==e&&"int8"!==e){const t=new Uint8Array(this.buffer.slice(r,r+i));t.reverse();const s=new Mt[e](t.buffer);return this.offset+=i,s.reverse(),s}const n=new Mt[e](s);return this.offset+=i,n}readInt16(){const t=this._data.getInt16(this.offset,this.littleEndian);return this.offset+=2,t}readUint16(){const t=this._data.getUint16(this.offset,this.littleEndian);return this.offset+=2,t}readInt32(){const t=this._data.getInt32(this.offset,this.littleEndian);return this.offset+=4,t}readUint32(){const t=this._data.getUint32(this.offset,this.littleEndian);return this.offset+=4,t}readFloat32(){const t=this._data.getFloat32(this.offset,this.littleEndian);return this.offset+=4,t}readFloat64(){const t=this._data.getFloat64(this.offset,this.littleEndian);return this.offset+=8,t}readBigInt64(){const t=this._data.getBigInt64(this.offset,this.littleEndian);return this.offset+=8,t}readBigUint64(){const t=this._data.getBigUint64(this.offset,this.littleEndian);return this.offset+=8,t}readChar(){return String.fromCharCode(this.readInt8())}readChars(t=1){let e="";for(let i=0;i<t;i++)e+=this.readChar();return e}readUtf8(t=1){return At(this.readBytes(t))}decodeText(t=1,e="utf-8"){return At(this.readBytes(t),e)}writeBoolean(t){return this.writeUint8(t?255:0),this}writeInt8(t){return this.ensureAvailable(1),this._data.setInt8(this.offset++,t),this._updateLastWrittenByte(),this}writeUint8(t){return this.ensureAvailable(1),this._data.setUint8(this.offset++,t),this._updateLastWrittenByte(),this}writeByte(t){return this.writeUint8(t)}writeBytes(t){this.ensureAvailable(t.length);for(let e=0;e<t.length;e++)this._data.setUint8(this.offset++,t[e]);return this._updateLastWrittenByte(),this}writeInt16(t){return this.ensureAvailable(2),this._data.setInt16(this.offset,t,this.littleEndian),this.offset+=2,this._updateLastWrittenByte(),this}writeUint16(t){return this.ensureAvailable(2),this._data.setUint16(this.offset,t,this.littleEndian),this.offset+=2,this._updateLastWrittenByte(),this}writeInt32(t){return this.ensureAvailable(4),this._data.setInt32(this.offset,t,this.littleEndian),this.offset+=4,this._updateLastWrittenByte(),this}writeUint32(t){return this.ensureAvailable(4),this._data.setUint32(this.offset,t,this.littleEndian),this.offset+=4,this._updateLastWrittenByte(),this}writeFloat32(t){return this.ensureAvailable(4),this._data.setFloat32(this.offset,t,this.littleEndian),this.offset+=4,this._updateLastWrittenByte(),this}writeFloat64(t){return this.ensureAvailable(8),this._data.setFloat64(this.offset,t,this.littleEndian),this.offset+=8,this._updateLastWrittenByte(),this}writeBigInt64(t){return this.ensureAvailable(8),this._data.setBigInt64(this.offset,t,this.littleEndian),this.offset+=8,this._updateLastWrittenByte(),this}writeBigUint64(t){return this.ensureAvailable(8),this._data.setBigUint64(this.offset,t,this.littleEndian),this.offset+=8,this._updateLastWrittenByte(),this}writeChar(t){return this.writeUint8(t.charCodeAt(0))}writeChars(t){for(let e=0;e<t.length;e++)this.writeUint8(t.charCodeAt(e));return this}writeUtf8(t){return this.writeBytes(function(t){return Ct.encode(t)}(t))}toArray(){return new Uint8Array(this.buffer,this.byteOffset,this.lastWrittenByte)}_updateLastWrittenByte(){this.offset>this.lastWrittenByte&&(this.lastWrittenByte=this.offset)}}const zt=1,Tt=2,Et=3,Lt=4,Ft=5,It=6;function Ot(t){switch(Number(t)){case zt:return"byte";case Tt:return"char";case Et:return"short";case Lt:return"int";case Ft:return"float";case It:return"double";default:return"undefined"}}function Dt(t){switch(Number(t)){case zt:case Tt:return 1;case Et:return 2;case Lt:case Ft:return 4;case It:return 8;default:return-1}}function kt(t){switch(String(t)){case"byte":return zt;case"char":return Tt;case"short":return Et;case"int":return Lt;case"float":return Ft;case"double":return It;default:return-1}}function Rt(t,e){if(1!==t){const i=new Array(t);for(let r=0;r<t;r++)i[r]=e();return i}return e()}function Pt(t,e,i){switch(e){case zt:return Array.from(t.readBytes(i));case Tt:return function(t){if(0===t.charCodeAt(t.length-1))return t.substring(0,t.length-1);return t}(t.readChars(i));case Et:return Rt(i,t.readInt16.bind(t));case Lt:return Rt(i,t.readInt32.bind(t));case Ft:return Rt(i,t.readFloat32.bind(t));case It:return Rt(i,t.readFloat64.bind(t));default:throw new Error(`non valid type ${e}`)}}function Ut(t,e){if(t)throw new TypeError(`Not a valid NetCDF v3.x file: ${e}`)}function Bt(t){t.offset%4!=0&&t.skip(4-t.offset%4)}function Nt(t){const e=t.readUint32(),i=t.readChars(e);return Bt(t),i}function Gt(t,e){const i={version:e},r={length:t.readUint32()},s=function(t){const e={};let i,r;const s=t.readUint32();let n;if(0===s)return Ut(0!==t.readUint32(),"wrong empty tag for list of dimensions"),[];{Ut(10!==s,"wrong tag for list of dimensions");const e=t.readUint32();n=new Array(e);for(let s=0;s<e;s++){const e=Nt(t),a=t.readUint32();0===a&&(i=s,r=e),n[s]={name:e,size:a}}}void 0!==i&&(e.recordId=i);void 0!==r&&(e.recordName=r);return e.dimensions=n,e}(t);Array.isArray(s)||(r.id=s.recordId,r.name=s.recordName,i.dimensions=s.dimensions),i.globalAttributes=Vt(t);const n=function(t,e,i){const r=t.readUint32();let s,n=0;if(0===r)return Ut(0!==t.readUint32(),"wrong empty tag for list of variables"),[];{Ut(11!==r,"wrong tag for list of variables");const a=t.readUint32();s=new Array(a);for(let r=0;r<a;r++){const a=Nt(t),o=t.readUint32(),l=new Array(o);for(let e=0;e<o;e++)l[e]=t.readUint32();const h=Vt(t),c=t.readUint32();Ut(c<1&&c>6,`non valid type ${c}`);const d=t.readUint32();let u=t.readUint32();2===i&&(Ut(u>0,"offsets larger than 4GB not supported"),u=t.readUint32());let f=!1;void 0!==e&&l[0]===e&&(n+=d,f=!0),s[r]={name:a,dimensions:l,attributes:h,type:Ot(c),size:d,offset:u,record:f}}}return{variables:s,recordStep:n}}(t,r?.id,e);return Array.isArray(n)||(i.variables=n.variables,r.recordStep=n.recordStep),i.recordDimension=r,i}function Vt(t){const e=t.readUint32();let i;if(0===e)return Ut(0!==t.readUint32(),"wrong empty tag for list of attributes"),[];{Ut(12!==e,"wrong tag for list of attributes");const r=t.readUint32();i=new Array(r);for(let e=0;e<r;e++){const r=Nt(t),s=t.readUint32();Ut(s<1||s>6,`non valid type ${s}`);const n=t.readUint32(),a=Pt(t,s,n);Bt(t),i[e]={name:r,type:Ot(s),value:a}}}return i}function jt(){const t=[];t.push("DIMENSIONS");for(const e of this.dimensions)t.push(`  ${e.name.padEnd(30)} = size: ${e.size}`);t.push(""),t.push("GLOBAL ATTRIBUTES");for(const e of this.globalAttributes)t.push(`  ${e.name.padEnd(30)} = ${e.value}`);const e=JSON.parse(JSON.stringify(this.variables));t.push(""),t.push("VARIABLES:");for(const i of e){i.value=this.getDataVariable(i);let e=JSON.stringify(i.value);e.length>50&&(e=e.substring(0,50)),isNaN(i.value.length)||(e+=` (length: ${i.value.length})`),t.push(`  ${i.name.padEnd(30)} = ${e}`)}return t.join("\n")}class NetCDFReader{constructor(t){this.toString=jt;const e=new IOBuffer(t);e.setBigEndian(),Ut("CDF"!==e.readChars(3),"should start with CDF");const i=e.readByte();Ut(i>2,"unknown version"),this.header=Gt(e,i),this.buffer=e}get version(){return 1===this.header.version?"classic format":"64-bit offset format"}get recordDimension(){return this.header.recordDimension}get dimensions(){return this.header.dimensions}get globalAttributes(){return this.header.globalAttributes}getAttribute(t){const e=this.globalAttributes.find((e=>e.name===t));return e?e.value:null}getDataVariableAsString(t){const e=this.getDataVariable(t);return e?e.join(""):null}get variables(){return this.header.variables}getDataVariable(t){let e;if(e="string"==typeof t?this.header.variables.find((e=>e.name===t)):t,void 0===e)throw new Error("Not a valid NetCDF v3.x file: variable not found");return this.buffer.seek(e.offset),e.record?function(t,e,i){const r=kt(e.type),s=e.size?e.size/Dt(r):1,n=i.length,a=new Array(n),o=i.recordStep;if(!o)throw new Error("recordDimension.recordStep is undefined");for(let e=0;e<n;e++){const i=t.offset;a[e]=Pt(t,r,s),t.seek(i+o)}return a}(this.buffer,e,this.header.recordDimension):function(t,e){const i=kt(e.type),r=e.size/Dt(i),s=new Array(r);for(let e=0;e<r;e++)s[e]=Pt(t,i,1);return s}(this.buffer,e)}dataVariableExists(t){return void 0!==this.header.variables.find((e=>e.name===t))}attributeExists(t){return void 0!==this.globalAttributes.find((e=>e.name===t))}}class GLModel{static sameObj(t,e){return t&&e?JSON.stringify(t)==JSON.stringify(e):t==e}constructor(t,e){this.atoms=[],this.frames=[],this.box=null,this.atomdfs=null,this.id=0,this.hidden=!1,this.molObj=null,this.renderedMolObj=null,this.lastColors=null,this.modelData={},this.modelDatas=null,this.idMatrix=new $.Matrix4,this.dontDuplicateAtoms=!0,this.defaultColor=s.elementColors.defaultColor,this.defaultStickRadius=.25,this.options=e||{},this.ElementColors=this.options.defaultcolors?this.options.defaultcolors:s.elementColors.defaultColors,this.defaultSphereRadius=this.options.defaultSphereRadius?this.options.defaultSphereRadius:1.5,this.defaultCartoonQuality=this.options.cartoonQuality?this.options.cartoonQuality:10,this.id=t}getRadiusFromStyle(t,e){var i=this.defaultSphereRadius;if(void 0!==e.radius)i=e.radius;else if(GLModel.vdwRadii[t.elem])i=GLModel.vdwRadii[t.elem];else if(t.elem.length>1){let e=t.elem;e=e[0].toUpperCase()+e[1].toLowerCase(),GLModel.vdwRadii[e]&&(i=GLModel.vdwRadii[e])}return void 0!==e.scale&&(i*=e.scale),i}drawAtomCross(t,e){if(t.style.cross){var i=t.style.cross;if(!i.hidden){var r=i.linewidth||GLModel.defaultlineWidth;e[r]||(e[r]=new n.Geometry);var s=e[r].updateGeoGroup(6),a=this.getRadiusFromStyle(t,i),o=[[a,0,0],[-a,0,0],[0,a,0],[0,-a,0],[0,0,a],[0,0,-a]],l=t.clickable||t.hoverable;l&&void 0===t.intersectionShape&&(t.intersectionShape={sphere:[],cylinder:[],line:[]});for(var h=(0,L.getColorFromStyle)(t,i),c=s.vertexArray,d=s.colorArray,u=0;u<6;u++){var f=3*s.vertices;if(s.vertices++,c[f]=t.x+o[u][0],c[f+1]=t.y+o[u][1],c[f+2]=t.z+o[u][2],d[f]=h.r,d[f+1]=h.g,d[f+2]=h.b,l){var p=new $.Vector3(o[u][0],o[u][1],o[u][2]);p.multiplyScalar(.1),p.set(p.x+t.x,p.y+t.y,p.z+t.z),t.intersectionShape.line.push(p)}}}}}getGoodCross(t,e,i,r){for(var s=null,n=-1,a=0,o=t.bonds.length;a<o;a++)if(t.bonds[a]!=e.index){let e=t.bonds[a],o=this.atoms[e],h=new $.Vector3(o.x,o.y,o.z).clone();h.sub(i);let c=h.clone();c.cross(r);var l=c.lengthSq();if(l>n&&(s=c,(n=l)>.1))return s}return s}getSideBondV(t,e,i){var r,s,n,a,o=new $.Vector3(t.x,t.y,t.z),l=new $.Vector3(e.x,e.y,e.z).clone(),h=null;if(l.sub(o),1===t.bonds.length)1===e.bonds.length?(h=l.clone(),Math.abs(h.x)>1e-4?h.y+=1:h.x+=1):(r=(i+1)%e.bonds.length,s=e.bonds[r],(n=this.atoms[s]).index==t.index&&(r=(r+1)%e.bonds.length,s=e.bonds[r],n=this.atoms[s]),(a=new $.Vector3(n.x,n.y,n.z).clone()).sub(o),(h=a.clone()).cross(l));else if((h=this.getGoodCross(t,e,o,l)).lengthSq()<.01){var c=this.getGoodCross(e,t,o,l);null!=c&&(h=c)}return h.lengthSq()<.01&&(h=l.clone(),Math.abs(h.x)>1e-4?h.y+=1:h.x+=1),h.cross(l),h.normalize(),h}addLine(t,e,i,r,s,n){t[i]=r.x,t[i+1]=r.y,t[i+2]=r.z,e[i]=n.r,e[i+1]=n.g,e[i+2]=n.b,t[i+3]=s.x,t[i+4]=s.y,t[i+5]=s.z,e[i+3]=n.r,e[i+4]=n.g,e[i+5]=n.b}drawBondLines(t,e,i){if(t.style.line){var r=t.style.line;if(!r.hidden){var a,o,l,h,c=r.linewidth||GLModel.defaultlineWidth;i[c]||(i[c]=new n.Geometry);for(var d=i[c].updateGeoGroup(6*t.bonds.length),u=d.vertexArray,f=d.colorArray,p=0;p<t.bonds.length;p++){var g=e[t.bonds[p]];if(g.style.line&&!(t.index>=g.index)){var m=new $.Vector3(t.x,t.y,t.z),v=new $.Vector3(g.x,g.y,g.z),_=m.clone().add(v).multiplyScalar(.5),y=!1,b=t.clickable||t.hoverable,x=g.clickable||g.hoverable;(b||x)&&(b&&(void 0===t.intersectionShape&&(t.intersectionShape={sphere:[],cylinder:[],line:[],triangle:[]}),t.intersectionShape.line.push(m),t.intersectionShape.line.push(_)),x&&(void 0===g.intersectionShape&&(g.intersectionShape={sphere:[],cylinder:[],line:[],triangle:[]}),g.intersectionShape.line.push(_),g.intersectionShape.line.push(v)));var w=(0,L.getColorFromStyle)(t,t.style.line),A=(0,L.getColorFromStyle)(g,g.style.line);if(t.bondStyles&&t.bondStyles[p]){var C=t.bondStyles[p];if(!C.iswire)continue;C.singleBond&&(y=!0),void 0!==C.color1&&(w=s.CC.color(C.color1)),void 0!==C.color2&&(A=s.CC.color(C.color2))}var S,M,z=3*d.vertices;if(t.bondOrder[p]>1&&t.bondOrder[p]<4&&!y){var T=this.getSideBondV(t,g,p),E=v.clone();E.sub(m),2==t.bondOrder[p]?(T.multiplyScalar(.1),(a=m.clone()).add(T),(o=m.clone()).sub(T),(l=a.clone()).add(E),(h=o.clone()).add(E),w==A?(d.vertices+=4,this.addLine(u,f,z,a,l,w),this.addLine(u,f,z+6,o,h,w)):(d.vertices+=8,E.multiplyScalar(.5),(S=a.clone()).add(E),(M=o.clone()).add(E),this.addLine(u,f,z,a,S,w),this.addLine(u,f,z+6,S,l,A),this.addLine(u,f,z+12,o,M,w),this.addLine(u,f,z+18,M,h,A))):3==t.bondOrder[p]&&(T.multiplyScalar(.1),(a=m.clone()).add(T),(o=m.clone()).sub(T),(l=a.clone()).add(E),(h=o.clone()).add(E),w==A?(d.vertices+=6,this.addLine(u,f,z,m,v,w),this.addLine(u,f,z+6,a,l,w),this.addLine(u,f,z+12,o,h,w)):(d.vertices+=12,E.multiplyScalar(.5),(S=a.clone()).add(E),(M=o.clone()).add(E),this.addLine(u,f,z,m,_,w),this.addLine(u,f,z+6,_,v,A),this.addLine(u,f,z+12,a,S,w),this.addLine(u,f,z+18,S,l,A),this.addLine(u,f,z+24,o,M,w),this.addLine(u,f,z+30,M,h,A)))}else w==A?(d.vertices+=2,this.addLine(u,f,z,m,v,w)):(d.vertices+=4,this.addLine(u,f,z,m,_,w),this.addLine(u,f,z+6,_,v,A))}}}}}drawAtomSphere(t,e){if(t.style.sphere){var i=t.style.sphere;if(!i.hidden){var r=(0,L.getColorFromStyle)(t,i),s=this.getRadiusFromStyle(t,i);if((!0===t.clickable||t.hoverable)&&void 0!==t.intersectionShape){var n=new $.Vector3(t.x,t.y,t.z);t.intersectionShape.sphere.push(new J.Sphere(n,s))}rt.drawSphere(e,t,s,r)}}}drawAtomClickSphere(t){if(t.style.clicksphere){var e=t.style.clicksphere;if(!e.hidden){var i=this.getRadiusFromStyle(t,e);if((!0===t.clickable||t.hoverable)&&void 0!==t.intersectionShape){var r=new $.Vector3(t.x,t.y,t.z);t.intersectionShape.sphere.push(new J.Sphere(r,i))}}}}drawAtomInstanced(t,e){if(t.style.sphere){var i=t.style.sphere;if(!i.hidden){var r=this.getRadiusFromStyle(t,i),s=(0,L.getColorFromStyle)(t,i),n=e.updateGeoGroup(1),a=n.vertices,o=3*a,l=n.vertexArray,h=n.colorArray,c=n.radiusArray;if(l[o]=t.x,l[o+1]=t.y,l[o+2]=t.z,h[o]=s.r,h[o+1]=s.g,h[o+2]=s.b,c[a]=r,(!0===t.clickable||t.hoverable)&&void 0!==t.intersectionShape){var d=new $.Vector3(t.x,t.y,t.z);t.intersectionShape.sphere.push(new J.Sphere(d,r))}n.vertices+=1}}}drawSphereImposter(t,e,i,r){var s,n=t.updateGeoGroup(4),a=n.vertices,o=3*a,l=n.vertexArray,h=n.colorArray;for(s=0;s<4;s++)l[o+3*s]=e.x,l[o+3*s+1]=e.y,l[o+3*s+2]=e.z;var c=n.normalArray;for(s=0;s<4;s++)h[o+3*s]=r.r,h[o+3*s+1]=r.g,h[o+3*s+2]=r.b;c[o+0]=-i,c[o+1]=i,c[o+2]=0,c[o+3]=-i,c[o+4]=-i,c[o+5]=0,c[o+6]=i,c[o+7]=-i,c[o+8]=0,c[o+9]=i,c[o+10]=i,c[o+11]=0,n.vertices+=4;var d=n.faceArray,u=n.faceidx;d[u+0]=a,d[u+1]=a+1,d[u+2]=a+2,d[u+3]=a+2,d[u+4]=a+3,d[u+5]=a,n.faceidx+=6}drawAtomImposter(t,e){if(t.style.sphere){var i=t.style.sphere;if(!i.hidden){var r=this.getRadiusFromStyle(t,i),s=(0,L.getColorFromStyle)(t,i);if((!0===t.clickable||t.hoverable)&&void 0!==t.intersectionShape){var n=new $.Vector3(t.x,t.y,t.z);t.intersectionShape.sphere.push(new J.Sphere(n,r))}this.drawSphereImposter(e,t,r,s)}}}calculateDashes(t,e,i,r,s){var n=Math.sqrt(Math.pow(t.x-e.x,2)+Math.pow(t.y-e.y,2)+Math.pow(t.z-e.z,2));i=Math.max(i,0),s=Math.max(s,0)+2*i,(r=Math.max(r,.001))+s>n&&(r=n,s=0);var a,o=Math.floor((n-r)/(r+s))+1;s=(n-o*r)/o;for(var l=new $.Vector3(t.x,t.y,t.z),h=new $.Vector3((e.x-t.x)/(n/s),(e.y-t.y)/(n/s),(e.z-t.z)/(n/s)),c=new $.Vector3((e.x-t.x)/(n/r),(e.y-t.y)/(n/r),(e.z-t.z)/(n/r)),d=[],u=0;u<o;u++)a=new $.Vector3(l.x+c.x,l.y+c.y,l.z+c.z),d.push({from:l,to:a}),l=new $.Vector3(a.x+h.x,a.y+h.y,a.z+h.z);return d}static drawStickImposter(t,e,i,r,s,n=0,a=0){for(var o,l=t.updateGeoGroup(4),h=l.vertices,c=3*h,d=l.vertexArray,u=l.colorArray,f=l.radiusArray,p=l.normalArray,g=s.r,m=s.g,v=s.b,_=c,y=0;y<4;y++)d[_]=e.x,p[_]=i.x,u[_]=g,d[++_]=e.y,p[_]=i.y,u[_]=m,d[++_]=e.z,p[_]=i.z,u[_]=y<2?v:(o=void 0,0==(o=-v)&&(o=-1e-4),o),_++;l.vertices+=4,f[h]=-r,f[h+1]=r,f[h+2]=-r,f[h+3]=r;var b=l.faceArray,x=l.faceidx;b[x+0]=h,b[x+1]=h+1,b[x+2]=h+2,b[x+3]=h+2,b[x+4]=h+3,b[x+5]=h,l.faceidx+=6}drawBondSticks(t,e,i){var r,n;if(t.style.stick){var a=t.style.stick;if(!a.hidden){var o,l,h,c,d,u,f,p,g,m,v,_,y,b,x=a.radius||this.defaultStickRadius,w=a.doubleBondScaling||.4,A=a.tripleBondScaling||.25,C=(null===(r=a.dashedBondConfig)||void 0===r?void 0:r.dashLength)||.1,S=(null===(n=a.dashedBondConfig)||void 0===n?void 0:n.gapLength)||.25,M=x,z=a.singleBonds||!1,T=a.dashedBonds||!1,E=0,F=0,I=(0,L.getColorFromStyle)(t,a);!t.capDrawn&&t.bonds.length<4&&(E=2);var O=t=>{var e=i.imposter?GLModel.drawStickImposter:rt.drawCylinder;return!T&&t>=1?e:(t,i,r,s,n,a=0,o=0,l=.1,h=.25)=>{this.calculateDashes(i,r,s,l,h).forEach((i=>{e(t,i.from,i.to,s,n,a,o)}))}};for(h=0;h<t.bonds.length;h++){var D=O(t.bondOrder[h]),k=e[t.bonds[h]];if(_=y=b=null,t.index<k.index){var R=k.style;if(!R.stick||R.stick.hidden)continue;var P=(0,L.getColorFromStyle)(k,R.stick);if(M=x,c=z,t.bondStyles&&t.bondStyles[h]){if((d=t.bondStyles[h]).iswire)continue;d.radius&&(M=d.radius),d.singleBond&&(c=!0),void 0!==d.color1&&(I=s.CC.color(d.color1)),void 0!==d.color2&&(P=s.CC.color(d.color2))}var U=new $.Vector3(t.x,t.y,t.z),B=new $.Vector3(k.x,k.y,k.z);if(t.bondOrder[h]<=1||c||t.bondOrder[h]>3){if(t.bondOrder[h]<1&&(M*=t.bondOrder[h]),!k.capDrawn&&k.bonds.length<4&&(F=2),I!=P?(_=(new $.Vector3).addVectors(U,B).multiplyScalar(.5),D(i,U,_,M,I,E,0,C,S),D(i,_,B,M,P,0,F,C,S)):D(i,U,B,M,I,E,F,C,S),o=t.clickable||t.hoverable,l=k.clickable||k.hoverable,o||l){if(_||(_=(new $.Vector3).addVectors(U,B).multiplyScalar(.5)),o){var N=new J.Cylinder(U,_,M),G=new J.Sphere(U,M);t.intersectionShape.cylinder.push(N),t.intersectionShape.sphere.push(G)}if(l){var V=new J.Cylinder(B,_,M),j=new J.Sphere(B,M);k.intersectionShape.cylinder.push(V),k.intersectionShape.sphere.push(j)}}}else if(t.bondOrder[h]>1){var H=0,W=0;M!=x&&(H=2,W=2);var q,Y,Z,X,K,Q=B.clone(),tt=null;Q.sub(U),tt=this.getSideBondV(t,k,h),2==t.bondOrder[h]?(q=M*w,tt.multiplyScalar(1.5*q),(Y=U.clone()).add(tt),(Z=U.clone()).sub(tt),(X=Y.clone()).add(Q),(K=Z.clone()).add(Q),I!=P?(_=(new $.Vector3).addVectors(Y,X).multiplyScalar(.5),y=(new $.Vector3).addVectors(Z,K).multiplyScalar(.5),D(i,Y,_,q,I,H,0),D(i,_,X,q,P,0,W),D(i,Z,y,q,I,H,0),D(i,y,K,q,P,0,W)):(D(i,Y,X,q,I,H,W),D(i,Z,K,q,I,H,W)),o=t.clickable||t.hoverable,l=k.clickable||k.hoverable,(o||l)&&(_||(_=(new $.Vector3).addVectors(Y,X).multiplyScalar(.5)),y||(y=(new $.Vector3).addVectors(Z,K).multiplyScalar(.5)),o&&(u=new J.Cylinder(Y,_,q),f=new J.Cylinder(Z,y,q),t.intersectionShape.cylinder.push(u),t.intersectionShape.cylinder.push(f)),l&&(g=new J.Cylinder(X,_,q),m=new J.Cylinder(K,y,q),k.intersectionShape.cylinder.push(g),k.intersectionShape.cylinder.push(m)))):3==t.bondOrder[h]&&(q=M*A,tt.cross(Q),tt.normalize(),tt.multiplyScalar(3*q),(Y=U.clone()).add(tt),(Z=U.clone()).sub(tt),(X=Y.clone()).add(Q),(K=Z.clone()).add(Q),I!=P?(_=(new $.Vector3).addVectors(Y,X).multiplyScalar(.5),y=(new $.Vector3).addVectors(Z,K).multiplyScalar(.5),b=(new $.Vector3).addVectors(U,B).multiplyScalar(.5),D(i,Y,_,q,I,H,0),D(i,_,X,q,P,0,W),D(i,U,b,q,I,E,0),D(i,b,B,q,P,0,F),D(i,Z,y,q,I,H,0),D(i,y,K,q,P,0,W)):(D(i,Y,X,q,I,H,W),D(i,U,B,q,I,E,F),D(i,Z,K,q,I,H,W)),o=t.clickable||t.hoverable,l=k.clickable||k.hoverable,(o||l)&&(_||(_=(new $.Vector3).addVectors(Y,X).multiplyScalar(.5)),y||(y=(new $.Vector3).addVectors(Z,K).multiplyScalar(.5)),b||(b=(new $.Vector3).addVectors(U,B).multiplyScalar(.5)),o&&(u=new J.Cylinder(Y.clone(),_.clone(),q),f=new J.Cylinder(Z.clone(),y.clone(),q),p=new J.Cylinder(U.clone(),b.clone(),q),t.intersectionShape.cylinder.push(u),t.intersectionShape.cylinder.push(f),t.intersectionShape.cylinder.push(p)),l&&(g=new J.Cylinder(X.clone(),_.clone(),q),m=new J.Cylinder(K.clone(),y.clone(),q),v=new J.Cylinder(B.clone(),b.clone(),q),k.intersectionShape.cylinder.push(g),k.intersectionShape.cylinder.push(m),k.intersectionShape.cylinder.push(v))))}}}var et=!1,it=0,st=!1;for(h=0;h<t.bonds.length;h++)c=z,t.bondStyles&&t.bondStyles[h]&&((d=t.bondStyles[h]).singleBond&&(c=!0),d.radius&&d.radius!=x&&(st=!0)),(c||1==t.bondOrder[h])&&it++;st?it>0&&(et=!0):0==it&&(t.bonds.length>0||a.showNonBonded)&&(et=!0),et&&(M=x,i.imposter?this.drawSphereImposter(i.sphereGeometry,t,M,I):rt.drawSphere(i,t,M,I))}}}createMolObj(t,e){e=e||{};var i,r,a,o,l=new n.Object3D,h=[],c={},d={},u=this.drawAtomSphere,f=null,p=null;e.supportsImposters?(u=this.drawAtomImposter,(f=new n.Geometry(!0)).imposter=!0,(p=new n.Geometry(!0,!0)).imposter=!0,p.sphereGeometry=new n.Geometry(!0),p.sphereGeometry.imposter=!0,p.drawnCaps={}):e.supportsAIA?(u=this.drawAtomInstanced,(f=new n.Geometry(!1,!0,!0)).instanced=!0,p=new n.Geometry(!0)):(f=new n.Geometry(!0),p=new n.Geometry(!0));var g,m={},v=[Number.POSITIVE_INFINITY,Number.NEGATIVE_INFINITY];for(i=0,a=t.length;i<a;i++){var _=t[i];if(_&&_.style){for(r in(_.clickable||_.hoverable)&&void 0===_.intersectionShape&&(_.intersectionShape={sphere:[],cylinder:[],line:[],triangle:[]}),o={line:void 0,cross:void 0,stick:void 0,sphere:void 0})_.style[r]?_.style[r].opacity?o[r]=parseFloat(_.style[r].opacity):o[r]=1:o[r]=void 0,m[r]?null!=o[r]&&m[r]!=o[r]&&(console.log("Warning: "+r+" opacity is ambiguous"),m[r]=1):m[r]=o[r];u.call(this,_,f),this.drawAtomClickSphere(_),this.drawAtomCross(_,d),this.drawBondLines(_,t,c),this.drawBondSticks(_,t,p),void 0===_.style.cartoon||_.style.cartoon.hidden||("spectrum"!==_.style.cartoon.color||"number"!=typeof _.resi||_.hetflag||(_.resi<v[0]&&(v[0]=_.resi),_.resi>v[1]&&(v[1]=_.resi)),h.push(_))}}if(h.length>0&&xt(l,h,v,this.defaultCartoonQuality),f&&f.vertices>0){f.initTypedArrays();var y=null,b=null;f.imposter?y=new n.SphereImposterMaterial({ambient:0,vertexColors:!0,reflectivity:0}):f.instanced?(b=new n.Geometry(!0),rt.drawSphere(b,{x:0,y:0,z:0},1,new s.Color(.5,.5,.5)),b.initTypedArrays(),y=new n.InstancedMaterial({sphereMaterial:new n.MeshLambertMaterial({ambient:0,vertexColors:!0,reflectivity:0}),sphere:b})):y=new n.MeshLambertMaterial({ambient:0,vertexColors:!0,reflectivity:0}),m.sphere<1&&m.sphere>=0&&(y.transparent=!0,y.opacity=m.sphere),b=new n.Mesh(f,y),l.add(b)}if(p.vertices>0){var x=null,w=null,A=p.sphereGeometry;A&&void 0!==A.vertices&&0!=A.vertices||(A=null),p.initTypedArrays(),A&&A.initTypedArrays();var C={ambient:0,vertexColors:!0,reflectivity:0};p.imposter?(x=new n.StickImposterMaterial(C),w=new n.SphereImposterMaterial(C)):(x=new n.MeshLambertMaterial(C),w=new n.MeshLambertMaterial(C),x.wireframe&&(p.setUpWireframe(),A&&A.setUpWireframe())),m.stick<1&&m.stick>=0&&(x.transparent=!0,x.opacity=m.stick,w.transparent=!0,w.opacity=m.stick);var S=new n.Mesh(p,x);if(l.add(S),A){var M=new n.Mesh(A,w);l.add(M)}}for(i in c)if(c.hasOwnProperty(i)){g=i;var z=new n.LineBasicMaterial({linewidth:g,vertexColors:!0});m.line<1&&m.line>=0&&(z.transparent=!0,z.opacity=m.line),c[i].initTypedArrays();var T=new n.Line(c[i],z,n.LineStyle.LinePieces);l.add(T)}for(i in d)if(d.hasOwnProperty(i)){g=i;var E=new n.LineBasicMaterial({linewidth:g,vertexColors:!0});m.cross<1&&m.cross>=0&&(E.transparent=!0,E.opacity=m.cross),d[i].initTypedArrays();var L=new n.Line(d[i],E,n.LineStyle.LinePieces);l.add(L)}if(this.dontDuplicateAtoms&&this.modelData.symmetries&&this.modelData.symmetries.length>0){var F,I=new n.Object3D;for(F=0;F<this.modelData.symmetries.length;F++){var O=new n.Object3D;(O=l.clone()).matrix.copy(this.modelData.symmetries[F]),O.matrixAutoUpdate=!1,I.add(O)}return I}return l}getInternalState(){return{atoms:this.atoms,frames:this.frames}}setInternalState(t){this.atoms=t.atoms,this.frames=t.frames,this.molObj=null}getCrystData(){if(this.modelData.cryst){if(!this.modelData.cryst.matrix){const t=this.modelData.cryst;this.modelData.cryst.matrix=(0,$.conversionMatrix3)(t.a,t.b,t.c,t.alpha,t.beta,t.gamma)}return this.modelData.cryst}return null}setCrystData(t,e,i,r,s,n){t=t||1,e=e||1,i=i||1,r=r||90,s=s||90,n=n||90;const a=(0,$.conversionMatrix3)(t,e,i,r,s,n);this.modelData.cryst={a:t,b:e,c:i,alpha:r,beta:s,gamma:n,matrix:a}}setCrystMatrix(t){t=t||new $.Matrix3(1,0,0,0,1,0,0,0,1),this.modelData.cryst={matrix:t}}getSymmetries(){return void 0===this.modelData.symmetries&&(this.modelData.symmetries=[this.idMatrix]),this.modelData.symmetries}setSymmetries(t){this.modelData.symmetries=void 0===t?[this.idMatrix]:t}getID(){return this.id}getNumFrames(){return null!=this.frames.numFrames?this.frames.numFrames:this.frames.length}adjustCoord(t,e,i,r){var s=e-t;return s<-i?e+r:s>i?e-r:e}adjustCoordinatesToBox(){if(this.box&&this.atomdfs)for(var t=this.box[0],e=this.box[1],i=this.box[2],r=.9*t,s=.9*e,n=.9*i,a=0;a<this.atomdfs.length;a++)for(var o=this.atomdfs[a],l=1;l<o.length;l++){var h=this.atoms[o[l][0]],c=this.atoms[o[l][1]];h.x=this.adjustCoord(c.x,h.x,r,t),h.y=this.adjustCoord(c.y,h.y,s,e),h.z=this.adjustCoord(c.z,h.z,n,i)}}setFrame(t,e){var i=this.getNumFrames();let r=this;return new Promise((function(s,n){if(0==i&&s(),(t<0||t>=i)&&(t=i-1),null!=r.frames.url){var a=r.frames.url;(0,L.getbin)(a+"/traj/frame/"+t+"/"+r.frames.path,void 0,"POST",void 0).then((function(t){for(var e=new Float32Array(t,44),i=0,n=0;n<r.atoms.length;n++)r.atoms[n].x=e[i++],r.atoms[n].y=e[i++],r.atoms[n].z=e[i++];r.box&&r.atomdfs&&r.adjustCoordinatesToBox(),s()})).catch(n)}else r.atoms=r.frames[t],s();r.molObj=null,r.modelDatas&&t<r.modelDatas.length&&(r.modelData=r.modelDatas[t],r.unitCellObjects&&e&&(e.removeUnitCell(r),e.addUnitCell(r)))}))}addFrame(t){this.frames.push(t)}vibrate(t=10,e=1,i=!1,r,s){var n=0,a=t;i&&(n=-t,a=t),void 0!==this.frames&&void 0!==this.frames.origIndex?this.setFrame(this.frames.origIndex):this.setFrame(0),n<a&&(this.frames=[]),i&&(this.frames.origIndex=t);for(var o=n;o<a;o++){var l=[],h=this.frames.length;if(0!=o||s){for(var c=0;c<this.atoms.length;c++){var d=(0,L.getAtomProperty)(this.atoms[c],"dx"),u=(0,L.getAtomProperty)(this.atoms[c],"dy"),f=(0,L.getAtomProperty)(this.atoms[c],"dz"),p=new $.Vector3(d,u,f),g=new $.Vector3(this.atoms[c].x,this.atoms[c].y,this.atoms[c].z),m=o*e/t;p.multiplyScalar(m),g.add(p);var v={};for(var _ in this.atoms[c])v[_]=this.atoms[c][_];if(v.x=g.x,v.y=g.y,v.z=g.z,l.push(v),r&&s){var y=(0,L.extend)({},s),b=new $.Vector3(d,u,f);if(b.multiplyScalar(e),b.add(g),y.start=g,y.end=b,y.frame=h,!y.color){var x=v.style.sphere;x||(x=v.style.stick),x||(x=v.style.line),y.color=(0,L.getColorFromStyle)(v,x)}r.addArrow(y)}}this.frames.push(l)}else this.frames.push(this.atoms)}}setAtomDefaults(t){for(let e=0;e<t.length;e++){let i=t[e];i&&(i.style=i.style||(0,L.deepCopy)(GLModel.defaultAtomStyle),i.color=i.color||this.ElementColors[i.elem]||this.defaultColor,i.model=this.id,(i.clickable||i.hoverable)&&(i.intersectionShape={sphere:[],cylinder:[],line:[],triangle:[]}))}}addMolData(t,e,i={}){var r=GLModel.parseMolData(t,e,i);this.dontDuplicateAtoms=!i.duplicateAssemblyAtoms;var s=r.modelData;if(s&&(Array.isArray(s)?(this.modelData=s[0],i.frames&&(this.modelDatas=s)):this.modelData=s),r.box?this.box=r.box:this.box=null,0==this.frames.length){for(let t=0;t<r.length;t++)0!=r[t].length&&this.frames.push(r[t]);this.frames[0]&&(this.atoms=this.frames[0])}else if(i.frames)for(let t=0;t<r.length;t++)this.frames.push(r[t]);else for(var n=0;n<r.length;n++)this.addAtoms(r[n]);for(let t=0;t<this.frames.length;t++)this.setAtomDefaults(this.frames[t]);i.vibrate&&i.vibrate.frames&&i.vibrate.amplitude&&this.vibrate(i.vibrate.frames,i.vibrate.amplitude),i.style&&this.setStyle({},i.style)}setDontDuplicateAtoms(t){this.dontDuplicateAtoms=t}setModelData(t){this.modelData=t}propertyMatches(t,e){if(t==e)return!0;if("string"==typeof e&&"number"==typeof t){var i=e.match(/(-?\d+)\s*-\s*(-?\d+)/);if(i){var r=parseInt(i[1]),s=parseInt(i[2]);if(i&&t>=r&&t<=s)return!0}}return!1}static deepCopyAndCache(t,e){if("object"!=typeof t||null==t)return t;if(t.__cache_created)return t;const i={};for(const r in t){const s=t[r];if(Array.isArray(s)){i[r]=[];for(let t=0;t<s.length;t++)i[r].push(GLModel.deepCopyAndCache(s[t],e))}else i[r]="object"==typeof s&&"properties"!=r&&"model"!=r?GLModel.deepCopyAndCache(s,e):s;if("and"==r||"or"==r){const t=[];for(const s of i[r]){const i=new Set;for(const t of e.selectedAtoms(s))i.add(t.index);t.push(i)}if("and"==r){const e=function(t,e){const i=new Set;for(const r of e)t.has(r)&&i.add(r);return i};let s=new Set(t[0]);for(const i of t.splice(1))s=e(s,i);i[r].__cached_results=s}else if("or"==r){const e=new Set;for(const i of t)for(const t of i)e.add(t);i[r].__cached_results=e}}}return i.__cache_created=!0,i}atomIsSelected(t,e){if(void 0===e)return!0;var i=!!e.invert,r=!0;for(var s in e)if("and"==s||"or"==s||"not"==s){if("not"==s){if(this.atomIsSelected(t,e[s])){r=!1;break}}else if(void 0===e[s].__cached_results&&(e=GLModel.deepCopyAndCache(e,this)),!(r=e[s].__cached_results.has(t.index)))break}else if("predicate"===s){if(!e.predicate(t)){r=!1;break}}else if("properties"==s&&t[s]){for(var n in e.properties)if(!n.startsWith("__cache")){if(void 0===t.properties[n]){r=!1;break}if(t.properties[n]!=e.properties[n]){r=!1;break}}}else if(e.hasOwnProperty(s)&&!GLModel.ignoredKeys.has(s)&&!s.startsWith("__cache")){if(void 0===t[s]){r=!1;break}var a=!1;if("bonds"===s){if(e[s]!=t.bonds.length){r=!1;break}}else if(Array.isArray(e[s])){var o=e[s],l=t[s];for(let t=0;t<o.length;t++)if(this.propertyMatches(l,o[t])){a=!0;break}if(!a){r=!1;break}}else{let i=e[s];if(!this.propertyMatches(t[s],i)){r=!1;break}}}return i?!r:r}static squaredDistance(t,e){var i=e.x-t.x,r=e.y-t.y,s=e.z-t.z;return i*i+r*r+s*s}expandAtomList(t,e){if(e<=0)return t;for(var i=(0,L.getExtent)(t,void 0),r=[[],[],[]],s=0;s<3;s++)r[0][s]=i[0][s]-e,r[1][s]=i[1][s]+e,r[2][s]=i[2][s];var n=[];for(let t=0;t<this.atoms.length;t++){var a=this.atoms[t].x,o=this.atoms[t].y,l=this.atoms[t].z;a>=r[0][0]&&a<=r[1][0]&&o>=r[0][1]&&o<=r[1][1]&&l>=r[0][2]&&l<=r[1][2]&&(a>=i[0][0]&&a<=i[1][0]&&o>=i[0][1]&&o<=i[1][1]&&l>=i[0][2]&&l<=i[1][2]||n.push(this.atoms[t]))}return n}static getFloat(t){return"number"==typeof t?t:parseFloat(t)}selectedAtoms(t,e){var i=[];t=GLModel.deepCopyAndCache(t||{},this),e||(e=this.atoms);for(var r=e.length,s=0;s<r;s++){var n=e[s];n&&this.atomIsSelected(n,t)&&i.push(n)}if(t.hasOwnProperty("expand")){const e=GLModel.getFloat(t.expand);let r=this.expandAtomList(i,e),s=i.length;const n=e*e;for(let t=0;t<r.length;t++)for(let e=0;e<s;e++){var a=GLModel.squaredDistance(r[t],i[e]);a<n&&a>0&&i.push(r[t])}}if(t.hasOwnProperty("within")&&t.within.hasOwnProperty("sel")&&t.within.hasOwnProperty("distance")){var o=this.selectedAtoms(t.within.sel,this.atoms),l={};const e=GLModel.getFloat(t.within.distance),r=e*e;for(let t=0;t<o.length;t++)for(let e=0;e<i.length;e++){let s=GLModel.squaredDistance(o[t],i[e]);s<r&&s>0&&(l[e]=1)}var h=[];if(t.within.invert)for(let t=0;t<i.length;t++)l[t]||h.push(i[t]);else for(let t in l)h.push(i[t]);i=h}if(t.hasOwnProperty("byres")){var c={},d=[],u=[];for(let t=0;t<i.length;t++){let e=i[t];var f=e.chain,p=e.resi;if(void 0===c[f]&&(c[f]={}),e.hasOwnProperty("resi")&&void 0===c[f][p])for(c[f][p]=!0,u.push(e);u.length>0;)if(e=u.pop(),f=e.chain,p=e.resi,void 0===d[e.index]){d[e.index]=!0;for(var g=0;g<e.bonds.length;g++){var m=this.atoms[e.bonds[g]];void 0===d[m.index]&&m.hasOwnProperty("resi")&&m.chain==f&&m.resi==p&&(u.push(m),i.push(m))}}}}return i}addAtoms(t){this.molObj=null;var e,i=this.atoms.length,r=[];for(e=0;e<t.length;e++)void 0===t[e].index&&(t[e].index=e),void 0===t[e].serial&&(t[e].serial=e),r[t[e].index]=i+e;for(e=0;e<t.length;e++){var s=t[e],n=r[s.index],a=(0,L.extend)({},s);a.index=n,a.bonds=[],a.bondOrder=[],a.model=this.id,a.style=a.style||(0,L.deepCopy)(GLModel.defaultAtomStyle),void 0===a.color&&(a.color=this.ElementColors[a.elem]||this.defaultColor);for(var o=s.bonds?s.bonds.length:0,l=0;l<o;l++){var h=r[s.bonds[l]];void 0!==h&&(a.bonds.push(h),a.bondOrder.push(s.bondOrder?s.bondOrder[l]:1))}this.atoms.push(a)}}assignBonds(){(0,u.assignBonds)(this.atoms,{assignBonds:!0})}removeAtoms(t){this.molObj=null;var e,i=[];for(e=0;e<t.length;e++)i[t[e].index]=!0;var r=[];for(e=0;e<this.atoms.length;e++){var s=this.atoms[e];i[s.index]||r.push(s)}this.atoms=[],this.addAtoms(r)}setStyle(t,e,i){void 0===e&&void 0===i&&(e=t,t={}),"string"==typeof e&&(e=(0,L.specStringToObject)(e));var r=!1,s=this,n=function(n){var a=s.selectedAtoms(t,n);for(let t=0;t<n.length;t++)n[t]&&(n[t].capDrawn=!1);for(let t=0;t<a.length;t++){r=!0,(a[t].clickable||a[t].hoverable)&&(a[t].intersectionShape={sphere:[],cylinder:[],line:[],triangle:[]}),i||(a[t].style={});for(let i in e)e.hasOwnProperty(i)&&(a[t].style[i]=a[t].style[i]||{},Object.assign(a[t].style[i],e[i]))}};if(void 0!==t.frame&&t.frame<this.frames.length){let e=t.frame;e<0&&(e=this.frames.length+e),n(this.frames[e])}else{n(this.atoms);for(var a=0;a<this.frames.length;a++)this.frames[a]!==this.atoms&&n(this.frames[a])}r&&(this.molObj=null)}setClickable(t,e,i){if(e=!!e,null!==(i=(0,L.makeFunction)(i))){var r=this.selectedAtoms(t,this.atoms),s=r.length;for(let t=0;t<s;t++)r[t].intersectionShape={sphere:[],cylinder:[],line:[],triangle:[]},r[t].clickable=e,i&&(r[t].callback=i);s>0&&(this.molObj=null)}else console.log("Callback is not a function")}setHoverable(t,e,i,r){if(e=!!e,i=(0,L.makeFunction)(i),r=(0,L.makeFunction)(r),null!==i)if(null!==r){var s=this.selectedAtoms(t,this.atoms),n=s.length;for(let t=0;t<n;t++)s[t].intersectionShape={sphere:[],cylinder:[],line:[],triangle:[]},s[t].hoverable=e,i&&(s[t].hover_callback=i),r&&(s[t].unhover_callback=r);n>0&&(this.molObj=null)}else console.log("Unhover_callback is not a function");else console.log("Hover_callback is not a function")}enableContextMenu(t,e){var i;e=!!e;var r=this.selectedAtoms(t,this.atoms),s=r.length;for(i=0;i<s;i++)r[i].intersectionShape={sphere:[],cylinder:[],line:[],triangle:[]},r[i].contextMenuEnabled=e;s>0&&(this.molObj=null)}setColorByElement(t,e){if(null===this.molObj||!GLModel.sameObj(e,this.lastColors)){this.lastColors=e;var i=this.selectedAtoms(t,i);i.length>0&&(this.molObj=null);for(var r=0;r<i.length;r++){var s=i[r];void 0!==e[s.elem]&&(s.color=e[s.elem])}}}setColorByProperty(t,e,i,s){var n,a,o=this.selectedAtoms(t,o);for(this.lastColors=null,o.length>0&&(this.molObj=null),"string"==typeof i&&void 0!==r.Gradient.builtinGradients[i]&&(i=new r.Gradient.builtinGradients[i]),s||(s=i.range()),s||(s=(0,L.getPropertyRange)(o,e)),n=0;n<o.length;n++){a=o[n],null!=(0,L.getAtomProperty)(a,e)&&(a.color=i.valueToHex(parseFloat(a.properties[e]),s))}}setColorByFunction(t,e){var i=this.selectedAtoms(t,i);if("function"==typeof e){this.lastColors=null,i.length>0&&(this.molObj=null);for(let t=0;t<i.length;t++){let r=i[t];r.color=e(r)}}}toCDObject(t=!1){var e={a:[],b:[]};t&&(e.s=[]);for(let r=0;r<this.atoms.length;r++){let s={},n=this.atoms[r];if(s.x=n.x,s.y=n.y,s.z=n.z,"C"!=n.elem&&(s.l=n.elem),t){for(var i=0;i<e.s.length&&JSON.stringify(n.style)!==JSON.stringify(e.s[i]);)i++;i===e.s.length&&e.s.push(n.style),0!==i&&(s.s=i)}e.a.push(s);for(let t=0;t<n.bonds.length;t++){let i=r,s=n.bonds[t];if(i>=s)continue;let a={b:i,e:s},o=n.bondOrder[t];1!=o&&(a.o=o),e.b.push(a)}}return e}globj(t,e){(null===this.molObj||e.regen)&&(this.molObj=this.createMolObj(this.atoms,e),this.renderedMolObj&&(t.remove(this.renderedMolObj),this.renderedMolObj=null),this.renderedMolObj=this.molObj.clone(),this.hidden&&(this.renderedMolObj.setVisible(!1),this.molObj.setVisible(!1)),t.add(this.renderedMolObj))}exportVRML(){return this.createMolObj(this.atoms,{supportsImposters:!1,supportsAIA:!1}).vrml()}removegl(t){this.renderedMolObj&&(void 0!==this.renderedMolObj.geometry&&this.renderedMolObj.geometry.dispose(),void 0!==this.renderedMolObj.material&&this.renderedMolObj.material.dispose(),t.remove(this.renderedMolObj),this.renderedMolObj=null),this.molObj=null}hide(){this.hidden=!0,this.renderedMolObj&&this.renderedMolObj.setVisible(!1),this.molObj&&this.molObj.setVisible(!1)}show(){this.hidden=!1,this.renderedMolObj&&this.renderedMolObj.setVisible(!0),this.molObj&&this.molObj.setVisible(!0)}addPropertyLabels(t,e,i,r){for(var s=this.selectedAtoms(e,s),n=(0,L.deepCopy)(r),a=0;a<s.length;a++){var o=s[a],l=null;void 0!==o[t]?l=String(o[t]):void 0!==o.properties[t]&&(l=String(o.properties[t])),null!=l&&(n.position=o,i.addLabel(l,n))}}addResLabels(t,e,i,r=!1){var s=[],n=function(r,n){for(var a=r.selectedAtoms(t,a),o={},l=0;l<a.length;l++){var h=a[l],c=h.chain,d=h.resn+""+h.resi;o[c]||(o[c]={}),o[c][d]||(o[c][d]=[]),o[c][d].push(h)}var u=(0,L.deepCopy)(i);for(let t in o)if(o.hasOwnProperty(t)){var f=o[t];for(let t in f)if(f.hasOwnProperty(t)){let i=f[t],r=new $.Vector3(0,0,0);for(let t=0;t<i.length;t++){let e=i[t];r.x+=e.x,r.y+=e.y,r.z+=e.z}r.divideScalar(i.length),u.position=r,u.frame=n;let a=e.addLabel(t,u,void 0,!0);s.push(a)}}};if(r){var a=this.getNumFrames();let t=this.atoms;for(let t=0;t<a;t++)this.frames[t]&&(this.atoms=this.frames[t],n(this,t));this.atoms=t}else n(this);return s}setupDFS(){this.atomdfs=[];var t=this,e=new Int8Array(this.atoms.length);e.fill(0);for(var i=function(r,s,n){n.push([r,s]);var a=t.atoms[r];e[r]=1;for(var o=0;o<a.bonds.length;o++){var l=a.bonds[o];t.atoms[l]&&!e[l]&&i(l,r,n)}},r=0;r<this.atoms.length;r++){if(this.atoms[r]&&!e[r]){var s=[];i(r,-1,s),this.atomdfs.push(s)}}}setCoordinatesFromURL(t,e){this.frames=[];var i=this;return this.box&&this.setupDFS(),t.startsWith("http")||(t="http://"+t),(0,L.get)(t+"/traj/numframes/"+e,(function(r){if(!isNaN(parseInt(r)))return i.frames.push(i.atoms),i.frames.numFrames=r,i.frames.url=t,i.frames.path=e,i.setFrame(0)}))}setCoordinates(t,e){if(!t)return[];if(/\.gz$/.test(e=e||"")){e=e.replace(/\.gz$/,"");try{t=(0,L.inflateString)(t)}catch(t){console.log(t)}}if({mdcrd:"",inpcrd:"",pdb:"",netcdf:"",array:""}.hasOwnProperty(e)){this.frames=[];for(var i=this.atoms.length,r=GLModel.parseCrd(t,e),s=0;s<r.length;){for(var n=[],a=0;a<i;a++){var o={};for(var l in this.atoms[a])o[l]=this.atoms[a][l];n[a]=o,n[a].x=r[s++],n[a].y=r[s++],n[a].z=r[s++]}this.frames.push(n)}return this.atoms=this.frames[0],this.frames}return[]}addAtomSpecs(t){}static parseCrd(t,e){var i=[],r=0;if("pdb"==e)for(var s=t.indexOf("\nATOM");-1!=s;){for(;"\nATOM"==t.slice(s,s+5)||"\nHETATM"==t.slice(s,s+7);)i[r++]=parseFloat(t.slice(s+31,s+39)),i[r++]=parseFloat(t.slice(s+39,s+47)),i[r++]=parseFloat(t.slice(s+47,s+55)),s=t.indexOf("\n",s+54),"\nTER"==t.slice(s,s+4)&&(s=t.indexOf("\n",s+5));s=t.indexOf("\nATOM",s)}else if("netcdf"==e){var n=new NetCDFReader(t);i=[].concat.apply([],n.getDataVariable("coordinates"))}else{if("array"==e||Array.isArray(t))return t.flat(2);{let r=t.indexOf("\n");"inpcrd"==e&&(r=t.indexOf("\n",r+1)),i=(t=t.slice(r+1)).match(/\S+/g).map(parseFloat)}}return i}static parseMolData(t,e="",i){if(!t)return[];if(/\.gz$/.test(e)){e=e.replace(/\.gz$/,"");try{t=e.match(/bcif/i)?(0,L.inflateString)(t,!1):(0,L.inflateString)(t)}catch(t){console.log(t)}}return void 0===K[e]&&(e=e.split(".").pop(),void 0===K[e]&&(console.log("Unknown format: "+e),e=t instanceof Uint8Array?"bcif":t.match(/^@<TRIPOS>MOLECULE/gm)?"mol2":t.match(/^data_/gm)&&t.match(/^loop_/gm)?"cif":t.match(/^HETATM/gm)||t.match(/^ATOM/gm)?"pdb":t.match(/ITEM: TIMESTEP/gm)?"lammpstrj":t.match(/^.*\n.*\n.\s*(\d+)\s+(\d+)/gm)?"sdf":t.match(/^%VERSION\s+VERSION_STAMP/gm)?"prmtop":"xyz",console.log("Best guess: "+e))),(0,K[e])(t,i)}}GLModel.defaultAtomStyle={line:{}},GLModel.defaultlineWidth=1,GLModel.vdwRadii={H:1.2,He:1.4,Li:1.82,Be:1.53,B:1.92,C:1.7,N:1.55,O:1.52,F:1.47,Ne:1.54,Na:2.27,Mg:1.73,Al:1.84,Si:2.1,P:1.8,S:1.8,Cl:1.75,Ar:1.88,K:2.75,Ca:2.31,Ni:1.63,Cu:1.4,Zn:1.39,Ga:1.87,Ge:2.11,As:1.85,Se:1.9,Br:1.85,Kr:2.02,Rb:3.03,Sr:2.49,Pd:1.63,Ag:1.72,Cd:1.58,In:1.93,Sn:2.17,Sb:2.06,Te:2.06,I:1.98,Xe:2.16,Cs:3.43,Ba:2.68,Pt:1.75,Au:1.66,Hg:1.55,Tl:1.96,Pb:2.02,Bi:2.07,Po:1.97,At:2.02,Rn:2.2,Fr:3.48,Ra:2.83,U:1.86},GLModel.ignoredKeys=new Set(["props","invert","model","frame","byres","expand","within","and","or","not"]);var Ht=i(111);const Wt=16;class GLViewer{getWidth(){let t=this.container,e=t.offsetWidth;if(0==e&&"none"===t.style.display){let i=t.style.position,r=t.style.visibility;t.style.display="block",t.style.visibility="hidden",t.style.position="absolute",e=t.offsetWidth,t.style.display="none",t.style.visibility=r,t.style.position=i}return e}getHeight(){let t=this.container,e=t.offsetHeight;if(0==e&&"none"===t.style.display){let i=t.style.position,r=t.style.visibility;t.style.display="block",t.style.visibility="hidden",t.style.position="absolute",e=t.offsetHeight,t.style.display="none",t.style.visibility=r,t.style.position=i}return e}setupRenderer(){this.renderer=new n.Renderer({antialias:this.config.antialias,preserveDrawingBuffer:!0,premultipliedAlpha:!1,id:this.config.id,row:this.config.row,col:this.config.col,rows:this.config.rows,cols:this.config.cols,canvas:this.config.canvas,containerWidth:this.WIDTH,containerHeight:this.HEIGHT,ambientOcclusion:this.config.ambientOcclusion,outline:this.config.outline}),this.renderer.domElement.style.width="100%",this.renderer.domElement.style.height="100%",this.renderer.domElement.style.padding="0",this.renderer.domElement.style.position="absolute",this.renderer.domElement.style.top="0px",this.renderer.domElement.style.left="0px",this.renderer.domElement.style.zIndex="0"}initializeScene(){this.scene=new n.Scene,this.scene.fog=new n.Fog(this.bgColor,100,200),this.modelGroup=new n.Object3D,this.rotationGroup=new n.Object3D,this.rotationGroup.useQuaternion=!0,this.rotationGroup.quaternion=new $.Quaternion(0,0,0,1),this.rotationGroup.add(this.modelGroup),this.scene.add(this.rotationGroup);var t=new n.Light(16777215);t.position=new $.Vector3(.2,.2,1).normalize(),t.intensity=1,this.scene.add(t)}_handleLostContext(t){const e=function(t){const e=t.getBoundingClientRect();return!(e.right<0||e.bottom<0||e.top>(window.innerHeight||document.documentElement.clientHeight)||e.left>(window.innerWidth||document.documentElement.clientWidth))};if(e(this.container)){let t=0;for(let i of document.getElementsByTagName("canvas"))if(e(i)&&null!=i._3dmol_viewer&&(i._3dmol_viewer.resize(),t+=1,t>=Wt))break}}initContainer(t){this.container=t,this.WIDTH=this.getWidth(),this.HEIGHT=this.getHeight(),this.ASPECT=this.renderer.getAspect(this.WIDTH,this.HEIGHT),this.renderer.setSize(this.WIDTH,this.HEIGHT),this.container.append(this.renderer.domElement),this.glDOM=this.renderer.domElement,this.glDOM._3dmol_viewer=this,this.glDOM.addEventListener("webglcontextlost",this._handleLostContext.bind(this)),this.nomouse||(this.glDOM.addEventListener("mousedown",this._handleMouseDown.bind(this),{passive:!1}),this.glDOM.addEventListener("touchstart",this._handleMouseDown.bind(this),{passive:!1}),this.glDOM.addEventListener("wheel",this._handleMouseScroll.bind(this),{passive:!1}),this.glDOM.addEventListener("mousemove",this._handleMouseMove.bind(this),{passive:!1}),this.glDOM.addEventListener("touchmove",this._handleMouseMove.bind(this),{passive:!1}),this.glDOM.addEventListener("contextmenu",this._handleContextMenu.bind(this),{passive:!1}))}decAnim(){this.animated--,this.animated<0&&(this.animated=0)}incAnim(){this.animated++}nextSurfID(){var t=0;for(let i in this.surfaces)if(this.surfaces.hasOwnProperty(i)){var e=parseInt(i);isNaN(e)||e>t&&(t=e)}return t+1}setSlabAndFog(){let t=this.camera.position.z-this.rotationGroup.position.z;t<1&&(t=1),this.camera.near=t+this.slabNear,!this.camera.ortho&&this.camera.near<1&&(this.camera.near=1),this.camera.far=t+this.slabFar,this.camera.near+1>this.camera.far&&(this.camera.far=this.camera.near+1),this.camera.fov=this.fov,this.camera.right=t*Math.tan(Math.PI/180*this.fov),this.camera.left=-this.camera.right,this.camera.top=this.camera.right/this.ASPECT,this.camera.bottom=-this.camera.top,this.camera.updateProjectionMatrix(),this.scene.fog.near=this.camera.near+this.fogStart*(this.camera.far-this.camera.near),this.scene.fog.far=this.camera.far,this.config.disableFog&&(this.scene.fog.near=this.scene.fog.far)}show(t){if(this.renderer.setViewport(),this.scene&&(this.setSlabAndFog(),this.renderer.render(this.scene,this.camera),this.viewChangeCallback&&this.viewChangeCallback(this._viewer.getView()),!t&&this.linkedViewers.length>0))for(var e=this._viewer.getView(),i=0;i<this.linkedViewers.length;i++){this.linkedViewers[i].setView(e,!0)}}updateClickables(){this.clickables.splice(0,this.clickables.length),this.hoverables.splice(0,this.hoverables.length),this.contextMenuEnabledObjects.splice(0,this.contextMenuEnabledObjects.length);for(let t=0,e=this.models.length;t<e;t++){let e=this.models[t];if(e){let t=e.selectedAtoms({clickable:!0}),i=e.selectedAtoms({hoverable:!0}),r=e.selectedAtoms({contextMenuEnabled:!0});for(let t=0;t<i.length;t++)this.hoverables.push(i[t]);for(let e=0;e<t.length;e++)this.clickables.push(t[e]);for(let t=0;t<r.length;t++)this.contextMenuEnabledObjects.push(r[t])}}for(let t=0,e=this.shapes.length;t<e;t++){let e=this.shapes[t];e&&e.clickable&&this.clickables.push(e),e&&e.hoverable&&this.hoverables.push(e),e&&e.contextMenuEnabled&&this.contextMenuEnabledObjects.push(e)}}handleClickSelection(t,e,i){let r=this.targetedObjects(t,e,this.clickables);if(r.length){var s=r[0].clickable;if(void 0!==s.callback&&("function"!=typeof s.callback&&(s.callback=(0,L.makeFunction)(s.callback)),"function"==typeof s.callback)){3===this.mouseButton&&this.contextMenuEnabledObjects.includes(s)&&this.userContextMenuHandler||s.callback(s,this._viewer,i,this.container,r)}}}canvasOffset(){let t=this.glDOM,e=t.getBoundingClientRect(),i=t.ownerDocument,r=i.documentElement,s=i.defaultView;return{top:e.top+s.pageYOffset-r.clientTop,left:e.left+s.pageXOffset-r.clientLeft}}setHover(t,e,i){this.current_hover!=t&&(this.current_hover&&("function"!=typeof this.current_hover.unhover_callback&&(this.current_hover.unhover_callback=(0,L.makeFunction)(this.current_hover.unhover_callback)),this.current_hover.unhover_callback(this.current_hover,this._viewer,e,this.container,i)),this.current_hover=t,t&&void 0!==t.hover_callback&&("function"!=typeof t.hover_callback&&(t.hover_callback=(0,L.makeFunction)(t.hover_callback)),"function"==typeof t.hover_callback&&t.hover_callback(t,this._viewer,e,this.container,i)))}handleHoverSelection(t,e,i){if(0==this.hoverables.length)return;let r=this.targetedObjects(t,e,this.hoverables);if(r.length){var s=r[0].clickable;this.setHover(s,i,r),this.current_hover=s}else this.setHover(null)}handleHoverContinue(t,e){let i=this.targetedObjects(t,e,this.hoverables);0!=i.length&&void 0!==i[0]||this.setHover(null),void 0!==i[0]&&i[0].clickable!==this.current_hover&&this.setHover(null)}closeEnoughForClick(t,{allowTolerance:e=t.targetTouches,tolerance:i=5}={}){const r=this.getX(t),s=this.getY(t);if(e){const t=Math.abs(r-this.mouseStartX),e=Math.abs(s-this.mouseStartY);return t<=i&&e<=i}return r===this.mouseStartX&&s===this.mouseStartY}calcTouchDistance(t){var e=t.targetTouches[0].pageX-t.targetTouches[1].pageX,i=t.targetTouches[0].pageY-t.targetTouches[1].pageY;return Math.hypot(e,i)}getX(t){var e=t.pageX;return null==e&&(e=t.pageX),t.targetTouches&&t.targetTouches[0]?e=t.targetTouches[0].pageX:t.changedTouches&&t.changedTouches[0]&&(e=t.changedTouches[0].pageX),e}getY(t){var e=t.pageY;return null==e&&(e=t.pageY),t.targetTouches&&t.targetTouches[0]?e=t.targetTouches[0].pageY:t.changedTouches&&t.changedTouches[0]&&(e=t.changedTouches[0].pageY),e}isInViewer(t,e){if(null!=this.viewers){var i=this.WIDTH/this.cols,r=this.HEIGHT/this.rows,s=this.canvasOffset(),n=t-s.left,a=e-s.top,o=this.rows-Math.floor(a/r)-1,l=Math.floor(n/i);if(o!=this.row||l!=this.col)return!1}return!0}adjustZoomToLimits(t){if(this.config.lowerZoomLimit&&this.config.lowerZoomLimit>0){let e=this.CAMERA_Z-this.config.lowerZoomLimit;t>e&&(t=e)}if(this.config.upperZoomLimit&&this.config.upperZoomLimit>0){let e=this.CAMERA_Z-this.config.upperZoomLimit;t<e&&(t=e)}return t>this.CAMERA_Z-1&&(t=this.CAMERA_Z-1),t}static slerp(t,e,i){if(1==i)return e.clone();if(0==i)return t.clone();let r=t.x*e.x+t.y*e.y+t.z*e.z+t.w*e.w;if(r>.9995){let r=new $.Quaternion(t.x+i*(e.x-t.x),t.y+i*(e.y-t.y),t.z+i*(e.z-t.z),t.w+i*(e.w-t.w));return r.normalize(),r}r<0&&(e=e.clone().multiplyScalar(-1),r=-r),r>1?r=1:r<-1&&(r=-1);var s=Math.acos(r)*i,n=e.clone();n.sub(t.clone().multiplyScalar(r)),n.normalize();var a=Math.cos(s),o=Math.sin(s),l=new $.Quaternion(t.x*a+n.x*o,t.y*a+n.y*o,t.z*a+n.z*o,t.w*a+n.w*o);return l.normalize(),l}constructor(t,e={}){if(this.nomouse=!1,this.glDOM=null,this.models=[],this.surfaces={},this.shapes=[],this.labels=[],this.clickables=[],this.hoverables=[],this.contextMenuEnabledObjects=[],this.current_hover=null,this.hoverDuration=500,this.longTouchDuration=1e3,this.viewer_frame=0,this.viewChangeCallback=null,this.stateChangeCallback=null,this.NEAR=1,this.FAR=800,this.CAMERA_Z=150,this.fov=20,this.linkedViewers=[],this.renderer=null,this.control_all=!1,this.scene=null,this.rotationGroup=null,this.modelGroup=null,this.fogStart=.4,this.slabNear=-50,this.slabFar=50,this.cq=new $.Quaternion(0,0,0,1),this.dq=new $.Quaternion(0,0,0,1),this.animated=0,this.animationTimers=new Set,this.isDragging=!1,this.mouseStartX=0,this.mouseStartY=0,this.touchDistanceStart=0,this.touchHold=!1,this.currentModelPos=0,this.cz=0,this.cslabNear=0,this.cslabFar=0,this.userContextMenuHandler=null,this.config=e,this.callback=this.config.callback,this.defaultcolors=this.config.defaultcolors,this.defaultcolors||(this.defaultcolors=s.elementColors.defaultColors),this.nomouse=Boolean(this.config.nomouse),this.bgColor=0,this.config.backgroundColor=this.config.backgroundColor||"#ffffff",void 0!==this.config.backgroundColor&&(this.bgColor=s.CC.color(this.config.backgroundColor).getHex()),this.config.backgroundAlpha=null==this.config.backgroundAlpha?1:this.config.backgroundAlpha,this.camerax=0,void 0!==this.config.camerax&&(this.camerax="string"==typeof this.config.camerax?parseFloat(this.config.camerax):this.config.camerax),this._viewer=this,this.container=t,null!=this.config.hoverDuration&&(this.hoverDuration=this.config.hoverDuration),void 0===this.config.antialias&&(this.config.antialias=!0),void 0===this.config.cartoonQuality&&(this.config.cartoonQuality=10),this.WIDTH=this.getWidth(),this.HEIGHT=this.getHeight(),this.setupRenderer(),this.row=null==this.config.row?0:this.config.row,this.col=null==this.config.col?0:this.config.col,this.cols=this.config.cols,this.rows=this.config.rows,this.viewers=this.config.viewers,this.control_all=this.config.control_all,this.ASPECT=this.renderer.getAspect(this.WIDTH,this.HEIGHT),this.camera=new n.Camera(this.fov,this.ASPECT,this.NEAR,this.FAR,this.config.orthographic),this.camera.position=new $.Vector3(this.camerax,0,this.CAMERA_Z),this.lookingAt=new $.Vector3,this.camera.lookAt(this.lookingAt),this.raycaster=new n.Raycaster(new $.Vector3(0,0,0),new $.Vector3(0,0,0)),this.projector=new n.Projector,this.initializeScene(),this.renderer.setClearColorHex(this.bgColor,this.config.backgroundAlpha),this.scene.fog.color=s.CC.color(this.bgColor),document.body.addEventListener("mouseup",this._handleMouseUp.bind(this)),document.body.addEventListener("touchend",this._handleMouseUp.bind(this)),this.initContainer(this.container),this.config.style&&this.setViewStyle(this.config),window.addEventListener("resize",this.resize.bind(this)),void 0!==window.ResizeObserver&&(this.divwatcher=new window.ResizeObserver(this.resize.bind(this)),this.divwatcher.observe(this.container)),void 0!==window.IntersectionObserver){let t=(t,e)=>{t.forEach((t=>{t.isIntersecting&&this.resize()}))};this.intwatcher=new window.IntersectionObserver(t),this.intwatcher.observe(this.container)}try{"function"==typeof this.callback&&this.callback(this)}catch(t){console.log("error with glviewer callback: "+t)}}targetedObjects(t,e,i){var r={x:t,y:e,z:-1};return Array.isArray(i)||(i=this.selectedAtoms(i)),0==i.length?[]:(this.raycaster.setFromCamera(r,this.camera),this.raycaster.intersectObjects(this.modelGroup,i))}modelToScreen(t){let e=!1;Array.isArray(t)||(t=[t],e=!0);let i=this.renderer.getXRatio(),r=this.renderer.getYRatio(),s=this.col,n=this.row,a=s*(this.WIDTH/i),o=(r-n-1)*(this.HEIGHT/r),l=[],h=this.canvasOffset();return t.forEach((t=>{let e=new $.Vector3(t.x,t.y,t.z);e.applyMatrix4(this.modelGroup.matrixWorld),this.projector.projectVector(e,this.camera);let s=this.WIDTH/i*(e.x+1)/2+h.left+a,n=-this.HEIGHT/r*(e.y-1)/2+h.top+o;l.push({x:s,y:n})})),e&&(l=l[0]),l}screenOffsetToModel(t,e,i){var r=t/this.WIDTH,s=e/this.HEIGHT,n=void 0===i?this.rotationGroup.position.z:i,a=this.rotationGroup.quaternion,o=new $.Vector3(0,0,n);return this.projector.projectVector(o,this.camera),o.x+=2*r,o.y-=2*s,this.projector.unprojectVector(o,this.camera),o.z=0,o.applyQuaternion(a),o}screenToModelDistance(t,e){let i=this.canvasOffset(),r=new $.Vector3(e.x,e.y,e.z);r.applyMatrix4(this.modelGroup.matrixWorld);let s=r.clone();this.projector.projectVector(r,this.camera);let n=new $.Vector3(2*(t.x-i.left)/this.WIDTH-1,2*(t.y-i.top)/-this.HEIGHT+1,r.z);return this.projector.unprojectVector(n,this.camera),n.distanceTo(s)}setViewChangeCallback(t){"function"!=typeof t&&null!=t||(this.viewChangeCallback=t)}setStateChangeCallback(t){"function"!=typeof t&&null!=t||(this.stateChangeCallback=t)}getConfig(){return this.config}setConfig(t){this.config=t,t.ambientOcclusion&&this.renderer.enableAmbientOcclusion(t.ambientOcclusion)}getInternalState(){var t={models:[],surfaces:[],shapes:[],labels:[]};for(let e=0;e<this.models.length;e++)this.models[e]&&(t.models[e]=this.models[e].getInternalState());return t}setInternalState(t){this.clear();var e=t.models;for(let t=0;t<e.length;t++)e[t]&&(this.models[t]=new GLModel(t),this.models[t].setInternalState(e[t]));this.render()}setZoomLimits(t,e){void 0!==t&&(this.config.lowerZoomLimit=t),e&&(this.config.upperZoomLimit=e),this.rotationGroup.position.z=this.adjustZoomToLimits(this.rotationGroup.position.z),this.show()}setCameraParameters(t){void 0!==t.fov&&(this.fov=t.fov,this.camera.fov=this.fov),void 0!==t.z&&(this.CAMERA_Z=t.z,this.camera.z=this.CAMERA_Z),void 0!==t.orthographic&&(this.camera.ortho=t.orthographic),this.setSlabAndFog()}_handleMouseDown(t){if(t.preventDefault(),!this.scene)return;var e=this.getX(t),i=this.getY(t);if(void 0===e)return;this.isDragging=!0,this.mouseButton=t.which,this.mouseStartX=e,this.mouseStartY=i,this.touchHold=!0,this.touchDistanceStart=0,t.targetTouches&&2==t.targetTouches.length&&(this.touchDistanceStart=this.calcTouchDistance(t)),this.cq=this.rotationGroup.quaternion.clone(),this.cz=this.rotationGroup.position.z,this.currentModelPos=this.modelGroup.position.clone(),this.cslabNear=this.slabNear,this.cslabFar=this.slabFar;let r=this;t.targetTouches&&1===t.targetTouches.length&&(this.longTouchTimeout=setTimeout((function(){if(1==r.touchHold){r.glDOM=r.renderer.domElement;const e=t.targetTouches[0],i=new PointerEvent("contextmenu",Object.assign(Object.assign({},t),{pageX:e.pageX,pageY:e.pageY,screenX:e.screenX,screenY:e.screenY,clientX:e.clientX,clientY:e.clientY}));r.glDOM.dispatchEvent(i)}}),this.longTouchDuration))}_handleMouseUp(t){if(this.touchHold=!1,this.isDragging&&this.scene){var e=this.getX(t),i=this.getY(t);if(this.closeEnoughForClick(t)&&this.isInViewer(e,i)){let r=this.mouseXY(e,i);this.handleClickSelection(r.x,r.y,t)}}this.isDragging=!1}_handleMouseScroll(t){if(t.preventDefault(),this.scene){var e=this.getX(t),i=this.getY(t);if(void 0!==e&&(this.control_all||this.isInViewer(e,i))){var r=.85*(this.CAMERA_Z-this.rotationGroup.position.z),s=1;if(t.ctrlKey&&(s=-1),t.detail)this.rotationGroup.position.z+=s*r*t.detail/10;else if(t.wheelDelta){let e=600*t.wheelDelta/(t.wheelDelta+600);this.rotationGroup.position.z-=s*r*e/400}this.rotationGroup.position.z=this.adjustZoomToLimits(this.rotationGroup.position.z),this.show()}}}pngURI(){return this.getCanvas().toDataURL("image/png")}apngURI(t){let e=this;return t=t||1,new Promise((function(i){let r=0,s=e.viewChangeCallback,n=[],a=[],o=Date.now();e.viewChangeCallback=function(){a.push(Date.now()-o),o=Date.now(),n.push(new Promise((t=>{e.getCanvas().toBlob((function(e){e.arrayBuffer().then(t)}),"image/png")}))),r+=1,r==t&&(e.viewChangeCallback=s,Promise.all(n).then((t=>{let r=[];for(let e=0;e<t.length;e++){let i=(0,Ht.decode)(t[e]);r.push((0,Ht.toRGBA8)(i)[0])}let s=e.getCanvas().width,n=e.getCanvas().height,o=(0,Ht.encode)(r,s,n,0,a),l=new Blob([o],{type:"image/png"}),h=new FileReader;h.onload=function(t){i(t.target.result)},h.readAsDataURL(l)})))}}))}getCanvas(){return this.glDOM}getRenderer(){return this.renderer}setHoverDuration(t){this.hoverDuration=t}mouseXY(t,e){let i=this.canvasOffset(),r=this.renderer.getXRatio(),s=this.renderer.getYRatio(),n=this.col,a=this.row,o=n*(this.WIDTH/r),l=(s-a-1)*(this.HEIGHT/s);return{x:(t-i.left-o)/(this.WIDTH/r)*2-1,y:-(e-i.top-l)/(this.HEIGHT/s)*2+1}}_handleMouseMove(t){clearTimeout(this.hoverTimeout),t.preventDefault();let e=this.getX(t),i=this.getY(t);if(void 0===e)return;let r=this.renderer.getXRatio(),s=this.renderer.getYRatio(),n=this.mouseXY(e,i),a=this;null!==this.current_hover&&this.handleHoverContinue(n.x,n.y);var o=0;if((this.control_all||this.isInViewer(e,i))&&this.scene&&(this.hoverables.length>0&&(this.hoverTimeout=setTimeout((function(){a.handleHoverSelection(n.x,n.y,t)}),this.hoverDuration)),this.isDragging)){t.targetTouches&&(t.targetTouches.length>1||1===t.targetTouches.length&&!this.closeEnoughForClick(t))&&clearTimeout(this.longTouchTimeout);var l=(e-this.mouseStartX)/this.WIDTH,h=(i-this.mouseStartY)/this.HEIGHT;if(0!=this.touchDistanceStart&&t.targetTouches&&2==t.targetTouches.length)o=2,h=2*(this.calcTouchDistance(t)-this.touchDistanceStart)/(this.WIDTH+this.HEIGHT);else t.targetTouches&&3==t.targetTouches.length&&(o=1);l*=r,h*=s;var c,d=Math.hypot(l,h);if(3==o||3==this.mouseButton&&t.ctrlKey)this.slabNear=this.cslabNear+100*l,this.slabFar=this.cslabFar-100*h;else if(2==o||3==this.mouseButton||t.shiftKey)(c=.85*(this.CAMERA_Z-this.rotationGroup.position.z))<80&&(c=80),this.rotationGroup.position.z=this.cz+h*c,this.rotationGroup.position.z=this.adjustZoomToLimits(this.rotationGroup.position.z);else if(1==o||2==this.mouseButton||t.ctrlKey){var u=this.screenOffsetToModel(r*(e-this.mouseStartX),s*(i-this.mouseStartY));this.modelGroup.position.addVectors(this.currentModelPos,u)}else if((0===o||1==this.mouseButton)&&0!==d){var f=Math.sin(d*Math.PI)/d;this.dq.x=Math.cos(d*Math.PI),this.dq.y=0,this.dq.z=f*l,this.dq.w=-f*h,this.rotationGroup.quaternion.set(1,0,0,0),this.rotationGroup.quaternion.multiply(this.dq),this.rotationGroup.quaternion.multiply(this.cq)}this.show()}}_handleContextMenu(t){if(t.preventDefault(),this.closeEnoughForClick(t)){var e=this.mouseStartX,i=this.mouseStartY,r=this.canvasOffset();let n=this.mouseXY(e,i),a=n.x,o=n.y,l=this.targetedObjects(a,o,this.contextMenuEnabledObjects);var s=null;l.length&&(s=l[0].clickable);r=this.canvasOffset(),e=this.mouseStartX-r.left,i=this.mouseStartY-r.top;this.userContextMenuHandler&&(this.userContextMenuHandler(s,e,i,l,t),this.isDragging=!1)}}setContainer(t){let e=(0,L.getElement)(t)||this.container;return this.initContainer(e),this}setBackgroundColor(t,e){(void 0===e||e<0||e>1)&&(e=1);var i=s.CC.color(t);return this.scene.fog.color=i,this.bgColor=i.getHex(),this.renderer.setClearColorHex(i.getHex(),e),this.show(),this}setProjection(t){this.camera.ortho="orthographic"===t,this.setSlabAndFog()}setViewStyle(t){if((t=t||{}).style=t.style||"",t.style.includes("outline")?this.renderer.enableOutline(t):this.renderer.disableOutline(),t.style.includes("ambientOcclusion")){var e={};t.strength&&(e.strength=t.strength),t.radius&&(e.radius=t.radius),this.renderer.enableAmbientOcclusion(e)}else this.renderer.disableAmbientOcclusion();return this}updateSize(){this.renderer.setSize(this.WIDTH,this.HEIGHT),this.ASPECT=this.renderer.getAspect(this.WIDTH,this.HEIGHT),this.renderer.setSize(this.WIDTH,this.HEIGHT),this.camera.aspect=this.ASPECT,this.camera.updateProjectionMatrix()}setWidth(t){return this.WIDTH=t||this.WIDTH,this.updateSize(),this}setHeight(t){return this.HEIGHT=t||this.HEIGHT,this.updateSize(),this}resize(){this.WIDTH=this.getWidth(),this.HEIGHT=this.getHeight();let t=!1;if(this.renderer.isLost()&&this.WIDTH>0&&this.HEIGHT>0){let e=!1,i=this.container.querySelector("canvas");i&&i!=this.renderer.getCanvas()?this.config.canvas=i:(i.remove(),this.config&&null!=this.config.canvas&&(delete this.config.canvas,e=!0)),this.setupRenderer(),this.initContainer(this.container),this.renderer.setClearColorHex(this.bgColor,this.config.backgroundAlpha),t=!0,e&&(this.config.canvas=this.renderer.getCanvas())}if(0==this.WIDTH||0==this.HEIGHT?this.animated&&this._viewer.pauseAnimate():this.animated&&this._viewer.resumeAnimate(),this.updateSize(),t){let t=this.renderer.supportedExtensions();if(t.regen=!0,this.viewers)for(let e=0,i=this.viewers.length;e<i;e++)for(let i=0,r=this.viewers[e].length;i<r;i++)this.viewers[e][i].render(null,t);this._viewer.render(null,t)}else this.show();return this}getModel(t){return void 0===t?0==this.models.length?null:this.models[this.models.length-1]:t instanceof GLModel?t:t in this.models?this.models[t]:0==this.models.length?null:this.models[this.models.length-1]}spin(t,e=1,i=!1){if(clearInterval(this.spinInterval),void 0===t&&(t="y"),"boolean"==typeof t){if(!t)return;t="y"}Array.isArray(t)&&(t={x:t[0],y:t[1],z:t[2]});var r=this;this.spinInterval=setInterval((function(){!r.getCanvas().isConnected&&r.renderer.isLost()&&clearInterval(r.spinInterval),(!i||r.container.checkVisibility&&r.container.checkVisibility())&&r.rotate(1*e,t)}),25)}animateMotion(t,e,i,r,s,n){var a=Math.ceil(t/20);a<1&&(a=1),this.incAnim();var o={mpos:this.modelGroup.position.clone(),rz:this.rotationGroup.position.z,rot:this.rotationGroup.quaternion.clone(),cam:this.lookingAt.clone()};if(e){let t=new Array(a);for(let e=0;e<a;e++){let l=(e+1)/a,h={mpos:o.mpos,rz:o.rz,rot:o.rot};h.mpos=i.clone().sub(o.mpos).multiplyScalar(l).add(o.mpos),h.rz=o.rz+l*(r-o.rz),h.rot=GLViewer.slerp(o.rot,s,l),h.cam=n.clone().sub(o.cam).multiplyScalar(l).add(o.cam),t[e]=h}let e=0,l=this,h=function(){var i=t[e];e+=1,l.modelGroup.position=i.mpos,l.rotationGroup.position.z=i.rz,l.rotationGroup.quaternion=i.rot,l.camera.lookAt(i.cam),e<t.length?setTimeout(h,20):l.decAnim(),l.show()};setTimeout(h,20)}else{var l={};let t=1/a;if(i&&(l.mpos=i.clone().sub(o.mpos).multiplyScalar(t)),void 0!==r&&null!=r&&(l.rz=t*(r-o.rz)),s){var h=GLViewer.slerp(o.rot,s,t);l.rot=o.rot.clone().inverse().multiply(h)}n&&(l.cam=n.clone().sub(o.cam).multiplyScalar(t));let e=0,c=this,d=function(){e+=1,l.mpos&&c.modelGroup.position.add(l.mpos),l.rz&&(c.rotationGroup.position.z+=l.rz),l.rot&&c.rotationGroup.quaternion.multiply(l.rot),l.cam&&(c.lookingAt.add(l.cam),c.camera.lookAt(c.lookingAt)),e<a?setTimeout(d,20):c.decAnim(),c.show()};setTimeout(d,20)}}rotate(t,e="y",i=0,r=!1){if("x"==e?e={x:1,y:0,z:0}:"y"==e?e={x:0,y:1,z:0}:"z"==e&&(e={x:0,y:0,z:1}),"vx"==e?e={vx:1,vy:0,vz:0}:"vy"==e?e={vx:0,vy:1,vz:0}:"vz"==e&&(e={vx:0,vy:0,vz:1}),void 0!==e.vx){var s=new $.Vector3(e.vx,e.vy,e.vz);s.applyQuaternion(this.rotationGroup.quaternion),e={x:s.x,y:s.y,z:s.z}}var n=function(t){var i,r,s,n=Math.sin(t/2),a=Math.cos(t/2);return i=e.x*n,r=e.y*n,s=e.z*n,new $.Quaternion(i,r,s,a).normalize()}(Math.PI*t/180);if(i){var a=(new $.Quaternion).copy(this.rotationGroup.quaternion).multiply(n);this.animateMotion(i,r,this.modelGroup.position,this.rotationGroup.position.z,a,this.lookingAt)}else this.rotationGroup.quaternion.multiply(n),this.show();return this}surfacesFinished(){for(var t in this.surfaces)if(!this.surfaces[t][0].done)return!1;return!0}getView(){if(!this.modelGroup)return[0,0,0,0,0,0,0,1];var t=this.modelGroup.position,e=this.rotationGroup.quaternion;return[t.x,t.y,t.z,this.rotationGroup.position.z,e.x,e.y,e.z,e.w]}setView(t,e){return void 0!==t&&(t instanceof Array||8!==t.length)&&this.modelGroup&&this.rotationGroup?(this.modelGroup.position.x=t[0],this.modelGroup.position.y=t[1],this.modelGroup.position.z=t[2],this.rotationGroup.position.z=t[3],this.rotationGroup.quaternion.x=t[4],this.rotationGroup.quaternion.y=t[5],this.rotationGroup.quaternion.z=t[6],this.rotationGroup.quaternion.w=t[7],void 0!==t[8]&&(this.rotationGroup.position.x=t[8],this.rotationGroup.position.y=t[9]),this.show(e),this):this}render(t,e){this.renderer.setViewport(),this.updateClickables();var i,r,s=this.getView();for(this.stateChangeCallback&&this.stateChangeCallback(this.getInternalState()),e||(e=this.renderer.supportedExtensions()),i=0;i<this.models.length;i++)this.models[i]&&this.models[i].globj(this.modelGroup,e);for(i=0;i<this.shapes.length;i++)this.shapes[i]&&(void 0===this.shapes[i].frame||this.viewer_frame<0||this.shapes[i].frame<0||this.shapes[i].frame==this.viewer_frame?this.shapes[i].globj(this.modelGroup,e):this.shapes[i].removegl(this.modelGroup));for(i=0;i<this.labels.length;i++)e.regen&&(this.labels[i].dispose(),this.modelGroup.remove(this.labels[i].sprite),this.labels[i].setContext(),this.modelGroup.add(this.labels[i].sprite)),this.labels[i]&&void 0!==this.labels[i].frame&&this.labels[i].frame>=0&&(this.modelGroup.remove(this.labels[i].sprite),(this.viewer_frame<0||this.labels[i].frame==this.viewer_frame)&&this.modelGroup.add(this.labels[i].sprite));for(i in this.surfaces)if(this.surfaces.hasOwnProperty(i)){var a=this.surfaces[i];for(r=0;r<a.length;r++)if(a.hasOwnProperty(r)){var o=a[r].geo;if(!a[r].finished||e.regen){o.verticesNeedUpdate=!0,o.elementsNeedUpdate=!0,o.normalsNeedUpdate=!0,o.colorsNeedUpdate=!0,o.buffersNeedUpdate=!0,a[r].mat.needsUpdate=!0,a[r].done&&(a[r].finished=!0),a[r].lastGL&&this.modelGroup.remove(a[r].lastGL);var l=null;if(l=a[r].mat instanceof n.LineBasicMaterial?new n.Line(o,a[r].mat):new n.Mesh(o,a[r].mat),a[r].mat.transparent&&0==a[r].mat.opacity?l.visible=!1:l.visible=!0,a[r].symmetries.length>1||1==a[r].symmetries.length&&!a[r].symmetries[r].isIdentity()){var h,c=new n.Object3D;for(h=0;h<a[r].symmetries.length;h++){var d=l.clone();d.matrix=a[r].symmetries[h],d.matrixAutoUpdate=!1,c.add(d)}a[r].lastGL=c,this.modelGroup.add(c)}else a[r].lastGL=l,this.modelGroup.add(l)}}}return this.setView(s),"function"==typeof t&&t(this),this}getModelList(t){let e=[];if(void 0===t||void 0===t.model)for(let t=0;t<this.models.length;t++)this.models[t]&&e.push(this.models[t]);else{let r=t.model;Array.isArray(r)||(r=[r]);for(let t=0;t<r.length;t++)if("number"==typeof r[t]){var i=r[t];i<0&&(i+=this.models.length),e.push(this.models[i])}else e.push(r[t])}return e}getAtomsFromSel(t){var e=[];void 0===t&&(t={});var i=this.getModelList(t);for(let r=0;r<i.length;r++)e=e.concat(i[r].selectedAtoms(t));return e}atomIsSelected(t,e){void 0===e&&(e={});for(var i=this.getModelList(e),r=0;r<i.length;r++)if(i[r].atomIsSelected(t,e))return!0;return!1}selectedAtoms(t){return this.getAtomsFromSel(t)}getUniqueValues(t,e){void 0===e&&(e={});var i=this.getAtomsFromSel(e),r={};for(var s in i){if(i[s].hasOwnProperty(t))r[i[s][t]]=!0}return Object.keys(r)}pdbData(t){for(var e=this.getAtomsFromSel(t),i="",r=0,s=e.length;r<s;++r)i+=e[r].pdbline+"\n";return i}zoom(t=2,e=0,i=!1){var r=(this.CAMERA_Z-this.rotationGroup.position.z)/t,s=this.CAMERA_Z-r;return e>0?this.animateMotion(e,i,this.modelGroup.position,this.adjustZoomToLimits(s),this.rotationGroup.quaternion,this.lookingAt):(this.rotationGroup.position.z=this.adjustZoomToLimits(s),this.show()),this}translate(t,e,i=0,r=!1){var s=t/this.WIDTH,n=e/this.HEIGHT,a=new $.Vector3(0,0,-this.CAMERA_Z);this.projector.projectVector(a,this.camera),a.x-=s,a.y-=n,this.projector.unprojectVector(a,this.camera),a.z=0;var o=this.lookingAt.clone().add(a);return i>0?this.animateMotion(i,r,this.modelGroup.position,this.rotationGroup.position.z,this.rotationGroup.quaternion,o):(this.lookingAt=o,this.camera.lookAt(this.lookingAt),this.show()),this}translateScene(t,e,i=0,r=!1){var s=this.screenOffsetToModel(t,e),n=this.modelGroup.position.clone().add(s);return i>0?this.animateMotion(i,r,this.modelGroup.position,this.rotationGroup.position.z,this.rotationGroup.quaternion,this.lookingAt):(this.modelGroup.position=n,this.show()),this}fitSlab(t){t=t||{};var e=this.getAtomsFromSel(t),i=(0,L.getExtent)(e),r=i[1][0]-i[0][0],s=i[1][1]-i[0][1],n=i[1][2]-i[0][2],a=Math.hypot(r,s,n);return a<5&&(a=5),this.slabNear=-a/1.9,this.slabFar=a/2,this}center(t={},e=0,i=!1){var r,s,n=this.getAtomsFromSel(t),a=(0,L.getExtent)(n);(0,L.isEmptyObject)(t)?(this.shapes.forEach((t=>{if(t&&t.boundingSphere&&t.boundingSphere.center){var e=t.boundingSphere.center,i=t.boundingSphere.radius;i>0?(n.push(new $.Vector3(e.x+i,e.y,e.z)),n.push(new $.Vector3(e.x-i,e.y,e.z)),n.push(new $.Vector3(e.x,e.y+i,e.z)),n.push(new $.Vector3(e.x,e.y-i,e.z)),n.push(new $.Vector3(e.x,e.y,e.z+i)),n.push(new $.Vector3(e.x,e.y,e.z-i))):n.push(e)}})),a=(0,L.getExtent)(n),r=n,s=a):(r=this.getAtomsFromSel({}),s=(0,L.getExtent)(r));var o=new $.Vector3(a[2][0],a[2][1],a[2][2]),l=s[1][0]-s[0][0],h=s[1][1]-s[0][1],c=s[1][2]-s[0][2],d=Math.hypot(l,h,c);d<5&&(d=5),this.slabNear=-d/1.9,this.slabFar=d/2,l=a[1][0]-a[0][0],h=a[1][1]-a[0][1],c=a[1][2]-a[0][2],(d=Math.hypot(l,h,c))<5&&(d=5);for(var u=25,f=0;f<n.length;f++)if(n[f]){var p=o.distanceToSquared(n[f]);p>u&&(u=p)}d=2*Math.sqrt(u);var g=o.clone().multiplyScalar(-1);return e>0?this.animateMotion(e,i,g,this.rotationGroup.position.z,this.rotationGroup.quaternion,this.lookingAt):(this.modelGroup.position=g,this.show()),this}zoomTo(t={},e=0,i=!1){let r=this.getAtomsFromSel(t),s=(0,L.getExtent)(r),n=s;if((0,L.isEmptyObject)(t)){let t=r&&r.length;if(this.shapes.forEach((t=>{if(t&&t.boundingSphere)if(t.boundingSphere.box){let e=t.boundingSphere.box;r.push(new $.Vector3(e.min.x,e.min.y,e.min.z)),r.push(new $.Vector3(e.max.x,e.max.y,e.max.z))}else if(t.boundingSphere.center){var e=t.boundingSphere.center,i=t.boundingSphere.radius;i>0?(r.push(new $.Vector3(e.x+i,e.y,e.z)),r.push(new $.Vector3(e.x-i,e.y,e.z)),r.push(new $.Vector3(e.x,e.y+i,e.z)),r.push(new $.Vector3(e.x,e.y-i,e.z)),r.push(new $.Vector3(e.x,e.y,e.z+i)),r.push(new $.Vector3(e.x,e.y,e.z-i))):r.push(e)}})),n=(0,L.getExtent)(r),!t)for(let t=0;t<3;t++)s[2][t]=(n[0][t]+n[1][t])/2}else{let t=this.getAtomsFromSel({});n=(0,L.getExtent)(t)}var a=new $.Vector3(s[2][0],s[2][1],s[2][2]),o=n[1][0]-n[0][0],l=n[1][1]-n[0][1],h=n[1][2]-n[0][2],c=Math.hypot(o,l,h);c<5&&(c=5),this.slabNear=-c/1.9,this.slabFar=c/2,0===Object.keys(t).length&&(this.slabNear=Math.min(2*-c,-50),this.slabFar=Math.max(2*c,50));var d=this.config.minimumZoomToDistance||5;o=s[1][0]-s[0][0],l=s[1][1]-s[0][1],h=s[1][2]-s[0][2],(c=Math.hypot(o,l,h))<d&&(c=d);for(var u=d*d,f=0;f<r.length;f++)if(r[f]){var p=a.distanceToSquared(r[f]);p>u&&(u=p)}c=2*Math.sqrt(u);var g=a.clone().multiplyScalar(-1),m=-(.5*c/Math.tan(Math.PI/180*this.camera.fov/2)-this.CAMERA_Z);return m=this.adjustZoomToLimits(m),e>0?this.animateMotion(e,i,g,m,this.rotationGroup.quaternion,this.lookingAt):(this.modelGroup.position=g,this.rotationGroup.position.z=m,this.show()),this}setSlab(t,e){this.slabNear=t,this.slabFar=e}getSlab(){return{near:this.slabNear,far:this.slabFar}}addLabel(t,e={},i,r=!1){if(i){var s=(0,L.getExtent)(this.getAtomsFromSel(i));e.position={x:s[2][0],y:s[2][1],z:s[2][2]}}var n=new Label(t,e);return n.setContext(),this.modelGroup.add(n.sprite),this.labels.push(n),r||this.show(),n}addResLabels(t,e,i=!1){let r=this.labels.length;return this.applyToModels("addResLabels",t,this,e,i),this.show(),this.labels.slice(r)}addPropertyLabels(t,e,i){return this.applyToModels("addPropertyLabels",t,e,this,i),this.show(),this}removeLabel(t){for(var e=0;e<this.labels.length;e++)if(this.labels[e]==t){this.labels.splice(e,1),t.dispose(),this.modelGroup.remove(t.sprite);break}return this.show(),this}removeAllLabels(){for(var t=0;t<this.labels.length;t++)this.labels[t]&&this.labels[t].sprite&&this.modelGroup.remove(this.labels[t].sprite);return this.labels.splice(0,this.labels.length),this.show(),this}hideAllLabels(){for(var t=0;t<this.labels.length;t++)this.labels[t]&&this.labels[t].hide();return this.show(),this}showAllLabels(){for(var t=0;t<this.labels.length;t++)this.labels[t]&&this.labels[t].show();return this.show(),this}setLabelStyle(t,e){return this.modelGroup.remove(t.sprite),t.dispose(),t.stylespec=e,t.setContext(),this.modelGroup.add(t.sprite),this.show(),t}setLabelText(t,e){return this.modelGroup.remove(t.sprite),t.dispose(),t.text=e,t.setContext(),this.modelGroup.add(t.sprite),this.show(),t}addShape(t){var e=new GLShape(t=t||{});return e.shapePosition=this.shapes.length,this.shapes.push(e),e}removeShape(t){if(!t)return this;for(t.removegl(this.modelGroup),delete this.shapes[t.shapePosition];this.shapes.length>0&&void 0===this.shapes[this.shapes.length-1];)this.shapes.pop();return this}removeAllShapes(){for(var t=0;t<this.shapes.length;t++){var e=this.shapes[t];e&&e.removegl(this.modelGroup)}return this.shapes.splice(0,this.shapes.length),this}getSelectionCenter(t){if(t.hasOwnProperty("x")&&t.hasOwnProperty("y")&&t.hasOwnProperty("z"))return t;var e=this.getAtomsFromSel(t);if(0==e.length)return{x:0,y:0,z:0};var i=(0,L.getExtent)(e);return{x:i[0][0]+(i[1][0]-i[0][0])/2,y:i[0][1]+(i[1][1]-i[0][1])/2,z:i[0][2]+(i[1][2]-i[0][2])/2}}addSphere(t){(t=t||{}).center=this.getSelectionCenter(t.center);var e=new GLShape(t);return e.shapePosition=this.shapes.length,e.addSphere(t),this.shapes.push(e),e.finalize(),e}addBox(t={}){null!=t.corner&&(t.corner=this.getSelectionCenter(t.corner)),null!=t.center&&(t.center=this.getSelectionCenter(t.center));var e=new GLShape(t);return e.shapePosition=this.shapes.length,e.addBox(t),this.shapes.push(e),e.finalize(),e}addArrow(t={}){t.start=this.getSelectionCenter(t.start),t.end=this.getSelectionCenter(t.end);var e=new GLShape(t);return e.shapePosition=this.shapes.length,e.addArrow(t),this.shapes.push(e),e.finalize(),e}addCylinder(t={}){t.start=this.getSelectionCenter(t.start),t.end=this.getSelectionCenter(t.end);var e=new GLShape(t);return e.shapePosition=this.shapes.length,t.dashed?e.addDashedCylinder(t):e.addCylinder(t),this.shapes.push(e),e.finalize(),e}addCurve(t={}){var e=new GLShape(t);return e.shapePosition=this.shapes.length,e.addCurve(t),this.shapes.push(e),e.finalize(),e}addLine(t={}){t.start=this.getSelectionCenter(t.start),t.end=this.getSelectionCenter(t.end),t.wireframe=!0;var e=new GLShape(t);return e.shapePosition=this.shapes.length,t.dashed?e=this.addLineDashed(t,e):e.addLine(t),this.shapes.push(e),e.finalize(),e}addUnitCell(t,e){t=this.getModel(t),(e=e||{alabel:"a",blabel:"b",clabel:"c"}).box=e.box||{},e.astyle=e.astyle||{color:"red",radius:.1,midpos:-1},e.bstyle=e.bstyle||{color:"green",radius:.1,midpos:-1},e.cstyle=e.cstyle||{color:"blue",radius:.1,midpos:-1},e.alabelstyle=e.alabelstyle||{fontColor:"red",showBackground:!1,alignment:"center",inFront:!1},e.blabelstyle=e.blabelstyle||{fontColor:"green",showBackground:!1,alignment:"center",inFront:!1},e.clabelstyle=e.clabelstyle||{fontColor:"blue",showBackground:!1,alignment:"center",inFront:!1},t.unitCellObjects&&this.removeUnitCell(t),t.unitCellObjects={shapes:[],labels:[]};var i=t.getCrystData(),r=null;if(i){if(i.matrix)r=i.matrix;else{var s,n,a,o=i.a,l=i.b,h=i.c,c=i.alpha,d=i.beta,u=i.gamma;c=c*Math.PI/180,d=d*Math.PI/180,u=u*Math.PI/180,s=Math.cos(d),n=(Math.cos(c)-Math.cos(d)*Math.cos(u))/Math.sin(u),a=Math.sqrt(Math.max(0,1-s*s-n*n)),r=new $.Matrix3(o,l*Math.cos(u),h*s,0,l*Math.sin(u),h*n,0,0,h*a)}var f=[new $.Vector3(0,0,0),new $.Vector3(1,0,0),new $.Vector3(0,1,0),new $.Vector3(0,0,1),new $.Vector3(1,1,0),new $.Vector3(0,1,1),new $.Vector3(1,0,1),new $.Vector3(1,1,1)];if(i.matrix4)for(let t=0;t<f.length;t++)i.size&&f[t].multiplyVectors(f[t],i.size),f[t]=f[t].applyMatrix4(i.matrix4);else for(let t=0;t<f.length;t++)f[t]=f[t].applyMatrix3(r);if(e.box&&!e.box.hidden){e.box.wireframe=!0;var p=new GLShape(e.box);p.shapePosition=this.shapes.length,p.addLine({start:f[0],end:f[1]}),p.addLine({start:f[0],end:f[2]}),p.addLine({start:f[1],end:f[4]}),p.addLine({start:f[2],end:f[4]}),p.addLine({start:f[0],end:f[3]}),p.addLine({start:f[3],end:f[5]}),p.addLine({start:f[2],end:f[5]}),p.addLine({start:f[1],end:f[6]}),p.addLine({start:f[4],end:f[7]}),p.addLine({start:f[6],end:f[7]}),p.addLine({start:f[3],end:f[6]}),p.addLine({start:f[5],end:f[7]}),this.shapes.push(p),t.unitCellObjects.shapes.push(p),p.finalize()}if(!e.astyle.hidden){e.astyle.start=f[0],e.astyle.end=f[1];let i=this.addArrow(e.astyle);t.unitCellObjects.shapes.push(i)}if(!e.bstyle.hidden){e.bstyle.start=f[0],e.bstyle.end=f[2];let i=this.addArrow(e.bstyle);t.unitCellObjects.shapes.push(i)}if(!e.cstyle.hidden){e.cstyle.start=f[0],e.cstyle.end=f[3];let i=this.addArrow(e.cstyle);t.unitCellObjects.shapes.push(i)}if(e.alabel){e.alabelstyle.position=f[1];let i=this.addLabel(e.alabel,e.alabelstyle);t.unitCellObjects.labels.push(i)}if(e.blabel){e.blabelstyle.position=f[2];let i=this.addLabel(e.blabel,e.blabelstyle);t.unitCellObjects.labels.push(i)}if(e.clabel){e.clabelstyle.position=f[3];let i=this.addLabel(e.clabel,e.clabelstyle);t.unitCellObjects.labels.push(i)}}}removeUnitCell(t){if((t=this.getModel(t)).unitCellObjects){let e=this;t.unitCellObjects.shapes.forEach((function(t){e.removeShape(t)})),t.unitCellObjects.labels.forEach((function(t){e.removeLabel(t)}))}delete t.unitCellObjects}replicateUnitCell(t=3,e=t,i=e,r,s,n){let a=(r=this.getModel(r)).getCrystData();if(a){const o=r.selectedAtoms({}),l=a.matrix;let h=function(t){return t%2==0?-t/2:Math.ceil(t/2)};t<=1&&e<=1&&i<=1&&(n=!0,t=e=i=3);let c=function(t,e,i){return!1};if(n){const t=(new $.Matrix3).getInverse3(l);c=function(e,i,r){let s=new $.Vector3(e,i,r).applyMatrix3(t);return!(s.x>-1e-4&&s.x<1.0001&&s.y>-1e-4&&s.y<1.0001&&s.z>-1e-4&&s.z<1.0001)}}for(let s=0;s<t;s++)for(let t=0;t<e;t++)for(let e=0;e<i;e++){if(0==s&&0==t&&0==e)continue;let i=new $.Vector3(h(s),h(t),h(e));i.applyMatrix3(l);let n=[];for(let t=0;t<o.length;t++){let e=o[t].x+i.x,r=o[t].y+i.y,s=o[t].z+i.z;if(c(e,r,s))continue;let a={};for(let e in o[t])a[e]=o[t][e];a.x=e,a.y=r,a.z=s,n.push(a)}r.addAtoms(n)}s&&r.assignBonds()}}addLineDashed(t,e){var i,r;t.dashLength=t.dashLength||.5,t.gapLength=t.gapLength||.5,i=t.start?new $.Vector3(t.start.x||0,t.start.y||0,t.start.z||0):new $.Vector3(0,0,0),r=t.end?new $.Vector3(t.end.x,t.end.y||0,t.end.z||0):new $.Vector3(0,0,0);var s,n,a,o=new $.Vector3,l=new $.Vector3,h=new $.Vector3,c=i.clone(),d=0;for(o.subVectors(r,i),s=o.length(),o.normalize(),l=o.clone(),h=o.clone(),l.multiplyScalar(t.dashLength),h.multiplyScalar(t.gapLength),n=l.length(),a=h.length();d<s;){if(d+n>s){t.start=i,t.end=r,e.addLine(t);break}c.addVectors(i,l),t.start=i,t.end=c,e.addLine(t),i=c.clone(),d+=n,c.addVectors(i,h),i=c.clone(),d+=a}return e.finalize(),e}addCustom(t){var e=new GLShape(t=t||{});return e.shapePosition=this.shapes.length,e.addCustom(t),this.shapes.push(e),e.finalize(),e}addVolumetricData(t,e,i={}){var r=new st.VolumeData(t,e);return i.hasOwnProperty("transferfn")?this.addVolumetricRender(r,i):this.addIsosurface(r,i)}addIsosurface(t,e={},i){var r=new GLShape(e);return r.shapePosition=this.shapes.length,r.addIsosurface(t,e,i,this),this.shapes.push(r),r}addVolumetricRender(t,e){var i=new GLVolumetricRender(t,e=e||{},this);return i.shapePosition=this.shapes.length,this.shapes.push(i),i}hasVolumetricRender(){return this.renderer.supportsVolumetric()}enableFog(t){t?this.scene.fog=new n.Fog(this.bgColor,100,200):(this.config.disableFog=!0,this.show())}setFrame(t){this.viewer_frame=t;let e=this;return new Promise((function(i){var r=e.models.map((function(i){return i.setFrame(t,e)}));Promise.all(r).then((function(){i()}))}))}getFrame(){return this.viewer_frame}getNumFrames(){var t=0;for(let e=0;e<this.models.length;e++)this.models[e].getNumFrames()>t&&(t=this.models[e].getNumFrames());for(let e=0;e<this.shapes.length;e++)this.shapes[e].frame&&this.shapes[e].frame>=t&&(t=this.shapes[e].frame+1);for(let e=0;e<this.labels.length;e++)this.labels[e].frame&&this.labels[e].frame>=t&&(t=this.labels[e].frame+1);return t}animate(t){this.incAnim();var e=100,i="forward",r=1/0;(t=t||{}).interval&&(e=t.interval),t.loop&&(i=t.loop),t.reps&&(r=t.reps);var s=this.getNumFrames(),n=this,a=0;t.startFrame&&(a=t.startFrame%s);var o=1;t.step&&(r/=o=t.step);var l,h,c=0,d=s*r,u=new Date,f=function(t){u=new Date,"forward"==t?n.setFrame(a).then((function(){a=(a+o)%s,l()})):"backward"==t?n.setFrame(s-1-a).then((function(){a=(a+o)%s,l()})):n.setFrame(a).then((function(){o*=(a+=o)%(s-1)==0?-1:1,l()}))};return l=function(){if(n.render(),n.getCanvas().isConnected)if(++c>=d||!n.isAnimated())h.cancel(),n.animationTimers.delete(h),n.decAnim();else{var t=e-((new Date).getTime()-u.getTime());t=t>0?t:0,n.animationTimers.delete(h),h=new L.PausableTimer(f,t,i),n.animationTimers.add(h)}else n.stopAnimate()},h=new L.PausableTimer(f,0,i),this.animationTimers.add(h),this}stopAnimate(){return this.animated=0,this.animationTimers.forEach((function(t){t.cancel()})),this.animationTimers=new Set,this}pauseAnimate(){return this.animationTimers.forEach((function(t){t.pause()})),this}resumeAnimate(){return this.animationTimers.forEach((function(t){t.resume()})),this}isAnimated(){return this.animated>0}getModelOpt(t){return t&&!t.defaultcolors?(t.defaultcolors=this.defaultcolors,t.cartoonQuality=t.cartoonQuality||this.config.cartoonQuality):void 0===t&&(t={defaultcolors:this.defaultcolors,cartoonQuality:this.config.cartoonQuality}),t}addModel(t,e="",i){i=this.getModelOpt(i);var r=new GLModel(this.models.length,i);return r.addMolData(t,e,i),this.models.push(r),r}addModels(t,e,i){(i=this.getModelOpt(i)).multimodel=!0,i.frames=!0;for(var r=GLModel.parseMolData(t,e,i),s=0;s<r.length;s++){var n=new GLModel(this.models.length,i);n.setAtomDefaults(r[s]),n.addFrame(r[s]),n.setFrame(0),r.modelData&&n.setModelData(r.modelData[s]),n.setDontDuplicateAtoms(!i.duplicateAssemblyAtoms),this.models.push(n)}return this.models}addModelsAsFrames(t,e,i){(i=this.getModelOpt(i)).multimodel=!0,i.frames=!0;var r=new GLModel(this.models.length,i);return r.addMolData(t,e,i),this.models.push(r),r}addAsOneMolecule(t,e,i){(i=this.getModelOpt(i)).multimodel=!0,i.onemol=!0;var r=new GLModel(this.models.length,i);return r.addMolData(t,e,i),this.models.push(r),r}removeModel(t){if(t=this.getModel(t)){for(t.removegl(this.modelGroup),delete this.models[t.getID()];this.models.length>0&&void 0===this.models[this.models.length-1];)this.models.pop();return this}}removeAllModels(){for(var t=0;t<this.models.length;t++){var e=this.models[t];e&&e.removegl(this.modelGroup)}return this.models.splice(0,this.models.length),this}exportJSON(t,e){var i={};return i.m=void 0===e?this.models.map((function(e){return e.toCDObject(t)})):[this.models[e].toCDObject()],JSON.stringify(i)}exportVRML(){var t=this.modelGroup;this.applyToModels("removegl",this.modelGroup),this.modelGroup=new n.Object3D,this.render(null,{supportsImposters:!1,supportsAIA:!1,regen:!0});var e="#VRML V2.0 utf8\n"+this.modelGroup.vrml()+"\n";return this.applyToModels("removegl",this.modelGroup),this.modelGroup=t,e}createModelFrom(t,e=!1){for(var i=new GLModel(this.models.length,this.defaultcolors),r=0;r<this.models.length;r++)if(this.models[r]){var s=this.models[r].selectedAtoms(t);i.addAtoms(s),e&&this.models[r].removeAtoms(s)}return this.models.push(i),i}applyToModels(t,e,i,r,s,n,a){for(var o=this.getModelList(e),l=0;l<o.length;l++)o[l][t](e,i,r,s,n,a)}setStyle(t,e){return void 0===e&&(e=t,t={}),this.applyToModels("setStyle",t,e,!1),this}addStyle(t,e){return void 0===e&&(e=t,t={}),this.applyToModels("setStyle",t,e,!0),this}setClickable(t,e,i){return this.applyToModels("setClickable",t,e,i),this}setHoverable(t,e,i,r){return this.applyToModels("setHoverable",t,e,i,r),this}enableContextMenu(t,e){return this.applyToModels("enableContextMenu",t,e),this}vibrate(t,e,i,r){return this.applyToModels("vibrate",t,e,i,this,r),this}setColorByProperty(t,e,i,r){return this.applyToModels("setColorByProperty",t,e,i,r),this}setColorByElement(t,e){return this.applyToModels("setColorByElement",t,e),this}static getAtomsWithin(t,e){var i=[];for(let s=0;s<t.length;s++){var r=t[s];void 0!==r&&(r.x<e[0][0]||r.x>e[1][0]||r.y<e[0][1]||r.y>e[1][1]||r.z<e[0][2]||r.z>e[1][2]||i.push(r))}return i}static volume(t){return(t[1][0]-t[0][0])*(t[1][1]-t[0][1])*(t[1][2]-t[0][2])}carveUpExtent(t,e,i){let r=[],s={};for(let t=0,i=e.length;t<i;t++)s[e[t].index]=t;let n=function(t){let e=[];for(let i=0,r=t.length;i<r;i++)t[i].index in s&&e.push(s[t[i].index]);return e},a=function(t){let e=[];return e[0]=[t[0][0],t[0][1],t[0][2]],e[1]=[t[1][0],t[1][1],t[1][2]],e},o=function(t){if(GLViewer.volume(t)<GLViewer.maxVolume)return[t];var e,i=t[1][0]-t[0][0],r=t[1][1]-t[0][1],s=t[1][2]-t[0][2];e=i>r&&i>s?0:r>i&&r>s?1:2;var n=a(t),l=a(t),h=(t[1][e]-t[0][e])/2+t[0][e];n[1][e]=h,l[0][e]=h;var c=o(n),d=o(l);return c.concat(d)},l=o(t);for(let t=0,s=l.length;t<s;t++){let s=a(l[t]);s[0][0]-=6,s[0][1]-=6,s[0][2]-=6,s[1][0]+=6,s[1][1]+=6,s[1][2]+=6;let o=GLViewer.getAtomsWithin(e,s),h=GLViewer.getAtomsWithin(i,l[t]);r.push({extent:l[t],atoms:n(o),toshow:n(h)})}return r}static generateSurfaceMesh(t,e,i){var r=new n.Geometry(!0),a=r.updateGeoGroup(0),o=[];for(let e=0,i=t.length;e<i;e++){var l=t[e];l&&(void 0!==l.surfaceColor?o[e]=l.surfaceColor:l.color&&(o[e]=s.CC.color(l.color)))}var h=a.vertexArray,c=e.vertices;for(let t=0,e=c.length;t<e;t++){let e=3*a.vertices;h[e]=c[t].x,h[e+1]=c[t].y,h[e+2]=c[t].z,a.vertices++}var d=a.colorArray;let u=a.atomArray;if(i.voldata&&i.volscheme){var f=i.volscheme,p=i.voldata,g=f.range()||[-1,1];for(let e=0,i=c.length;e<i;e++){let i=c[e].atomid,r=p.getVal(c[e].x,c[e].y,c[e].z),n=s.CC.color(f.valueToHex(r,g)),a=3*e;d[a]=n.r,d[a+1]=n.g,d[a+2]=n.b,u[e]=t[i]}}else if(o.length>0)for(let e=0,i=c.length;e<i;e++){let i=c[e].atomid,r=3*e;d[r]=o[i].r,d[r+1]=o[i].g,d[r+2]=o[i].b,u[e]=t[i]}var m=e.faces;a.faceidx=m.length,r.initTypedArrays();var v,_,y,b,x=a.vertexArray,w=a.normalArray;for(let t=0,e=m.length;t<e;t+=3){var A=3*m[t],C=3*m[t+1],S=3*m[t+2];v=new $.Vector3(x[A],x[A+1],x[A+2]),_=new $.Vector3(x[C],x[C+1],x[C+2]),(y=new $.Vector3(x[S],x[S+1],x[S+2])).subVectors(y,_),v.subVectors(v,_),y.cross(v),(b=y).normalize(),w[A]+=b.x,w[C]+=b.x,w[S]+=b.x,w[A+1]+=b.y,w[C+1]+=b.y,w[S+1]+=b.y,w[A+2]+=b.z,w[C+2]+=b.z,w[S+2]+=b.z}return a.faceArray=new Uint16Array(m),new n.Mesh(r,i)}static generateMeshSyncHelper(t,e,i,r,s,n){var a=new ProteinSurface;return a.initparm(e,1!==t,n),a.fillvoxels(s,i),a.buildboundary(),t!=Q.SES&&t!=Q.MS||(a.fastdistancemap(),a.boundingatom(!1),a.fillvoxelswaals(s,i)),a.marchingcube(t),a.getFacesAndVertices(r)}static getMatWithStyle(t){let e=null;for(var i in e=t.onesided?new n.MeshLambertMaterial:new n.MeshDoubleLambertMaterial,e.vertexColors=n.Coloring.VertexColors,t)"color"===i||"map"===i||t.hasOwnProperty(i)&&(e[i]=t[i]);return void 0!==t.opacity&&(1===t.opacity?e.transparent=!1:e.transparent=!0),e}addMesh(t){var e={geo:t.geometry,mat:t.material,done:!0,finished:!1},i=this.nextSurfID();return this.surfaces[i]=[e],i}static shallowCopy(t){var e=[];let i=t.length;for(let r=0;r<i;r++)e[r]=(0,L.extend)({},t[r]);return e}addSurface(t,e={},i={},s,a,o){let l=this.nextSurfID(),h=null,c=this,d=Q.VDW;"string"==typeof t?void 0!==GLViewer.surfaceTypeMap[t.toUpperCase()]?d=GLViewer.surfaceTypeMap[t]:console.log("Surface type : "+t+" is not recognized"):"number"==typeof t&&(d=t);var u=null,f=null,p=GLViewer.shallowCopy(this.getAtomsFromSel(i));u=s?GLViewer.shallowCopy(this.getAtomsFromSel(s)):p,(0,L.adjustVolumeStyle)(e);var g,m=!1;for(g=0;g<this.models.length;g++)if(this.models[g]){var v=this.models[g].getSymmetries();if(v.length>1||1==v.length&&!v[0].isIdentity()){m=!0;break}}var _=function(t,i,s){var n;f=a?GLViewer.shallowCopy(c.getAtomsFromSel(a)):s;var o=(0,L.getExtent)(s,!0);if(e.map&&e.map.prop){var u=e.map.prop;let t=(0,r.getGradient)(e.map.scheme||e.map.gradient||new r.Gradient.RWB),i=t.range();i||(i=(0,L.getPropertyRange)(s,u)),e.colorscheme={prop:u,gradient:t}}for(let t=0,r=i.length;t<r;t++)(n=i[t]).surfaceColor=(0,L.getColorFromStyle)(n,e);var p=GLViewer.volume(o),g=c.carveUpExtent(o,i,s);if(f&&f.length&&f.length>0){var m=(0,L.getExtent)(f,!0);g.sort((function(t,e){var i=function(t,e){var i=t.extent,r=i[1][0]-i[0][0],s=i[1][1]-i[0][1],n=i[1][2]-i[0][2],a=r-e[2][0];a*=a;var o=s-e[2][1];o*=o;var l=n-e[2][2];return a+o+(l*=l)};return i(t,m)-i(e,m)}))}var v=[];for(let t=0,e=i.length;t<e;t++)n=i[t],v[t]={x:n.x,y:n.y,z:n.z,serial:t,elem:n.elem};if(!!tt){var _=function(e){return new Promise((function(r){for(var s=GLViewer.generateMeshSyncHelper(d,g[e].extent,g[e].atoms,g[e].toshow,v,p),n=wt({vertexArr:s.vertices,faceArr:s.faces}),a=0,o=n.length;a<o;a++){s={vertices:n[a].vertexArr,faces:n[a].faceArr};var l=GLViewer.generateSurfaceMesh(i,s,h);(0,L.mergeGeos)(t.geo,l)}c.render(),r()}))},y=[];for(let t=0;t<g.length;t++)y.push(_(t));return Promise.all(y).then((function(){return t.done=!0,Promise.resolve(l)}))}var b=[];d<0&&(d=0);for(let t=0,e=GLViewer.numWorkers;t<e;t++){var x=new Worker($3Dmol.SurfaceWorker);b.push(x),x.postMessage({type:-1,atoms:v,volume:p})}return new Promise((function(e,r){var s=0,n=function(){b&&b.length&&b.forEach((function(t){t&&t.terminate&&t.terminate()}))},a=function(r){for(var a=wt({vertexArr:r.data.vertices,faceArr:r.data.faces}),o=0,d=a.length;o<d;o++){var u={vertices:a[o].vertexArr,faces:a[o].faceArr},f=GLViewer.generateSurfaceMesh(i,u,h);(0,L.mergeGeos)(t.geo,f)}c.render(),++s==g.length&&(t.done=!0,n(),e(l))},o=function(t){n(),console.log(t.message+" ("+t.filename+":"+t.lineno+")"),r(t)};for(let t=0;t<g.length;t++){var u=b[t%b.length];u.onmessage=a,u.onerror=o,u.postMessage({type:d,expandedExtent:g[t].extent,extendedAtoms:g[t].atoms,atomsToShow:g[t].toshow})}}))};e=e||{},h=GLViewer.getMatWithStyle(e);var y=[];y.style=e,y.atomsel=i,y.allsel=s,y.focus=a;var b=null;if(m){var x={},w={};for(g=0;g<this.models.length;g++)x[g]=[],w[g]=[];for(g=0;g<u.length;g++)x[u[g].model].push(u[g]);for(g=0;g<p.length;g++)w[p[g].model].push(p[g]);var A=[];for(g=0;g<this.models.length;g++)w[g].length>0&&(y.push({geo:new n.Geometry(!0),mat:h,done:!1,finished:!1,symmetries:this.models[g].getSymmetries()}),A.push(_(y[y.length-1],x[g],w[g])));b=Promise.all(A)}else y.push({geo:new n.Geometry(!0),mat:h,done:!1,finished:!1,symmetries:[new $.Matrix4]}),b=_(y[y.length-1],u,p);return this.surfaces[l]=y,b.surfid=l,o&&"function"==typeof o?(b.then((function(t){o(t)})),l):b}setSurfaceMaterialStyle(t,e){if((0,L.adjustVolumeStyle)(e),this.surfaces[t]){var i=this.surfaces[t];for(let t=0;t<i.length;t++){var r=i[t].mat=GLViewer.getMatWithStyle(e);if(i[t].mat.side=n.FrontSide,e.color){i[t].mat.color=s.CC.color(e.color),i[t].geo.colorsNeedUpdate=!0;const r=s.CC.color(e.color);i[t].geo.setColor(r)}else if(r.voldata&&r.volscheme){const e=r.volscheme,n=r.voldata,a=s.CC,o=e.range()||[-1,1];i[t].geo.setColors((function(t,i,r){let s=n.getVal(t,i,r);return a.color(e.valueToHex(s,o))}))}else{i[t].geo.colorsNeedUpdate=!0;for(let r of i[t].geo.geometryGroups)for(let t=0;t<r.vertices;t++){let i=(0,L.getColorFromStyle)(r.atomArray[t],e),s=3*t;r.colorArray[s]=i.r,r.colorArray[s+1]=i.g,r.colorArray[s+2]=i.b}}i[t].finished=!1}}return this}getSurface(t){return this.surfaces[t]}removeSurface(t){for(var e=this.surfaces[t],i=0;i<e.length;i++)e[i]&&e[i].lastGL&&(void 0!==e[i].geo&&e[i].geo.dispose(),void 0!==e[i].mat&&e[i].mat.dispose(),this.modelGroup.remove(e[i].lastGL));return delete this.surfaces[t],this.show(),this}removeAllSurfaces(){for(var t in this.surfaces)if(this.surfaces.hasOwnProperty(t)){for(var e=this.surfaces[t],i=0;i<e.length;i++)e[i]&&e[i].lastGL&&(void 0!==e[i].geo&&e[i].geo.dispose(),void 0!==e[i].mat&&e[i].mat.dispose(),this.modelGroup.remove(e[i].lastGL));delete this.surfaces[t]}return this.show(),this}jmolMoveTo(){var t=this.modelGroup.position,e="center { "+-t.x+" "+-t.y+" "+-t.z+" }; ",i=this.rotationGroup.quaternion;return e+="moveto .5 quaternion { "+i.x+" "+i.y+" "+i.z+" "+i.w+" };"}clear(){return this.removeAllSurfaces(),this.removeAllModels(),this.removeAllLabels(),this.removeAllShapes(),this.show(),this}mapAtomProperties(t,e){e=e||{};var i=this.getAtomsFromSel(e);if("function"==typeof t)for(let e=0,r=i.length;e<r;e++){t(i[e])}else for(let e=0,n=i.length;e<n;e++){var r=i[e];for(let e=0,i=t.length;e<i;e++){let i=t[e];if(i.props)for(var s in i.props)i.props.hasOwnProperty(s)&&this.atomIsSelected(r,i)&&(r.properties||(r.properties={}),r.properties[s]=i.props[s])}}return this}linkViewer(t){return this.linkedViewers.push(t),this}getPerceivedDistance(){return this.CAMERA_Z-this.rotationGroup.position.z}setPerceivedDistance(t){this.rotationGroup.position.z=this.CAMERA_Z-t}setAutoEyeSeparation(t,e){var i=this.getPerceivedDistance();return e||(e=5),t||this.camera.position.x>0?this.camera.position.x=i*Math.tan(Math.PI/180*e):this.camera.position.x=-i*Math.tan(Math.PI/180*e),this.camera.lookAt(new $.Vector3(0,0,this.rotationGroup.position.z)),this.camera.position.x}setDefaultCartoonQuality(t){this.config.cartoonQuality=t}}function qt(t,e){if(t=(0,L.getElement)(t)){e=e||{};try{return new GLViewer(t,e)}catch(t){throw"error creating viewer: "+t}}}function Yt(t,e={},i={}){if(t=(0,L.getElement)(t)){var r=[],s=document.createElement("canvas");i.rows=e.rows,i.cols=e.cols,i.control_all=null!=e.control_all&&e.control_all,t.appendChild(s);try{for(var n=0;n<e.rows;n++){for(var a=[],o=0;o<e.cols;o++){i.row=n,i.col=o,i.canvas=s,i.viewers=r,i.control_all=e.control_all;var l=qt(t,(0,L.extend)({},i));a.push(l)}r.unshift(a)}}catch(t){throw"error creating viewer grid: "+t}return r}}function Zt(t){var e=this;if(t=(0,L.getElement)(t)){var i=Yt(t,{rows:1,cols:2,control_all:!0});this.glviewer1=i[0][0],this.glviewer2=i[0][1],this.glviewer1.setAutoEyeSeparation(!1),this.glviewer2.setAutoEyeSeparation(!0),this.glviewer1.linkViewer(this.glviewer2),this.glviewer2.linkViewer(this.glviewer1);for(var r=Object.getOwnPropertyNames(this.glviewer1.__proto__).filter((function(t){return"function"==typeof e.glviewer1[t]})),s=0;s<r.length;s++)this[r[s]]=function(t){return function(){return[this.glviewer1[t].apply(this.glviewer1,arguments),this.glviewer2[t].apply(this.glviewer2,arguments)]}}(r[s]);this.setCoordinates=function(t,e,i){for(var r=0;r<t.length;r++)t[r].setCoordinates(e,i)},this.surfacesFinished=function(){return this.glviewer1.surfacesFinished()&&this.glviewer2.surfacesFinished()},this.isAnimated=function(){return this.glviewer1.isAnimated()||this.glviewer2.isAnimated()},this.render=function(t){this.glviewer1.render(),this.glviewer2.render(),t&&t(this)},this.getCanvas=function(){return this.glviewer1.getCanvas()}}}GLViewer.numWorkers=4,GLViewer.maxVolume=64e3,GLViewer.surfaceTypeMap={VDW:Q.VDW,MS:Q.MS,SAS:Q.SAS,SES:Q.SES};var Xt=!1,Kt=!1,Qt={};function $t(t,e){var i,r,n;if(null!=document.querySelector(".viewer_3Dmoljs")&&(Xt=!0),Xt){Kt=!0,t=null!=t?t:null;var a=0;document.querySelectorAll(".viewer_3Dmoljs").forEach((o=>{var l=[],h=[],c="";"static"==o.style.position&&(o.style.position="relative");var d=null;if(n=null,o.dataset.pdb)l.push("https://files.rcsb.org/view/"+o.dataset.pdb+".pdb"),h.push("pdb");else if(o.dataset.cid)h.push("sdf"),l.push("https://pubchem.ncbi.nlm.nih.gov/rest/pug/compound/cid/"+o.dataset.cid+"/SDF?record_type=3d");else if(o.dataset.href||o.dataset.url){if(c=o.dataset.href?o.dataset.href:o.dataset.url,l.push(c),"gz"==(n=c.substring(c.lastIndexOf(".")+1))){let t=c.substring(0,c.lastIndexOf(".")).lastIndexOf(".");n=c.substring(t+1)}h.push(n);var u=c.substring(c.lastIndexOf("/")+1,c.lastIndexOf("."));"/"==u&&(u=c.substring(c.lastIndexOf("/")+1)),o.dataset[h[h.length-1]]=u}let f=o.dataset;for(i in f)"pdb"===i.substring(0,3)&&"pdb"!==i?(l.push("https://files.rcsb.org/view/"+f[i]+".pdb"),h.push("pdb")):"href"===i.substring(0,4)&&"href"!==i?(c=f[i],l.push(c),h.push(c.substring(c.lastIndexOf(".")+1))):"cid"===i.substring(0,3)&&"cid"!==i&&(l.push("https://pubchem.ncbi.nlm.nih.gov/rest/pug/compound/cid/"+f[i]+"/SDF?record_type=3d"),h.push("sdf"));var p={};o.dataset.options&&(p=(0,L.specStringToObject)(o.dataset.options));var g=s.CC.color(o.dataset.backgroundcolor),m=o.dataset.backgroundalpha;m=null==m?1:parseFloat(m);var v={line:{}};o.dataset.style&&(v=(0,L.specStringToObject)(o.dataset.style));var _={};o.dataset.select&&(_=(0,L.specStringToObject)(o.dataset.select));var y=[],b=[],x=[],w={},A=null,C=o.dataset,S=/style(.+)/,M=/surface(.*)/,z=/labelres(.*)/,T=[];for(r in C)Object.prototype.hasOwnProperty.call(C,r)&&T.push(r);for(T.sort(),i=0;i<T.length;i++){r=T[i];var E,F,I,O=S.exec(r);O&&(E="select"+O[1],F=(0,L.specStringToObject)(C[E]),I=(0,L.specStringToObject)(C[r]),y.push([F,I])),(O=M.exec(r))&&(E="select"+O[1],F=(0,L.specStringToObject)(C[E]),I=(0,L.specStringToObject)(C[r]),b.push([F,I])),(O=z.exec(r))&&(E="select"+O[1],F=(0,L.specStringToObject)(C[E]),I=(0,L.specStringToObject)(C[r]),x.push([F,I])),"zoomto"==r&&(w=(0,L.specStringToObject)(C[r])),"spin"==r&&(A=(0,L.specStringToObject)(C[r]))}var D=function(t){for(t.setStyle(_,v),d&&d.createSelectionAndStyle(_,v),i=0;i<y.length;i++){let e=y[i][0]||{},r=y[i][1]||{line:{}};t.setStyle(e,r),d&&d.createSelectionAndStyle(_,v)}for(i=0;i<b.length;i++){let e=b[i][0]||{},r=b[i][1]||{};d?t.addSurface(Q.VDW,r,e,e).then((t=>{d.loadSurface("VDW",e,r,t)})):t.addSurface(Q.VDW,r,e,e)}for(i=0;i<x.length;i++){let e=x[i][0]||{},r=x[i][1]||{};t.addResLabels(e,r)}t.render(),t.zoomTo(w),A&&t.spin(A.axis,A.speed)};let k=t;try{var R=(0,L.specStringToObject)(o.dataset.config)||{};void 0===R.backgroundColor&&(R.backgroundColor=g),void 0===R.backgroundAlpha&&(R.backgroundAlpha=m),null==k?k=Qt[o.id||a++]=qt(o,R):(k.setBackgroundColor(g,m),k.setConfig(R),d&&d.initiateUI()),o.dataset.ui&&$3Dmol.StateManager&&(d=new $3Dmol.StateManager(k))}catch(t){console.log(t),o.textContent="WebGL appears to be disabled."}if(0!=l.length){let t=0,i=((r,s)=>function(n){c=l[t];var a=r.dataset.type||r.dataset.datatype||h[t];if(s.addModel(n,a,p),d){var o=r.dataset[h[t]];d.setModelTitle(o)}if(t+=1,t<l.length)(0,L.get)(l[t]).then(i);else{if(D(s),r.dataset.callback){(0,L.makeFunction)(r.dataset.callback)(s)}Kt=!1,e&&e(s)}})(o,k);n&&n.endsWith("gz")?(0,L.getbin)(l[0]).then(i):(0,L.get)(l[0]).then(i)}else{if(o.dataset.element){var P="#"+o.dataset.element,U=document.querySelector(P),B=U?U.textContent:"";n=o.dataset.type||o.dataset.datatype,k.addModel(B,n,p)}if(D(k),o.dataset.callback){(0,L.makeFunction)(o.dataset.callback)(k)}Kt=!1,e&&e(k)}}))}}document.onreadystatechange=()=>{"complete"===document.readyState&&$t()},window&&(window.$3Dmol=e)},865:(t,e,i)=>{"use strict";i.r(e),i.d(e,{CUBE:()=>a});var r=i(638),s=i(392);const n={1:"H",2:"He",3:"Li",4:"Be",5:"B",6:"C",7:"N",8:"O",9:"F",10:"Ne",11:"Na",12:"Mg",13:"Al",14:"Si",15:"P",16:"S",17:"Cl",18:"Ar",19:"K",20:"Ca",21:"Sc",22:"Ti",23:"V",24:"Cr",25:"Mn",26:"Fe",27:"Co",28:"Ni",29:"Cu",30:"Zn",31:"Ga",32:"Ge",33:"As",34:"Se",35:"Br",36:"Kr",37:"Rb",38:"Sr",39:"Y",40:"Zr",41:"Nb",42:"Mo",43:"Tc",44:"Ru",45:"Rh",46:"Pd",47:"Ag",48:"Cd",49:"In",50:"Sn",51:"Sb",52:"Te",53:"I",54:"Xe",55:"Cs",56:"Ba",71:"Lu",72:"Hf",73:"Ta",74:"W",75:"Re",76:"Os",77:"Ir",78:"Pt",79:"Au",80:"Hg",81:"Tl",82:"Pb",83:"Bi",84:"Po",85:"At",86:"Rn",87:"Fr",88:"Ra",104:"Rf",105:"Db",106:"Sg",107:"Bh",108:"Hs",109:"Mt",110:"Ds",111:"Rg",112:"Cn",113:"Nh",114:"Fl",115:"Mc",116:"Lv",117:"Ts",118:"Og",57:"La",58:"Ce",59:"Pr",60:"Nd",61:"Pm",62:"Sm",63:"Eu",64:"Gd",65:"Tb",66:"Dy",67:"Ho",68:"Er",69:"Tm",70:"Yb",89:"Ac",90:"Th",91:"Pa",92:"U",93:"Np",94:"Pu",95:"Am",96:"Cm",97:"Bk",98:"Cf",99:"Es",100:"Fm",101:"Md",102:"No"};function a(t,e){e=e||{};const i=[[]];let a=t.split(/\r?\n/);const o=void 0===e.assignBonds||e.assignBonds;if(a.length<6)return i;let l=a[2].replace(/^\s+/,"").replace(/\s+/g," ").split(" ");const h=Math.abs(parseFloat(l[0]));let c={origin:void 0,size:void 0,unit:void 0,matrix4:void 0,matrix:void 0};const d=c.origin=new r.Vector3(parseFloat(l[1]),parseFloat(l[2]),parseFloat(l[3]));l=a[3].replace(/^\s+/,"").replace(/\s+/g," ").split(" "),l=a[3].replace(/^\s+/,"").replace(/\s+/g," ").split(" ");const u=l[0]>0?.529177:1;d.multiplyScalar(u);const f=Math.abs(l[0]),p=new r.Vector3(parseFloat(l[1]),parseFloat(l[2]),parseFloat(l[3])).multiplyScalar(u);l=a[4].replace(/^\s+/,"").replace(/\s+/g," ").split(" ");const g=Math.abs(l[0]),m=new r.Vector3(parseFloat(l[1]),parseFloat(l[2]),parseFloat(l[3])).multiplyScalar(u);l=a[5].replace(/^\s+/,"").replace(/\s+/g," ").split(" ");const v=Math.abs(l[0]),_=new r.Vector3(parseFloat(l[1]),parseFloat(l[2]),parseFloat(l[3])).multiplyScalar(u);if(c.size={x:f,y:g,z:v},c.unit=new r.Vector3(p.x,m.y,_.z),0!=p.y||0!=p.z||0!=m.x||0!=m.z||0!=_.x||0!=_.y){c.matrix4=new r.Matrix4(p.x,m.x,_.x,0,p.y,m.y,_.y,0,p.z,m.z,_.z,0,0,0,0,1);let t=(new r.Matrix4).makeTranslation(d.x,d.y,d.z);c.matrix4=c.matrix4.multiplyMatrices(t,c.matrix4),c.matrix=c.matrix4.matrix3FromTopLeft(),c.origin=new r.Vector3(0,0,0),c.unit=new r.Vector3(1,1,1)}i.modelData=[{cryst:c}],a=a.splice(6,h);for(var y=i[i.length-1].length,b=y+a.length,x=y;x<b;++x){var w={};w.serial=x;var A=a[x-y].replace(/^\s+/,"").replace(/\s+/g," ").split(" ");w.elem=n[A[0]],w.x=parseFloat(A[2])*u,w.y=parseFloat(A[3])*u,w.z=parseFloat(A[4])*u,w.hetflag=!0,w.bonds=[],w.bondOrder=[],w.properties={},i[i.length-1].push(w)}if(o)for(let t=0;t<i.length;t++)(0,s.assignBonds)(i[t],e);return i}},797:(t,e,i)=>{"use strict";i.r(e),i.d(e,{VASP:()=>n});var r=i(638),s=i(392);function n(t,e={}){var i=[[]],n={};const a=void 0===e.assignBonds||e.assignBonds;var o=t.replace(/^\s+/,"").split(/\r?\n/);if(o.length<3)return i;if(!o[1].match(/\d+/))return console.log("Warning: second line of the vasp structure file must be a number"),i;if(n.length=parseFloat(o[1]),n.length<0)return console.log("Warning: Vasp implementation for negative lattice lengths is not yet available"),i;n.xVec=new Float32Array(o[2].replace(/^\s+/,"").split(/\s+/)),n.yVec=new Float32Array(o[3].replace(/^\s+/,"").split(/\s+/)),n.zVec=new Float32Array(o[4].replace(/^\s+/,"").split(/\s+/));var l=new r.Matrix3(n.xVec[0],n.xVec[1],n.xVec[2],n.yVec[0],n.yVec[1],n.yVec[2],n.zVec[0],n.zVec[1],n.zVec[2]);l.multiplyScalar(n.length),i.modelData=[{symmetries:[],cryst:{matrix:l}}];var h=o[5].trim().split(/\s+/),c=new Int16Array(o[6].trim().split(/\s+/)),d=o[7].trim(),u=!1;if(d.match(/S/)&&(u=!0,d=o[8].trim()),"c"==d.toLowerCase()[0])d="cartesian";else{if("d"!=d.toLowerCase()[0])return console.log("Warning: Unknown vasp mode in POSCAR file: mode must be either C(artesian) or D(irect)"),i;d="direct"}if(h.length!=c.length)return console.log("Warning: declaration of atomary species wrong:"),console.log(h),console.log(c),i;u?o.splice(0,9):o.splice(0,8);for(var f=0,p=0,g=h.length;p<g;p++){for(var m=h[p],v=0,_=c[p];v<_;v++){var y=new Float32Array(o[f+v].trim().split(/\s+/)),b={};b.elem=m,"cartesian"==d?(b.x=n.length*y[0],b.y=n.length*y[1],b.z=n.length*y[2]):(b.x=n.length*(y[0]*n.xVec[0]+y[1]*n.yVec[0]+y[2]*n.zVec[0]),b.y=n.length*(y[0]*n.xVec[1]+y[1]*n.yVec[1]+y[2]*n.zVec[1]),b.z=n.length*(y[0]*n.xVec[2]+y[1]*n.yVec[2]+y[2]*n.zVec[2])),b.bonds=[],b.bondOrder=[],i[0].push(b)}f+=c[p]}if(a)for(let t=0;t<i.length;t++)(0,s.assignBonds)(i[t],e);return i}},408:(t,e,i)=>{"use strict";i.r(e),i.d(e,{areConnected:()=>n});var r=i(40);const s=new Set(["Na","K","Ca","Mg","Mn","Sr"]);function n(t,e,i){if(i&&i.unboundCations&&(s.has(t.elem)||s.has(e.elem)))return!1;let n=(0,r.bondLength)(t.elem)+(0,r.bondLength)(e.elem);n+=.25,n*=n;let a=t.x-e.x;if(a*=a,a>n)return!1;let o=t.y-e.y;if(o*=o,o>n)return!1;let l=t.z-e.z;if(l*=l,l>n)return!1;const h=a+o+l;return!(isNaN(h)||h<.5||h>n||t.altLoc!==e.altLoc&&""!==t.altLoc.trim()&&""!==e.altLoc.trim())}},392:(t,e,i)=>{"use strict";i.r(e),i.d(e,{assignBonds:()=>a});var r=i(408);const s=[{x:0,y:0,z:1},{x:0,y:1,z:-1},{x:0,y:1,z:0},{x:0,y:1,z:1},{x:1,y:-1,z:-1},{x:1,y:-1,z:0},{x:1,y:-1,z:1},{x:1,y:0,z:-1},{x:1,y:0,z:0},{x:1,y:0,z:1},{x:1,y:1,z:-1},{x:1,y:1,z:0},{x:1,y:1,z:1}],n=4.95;function a(t,e){for(let e=0,i=t.length;e<i;e++)t[e].index||(t[e].index=e);const i={x:{y:{z:[]}}};for(let e=0;e<t.length;e++){const r=t[e],s=Math.floor(r.x/n),a=Math.floor(r.y/n),o=Math.floor(r.z/n);i[s]||(i[s]={}),i[s][a]||(i[s][a]={}),i[s][a][o]||(i[s][a][o]=[]),i[s][a][o].push(r)}function a(t,i){for(let s=0;s<t.length;s++){const n=t[s];for(let t=0;t<i.length;t++){const s=i[t];if((0,r.areConnected)(n,s,e)){const t=n.bonds.indexOf(s.index),e=s.bonds.indexOf(n.index);-1===t&&-1===e?(n.bonds.push(s.index),n.bondOrder.push(1),s.bonds.push(n.index),s.bondOrder.push(1)):-1===t?(n.bonds.push(s.index),n.bondOrder.push(s.bondOrder[e])):-1===e&&(s.bonds.push(n.index),s.bondOrder.push(n.bondOrder[t]))}}}}for(let t in i){const n=parseInt(t);for(let t in i[n]){const o=parseInt(t);for(let t in i[n][o]){const l=parseInt(t),h=i[n][o][l];for(let t=0;t<h.length;t++){const i=h[t];for(let s=t+1;s<h.length;s++){const t=h[s];(0,r.areConnected)(i,t,e)&&-1==i.bonds.indexOf(t.index)&&(i.bonds.push(t.index),i.bondOrder.push(1),t.bonds.push(i.index),t.bondOrder.push(1))}}for(let t=0;t<s.length;t++){const e=s[t];if(!i[n+e.x]||!i[n+e.x][o+e.y]||!i[n+e.x][o+e.y][l+e.z])continue;a(h,i[n+e.x][o+e.y][l+e.z])}}}}}},40:(t,e,i)=>{"use strict";i.r(e),i.d(e,{bondLength:()=>s,bondTable:()=>r,setBondLength:()=>n});let r={H:.37,He:.32,Li:1.34,Be:.9,B:.82,C:.77,N:.75,O:.73,F:.71,Ne:.69,Na:1.54,Mg:1.3,Al:1.18,Si:1.11,P:1.06,S:1.02,Cl:.99,Ar:.97,K:1.96,Ca:1.74,Sc:1.44,Ti:1.56,V:1.25,Mn:1.39,Fe:1.25,Co:1.26,Ni:1.21,Cu:1.38,Zn:1.31,Ga:1.26,Ge:1.22,Se:1.16,Br:1.14,Kr:1.1,Rb:2.11,Sr:1.92,Y:1.62,Zr:1.48,Nb:1.37,Mo:1.45,Tc:1.56,Ru:1.26,Rh:1.35,Pd:1.31,Ag:1.53,Cd:1.48,In:1.44,Sn:1.41,Sb:1.38,Te:1.35,I:1.33,Xe:1.3,Cs:2.25,Ba:1.98,Lu:1.6,Hf:1.5,Ta:1.38,W:1.46,Re:1.59,Os:1.44,Ir:1.37,Pt:1.28,Au:1.44,Hg:1.49,Tl:1.48,Pb:1.47,Bi:1.46,Rn:1.45};function s(t){return r[t]||1.6}function n(t,e){e<0&&(e=0),r[t]=e}},864:(__unused_webpack_module,__webpack_exports__,__webpack_require__)=>{"use strict";__webpack_require__.r(__webpack_exports__),__webpack_require__.d(__webpack_exports__,{PausableTimer:()=>PausableTimer,adjustVolumeStyle:()=>adjustVolumeStyle,base64ToArray:()=>base64ToArray,deepCopy:()=>deepCopy,download:()=>download,extend:()=>extend,get:()=>get,getAtomProperty:()=>getAtomProperty,getColorFromStyle:()=>getColorFromStyle,getElement:()=>getElement,getExtent:()=>getExtent,getPropertyRange:()=>getPropertyRange,getbin:()=>getbin,inflateString:()=>inflateString,isEmptyObject:()=>isEmptyObject,isNumeric:()=>isNumeric,makeFunction:()=>makeFunction,mergeGeos:()=>mergeGeos,specStringToObject:()=>specStringToObject});var _Gradient__WEBPACK_IMPORTED_MODULE_0__=__webpack_require__(546),_VolumeData__WEBPACK_IMPORTED_MODULE_1__=__webpack_require__(848),_colors__WEBPACK_IMPORTED_MODULE_2__=__webpack_require__(222),pako__WEBPACK_IMPORTED_MODULE_3__=__webpack_require__(75);function extend(t,e){for(var i in e)e.hasOwnProperty(i)&&void 0!==e[i]&&(t[i]=e[i]);return t}function deepCopy(t){let e,i,r;if(null==t)return{};if("object"!=typeof t||null===t)return t;for(r in e=Array.isArray(t)?[]:{},t)i=t[r],e[r]=deepCopy(i);return e}function isNumeric(t){var e=typeof t;return("number"===e||"string"===e)&&!isNaN(t-parseFloat(t))}function isEmptyObject(t){var e;for(e in t)return!1;return!0}function makeFunction(callback){return callback&&"string"==typeof callback&&(callback=eval("("+callback+")")),callback&&"function"!=typeof callback?(console.warn("Invalid callback provided."),()=>{}):callback}function adjustVolumeStyle(t){t&&(!t.volformat||t.voldata instanceof _VolumeData__WEBPACK_IMPORTED_MODULE_1__.VolumeData||(t.voldata=new _VolumeData__WEBPACK_IMPORTED_MODULE_1__.VolumeData(t.voldata,t.volformat)),t.volscheme&&(t.volscheme=_Gradient__WEBPACK_IMPORTED_MODULE_0__.Gradient.getGradient(t.volscheme)))}function getExtent(t,e){var i,r,s,n,a,o,l,h,c,d,u=!e;if(i=r=s=9999,n=a=o=-9999,l=h=c=d=0,0===t.length)return[[0,0,0],[0,0,0],[0,0,0]];for(var f=0;f<t.length;f++){var p=t[f];if(void 0!==p&&isFinite(p.x)&&isFinite(p.y)&&isFinite(p.z)&&(d++,l+=p.x,h+=p.y,c+=p.z,i=i<p.x?i:p.x,r=r<p.y?r:p.y,s=s<p.z?s:p.z,n=n>p.x?n:p.x,a=a>p.y?a:p.y,o=o>p.z?o:p.z,p.symmetries&&u))for(var g=0;g<p.symmetries.length;g++)d++,l+=p.symmetries[g].x,h+=p.symmetries[g].y,c+=p.symmetries[g].z,i=i<p.symmetries[g].x?i:p.symmetries[g].x,r=r<p.symmetries[g].y?r:p.symmetries[g].y,s=s<p.symmetries[g].z?s:p.symmetries[g].z,n=n>p.symmetries[g].x?n:p.symmetries[g].x,a=a>p.symmetries[g].y?a:p.symmetries[g].y,o=o>p.symmetries[g].z?o:p.symmetries[g].z}return[[i,r,s],[n,a,o],[l/d,h/d,c/d]]}function getPropertyRange(t,e){for(var i=Number.POSITIVE_INFINITY,r=Number.NEGATIVE_INFINITY,s=0,n=t.length;s<n;s++){var a=getAtomProperty(t[s],e);null!=a&&(a<i&&(i=a),a>r&&(r=a))}return isFinite(i)||isFinite(r)?isFinite(i)?isFinite(r)||(r=i):i=r:i=r=0,[i,r]}class PausableTimer{constructor(t,e,i){this.total_time_run=0,this.fn=t,this.arg=i,this.countdown=e,this.start_time=(new Date).getTime(),this.ident=setTimeout(t,e,i)}cancel(){clearTimeout(this.ident)}pause(){clearTimeout(this.ident),this.total_time_run=(new Date).getTime()-this.start_time}resume(){this.ident=setTimeout(this.fn,Math.max(0,this.countdown-this.total_time_run),this.arg)}}function base64ToArray(t){for(var e=window.atob(t),i=e.length,r=new Uint8Array(i),s=0;s<i;s++)r[s]=e.charCodeAt(s);return r}function getAtomProperty(t,e){var i=null;return t.properties&&void 0!==t.properties[e]?i=t.properties[e]:void 0!==t[e]&&(i=t[e]),i}function mergeGeos(t,e){var i=e.geometry;void 0!==i&&t.geometryGroups.push(i.geometryGroups[0])}function specStringToObject(t){if("object"==typeof t)return t;if(void 0===t||null==t)return t;try{return JSON.parse(t)}catch(t){}var e=function(t){return isNumeric(t)?Math.floor(parseFloat(t))==parseInt(t)||t.indexOf(".")>=0?parseFloat(t):parseInt(t):"true"===t||"false"!==t&&t},i={};if("all"===(t=t.replace(/%7E/g,"~")))return i;for(var r=t.split(";"),s=0;s<r.length;s++){var n=r[s].split(":"),a=n[0],o={},l=n[1];if(l)if(-1!==(l=l.replace(/~/g,"=")).indexOf("="))for(var h=l.split(","),c=0;c<h.length;c++){var d=h[c].split("=",2);o[d[0]]=e(d[1])}else o=-1!==l.indexOf(",")?l.split(","):e(l);i[a]=o}return i}function checkStatus(t){if(!t.ok)throw new Error(`HTTP ${t.status} - ${t.statusText}`);return t}function get(t,e){var i=fetch(t).then(checkStatus).then((t=>t.text()));return e?i.then(e):i}function getbin(t,e,i,r){var s;return s="POST"==i?fetch(t,{method:"POST",body:r}).then((t=>checkStatus(t))).then((t=>t.arrayBuffer())):fetch(t).then((t=>checkStatus(t))).then((t=>t.arrayBuffer())),e?s.then(e):s}function download(t,e,i,r){var s="",n="",a="",o=null,l=e.addModel();if(t.indexOf(":")<0&&(t=4==t.length?"pdb:"+t:isNaN(t)?"url:"+t:"cid:"+t),"mmtf:"==t.substring(0,5)&&(console.warn("WARNING: MMTF now deprecated.  Reverting to bcif."),t="bcif:"+t.slice(5)),"bcif:"===t.substring(0,5))t=t.substring(5).toUpperCase(),a="https://models.rcsb.org/"+t+".bcif.gz",i&&void 0===i.noComputeSecondaryStructure&&(i.noComputeSecondaryStructure=!0),o=new Promise((function(t){getbin(a).then((function(r){l.addMolData(r,"bcif.gz",i),e.zoomTo(),e.render(),t(l)}),(function(){console.error("fetch of "+a+" failed.")}))}));else{if("pdb:"===t.substring(0,4)){if(s="bcif",i&&i.format&&(s=i.format),i&&void 0===i.noComputeSecondaryStructure&&(i.noComputeSecondaryStructure=!0),!(t=t.substring(4).toUpperCase()).match(/^[1-9][A-Za-z0-9]{3}$/))return void alert("Wrong PDB ID");"bcif"==s?a="https://models.rcsb.org/"+t.toUpperCase()+".bcif.gz":(n=i&&i.pdbUri?i.pdbUri:"https://files.rcsb.org/view/",a=n+t+"."+s)}else if("cid:"==t.substring(0,4)){if(s="sdf",!(t=t.substring(4)).match(/^[0-9]+$/))return void alert("Wrong Compound ID");a="https://pubchem.ncbi.nlm.nih.gov/rest/pug/compound/cid/"+t+"/SDF?record_type=3d"}else"url:"==t.substring(0,4)&&(a=t.substring(4),s=a);var h=function(t){l.addMolData(t,s,i),e.zoomTo(),e.render()};o=new Promise((function(e){"bcif"==s?getbin(a).then((function(t){h(t),e(l)})).catch((function(){n=i&&i.pdbUri?i.pdbUri:"https://files.rcsb.org/view/",a=n+t+".pdb",s="pdb",console.warn("falling back to pdb format"),get(a).then((function(t){h(t),e(l)})).catch((function(t){h(""),e(l),console.error("fetch of "+a+" failed: "+t.statusText)}))})):get(a).then((function(t){h(t),e(l)})).catch((function(t){h(""),e(l),console.error("fetch of "+a+" failed: "+t.statusText)}))}))}return r?(o.then((function(t){r(t)})),l):o}function getColorFromStyle(t,e){let i=e.colorscheme;if(void 0!==_colors__WEBPACK_IMPORTED_MODULE_2__.builtinColorSchemes[i])i=_colors__WEBPACK_IMPORTED_MODULE_2__.builtinColorSchemes[i];else if("string"==typeof i&&i.endsWith("Carbon")){let t=i.substring(0,i.lastIndexOf("Carbon")).toLowerCase();if(void 0!==_colors__WEBPACK_IMPORTED_MODULE_2__.htmlColors[t]){let e=Object.assign({},_colors__WEBPACK_IMPORTED_MODULE_2__.elementColors.defaultColors);e.C=_colors__WEBPACK_IMPORTED_MODULE_2__.htmlColors[t],_colors__WEBPACK_IMPORTED_MODULE_2__.builtinColorSchemes[i]={prop:"elem",map:e},i=_colors__WEBPACK_IMPORTED_MODULE_2__.builtinColorSchemes[i]}}let r=t.color;if(void 0!==e.color&&"spectrum"!=e.color&&(r=e.color),void 0!==i){let n,a;if(void 0!==_colors__WEBPACK_IMPORTED_MODULE_2__.elementColors[i])i=_colors__WEBPACK_IMPORTED_MODULE_2__.elementColors[i],void 0!==i[t[i.prop]]&&(r=i.map[t[i.prop]]);else if(void 0!==i[t[i.prop]])r=i.map[t[i.prop]];else if(void 0!==i.prop&&void 0!==i.gradient){n=i.prop;var s=i.gradient;s instanceof _Gradient__WEBPACK_IMPORTED_MODULE_0__.GradientType||(s=(0,_Gradient__WEBPACK_IMPORTED_MODULE_0__.getGradient)(i));let e=s.range()||[-1,1];a=getAtomProperty(t,n),null!=a&&(r=s.valueToHex(a,e))}else void 0!==i.prop&&void 0!==i.map?(n=i.prop,a=getAtomProperty(t,n),void 0!==i.map[a]&&(r=i.map[a])):void 0!==e.colorscheme[t.elem]?r=e.colorscheme[t.elem]:console.warn("Could not interpret colorscheme "+i)}else void 0!==e.colorfunc&&(r=e.colorfunc(t));return _colors__WEBPACK_IMPORTED_MODULE_2__.CC.color(r)}function getElement(t){let e=t;return"string"==typeof t?e=document.querySelector("#"+t):"object"==typeof t&&t.get&&(e=t.get(0)),e}function inflateString(t,e=!0){let i;if("string"==typeof t){i=(new TextEncoder).encode(t)}else i=new Uint8Array(t);return(0,pako__WEBPACK_IMPORTED_MODULE_3__.inflate)(i,{to:e?"string":null})}},111:(t,e,i)=>{var r;r={},t.exports=r,function(t,e){t.toRGBA8=function(e){var i=e.width,r=e.height;if(null==e.tabs.acTL)return[t.toRGBA8.decodeImage(e.data,i,r,e).buffer];var s=[];null==e.frames[0].data&&(e.frames[0].data=e.data);for(var n,a=new Uint8Array(i*r*4),o=0;o<e.frames.length;o++){var l=e.frames[o],h=l.rect.x,c=l.rect.y,d=l.rect.width,u=l.rect.height,f=t.toRGBA8.decodeImage(l.data,d,u,e);if(0==o?n=f:0==l.blend?t._copyTile(f,d,u,n,i,r,h,c,0):1==l.blend&&t._copyTile(f,d,u,n,i,r,h,c,1),s.push(n.buffer),n=n.slice(0),0==l.dispose);else if(1==l.dispose)t._copyTile(a,d,u,n,i,r,h,c,0);else if(2==l.dispose){for(var p=o-1;2==e.frames[p].dispose;)p--;n=new Uint8Array(s[p]).slice(0)}}return s},t.toRGBA8.decodeImage=function(e,i,r,s){var n=i*r,a=t.decode._getBPP(s),o=Math.ceil(i*a/8),l=new Uint8Array(4*n),h=new Uint32Array(l.buffer),c=s.ctype,d=s.depth,u=t._bin.readUshort;if(6==c){var f=n<<2;if(8==d)for(var p=0;p<f;p++)l[p]=e[p];if(16==d)for(p=0;p<f;p++)l[p]=e[p<<1]}else if(2==c){var g=s.tabs.tRNS,m=-1,v=-1,_=-1;if(g&&(m=g[0],v=g[1],_=g[2]),8==d)for(p=0;p<n;p++){var y=3*p;l[M=p<<2]=e[y],l[M+1]=e[y+1],l[M+2]=e[y+2],l[M+3]=255,-1!=m&&e[y]==m&&e[y+1]==v&&e[y+2]==_&&(l[M+3]=0)}if(16==d)for(p=0;p<n;p++)y=6*p,l[M=p<<2]=e[y],l[M+1]=e[y+2],l[M+2]=e[y+4],l[M+3]=255,-1!=m&&u(e,y)==m&&u(e,y+2)==v&&u(e,y+4)==_&&(l[M+3]=0)}else if(3==c){var b=s.tabs.PLTE,x=s.tabs.tRNS,w=x?x.length:0;if(1==d)for(var A=0;A<r;A++){var C=A*o,S=A*i;for(p=0;p<i;p++){var M=S+p<<2,z=3*(T=e[C+(p>>3)]>>7-(7&p)&1);l[M]=b[z],l[M+1]=b[z+1],l[M+2]=b[z+2],l[M+3]=T<w?x[T]:255}}if(2==d)for(A=0;A<r;A++)for(C=A*o,S=A*i,p=0;p<i;p++)M=S+p<<2,z=3*(T=e[C+(p>>2)]>>6-((3&p)<<1)&3),l[M]=b[z],l[M+1]=b[z+1],l[M+2]=b[z+2],l[M+3]=T<w?x[T]:255;if(4==d)for(A=0;A<r;A++)for(C=A*o,S=A*i,p=0;p<i;p++)M=S+p<<2,z=3*(T=e[C+(p>>1)]>>4-((1&p)<<2)&15),l[M]=b[z],l[M+1]=b[z+1],l[M+2]=b[z+2],l[M+3]=T<w?x[T]:255;if(8==d)for(p=0;p<n;p++){var T;M=p<<2,z=3*(T=e[p]),l[M]=b[z],l[M+1]=b[z+1],l[M+2]=b[z+2],l[M+3]=T<w?x[T]:255}}else if(4==c){if(8==d)for(p=0;p<n;p++){M=p<<2;var E=e[L=p<<1];l[M]=E,l[M+1]=E,l[M+2]=E,l[M+3]=e[L+1]}if(16==d)for(p=0;p<n;p++){var L;M=p<<2,E=e[L=p<<2],l[M]=E,l[M+1]=E,l[M+2]=E,l[M+3]=e[L+2]}}else if(0==c){if(m=s.tabs.tRNS?s.tabs.tRNS:-1,1==d)for(p=0;p<n;p++){var F=(E=255*(e[p>>3]>>7-(7&p)&1))==255*m?0:255;h[p]=F<<24|E<<16|E<<8|E}if(2==d)for(p=0;p<n;p++)F=(E=85*(e[p>>2]>>6-((3&p)<<1)&3))==85*m?0:255,h[p]=F<<24|E<<16|E<<8|E;if(4==d)for(p=0;p<n;p++)F=(E=17*(e[p>>1]>>4-((1&p)<<2)&15))==17*m?0:255,h[p]=F<<24|E<<16|E<<8|E;if(8==d)for(p=0;p<n;p++)F=(E=e[p])==m?0:255,h[p]=F<<24|E<<16|E<<8|E;if(16==d)for(p=0;p<n;p++)E=e[p<<1],F=u(e,p<<1)==m?0:255,h[p]=F<<24|E<<16|E<<8|E}return l},t.decode=function(e){for(var i,r=new Uint8Array(e),s=8,n=t._bin,a=n.readUshort,o=n.readUint,l={tabs:{},frames:[]},h=new Uint8Array(r.length),c=0,d=0,u=[137,80,78,71,13,10,26,10],f=0;f<8;f++)if(r[f]!=u[f])throw"The input is not a PNG file!";for(;s<r.length;){var p=n.readUint(r,s);s+=4;var g=n.readASCII(r,s,4);if(s+=4,"IHDR"==g)t.decode._IHDR(r,s,l);else if("IDAT"==g){for(f=0;f<p;f++)h[c+f]=r[s+f];c+=p}else if("acTL"==g)l.tabs[g]={num_frames:o(r,s),num_plays:o(r,s+4)},i=new Uint8Array(r.length);else if("fcTL"==g){0!=d&&((C=l.frames[l.frames.length-1]).data=t.decode._decompress(l,i.slice(0,d),C.rect.width,C.rect.height),d=0);var m={x:o(r,s+12),y:o(r,s+16),width:o(r,s+4),height:o(r,s+8)},v=a(r,s+22);v=a(r,s+20)/(0==v?100:v);var _={rect:m,delay:Math.round(1e3*v),dispose:r[s+24],blend:r[s+25]};l.frames.push(_)}else if("fdAT"==g){for(f=0;f<p-4;f++)i[d+f]=r[s+f+4];d+=p-4}else if("pHYs"==g)l.tabs[g]=[n.readUint(r,s),n.readUint(r,s+4),r[s+8]];else if("cHRM"==g)for(l.tabs[g]=[],f=0;f<8;f++)l.tabs[g].push(n.readUint(r,s+4*f));else if("tEXt"==g){null==l.tabs[g]&&(l.tabs[g]={});var y=n.nextZero(r,s),b=n.readASCII(r,s,y-s),x=n.readASCII(r,y+1,s+p-y-1);l.tabs[g][b]=x}else if("iTXt"==g){null==l.tabs[g]&&(l.tabs[g]={}),y=0;var w=s;y=n.nextZero(r,w),b=n.readASCII(r,w,y-w),r[w=y+1],r[w+1],w+=2,y=n.nextZero(r,w),n.readASCII(r,w,y-w),w=y+1,y=n.nextZero(r,w),n.readUTF8(r,w,y-w),w=y+1,x=n.readUTF8(r,w,p-(w-s)),l.tabs[g][b]=x}else if("PLTE"==g)l.tabs[g]=n.readBytes(r,s,p);else if("hIST"==g){var A=l.tabs.PLTE.length/3;for(l.tabs[g]=[],f=0;f<A;f++)l.tabs[g].push(a(r,s+2*f))}else if("tRNS"==g)3==l.ctype?l.tabs[g]=n.readBytes(r,s,p):0==l.ctype?l.tabs[g]=a(r,s):2==l.ctype&&(l.tabs[g]=[a(r,s),a(r,s+2),a(r,s+4)]);else if("gAMA"==g)l.tabs[g]=n.readUint(r,s)/1e5;else if("sRGB"==g)l.tabs[g]=r[s];else if("bKGD"==g)0==l.ctype||4==l.ctype?l.tabs[g]=[a(r,s)]:2==l.ctype||6==l.ctype?l.tabs[g]=[a(r,s),a(r,s+2),a(r,s+4)]:3==l.ctype&&(l.tabs[g]=r[s]);else if("IEND"==g){var C;0!=d&&((C=l.frames[l.frames.length-1]).data=t.decode._decompress(l,i.slice(0,d),C.rect.width,C.rect.height),d=0),l.data=t.decode._decompress(l,h,l.width,l.height);break}s+=p,n.readUint(r,s),s+=4}return delete l.compress,delete l.interlace,delete l.filter,l},t.decode._decompress=function(e,i,r,s){return 0==e.compress&&(i=t.decode._inflate(i)),0==e.interlace?i=t.decode._filterZero(i,e,0,r,s):1==e.interlace&&(i=t.decode._readInterlace(i,e)),i},t.decode._inflate=function(t){return e.inflate(t)},t.decode._readInterlace=function(e,i){for(var r=i.width,s=i.height,n=t.decode._getBPP(i),a=n>>3,o=Math.ceil(r*n/8),l=new Uint8Array(s*o),h=0,c=[0,0,4,0,2,0,1],d=[0,4,0,2,0,1,0],u=[8,8,8,4,4,2,2],f=[8,8,4,4,2,2,1],p=0;p<7;){for(var g=u[p],m=f[p],v=0,_=0,y=c[p];y<s;)y+=g,_++;for(var b=d[p];b<r;)b+=m,v++;var x=Math.ceil(v*n/8);t.decode._filterZero(e,i,h,v,_);for(var w=0,A=c[p];A<s;){for(var C=d[p],S=h+w*x<<3;C<r;){var M;if(1==n&&(M=(M=e[S>>3])>>7-(7&S)&1,l[A*o+(C>>3)]|=M<<7-(3&C)),2==n&&(M=(M=e[S>>3])>>6-(7&S)&3,l[A*o+(C>>2)]|=M<<6-((3&C)<<1)),4==n&&(M=(M=e[S>>3])>>4-(7&S)&15,l[A*o+(C>>1)]|=M<<4-((1&C)<<2)),n>=8)for(var z=A*o+C*a,T=0;T<a;T++)l[z+T]=e[(S>>3)+T];S+=n,C+=m}w++,A+=g}v*_!=0&&(h+=_*(1+x)),p+=1}return l},t.decode._getBPP=function(t){return[1,null,3,1,2,null,4][t.ctype]*t.depth},t.decode._filterZero=function(e,i,r,s,n){var a=t.decode._getBPP(i),o=Math.ceil(s*a/8),l=t.decode._paeth;a=Math.ceil(a/8);for(var h=0;h<n;h++){var c=r+h*o,d=c+h+1,u=e[d-1];if(0==u)for(var f=0;f<o;f++)e[c+f]=e[d+f];else if(1==u){for(f=0;f<a;f++)e[c+f]=e[d+f];for(f=a;f<o;f++)e[c+f]=e[d+f]+e[c+f-a]&255}else if(0==h){for(f=0;f<a;f++)e[c+f]=e[d+f];if(2==u)for(f=a;f<o;f++)e[c+f]=255&e[d+f];if(3==u)for(f=a;f<o;f++)e[c+f]=e[d+f]+(e[c+f-a]>>1)&255;if(4==u)for(f=a;f<o;f++)e[c+f]=e[d+f]+l(e[c+f-a],0,0)&255}else{if(2==u)for(f=0;f<o;f++)e[c+f]=e[d+f]+e[c+f-o]&255;if(3==u){for(f=0;f<a;f++)e[c+f]=e[d+f]+(e[c+f-o]>>1)&255;for(f=a;f<o;f++)e[c+f]=e[d+f]+(e[c+f-o]+e[c+f-a]>>1)&255}if(4==u){for(f=0;f<a;f++)e[c+f]=e[d+f]+l(0,e[c+f-o],0)&255;for(f=a;f<o;f++)e[c+f]=e[d+f]+l(e[c+f-a],e[c+f-o],e[c+f-a-o])&255}}}return e},t.decode._paeth=function(t,e,i){var r=t+e-i,s=Math.abs(r-t),n=Math.abs(r-e),a=Math.abs(r-i);return s<=n&&s<=a?t:n<=a?e:i},t.decode._IHDR=function(e,i,r){var s=t._bin;r.width=s.readUint(e,i),i+=4,r.height=s.readUint(e,i),i+=4,r.depth=e[i],i++,r.ctype=e[i],i++,r.compress=e[i],i++,r.filter=e[i],i++,r.interlace=e[i],i++},t._bin={nextZero:function(t,e){for(;0!=t[e];)e++;return e},readUshort:function(t,e){return t[e]<<8|t[e+1]},writeUshort:function(t,e,i){t[e]=i>>8&255,t[e+1]=255&i},readUint:function(t,e){return 16777216*t[e]+(t[e+1]<<16|t[e+2]<<8|t[e+3])},writeUint:function(t,e,i){t[e]=i>>24&255,t[e+1]=i>>16&255,t[e+2]=i>>8&255,t[e+3]=255&i},readASCII:function(t,e,i){for(var r="",s=0;s<i;s++)r+=String.fromCharCode(t[e+s]);return r},writeASCII:function(t,e,i){for(var r=0;r<i.length;r++)t[e+r]=i.charCodeAt(r)},readBytes:function(t,e,i){for(var r=[],s=0;s<i;s++)r.push(t[e+s]);return r},pad:function(t){return t.length<2?"0"+t:t},readUTF8:function(e,i,r){for(var s,n="",a=0;a<r;a++)n+="%"+t._bin.pad(e[i+a].toString(16));try{s=decodeURIComponent(n)}catch(s){return t._bin.readASCII(e,i,r)}return s}},t._copyTile=function(t,e,i,r,s,n,a,o,l){for(var h=Math.min(e,s),c=Math.min(i,n),d=0,u=0,f=0;f<c;f++)for(var p=0;p<h;p++)if(a>=0&&o>=0?(d=f*e+p<<2,u=(o+f)*s+a+p<<2):(d=(-o+f)*e-a+p<<2,u=f*s+p<<2),0==l)r[u]=t[d],r[u+1]=t[d+1],r[u+2]=t[d+2],r[u+3]=t[d+3];else if(1==l){var g=t[d+3]*(1/255),m=t[d]*g,v=t[d+1]*g,_=t[d+2]*g,y=r[u+3]*(1/255),b=r[u]*y,x=r[u+1]*y,w=r[u+2]*y,A=1-g,C=g+y*A,S=0==C?0:1/C;r[u+3]=255*C,r[u+0]=(m+b*A)*S,r[u+1]=(v+x*A)*S,r[u+2]=(_+w*A)*S}else if(2==l)g=t[d+3],m=t[d],v=t[d+1],_=t[d+2],y=r[u+3],b=r[u],x=r[u+1],w=r[u+2],g==y&&m==b&&v==x&&_==w?(r[u]=0,r[u+1]=0,r[u+2]=0,r[u+3]=0):(r[u]=m,r[u+1]=v,r[u+2]=_,r[u+3]=g);else if(3==l){if(g=t[d+3],m=t[d],v=t[d+1],_=t[d+2],y=r[u+3],b=r[u],x=r[u+1],w=r[u+2],g==y&&m==b&&v==x&&_==w)continue;if(g<220&&y>20)return!1}return!0},t.encode=function(e,i,r,s,n,a){null==s&&(s=0),null==a&&(a=!1);for(var o=new Uint8Array(e[0].byteLength*e.length+100),l=[137,80,78,71,13,10,26,10],h=0;h<8;h++)o[h]=l[h];var c=8,d=t._bin,u=t.crc.crc,f=d.writeUint,p=d.writeUshort,g=d.writeASCII,m=t.encode.compressPNG(e,i,r,s,a);f(o,c,13),g(o,c+=4,"IHDR"),f(o,c+=4,i),f(o,c+=4,r),o[c+=4]=m.depth,o[++c]=m.ctype,o[++c]=0,o[++c]=0,o[++c]=0,f(o,++c,u(o,c-17,17)),f(o,c+=4,1),g(o,c+=4,"sRGB"),o[c+=4]=1,f(o,++c,u(o,c-5,5)),c+=4;var v=e.length>1;if(v&&(f(o,c,8),g(o,c+=4,"acTL"),f(o,c+=4,e.length),f(o,c+=4,0),f(o,c+=4,u(o,c-12,12)),c+=4),3==m.ctype){for(f(o,c,3*(M=m.plte.length)),g(o,c+=4,"PLTE"),c+=4,h=0;h<M;h++){var
```
