## Additional file 1. Evaluated software source and dependencies for "Helix Design Studio: designing antimicrobial peptide candidates for membrane environments": app.js

```
const $ = (sel, root = document) => root.querySelector(sel);
const $$ = (sel, root = document) => [...root.querySelectorAll(sel)];
const ANON_STORAGE_KEY = 'helix-design-anonymous-id';
let usageLoggingEnabled = false;
const STANDARD_RESIDUE_ORDER = ['A', 'R', 'N', 'D', 'C', 'Q', 'E', 'G', 'H', 'I', 'L', 'K', 'M', 'F', 'P', 'S', 'T', 'W', 'Y', 'V'];
const HYDRO_SCALE = {
  D: -0.77, E: -0.64, K: -0.99, R: -1.01, H: 0.13,
  G: 0.00, A: 0.31, V: 1.22, L: 1.70, I: 1.80,
  P: 0.72, M: 1.23, F: 1.79, W: 2.25, Y: 0.96,
  T: 0.26, S: -0.04, C: 1.54, N: -0.60, Q: -0.22,
};
const REFERENCE_PEPTIDES = {
  eaaak3: {
    name: '(EAAAK)3',
    category: 'Water-stable designed helices',
    sequence: 'EAAAKEAAAKEAAAK',
    family: 'de novo Glu-Lys stabilized helix motif',
    profile: {
      water: 'strong, canonical helix in water',
      apolar: 'still helical, but not a membrane-specific sequence',
      anionic: 'moderately amphipathic; not a canonical AMP',
    },
    note: 'One of the cleanest aqueous alpha-helix benchmarks. The repeating Glu-Lys pattern stabilizes helical structure through intrahelical i,i+4 electrostatic pairing.',
  },
  aakaa3: {
    name: '(AAKAA)3',
    category: 'Water-stable designed helices',
    sequence: 'AAKAAAAKAAAAKAA',
    family: 'alanine-rich model helix',
    profile: {
      water: 'strong alanine-driven helix',
      apolar: 'very helical, but lacks membrane-targeting patterning',
      anionic: 'weak membrane selectivity; mainly a helicity control',
    },
    note: 'Alanine-rich benchmark that emphasizes intrinsic helical propensity in water more than membrane amphipathicity.',
  },
  ll37: {
    name: 'LL-37',
    category: 'Membrane-active amphipathic helices',
    sequence: 'LLGDFFRKSKEKIGKEFKRIVQRIKDFLRNLVPRTES',
    family: 'human cathelicidin host-defense peptide',
    profile: {
      water: 'dynamic / partly helical',
      apolar: 'membrane-mimetic helix, but not a hydrophobic transmembrane peptide',
      anionic: 'strong surface-bound amphipathic helix',
    },
    note: 'Canonical human AMP that becomes substantially more helical in membrane-mimetic and anionic environments than in bulk water.',
  },
  melittin: {
    name: 'Melittin',
    category: 'Membrane-active amphipathic helices',
    sequence: 'GIGAVLKVLTTGLPALISWIKRKRQQ',
    family: 'bee-venom membrane-active peptide',
    profile: {
      water: 'flexible / partly structured',
      apolar: 'strong amphipathic helix',
      anionic: 'strong membrane-active helix',
    },
    note: 'Classic amphipathic helix benchmark: highly membrane-active and strongly helical in membrane-like or apolar media.',
  },
  magainin2: {
    name: 'Magainin 2',
    category: 'Membrane-active amphipathic helices',
    sequence: 'GIGKFLHSAKKFGKAFVGEIMNS',
    family: 'frog antimicrobial peptide',
    profile: {
      water: 'mostly disordered',
      apolar: 'helix in membrane mimics',
      anionic: 'surface-bound amphipathic helix',
    },
    note: 'Well-known AMP that is weakly structured in water and becomes helical upon binding to negatively charged membrane mimics.',
  },
  cecropin_a: {
    name: 'Cecropin A',
    category: 'Membrane-active amphipathic helices',
    sequence: 'KWKLFKKIEKVGQNIRDGIIKAGPAVAVVGQATQIAK',
    family: 'insect antimicrobial peptide',
    profile: {
      water: 'limited helix / flexible',
      apolar: 'helix in membrane-mimetic environments',
      anionic: 'strong amphipathic helix',
    },
    note: 'Reference cationic AMP with strong membrane-induced helicity and a long amphipathic sequence often used in comparative studies.',
  },
  pgla: {
    name: 'PGLa',
    category: 'Membrane-active amphipathic helices',
    sequence: 'GMASKAGAIAGKIAKVALKAL',
    family: 'frog membrane-active peptide',
    profile: {
      water: 'weakly helical',
      apolar: 'helix in membrane mimics',
      anionic: 'surface-active helix; synergistic with magainin 2',
    },
    note: 'Short amphipathic helix that prefers negatively charged bilayers and is often discussed together with magainin 2.',
  },
  piscidin1: {
    name: 'Piscidin 1',
    category: 'Membrane-active amphipathic helices',
    sequence: 'FFHHIFRGIVHVGKTIHRLVTG',
    family: 'fish antimicrobial peptide',
    profile: {
      water: 'partly structured / dynamic',
      apolar: 'strong helix in membrane mimics',
      anionic: 'strong amphipathic surface-bound helix',
    },
    note: 'A strong amphipathic membrane-active helix benchmark with pronounced aromatic and histidine content.',
  },
  bp100: {
    name: 'BP100',
    category: 'Membrane-active amphipathic helices',
    sequence: 'KKLFKKILKYL',
    family: 'short synthetic cationic AMP',
    profile: {
      water: 'weakly structured in bulk water',
      apolar: 'readily helical in membrane mimics',
      anionic: 'very strong short amphipathic helix',
    },
    note: 'A short, simulation-friendly amphipathic AMP benchmark often used as a compact membrane-active helical reference.',
  },
  polyala15: {
    name: 'Poly-Ala 15',
    category: 'Apolar / helix-prone controls',
    sequence: 'AAAAAAAAAAAAAAA',
    family: 'alanine helix propensity control',
    profile: {
      water: 'helix-prone but less stabilized than charged aqueous designs',
      apolar: 'near-ideal helix in helix-promoting media',
      anionic: 'poor membrane selectivity; mostly a helicity benchmark',
    },
    note: 'A simple control highlighting strong intrinsic helical propensity, especially in helix-promoting or apolar environments.',
  },
  kalp17: {
    name: 'KALP17',
    category: 'Transmembrane/apolar model helices',
    sequence: 'GKKLALALALALALKKA',
    family: 'model membrane-spanning helix',
    profile: {
      water: 'unfavorable in water',
      apolar: 'strong hydrophobic helix benchmark',
      anionic: 'hydrophobic membrane-insertion control rather than AMP archetype',
    },
    note: 'A compact KALP-like model helix used as a transmembrane/apolar reference rather than a surface-active AMP.',
  },
  walp23: {
    name: 'WALP23',
    category: 'Transmembrane/apolar model helices',
    sequence: 'GWWLALALALALALALALALWWA',
    family: 'hydrophobic transmembrane model peptide',
    profile: {
      water: 'very unfavorable',
      apolar: 'very strong transmembrane helix',
      anionic: 'hydrophobic benchmark, not a canonical AMP',
    },
    note: 'Useful negative-control/benchmark for aqueous AMP behavior and a positive benchmark for hydrophobic transmembrane helicity.',
  },
};

function referencePeptideOptionsHtml() {
  const groups = {};
  Object.entries(REFERENCE_PEPTIDES).forEach(([key, ref]) => {
    const category = ref.category || 'Reference helices';
    if (!groups[category]) groups[category] = [];
    groups[category].push({ key, name: ref.name });
  });
  return `
    <option value="">Custom sequence</option>
    ${Object.entries(groups).map(([category, entries]) => `
      <optgroup label="${category}">
        ${entries.map((entry) => `<option value="${entry.key}">${entry.name}</option>`).join('')}
      </optgroup>
    `).join('')}
  `;
}

function forgetAnonymousUserId() {
  try { window.localStorage.removeItem(ANON_STORAGE_KEY); } catch (_) { /* Storage is optional. */ }
}

function getAnonymousUserId() {
  if (!usageLoggingEnabled) return null;
  let id;
  try { id = window.localStorage.getItem(ANON_STORAGE_KEY); } catch (_) { /* Storage is optional. */ }
  if (id) return id;
  if (window.crypto?.randomUUID) {
    id = window.crypto.randomUUID();
  } else {
    id = `anon-${Math.random().toString(36).slice(2)}-${Date.now().toString(36)}`;
  }
  try { window.localStorage.setItem(ANON_STORAGE_KEY, id); } catch (_) { /* Keep the app usable without storage. */ }
  return id;
}

function usageHeaders() {
  const id = getAnonymousUserId();
  return id ? { 'X-Anonymous-Id': id } : {};
}

function renderReferenceNote(reference) {
  if (!reference) return '';
  const entries = [
    ['Water', reference.profile.water],
    ['Apolar medium', reference.profile.apolar],
    ['Anionic membrane', reference.profile.anionic],
  ];
  return `
    <div class="reference-note-head">
      <strong>${reference.name}</strong>
      <span>${reference.family}</span>
    </div>
    <div class="reference-note-head">
      <span><strong>Category:</strong> ${reference.category || 'Reference helix'}</span>
    </div>
    <div class="reference-profile-tags">
      ${entries.map(([label, value]) => `
        <span class="reference-tag">
          <em>${label}</em>
          <strong>${value}</strong>
        </span>
      `).join('')}
    </div>
    <p>${reference.note}</p>
    <p class="reference-note-foot">Literature-inspired qualitative profile. The sequence is auto-filled; the allowed residues must include all of its residues.</p>
  `;
}

function bindReferencePeptideSelectors() {
  $$('.reference-peptide-select').forEach((select) => {
    select.innerHTML = referencePeptideOptionsHtml();
  });

  $$('.reference-peptide-select').forEach((select) => {
    const scope = select.dataset.referenceScope;
    const form = select.closest('form');
    const sequenceField = $('[name="sequence"]', form);
    const residuesField = $('[name="residues"]', form);
    const noteField = $(`[data-reference-note="${scope}"]`);
    if (!form || !sequenceField || !residuesField || !noteField) return;

    const sync = () => {
      const reference = REFERENCE_PEPTIDES[select.value];
      if (!reference) {
        noteField.classList.add('hidden-field');
        noteField.innerHTML = '';
        return;
      }
      sequenceField.value = reference.sequence;
      noteField.innerHTML = renderReferenceNote(reference);
      noteField.classList.remove('hidden-field');
    };

    select.addEventListener('change', sync);
    sequenceField.addEventListener('input', () => {
      if (select.value && sequenceField.value.trim().toUpperCase() !== REFERENCE_PEPTIDES[select.value]?.sequence) {
        select.value = '';
        sync();
      }
    });
    sync();
  });
}

function penetrationLabel(percent) {
  if (percent <= 0) return '0% (water)';
  if (percent >= 100) return '100% (apolar)';
  return `${percent}%`;
}

function isInterfacialPreset(preset) {
  return String(preset || '').startsWith('interfacial_');
}

function penetrationPercentToHalfWidth(percent) {
  const clamped = Math.max(0, Math.min(100, Number(percent) || 0));
  return 40 + (clamped / 100) * 110;
}

function buildEnvironmentFromPresetAndPenetration(preset, penetrationPercent, exactHalfWidth = null) {
  if (!isInterfacialPreset(preset)) return { preset };
  // Same mapping as the server-side penetration scan: the end points are the homogeneous media.
  const percent = Math.max(0, Math.min(100, Number(penetrationPercent ?? 50)));
  if (percent <= 0) return { preset: 'homogeneous_polar' };
  if (percent >= 100) return { preset: 'homogeneous_apolar' };
  const halfWidth = exactHalfWidth === null || exactHalfWidth === undefined || exactHalfWidth === ''
    ? penetrationPercentToHalfWidth(percent) : Number(exactHalfWidth);
  if (!Number.isFinite(halfWidth) || halfWidth < 0 || halfWidth > 180) {
    throw new Error('The apolar-sector half-width must be between 0 and 180 degrees.');
  }
  return { preset, wheel_halfwidth_deg: halfWidth };
}

function penetrationDescription(percent) {
  const value = Number(percent);
  if (value <= 0) return '0% (homogeneous polar medium)';
  if (value >= 100) return '100% (homogeneous apolar medium)';
  return `${value}% (apolar sector half-width ${penetrationPercentToHalfWidth(value).toFixed(0)}°)`;
}

function buildEnvironmentFromFormData(fd, presetFieldName, penetrationFieldName) {
  const preset = fd.get(presetFieldName);
  return buildEnvironmentFromPresetAndPenetration(
    preset, fd.get(penetrationFieldName) || 50, fd.get(`exact_halfwidth_${penetrationFieldName}`),
  );
}

function calculationSettingsFromFormData(fd, defaultCalibration = 2000) {
  const calibration = fd.get('n_decoys');
  const seed = fd.get('seed');
  const nDecoys = calibration === null || calibration === undefined || calibration === ''
    ? defaultCalibration : Number(calibration);
  const seedValue = seed === null || seed === undefined || seed === '' ? 42 : Number(seed);
  if (!Number.isInteger(nDecoys) || nDecoys < 100 || nDecoys > 10000) {
    throw new Error('The calibration sample size must be an integer between 100 and 10,000.');
  }
  if (!Number.isInteger(seedValue) || seedValue < 0 || seedValue > 4294967295) {
    throw new Error('The random seed must be an integer between 0 and 4,294,967,295.');
  }
  return { n_decoys: nDecoys, seed: seedValue };
}

function calculationSettingsHtml(defaultCalibration, angles) {
  return `
    <label>Calibration sample size
      <input type="number" name="n_decoys" min="100" max="10000" step="1" value="${defaultCalibration}" required>
      <span class="field-hint">Random sequences used to standardize the score components before scoring or design. Keep this count unchanged when reproducing a calculation.</span>
    </label>
    <label>Random seed
      <input type="number" name="seed" min="0" max="4294967295" step="1" value="42" required>
      <span class="field-hint">Controls calibration, reference sampling and search. The same seed and settings reproduce the same calculation.</span>
    </label>
    ${angles.map(({ field, scope, preset, label }) => `
      <label>${label}
        <input type="number" name="exact_halfwidth_${field}" min="0" max="180" step="any" placeholder="Use exposure slider" data-exact-halfwidth data-penetration-name="${field}" data-exposure-scope="${scope}" data-preset-name="${preset}">
        <span class="field-hint">Optional angle from the centre to the edge of the non-polar sector: 90° gives equal polar and non-polar angular sectors. It overrides intermediate exposure settings for interfacial media. Leave blank to use the slider; moving the slider clears the override. The 0% and 100% limits remain homogeneous media.</span>
      </label>
    `).join('')}
  `;
}

function initializeCalculationSettings() {
  const singleAngle = (scope, preset = 'preset') => [{
    field: 'penetration_depth', scope, preset, label: 'Exact apolar-sector half-width (°)',
  }];
  const pairedAngles = (scope) => ['a', 'b'].map((side) => ({
    field: `penetration_depth_${side}`, scope: `${scope}_${side}`, preset: `preset_${side}`,
    label: `Exact apolar-sector half-width, environment ${side.toUpperCase()} (°)`,
  }));
  const configurations = [
    ['score', 2000, singleAngle('score')],
    ['design', 2000, singleAngle('design')],
    ['family', 2000, singleAngle('family')],
    ['specificity', 10000, singleAngle('specificity_shared', 'target_preset')],
    ['compare', 2000, pairedAngles('compare')],
    ['cross', 2000, pairedAngles('cross')],
    ['penetration', 2000, []],
  ];
  configurations.forEach(([scope, calibration, angles]) => {
    const grid = $(`#${scope}-form .advanced-settings .fields-grid`);
    if (grid && !$('[name="n_decoys"]', grid)) {
      grid.insertAdjacentHTML('beforeend', calculationSettingsHtml(calibration, angles));
    }
  });
  $$('[data-exact-halfwidth]').forEach((input) => input.addEventListener('input', syncPenetrationControls));
}

function syncExactHalfWidthControls() {
  $$('[data-exact-halfwidth]').forEach((input) => {
    const form = input.closest('form');
    const slider = $(`[name="${input.dataset.penetrationName}"]`, form);
    const preset = $(`[name="${input.dataset.presetName}"]`, form);
    const percent = Number(slider.value);
    const hasInterface = isInterfacialPreset(preset.value)
      || (form.id === 'specificity-form' && $$('input[name="off_target_preset"]', form)
        .some((choice) => choice.checked && !choice.disabled && isInterfacialPreset(choice.value)));
    input.disabled = !hasInterface || percent <= 0 || percent >= 100;
    const display = $(`[data-range-value="${input.dataset.exposureScope}"]`);
    if (display && !input.disabled && input.value !== '' && input.validity.valid) {
      display.textContent = `Exact apolar half-width: ${Number(input.value)}°`;
    }
  });
}

function syncPenetrationControls() {
  $$('.env-preset').forEach((select) => {
    const scope = select.dataset.envScope;
    const field = $(`[data-penetration-field="${scope}"]`);
    if (!field) return;
    field.classList.toggle('hidden-field', !isInterfacialPreset(select.value));
  });

  $$('.penetration-slider').forEach((slider) => {
    const scope = slider.dataset.rangeOutput;
    const target = $(`[data-range-value="${scope}"]`);
    if (target) target.textContent = penetrationDescription(slider.value);
  });
  syncSpecificityControls();
  syncExactHalfWidthControls();
}

function syncLambdaBalanceControls() {
  $$('.lambda-balance-slider').forEach((slider) => {
    const scope = slider.dataset.rangeOutput;
    const target = $(`[data-range-value="${scope}"]`);
    if (target) target.textContent = Number(slider.value).toFixed(2);
  });
}

function syncSpecificityControls() {
  const targetSelect = $('[name="target_preset"]');
  const sharedPenetrationField = $('[data-penetration-field="specificity_shared"]');
  const specificityCheckboxes = $$('input[name="off_target_preset"]');
  if (!targetSelect || !sharedPenetrationField || !specificityCheckboxes.length) return;

  const targetPreset = String(targetSelect.value || '');
  const anyInterfacialOffTarget = specificityCheckboxes.some((input) => input.checked && isInterfacialPreset(input.value));
  const needsSharedPenetration = isInterfacialPreset(targetPreset) || anyInterfacialOffTarget;
  sharedPenetrationField.classList.toggle('hidden-field', !needsSharedPenetration);

  specificityCheckboxes.forEach((input) => {
    const option = input.closest('.selector-option');
    const sameAsTarget = input.value === targetPreset;
    if (sameAsTarget && !input.disabled) {
      input.dataset.wasChecked = input.checked ? 'true' : 'false';
      input.checked = false;
    } else if (!sameAsTarget && input.disabled) {
      input.checked = input.dataset.wasChecked === 'true';
    }
    input.disabled = sameAsTarget;
    option?.classList.toggle('selector-option-disabled', sameAsTarget);
  });
}

function activatePanel(btn) {
  $$('.nav-btn').forEach((b) => {
    b.classList.remove('active');
    b.removeAttribute('aria-current');
  });
  $$('.panel').forEach((p) => p.classList.remove('active'));
  btn.classList.add('active');
  btn.setAttribute('aria-current', 'page');
  $(`#panel-${btn.dataset.panel}`).classList.add('active');
}
$$('.nav-btn').forEach((btn) => btn.addEventListener('click', () => activatePanel(btn)));
const initialPanelButton = $('.nav-btn.active');
if (initialPanelButton) activatePanel(initialPanelButton);
$$('.result').forEach((node) => node.setAttribute('aria-live', 'polite'));
initializeCalculationSettings();
$$('.env-preset').forEach((select) => select.addEventListener('change', syncPenetrationControls));
$$('.penetration-slider').forEach((slider) => slider.addEventListener('input', () => {
  const exact = $(`[name="exact_halfwidth_${slider.name}"]`, slider.closest('form'));
  if (exact) exact.value = '';
  syncPenetrationControls();
}));
$$('input[name="off_target_preset"]').forEach((checkbox) => checkbox.addEventListener('change', syncPenetrationControls));
$$('.lambda-balance-slider').forEach((slider) => slider.addEventListener('input', syncLambdaBalanceControls));
syncPenetrationControls();
syncSpecificityControls();
syncLambdaBalanceControls();
bindReferencePeptideSelectors();

function parseResidues(value) {
  return value.split(',').map((v) => v.trim().toUpperCase()).filter(Boolean);
}

function loadingNode() {
  return $('#loading-template').content.firstElementChild.cloneNode(true);
}

function errorDetail(data, fallback) {
  const detail = data && data.detail;
  if (Array.isArray(detail)) {
    const messages = detail.map((item) => {
      const field = Array.isArray(item?.loc) ? item.loc.filter((part) => part !== 'body').join('.') : '';
      const message = String(item?.msg || 'Invalid value');
      return field ? `${field}: ${message}` : message;
    });
    return messages.join('; ') || fallback;
  }
  return detail ? String(detail) : fallback;
}

async function postJSON(url, payload, options = {}) {
  const res = await fetch(url, {
    method: 'POST',
    headers: {
      'Content-Type': 'application/json',
      ...usageHeaders(),
    },
    body: JSON.stringify(payload),
    signal: options.signal,
  });
  const data = await res.json().catch(() => null);
  if (!res.ok) throw new Error(errorDetail(data, `Request failed (HTTP ${res.status})`));
  if (data === null) throw new Error('The server returned an invalid response.');
  if (url !== '/api/visit') {
    refreshUsageCounters().catch(() => {});
  }
  return data;
}

async function postNDJSON(url, payload, onMessage, options = {}) {
  const res = await fetch(url, {
    method: 'POST',
    headers: {
      'Content-Type': 'application/json',
      ...usageHeaders(),
    },
    body: JSON.stringify(payload),
    signal: options.signal,
  });
  if (!res.ok) {
    const data = await res.json().catch(() => null);
    throw new Error(errorDetail(data, `Streaming request failed (HTTP ${res.status})`));
  }
  if (!res.body) {
    throw new Error('Streaming not supported by this browser');
  }

  const reader = res.body.getReader();
  const decoder = new TextDecoder();
  let buffer = '';

  while (true) {
    const { value, done } = await reader.read();
    if (done) break;
    buffer += decoder.decode(value, { stream: true });
    const lines = buffer.split('\n');
    buffer = lines.pop() || '';
    for (const line of lines) {
      const trimmed = line.trim();
      if (!trimmed) continue;
      onMessage(parseStreamLine(trimmed));
    }
  }
  if (buffer.trim()) {
    onMessage(parseStreamLine(buffer.trim()));
  }
}

function parseStreamLine(line) {
  try {
    return JSON.parse(line);
  } catch (_) {
    throw new Error('Received an incomplete response from the server.');
  }
}

function compareSpecificityCandidates(a, b) {
  // Same ordering as the server: objective, larger ΔZ, lower target Z, sequence.
  const objective = (item) => Math.round(Number(item.specificity_design?.objective || 0) * 1e8) / 1e8;
  return objective(a) - objective(b)
    || Number(b.specificity_design?.z_score_margin || 0) - Number(a.specificity_design?.z_score_margin || 0)
    || Number(designZ(a.target_environment)) - Number(designZ(b.target_environment))
    || String(a.sequence).localeCompare(String(b.sequence));
}

async function registerVisitCounter() {
  try {
    const response = await fetch('/api/health', { cache: 'no-store' });
    const config = await response.json();
    usageLoggingEnabled = config.usage_logging === true;
  } catch (_) {
    usageLoggingEnabled = false;
  }
  await refreshUsageCounters(true);
}

async function refreshUsageCounters(logVisit = false) {
  const counters = document.getElementById('footer-stats');
  if (counters) counters.style.display = usageLoggingEnabled ? '' : 'none';
  if (!usageLoggingEnabled) {
    forgetAnonymousUserId();
    return;
  }
  const totalEl = document.getElementById('visit-counter-total');
  const uniqueEl = document.getElementById('visit-counter-unique');
  const runsEl = document.getElementById('visit-counter-runs');
  if (!totalEl || !uniqueEl || !runsEl) return;
  try {
    const data = logVisit
      ? await postJSON('/api/visit', {})
      : await fetch('/api/usage-metrics', {
          headers: {
            ...usageHeaders(),
          },
        }).then(async (res) => {
          const payload = await res.json().catch(() => ({}));
          if (!res.ok) throw new Error(payload.detail || 'Unable to refresh usage metrics');
          return payload;
        });
    totalEl.textContent = `Visits: ${Number(data.total_visits || 0).toLocaleString()}`;
    uniqueEl.textContent = `Unique visitors: ${Number(data.unique_visitors || 0).toLocaleString()}`;
    runsEl.textContent = `Runs: ${Number(data.total_runs || 0).toLocaleString()}`;
  } catch (error) {
    totalEl.textContent = 'Visits: —';
    uniqueEl.textContent = 'Unique visitors: —';
    runsEl.textContent = 'Runs: —';
  }
}

function escapeHtml(text) {
  return String(text ?? '').replace(/[&<>"']/g, (m) => ({ '&': '&amp;', '<': '&lt;', '>': '&gt;', '"': '&quot;', "'": '&#39;' }[m]));
}

function downloadJson(filename, data) {
  const blob = new Blob([JSON.stringify(data, null, 2)], { type: 'application/json' });
  const url = URL.createObjectURL(blob);
  const a = document.createElement('a');
  a.href = url;
  a.download = filename;
  document.body.appendChild(a);
  a.click();
  a.remove();
  URL.revokeObjectURL(url);
}

function downloadText(filename, text, type = 'text/plain') {
  const blob = new Blob([text], { type });
  const url = URL.createObjectURL(blob);
  const a = document.createElement('a');
  a.href = url;
  a.download = filename;
  document.body.appendChild(a);
  a.click();
  a.remove();
  URL.revokeObjectURL(url);
}

let viewerCounter = 0;
let svgIdCounter = 0;

function nextSvgId(prefix) {
  svgIdCounter += 1;
  return `${prefix}-${svgIdCounter}`;
}

function nextViewerId(prefix = 'pdb-viewer') {
  viewerCounter += 1;
  return `${prefix}-${viewerCounter}`;
}

function ensure3Dmol() {
  if (window.$3Dmol) return Promise.resolve(window.$3Dmol);
  return new Promise((resolve, reject) => {
    const interval = setInterval(() => {
      if (window.$3Dmol) { clearInterval(interval); resolve(window.$3Dmol); }
    }, 50);
    setTimeout(() => { clearInterval(interval); reject(new Error('3D viewer could not be loaded')); }, 10000);
  });
}

let activePdbViewer = null;

function releasePdbViewer() {
  if (!activePdbViewer) return;
  try {
    const canvas = activePdbViewer.getCanvas?.();
    const gl = canvas?.getContext('webgl2') || canvas?.getContext('webgl');
    gl?.getExtension('WEBGL_lose_context')?.loseContext();
  } catch (_) { /* The old viewer is being discarded anyway. */ }
  activePdbViewer = null;
}

async function mountPdbViewer(containerId, pdb) {
  const container = document.getElementById(containerId);
  if (!container) return;
  try {
    const $3Dmol = await ensure3Dmol();
    releasePdbViewer();
    const viewer = $3Dmol.createViewer(container, {
      backgroundColor: '#fbf6ec',
    });
    viewer.clear();
    viewer.addModel(pdb, 'pdb');
    viewer.setStyle({}, {
      cartoon: {
        color: '#dde6eb',
        opacity: 0.96,
        thickness: 0.45,
        style: 'oval',
      },
    });
    viewer.addStyle(
      { not: { atom: ['N', 'C', 'O', 'OXT'] } },
      { stick: { colorscheme: 'Jmol', radius: 0.18 } },
    );
    viewer.addStyle(
      { atom: ['N', 'C', 'O', 'OXT'] },
      { stick: { color: '#7c8e97', radius: 0.08 } },
    );
    viewer.zoomTo();
    viewer.zoom(1.12);
    viewer.rotate(90, 'y');
    viewer.rotate(10, 'z');
    viewer.render();
    activePdbViewer = viewer;
  } catch (error) {
    container.innerHTML = `<div class="viewer-error">${escapeHtml(error.message)}</div>`;
  }
}

function wheelPoint(thetaDeg, radius, cx, cy) {
  const theta = (thetaDeg * Math.PI) / 180;
  return {
    x: cx + radius * Math.sin(theta),
    y: cy + radius * Math.cos(theta),
  };
}

function polarLabelSvg(cx, cy, radius, halfWidth) {
  const badge = (label, y, fill, stroke, textFill) => `
    <g>
      <rect x="${cx - 58}" y="${y - 18}" rx="13" ry="13" width="116" height="28" fill="${fill}" stroke="${stroke}" />
      <text x="${cx}" y="${y - 1}" text-anchor="middle" dominant-baseline="middle" font-size="14" font-weight="700" fill="${textFill}">${label}</text>
    </g>
  `;
  if (halfWidth >= 180) {
    return badge('non-polar', cy + 6, 'rgba(176,90,27,.12)', 'rgba(120,60,18,.18)', 'rgba(120,60,18,.86)');
  }
  if (halfWidth <= 0) {
    return badge('polar', cy + 6, 'rgba(38,93,203,.10)', 'rgba(24,76,164,.18)', 'rgba(24,76,164,.88)');
  }
  return `
    ${badge('polar', cy - radius - 110, 'rgba(38,93,203,.10)', 'rgba(24,76,164,.18)', 'rgba(24,76,164,.88)')}
    ${badge('non-polar', cy + radius + 110, 'rgba(176,90,27,.12)', 'rgba(120,60,18,.18)', 'rgba(120,60,18,.86)')}
  `;
}

function hydrophobicMomentArrowSvg(nodeGeometry, cx, cy, arrowLength) {
  let mx = 0;
  let my = 0;
  for (const point of nodeGeometry) {
    const h = HYDRO_SCALE[point.item.residue] ?? 0;
    const dx = point.x - cx;
    const dy = point.y - cy;
    const norm = Math.hypot(dx, dy) || 1;
    mx += h * (dx / norm);
    my += h * (dy / norm);
  }
  const magnitude = Math.hypot(mx, my);
  if (magnitude < 1e-8) return '';
  const ux = mx / magnitude;
  const uy = my / magnitude;
  const x2 = cx + ux * arrowLength;
  const y2 = cy + uy * arrowLength;
  const px = -uy;
  const py = ux;
  const labelX = cx + ux * (arrowLength * 0.62) + px * 24;
  const labelY = cy + uy * (arrowLength * 0.62) + py * 24;
  const markerId = nextSvgId('mu-arrow-head');
  return `
    <defs>
      <marker id="${markerId}" markerWidth="10" markerHeight="10" refX="8" refY="4" orient="auto" markerUnits="strokeWidth">
        <path d="M0,0 L8,4 L0,8 z" fill="rgba(174,25,122,.96)" />
      </marker>
    </defs>
    <line
      x1="${cx}"
      y1="${cy}"
      x2="${x2.toFixed(2)}"
      y2="${y2.toFixed(2)}"
      stroke="rgba(255,255,255,.82)"
      stroke-width="8"
      stroke-linecap="round"
    />
    <line
      x1="${cx}"
      y1="${cy}"
      x2="${x2.toFixed(2)}"
      y2="${y2.toFixed(2)}"
      stroke="rgba(174,25,122,.96)"
      stroke-width="5.5"
      stroke-linecap="round"
      marker-end="url(#${markerId})"
    />
    <circle cx="${cx}" cy="${cy}" r="5" fill="rgba(174,25,122,.96)" stroke="rgba(255,255,255,.92)" stroke-width="2" />
    <rect
      x="${(labelX - 20).toFixed(2)}"
      y="${(labelY - 14).toFixed(2)}"
      width="40"
      height="24"
      rx="12"
      ry="12"
      fill="rgba(255,255,255,.92)"
      stroke="rgba(174,25,122,.22)"
    />
    <text
      x="${labelX.toFixed(2)}"
      y="${(labelY + 1).toFixed(2)}"
      text-anchor="middle"
      dominant-baseline="middle"
      font-size="16"
      font-weight="800"
      fill="rgba(174,25,122,.96)"
    >μ<tspan baseline-shift="sub" font-size="11">H</tspan></text>
  `;
}

function wheelSvg(wheel, environment) {
  const boundaryRadius = 215;
  const nodeBaseRadius = 136;
  const radialStepPerTurn = 22;
  const cx = 300;
  const cy = 340;
  const halfWidth = Number(environment.wheel_halfwidth_deg);
  // Same rule as the model: a residue faces the apolar side when |angle| <= half-width,
  // with angle 0 at the bottom of the wheel. The sector is therefore an angular wedge.
  let background = `<circle cx="${cx}" cy="${cy}" r="${boundaryRadius}" fill="rgba(38,93,203,.10)" />`;
  if (halfWidth >= 180) {
    background = `<circle cx="${cx}" cy="${cy}" r="${boundaryRadius}" fill="rgba(176,90,27,.12)" />`;
  } else if (halfWidth > 0) {
    const start = wheelPoint(-halfWidth, boundaryRadius, cx, cy);
    const end = wheelPoint(halfWidth, boundaryRadius, cx, cy);
    const largeArc = halfWidth > 90 ? 1 : 0;
    background = `
      <circle cx="${cx}" cy="${cy}" r="${boundaryRadius}" fill="rgba(38,93,203,.10)" />
      <path d="M ${cx} ${cy} L ${start.x.toFixed(2)} ${start.y.toFixed(2)} A ${boundaryRadius} ${boundaryRadius} 0 ${largeArc} 0 ${end.x.toFixed(2)} ${end.y.toFixed(2)} Z"
        fill="rgba(176,90,27,.12)" stroke="rgba(29,39,51,.18)" stroke-width="1.5" />
    `;
  }
  const nodeGeometry = wheel.map((item, idx) => {
    const turnIndex = idx / 3.6;
    const radialOffset = nodeBaseRadius + turnIndex * radialStepPerTurn;
    const nodeRadius = Math.max(11.5, 17 - turnIndex * 0.75);
    const { x, y } = wheelPoint(item.angle_deg, radialOffset, cx, cy);
    const labelY = y + Math.max(20, nodeRadius + 10);
    return { item, x, y, nodeRadius, labelY, turnIndex };
  });
  const sequenceSegments = nodeGeometry.slice(1).map((point, idx) => {
    const prev = nodeGeometry[idx];
    const segmentTurn = (prev.turnIndex + point.turnIndex) / 2;
    const fade = Math.min(1, segmentTurn / Math.max(1, (wheel.length - 1) / 3.6));
    const opacity = 0.34 - fade * 0.18;
    const width = 2.8 - fade * 1.25;
    return `
      <line
        x1="${prev.x.toFixed(2)}"
        y1="${prev.y.toFixed(2)}"
        x2="${point.x.toFixed(2)}"
        y2="${point.y.toFixed(2)}"
        stroke="rgba(29,39,51,${opacity.toFixed(3)})"
        stroke-width="${width.toFixed(2)}"
        stroke-linecap="round"
      />
    `;
  }).join('');
  const nodes = nodeGeometry.map(({ item, x, y, nodeRadius, labelY, turnIndex }) => {
    const ring = item.environment === 'membrane' ? '#9a4c12' : '#1c4fb0';
    return `
      <circle cx="${x}" cy="${y}" r="${nodeRadius + 3}" fill="none" stroke="${ring}" stroke-width="2.5" />
      <circle cx="${x}" cy="${y}" r="${nodeRadius}" fill="${item.color}" />
      <text x="${x}" y="${y + 4}" text-anchor="middle" font-size="${Math.max(10, 12 - turnIndex * 0.35)}" font-weight="800" fill="white">${item.residue}</text>
      <text x="${x}" y="${labelY}" text-anchor="middle" font-size="11" font-weight="600" fill="#5d6a74">${item.index}</text>
    `;
  }).join('');
  const hydroMomentArrow = hydrophobicMomentArrowSvg(nodeGeometry, cx, cy, boundaryRadius * 0.62);
  return `
    <div class="wheel-wrap">
      <svg class="wheel" viewBox="0 -40 600 860" role="img" aria-label="Helical wheel: residues facing the apolar side are ringed in brown, residues facing the polar side in blue">
        ${background}
        ${polarLabelSvg(cx, cy, boundaryRadius, halfWidth)}
        <circle cx="${cx}" cy="${cy}" r="${boundaryRadius}" fill="none" stroke="rgba(29,39,51,.16)" stroke-width="1.5" />
        ${hydroMomentArrow}
        ${sequenceSegments}
        ${nodes}
      </svg>
      <div class="wheel-legend">
        <span><i class="ring ring-apolar"></i>faces apolar side</span>
        <span><i class="ring ring-polar"></i>faces polar side</span>
        ${Object.entries(RESIDUE_FAMILY_LABELS).map(([family, label]) => `<span><i class="dot dot-${family}"></i>${label}</span>`).join('')}
      </div>
    </div>
  `;
}

const RESIDUE_FAMILY_LABELS = {
  hydrophobic: 'hydrophobic',
  polar: 'polar',
  positive: 'positive',
  negative: 'negative',
  special: 'histidine',
};

function zScorePlot(data) {
  const z = Number(data.z_score);
  const bound = Math.max(4, Math.ceil(Math.abs(z)));
  const x = 24 + ((z + bound) / (2 * bound)) * 312;
  const gradientId = nextSvgId('zBand');
  const tick = (value) => (value === 0 ? '0' : `${value > 0 ? '+' : ''}${value}σ`);
  const percentile = data.percentile_lower_is_better.toFixed(1);
  const mean = Number(data.random_reference.mean).toFixed(2);
  const std = Number(data.random_reference.std).toFixed(2);
  return `
    <div class="score-plot">
      <h3>Z-score position</h3>
      <svg viewBox="0 0 360 120" width="100%" aria-label="Z-score plot">
        <defs>
          <linearGradient id="${gradientId}" x1="0%" y1="0%" x2="100%" y2="0%">
            <stop offset="0%" stop-color="rgba(203,58,43,.20)" />
            <stop offset="35%" stop-color="rgba(187,90,36,.18)" />
            <stop offset="50%" stop-color="rgba(14,90,109,.10)" />
            <stop offset="65%" stop-color="rgba(14,90,109,.18)" />
            <stop offset="100%" stop-color="rgba(14,90,109,.26)" />
          </linearGradient>
        </defs>
        <rect x="24" y="44" width="312" height="16" rx="8" fill="url(#${gradientId})" />
        <line x1="180" y1="34" x2="180" y2="76" stroke="rgba(29,39,51,.35)" stroke-dasharray="4 4" />
        <line x1="${x}" y1="28" x2="${x}" y2="84" stroke="#1d2733" stroke-width="3" />
        <circle cx="${x}" cy="52" r="8" fill="#1d2733" />
        <text x="24" y="98" font-size="11" fill="#5d6a74">${tick(-bound)}</text>
        <text x="102" y="98" font-size="11" fill="#5d6a74" text-anchor="middle">${tick(-bound / 2)}</text>
        <text x="180" y="98" font-size="11" fill="#5d6a74" text-anchor="middle">0</text>
        <text x="258" y="98" font-size="11" fill="#5d6a74" text-anchor="middle">${tick(bound / 2)}</text>
        <text x="336" y="98" font-size="11" fill="#5d6a74" text-anchor="end">${tick(bound)}</text>
      </svg>
      <p class="plot-caption">Mean random energy ${mean} with σ ${std}. This sequence sits at Z = ${z.toFixed(3)} and percentile ${percentile}% (lower is better).</p>
    </div>
  `;
}

function penetrationProfilePlot(data) {
  const profile = data.profile || [];
  if (!profile.length) return '';
  const width = 720;
  const height = 220;
  const padL = 54;
  const padR = 96;
  const padT = 20;
  const padB = 40;
  const zValues = profile.map((p) => Number(p.z_score));
  const eValues = profile.map((p) => Number(p.energy));
  const minZ = Math.min(...zValues);
  const maxZ = Math.max(...zValues);
  const zSpan = Math.max(1e-6, maxZ - minZ);
  const minE = Math.min(...eValues);
  const maxE = Math.max(...eValues);
  const eSpan = Math.max(1e-6, maxE - minE);
  const xFor = (percent) => padL + ((percent - profile[0].penetration_percent) / Math.max(1, profile[profile.length - 1].penetration_percent - profile[0].penetration_percent)) * (width - padL - padR);
  const yFor = (z) => padT + ((maxZ - z) / zSpan) * (height - padT - padB);
  const yForEnergy = (energy) => padT + ((maxE - energy) / eSpan) * (height - padT - padB);
  const path = profile.map((p, idx) => `${idx === 0 ? 'M' : 'L'} ${xFor(p.penetration_percent).toFixed(2)} ${yFor(p.z_score).toFixed(2)}`).join(' ');
  const energyPath = profile.map((p, idx) => `${idx === 0 ? 'M' : 'L'} ${xFor(p.penetration_percent).toFixed(2)} ${yForEnergy(p.energy).toFixed(2)}`).join(' ');
  const best = profile.find((p) => p.penetration_percent === data.best_penetration_percent) || profile[0];
  const xTicks = profile.map((p) => p.penetration_percent);
  const yTicks = [minZ, minZ + zSpan / 2, maxZ];
  const yEnergyTicks = [minE, minE + eSpan / 2, maxE];
  return `
    <div class="score-plot penetration-plot">
      <h3>Z-score vs penetration</h3>
      <p class="plot-caption"><span style="display:inline-flex;align-items:center;gap:6px;margin-right:14px;"><span style="display:inline-block;width:14px;height:3px;background:#0e5a6d;border-radius:999px;"></span>Z-score</span><span style="display:inline-flex;align-items:center;gap:6px;"><span style="display:inline-block;width:14px;height:3px;background:#bb5a24;border-radius:999px;"></span>Energy</span></p>
      <svg viewBox="0 0 ${width} ${height}" width="100%" aria-label="penetration profile plot">
        ${yTicks.map((tick) => `
          <line x1="${padL}" y1="${yFor(tick)}" x2="${width - padR}" y2="${yFor(tick)}" stroke="rgba(29,39,51,.10)" />
          <text x="${padL - 10}" y="${yFor(tick) + 4}" text-anchor="end" font-size="12" fill="#5d6a74">${tick.toFixed(2)}</text>
        `).join('')}
        ${yEnergyTicks.map((tick) => `
          <text x="${width - 8}" y="${yForEnergy(tick) + 4}" text-anchor="end" font-size="12" fill="#bb5a24">${tick.toFixed(2)}</text>
        `).join('')}
        ${xTicks.map((tick) => `
          <line x1="${xFor(tick)}" y1="${padT}" x2="${xFor(tick)}" y2="${height - padB}" stroke="rgba(29,39,51,.06)" />
          <text x="${xFor(tick)}" y="${height - 12}" text-anchor="middle" font-size="11" fill="#5d6a74">${tick === 0 ? 'water' : (tick === 100 ? 'apolar' : `${tick}%`)}</text>
        `).join('')}
        <path d="${energyPath}" fill="none" stroke="#bb5a24" stroke-width="2.5" stroke-linecap="round" stroke-linejoin="round" opacity="0.95" />
        <path d="${path}" fill="none" stroke="#0e5a6d" stroke-width="3" stroke-linecap="round" stroke-linejoin="round" />
        ${profile.map((p) => `
          <circle cx="${xFor(p.penetration_percent)}" cy="${yForEnergy(p.energy)}" r="3.5" fill="#bb5a24" />
        `).join('')}
        ${profile.map((p) => `
          <circle cx="${xFor(p.penetration_percent)}" cy="${yFor(p.z_score)}" r="${p.penetration_percent === data.best_penetration_percent ? 6 : 4}" fill="${p.penetration_percent === data.best_penetration_percent ? '#cb3a2b' : '#0e5a6d'}" />
        `).join('')}
      </svg>
      <p class="plot-caption">The scan runs from homogeneous polar medium at 0% to homogeneous apolar medium at 100%, with the selected interfacial preset applied only at intermediate points. Z-score is read on the left axis and energy on the right axis. Current best by Z-score: ${penetrationLabel(best.penetration_percent)} with Z = ${best.z_score.toFixed(3)} and E = ${best.energy.toFixed(3)}.</p>
    </div>
  `;
}

const HELP_TEXT = {
  stats: {
    'Model energy': 'Hamiltonian energy in dimensionless model units. Lower is better, but values are only meaningful comparatively within the same scoring setup.',
    'Z-score': 'Energy expressed relative to a random-sequence background. More negative means more exceptional.',
    'Percentile (lower better)': 'Fraction of random sequences with energy equal to or lower than this one. Lower is better.',
    'Net charge': 'Net charge under the model convention: Lys and Arg +1, Asp and Glu -1; histidine and the termini are treated as neutral.',
    'Joint energy': 'Sum of the two environment energies in Compare mode.',
    'Energy gap': 'Absolute difference between the sequence model energies in the two environments.',
    'Z-score gap': 'Absolute difference between the two Z-scores.',
    'Length': 'Number of residues in the sequence.',
    'Best penetration': 'Sampled point that gives the most negative Z-score along the scan from homogeneous polar medium (0%) to homogeneous apolar medium (100%). The same percentage reproduces this environment in the other panels.',
    'Sampled depths': 'Number of penetration-depth values tested in the scan.',
    'Interface': 'Selected interfacial preset used for the penetration scan.',
    'Optimization target': 'Criterion used to define the optimum. Here it is the minimum Z-score across the scan.',
    'Requested': 'Requested family size.',
    'Returned': 'Number of unique candidates actually found.',
    'Environment': 'Selected environment preset.',
    'Alphabet size': 'Number of amino-acid types allowed during scoring or design.',
    'Joint objective': 'Cross-design objective combining both environments and the λgap penalty.',
    'λgap': 'Penalty weight that discourages sequences from behaving too differently across environments.',
    'Specificity objective': 'Optimization target for one-vs-rest design. Lower values mean the target stays more favorable relative to the best competing off-target.',
    'λ balance': 'Interpolation between the hard one-vs-rest limit and target-only optimization. λ=0 gives the strongest selectivity pressure; λ=1 reduces the objective to Htarget alone.',
    'Energy margin': 'Difference between the worst competing off-target model energy and the target model energy. Larger positive values mean cleaner target-vs-off-target separation.',
    'Z-score margin': 'Lowest off-target Z-score minus the target Z-score. Larger positive values mean cleaner target-vs-off-target separation.',
    'Worst off-target': 'Off-target environment with the lowest model energy for the designed sequence (used for the energy margin).',
  },
  breakdown: {
    env: 'Environment term: preference of residue types for the chosen medium or sector.',
    charge: 'Charge term: compatibility with membrane surface charge in interfacial modes.',
    local: 'Local helix term: residue-level alpha-helical propensity.',
    pairwise: 'Pairwise term: sequence-dependent interaction preferences between positions.',
    electrostatic: 'Electrostatic term: attractive and repulsive charge interactions.',
    neigh: 'Helix-neighbor term: short-range helical cooperativity along the sequence.',
    total: 'Weighted sum of all Hamiltonian contributions.',
  },
  observables: {
    abs_mu_h: 'Absolute transverse hydrophobic moment computed on the helical wheel. It is shown as a geometric descriptor and is not part of the current Hamiltonian energy.',
    abs_mu_h_per_residue: 'Absolute transverse hydrophobic moment divided by sequence length, useful for comparing helices of different sizes.',
  },
};

function settingsLine(data) {
  const env = data.environment || {};
  const parts = [humanizePreset(env.preset)];
  if (isInterfacialPreset(env.preset)) {
    parts.push(`apolar half-width ${Number(env.wheel_halfwidth_deg).toFixed(1)}°`);
    parts.push(`wheel phase ${Number(env.wheel_phase_deg).toFixed(1)}°`);
  }
  if (data.alphabet) parts.push(`alphabet ${data.alphabet.join('')}`);
  if (data.calibration) parts.push(`calibration ${data.calibration.n_decoys} decoys, seed ${data.calibration.seed_entropy}`);
  if (data.random_reference?.n) parts.push(`random reference n = ${data.random_reference.n}`);
  if (data.design_settings) {
    parts.push(`steps ${data.design_settings.steps}, restarts ${data.design_settings.restarts}, seed ${data.design_settings.seed}`);
  }
  return `<div class="settings-line">Settings: ${escapeHtml(parts.join(' · '))}</div>`;
}

function energyUnitNote() {
  return `<div class="energy-unit-note">Model energies are reported in dimensionless Hamiltonian units, not in physical units such as kcal/mol. Use them comparatively within the same environment setup, alphabet and weight set.</div>`;
}

function statCard(label, value) {
  const hint = HELP_TEXT.stats[label] || '';
  const titleAttr = hint ? ` title="${escapeHtml(hint)}"` : '';
  return `<div class="stat"${titleAttr}><span>${label}</span><strong>${value}</strong></div>`;
}

function humanizePreset(preset) {
  const labels = {
    homogeneous_polar: 'Homogeneous polar',
    homogeneous_apolar: 'Homogeneous apolar',
    interfacial_neg: 'Interfacial anionic',
    interfacial_neu: 'Interfacial neutral',
    interfacial_pos: 'Interfacial cationic',
  };
  return labels[preset] || String(preset || 'Environment');
}

function breakdownGrid(breakdown) {
  const labels = {
    env: { name: 'Environment', symbol: 'H<sub>env</sub>' },
    charge: { name: 'Charge', symbol: 'H<sub>chg</sub>' },
    local: { name: 'Local helix', symbol: 'H<sub>local</sub>' },
    pairwise: { name: 'Pairwise', symbol: 'H<sub>pair</sub>' },
    electrostatic: { name: 'Electrostatics', symbol: 'H<sub>elec</sub>' },
    neigh: { name: 'Helix neighbors', symbol: 'H<sub>neigh</sub>' },
    total: { name: 'Total', symbol: 'H<sub>tot</sub>' },
  };
  return `
    <div class="breakdown-section">
      <div class="breakdown-head">
        <h3>Hamiltonian Contributions</h3>
        <div class="mini-equation">H = Σ λ<sub>i</sub> H<sub>i</sub></div>
        <p>Weighted contributions of each term to the total model energy for this sequence and environment.</p>
      </div>
      <div class="breakdown">${Object.entries(breakdown).map(([k, v]) => `
        <div class="breakdown-item" title="${escapeHtml(HELP_TEXT.breakdown[k] || '')}">
          <div class="key">
            <span class="term-symbol">${labels[k]?.symbol || k}</span>
            <span class="term-name">${labels[k]?.name || k}</span>
          </div>
          <div class="val">${Number(v).toFixed(3)}</div>
        </div>
      `).join('')}</div>
    </div>
  `;
}

function observableGrid(observables = {}) {
  const items = [
    {
      key: 'abs_mu_h',
      symbol: '|μ_H|',
      name: 'Hydrophobic moment descriptor',
      value: Number(observables.abs_mu_h ?? 0).toFixed(3),
    },
    {
      key: 'abs_mu_h_per_residue',
      symbol: '|μ_H| / L',
      name: 'Per-residue hydrophobic moment',
      value: Number(observables.abs_mu_h_per_residue ?? 0).toFixed(4),
    },
  ];
  return `
    <div class="observable-strip">
      ${items.map((item) => `
        <div class="observable-item" title="${escapeHtml(HELP_TEXT.observables[item.key] || '')}">
          <div class="key">
            <span class="observable-symbol">${item.symbol}</span>
            <span>${item.name}</span>
          </div>
          <div class="val">${item.value}</div>
        </div>
      `).join('')}
    </div>
  `;
}

function renderScore(data) {
  const chargeValue = `${data.model_net_charge}`;
  return `
    <div class="result-card">
      <div class="sequence-banner">${data.sequence}</div>
      <div class="result-grid">
        ${statCard('Model energy', data.energy.toFixed(3))}
        ${statCard('Z-score', data.z_score.toFixed(3))}
        ${statCard('Percentile (lower better)', `${data.percentile_lower_is_better.toFixed(1)}%`)}
        ${statCard('Net charge', chargeValue)}
      </div>
      ${settingsLine(data)}
      ${energyUnitNote()}
      <div class="result-note">Interpretation: lower energy and more negative Z-score indicate stronger compatibility with the selected helical state within this model.</div>
      <div class="viz-grid">
        <div>
          ${zScorePlot(data)}
          ${observableGrid(data.observables)}
          ${breakdownGrid(data.breakdown)}
        </div>
        ${wheelSvg(data.wheel, data.environment)}
      </div>
    </div>
  `;
}

function renderCompareEnvironmentCard(data, label) {
  const chargeValue = `${data.model_net_charge}`;
  return `
    <div class="compare-env-card">
      <div class="compare-env-head">
        <div>
          <h3>${label}</h3>
          <p>${humanizePreset(data.environment?.preset)}</p>
        </div>
      </div>
      <div class="compare-env-stats">
        ${statCard('Model energy', data.energy.toFixed(3))}
        ${statCard('Z-score', data.z_score.toFixed(3))}
        ${statCard('Percentile (lower better)', `${data.percentile_lower_is_better.toFixed(1)}%`)}
        ${statCard('Net charge', chargeValue)}
      </div>
      ${settingsLine(data)}
      <div class="compare-env-note">Lower energy and more negative Z-score indicate stronger compatibility with this environment within the model.</div>
      <div class="compare-env-viz">
        ${zScorePlot(data)}
        ${observableGrid(data.observables)}
        ${breakdownGrid(data.breakdown)}
      </div>
      <div class="compare-env-wheel">
        ${wheelSvg(data.wheel, data.environment)}
      </div>
    </div>
  `;
}

function renderCompare(data, header = '') {
  return `
    <div class="result-card">
      <div class="sequence-banner">${data.sequence}</div>
      ${header}
      <div class="result-grid">
        ${statCard('Joint energy', data.joint_energy.toFixed(3))}
        ${statCard('Energy gap', data.energy_gap.toFixed(3))}
        ${statCard('Z-score gap', data.z_score_gap.toFixed(3))}
        ${statCard('Length', data.sequence.length)}
      </div>
      ${energyUnitNote()}
      <div class="compare-grid">
        ${renderCompareEnvironmentCard(data.environment_a, 'Environment A')}
        ${renderCompareEnvironmentCard(data.environment_b, 'Environment B')}
      </div>
    </div>
  `;
}

function renderSpecificity(data) {
  const candidates = data.candidates && data.candidates.length ? data.candidates : [data];
  const selected = candidates[0];
  const target = selected.target_environment;
  const offTargets = selected.off_target_environments || [];
  const specificity = selected.specificity_design || {};
  const lambdaBalance = Number(specificity.lambda_balance ?? 0.0);
  const bestObjective = Number(candidates[0]?.specificity_design?.objective || 0);
  const offTargetOrder = (data.specificity_collection?.off_target_presets || offTargets.map((item) => item.environment?.preset)).filter(Boolean);
  const tableHeaders = [
    { key: 'rank', label: 'Rank' },
    { key: 'sequence', label: 'Sequence' },
    { key: 'objective_delta', label: 'Δobjective vs best' },
    { key: 'target_z', label: `Z ${humanizePreset(specificity.target_preset || target.environment?.preset)}` },
    ...offTargetOrder.map((preset) => ({ key: preset, label: `Z ${humanizePreset(preset)}` })),
    { key: 'worst', label: 'Worst off-target' },
    { key: 'margin', label: 'ΔZ' },
  ];
  return `
    <div class="result-card">
      <div class="toolbar">
        <button class="ghost-btn" type="button" data-download-specificity-csv>Download CSV</button>
        <button class="ghost-btn" type="button" data-download-specificity-json>Download JSON</button>
      </div>
      ${data.specificity_collection?.partial ? `<div class="result-note partial-note" role="status">Partial result (${escapeHtml(data.specificity_collection.partial_reason || 'stopped early')}): ${candidates.length} of ${Number(data.specificity_collection.requested || candidates.length)} requested candidates.</div>` : ''}
      <div class="sequence-banner">${selected.sequence}</div>
      <div class="result-grid">
        ${statCard('Requested candidates', Number(data.specificity_collection?.requested || candidates.length))}
        ${statCard('Returned candidates', candidates.length)}
        ${statCard('λ balance', lambdaBalance.toFixed(2))}
        ${statCard('Best objective', bestObjective.toFixed(6))}
        ${statCard('Selected ΔZ', Number(specificity.z_score_margin || 0).toFixed(3))}
      </div>
      ${energyUnitNote()}
      <div class="result-note">This mode minimizes L<sub>&lambda;</sub> = &lambda;H<sub>target</sub> + (1-&lambda;)(H<sub>target</sub> - min(H<sub>off-targets</sub>)). Lower objective values are better. Every proposed sequence is aligned by its hydrophobic moment in interfacial environments, using the same scoring rule as the Score panel. The objective and margins are calculated from the environment scores shown below.</div>
      <div class="specificity-multi">
        <div class="specificity-table-wrap">
          <table class="specificity-table">
            <thead>
              <tr>${tableHeaders.map((header) => `<th>${escapeHtml(header.label)}</th>`).join('')}</tr>
            </thead>
            <tbody>
              ${candidates.map((candidate, idx) => {
                const envMap = Object.fromEntries(
                  (candidate.off_target_environments || []).map((item) => [item.environment?.preset, item])
                );
                return `
                  <tr class="${idx === 0 ? 'is-selected' : ''}" data-specificity-row="${idx}">
                    <td>${idx + 1}</td>
                    <td>
                      <button type="button" class="specificity-pick" data-specificity-pick="${idx}">
                        ${escapeHtml(candidate.sequence)}
                      </button>
                    </td>
                    <td>${formatObjectiveDelta(candidate.specificity_design?.objective, bestObjective)}</td>
                    <td>${Number(designZ(candidate.target_environment)).toFixed(3)}</td>
                    ${offTargetOrder.map((preset) => `<td>${Number(designZ(envMap[preset])).toFixed(3)}</td>`).join('')}
                    <td>${escapeHtml(humanizePreset(candidate.specificity_design?.worst_off_target_preset))}</td>
                    <td>${Number(candidate.specificity_design?.z_score_margin || 0).toFixed(3)}</td>
                  </tr>
                `;
              }).join('')}
            </tbody>
          </table>
        </div>
        <div class="specificity-detail" data-specificity-detail>
          ${renderSpecificityCandidateDetail(selected)}
        </div>
      </div>
    </div>
  `;
}

function renderSpecificityCandidateDetail(candidate) {
  const target = candidate.target_environment;
  const offTargets = candidate.off_target_environments || [];
  const specificity = candidate.specificity_design || {};
  return `
    <div class="result-grid">
      ${statCard('Selected sequence', `<code>${escapeHtml(candidate.sequence)}</code>`)}
      ${statCard('Specificity objective', Number(specificity.objective || 0).toFixed(3))}
      ${statCard('Energy margin', Number(specificity.energy_margin || 0).toFixed(3))}
      ${statCard('Z-score margin', Number(specificity.z_score_margin || 0).toFixed(3))}
      ${statCard('Worst off-target', humanizePreset(specificity.worst_off_target_preset))}
    </div>
    <div class="compare-grid">
      ${renderCompareEnvironmentCard(target, 'Target')}
      ${offTargets.map((item, idx) => renderCompareEnvironmentCard(item, `Off-target ${idx + 1}`)).join('')}
    </div>
  `;
}

function buildSpecificityCsv(data) {
  const candidates = data.candidates && data.candidates.length ? data.candidates : [data];
  const bestObjective = Number(candidates[0]?.specificity_design?.objective || 0);
  const offTargetOrder = (data.specificity_collection?.off_target_presets || []).filter(Boolean);
  const headers = [
    'rank',
    'sequence',
    'objective',
    'delta_objective_vs_best',
    'target_preset',
    'target_z_score',
    ...offTargetOrder.map((preset) => `${preset}_z_score`),
    'worst_off_target_preset',
    'z_score_margin',
    'energy_margin',
    'lambda_balance',
    'partial',
  ];

  const rows = candidates.map((candidate, idx) => {
    const envMap = Object.fromEntries(
      (candidate.off_target_environments || []).map((item) => [item.environment?.preset, item]),
    );
    return [
      idx + 1,
      candidate.sequence,
      Number(candidate.specificity_design?.objective || 0).toFixed(6),
      formatObjectiveDeltaCsv(candidate.specificity_design?.objective, bestObjective),
      candidate.target_environment?.environment?.preset || '',
      Number(designZ(candidate.target_environment)).toFixed(6),
      ...offTargetOrder.map((preset) => Number(designZ(envMap[preset])).toFixed(6)),
      candidate.specificity_design?.worst_off_target_preset || '',
      Number(candidate.specificity_design?.z_score_margin || 0).toFixed(6),
      Number(candidate.specificity_design?.energy_margin || 0).toFixed(6),
      Number(candidate.specificity_design?.lambda_balance || 0).toFixed(2),
      data.specificity_collection?.partial ? 'true' : 'false',
    ];
  });

  const csvEscape = (value) => {
    const text = String(value ?? '');
    if (/[",\n]/.test(text)) return `"${text.replaceAll('"', '""')}"`;
    return text;
  };

  return [headers, ...rows].map((row) => row.map(csvEscape).join(',')).join('\n');
}

function designZ(scored) {
  return scored?.z_score ?? 0;
}

function formatObjectiveDelta(value, bestObjective) {
  const delta = Number(value || 0) - Number(bestObjective || 0);
  if (Math.abs(delta) < 5e-13) return '0';
  return delta.toExponential(2);
}

function formatObjectiveDeltaCsv(value, bestObjective) {
  const delta = Number(value || 0) - Number(bestObjective || 0);
  if (Math.abs(delta) < 5e-13) return '0';
  return delta.toExponential(6);
}

function bindSpecificityInteractions(root, data) {
  if (!root) return;
  const candidates = data.candidates && data.candidates.length ? data.candidates : [data];
  const detailNode = $('[data-specificity-detail]', root);
  const rows = $$('[data-specificity-row]', root);
  const pickButtons = $$('[data-specificity-pick]', root);
  if (!detailNode || !pickButtons.length) return;

  const selectCandidate = (index) => {
    const candidate = candidates[index];
    if (!candidate) return;
    detailNode.innerHTML = renderSpecificityCandidateDetail(candidate);
    rows.forEach((row) => {
      row.classList.toggle('is-selected', Number(row.dataset.specificityRow) === index);
    });
  };

  pickButtons.forEach((button) => {
    button.addEventListener('click', () => {
      selectCandidate(Number(button.dataset.specificityPick));
    });
  });

  const jsonBtn = $('[data-download-specificity-json]', root);
  if (jsonBtn) jsonBtn.addEventListener('click', () => downloadJson('specificity_result.json', data));
  const downloadBtn = $('[data-download-specificity-csv]', root);
  if (downloadBtn) {
    downloadBtn.addEventListener('click', () => {
      const csv = buildSpecificityCsv(data);
      downloadText('specificity_candidates.csv', csv, 'text/csv;charset=utf-8');
    });
  }
}

function renderSpecificityState(state) {
  const progressPercent = Math.max(0, Math.min(100, Math.round((state.progressFraction || 0) * 100)));
  const annealingDetail = `Restart ${state.restart || 0}/${state.restarts || 0}${state.step != null ? ` · step ${state.step}/${state.steps}` : ''}`;
  const detail = state.phase === 'annealing'
    ? annealingDetail
    : state.phase === 'calibrating'
      ? `Prepared ${state.completed || 0}/${state.total || 0} environment models`
      : state.phase === 'collecting'
        ? `Recovered ${state.accepted || 0}/${state.targetCount || 0} unique candidates${state.attempt ? ` · attempt ${state.attempt}/${state.attemptsTotal || 0}` : ''}${state.restart ? ` · ${annealingDetail}` : ''}`
      : state.phase === 'evaluating'
        ? `Scored ${state.completed || 0}/${state.total || 0} environments`
        : 'Preparing calculation';
  const latestLine = state.bestSequence
    ? `<div class="family-progress-seq">Current best: <code>${escapeHtml(state.bestSequence)}</code>${state.bestObjective != null ? ` · objective ${Number(state.bestObjective).toFixed(3)}` : ''}</div>`
    : state.latestSequence
      ? `<div class="family-progress-seq">Latest sequence: <code>${escapeHtml(state.latestSequence)}</code></div>`
      : '';
  const offTargetsLabel = (state.offTargets || []).length
    ? state.offTargets.map((preset) => humanizePreset(preset)).join(', ')
    : '—';

  return `
      <div class="result-grid">
        ${statCard('Phase', state.phaseLabel || 'Initializing')}
        ${statCard('λ balance', Number(state.lambdaBalance || 0).toFixed(2))}
        ${statCard('Candidates', `${state.accepted || 0}/${state.targetCount || 0}`)}
        ${statCard('Restart budget', state.totalRestartBudget || 0)}
        ${statCard('Target', humanizePreset(state.targetPreset))}
        ${statCard('Off-targets', (state.offTargets || []).length)}
      </div>
      <div class="family-progress specificity-progress">
        <div class="family-progress-head">
          <strong>${escapeHtml(state.note || 'Running specificity design')}</strong>
          <span>${progressPercent}%</span>
        </div>
        <div class="family-progress-bar">
          <div class="family-progress-fill" style="width:${progressPercent}%"></div>
        </div>
        <div class="specificity-progress-detail">${escapeHtml(detail)}</div>
        ${latestLine}
      </div>
      <div class="family-note">Target environment: <strong>${escapeHtml(humanizePreset(state.targetPreset))}</strong>. Off-target set: ${escapeHtml(offTargetsLabel)}.</div>
  `;
}

function partialExportMetadata(state, members) {
  const provenance = state.provenance || members.find((member) => member.provenance)?.provenance;
  return { provenance, request: provenance?.request || state.request };
}

function partialSpecificityPayload(state, reason) {
  const candidates = [...(state.acceptedCandidates || [])]
    .sort(compareSpecificityCandidates)
    .map((candidate, idx) => ({
      ...candidate,
      specificity_design: { ...candidate.specificity_design, rank: idx + 1 },
    }));
  return {
    ...candidates[0],
    ...partialExportMetadata(state, candidates),
    candidates,
    specificity_collection: {
      requested: state.targetCount || candidates.length,
      returned: candidates.length,
      lambda_balance: state.lambdaBalance || 0,
      target_preset: state.targetPreset,
      off_target_presets: state.offTargets || [],
      partial: true,
      partial_reason: reason,
    },
  };
}

function partialFamilyPayload(state, reason) {
  const members = [...state.members].sort((a, b) =>
    Number(a.design_geometry?.energy ?? a.energy) - Number(b.design_geometry?.energy ?? b.energy));
  return {
    ...partialExportMetadata(state, members),
    members,
    family_size_requested: state.target,
    family_size_returned: members.length,
    partial: true,
    partial_reason: reason,
    status_message: `Retained ${members.length} fully evaluated candidates before the calculation was ${reason}.`,
  };
}

function renderOptimalPenetration(data) {
  const best = data.best_score;
  const chargeValue = `${best.model_net_charge}`;
  return `
    <div class="result-card">
      <div class="sequence-banner">${data.sequence}</div>
      <div class="result-grid">
        ${statCard('Best penetration', penetrationLabel(data.best_penetration_percent))}
        ${statCard('Z-score', best.z_score.toFixed(3))}
        ${statCard('Model energy', best.energy.toFixed(3))}
        ${statCard('Sampled depths', data.profile.length)}
      </div>
      ${energyUnitNote()}
      <div class="result-note">Optimal penetration is defined here as the sampled point that yields the most negative Z-score while scanning from homogeneous polar medium (0%) to homogeneous apolar medium (100%), using the selected interfacial preset only at intermediate depths.</div>
      ${penetrationProfilePlot(data)}
      <div class="viz-grid">
        <div>
          ${zScorePlot(best)}
          ${observableGrid(best.observables)}
          ${breakdownGrid(best.breakdown)}
        </div>
        ${wheelSvg(best.wheel, best.environment)}
      </div>
      <div class="result-grid">
        ${statCard('Percentile (lower better)', `${best.percentile_lower_is_better.toFixed(1)}%`)}
        ${statCard('Net charge', chargeValue)}
        ${statCard('Interface', humanizePreset(data.interface_preset))}
        ${statCard('Optimization target', 'Minimum Z-score')}
      </div>
    </div>
  `;
}

function renderFamilyState(state) {
  const note = state.note ? `<div class="family-note">${escapeHtml(state.note)}</div>` : '';
  const progress = `
    <div class="family-progress">
      <div class="family-progress-head">
        <strong>Generating candidates</strong>
        <span>${state.accepted}/${state.target} accepted${state.attempt ? ` after ${state.attempt} attempts` : ''}</span>
      </div>
      <div class="family-progress-bar">
        <div class="family-progress-fill" style="width:${Math.min(100, (state.accepted / Math.max(1, state.target)) * 100)}%"></div>
      </div>
      ${state.latestSequence ? `<div class="family-progress-seq">Latest attempt: <code>${escapeHtml(state.latestSequence)}</code></div>` : ''}
    </div>
  `;
  const members = state.members.length
    ? state.members.map((member, idx) => `
      <div class="family-item">
        <div class="family-item-head">
          <strong>Candidate ${idx + 1}</strong>
          <span>Score: energy ${member.energy.toFixed(3)} | Z ${member.z_score.toFixed(3)}</span>
        </div>
        <div class="family-seq">${member.sequence}</div>
      </div>
    `).join('')
    : `<div class="family-empty">No accepted candidates yet. They will appear here as they are found.</div>`;
  const downloadBtn = state.finalData && state.members.length
    ? `<button class="ghost-btn" type="button" data-download-family>Download JSON</button>`
    : '';
  const stopBtn = state.running
    ? `<button class="ghost-btn danger-ghost" type="button" data-stop-family>Stop and keep accepted candidates</button>`
    : '';
  const partial = state.partialReason
    ? `<div class="result-note partial-note" role="status">Partial result (${escapeHtml(state.partialReason)}): ${state.members.length} of ${state.target} requested candidates.</div>`
    : '';
  return `
    <div class="result-card">
      <div class="toolbar">${downloadBtn}${stopBtn}</div>
      ${partial}
      <div class="result-grid">
        ${statCard('Requested', state.target)}
        ${statCard('Returned', state.accepted)}
        ${statCard('Environment', state.environment ? humanizePreset(state.environment) : '—')}
        ${statCard('Alphabet size', state.alphabetSize || '—')}
      </div>
      ${progress}
      ${note}
      <div class="family-list">${members}</div>
    </div>
  `;
}

function errorHtml(message) {
  return `<div class="error" role="alert">${escapeHtml(message)}</div>`;
}

function setFormBusy(form, busy) {
  const submit = $('button[type="submit"]', form);
  if (submit) submit.disabled = busy;
  form.setAttribute('aria-busy', busy ? 'true' : 'false');
}

function bindForm(formId, resultId, builder, renderer, onReset = null) {
  const form = $(`#${formId}`);
  const result = $(`#${resultId}`);
  let controller = null;
  form.addEventListener('submit', async (event) => {
    event.preventDefault();
    if (controller) return;  // one calculation per form at a time
    const current = new AbortController();
    controller = current;
    setFormBusy(form, true);
    const loading = loadingNode();
    const cancel = document.createElement('button');
    cancel.type = 'button';
    cancel.className = 'ghost-btn';
    cancel.textContent = 'Cancel';
    cancel.addEventListener('click', () => current.abort());
    loading.appendChild(cancel);
    if (onReset) onReset();
    result.innerHTML = '';
    result.appendChild(loading);
    try {
      const data = await builder(new FormData(form), current.signal);
      result.innerHTML = renderer(data);
      if (formId !== 'pdb-form') {
        const download = document.createElement('button');
        download.type = 'button';
        download.className = 'ghost-btn';
        download.textContent = 'Download JSON';
        download.addEventListener('click', () => downloadJson(`${formId.replace(/-form$/, '')}_result.json`, data));
        const toolbar = document.createElement('div');
        toolbar.className = 'toolbar';
        toolbar.appendChild(download);
        result.prepend(toolbar);
      }
    } catch (error) {
      result.innerHTML = error.name === 'AbortError'
        ? '<div class="result-note">Calculation cancelled.</div>'
        : errorHtml(error.message);
    } finally {
      controller = null;
      setFormBusy(form, false);
    }
  });
}

registerVisitCounter();

bindForm('score-form', 'score-result', async (fd, signal) => postJSON('/api/score', {
  ...calculationSettingsFromFormData(fd),
  sequence: fd.get('sequence'),
  residues: parseResidues(fd.get('residues')),
  environment: buildEnvironmentFromFormData(fd, 'preset', 'penetration_depth'),
  n_random: Number(fd.get('n_random') || 500),
}, { signal }), renderScore);

bindForm('design-form', 'design-result', async (fd, signal) => postJSON('/api/design', {
  ...calculationSettingsFromFormData(fd),
  length: Number(fd.get('length') || 20),
  residues: parseResidues(fd.get('residues')),
  environment: buildEnvironmentFromFormData(fd, 'preset', 'penetration_depth'),
  steps: Number(fd.get('steps') || 12000),
  restarts: Number(fd.get('restarts') || 8),
}, { signal }), renderScore);

bindForm('compare-form', 'compare-result', async (fd, signal) => postJSON('/api/compare', {
  ...calculationSettingsFromFormData(fd),
  sequence: fd.get('sequence'),
  residues: parseResidues(fd.get('residues')),
  environment_a: buildEnvironmentFromFormData(fd, 'preset_a', 'penetration_depth_a'),
  environment_b: buildEnvironmentFromFormData(fd, 'preset_b', 'penetration_depth_b'),
}, { signal }), renderCompare);

bindForm('penetration-form', 'penetration-result', async (fd, signal) => postJSON('/api/optimal-penetration', {
  ...calculationSettingsFromFormData(fd),
  sequence: fd.get('sequence'),
  residues: parseResidues(fd.get('residues')),
  environment: { preset: fd.get('preset') },
  n_random: Number(fd.get('n_random') || 400),
  percent_min: 0,
  percent_max: 100,
  percent_step: 5,
}, { signal }), renderOptimalPenetration);

bindForm('cross-form', 'cross-result', async (fd, signal) => postJSON('/api/cross-design', {
  ...calculationSettingsFromFormData(fd),
  length: Number(fd.get('length') || 20),
  residues: parseResidues(fd.get('residues')),
  environment_a: buildEnvironmentFromFormData(fd, 'preset_a', 'penetration_depth_a'),
  environment_b: buildEnvironmentFromFormData(fd, 'preset_b', 'penetration_depth_b'),
  lambda_gap: Number(fd.get('lambda_gap') || 0.5),
  steps: Number(fd.get('steps') || 16000),
  restarts: Number(fd.get('restarts') || 10),
}, { signal }), (data) => renderCompare(data, `
      <div class="result-grid">
        ${statCard('Joint objective', data.cross_design.objective.toFixed(3))}
        ${statCard('λgap', data.cross_design.lambda_gap.toFixed(2))}
      </div>
`));

const specificityForm = $('#specificity-form');
const specificityResult = $('#specificity-result');
let specificityController = null;
specificityForm.addEventListener('submit', async (event) => {
  event.preventDefault();
  if (specificityController) return;  // one run at a time; use Stop to end it
  const controller = new AbortController();
  specificityController = controller;
  setFormBusy(specificityForm, true);
  const fd = new FormData(specificityForm);
  const targetPreset = String(fd.get('target_preset'));
  const sharedPenetration = Number(fd.get('penetration_depth') || 25);
  const selectedOffTargets = fd.getAll('off_target_preset').map((value) => String(value));
  const exactHalfWidth = fd.get('exact_halfwidth_penetration_depth');
  const payload = {
    ...calculationSettingsFromFormData(fd, 10000),
    length: Number(fd.get('length') || 20),
    residues: parseResidues(fd.get('residues')),
    target_environment: buildEnvironmentFromPresetAndPenetration(targetPreset, sharedPenetration, exactHalfWidth),
    off_target_environments: selectedOffTargets.map((preset) => buildEnvironmentFromPresetAndPenetration(preset, sharedPenetration, exactHalfWidth)),
    num_sequences: Number(fd.get('num_sequences') || 10),
    lambda_balance: Number(fd.get('lambda_balance') || 0.0),
    steps: Number(fd.get('steps') || 60000),
    restarts: Number(fd.get('restarts') || 20),
  };

  const state = {
    phase: 'initializing',
    phaseLabel: 'Initializing',
    note: 'Preparing the specificity calculation…',
    progressFraction: 0,
    targetPreset,
    offTargets: selectedOffTargets,
    targetCount: payload.num_sequences,
    accepted: 0,
    attempt: 0,
    attemptsTotal: 0,
    totalRestartBudget: payload.restarts,
    lambdaBalance: payload.lambda_balance,
    restart: 0,
    restarts: 0,
    step: 0,
    steps: payload.steps,
    latestSequence: '',
    bestSequence: '',
    bestObjective: null,
    completed: 0,
    total: 0,
    acceptedCandidates: [],
    request: payload,
    provenance: null,
  };

  // The Stop button stays in place (and keeps keyboard focus); only the progress block is redrawn.
  specificityResult.innerHTML = `
    <div class="result-card">
      <div class="toolbar">
        <button class="ghost-btn danger-ghost" type="button" data-stop-specificity>Stop and keep partial results</button>
      </div>
      <div data-specificity-state role="status"></div>
    </div>
  `;
  $('[data-stop-specificity]', specificityResult).addEventListener('click', () => controller.abort(), { once: true });
  const stateNode = $('[data-specificity-state]', specificityResult);
  const refresh = () => {
    stateNode.innerHTML = renderSpecificityState(state);
  };
  const showPartial = (reason, message) => {
    const partial = partialSpecificityPayload(state, reason);
    const notice = message ? errorHtml(message) : '';
    if (!partial.candidates.length) {
      specificityResult.innerHTML = notice || '<div class="result-note">Specificity run stopped before any candidate had been fully evaluated.</div>';
      return;
    }
    specificityResult.innerHTML = notice + renderSpecificity(partial);
    bindSpecificityInteractions(specificityResult, partial);
  };
  let finished = false;

  refresh();

  try {
    await postNDJSON('/api/specificity-design-stream', payload, (message) => {
      state.provenance = message.payload?.provenance || state.provenance || message.payload?.member?.provenance || null;
      if (message.type === 'meta') {
        state.phase = 'initializing';
        state.phaseLabel = 'Initializing';
        state.note = 'Building target and off-target models…';
        state.targetCount = Number(message.payload.num_sequences || state.targetCount);
        state.totalRestartBudget = Number(message.payload.total_restart_budget || state.totalRestartBudget);
        state.restarts = Number(message.payload.restarts_per_attempt || message.payload.restarts || state.restarts);
      } else if (message.type === 'phase') {
        state.phase = message.payload.phase;
        state.progressFraction = Number(message.payload.progress_fraction || 0);
        state.completed = Number(message.payload.completed || 0);
        state.total = Number(message.payload.total || 0);
        if (message.payload.phase === 'calibrating') {
          state.phaseLabel = 'Calibrating';
          state.note = 'Preparing the target and off-target background models…';
        } else if (message.payload.phase === 'collecting') {
          state.phaseLabel = 'Collecting candidates';
          state.note = 'Running repeated specificity searches to recover several unique sequences…';
          state.accepted = Number(message.payload.completed || state.accepted);
          state.attempt = Number(message.payload.attempt || state.attempt);
          state.attemptsTotal = Number(message.payload.attempts_total || state.attemptsTotal);
        } else if (message.payload.phase === 'evaluating') {
          state.phaseLabel = 'Evaluating';
          state.note = 'Scoring the final candidate against the target and all off-targets…';
        }
      } else if (message.type === 'candidate') {
        state.phase = 'collecting';
        state.phaseLabel = 'Collecting candidates';
        state.note = 'Accepted a new unique specificity candidate.';
        state.accepted = Number(message.payload.accepted || state.accepted);
        state.targetCount = Number(message.payload.target || state.targetCount);
        state.attempt = Number(message.payload.attempt || state.attempt);
        if (message.payload.member) {
          state.acceptedCandidates.push(message.payload.member);
        }
      } else if (message.type === 'progress') {
        state.phase = message.payload.phase || 'annealing';
        state.phaseLabel = state.phase === 'collecting' ? 'Collecting candidates' : 'Searching';
        state.note = state.phase === 'collecting'
          ? 'Running repeated specificity searches to recover several unique sequences…'
          : 'Optimizing the sequence with simulated annealing…';
        state.progressFraction = Number(message.payload.progress_fraction || 0);
        state.restart = Number(message.payload.restart || 0);
        state.restarts = Number(message.payload.restarts || state.restarts);
        state.step = Number(message.payload.step || 0);
        state.steps = Number(message.payload.steps || state.steps);
        state.accepted = Number(message.payload.accepted || state.accepted);
        state.targetCount = Number(message.payload.target_count || state.targetCount);
        state.attempt = Number(message.payload.attempt || state.attempt);
        state.attemptsTotal = Number(message.payload.attempts_total || state.attemptsTotal);
        state.latestSequence = message.payload.latest_sequence || '';
        state.bestSequence = message.payload.best_sequence || '';
        state.bestObjective = message.payload.best_objective ?? null;
      } else if (message.type === 'final') {
        finished = true;
        specificityResult.innerHTML = renderSpecificity(message.payload);
        bindSpecificityInteractions(specificityResult, message.payload);
        refreshUsageCounters().catch(() => {});
        return;
      } else if (message.type === 'error') {
        throw new Error(message.detail || 'Specificity design failed');
      }
      refresh();
    }, { signal: controller.signal });
    if (!finished) throw new Error('The calculation ended without a final result.');
  } catch (error) {
    if (finished) return;
    if (error.name === 'AbortError') {
      showPartial('stopped by the user');
    } else {
      showPartial('interrupted by an error', error.message);
    }
  } finally {
    specificityController = null;
    setFormBusy(specificityForm, false);
  }
});

bindForm('pdb-form', 'pdb-result', async (fd, signal) => postJSON('/api/pdb', {
  sequence: fd.get('sequence'),
}, { signal }), (data) => {
  const viewerId = nextViewerId();
  setTimeout(() => {
    const downloadBtn = document.querySelector('[data-download-pdb]');
    if (downloadBtn) {
      downloadBtn.onclick = () => downloadText(`${data.sequence}.pdb`, data.pdb, 'chemical/x-pdb');
    }
    mountPdbViewer(viewerId, data.pdb);
  }, 0);
  return `
    <div class="result-card">
      <div class="sequence-banner">${data.sequence}</div>
      <div class="toolbar">
        <button class="ghost-btn" type="button" data-download-pdb>Download PDB</button>
      </div>
      <div class="pdb-grid">
        <div class="viewer-card">
          <div class="viewer-head">
            <strong>3D structure</strong>
            <span>Ideal alpha helix</span>
          </div>
          <div class="pdb-viewer" id="${viewerId}"></div>
        </div>
        <div>
          <pre class="pdb">${escapeHtml(data.pdb)}</pre>
        </div>
      </div>
    </div>
  `;
}, releasePdbViewer);

const familyForm = $('#family-form');
const familyResult = $('#family-result');
let familyController = null;
familyForm.addEventListener('submit', async (event) => {
  event.preventDefault();
  if (familyController) return;  // one run at a time
  const controller = new AbortController();
  familyController = controller;
  setFormBusy(familyForm, true);
  const fd = new FormData(familyForm);
  const payload = {
    ...calculationSettingsFromFormData(fd),
    length: Number(fd.get('length') || 20),
    residues: parseResidues(fd.get('residues')),
    environment: buildEnvironmentFromFormData(fd, 'preset', 'penetration_depth'),
    family_size: Number(fd.get('family_size') || 6),
    steps: Number(fd.get('steps') || 10000),
    restarts: Number(fd.get('restarts') || 6),
  };

  const state = {
    target: payload.family_size,
    accepted: 0,
    attempt: 0,
    latestSequence: '',
    members: [],
    environment: payload.environment.preset,
    alphabetSize: payload.residues.length,
    note: 'Starting family generation…',
    finalData: null,
    running: true,
    partialReason: '',
    request: payload,
    provenance: null,
  };

  const refresh = () => {
    const hadFocus = document.activeElement?.matches?.('[data-stop-family]');
    familyResult.innerHTML = renderFamilyState(state);
    const btn = familyResult.querySelector('[data-download-family]');
    if (btn && state.finalData) {
      btn.onclick = () => downloadJson('helix_family.json', state.finalData);
    }
    const stop = familyResult.querySelector('[data-stop-family]');
    if (stop) {
      stop.addEventListener('click', () => controller.abort(), { once: true });
      if (hadFocus) stop.focus();
    }
  };

  refresh();

  try {
    await postNDJSON('/api/design-family-stream', payload, (message) => {
      state.provenance = message.payload?.provenance || state.provenance || message.payload?.member?.provenance || null;
      if (message.type === 'meta') {
        state.note = `Searching for ${state.target} distinct low-energy candidates. Minimum enforced Hamming diversity: ${message.payload.min_distance}.`;
        state.environment = message.payload.environment.preset;
      } else if (message.type === 'progress') {
        state.attempt = message.payload.attempt;
        state.accepted = message.payload.accepted;
        state.latestSequence = message.payload.latest_sequence || '';
      } else if (message.type === 'candidate') {
        state.attempt = message.payload.attempt;
        state.accepted = message.payload.accepted;
        state.members.push(message.payload.member);
        state.latestSequence = message.payload.member.sequence;
        state.note = message.payload.min_hamming_to_previous == null
          ? 'Accepted first family member.'
          : `Accepted candidate ${state.accepted} with minimum Hamming distance ${message.payload.min_hamming_to_previous} from previous members.`;
      } else if (message.type === 'final') {
        state.finalData = message.payload;
        state.members = message.payload.members;
        state.accepted = message.payload.family_size_returned;
        state.note = message.payload.status_message;
      } else if (message.type === 'error') {
        throw new Error(message.detail || 'Family generation failed');
      }
      refresh();
    }, { signal: controller.signal });
    if (!state.finalData) throw new Error('The calculation ended without a final result.');
    state.running = false;
    refresh();
  } catch (error) {
    state.running = false;
    if (state.finalData) {
      refresh();
      return;
    }
    const aborted = error.name === 'AbortError';
    state.partialReason = aborted ? 'stopped by the user' : 'interrupted by an error';
    if (state.members.length) {
      state.finalData = partialFamilyPayload(state, state.partialReason);
      state.members = state.finalData.members;
      state.accepted = state.finalData.family_size_returned;
      state.note = state.finalData.status_message;
      refresh();
      if (!aborted) familyResult.insertAdjacentHTML('afterbegin', errorHtml(error.message));
    } else {
      familyResult.innerHTML = aborted ? '<div class="result-note">Family generation stopped.</div>' : errorHtml(error.message);
    }
  } finally {
    familyController = null;
    setFormBusy(familyForm, false);
  }
});
```
