## Additional file 1. Evaluated software source and dependencies for "Helix Design Studio: designing antimicrobial peptide candidates for membrane environments": home-visual.js

```
/* Homepage illustration only. Environment previews do not change tool inputs. */
(() => {
  const visual = document.querySelector('.workflow-visual');
  if (!visual) return;
  const captions = {
    water: ['An assumed alpha-helix in a water-like medium', 'A water-like, polar environment around the helix.'],
    membrane: ['An assumed alpha-helix at a membrane-like interface', 'An interface between water-like and non-polar regions.'],
    apolar: ['An assumed alpha-helix in a non-polar medium', 'A non-polar environment around the helix.'],
  };
  const buttons = [...visual.querySelectorAll('[data-preview-environment]')];
  buttons.forEach((button) => button.addEventListener('click', () => {
    const choice = button.dataset.previewEnvironment;
    visual.dataset.environment = choice;
    buttons.forEach((item) => item.setAttribute('aria-pressed', String(item === button)));
    visual.querySelector('#environment-scene-title').textContent = captions[choice][0];
    visual.querySelector('#environment-scene-desc').textContent = `${captions[choice][1]} This illustrates the environment model, not a predicted structure or insertion depth.`;
    visual.querySelector('.environment-preview-caption').textContent = captions[choice][1];
  }));
})();
```
