## Additional file 1. Evaluated software source and dependencies for "Helix Design Studio: designing antimicrobial peptide candidates for membrane environments": index.html


Skip to workspace


☰


Helix
DESIGN STUDIO


Peptide design & analysis

By SIMBIOS
↗


Behind the model


WORKSPACE

Overview

ANALYZE A SEQUENCE

Check a sequence


Compare environments


Explore exposure

DESIGN NEW SEQUENCES

Design a peptide


Create a family


Design for selectivity


Design for two environments

STRUCTURE

Build a 3D helix

Behind the model


About the project

Open science. Open possibilities.

PEPTIDE DESIGN & ANALYSIS

### Explore and design *helical peptides.*

See how a sequence of amino acids fits different environments — or design a new one.

One workspace. Two ways to explore peptides.

How it works

#### 01 Your starting point

ANALYZE


W

F

E

K

W

E
 **A sequence you have** 

or

DESIGN


 *↔* 
 **A sequence you need** 

Set length & amino acids

#### 02 Choose its surroundings

Water-like


Membrane


Non-polar

An assumed alpha-helix at a membrane-like interface

A schematic of a single helical peptide with water-like and non-polar surroundings. This illustrates the environment model, not a predicted structure or insertion depth.


Water-like medium


Water-like

Non-polar


Non-polar medium


Assumed α-helix

An interface between water-like and non-polar regions.

#### 03 Explore the results

ANALYZE
 **Score & compare** 

How well does it fit?

DESIGN
 **New candidates** 

One sequence or a family

Export results & an ideal 3D helix

Schematic overview · Assumes an α-helix · Model-based compatibility

#### What would you like to do?

Start here. No setup needed.

I HAVE A SEQUENCE
 **Check a sequence** 

Find out how well your peptide fits a chosen environment.

Analyze a sequence


I WANT A NEW SEQUENCE
 **Design a peptide** 

Choose a length and an environment. Explore a new candidate.

Create a sequence

#### Take your exploration further

All tools, one workspace

**Compare environments** 

One sequence. Two environments.

↗


 **Create a family** 

Find several different candidates.

↗


 **Explore exposure** 

Scan from water-like to non-polar.

↗


 **Design for selectivity** 

Favor one environment over others.

↗


 **Design for two environments** 

Find a balance between two settings.

↗


 **Build a 3D helix** 

View and export an ideal structure.

↗

**A model for exploration.** 
Results describe compatibility with an assumed α-helix, not a prediction of folding or biological activity.

How it works
↗

THE SOFTWARE & THE SCIENCE

#### About Helix Design Studio

Explore how a peptide sequence fits different environments, or design new candidates for the environment you choose.

- 01 Score & compare sequences
- 02 Design candidates & families
- 03 Export ideal 3D helices

The model assumes an α-helix and measures compatibility. It does not predict folding or biological activity.

THE SCIENCE BEHIND HELIX

##### Environment-Conditioned Design of α-Helical Peptides

Read the scientific method behind sequence scoring, environment comparisons and peptide design.

Daniel Conde-Torres · Rebeca García-Fandiño · Ángel Piñeiro

Journal of Chemical Theory and Computation  
2026 · Volume 22 · Pages 9776–9789

Read the paper ↗
DOI: 10.1021/acs.jctc.6c01278

DEVELOPED BY

##### SIMBIOS

Simulating life. Atom by atom.

Computational biophysics research on molecular simulation, membrane-active systems, biomolecular recognition and data-driven analysis.

Universidade de  
Santiago de Compostela

Meet the research group ↗

##### Acknowledgements

We gratefully acknowledge the support of the following research projects.

###### Consolidación e estruturación 2025 · GRC GI-2132

Sistemas Supramoleculares, Nanobiomimética e Biofísica Molecular – SupraNanoBioMol.

Xunta de Galicia · ED431C 2025/15

View project details ↗

###### Generación de Conocimiento 2022

Proyectos de investigación orientada

Descifrando la conexión del código lipídico entre cáncer, infección y envejecimiento: hacia herramientas teranósticas no convencionales y vacunas basadas en la memoria innata.

Project PID2022-141534OB-I00 funded by MICIU/AEI/10.13039/501100011033 and by ERDF, EU.

View project details ↗

A LITTLE GUIDANCE

#### Behind the model

Understand the model, explore the concepts and interpret your results. Open any topic for more detail.

01

##### Choose a sequence

Use your own, try a reference, or design a new one.

02

##### Choose its surroundings

Water-like, non-polar, or a charged or neutral membrane interface.

03

##### Explore compatibility

A more negative Z-score is better relative to the random reference.

What the app does

Use
 **Check a sequence** 
to evaluate one sequence,
 **Design a peptide** 
to search for a low-energy sequence in one environment,
 **Create a family** 
to generate several diverse candidates,
 **Compare environments** 
to inspect the same sequence in two environments,
 **Design for selectivity** 
to design one-vs-rest selective candidates, and
 **Design for two environments** 
to search for trade-offs between environments.


How to read the results

**Lower model energy** 
means better compatibility with the chosen helical state within the model. These energies are reported in
 **dimensionless Hamiltonian units** 
, not in physical units such as kcal/mol, so they should be interpreted comparatively within the same scoring setup. A more negative
 **Z-score** 
means the sequence is better than the random-sequence background.
 **Percentile** 
tells you where the sequence sits relative to that random ensemble, with lower being better.
 **Net charge** 
follows the current model convention; in this version histidine is treated as neutral in the charge-dependent terms.


What environments mean

**Homogeneous polar** 
approximates a water-like medium.
 **Homogeneous apolar** 
approximates a non-polar medium.
 **Interfacial** 
modes impose a membrane-facing and water-facing sector on the helical wheel, plus an anionic, neutral or cationic membrane surface charge. In the app,
 **penetration depth** 
is a percentage with the same meaning in every panel: 0% is the homogeneous polar medium, 100% the homogeneous apolar medium, and intermediate values use the interfacial preset with an apolar sector half-width of 40° + 1.1° per percent. It is an angular exposure setting, not a physical insertion depth. In interfacial environments, scoring and design both align each sequence by its own hydrophobic moment toward the apolar sector. Calibration and random reference sequences follow the same rule. If the hydrophobic moment vanishes, the preset wheel phase is used. Orientation has no effect in homogeneous media.

In the workspace, “Non-polar exposure” and “Explore exposure” refer to the original penetration-depth setting: an angular exposure coordinate, not a physical insertion depth. Interface labels indicate membrane surface charge.


What λgap means

**λgap** 
is only used in
 **Design for two environments** 
. It penalizes sequences that behave very differently in the two chosen environments. Small values favor the best overall compromise; larger values push the search toward more balanced cross-environment compatibility.


Hamiltonian terms

H = λ
 _env_ 
H
 _env_ 
+ λ
 _chg_ 
H
 _chg_ 
+ λ
 _local_ 
H
 _local_ 
+ λ
 _pair_ 
H
 _pair_ 
+ λ
 _elec_ 
H
 _elec_ 
+ λ
 _neigh_ 
H
 _neigh_

**Environment** 
rewards residues that match the target medium.
 **Charge** 
captures residue preference with membrane surface charge.
 **Local helix** 
and
 **Helix neighbors** 
reward helical propensity.
 **Pairwise** 
adds sequence-dependent interaction preferences.
 **Electrostatics** 
captures attractive and repulsive charge interactions along the helix. The transverse hydrophobic moment is a geometric descriptor displayed on the helical wheel and used for interfacial alignment. It is not an energy term. Histidine contributes through the hydrophobicity and helical-propensity terms and is neutral in the charge-dependent terms.


API and programmatic examples

Use the HTTP API to run scoring, sequence design, environment comparisons and exposure scans from scripts. Open
interactive API documentation
, download the
OpenAPI schema
, or follow the
API guide with curl and Python examples
. Check the serving version in the schema and retain each result's provenance when reporting calculations.


Practical limitations

The model scores compatibility with a predefined helical scaffold. It is best interpreted as a relative design and ranking tool. Experimental activity, folding kinetics, insertion depth and membrane disruption require additional validation.

ANALYZE

#### Check a sequence

How well does your peptide fit its environment?

01
Your sequence

Try a reference peptide

Custom sequence

Choose an example to fill the sequence below.

Amino-acid sequence
WFEKWEKEWKEFWKEWKDFW

An example is ready. Replace it with your own one-letter amino-acid sequence (up to 40 residues).


02
Choose an environment

Environment

Water-like · polar

Non-polar · apolar

Membrane interface · negative

Membrane interface · neutral

Membrane interface · positive

Non-polar exposure

 **50%** 
· Angular exposure, not physical insertion depth.


Water-like · 0%

Non-polar · 100%


Advanced settings

Optional · defaults are ready to use

Allowed residues

Random reference size

Number of random sequences used to estimate the background mean and standard deviation for the Z-score. Web limit: 2,000.

Check sequence →


WHAT YOU’LL GET

##### Understand your sequence

A compatibility score, a comparison with random sequences, and a helical wheel.

Scores describe an assumed α-helix. Lower model energy means better compatibility within the same scoring setup.

Understand the model

**Your results will appear here** 

Choose your settings above, then start the calculation.

ANALYZE

#### Explore exposure

Explore the balance between water-facing and non-polar-facing parts of your helix.

01
Your sequence

Try a reference peptide

Custom sequence

Choose an example to fill the sequence below.

Amino-acid sequence
WFEKWEKEWKEFWKEWKDFW

An example is ready. Replace it with your own one-letter amino-acid sequence (up to 40 residues).


02
Choose an environment

Environment

Membrane interface · negative

Membrane interface · neutral

Membrane interface · positive

**0%
⟶
100%** 

21 exposure settings, from water-like to non-polar.


Advanced settings

Optional · defaults are ready to use

Allowed residues

Random reference size

Background size used for each sampled penetration. Web limit: 2,000.

Scan exposure →


WHAT YOU’LL GET

##### Explore exposure

A profile across 21 exposure settings, with the lowest Z-score highlighted.

Scores describe an assumed α-helix. Lower model energy means better compatibility within the same scoring setup.

Understand the model

**Your results will appear here** 

Choose your settings above, then start the calculation.

DESIGN

#### Design a peptide

Choose a length and an environment. Find a sequence that fits.

01
Set your peptide

Peptide length

Number of amino acids, from 4 to 40.


02
Choose an environment

Environment

Water-like · polar

Non-polar · apolar

Membrane interface · negative

Membrane interface · neutral

Membrane interface · positive

Non-polar exposure

 **50%** 
· Angular exposure, not physical insertion depth.


Water-like · 0%

Non-polar · 100%


Advanced settings

Optional · defaults are ready to use

Allowed residues

Optimization steps (classical simulated annealing)

Number of Monte Carlo mutation steps used during the simulated-annealing search. Higher values let the optimizer test more sequence changes, usually improving exploration at the cost of longer runtime. Web limit: 50,000.

Restarts

Independent annealing runs from different random starting points. Web limit: 20.

Need heavier searches or custom large-scale runs? Please contact
SIMBIOS
.

Design sequence →


WHAT YOU’LL GET

##### Design a peptide

One candidate sequence, its compatibility score, and a helical wheel.

Scores describe an assumed α-helix. Lower model energy means better compatibility within the same scoring setup.

Understand the model

**Your results will appear here** 

Choose your settings above, then start the calculation.

DESIGN

#### Create a family

Explore several different sequences for the same environment.

01
Set your peptide

Peptide length

Number of amino acids, from 4 to 40.

Number of candidates

Request 2–12 unique candidates. The search may return fewer.


02
Choose an environment

Environment

Water-like · polar

Non-polar · apolar

Membrane interface · negative

Membrane interface · neutral

Membrane interface · positive

Non-polar exposure

 **50%** 
· Angular exposure, not physical insertion depth.


Water-like · 0%

Non-polar · 100%


Advanced settings

Optional · defaults are ready to use

Allowed residues

Optimization steps (classical simulated annealing)

Number of Monte Carlo mutation steps used in each annealing run. More steps generally help each candidate settle into a better low-energy sequence, but they also make family generation slower. Web limit: 20,000.

Restarts per candidate

More restarts improve robustness but increase runtime. Web limit: 20.

Need larger families or heavier searches? Please contact
SIMBIOS
.

Generate family →


WHAT YOU’LL GET

##### Create a family

A set of unique candidates, each with scores and a helical wheel.

Scores describe an assumed α-helix. Lower model energy means better compatibility within the same scoring setup.

Understand the model

**Your results will appear here** 

Choose your settings above, then start the calculation.

ANALYZE

#### Compare environments

One sequence, two environments. See how its scores change.

01
Your sequence

Try a reference peptide

Custom sequence

Choose an example to fill the sequence below.

Amino-acid sequence
WFEKWEKEWKEFWKEWKDFW

An example is ready. Replace it with your own one-letter amino-acid sequence (up to 40 residues).


02
Choose environments

Environment A

Water-like · polar

Non-polar · apolar

Membrane interface · negative

Membrane interface · neutral

Membrane interface · positive

Environment B

Water-like · polar

Non-polar · apolar

Membrane interface · negative

Membrane interface · neutral

Membrane interface · positive

Non-polar exposure A

 **50%** 
· Angular exposure, not physical insertion depth.


Water-like · 0%

Non-polar · 100%

Non-polar exposure B

 **50%** 
· Angular exposure, not physical insertion depth.


Water-like · 0%

Non-polar · 100%


Advanced settings

Optional · defaults are ready to use

Allowed residues

Compare environments →


WHAT YOU’LL GET

##### Compare environments

Side-by-side scores and helical wheels for the same sequence.

Scores describe an assumed α-helix. Lower model energy means better compatibility within the same scoring setup.

Understand the model

**Your results will appear here** 

Choose your settings above, then start the calculation.

DESIGN

#### Design for two environments

Find a sequence with balanced compatibility across two environments.

01
Set your peptide

Peptide length

Number of amino acids, from 4 to 40.


02
Choose environments

Environment A

Water-like · polar

Non-polar · apolar

Membrane interface · negative

Membrane interface · neutral

Membrane interface · positive

Environment B

Water-like · polar

Non-polar · apolar

Membrane interface · negative

Membrane interface · neutral

Membrane interface · positive

Non-polar exposure A

 **50%** 
· Angular exposure, not physical insertion depth.


Water-like · 0%

Non-polar · 100%

Non-polar exposure B

 **50%** 
· Angular exposure, not physical insertion depth.


Water-like · 0%

Non-polar · 100%

Balance between environments (λgap)

Penalty applied when the same sequence behaves too differently in the two environments. Increase it to favor more balanced solutions. Web limit: 10.


Advanced settings

Optional · defaults are ready to use

Allowed residues

Optimization steps (classical simulated annealing)

Number of Monte Carlo mutation steps in the joint simulated-annealing search. Larger values give the optimizer more chances to improve the sequence across both environments, but increase runtime. Web limit: 60,000.

Restarts

Independent optimization runs from different starting points. Web limit: 20.

For heavier cross-environment searches or custom campaigns, please contact
SIMBIOS
.

Design for both environments →


WHAT YOU’LL GET

##### Design for two environments

One candidate with a score and helical wheel for each environment.

Scores describe an assumed α-helix. Lower model energy means better compatibility within the same scoring setup.

Understand the model

**Your results will appear here** 

Choose your settings above, then start the calculation.

DESIGN

#### Design for selectivity

Find sequences that favor your target environment over competing environments.

01
Set your peptide

Peptide length

Number of amino acids, from 4 to 40.

Number of candidates

Request 1–10 unique candidates.


02
Choose environments

Target environment

Water-like · polar

Non-polar · apolar

Membrane interface · negative

Membrane interface · neutral

Membrane interface · positive

Non-polar exposure

 **25%** 
· Angular exposure, not physical insertion depth.


Water-like · 0%

Non-polar · 100%


Environments to compare against

Membrane interface · neutral


Membrane interface · positive


Water-like · polar


Non-polar · apolar


Membrane interface · negative

How is selectivity calculated?

The specificity target is
 **L
 _λ_ 
= λH
 _target_ 
+ (1-λ)(H
 _target_ 
- min(H
 _off-targets_ 
))** 
. Lower values are better. Use
 **λ = 0** 
for the hard one-vs-rest limit, and larger λ values when you want to give progressively more weight to absolute target favorability.


Selectivity balance (λ)

 **0.00** 
· Lower favors selectivity; higher favors target compatibility.


Advanced settings

Optional · defaults are ready to use

Allowed residues

Optimization steps (classical simulated annealing)

Number of Monte Carlo mutation steps in the one-vs-rest simulated-annealing search. More steps increase the search budget and execution time. Web limit: 60,000.

Total restart budget

Total number of annealing restarts allocated to the whole run. This budget is distributed across the requested candidate sequences, so asking for more candidates gives fewer restarts per candidate. Web limit: 20.

This mode is useful for designing candidates that prefer one target state against several competing media, for example an anionic membrane against neutral/cationic interfaces plus homogeneous polar and apolar media.

Design selective sequences →


WHAT YOU’LL GET

##### Design for selectivity

Ranked candidates, detailed comparisons, and CSV or JSON exports.

Scores describe an assumed α-helix. Lower model energy means better compatibility within the same scoring setup.

Understand the model

**Your results will appear here** 

Choose your settings above, then start the calculation.

STRUCTURE

#### Build a 3D helix

Turn your sequence into an ideal α-helical model.

01
Your sequence

Amino-acid sequence
WFEKWEKEWKEFWKEWKDFW

An example is ready. Replace it with your own one-letter amino-acid sequence (up to 40 residues).

Build 3D helix →


WHAT YOU’LL GET

##### Build a 3D helix

An interactive 3D model and a downloadable PDB file.

This is an ideal structure, not a folding prediction.

Understand the model

**Your results will appear here** 

Build your helix to explore it in 3D.


Developed by

**SIMBIOS**Simulating life. Atom by atom.


Universidade de  
Santiago de Compostela

© 2026 Helix Design Studio contributors. Original code: MIT license.

Visits: —

Unique visitors: —

Runs: —

Anonymous usage statistics are collected; a random identifier is stored in this browser.


Calculating your results…
