## Additional file 1. Evaluated software source and dependencies for "Helix Design Studio: designing antimicrobial peptide candidates for membrane environments": interface.js

```
/* Presentation only: navigation, disclosure and accessible result affordances.
 * Scientific requests, defaults, rendering and exports remain in app.js.
 */
(() => {
  'use strict';
  const navigation = document.getElementById('workspace-navigation');
  const menu = document.querySelector('.mobile-menu');
  const navButtons = [...document.querySelectorAll('.nav-btn')];
  let restoringHistory = false;
  const closeMenu = () => {
    navigation.classList.remove('is-open');
    menu.setAttribute('aria-expanded', 'false');
    menu.setAttribute('aria-label', 'Open navigation');
  };
  const openPanel = (name) => {
    const button = navButtons.find((item) => item.dataset.panel === name);
    if (button) button.click();
  };

  menu.addEventListener('click', () => {
    const expanded = navigation.classList.toggle('is-open');
    menu.setAttribute('aria-expanded', String(expanded));
    menu.setAttribute('aria-label', expanded ? 'Close navigation' : 'Open navigation');
  });
  document.addEventListener('keydown', (event) => {
    if (event.key === 'Escape' && navigation.classList.contains('is-open')) {
      closeMenu();
      menu.focus();
    }
  });
  document.addEventListener('click', (event) => {
    if (!navigation.contains(event.target) && !menu.contains(event.target)) closeMenu();
  });
  document.querySelectorAll('[data-open-panel]').forEach((button) => {
    button.addEventListener('click', () => openPanel(button.dataset.openPanel));
  });
  navButtons.forEach((button) => {
    button.addEventListener('click', () => {
      closeMenu();
      const panel = document.getElementById(`panel-${button.dataset.panel}`);
      const heading = panel.querySelector('h1, h2');
      if (heading) {
        heading.setAttribute('tabindex', '-1');
        heading.focus({ preventScroll: true });
      }
      window.scrollTo({ top: 0, behavior: 'instant' });
      const hash = `#${button.dataset.panel}`;
      if (!restoringHistory && location.hash !== hash) history.pushState(null, '', hash);
    });
  });
  window.addEventListener('popstate', () => {
    restoringHistory = true;
    try { openPanel(location.hash.slice(1) || 'home'); }
    finally { restoringHistory = false; }
  });
  if (location.hash) openPanel(location.hash.slice(1));

  // Closed advanced settings still submit normally; reveal them for native validation.
  document.querySelectorAll('form').forEach((form) => {
    form.addEventListener('invalid', (event) => {
      const disclosure = event.target.closest('details');
      if (disclosure) disclosure.open = true;
    }, true);
    form.addEventListener('submit', () => {
      const result = document.getElementById(form.id.replace('-form', '-result'));
      requestAnimationFrame(() => {
        if (form.closest('.panel').classList.contains('active') && result.firstChild) {
          result.scrollIntoView({ behavior: 'smooth', block: 'start' });
        }
      });
    });
  });

  const disclose = (node, title) => {
    if (node.parentElement.classList.contains('result-details')) return;
    const details = document.createElement('details');
    details.className = 'result-details';
    const summary = document.createElement('summary');
    summary.textContent = title;
    node.before(details);
    details.append(summary, node);
  };
  document.querySelectorAll('.result').forEach((result) => {
    const options = { childList: true, subtree: true };
    const enhance = () => {
      observer.disconnect();
      result.querySelectorAll('.settings-line').forEach((node) => disclose(node, 'Calculation settings & reproducibility'));
      result.querySelectorAll('.breakdown-section').forEach((node) => disclose(node, 'Explore the energy contributions'));
      result.querySelectorAll('pre.pdb').forEach((node) => disclose(node, 'View PDB file contents'));
      result.querySelectorAll('.stat[title]').forEach((node) => {
        if (node.querySelector('.stat-help')) return;
        const detail = document.createElement('details');
        detail.className = 'stat-help';
        const summary = document.createElement('summary');
        summary.textContent = 'What does this mean?';
        const explanation = document.createElement('p');
        explanation.textContent = node.title;
        detail.append(summary, explanation);
        node.append(detail);
      });
      observer.observe(result, options);
    };
    const observer = new MutationObserver(enhance);
    enhance();
  });
})();
```
