## Additional file 1. Evaluated software source and dependencies for "Helix Design Studio: designing antimicrobial peptide candidates for membrane environments": local-entry.js

```
// Direct file previews need the local app server for assets and calculations.
// The relative script URL works from both app/static/index.html and the site root.
if (window.location.protocol === 'file:') {
  window.location.replace(
    'http://127.0.0.1:8000/' + window.location.search + window.location.hash
  );
}
```
