## Additional file 1. Evaluated software source and dependencies for "Helix Design Studio: designing antimicrobial peptide candidates for membrane environments": CHANGELOG.md

# Changelog

Use `v0.1.0` for the original comparison baseline, `v1.0.0rc2` for the source-corrected parameter tables, `v1.0.0rc4` for the last release with fixed-phase search objectives, `v1.0.0rc5` for aligned searches with positive M1, and `v1.0.0rc6` for aligned searches with the corrected negative M1 contribution. Each result reports the version, the parameter-set checksum, the complete request and the seed protocol under `provenance`; `GET /api/health` reports the running version. Results from different versions are not interchangeable.

## 1.0.0rc6 — 2026-10-17

### Helical pairing propensity in the minimized score

- The raw neighbor score is now `−mean(M1) + mean(M3) + mean(M4)`. Nacar (2020), Figure 3, defines M1 as observed/expected helical-pair enrichment: larger values indicate greater enrichment. Its negative contribution now rewards that enrichment in the minimized score. Earlier releases added M1 positively, opposing this intended contribution.
- The 400 M1 source values, their axis mapping and their empirical reuse at sequence separation i,i+1 are preserved. The source itself measures hydrogen-bonded n,n+4 pairs in globular proteins. A source zero contributes zero to the M1 sum. The M3/M4 coefficients and their negative-is-favorable energy convention are unchanged.
- The three separation means retain fixed internal coefficients −1, +1 and +1 and are standardized as one component. This combination is an empirical descriptor with fixed internal scaling across heterogeneous source quantities; it does not constitute a conversion of M1 into thermodynamic energies. Other component equations and configured weights are unchanged.
- Calibration-cache format advances to 5. The explicit convention `negative_m1_plus_m3_plus_m4` and internal coefficients enter cache metadata, matrix fingerprints and the public parameter-set checksum. Raw table checksums are unchanged, while the parameter-set checksum distinguishes the scoring change. Cached calibrations from the positive-M1 convention are not reused.
- Sequence scores, random-reference distributions, search results and derived rankings must be recalculated for this release. A fixed seed alone does not preserve search trajectories when the scoring convention changes. Historical comparison files remain associated with their recorded release and inputs.
- Regression checks cover the direction of the M1 reward, source zeros, the preserved M3/M4 signs, calibration-cache separation and exact agreement with all published matrix entries.
- The Table 1 rerun tool identifies the current formula and optional positive-M1 control separately, with per-variant provenance. Its optional negative-log control adds `−ln(M1)` with the correct sign and explicitly rejects source zeros without pseudocounts. Archived numerical JSON reports are preserved under their original release conventions.

- Browser Advanced settings now expose calibration count, seed and an exact interfacial half-width, preserving previous defaults. These controls allow the documented calculation protocols to be entered directly.
- Specificity progress events now pair the displayed objective with the selected candidate; final selection and objective formulas are unchanged.

## 1.0.0rc5 — 2026-10-15

### Hydrophobic-moment alignment during sequence search

- Design, Family, Cross-Design and Specificity now orient **every proposed sequence** by its own hydrophobic moment before computing its objective in an interfacial environment. The moment points to the center of the apolar sector. Calibration decoys and random reference sequences follow that same rule. Homogeneous environments do not require rotation, and a numerically zero hydrophobic moment retains the preset phase.
- The search, the displayed score, family ranking, cross-environment compromise and specificity margins now use the same calibrated Hamiltonian and reference distribution for each environment. Completed candidates are not rescored under a second orientation convention. The `design_geometry` response field remains for compatibility and repeats the associated score, Z-score, components and actual phase; its orientation metadata describes the rule actually used.
- The change is a scientific extension of the fixed-geometry sequence optimization described in the [method article](https://doi.org/10.1021/acs.jctc.6c01278) and retained through 1.0.0rc4. Earlier searches held the wheel phase fixed and then also reported a separate hydrophobic-moment-aligned score. In 1.0.0rc5, residue substitutions can rotate the complete helix during the search and thereby change the environmental contributions of other residues.
- Hydrophobic-moment alignment is a prescribed geometric approximation. It does not minimize the full score over all rotation angles, predict equilibrium orientation or establish antimicrobial activity. Multi-environment objectives apply the rule separately in each environment; their scores are not a calculation of a single shared physical orientation across membranes.
- The six component equations, weights, neighbor tables and residue scales are unchanged. Parameter-set checksums therefore remain unchanged; the software version and orientation metadata identify the revised objective. Calibrations and random-reference distributions must match the orientation rule. Aligned scoring reuses only matching aligned calibrations, not the earlier fixed-phase search calibrations.
- Previous **design sequences, rankings, cross-environment compromises and specificity margins must be recalculated** before being attributed to this release. Reusing a seed does not preserve a search trajectory when the objective changes. The standard scoring convention used by Score, Compare and Penetration was already individually aligned; this release unifies searches with that convention.
- The README describes browser workflows and downloadable reproducibility records. Backend request examples remain available as technical documentation for local installations.

## 1.0.0rc4 — 2026-10-14

- The standard-library Python client identifies requests with `User-Agent: Helix-Design-Studio-API/1.0`, allowing reverse proxies to distinguish it from the default Python client.
- Documented the client identification and how to report HTTP errors generated by a reverse proxy before a request reaches the API.
- Scoring equations, numerical parameters, search algorithms and request payloads are unchanged from 1.0.0rc3. The response version identifies this client and documentation release.

## 1.0.0rc3 — 2026-10-14

- Added a user-oriented API guide covering the computational endpoints, score interpretation, provenance, orientation and errors.
- Added executable JSON requests for Design, Compare and Penetration alongside the existing Score example, plus a standard-library Python client. All four examples were executed against the local API.
- Linked Swagger, ReDoc, the OpenAPI schema and the API guide from the browser Guide and README.
- Scoring equations, numerical parameters and search algorithms are unchanged from 1.0.0rc2. The release version in responses identifies this documented source revision.

## 1.0.0rc2 — 2026-10-14

### Neighbor parameter sources and corrections

- **M1:** replaced the supplied table with the 400 printed entries in Nacar (2020), Figure 3 ([doi:10.1007/s10930-020-09880-6](https://doi.org/10.1007/s10930-020-09880-6)). The previous table differs from the source in **303/400 cells when compared in the source's visual layout**. Figure 3 rows are the later residue and columns the earlier residue; the runtime uses rows for the earlier residue and columns for the later residue. After applying this axis mapping, **392/400 runtime entries change** relative to 1.0.0rc1. The 303 count describes discrepancies between the numeric tables in the source layout; the 392 count describes the complete replacement, including orientation. They compare different layouts and must not be added together.
- **M1 model scope:** the published source measures hydrogen-bonded n,n+4 pairs. Its reuse at i,i+1 remains an explicitly documented empirical model adaptation. No logarithmic transformation is introduced. The erratum changes a section cross-reference, not values.
- **M3:** all **324/324 published entries already agree** with Muñoz–Serrano (1994), Table 2 ([doi:10.1038/nsb0694-399](https://doi.org/10.1038/nsb0694-399)); no numerical entries change.
- **M4:** corrected two entries against that table: **F→H −0.850 to −0.800**, and **Y→H −0.800 to −0.850 kcal/mol**. The published row order differs from its column order at F/Y. The resulting runtime table matches all **324/324 published entries**.
- **Cys/Pro in M3/M4:** retained the 76 zero defaults in each 20×20 matrix. The source contains only 18 residues, so these entries are documented as an extension of the model rather than published measurements.
- Included the primary-source CSV tables, bibliographic and axis metadata, source and runtime checksums, `tools/verify_parameter_sources.py`, and regression tests covering all published values, the asymmetric M1 direction and the Cys/Pro extension.
- Updated citation metadata to the software authors Daniel Conde-Torres, Rebeca García-Fandiño and Ángel Piñeiro.

The scoring equation, weights, separation averaging, alphabet, calibration sample protocol and search random streams are unchanged. Parameter checksums change, so the calibration cache automatically separates this parameter set from previous results. Scores and designs must be recalculated with the corrected data; 1.0.0rc1 numerical measurements below describe that earlier candidate.

### Reproducing the parameter comparison

```bash
python tools/verify_parameter_sources.py --output parameter_source_checks.json
```

The report distinguishes agreement with published values (400 M1 + 324 M3 + 324 M4 entries) from the additional Cys/Pro defaults.

### Numerical effects relative to 1.0.0rc1

The [complete parameter comparison](tools/data/parameter_correction_comparison.json) holds the engine, weights, requests and random streams fixed while restoring M1, M4 or both; calibration is recomputed with the same sampled sequences for every parameter case. The benchmark uses the 12 reference sequences in the interface across five presets (60 Z-scores), the full alphabet, 1,000 calibration decoys, 500 random references and seed 42. Six 18-residue designs use 4,000 steps, two restarts, seeds 1–3 in the anionic interface and polar homogeneous medium.

| Parameter correction | Median / max \|ΔZ\| | Spearman of Z | Design positions changed |
| --- | --- | --- | --- |
| M1 only | 0.182503 / 0.430724 | 0.992831 | 58/108 (53.7%) |
| M4 only | 0.000144 / 0.000686 | 1.000000 | 18/108 (16.7%) |
| M1 and M4 | 0.182471 / 0.430694 | 0.992831 | 61/108 (56.5%) |

The largest combined score change is BP100 in homogeneous apolar medium. The mean percentile of 150 random 18-residue sequences in the anionic interface is 51.245 in the previous parameter set and 51.483 after both corrections. Design sequence differences reflect the sensitivity of these paired stochastic searches to changed objectives; the position percentages do not estimate biological effects or typical differences for arbitrary inputs.

To reproduce this historical paired parameter comparison, run the tool from `v1.0.0rc2` and create a separate checkout of the preceding tag. Running it from a later release with changed search objectives would not isolate the parameter corrections:

```bash
git checkout v1.0.0rc2
git worktree add ../helix-1.0.0rc1 v1.0.0rc1
python tools/measure_parameter_corrections.py --baseline ../helix-1.0.0rc1 --output parameter_correction_comparison.json
```

The output includes every score and designed sequence, the complete protocol, parameter and source-code hashes, and Python/NumPy versions. All manuscript demonstrations and measurements are recalculated with the corrected parameter set.

### Rerun of the published method article

The seven sequence strings, printed lengths and fourteen rounded reference Z-scores in `tools/data/article_table1.json` were independently checked against the final published Table 1 (p. 9777). The [1.0.0rc2 direct-coefficient rerun](tools/data/article_table1_comparison_1.0.0rc2.json), with 2,000 calibration sequences, 10,000 random references per environment and seed 42, has mean absolute Z-score difference **0.303771** and maximum **0.439849**. These measure agreement under the declared rerun protocol; they do not establish the historical computation settings or experimental accuracy. The previous candidate output remains archived separately.

```bash
python tools/reproduce_article_table1.py --output article_table1_comparison_1.0.0rc2.json
```

The default command uses direct M1 entries. Its optional `--include-negative-log` mode requires a strictly positive matrix; the source-restored M1 contains zeros, so that transformation is undefined and no pseudocount is introduced.

## 1.0.0rc1 — 2026-10-12

The following historical measurements apply to commit `ce4d718b35212b5994c3ec1c22c6ab5e51e2b8b5` (`v1.0.0rc1`) and the protocols below; they do not describe the corrected 1.0.0rc2 parameter set.

### Scoring model

Measured with `tools/measure_changes.py`: each row restores one 0.1.0 behaviour on top of the candidate, with the other settings held fixed. Benchmark: the 12 reference peptides of the interface in the 5 environment presets (60 Z-scores, full alphabet, 1000 calibration decoys, 500 random references, seed 42); the mean percentile of 150 random 18-residue sequences in the anionic interface (50 is the population expectation for a continuous, exchangeable query/reference distribution); and 6 designs of 18 residues with fixed seeds. The "design positions changed" column describes those six runs. It does not separate sensitivity to model parameters from search variability. The script also reports a separate search control: it changes search entropy from [seed,2] to [seed+100,2], while holding the calibration and random-reference streams fixed. This control changed 41% of design positions over the six runs; it is a descriptive sensitivity check, not an attribution of parameter effects.

| Change | 0.1.0 | 1.0.0rc1 | Reason | Median / max \|ΔZ\| | Spearman of Z | Design positions changed |
|---|---|---|---|---|---|---|
| Helix-neighbor tables | M1 (i,i+1), M3 (i,i+3) and M4 (i,i+4) indexed by position in the requested alphabet | Indexed by residue label, as the Miyazawa–Jernigan table already was | Positional indexing assigned the parameters of other residues whenever the alphabet order differed from the file order (always for M3/M4) | 0.173 / 0.499 | 0.990 | 47% |
| Fauchère–Pliska hydrophobicity | Ser 0.26, Thr −0.04 | Ser −0.04, Thr 0.26 | Values were swapped relative to the published scale | 0.033 / 0.145 | 0.999 | 59% |
| Pace–Scholtz helix propensity | Pro 3.15 | Pro 3.16 | Published value | 0.001 / 0.002 | 1.000 | 31% |
| Miyazawa–Jernigan contacts | i to i+1 (d_max = 1) | i to i+1…i+4 (d_max = 4) | Contact range of the method article (default d_max = 4) | 0.001 / 0.008 | 1.000 | 26% |
| Orientation in scoring | Only the query was oriented by its hydrophobic moment; calibration decoys and random references were evaluated at the query's phase | Query, calibration decoys and random references are oriented by the same rule | The query had an orientation advantage over its background: random sequences scored at a mean percentile of 24.5 (51.2 in the candidate) | 0.363 / 1.626 | 0.951 | scoring only |

The i,i+1, i,i+3 and i,i+4 tables are summed directly, with their stored sign and no per-matrix transformation; 0.1.0 used the same convention. The combined neighbor term is standardized after summation. Eq. 3 motivates an additive term, but does not verify the origin, preprocessing or residue ordering of the supplied numerical tables. The supplied source parameters in 1.0.0rc1 were subsequently corrected in 1.0.0rc2 as documented above.

All changes together (complete 0.1.0 source tree against the candidate, same benchmark): median |ΔZ| 0.491, maximum 1.543 (Poly-Ala 15, neutral interface), Spearman 0.944; mean percentile of random sequences 22.1 in 0.1.0 and 51.2 in the candidate. The orientation comparison above produces the largest median score change among the individual variants in this benchmark. These figures compare software versions on the declared inputs; assessing results obtained with earlier versions requires their original inputs and settings.

Also in the model:

- The i,i+3 and i,i+4 table headers listed 22 labels for 20 columns (Y and V repeated after L). They now list 20 labels: the first 18 as given, and C and P for the last row and column, which are all zero.
- Searches (Design, Family, Cross-Design, Specificity) optimize the fixed-geometry Hamiltonian, as in 0.1.0. Design results now report both the design-geometry energy and the standard score. Cross-Design and Specificity previously reported energies and margins from a separate re-scoring (other calibration seeds, sample sizes and orientation); they now report the optimized quantities, and rankings use them.
- The penetration scan uses the same seed at every depth (previously one seed per depth).
- Calibration, random references and searches draw from independent random streams (`SeedSequence`); 0.1.0 used seed offsets that could coincide. The same user seed therefore selects different random draws across versions.
- The calibration no longer evaluates a hydrophobic-moment term that was not part of the energy (no effect on results).
- Calibration caches are keyed by the parameter tables; caches from 0.1.0 are not reused.

### API and interface

- Stricter validation: standard residues only, the sequence must use the allowed residues, known presets and membrane charges, bounded angles, weights and seeds, no NaN/Infinity, no unknown fields, at most four off-target environments. Physically identical target and off-target environments are rejected. `wheel_phase_deg` is no longer accepted. `/api/design` uses the 20-residue alphabet by default, like the interface.
- Client errors return 400/422 and internal failures 500; `/api/health` only reports server-side failures and includes the version.
- Calculations have a concurrency limit, a time limit and a request-size limit, and stop when the client disconnects (including streams and calibration).
- Results include `provenance` and `calibration` blocks. Stream metadata and every fully evaluated candidate carry provenance, so interrupted Family and Specificity downloads preserve the complete request and version. Both workflows export their partial JSON results after stop or error.
- Interface fixes: cancel buttons, partial results kept on stop or error, consistent meaning of the penetration slider, helical wheel drawn with the model's angular sector, escaped error messages, accessibility and layout fixes. Two reference peptides whose sequences did not match the cited proteins (GB1 helix fragment, α3D segment) were removed and KALP17 follows the GKK(LA)ₙLKKA pattern.

### Reproducing the measurements

Run this historical benchmark from `v1.0.0rc1` to reproduce the measurements above.

```bash
git checkout v1.0.0rc1
git worktree add ../helix-0.1.0 v0.1.0
python tools/measure_changes.py --baseline ../helix-0.1.0
```

Add `--output model_change_comparison.json` to save the full input protocol, scores, designs and numerical summary, including the control that changes only the search stream.

### Comparison with the method article's Table 1

`tools/data/article_table1.json` contains seven exact sequence strings and fourteen Z-scores transcribed from the identified author manuscript's Table 1, with its DOI, source hash and the original table rows. GWALP23 is printed with length 23 but contains 21 residues in that sequence string; the rerun evaluates the printed sequence and records both lengths. Terminal modifications are omitted, as in the sequence-only model.

Run the historical two-convention comparison from `v1.0.0rc1`. In 1.0.0rc2, M1 retains source zero values and −ln(M1) is undefined for those entries; the current tool reports this rather than introducing an arbitrary pseudocount.

```bash
git checkout v1.0.0rc1
python tools/reproduce_article_table1.py --output article_table1_comparison.json
```

The archived 1.0.0rc1 rerun uses the full 20-residue alphabet in its recorded order, default weights, 2,000 calibration sequences, 10,000 random references per environment, seed 42 and the candidate's independent random streams. It compares direct M1 entries with −ln(M1), holding every other setting and the random sequences fixed while recalibrating the combined neighbor term. The output includes all fourteen scores for each convention, error summaries, input and source-file hashes, and runtime versions. Agreement with the rounded reference values does not reproduce the historical calculation protocol. This earlier rerun uses the 1.0.0rc1 parameter set, before the source corrections above.

The [archived numerical output](tools/data/article_table1_comparison.json) gives a mean absolute Z-score difference of **0.444852** for direct M1 entries and **1.313673** for −ln(M1), over the fourteen transcribed reference values. The maximum absolute differences are 0.670831 and 1.666431, respectively. These are comparisons with the rounded values under this rerun protocol, not estimates of experimental accuracy.

## 0.1.0

Web application source before the changes above (API version 0.1.0, commit `a8cdf43`).
