## Additional file 1. Evaluated software source and dependencies for "Helix Design Studio: designing antimicrobial peptide candidates for membrane environments": hamiltonian_builder.py

```
"""Classical sequence scoring and random-ensemble calibration for Helix Design Studio."""

import hashlib
import json
import logging
import math
import os
import tempfile
from typing import Any, Dict, List, Optional, Sequence, Tuple

import numpy as np

from data_loaders.energy_matrix_loader import (
    _load_first_neighbors_matrix_file,
    _load_third_neighbors_matrix_file,
    _load_fourth_neighbors_matrix_file,
    _load_energy_matrix_file
)

# Propensión a hélice de Pace y Scholtz (1998), kcal/mol: menor valor = mayor propensión.
PACE_SCHOLTZ_HELIX_PROPENSITY = {
    'A': 0.00, 'L': 0.21, 'R': 0.21, 'M': 0.24, 'K': 0.26, 'Q': 0.39,
    'E': 0.40, 'I': 0.41, 'W': 0.49, 'S': 0.50, 'Y': 0.53, 'F': 0.54,
    'V': 0.61, 'H': 0.61, 'N': 0.65, 'T': 0.66, 'C': 0.68, 'D': 0.69,
    'G': 1.00, 'P': 3.16
}
# Hidrofobicidad de Fauchère y Pliska (1983), π: >0 apolar, <0 polar.
FAUCHERE_PLISKA_HYDROPHOBICITY = {
    'D': -0.77, 'E': -0.64, 'K': -0.99, 'R': -1.01, 'H':  0.13,
    'G':  0.00, 'A':  0.31, 'V':  1.22, 'L':  1.70, 'I':  1.80,
    'P':  0.72, 'M':  1.23, 'F':  1.79, 'W':  2.25, 'Y':  0.96,
    'T':  0.26, 'S': -0.04, 'C':  1.54, 'N': -0.60, 'Q': -0.22
}
RESIDUE_ANGLE_DEG = 100.0  # giro por residuo en una hélice alfa
DEFAULT_STATS_CACHE_MAX_FILES = 2000

logger = logging.getLogger(__name__)


def hydrophobic_moment(sequence: Sequence[str]) -> Tuple[float, float]:
    """Componentes (x, y) del momento hidrofóbico; el residuo i (desde 1) está en (i-1)*100°."""
    mx = my = 0.0
    for idx, aa in enumerate(sequence):
        hydro = FAUCHERE_PLISKA_HYDROPHOBICITY.get(aa, 0.0)
        theta = math.radians(idx * RESIDUE_ANGLE_DEG)
        mx += hydro * math.cos(theta)
        my += hydro * math.sin(theta)
    return mx, my


RESOURCE_FILES = ("mj_matrix.txt", "helix_pairs_prop.txt", "helix_sd_i_3.txt", "helix_sd_i_4.txt")
NEIGHBOR_SCORING_CONVENTION = "negative_m1_plus_m3_plus_m4"
NEIGHBOR_MATRIX_WEIGHTS = {1: -1.0, 3: 1.0, 4: 1.0}


def parameter_set_sha256() -> str:
    """Huella de tablas, escalas y convención de vecinos, independiente del alfabeto."""
    digest = hashlib.sha256()
    resources = os.path.join(os.path.dirname(__file__), "resources")
    for name in RESOURCE_FILES:
        with open(os.path.join(resources, name), "rb") as fh:
            digest.update(name.encode("utf-8") + b"\0" + fh.read())
    scales = {
        "pace_scholtz": PACE_SCHOLTZ_HELIX_PROPENSITY,
        "fauchere_pliska": FAUCHERE_PLISKA_HYDROPHOBICITY,
        "neighbor_score_convention": NEIGHBOR_SCORING_CONVENTION,
        "neighbor_matrix_weights": NEIGHBOR_MATRIX_WEIGHTS,
    }
    digest.update(json.dumps(scales, sort_keys=True).encode("utf-8"))
    return digest.hexdigest()


def aligned_phase_deg(sequence: Sequence[str]) -> Optional[float]:
    """Fase de la rueda que centra el momento hidrofóbico en el sector de membrana (None si es nulo)."""
    mx, my = hydrophobic_moment(sequence)
    if math.hypot(mx, my) < 1e-10:
        return None
    return (-math.degrees(math.atan2(my, mx))) % 360.0


class HamiltonianBuilder:
    """Length-scaled Hamiltonian components with random-ensemble Z-score calibration.

    Con align_phase=True, cada secuencia evaluada (consulta, señuelos de calibración, referencias
    aleatorias o propuestas de búsqueda) se orienta con su propio momento hidrofóbico, de modo que
    todas se comparan bajo el mismo criterio.
    """

    TERMS = ('helix_local', 'env_pol', 'env_chg', 'pw_int', 'electrostatic', 'helix_neigh')

    def __init__(
        self,
        L: int,
        amino_acids: List[str],
        **kwargs
    ):
        self.L = L
        self.amino_acids = amino_acids
        self.n_aa = len(amino_acids)
        self.kwargs = kwargs
        self.stats_cache_version = 5
        self.align_phase = bool(kwargs.get('align_phase', False))

        # 1. Physicochemical properties
        self._init_properties()

        # 2. Matrices MJ y vecinos estadísticos k=1,3,4
        self._load_matrices_from_files()

        # 3. Calibración Z-score real (Goldstein et al. 1992)
        #    Evalúa n_decoys secuencias aleatorias para estimar μ y σ de cada término
        n_decoys = self.kwargs.get('n_decoys', 10000)
        self.stats = self._load_or_calibrate_z_scores(n_decoys=n_decoys)

    def _stats_cache_payload(self, n_decoys: int) -> Dict[str, Any]:
        return {
            "stats_cache_version": self.stats_cache_version,
            "neighbor_score_convention": NEIGHBOR_SCORING_CONVENTION,
            "neighbor_matrix_weights": NEIGHBOR_MATRIX_WEIGHTS,
            "L": self.L,
            "amino_acids": self.amino_acids,
            "n_decoys": int(n_decoys),
            "zscore_seed": int(self.kwargs.get('zscore_seed', 42)),
            "wheel_phase_deg": float(self.kwargs.get('wheel_phase_deg', 0.0)),
            "wheel_halfwidth_deg": float(self.kwargs.get('wheel_halfwidth_deg', 90.0)),
            "align_phase": bool(self.kwargs.get('align_phase', False)),
            "membrane_charge": self.kwargs.get('membrane_charge', 'neg'),
            "max_interaction_dist": int(self.kwargs.get('max_interaction_dist', 1)),
            "electrostatic_cutoff": int(self.kwargs.get('electrostatic_cutoff', 8)),
            "hydro": self.hydro,
            "charges": self.charges,
            "h_alpha": self.h_alpha,
            "parameters_sha256": self._parameters_fingerprint(),
        }

    def _stats_cache_path(self, n_decoys: int) -> str:
        payload = self._stats_cache_payload(n_decoys)
        cache_key = hashlib.sha256(
            json.dumps(payload, sort_keys=True, separators=(",", ":")).encode("utf-8")
        ).hexdigest()
        cache_dir = self.kwargs.get('stats_cache_dir') or os.path.join(tempfile.gettempdir(), "helix-design-studio-cache")
        os.makedirs(cache_dir, mode=0o700, exist_ok=True)
        return os.path.join(cache_dir, f"zscore_stats_{cache_key}.json")

    @staticmethod
    def _cache_dir_is_private(cache_dir: str) -> bool:
        """Solo se confía en una caché propia y no escribible por otros usuarios."""
        info = os.stat(cache_dir)
        if hasattr(os, "getuid") and info.st_uid != os.getuid():
            return False
        return not info.st_mode & 0o022

    def _prune_stats_cache(self, cache_dir: str) -> None:
        """Mantiene la caché acotada eliminando las calibraciones usadas hace más tiempo."""
        max_files = int(self.kwargs.get('stats_cache_max_files') or DEFAULT_STATS_CACHE_MAX_FILES)
        entries = []
        with os.scandir(cache_dir) as listing:
            for entry in listing:
                if entry.name.startswith("zscore_stats_") and entry.name.endswith(".json"):
                    entries.append((entry.stat().st_mtime, entry.path))
        for _, path in sorted(entries)[:max(0, len(entries) - max_files)]:
            try:
                os.unlink(path)
            except OSError:
                pass

    def _load_or_calibrate_z_scores(self, n_decoys: int) -> Dict[str, Tuple[float, float]]:
        cache_path = self._stats_cache_path(n_decoys)
        cache_dir = os.path.dirname(cache_path)
        use_cache = self._cache_dir_is_private(cache_dir)
        if not use_cache:
            logger.warning("Ignoring Z-score cache directory not private to this user: %s", cache_dir)
        if use_cache and os.path.exists(cache_path):
            cached_stats = self._read_stats_cache(cache_path, n_decoys)
            if cached_stats is not None:
                logger.debug("Reusing cached Z-score calibration: %s", cache_path)
                try:
                    os.utime(cache_path)
                except OSError:
                    pass
                return cached_stats

        logger.debug("Calibrating Z-score statistics with %d decoys", n_decoys)
        stats = self._calibrate_z_scores(n_decoys=n_decoys)
        if not use_cache:
            return stats
        serializable = {
            key: {"mu": float(mu), "sigma": float(sigma)}
            for key, (mu, sigma) in stats.items()
        }
        payload = {
            "metadata": self._stats_cache_payload(n_decoys),
            "stats": serializable,
        }
        # Escritura atómica: una petición concurrente nunca lee un fichero a medio escribir.
        tmp_path = None
        try:
            fd, tmp_path = tempfile.mkstemp(dir=os.path.dirname(cache_path), prefix=".zscore_", suffix=".tmp")
            with os.fdopen(fd, "w", encoding="utf-8") as fh:
                json.dump(payload, fh, indent=2, sort_keys=True)
            os.replace(tmp_path, cache_path)
            tmp_path = None
            logger.debug("Saved Z-score calibration cache: %s", cache_path)
            self._prune_stats_cache(cache_dir)
        except OSError as exc:
            logger.warning("Could not save Z-score calibration cache: %s", exc)
        finally:
            if tmp_path is not None and os.path.exists(tmp_path):
                os.unlink(tmp_path)
        return stats

    def _read_stats_cache(self, cache_path: str, n_decoys: int):
        """Devuelve las estadísticas cacheadas, o None si el fichero es ilegible o no coincide."""
        expected_metadata = json.loads(json.dumps(self._stats_cache_payload(n_decoys)))
        try:
            with open(cache_path, "r", encoding="utf-8") as fh:
                cached = json.load(fh)
            if cached.get("metadata") != expected_metadata:
                return None
            stats = {
                key: (float(values["mu"]), float(values["sigma"]))
                for key, values in cached["stats"].items()
            }
        except (OSError, ValueError, KeyError, TypeError, AttributeError) as exc:
            logger.warning("Ignoring unreadable Z-score cache %s: %s", cache_path, exc)
            return None
        # Una σ nula o negativa dividiría por cero o invertiría el signo de un término.
        valid = all(math.isfinite(mu) and math.isfinite(sigma) and sigma > 0 for mu, sigma in stats.values())
        if set(stats) != set(self.TERMS) or not valid:
            logger.warning("Ignoring inconsistent Z-score cache %s", cache_path)
            return None
        return stats

    # ------------------------------------------------------------------
    # Inicialización de propiedades
    # ------------------------------------------------------------------

    def _init_properties(self):
        """
        Pace-Scholtz (hélice) y Fauchère-Pliska (hidrofobicidad).
        """
        self.h_alpha = dict(PACE_SCHOLTZ_HELIX_PROPENSITY)
        self.hydro = dict(FAUCHERE_PLISKA_HYDROPHOBICITY)
        # Cargas formales
        self.charges = {aa: 0 for aa in self.amino_acids}
        for aa in ['R', 'K']:
            self.charges[aa] = 1
        for aa in ['D', 'E']:
            self.charges[aa] = -1

    def _load_matrices_from_files(self):
        """Carga MJ y matrices de vecinos estadísticos k=1,3,4 en el orden del alfabeto."""
        self.mj_matrix = self._matrix_for_alphabet(*_load_energy_matrix_file())
        # Keep the published coefficients intact. The neighbor score subtracts M1,
        # so larger observed/expected helix propensities give a lower model score.
        # M3/M4 retain their source energy signs (negative contributions are favorable).
        self.M1 = self._matrix_for_alphabet(*_load_first_neighbors_matrix_file())
        self.M3 = self._matrix_for_alphabet(*_load_third_neighbors_matrix_file())
        self.M4 = self._matrix_for_alphabet(*_load_fourth_neighbors_matrix_file())

    def _matrix_for_alphabet(self, matrix: np.ndarray, symbols: List[str]) -> np.ndarray:
        """Reindexa una tabla por etiqueta de residuo: fila/columna i = self.amino_acids[i]."""
        aa_to_idx = {aa: i for i, aa in enumerate(symbols)}
        missing = [aa for aa in self.amino_acids if aa not in aa_to_idx]
        if missing:
            raise ValueError(f"Residues {missing} are not parameterized (available: {symbols})")
        order = [aa_to_idx[aa] for aa in self.amino_acids]
        return np.array(matrix, dtype=float)[np.ix_(order, order)]

    def _parameters_fingerprint(self) -> str:
        """Huella de las tablas usadas, para invalidar la caché si cambian los parámetros."""
        digest = hashlib.sha256()
        for matrix in (self.mj_matrix, self.M1, self.M3, self.M4):
            digest.update(np.ascontiguousarray(matrix, dtype=np.float64).tobytes())
        convention = {
            "neighbor_score_convention": NEIGHBOR_SCORING_CONVENTION,
            "neighbor_matrix_weights": NEIGHBOR_MATRIX_WEIGHTS,
        }
        digest.update(json.dumps(convention, sort_keys=True).encode("utf-8"))
        return digest.hexdigest()

    # ------------------------------------------------------------------
    # Entorno (Helical Wheel)
    # ------------------------------------------------------------------

    def phase_for(self, seq: List[int]) -> float:
        """Fase de la rueda con la que se evalúa una secuencia."""
        base = float(self.kwargs.get('wheel_phase_deg', 0.0))
        if not self.align_phase:
            return base
        aligned = aligned_phase_deg([self.amino_acids[a] for a in seq])
        return base if aligned is None else aligned

    def _membrane_mask(self, phase_deg: float) -> List[bool]:
        """True para las posiciones cuya cara queda dentro del sector de membrana."""
        half_width = float(self.kwargs.get('wheel_halfwidth_deg', 90.0))
        mask = []
        for i in range(self.L):
            theta = (i * RESIDUE_ANGLE_DEG + phase_deg) % 360.0
            angle = theta if theta <= 180.0 else theta - 360.0
            mask.append(abs(angle) <= half_width)
        return mask

    def _mask_for(self, seq: List[int], phase_deg: Optional[float]) -> List[bool]:
        return self._membrane_mask(self.phase_for(seq) if phase_deg is None else phase_deg)

    # ------------------------------------------------------------------
    # Random-ensemble Z-score calibration
    # ------------------------------------------------------------------

    def _raw_helix_local(self, seq: List[int]) -> float:
        scale = 1.0 / self.L
        return sum(self.h_alpha[self.amino_acids[a]] for a in seq) * scale

    def _raw_env_pol(self, seq: List[int], phase_deg: Optional[float] = None) -> float:
        """
        Actua en AMBOS entornos (simetria anfipática):
          Membrana: -h -> premia apolares (h>0), penaliza polares (h<0)
          Agua:     +h -> premia polares  (h<0), penaliza apolares (h>0)
        """
        scale = 1.0 / self.L
        total = 0.0
        for a, in_membrane in zip(seq, self._mask_for(seq, phase_deg)):
            h = self.hydro[self.amino_acids[a]]
            total += -h if in_membrane else +h
        return total * scale

    def _raw_env_chg(self, seq: List[int], phase_deg: Optional[float] = None) -> float:
        """
        Actua SOLO en agua. Premio UNIDIRECCIONAL: solo favorece la carga
        OPUESTA a sigma, nunca penaliza la carga del mismo signo.
          neg (sigma=-1): K,R   en agua -> min((-1)(+1), 0) = -1  FAV
                          D,E   en agua -> min((-1)(-1), 0) =  0  NEUTRO
          pos (sigma=+1): D,E   en agua -> min((+1)(-1), 0) = -1  FAV
                          K,R   en agua -> min((+1)(+1), 0) =  0  NEUTRO
        Residuos neutros (q=0): sin efecto en cualquier caso.
        """
        scale = 1.0 / self.L
        m_charge = self.kwargs.get('membrane_charge', 'neg').lower()
        sigma = {'neg': -1.0, 'pos': 1.0, 'neu': 0.0}[m_charge]
        total = 0.0
        for a, in_membrane in zip(seq, self._mask_for(seq, phase_deg)):
            if not in_membrane:   # solo en agua
                raw = sigma * self.charges[self.amino_acids[a]]
                total += min(raw, 0.0)            # solo el componente favorable
        return total * scale

    def _raw_pw_int(self, seq: List[int]) -> float:
        d_max = self.kwargs.get('max_interaction_dist', 1)
        scale = 1.0 / (self.L * d_max)
        total = 0.0
        for i in range(self.L):
            for j in range(i + 1, min(i + d_max + 1, self.L)):
                total += self.mj_matrix[seq[i], seq[j]]
        return total * scale

    def _raw_electrostatic(self, seq: List[int], phase_deg: Optional[float] = None) -> float:
        r_c = self.kwargs.get('electrostatic_cutoff', 8)
        scale = 2.0 / (self.L * (self.L - 1)) if self.L > 1 else 1.0
        mask = self._mask_for(seq, phase_deg)
        total = 0.0
        for i in range(self.L):
            qi = self.charges[self.amino_acids[seq[i]]]
            if not qi:
                continue
            for j in range(i + 1, min(self.L, i + r_c + 1)):
                qj = self.charges[self.amino_acids[seq[j]]]
                if not qj:
                    continue
                if mask[i] and mask[j]:
                    eps = 0.1
                elif not mask[i] and not mask[j]:
                    eps = 1.0
                else:
                    eps = 0.3
                dist = j - i
                total += (qi * qj) / ((1 + dist) * eps)
        return total * scale

    def _raw_helix_neigh(self, seq: List[int]) -> float:
        """Empirical neighbor score: -mean(M1) + mean(M3) + mean(M4)."""
        matrices = {1: self.M1, 3: self.M3, 4: self.M4}
        total = 0.0
        for k, matrix in matrices.items():
            if self.L <= k:
                continue
            scale = 1.0 / (self.L - k)
            for i in range(self.L - k):
                total += NEIGHBOR_MATRIX_WEIGHTS[k] * matrix[seq[i], seq[i + k]] * scale
        return total

    def raw_terms(self, seq: List[int]) -> Dict[str, float]:
        """Términos brutos de una secuencia, evaluados con su fase (alineada si align_phase)."""
        phase = self.phase_for(seq)
        return {
            'helix_local': self._raw_helix_local(seq),
            'env_pol': self._raw_env_pol(seq, phase),
            'env_chg': self._raw_env_chg(seq, phase),
            'pw_int': self._raw_pw_int(seq),
            'electrostatic': self._raw_electrostatic(seq, phase),
            'helix_neigh': self._raw_helix_neigh(seq),
        }

    def _calibrate_z_scores(self, n_decoys: int) -> Dict[str, Tuple[float, float]]:
        """
        Estima μ y σ para cada término evaluando n_decoys secuencias aleatorias.
        """
        rng = np.random.default_rng(self.kwargs.get('zscore_seed', 42))
        accum: Dict[str, List[float]] = {key: [] for key in self.TERMS}
        # Permite detener la calibración (cancelación o plazo) sin esperar a que termine.
        interrupt_check = self.kwargs.get('interrupt_check')

        for index in range(n_decoys):
            if interrupt_check is not None and index % 100 == 0:
                interrupt_check()
            seq = rng.integers(0, self.n_aa, size=self.L).tolist()
            for key, value in self.raw_terms(seq).items():
                accum[key].append(value)

        stats = {}
        for key, values in accum.items():
            arr = np.array(values)
            mu = float(np.mean(arr))
            sigma = float(np.std(arr))
            # Evitar división por cero: si σ≈0 el término es constante → no aporta gradiente
            stats[key] = (mu, sigma if sigma > 1e-12 else 1.0)

        return stats

    def _apply_z_score(self, term_name: str, raw_energy: float) -> float:
        """H' = (H - μ) / σ."""
        mu, sigma = self.stats[term_name]
        return (raw_energy - mu) / sigma
```
