## Additional file 1. Evaluated software source and dependencies for "Helix Design Studio: designing antimicrobial peptide candidates for membrane environments": __init__.py

```
# (C) Copyright IBM 2021, 2022.
#
# This code is licensed under the Apache License, Version 2.0. You may
# obtain a copy of this license in the LICENSE.txt file in the root directory
# of this source tree or at http://www.apache.org/licenses/LICENSE-2.0.
#
# Any modifications or derivative works of this code must retain this
# copyright notice, and modified files need to carry a notice indicating
# that they have been altered from the originals.
"""
The mj_matrix.txt file stores knowledge-based (statistical) potentials for the description of
inter-residue contacts for proteins derived by using quasi-chemical approximation. The potentials
used here are introduced by Miyazawa, S. and Jernigan, R. L. in J. Mol. Biol.256, 623–644 (1996)
in Table 3.
"""
```
