## Additional file 1. Evaluated software source and dependencies for "Helix Design Studio: designing antimicrobial peptide candidates for membrane environments": energy_matrix_loader.py

```
# (C) Copyright IBM 2021, 2022.
#
# This code is licensed under the Apache License, Version 2.0. You may
# obtain a copy of this license in the LICENSE.txt file in the root directory
# of this source tree or at http://www.apache.org/licenses/LICENSE-2.0.
#
# Any modifications or derivative works of this code must retain this
# copyright notice, and modified files need to carry a notice indicating
# that they have been altered from the originals.
#
# Modified by the Helix Design Studio contributors: loaders for additional
# parameter tables, rewritten parsing, symmetrisation and validation, caching
# and logging. A copy of the license is included in LICENSES/Apache-2.0.txt.

import functools
import logging
import numpy as np
import os
from typing import List, Tuple

logger = logging.getLogger(__name__)

def _construct_resource_path(filename: str) -> str:
    """
    Constructs the file path for a given resource file.
    This function navigates up one directory and then down into 'core/resources'.
    """
    path = os.path.realpath(
        os.path.join(
            os.path.dirname(__file__),
            "..",
            "core",
            "resources",
            filename,
        )
    )
    return os.path.normpath(path)


STANDARD_RESIDUES = frozenset("ACDEFGHIKLMNPQRSTVWY")


def _validate_labeled_matrix(matrix: np.ndarray, symbols: List[str], name: str) -> None:
    """
    Checks that a parameter table is a finite 20x20 matrix labelled once with each
    of the 20 standard residues, so that callers can index it by residue label.
    """
    if len(set(symbols)) != len(symbols):
        raise ValueError(f"{name} header contains repeated residue labels: {symbols}")
    if set(symbols) != STANDARD_RESIDUES:
        raise ValueError(f"{name} header must list the 20 standard residues, got {symbols}")
    if matrix.shape != (len(symbols), len(symbols)):
        raise ValueError(
            f"{name} shape {matrix.shape} does not match its {len(symbols)} header labels"
        )
    if not np.isfinite(matrix).all():
        raise ValueError(f"{name} contains non-finite values")


@functools.lru_cache(maxsize=None)
def _load_energy_matrix_file() -> Tuple[np.ndarray, Tuple[str, ...]]:
    """
    Loads the Miyazawa-Jernigan interaction energy matrix from file.
    The file is read as a full 20x20 matrix where the upper triangle
    and diagonal contain the correct interaction values.

    Constructs the symmetric matrix (M=M_diag + M_upper + M_upper.T).
    """
    path = _construct_resource_path("mj_matrix.txt")

    try:
        with open(path, 'r') as f:
            lines = f.readlines()

        # First line contains amino acid symbols
        symbols = lines[0].strip().split()
        n_aa = len(symbols)

        # Read the raw 20x20 matrix from file
        data_lines = [line.strip() for line in lines[1:] if line.strip()]
        energy_matrix_raw = np.loadtxt(data_lines, dtype=float, ndmin=2)

        if energy_matrix_raw.shape != (n_aa, n_aa):
             raise ValueError(f"Matrix shape error. Expected ({n_aa}, {n_aa}), but got {energy_matrix_raw.shape}")

        # --- Simetrización Correcta ---
        # 1. Obtener el triángulo superior estricto (sin la diagonal, k=1)
        M_upper = np.triu(energy_matrix_raw, k=1)

        # 2. Obtener la diagonal
        M_diag = np.diag(np.diag(energy_matrix_raw))

        # 3. Construir la matriz simétrica final: M_diag + M_upper + M_upper.T
        energy_matrix = M_diag + M_upper + M_upper.T

        # -----------------------------

        _validate_labeled_matrix(energy_matrix, symbols, "MJ matrix")
        logger.debug("Loaded MJ matrix %s with symbols %s", energy_matrix.shape, symbols)
        energy_matrix.setflags(write=False)

        return energy_matrix, tuple(symbols)

    except FileNotFoundError:
        raise FileNotFoundError(f"Error: The file {path} was not found. Check the path and filename.") from None
    except Exception as e:
        raise RuntimeError(f"Error loading Miyazawa-Jernigan matrix: {e}") from e

@functools.lru_cache(maxsize=None)
def _load_first_neighbors_matrix_file() -> Tuple[np.ndarray, Tuple[str, ...]]:
    """
    Loads the helix propensity matrix from file.
    Expected format:
    - First line: amino acid symbols (tab/space separated)
    - Following lines: full square matrix (n_aa x n_aa values)
    """
    path = _construct_resource_path("helix_pairs_prop.txt")

    try:
        with open(path, 'r') as f:
            lines = f.readlines()

        # First line contains amino acid symbols; the rest is the square matrix
        symbols = lines[0].strip().split()
        data_lines = [line.strip() for line in lines[1:] if line.strip()]
        helix_matrix = np.loadtxt(data_lines, dtype=float, ndmin=2)

        _validate_labeled_matrix(helix_matrix, symbols, "Helix propensity matrix")
        logger.debug("Loaded helix propensity matrix %s with symbols %s", helix_matrix.shape, symbols)
        helix_matrix.setflags(write=False)
        return helix_matrix, tuple(symbols)

    except FileNotFoundError:
        raise FileNotFoundError(f"Error: The file {path} was not found. Check the path and filename.") from None
    except Exception as e:
        raise RuntimeError(f"Error loading helix propensity matrix: {e}") from e

@functools.lru_cache(maxsize=None)
def _load_third_neighbors_matrix_file() -> Tuple[np.ndarray, Tuple[str, ...]]:
    """Returns the helix i,i+3 interaction matrix from file."""
    path = _construct_resource_path("helix_sd_i_3.txt")

    try:
        with open(path, 'r') as f:
            lines = f.readlines()

        # The first line contains the symbols
        symbols = lines[0].strip().split()

        # The rest of lines contain the matrix data
        data_lines = [line.strip() for line in lines[1:] if line.strip()]
        helix_matrix = np.loadtxt(data_lines, dtype=float, ndmin=2)

        _validate_labeled_matrix(helix_matrix, symbols, "Helix i,i+3 matrix")
        logger.debug("Loaded helix i,i+3 matrix %s with symbols %s", helix_matrix.shape, symbols)
        helix_matrix.setflags(write=False)
        return helix_matrix, tuple(symbols)

    except FileNotFoundError:
        raise FileNotFoundError(f"Error: The file {path} was not found. Check the path and filename.") from None
    except Exception as e:
        raise RuntimeError(f"Error loading helix i,i+3 matrix: {e}") from e

@functools.lru_cache(maxsize=None)
def _load_fourth_neighbors_matrix_file() -> Tuple[np.ndarray, Tuple[str, ...]]:
    """Returns the helix i,i+4 interaction matrix from file."""
    path = _construct_resource_path("helix_sd_i_4.txt")

    try:
        with open(path, 'r') as f:
            lines = f.readlines()

        # The first line contains the symbols
        symbols = lines[0].strip().split()

        # The rest of lines contain the matrix data
        data_lines = [line.strip() for line in lines[1:] if line.strip()]
        helix_matrix = np.loadtxt(data_lines, dtype=float, ndmin=2)

        _validate_labeled_matrix(helix_matrix, symbols, "Helix i,i+4 matrix")
        logger.debug("Loaded helix i,i+4 matrix %s with symbols %s", helix_matrix.shape, symbols)
        helix_matrix.setflags(write=False)
        return helix_matrix, tuple(symbols)

    except FileNotFoundError:
        raise FileNotFoundError(f"Error: The file {path} was not found. Check the path and filename.") from None
    except Exception as e:
        raise RuntimeError(f"Error loading helix i,i+4 matrix: {e}") from e
```
