## Additional file 1. Evaluated software source and dependencies for "Helix Design Studio: designing antimicrobial peptide candidates for membrane environments": Dockerfile

```
FROM python:3.12-slim

ENV PYTHONUNBUFFERED=1 \
    PYTHONDONTWRITEBYTECODE=1 \
    HELIX_CACHE_DIR=/tmp/helix-cache \
    HELIX_VAR_DIR=/tmp/helix-usage \
    HELIX_USAGE_LOGGING=0
WORKDIR /app
COPY requirements.txt ./
RUN pip install --no-cache-dir -r requirements.txt \
    && useradd --create-home --uid 10001 helix
COPY app ./app
COPY Code ./Code
COPY LICENSE THIRD_PARTY_NOTICES.md ./
COPY LICENSES ./LICENSES
USER helix
EXPOSE 8000
HEALTHCHECK --interval=30s --timeout=5s --start-period=20s --retries=3 \
    CMD ["python", "-c", "import os, urllib.request; urllib.request.urlopen('http://127.0.0.1:%s/' % os.environ.get('PORT', '8000'), timeout=4)"]
# exec replaces the shell so that uvicorn receives SIGTERM and shuts down cleanly.
CMD ["sh", "-c", "exec uvicorn app.main:app --host 0.0.0.0 --port ${PORT:-8000} --workers ${WEB_CONCURRENCY:-1} --proxy-headers --forwarded-allow-ips=${FORWARDED_ALLOW_IPS:-127.0.0.1} --timeout-graceful-shutdown 30"]
```
