## Additional file 1. Evaluated software source and dependencies for "Helix Design Studio: designing antimicrobial peptide candidates for membrane environments": API.md

# API guide

This is technical documentation for the backend of a local Helix Design Studio installation. The browser interface is the application workflow for scoring and designing peptide sequences. When testing the backend, use the intended software version and retain the complete JSON responses for reproducibility.

## Interactive documentation

On a local installation, open [Swagger UI](http://127.0.0.1:8000/docs), [ReDoc](http://127.0.0.1:8000/redoc), or the [OpenAPI schema](http://127.0.0.1:8000/openapi.json). Swagger UI shows request fields and validation limits and lets you execute requests with **Try it out**.

Check the local schema's `info.version` and each result's `provenance.version` before comparing calculations. For a specified release, install its Git tag using the [installation instructions](../README.md#installation).

## Available calculations

All calculation endpoints below accept HTTP `POST` with `Content-Type: application/json`.

| Endpoint | Purpose | Main response fields |
|---|---|---|
| `/api/score` | Score one sequence in one environment | `sequence`, `energy`, `z_score`, `breakdown`, `environment`, `calibration`, `random_reference` |
| `/api/design` | Search for one sequence in a target environment | Score fields, compatible `design_geometry` copy, `design_settings` |
| `/api/compare` | Evaluate the same sequence in two environments | `environment_a`, `environment_b`, `energy_gap`, `z_score_gap` |
| `/api/optimal-penetration` | Scan angular membrane exposure from 0 to 100% | `profile`, `best_penetration_percent`, `best_score`, `scan_settings` |
| `/api/design-family` | Generate a sequence family in one environment | `members`, `family_size_returned` |
| `/api/design-family-stream` | Receive family candidates as they become available | Newline-delimited JSON events |
| `/api/cross-design` | Search for compatibility with two environments | `sequence`, `environment_a`, `environment_b`, `cross_design` |
| `/api/specificity-design` | Search for candidates for a target versus off-target environments | `candidates`, `specificity_collection`; top-ranked candidate also exposed as `sequence`, `target_environment`, `off_target_environments`, `specificity_design` |
| `/api/specificity-design-stream` | Receive specificity candidates as they become available | Newline-delimited JSON events |
| `/api/pdb` | Construct an ideal alpha-helical PDB for a sequence | `sequence`, `pdb` |

Calculation responses include `provenance` with the software version, parameter checksum, validated request, seed protocol and runtime versions. Streaming responses attach provenance to the metadata, candidate and final payloads. The streaming content type is `application/x-ndjson`: read one JSON object per line and inspect the `type` field (`meta`, `phase`, `progress`, `candidate`, `final`, or `error`). An error event can occur after the HTTP response has started, so HTTP status alone does not establish that a streamed calculation completed.

For family design, iterate over `members`. For specificity design, iterate over `candidates`, whose entries each contain a sequence, target/off-target scores and `specificity_design` metrics. `specificity_collection.requested` and `.returned` identify the requested and recovered collection sizes; the top-level candidate fields duplicate the highest-ranked entry for convenient inspection. In both streaming workflows, each `candidate` event carries that candidate in `payload.member`, and the `final` event carries the complete collection in `payload`.

## Inputs and scoring conventions

- `sequence` uses standard one-letter amino-acid codes. The `residues` list specifies the allowed alphabet and must include every residue in a supplied sequence. Its order is part of the random sampling protocol.
- `environment.preset` is one of `homogeneous_polar`, `homogeneous_apolar`, `interfacial_neg`, `interfacial_neu`, or `interfacial_pos`. Interfacial presets represent anionic, neutral or cationic membrane surfaces. Their default apolar-sector half-width is 90°. `wheel_halfwidth_deg` can override this angular exposure. `membrane_charge` can override the surface-charge convention with `neg`, `neu`, or `pos`.
- `weights` can override the six Hamiltonian weights. Omitted weights use the release defaults shown in the OpenAPI schema.
- `n_decoys` sets the calibration sample for standardizing Hamiltonian components. `n_random` sets the separate random-sequence reference used by Score and Penetration for the total-score Z-score. Design and Compare use a fixed reference of 500 sequences per environment in this version.
- `seed` makes a request reproducible under the same software, parameters, dependency versions and complete protocol. Calibration, reference sampling and searches use separate random streams.
- `length`, `steps`, and `restarts` control sequence searches. Additional family, cross-environment and specificity options are described in the interactive schema.

In interfacial environments, **each sequence is rotated by its own hydrophobic moment**, pointing that moment at the center of the apolar sector. This includes every search proposal, query, calibration decoy and random reference sequence. If the moment is numerically zero, the preset phase is retained; homogeneous media do not require alignment. Each environment's search and reported score share one calibrated Hamiltonian and one reference distribution. A mutation can change the phase of the entire sequence, so its environmental score must be reevaluated under that new phase.

From 1.0.0rc5, `design_geometry` is a compatibility copy of the containing score's `energy`, `z_score`, `percentile_lower_is_better` and `breakdown`, plus the actual `environment.wheel_phase_deg` and orientation metadata; it does not represent a second scoring convention. Family ranking, cross-environment objectives and specificity margins use these same aligned scores. In 1.0.0rc4 and earlier, the field instead identified the separate fixed-phase search objective; retain the release version when interpreting saved results.

Alignment is a geometric scoring rule, not minimization of the complete Hamiltonian over rotation angle or a prediction of equilibrium orientation. Lower scores indicate better compatibility with the assumed helical state within the specified protocol. The scores are dimensionless comparisons and do not establish antimicrobial activity or membrane selectivity.

In 1.0.0rc6, the neighbor component uses `−mean(M1) + mean(M3) + mean(M4)` before calibration and weighting. Each mean covers the L−k available sequence pairs at its separation. Larger Nacar helical pairing propensities therefore lower the model score, while M3/M4 retain their published energy signs. The raw tables are preserved. The convention `negative_m1_plus_m3_plus_m4` and its fixed internal coefficients are part of the parameter-set checksum and calibration-cache metadata; calibration statistics from the positive-M1 convention are recomputed.

The Penetration endpoint scans 21 exposure settings in 5% increments. Zero percent is a homogeneous polar environment; 100% is a homogeneous apolar environment. Intermediate settings use an apolar-sector half-width of `40 + 1.1 × percent` degrees and the same hydrophobic-moment alignment rule. This percentage describes an **angular exposure mask**, not a physical insertion depth.

## Examples with curl

Start the server as described in the [README](../README.md#installation), then run these commands from the repository root. The example files use short searches and only 100 calibration sequences to demonstrate the interface quickly; these settings do not establish sampling convergence for scientific use.

**Score a short sequence:**

```bash
curl --fail-with-body http://127.0.0.1:8000/api/score \
  -H 'Content-Type: application/json' \
  --data @examples/score.json > score-result.json
```

Inspect `energy`, `z_score`, and the six standardized components in `breakdown`. The response's `environment.wheel_phase_deg` records the phase used for this sequence. `calibration.orientation` identifies its orientation protocol.

**Design a 12-residue candidate for an anionic membrane environment:**

```bash
curl --fail-with-body http://127.0.0.1:8000/api/design \
  -H 'Content-Type: application/json' \
  --data @examples/design.json > design-result.json
```

The example allows A, L, K, E, G, F and S and runs one 1,000-step search. Read `sequence` for the candidate and `energy` and `z_score` for its aligned score. `design_geometry.energy` and `.z_score` duplicate these values for compatibility; the objective optimized in this single-environment search is `energy`.

**Compare magainin 2 in an anionic interface and a polar medium:**

```bash
curl --fail-with-body http://127.0.0.1:8000/api/compare \
  -H 'Content-Type: application/json' \
  --data @examples/compare.json > compare-result.json
```

The response contains the complete scores under `environment_a` and `environment_b`, each with its own calibration and reference distribution. Their absolute differences are reported as `energy_gap` and `z_score_gap`. These are model comparisons; they are not experimental activity or binding measurements.

**Scan angular exposure for the same sequence:**

```bash
curl --fail-with-body http://127.0.0.1:8000/api/optimal-penetration \
  -H 'Content-Type: application/json' \
  --data @examples/penetration.json > penetration-result.json
```

Read the 21 records in `profile` to plot score versus exposure. `best_penetration_percent` minimizes the model Z-score among those settings, and `best_score` contains that setting's complete score. It does not predict a physical insertion depth.

## Python client

The supplied [client](../examples/api_client.py) uses only Python's standard library. It reads the same JSON requests and saves the complete response:

```bash
python examples/api_client.py score --output score-result.json
python examples/api_client.py design --output design-result.json
python examples/api_client.py compare --output compare-result.json
python examples/api_client.py penetration --output penetration-result.json
```

Choose another local installation with `--base-url`, for example `--base-url http://127.0.0.1:8080`. Check that installation's software version before using or comparing its numerical results. The client identifies its requests with `User-Agent: Helix-Design-Studio-API/1.0`.

## Handling errors

Invalid requests return a non-success HTTP status and a JSON `detail` field identifying the issue. Read the schema for the serving release's limits. A busy server or a calculation exceeding its execution limit also returns an error. The Python client and `curl --fail-with-body` report HTTP failures. A reverse proxy can also reject a request before it reaches the API; such a response may contain HTML rather than a JSON `detail`. Preserve its HTTP status and message when reporting an access problem. For streaming endpoints, inspect every event and require a `final` event before treating the run as complete.
