## Additional file 1. Evaluated software source and dependencies for "Helix Design Studio: designing antimicrobial peptide candidates for membrane environments": api_client.py

```
"""Run one of the supplied JSON examples using Python's standard library."""

import argparse
import json
import sys
from pathlib import Path
from urllib.error import HTTPError, URLError
from urllib.request import Request, urlopen


ENDPOINTS = {
    "score": "/api/score",
    "design": "/api/design",
    "compare": "/api/compare",
    "penetration": "/api/optimal-penetration",
}


def main():
    parser = argparse.ArgumentParser(description=__doc__)
    parser.add_argument("operation", choices=ENDPOINTS, nargs="?", default="score")
    parser.add_argument("--base-url", default="http://127.0.0.1:8000")
    parser.add_argument("--output", type=Path, help="Save the complete JSON response, including provenance.")
    args = parser.parse_args()
    payload_path = Path(__file__).with_name(f"{args.operation}.json")
    payload = json.loads(payload_path.read_text(encoding="utf-8"))
    request = Request(
        args.base_url.rstrip("/") + ENDPOINTS[args.operation],
        data=json.dumps(payload).encode("utf-8"),
        headers={
            "Content-Type": "application/json",
            "Accept": "application/json",
            "User-Agent": "Helix-Design-Studio-API/1.0",
        },
        method="POST",
    )
    try:
        with urlopen(request, timeout=300) as response:
            result = json.load(response)
    except HTTPError as exc:
        print(f"HTTP {exc.code}: {exc.read().decode('utf-8', errors='replace')}", file=sys.stderr)
        return 1
    except URLError as exc:
        print(f"Could not reach the API: {exc.reason}", file=sys.stderr)
        return 1
    rendered = json.dumps(result, indent=2, ensure_ascii=False) + "\n"
    if args.output:
        args.output.write_text(rendered, encoding="utf-8")
        print(f"Saved {args.output}")
    else:
        print(rendered, end="")
    return 0


if __name__ == "__main__":
    raise SystemExit(main())
```
