## Additional file 1. Evaluated software source and dependencies for "Helix Design Studio: designing antimicrobial peptide candidates for membrane environments": 3Dmol.txt

3Dmol.js incorporates code from GLmol, Three.js, and jQuery and
is licensed under a BSD-3-Clause license.

* 3Dmol.js
	Copyright (c) 2014, University of Pittsburgh and contributors
	All rights reserved.

	Redistribution and use in source and binary forms, with or without 
	modification, are permitted provided that the following conditions are met:

	1. Redistributions of source code must retain the above copyright 
	notice, this list of conditions and the following disclaimer.

	2. Redistributions in binary form must reproduce the above copyright 
	notice, this list of conditions and the following disclaimer in the 
	documentation and/or other materials provided with the distribution.

	3. Neither the name of the copyright holder nor the names of its 
	contributors may be used to endorse or promote products derived from 
	this software without specific prior written permission.

	THIS SOFTWARE IS PROVIDED BY THE COPYRIGHT HOLDERS AND CONTRIBUTORS 
	"AS IS" AND ANY EXPRESS OR IMPLIED WARRANTIES, INCLUDING, BUT NOT 
	LIMITED TO, THE IMPLIED WARRANTIES OF MERCHANTABILITY AND FITNESS FOR A 
	PARTICULAR PURPOSE ARE DISCLAIMED. IN NO EVENT SHALL THE COPYRIGHT
	HOLDER OR CONTRIBUTORS BE LIABLE FOR ANY DIRECT, INDIRECT, INCIDENTAL, 
	SPECIAL, EXEMPLARY, OR CONSEQUENTIAL DAMAGES (INCLUDING, BUT NOT LIMITED 
	TO, PROCUREMENT OF SUBSTITUTE GOODS OR SERVICES; LOSS OF USE, DATA, OR 
	PROFITS; OR BUSINESS INTERRUPTION) HOWEVER CAUSED AND ON ANY THEORY OF 
	LIABILITY, WHETHER IN CONTRACT, STRICT LIABILITY, OR TORT (INCLUDING 
	NEGLIGENCE OR OTHERWISE) ARISING IN ANY WAY OUT OF THE USE OF THIS 
	SOFTWARE, EVEN IF ADVISED OF THE POSSIBILITY OF SUCH DAMAGE.

* GLmol
 GLmol - Molecular Viewer on WebGL/Javascript (0.47)
  (C) Copyright 2011-2012, biochem_fan
      License: dual license of MIT or LGPL3

  Contributors:
    Robert Hanson for parseXYZ, deferred instantiation


* Three.js 
     https://github.com/mrdoob/three.js

     Copyright (c) 2010-2012 three.js Authors. All rights reserved.

     Permission is hereby granted, free of charge, to any person obtaining a copy
     of this software and associated documentation files (the "Software"), to deal
     in the Software without restriction, including without limitation the rights
     to use, copy, modify, merge, publish, distribute, sublicense, and/or sell
     copies of the Software, and to permit persons to whom the Software is
     furnished to do so, subject to the following conditions:

     The above copyright notice and this permission notice shall be included in
     all copies or substantial portions of the Software.

     THE SOFTWARE IS PROVIDED "AS IS", WITHOUT WARRANTY OF ANY KIND, EXPRESS OR
     IMPLIED, INCLUDING BUT NOT LIMITED TO THE WARRANTIES OF MERCHANTABILITY,
     FITNESS FOR A PARTICULAR PURPOSE AND NONINFRINGEMENT. IN NO EVENT SHALL THE
     AUTHORS OR COPYRIGHT HOLDERS BE LIABLE FOR ANY CLAIM, DAMAGES OR OTHER
     LIABILITY, WHETHER IN AN ACTION OF CONTRACT, TORT OR OTHERWISE, ARISING FROM,
     OUT OF OR IN CONNECTION WITH THE SOFTWARE OR THE USE OR OTHER DEALINGS IN
     THE SOFTWARE.

* jQuery
     http://jquery.org/

     Copyright (c) 2011 John Resig

     Permission is hereby granted, free of charge, to any person obtaining
     a copy of this software and associated documentation files (the
     "Software"), to deal in the Software without restriction, including
     without limitation the rights to use, copy, modify, merge, publish,
     distribute, sublicense, and/or sell copies of the Software, and to
     permit persons to whom the Software is furnished to do so, subject to
     the following conditions:

     The above copyright notice and this permission notice shall be
     included in all copies or substantial portions of the Software.

     THE SOFTWARE IS PROVIDED "AS IS", WITHOUT WARRANTY OF ANY KIND,
     EXPRESS OR IMPLIED, INCLUDING BUT NOT LIMITED TO THE WARRANTIES OF
     MERCHANTABILITY, FITNESS FOR A PARTICULAR PURPOSE AND
     NONINFRINGEMENT. IN NO EVENT SHALL THE AUTHORS OR COPYRIGHT HOLDERS BE
     LIABLE FOR ANY CLAIM, DAMAGES OR OTHER LIABILITY, WHETHER IN AN ACTION
     OF CONTRACT, TORT OR OTHERWISE, ARISING FROM, OUT OF OR IN CONNECTION
     WITH THE SOFTWARE OR THE USE OR OTHER DEALINGS IN THE SOFTWARE.
