## Additional file 1. Evaluated software source and dependencies for "Helix Design Studio: designing antimicrobial peptide candidates for membrane environments": README.md

# Helix Design Studio

[Web server](https://helix.simbioslab.com/) · [Method article](https://doi.org/10.1021/acs.jctc.6c01278) · [Citation](CITATION.cff)

Helix Design Studio scores and designs peptide sequences in a **predefined alpha-helical state**, with polar, apolar and membrane-like environments. Its browser interface supports sequence scoring, single and family design, environmental comparisons, specificity design, exposure scans and idealized PDB export.

**Current version: 1.0.0rc6**, a release candidate for evaluation. Searches and sequence scoring use the same hydrophobic-moment alignment rule. The neighbor score rewards larger helical pairing propensities through a negative M1 contribution. The published parameter tables and model adaptations are described below.

## Installation

Use **Python 3.12** and the pinned dependencies. Interactive molecular rendering requires a WebGL-capable browser; the viewer is included in the application. Local execution has been tested on macOS with Python 3.12.

```bash
git clone https://github.com/SIMBIOS-USC/helix-design-studio-web.git
cd helix-design-studio-web
git checkout v1.0.0rc6
python3.12 -m venv .venv
source .venv/bin/activate
python -m pip install -r requirements.txt
python -m uvicorn app.main:app --host 127.0.0.1 --port 8000
```

On Windows, activate the environment with `.venv\Scripts\activate`.

Open **http://127.0.0.1:8000**. The interface includes a Guide and examples. The first calculation for a calibration configuration samples random sequences and caches the component statistics for subsequent requests.

Opening `app/static/index.html` directly redirects to this local address. Keep the server running on port 8000; calculations require the server.

## Docker

```bash
docker build -t helix-design-studio .
docker run --rm -p 8000:8000 helix-design-studio
```

The image runs as an unprivileged user and defines a health check. Set `PORT` to change its internal listening port and adjust the port mapping accordingly, `WEB_CONCURRENCY` to run several worker processes, and `FORWARDED_ALLOW_IPS` to the address of the reverse proxy whose `X-Forwarded-For` header should be trusted. Calibration caches and optional usage logs are written to `/tmp/helix-cache` and `/tmp/helix-usage`; mount volumes there (for example `-v helix-cache:/tmp/helix-cache`) to keep them across container restarts. Container startup and the scoring example have been verified on Linux.

## Reproducible calculations

Download the complete JSON calculation records from the browser interface and retain them with reported results. Each record includes a `provenance` block with the software version, parameter-set checksum, complete request, seed protocol, and Python and NumPy versions. The parameter-set checksum identifies the raw tables, residue scales and neighbor-scoring convention. Scientific applications require calibration and reference sample sizes appropriate to the question. The residue alphabet order determines the random streams used for calibration and search. Changes between versions and their measured effects are listed in the [changelog](CHANGELOG.md).

Check the software version in the downloaded record before comparing numerical results from the public server with a specified release. Technical documentation and local request examples for the backend are provided in [docs/API.md](docs/API.md).

## Reproducing a calculation in the browser

Each calculation panel provides Advanced settings for the calibration sample count and random seed. Interfacial calculations also accept an optional exact apolar-sector half-width in degrees; enter 90 to reproduce the standard half-plane geometry independently of the exposure slider. Exported records preserve these inputs. Changing the exposure slider clears the angle override; homogeneous polar/apolar endpoints retain their preset geometry.

## Model scope and limitations

- Scores measure compatibility with an assumed alpha-helical state. They do not establish folding, binding, membrane insertion, biological activity or experimental stability.
- In interfacial environments, every search proposal (Design, Family, Cross-Design and Specificity) and every scored sequence is rotated so that its own hydrophobic moment points to the center of the apolar sector. Calibration decoys and random reference sequences are oriented individually by the same rule. A sequence with a numerically zero moment retains the preset phase; rotation has no effect in homogeneous media. This alignment is a prescribed orientation rule, not a search for the angle that minimizes the complete Hamiltonian or a prediction of equilibrium membrane orientation.
- Searches, rankings and specificity margins use this aligned Hamiltonian and its matching calibration and reference distributions. A residue substitution can change the orientation of the complete helix, so the environmental contributions depend on the whole sequence. Design results report the same score used during the search; the retained `design_geometry` field duplicates that score and records its actual orientation for compatibility with existing result readers.
- The helix-neighbor score is `−mean(M1) + mean(M3) + mean(M4)`, where the mean for separation k is over the L−k available sequence pairs. Larger M1 propensities therefore lower the minimized score; negative M3/M4 interaction contributions also lower it. The combined `helix_neigh` component is standardized against calibration sequences and given its configured component weight. The internal coefficients −1, +1 and +1 define a fixed empirical combination of dimensionless relative propensities and numerical side-chain interaction parameters expressed in kcal/mol. It is a dimensionless comparative descriptor after standardization, rather than an absolute thermodynamic energy. All tables are indexed by residue label. Miyazawa–Jernigan coefficients are applied to residues i and i+1…i+4.
- M1 uses the 400 values in Nacar (2020), Figure 3. The odds ratios express enrichment of hydrogen-bonded n,n+4 pairs over a composition-based random background in globular proteins. A larger ratio indicates greater helical enrichment. The software subtracts these values in its empirical i,i+1 score; a zero entry contributes zero to that M1 sum. Reusing the source at i,i+1 remains a model adaptation rather than evidence for adjacent-residue pair propensities. Source columns identify the earlier carbonyl acceptor and rows the later amide donor. The runtime table is transposed so that rows identify residue i and columns residue i+1.
- M3 and M4 use the 324 values per separation in Muñoz–Serrano (1994), Table 2, in kcal/mol. The published table contains 18 residues and omits Cys and Pro. The application assigns zero contribution to any M3/M4 pair containing Cys or Pro (76 additional entries per matrix); these are model defaults, not measured zero interactions. This convention does not remove the separate single-residue helix propensity for Pro.
- The membrane-charge term acts only on water-facing residues of charged interfaces; it is zero in neutral interfaces and homogeneous media.
- PDB export constructs an ideal helix; it does not predict or minimize a structure.
- The Penetration scan changes an angular exposure mask. Its coordinate is not a physical insertion depth or a free-energy reaction coordinate.
- Calibration and reference distributions depend on sequence length, alphabet, environment, orientation rule, sample sizes and seeds. Scores from different protocols are not directly interchangeable.

Source citations, axis definitions and the complete comparison procedure are given in [parameter provenance](THIRD_PARTY_NOTICES.md#parameter-provenance).

The [archived 1.0.0rc2 Table 1 rerun](tools/data/article_table1_comparison_1.0.0rc2.json) evaluates the seven exact sequence strings and fourteen rounded Z-scores of the published method article with the positive-M1 convention used by that release. Its mean and maximum absolute Z-score differences are 0.303771 and 0.439849. These historical measurements describe that specific release and rerun protocol. The [changelog](CHANGELOG.md) records the subsequent alignment and M1-scoring changes; archived numerical results must be associated with their recorded software version.

### Parameter table checksums

SHA-256 of the parameter tables in `Code/core/resources/`, so a result's parameter set can be identified independently of the checksum recorded in each result's `provenance` block:

```text
mj_matrix.txt         53af4769e5cd9c185938a5f2dd23b0374d8f4dc5e7ca2e32e825b8f79f2c6171
helix_pairs_prop.txt  266410b1335e1de1341e39e031ecd23054e174d3017322d51be1bb20c4e19684
helix_sd_i_3.txt      ac0fcc7c17ffeaeea6664906b5473c1c9ddd2f2145d0d75626e674b7eb0505c5
helix_sd_i_4.txt      492bb545cc304bd51a417ad7ffa4e9cac6babf8cf4b256a828406551ccab4897
```

## Tests

```bash
python -m pip install -r requirements-dev.txt
python -m unittest discover -s tests -v
```

The tests cover the computational API, streaming responses, static assets, idealized PDB export, input validation, request limits, usage logging, calibration caching, label-based parameter indexing, literature residue scales, all published neighbor parameters and consistency between workflows. They do not establish sampling convergence or experimental validity.

To check the parameter data independently of the API, run:

```bash
python tools/verify_parameter_sources.py
```

This compares all 400 M1 values and all 324 published values in each of M3/M4, using the source axis labels. The 76 Cys/Pro defaults per M3/M4 matrix are checked separately and are not counted as published data. Source CSVs, citations and checksums are included in `tools/data/parameter_sources/`.

The browser-side partial-result export tests additionally use Node.js 18 or later with its built-in test runner (no npm packages required):

```bash
node --test tests/test_partial_exports.js
```

## Configuration and data

| Variable | Default | Purpose |
| --- | --- | --- |
| `HELIX_CACHE_DIR` | `helix-design-studio-cache` in the OS temporary directory | Generated calibration statistics (private to the service user). |
| `HELIX_CACHE_MAX_FILES` | `2000` | Maximum number of cached calibrations; the least recently used are removed. |
| `HELIX_MAX_CONCURRENT_JOBS` | `4` | Calculations run at the same time per server process; further requests receive HTTP 503. |
| `HELIX_MAX_JOB_SECONDS` | `1800` | Time limit for a single calculation. |
| `HELIX_CORS_ORIGINS` | `https://simbios.usc.es` | Comma-separated origins allowed to call the API from other sites; empty disables CORS. |
| `HELIX_USAGE_LOGGING` | `0` | Set to `1` to enable local usage-event logging. |
| `HELIX_VAR_DIR` | A directory in the OS temporary directory | Optional usage logs; choose a persistent directory if required. |
| `HELIX_LOG_HASH_KEY` | Random per process | Secret key for the hashes stored in usage logs; set it to keep hashes comparable across restarts. |
| `HELIX_CODE_DIR` | This repository's `Code/` | Override the location of scoring modules and parameter tables (`QFOLD_CODE_DIR` is accepted as a legacy name). |
| `PORT` | `8000` | Container listening port. |

With logging disabled, the application creates no usage-event log or persistent browser visitor identifier. The visit and run counters shown when logging is enabled are informational: they are reported by browsers and are not protected against inflation. When enabled, events include a random browser identifier, keyed hashes (HMAC-SHA256) of IP address, user agent and sequences, request settings, truncated origin/referrer, timing, HTTP status and error messages. Hashes do not guarantee anonymity. Events are stored on the server; server/proxy access logs are configured separately.

`GET /api/health` reports service configuration and, when logging is enabled, recent server-side failures (HTTP 5xx); it returns 503 while such failures are present in the requested window. Calibration caches are keyed by the parameter tables and settings; they are disposable and can be cleared at any time.

For a public deployment, run the service behind a reverse proxy that terminates TLS and applies per-client rate limits, start uvicorn with `--proxy-headers --forwarded-allow-ips=<proxy address>` so that client addresses are taken from the trusted proxy, and use `--workers` to serve several calculations in parallel.

## Source layout

- `app/`: API, search workflows and static browser interface.
- `Code/`: scoring, calibration and four parameter tables.
- `tests/`, `examples/` and `docs/`: automated checks, API request examples, a Python client and the API guide.
- `tools/measure_changes.py`: measures the numerical effect of each model change between versions, with a separate control that changes only the search random stream.
- `tools/reproduce_article_table1.py`: reruns the method article's Table 1 sequences with the current model. `--include-positive-m1` adds an explicitly labelled earlier-sign control; each variant records its formula, version and parameter checksum. The optional negative-log control rejects the published zeros without smoothing. `tools/data/` retains the separately versioned historical comparisons.
- `tools/verify_parameter_sources.py` and `tools/data/parameter_sources/`: verify runtime neighbor parameters against the transcribed primary tables, with explicit source axes and model extensions.
- `LICENSES/`, `THIRD_PARTY_NOTICES.md`, `CITATION.cff`: licenses, attribution and citation metadata.

## Citation and license

Please cite the **software version or commit** used and the underlying method:

Daniel Conde-Torres, Rebeca García-Fandiño and Ángel Piñeiro. *Environment-Conditioned Design of α-Helical Peptides*. Journal of Chemical Theory and Computation **22** (18), 9776–9789 (2026). [doi:10.1021/acs.jctc.6c01278](https://doi.org/10.1021/acs.jctc.6c01278).

Software contributors are listed in [CITATION.cff](CITATION.cff). Original project code is distributed under the [MIT license](LICENSE). Included components retain their own terms; see [third-party notices](THIRD_PARTY_NOTICES.md). Institutional names and logos are excluded from the MIT grant. Citation is requested as scholarly practice, not as an additional license condition.

Report problems through the [issue tracker](https://github.com/SIMBIOS-USC/helix-design-studio-web/issues), including the commit, request parameters and steps to reproduce. Omit private sequences and usage logs from public reports.
