## Additional file 1. Evaluated software source and dependencies for "Helix Design Studio: designing antimicrobial peptide candidates for membrane environments": test_calculation_settings.js

```
// Run with: node --test tests/test_calculation_settings.js
const test = require('node:test');
const assert = require('node:assert/strict');
const fs = require('node:fs');
const path = require('node:path');
const vm = require('node:vm');

const source = fs.readFileSync(path.join(__dirname, '../app/static/app.js'), 'utf8');
const names = [
  'isInterfacialPreset', 'penetrationPercentToHalfWidth', 'penetrationDescription',
  'buildEnvironmentFromPresetAndPenetration', 'buildEnvironmentFromFormData',
  'calculationSettingsFromFormData', 'calculationSettingsHtml',
  'initializeCalculationSettings', 'syncExactHalfWidthControls', 'parseResidues',
];
const helpers = names.map((name) => {
  const match = source.match(new RegExp(`^function ${name}\\([^]*?^}`, 'm'));
  assert.ok(match, `Missing application function: ${name}`);
  return match[0];
}).join('\n');
const plain = (value) => JSON.parse(JSON.stringify(value));
const formData = (values) => ({ get: (name) => values[name] ?? null });

function context(overrides = {}) {
  const ctx = vm.createContext(overrides);
  vm.runInContext(helpers, ctx);
  return ctx;
}

test('exact angular settings coexist with exposure endpoints and homogeneous environments', () => {
  const ctx = context();
  for (const [percent, expected] of [[0, 'homogeneous_polar'], [100, 'homogeneous_apolar']]) {
    assert.deepEqual(plain(ctx.buildEnvironmentFromPresetAndPenetration('interfacial_neg', percent, '90')), { preset: expected });
  }
  for (const preset of ['homogeneous_polar', 'homogeneous_apolar']) {
    assert.deepEqual(plain(ctx.buildEnvironmentFromPresetAndPenetration(preset, 50, '90')), { preset });
  }
  assert.equal(ctx.buildEnvironmentFromPresetAndPenetration('interfacial_neg', 50, '90').wheel_halfwidth_deg, 90);
  assert.equal(ctx.buildEnvironmentFromPresetAndPenetration('interfacial_neg', 25, '').wheel_halfwidth_deg, 67.5);
  assert.equal(ctx.buildEnvironmentFromPresetAndPenetration('interfacial_neg', 50).wheel_halfwidth_deg, 95);
  assert.throws(() => ctx.buildEnvironmentFromPresetAndPenetration('interfacial_neg', 50, '181'), /between 0 and 180/);
});

test('sampling controls accept seed zero and the full backend ranges', () => {
  const ctx = context();
  assert.deepEqual(plain(ctx.calculationSettingsFromFormData(formData({ n_decoys: '100', seed: '0' }))), { n_decoys: 100, seed: 0 });
  assert.deepEqual(plain(ctx.calculationSettingsFromFormData(formData({ n_decoys: '10000', seed: '4294967295' }))), { n_decoys: 10000, seed: 4294967295 });
  assert.deepEqual(plain(ctx.calculationSettingsFromFormData(formData({}), 10000)), { n_decoys: 10000, seed: 42 });
  for (const bad of ['99', '10001', '2000.5', 'abc']) {
    assert.throws(() => ctx.calculationSettingsFromFormData(formData({ n_decoys: bad })), /calibration sample size/);
  }
  for (const bad of ['-1', '4294967296', '1.5', 'abc']) {
    assert.throws(() => ctx.calculationSettingsFromFormData(formData({ seed: bad })), /random seed/);
  }
});

test('all calculation forms receive controls inside their existing advanced settings', () => {
  const forms = new Map();
  const ctx = context({
    $: (selector, root) => {
      if (root) return null;
      const match = selector.match(/^#([a-z]+)-form \.advanced-settings \.fields-grid$/);
      if (!match) return null;
      const grid = { insertAdjacentHTML: (_position, html) => { grid.html = html; } };
      forms.set(match[1], grid);
      return grid;
    },
    $$: () => [],
  });
  ctx.initializeCalculationSettings();
  assert.deepEqual([...forms.keys()].sort(), ['compare', 'cross', 'design', 'family', 'penetration', 'score', 'specificity']);
  for (const [name, grid] of forms) {
    assert.match(grid.html, new RegExp(`name="n_decoys"[^>]*value="${name === 'specificity' ? 10000 : 2000}"`));
    assert.match(grid.html, /name="seed"[^>]*value="42"/);
    if (name === 'penetration') assert.doesNotMatch(grid.html, /data-exact-halfwidth/);
    else assert.match(grid.html, /data-exact-halfwidth/);
  }
});

test('exact-angle controls activate only for interfaces at intermediate exposures', () => {
  const display = {};
  const slider = { value: '50' };
  const preset = { value: 'interfacial_neg' };
  const form = { id: 'score-form' };
  const input = {
    dataset: { penetrationName: 'penetration_depth', presetName: 'preset', exposureScope: 'score' },
    closest: () => form, value: '90', validity: { valid: true },
  };
  const ctx = context({
    $: (selector) => selector === '[name="penetration_depth"]' ? slider
      : selector === '[name="preset"]' ? preset : display,
    $$: (selector) => selector === '[data-exact-halfwidth]' ? [input] : [],
  });
  ctx.syncExactHalfWidthControls();
  assert.equal(input.disabled, false);
  assert.equal(display.textContent, 'Exact apolar half-width: 90°');
  preset.value = 'homogeneous_polar';
  ctx.syncExactHalfWidthControls();
  assert.equal(input.disabled, true);
  preset.value = 'interfacial_neg';
  for (const value of ['0', '100']) {
    slider.value = value;
    ctx.syncExactHalfWidthControls();
    assert.equal(input.disabled, true);
  }
  form.id = 'specificity-form';
  slider.value = '25';
  preset.value = 'homogeneous_polar';
  ctx.$$ = (selector) => selector === '[data-exact-halfwidth]' ? [input]
    : [{ checked: true, disabled: false, value: 'interfacial_neu' }];
  ctx.syncExactHalfWidthControls();
  assert.equal(input.disabled, false, 'An interfacial competitor uses the shared exact angle');
});

function boundCalculations(ctx) {
  const builders = new Map();
  ctx.bindForm = (formId, _resultId, builder) => builders.set(formId, builder);
  ctx.renderScore = () => '';
  ctx.renderCompare = () => '';
  ctx.renderOptimalPenetration = () => '';
  ctx.postJSON = async (_url, payload) => payload;
  const begin = source.indexOf("bindForm('score-form'");
  const end = source.indexOf("const specificityForm = $('#specificity-form');", begin);
  vm.runInContext(source.slice(begin, end), ctx);
  return builders;
}

test('Score, Design, Compare, Cross and Penetration submit the selected reproducibility settings', async () => {
  const builders = boundCalculations(context());
  const values = {
    sequence: 'LKKL', residues: 'A,R,N,D,C,Q,E,G,H,I,L,K,M,F,P,S,T,W,Y,V', length: '4',
    preset: 'interfacial_neg', preset_a: 'interfacial_neg', preset_b: 'interfacial_neu',
    penetration_depth: '50', penetration_depth_a: '50', penetration_depth_b: '25',
    exact_halfwidth_penetration_depth: '90', exact_halfwidth_penetration_depth_a: '90',
    exact_halfwidth_penetration_depth_b: '70.25', n_decoys: '2500', seed: '7',
  };
  for (const name of ['score', 'design', 'compare', 'cross', 'penetration']) {
    const data = await builders.get(`${name}-form`)(formData(values), null);
    assert.equal(data.n_decoys, 2500, name);
    assert.equal(data.seed, 7, name);
    assert.equal(data.residues.join(''), values.residues.replaceAll(',', ''), 'Preserve the residue order');
    if (name === 'score' || name === 'design') assert.equal(data.environment.wheel_halfwidth_deg, 90);
    if (name === 'compare' || name === 'cross') {
      assert.equal(data.environment_a.wheel_halfwidth_deg, 90);
      assert.equal(data.environment_b.wheel_halfwidth_deg, 70.25);
    }
    if (name === 'penetration') {
      assert.deepEqual(plain(data.environment), { preset: 'interfacial_neg' }, 'The scan selects its own exposure at each point');
      assert.equal(data.percent_min, 0);
      assert.equal(data.percent_max, 100);
    }
  }
});
```
