## Additional file 1. Evaluated software source and dependencies for "Helix Design Studio: designing antimicrobial peptide candidates for membrane environments": test_neighbor_scoring.py

```
"""Scientific direction and provenance of the empirical neighbor score."""

import hashlib
import json
from pathlib import Path
import sys
import tempfile
import unittest
from unittest.mock import patch

import numpy as np


ROOT = Path(__file__).resolve().parents[1]
sys.path.insert(0, str(ROOT / "Code"))
from core import hamiltonian_builder as model  # noqa: E402


class NeighborScoringTests(unittest.TestCase):
    def bare_builder(self, length):
        """Use the actual parameter loaders without constructing a calibration cache."""
        builder = model.HamiltonianBuilder.__new__(model.HamiltonianBuilder)
        builder.L = length
        builder.amino_acids = list("ARNDCQEGHILKMFPSTWYV")
        builder.n_aa = len(builder.amino_acids)
        builder.kwargs = {}
        builder.stats_cache_version = 5
        builder.align_phase = False
        builder._init_properties()
        builder._load_matrices_from_files()
        return builder

    def codes(self, builder, sequence):
        return [builder.amino_acids.index(residue) for residue in sequence]

    def test_larger_m1_propensity_lowers_raw_and_standardized_score(self):
        # L=2 isolates the nearest-neighbor contribution from M3/M4.
        builder = self.bare_builder(2)
        alanine = builder._raw_helix_neigh(self.codes(builder, "AA"))
        glycine = builder._raw_helix_neigh(self.codes(builder, "GG"))
        self.assertAlmostEqual(alanine, -2.734)
        self.assertAlmostEqual(glycine, -0.370)
        self.assertLess(alanine, glycine)
        builder.stats = {"helix_neigh": (-1.0, 0.5)}
        self.assertLess(
            1.5 * builder._apply_z_score("helix_neigh", alanine),
            1.5 * builder._apply_z_score("helix_neigh", glycine),
        )

    def test_zero_source_entry_gives_no_m1_reward(self):
        builder = self.bare_builder(2)
        self.assertEqual(builder._raw_helix_neigh(self.codes(builder, "AP")), 0.0)
        self.assertLess(builder._raw_helix_neigh(self.codes(builder, "AA")), 0.0)
        # Source zeros remain zeros; only their additive scoring coefficient changes.
        self.assertEqual(builder.M1[builder.amino_acids.index("A"), builder.amino_acids.index("P")], 0.0)

    def test_m3_m4_keep_their_favorable_negative_contributions(self):
        builder = self.bare_builder(4)
        builder.M1.fill(0.0)
        self.assertAlmostEqual(builder._raw_helix_neigh(self.codes(builder, "IAAF")), -0.450)
        builder = self.bare_builder(5)
        builder.M1.fill(0.0)
        builder.M3.fill(0.0)
        self.assertAlmostEqual(builder._raw_helix_neigh(self.codes(builder, "WAAAH")), -0.900)

    def test_scoring_convention_changes_public_and_matrix_fingerprints(self):
        builder = self.bare_builder(5)
        public_fingerprint = model.parameter_set_sha256()
        matrix_fingerprint = builder._parameters_fingerprint()
        with patch.object(model, "NEIGHBOR_SCORING_CONVENTION", "positive_m1_plus_m3_plus_m4"):
            self.assertNotEqual(model.parameter_set_sha256(), public_fingerprint)
            self.assertNotEqual(builder._parameters_fingerprint(), matrix_fingerprint)
        with patch.dict(model.NEIGHBOR_MATRIX_WEIGHTS, {1: 1.0}):
            self.assertNotEqual(model.parameter_set_sha256(), public_fingerprint)
            self.assertNotEqual(builder._parameters_fingerprint(), matrix_fingerprint)

    def test_previous_positive_m1_cache_is_not_reused(self):
        builder = self.bare_builder(5)
        with tempfile.TemporaryDirectory(prefix="helix-neighbor-cache-") as directory:
            builder.kwargs = {"stats_cache_dir": directory, "n_decoys": 100, "zscore_seed": 42}
            current_metadata = builder._stats_cache_payload(100)
            self.assertEqual(current_metadata["stats_cache_version"], 5)
            self.assertEqual(current_metadata["neighbor_score_convention"], model.NEIGHBOR_SCORING_CONVENTION)
            legacy_metadata = dict(current_metadata)
            legacy_metadata["stats_cache_version"] = 4
            legacy_metadata.pop("neighbor_score_convention")
            legacy_metadata.pop("neighbor_matrix_weights")
            legacy_digest = hashlib.sha256()
            for matrix in (builder.mj_matrix, builder.M1, builder.M3, builder.M4):
                legacy_digest.update(np.ascontiguousarray(matrix, dtype=np.float64).tobytes())
            legacy_metadata["parameters_sha256"] = legacy_digest.hexdigest()
            legacy_key = hashlib.sha256(json.dumps(
                legacy_metadata, sort_keys=True, separators=(",", ":")
            ).encode("utf-8")).hexdigest()
            legacy_path = Path(directory) / f"zscore_stats_{legacy_key}.json"
            legacy_path.write_text(json.dumps({
                "metadata": legacy_metadata,
                "stats": {term: {"mu": 777.0, "sigma": 1.0} for term in builder.TERMS},
            }), encoding="utf-8")
            self.assertIsNone(builder._read_stats_cache(str(legacy_path), 100))
            calibrated = builder._load_or_calibrate_z_scores(100)
            self.assertNotEqual(calibrated["helix_neigh"][0], 777.0)
            current_path = Path(builder._stats_cache_path(100))
            self.assertNotEqual(current_path, legacy_path)
            self.assertTrue(current_path.is_file())
            saved_metadata = json.loads(current_path.read_text())["metadata"]
            self.assertEqual(saved_metadata["neighbor_score_convention"], model.NEIGHBOR_SCORING_CONVENTION)
            self.assertTrue(all(sigma > 0 for _, sigma in calibrated.values()))


if __name__ == "__main__":
    unittest.main()
```
