## Additional file 1. Evaluated software source and dependencies for "Helix Design Studio: designing antimicrobial peptide candidates for membrane environments": test_parameter_sources.py

```
"""Regression against published neighbor parameters, independent of API scoring."""

from decimal import Decimal
from pathlib import Path
import shutil
import tempfile
import unittest

from tools import verify_parameter_sources as sources


class PublishedParameterTests(unittest.TestCase):
    def test_all_published_cells_and_model_extensions(self):
        result = sources.verify()
        self.assertTrue(result["verified"], result)
        for name, count, extension in [("M1", 400, 0), ("M3", 324, 76), ("M4", 324, 76)]:
            with self.subTest(matrix=name):
                matrix = result["matrices"][name]
                self.assertEqual(matrix["published_cells_matching"], count)
                self.assertEqual(matrix["model_zero_extension_cells"], extension)
                self.assertFalse(matrix["published_cell_differences"])
                self.assertFalse(matrix["model_zero_extension_errors"])

    def test_nacar_sequence_direction_and_proline_donor(self):
        _, runtime = sources.load_runtime(sources.RESOURCES_DIRECTORY / "helix_pairs_prop.txt")
        # Figure 3 columns are the earlier acceptor and rows the later donor.
        self.assertEqual(runtime["A", "R"], Decimal("1.724"))
        self.assertEqual(runtime["R", "A"], Decimal("1.648"))
        self.assertEqual(runtime["C", "C"], Decimal("1.800"))
        self.assertEqual(runtime["M", "M"], Decimal("2.878"))
        self.assertTrue(all(runtime[residue, "P"] == 0 for residue in sources.STANDARD_RESIDUES))
        self.assertNotEqual(runtime["P", "A"], 0)
        result = sources.verify()["matrices"]["M1"]
        self.assertEqual(result["source_separation"], 4)
        self.assertEqual(result["runtime_separation"], 1)

    def test_munoz_distinct_row_column_orders_and_m4_values(self):
        rows, columns, published = sources.load_source_csv(
            sources.SOURCE_DIRECTORY / "munoz_serrano1994_table2_i4.csv")
        self.assertLess(rows.index("F"), rows.index("Y"))
        self.assertLess(columns.index("Y"), columns.index("F"))
        self.assertEqual(published["F", "H"], Decimal("-0.800"))
        self.assertEqual(published["Y", "H"], Decimal("-0.850"))
        _, runtime = sources.load_runtime(sources.RESOURCES_DIRECTORY / "helix_sd_i_4.txt")
        self.assertEqual(runtime["F", "H"], published["F", "H"])
        self.assertEqual(runtime["Y", "H"], published["Y", "H"])

    def test_a_wrong_published_cell_is_reported(self):
        with tempfile.TemporaryDirectory(prefix="helix-source-regression-") as temporary:
            directory = Path(temporary)
            for path in sources.RESOURCES_DIRECTORY.glob("helix*.txt"):
                shutil.copyfile(path, directory / path.name)
            path = directory / "helix_sd_i_4.txt"
            lines = path.read_text().splitlines()
            header = "".join(lines[0].split())
            row_index = header.index("F") + 1
            tokens = lines[row_index].split()
            tokens[header.index("H")] = "-0.850"
            lines[row_index] = " ".join(tokens)
            path.write_text("\n".join(lines) + "\n")
            result = sources.verify(directory)
            self.assertFalse(result["verified"])
            self.assertEqual(result["matrices"]["M4"]["published_cells_matching"], 323)
            self.assertEqual(result["matrices"]["M4"]["published_cell_differences"][0]["residue_i"], "F")

    def test_a_nonzero_model_extension_is_not_counted_as_published_data(self):
        with tempfile.TemporaryDirectory(prefix="helix-source-regression-") as temporary:
            directory = Path(temporary)
            for path in sources.RESOURCES_DIRECTORY.glob("helix*.txt"):
                shutil.copyfile(path, directory / path.name)
            path = directory / "helix_sd_i_3.txt"
            lines = path.read_text().splitlines()
            header = "".join(lines[0].split())
            row_index = header.index("C") + 1
            tokens = lines[row_index].split()
            tokens[header.index("A")] = "0.1"
            lines[row_index] = " ".join(tokens)
            path.write_text("\n".join(lines) + "\n")
            result = sources.verify(directory)["matrices"]["M3"]
            self.assertFalse(result["verified"])
            self.assertEqual(result["published_cells_matching"], 324)
            self.assertEqual(len(result["model_zero_extension_errors"]), 1)


if __name__ == "__main__":
    unittest.main()
```
