## Additional file 1. Evaluated software source and dependencies for "Helix Design Studio: designing antimicrobial peptide candidates for membrane environments": test_partial_exports.js

```
// Run with: node --test tests/test_partial_exports.js
// Execute the application helpers and stream handlers, without browser dependencies.
const test = require('node:test');
const assert = require('node:assert/strict');
const fs = require('node:fs');
const path = require('node:path');
const vm = require('node:vm');

const source = fs.readFileSync(path.join(__dirname, '../app/static/app.js'), 'utf8');
const helperNames = [
  'partialExportMetadata', 'partialSpecificityPayload', 'partialFamilyPayload',
  'compareSpecificityCandidates', 'designZ', 'escapeHtml', 'statCard', 'humanizePreset',
  'isInterfacialPreset', 'renderFamilyState', 'renderSpecificity', 'renderSpecificityState',
  'renderSpecificityCandidateDetail', 'bindSpecificityInteractions', 'buildSpecificityCsv',
  'formatObjectiveDelta', 'formatObjectiveDeltaCsv', 'energyUnitNote', 'errorHtml',
  'setFormBusy', 'parseResidues', 'buildEnvironmentFromFormData',
  'buildEnvironmentFromPresetAndPenetration', 'penetrationPercentToHalfWidth',
  'calculationSettingsFromFormData',
];
const helpers = helperNames.map((name) => {
  // Function bodies in app.js end with an unindented closing brace.
  const match = source.match(new RegExp(`^function ${name}\\([^]*?^}`, 'm'));
  assert.ok(match, `Missing application function: ${name}`);
  return match[0];
}).join('\n');
const plain = (value) => JSON.parse(JSON.stringify(value));

function provenance(mode) {
  return {
    software: 'Helix Design Studio', version: '1.0.0rc5',
    parameters_sha256: 'parameters-fixture', seed_protocol: 'seed-sequence-fixture',
    python: '3.12.8', numpy: '2.2.4',
    request: {
      length: 4, residues: ['A', 'E', 'K'], steps: 10, restarts: 2,
      seed: 123, n_decoys: 2000, weights: { M1: 1, M2: 1, M3: 1, M4: 1 },
      ...(mode === 'family'
        ? { family_size: 3, environment: { preset: 'interfacial_neg' }, oversample_factor: 4 }
        : { num_sequences: 3, target_environment: { preset: 'interfacial_neg' },
          off_target_environments: [{ preset: 'homogeneous_polar' }], lambda_balance: 0 }),
    },
  };
}

function scored(sequence, energy) {
  return {
    sequence, energy, z_score: energy / 10,
    environment: { preset: 'interfacial_neg' },
    design_geometry: { energy, z_score: energy / 10, orientation: 'own hydrophobic moment' },
  };
}

function specific(sequence, objective, designZ) {
  return {
    sequence,
    target_environment: scored(sequence, designZ * 10),
    off_target_environments: [],
    specificity_design: { objective, z_score_margin: 1, rank: 99 },
  };
}

function context(overrides = {}) {
  const ctx = vm.createContext({ HELP_TEXT: { stats: {} }, ...overrides });
  vm.runInContext(helpers, ctx);
  return ctx;
}

test('specificity partial retains canonical metadata and ranks without changing accepted candidates', () => {
  const ctx = context();
  const meta = provenance('specificity');
  const accepted = [specific('AAAA', -2, -1), specific('EEEE', -2, -3)];
  accepted.forEach((member) => { Object.freeze(member.specificity_design); Object.freeze(member); });
  const data = ctx.partialSpecificityPayload({
    acceptedCandidates: Object.freeze(accepted), targetCount: 3,
    targetPreset: 'interfacial_neg', offTargets: ['homogeneous_polar'],
    provenance: meta, request: { length: 4 },
  }, 'stopped by the user');
  assert.equal(data.provenance, meta);
  assert.equal(data.request, meta.request);
  assert.equal(data.request.seed, 123);
  assert.deepEqual(plain(data.candidates.map((member) => member.sequence)), ['EEEE', 'AAAA']);
  assert.deepEqual(plain(data.candidates.map((member) => member.specificity_design.rank)), [1, 2]);
  assert.deepEqual(accepted.map((member) => member.specificity_design.rank), [99, 99]);
  assert.equal(data.specificity_collection.requested, 3);
  assert.equal(data.specificity_collection.returned, 2);
  assert.equal(data.specificity_collection.partial, true);
  assert.equal(data.specificity_collection.partial_reason, 'stopped by the user');
});

test('partial metadata falls back to an accepted member, then to the submitted request', () => {
  const ctx = context();
  const meta = provenance('specificity');
  const member = { ...specific('AAAA', -2, -1), provenance: meta };
  const state = { acceptedCandidates: [member], request: { length: 4 } };
  const data = ctx.partialSpecificityPayload(state, 'interrupted by an error');
  assert.equal(data.provenance, meta);
  assert.equal(data.request, meta.request);
  const withoutMeta = ctx.partialSpecificityPayload({ ...state, acceptedCandidates: [] }, 'stopped by the user');
  assert.equal(withoutMeta.request, state.request);
  assert.equal(withoutMeta.candidates.length, 0);
});

test('family partial sorts by the displayed energy and preserves complete scored members', () => {
  const ctx = context();
  const meta = provenance('family');
  const members = [scored('AAAA', -1), { ...scored('EEEE', -3), provenance: meta }];
  const data = ctx.partialFamilyPayload({ members: Object.freeze(members), target: 3 }, 'interrupted by an error');
  assert.equal(data.provenance, meta);
  assert.equal(data.request, meta.request);
  assert.equal(data.members[0], members[1]);
  assert.equal(data.members[1], members[0]);
  assert.equal(data.family_size_requested, 3);
  assert.equal(data.family_size_returned, 2);
  assert.equal(data.partial, true);
  assert.equal(data.partial_reason, 'interrupted by an error');
  assert.equal(members[0].sequence, 'AAAA');
});

// Minimal DOM surface for the actual submit, Stop and download handlers. Replacing
// innerHTML replaces its nodes, so a lost listener after a refresh fails the test.
class Element {
  constructor() { this.listeners = {}; this.nodes = []; this.attributes = {}; this._html = ''; }
  set innerHTML(value) {
    this._html = value;
    this.nodes = [...value.matchAll(/\b(data-[a-z-]+)(?:="([^"]*)")?/g)].map((match) => {
      const node = new Element();
      node.attribute = match[1];
      node.dataset = { [match[1].slice(5).replace(/-([a-z])/g, (_, letter) => letter.toUpperCase())]: match[2] || '' };
      node.classList = { toggle() {} };
      return node;
    });
  }
  get innerHTML() { return this._html; }
  querySelector(selector) { return this.querySelectorAll(selector)[0] || null; }
  querySelectorAll(selector) { return this.nodes.filter((node) => `[${node.attribute}]` === selector); }
  addEventListener(type, fn, options = {}) { (this.listeners[type] ||= []).push({ fn, once: options.once }); }
  async dispatch(type) {
    for (const item of [...(this.listeners[type] || [])]) {
      if (item.once) this.listeners[type] = this.listeners[type].filter((other) => other !== item);
      await item.fn({ preventDefault() {} });
    }
    if (type === 'click' && this.onclick) await this.onclick();
  }
  click() { return this.dispatch('click'); }
  focus() {}
  setAttribute(name, value) { this.attributes[name] = value; }
  insertAdjacentHTML(_position, value) { this._html = value + this._html; }
}

function streamHarness(mode, runStream, extraValues = {}) {
  const result = new Element();
  const form = new Element();
  const submit = new Element();
  form.querySelector = (selector) => selector === 'button[type="submit"]' ? submit : null;
  const values = {
    length: '4', residues: 'A,E,K', steps: '10', restarts: '2', family_size: '3',
    preset: 'interfacial_neg', penetration_depth: '25', num_sequences: '3',
    target_preset: 'interfacial_neg', off_target_preset: ['homogeneous_polar'], lambda_balance: '0',
    ...extraValues,
  };
  const document = { activeElement: null, querySelector: (selector) => {
    if (selector === `#${mode}-form`) return form;
    if (selector === `#${mode}-result`) return result;
    return null;
  } };
  const downloads = [];
  let streamCalls = 0;
  const ctx = context({
    document, AbortController,
    $: (selector, root = document) => root.querySelector(selector),
    $$: (selector, root = document) => root.querySelectorAll(selector),
    FormData: class { get(key) { return values[key]; } getAll(key) { return values[key] || []; } },
    downloadJson: (filename, data) => downloads.push({ filename, data }),
    refreshUsageCounters: async () => {},
    renderCompareEnvironmentCard: () => '', // unrelated score plots
    postNDJSON: async (_url, request, emit, options) => {
      streamCalls += 1;
      await runStream({ request, emit, signal: options.signal, result });
    },
  });
  const handler = mode === 'family'
    ? source.slice(source.indexOf("const familyForm = $('#family-form');"))
    : source.slice(source.indexOf("const specificityForm = $('#specificity-form');"), source.indexOf("bindForm('pdb-form'"));
  vm.runInContext(handler, ctx);
  return { form, result, submit, downloads, streamCalls: () => streamCalls };
}

for (const mode of ['family', 'specificity']) {
  test(`${mode}: advanced settings reach the calculation and keep homogeneous competitors`, async () => {
    let submitted;
    const harness = streamHarness(mode, async ({ request }) => { submitted = request; }, {
      n_decoys: '2000', seed: '0', exact_halfwidth_penetration_depth: '90',
      off_target_preset: ['interfacial_neu', 'homogeneous_polar'],
    });
    await harness.form.dispatch('submit');
    assert.equal(submitted.n_decoys, 2000);
    assert.equal(submitted.seed, 0);
    const target = mode === 'family' ? submitted.environment : submitted.target_environment;
    assert.equal(target.wheel_halfwidth_deg, 90);
    if (mode === 'specificity') {
      assert.deepEqual(plain(submitted.off_target_environments), [
        { preset: 'interfacial_neu', wheel_halfwidth_deg: 90 },
        { preset: 'homogeneous_polar' },
      ]);
    }
  });

  test(`${mode}: absent advanced settings preserve existing sampling defaults`, async () => {
    let submitted;
    const harness = streamHarness(mode, async ({ request }) => { submitted = request; });
    await harness.form.dispatch('submit');
    assert.equal(submitted.n_decoys, mode === 'specificity' ? 10000 : 2000);
    assert.equal(submitted.seed, 42);
    const target = mode === 'family' ? submitted.environment : submitted.target_environment;
    assert.equal(target.wheel_halfwidth_deg, 67.5);
  });

  for (const ending of ['stop', 'error', 'eof']) {
    test(`${mode}: ${ending} keeps accepted results downloadable with canonical provenance`, async () => {
      const meta = provenance(mode);
      const member = mode === 'family' ? scored('AAAA', -2) : specific('AAAA', -2, -3);
      const harness = streamHarness(mode, async ({ emit, signal, result }) => {
        emit({ type: 'meta', payload: { provenance: meta, environment: { preset: 'interfacial_neg' }, min_distance: 2 } });
        emit({ type: 'candidate', payload: { accepted: 1, attempt: 1, target: 3, member } });
        if (ending === 'stop') {
          const stop = result.querySelector(`[data-stop-${mode}]`);
          assert.ok(stop, 'Stop remains bound after a candidate refresh');
          await stop.click();
          assert.equal(signal.aborted, true);
          throw Object.assign(new Error('Aborted'), { name: 'AbortError' });
        }
        if (ending === 'error') throw new Error('Connection interrupted');
      });
      await harness.form.dispatch('submit');
      assert.equal(harness.submit.disabled, false);
      const button = harness.result.querySelector(`[data-download-${mode === 'family' ? 'family' : 'specificity-json'}]`);
      assert.ok(button, 'Partial JSON button is visible');
      await button.click();
      assert.equal(harness.downloads.length, 1);
      const data = harness.downloads[0].data;
      assert.equal(data.provenance, meta);
      assert.equal(data.request, meta.request);
      assert.equal(data.request.seed, 123);
      const collection = mode === 'family' ? data : data.specificity_collection;
      assert.equal(collection.partial, true);
      assert.equal(collection.partial_reason, ending === 'stop' ? 'stopped by the user' : 'interrupted by an error');
      assert.equal(mode === 'family' ? data.family_size_requested : collection.requested, 3);
      assert.equal(mode === 'family' ? data.family_size_returned : collection.returned, 1);
      assert.equal(harness.result.querySelector(`[data-stop-${mode}]`), null);
      // A later submission must not be blocked by the previous controller.
      await harness.form.dispatch('submit');
      assert.equal(harness.streamCalls(), 2);
    });
  }

  test(`${mode}: candidate provenance is retained if meta is absent`, async () => {
    const meta = provenance(mode);
    const member = { ...(mode === 'family' ? scored('AAAA', -2) : specific('AAAA', -2, -3)), provenance: meta };
    const harness = streamHarness(mode, async ({ emit }) => {
      emit({ type: 'candidate', payload: { accepted: 1, attempt: 1, target: 3, member } });
      throw new Error('Connection interrupted');
    });
    await harness.form.dispatch('submit');
    await harness.result.querySelector(`[data-download-${mode === 'family' ? 'family' : 'specificity-json'}]`).click();
    assert.equal(harness.downloads[0].data.provenance, meta);
    assert.equal(harness.downloads[0].data.request, meta.request);
  });

  test(`${mode}: no accepted candidates means no download button`, async () => {
    const harness = streamHarness(mode, async () => { throw new Error('Connection interrupted'); });
    await harness.form.dispatch('submit');
    assert.doesNotMatch(harness.result.innerHTML, /data-download-/);
    assert.equal(harness.downloads.length, 0);
  });

  for (const lateError of [false, true]) {
    test(`${mode}: complete JSON is unchanged${lateError ? ' after a later connection error' : ''}`, async () => {
      const member = mode === 'family' ? scored('AAAA', -2) : specific('AAAA', -2, -3);
      const finalData = {
        ...(mode === 'family'
          ? { members: [member], family_size_requested: 3, family_size_returned: 1, status_message: 'Completed' }
          : { ...member, candidates: [member], specificity_collection: { requested: 3, returned: 1 } }),
        provenance: provenance(mode), server_field: { preserve: true },
      };
      const original = JSON.stringify(finalData);
      const harness = streamHarness(mode, async ({ emit }) => {
        emit({ type: 'final', payload: finalData });
        if (lateError) throw new Error('Connection closed after final');
      });
      await harness.form.dispatch('submit');
      await harness.result.querySelector(`[data-download-${mode === 'family' ? 'family' : 'specificity-json'}]`).click();
      assert.equal(harness.downloads[0].data, finalData);
      assert.equal(JSON.stringify(harness.downloads[0].data), original);
      assert.doesNotMatch(harness.result.innerHTML, /Partial result/);
    });
  }
}

test('family renderer never offers an empty JSON collection', () => {
  const html = context().renderFamilyState({ members: [], target: 3, accepted: 0, finalData: { members: [] } });
  assert.doesNotMatch(html, /data-download-family/);
});


test('specificity CSV exports a single scoring convention with no duplicate design columns', () => {
  const candidate = specific('AAAA', -2, -3);
  const off = { ...scored('AAAA', -20), environment: { preset: 'homogeneous_polar' } };
  candidate.off_target_environments = [off];
  const csv = context().buildSpecificityCsv({
    ...candidate, candidates: [candidate],
    specificity_collection: { off_target_presets: ['homogeneous_polar'] },
  });
  const [header, row] = csv.split('\n').map((line) => line.split(','));
  assert.equal(header.length, row.length);
  assert.ok(header.includes('target_z_score'));
  assert.ok(header.includes('homogeneous_polar_z_score'));
  assert.doesNotMatch(header.join(','), /design_geometry|standard/);
  assert.equal(row[header.indexOf('target_z_score')], '-3.000000');
  assert.equal(row[header.indexOf('homogeneous_polar_z_score')], '-2.000000');
});

test('family renderer shows only the single optimized score', () => {
  const html = context().renderFamilyState({ members: [scored('AAAA', -2)], target: 1, accepted: 1 });
  assert.match(html, /Score: energy -2.000 \| Z -0.200/);
  assert.doesNotMatch(html, /design geometry|fixed wheel/);
});
```
