## Additional file 1. Evaluated software source and dependencies for "Helix Design Studio: designing antimicrobial peptide candidates for membrane environments": test_smoke.py

```
"""Offline runtime checks, not validation of the scientific model.

Run from the repository root: python -m unittest discover -s tests -v -b
Parameter-table checks verify label-based indexing, not parameter provenance.
"""

import io
import json
import math
import os
from pathlib import Path
import re
import tempfile
import unittest


class WebSmokeTests(unittest.TestCase):
    @classmethod
    def setUpClass(cls):
        cls.temporary = tempfile.TemporaryDirectory(prefix="helix-tests-")
        cls.addClassCleanup(cls.temporary.cleanup)
        cls.previous_env = {
            key: os.environ.get(key)
            for key in ("HELIX_CACHE_DIR", "HELIX_VAR_DIR", "HELIX_USAGE_LOGGING")
        }
        os.environ["HELIX_CACHE_DIR"] = str(Path(cls.temporary.name) / "cache")
        os.environ["HELIX_VAR_DIR"] = str(Path(cls.temporary.name) / "var")
        os.environ["HELIX_USAGE_LOGGING"] = "0"
        cls.addClassCleanup(cls.restore_environment)
        # Configuration is read at import time, after setting temporary paths.
        from fastapi.testclient import TestClient
        from app.main import app
        from app import engine

        cls.engine = engine
        cls.client = TestClient(app)
        cls.addClassCleanup(cls.client.close)
        cls.common = {"residues": engine.DEFAULT_ALPHABET_16, "n_decoys": 100, "seed": 42}
        cls.design = {**cls.common, "length": 4, "steps": 1000, "restarts": 1}
        cls.env_a = {"preset": "interfacial_neg"}
        cls.env_b = {"preset": "homogeneous_polar"}

    @classmethod
    def restore_environment(cls):
        for key, value in cls.previous_env.items():
            if value is None:
                os.environ.pop(key, None)
            else:
                os.environ[key] = value

    def assert_finite(self, value):
        if isinstance(value, dict):
            for child in value.values():
                self.assert_finite(child)
        elif isinstance(value, list):
            for child in value:
                self.assert_finite(child)
        elif isinstance(value, (int, float)):
            self.assertTrue(math.isfinite(value), value)

    def post(self, route, payload):
        response = self.client.post(route, json=payload)
        self.assertEqual(response.status_code, 200, response.text)
        result = response.json()
        self.assert_finite(result)
        return result

    def stream(self, route, payload):
        with self.client.stream("POST", route, json=payload) as response:
            self.assertEqual(response.status_code, 200)
            self.assertIn("application/x-ndjson", response.headers["content-type"])
            events = [json.loads(line) for line in response.iter_lines() if line]
        self.assertTrue(events)
        self.assertFalse([event for event in events if event["type"] == "error"], events)
        self.assertEqual(events[-1]["type"], "final")
        self.assertEqual(sum(event["type"] == "final" for event in events), 1)
        self.assert_finite(events)
        return events[-1]["payload"]

    def assert_score(self, result):
        self.assertEqual(len(result["sequence"]), 4)
        self.assertEqual(result["length"], 4)
        self.assertTrue(set(result["sequence"]) <= set(self.common["residues"]))
        self.assertAlmostEqual(result["energy"], result["breakdown"]["total"])
        self.assertEqual(result["wheel"][0]["angle_deg"] % 360, result["environment"]["wheel_phase_deg"] % 360)
        self.assertGreater(result["random_reference"]["std"], 0)
        self.assertLessEqual(0, result["percentile_lower_is_better"])
        self.assertLessEqual(result["percentile_lower_is_better"], 100)
        self.assertEqual(len(result["wheel"]), 4)

    def test_page_assets_and_health(self):
        response = self.client.get("/")
        self.assertEqual(response.status_code, 200)
        assets = set(re.findall(r'(?:src|href)="(/static/[^"?#]+)', response.text))
        self.assertTrue(assets)
        for asset in assets:
            with self.subTest(asset=asset):
                response = self.client.get(asset)
                self.assertEqual(response.status_code, 200)
                self.assertTrue(response.content)
        health = self.client.get("/api/health")
        self.assertEqual(health.status_code, 200)
        self.assertEqual(health.json()["status"], "ok")
        self.assertEqual(self.client.get("/openapi.json").status_code, 200)

    def test_score_and_reproducibility(self):
        payload = {**self.common, "sequence": "ALKD", "n_random": 100}
        first = self.post("/api/score", payload)
        self.assert_score(first)
        self.assertEqual(first, self.post("/api/score", payload))

    def test_single_design(self):
        self.assert_score(self.post("/api/design", self.design))

    def test_comparison_and_cross_design(self):
        environments = {"environment_a": self.env_a, "environment_b": self.env_b}
        requests = {
            "/api/compare": {**self.common, **environments, "sequence": "ALKD"},
            "/api/cross-design": {**self.design, **environments},
        }
        for route, payload in requests.items():
            with self.subTest(route=route):
                result = self.post(route, payload)
                self.assert_score(result["environment_a"])
                self.assert_score(result["environment_b"])
                self.assertEqual(result["environment_a"]["sequence"], result["sequence"])
                self.assertEqual(result["environment_b"]["sequence"], result["sequence"])

    def test_family_and_stream(self):
        payload = {**self.design, "family_size": 2, "oversample_factor": 1}
        result = self.post("/api/design-family", payload)
        self.assertEqual(result["family_size_returned"], 2)
        self.assertEqual(len({member["sequence"] for member in result["members"]}), 2)
        for member in result["members"]:
            self.assert_score(member)
        self.assertEqual(result, self.stream("/api/design-family-stream", payload))

    def test_specificity_and_stream(self):
        payload = {
            **self.design,
            "target_environment": self.env_a,
            "off_target_environments": [self.env_b],
            "num_sequences": 1,
        }
        result = self.post("/api/specificity-design", payload)
        self.assertEqual(result["specificity_collection"]["returned"], 1)
        self.assertEqual(len(result["candidates"]), 1)
        self.assert_score(result["target_environment"])
        self.assertEqual(len(result["off_target_environments"]), 1)
        self.assert_score(result["off_target_environments"][0])
        self.assertEqual(result, self.stream("/api/specificity-design-stream", payload))

    def test_penetration_scan(self):
        result = self.post("/api/optimal-penetration", {
            **self.common, "sequence": "ALKD", "environment": self.env_a, "n_random": 100,
        })
        self.assertEqual([point["penetration_percent"] for point in result["profile"]], list(range(0, 101, 5)))
        self.assert_score(result["best_score"])
        best = min(result["profile"], key=lambda point: point["z_score"])
        self.assertEqual(result["best_penetration_percent"], best["penetration_percent"])

    def test_pdb_export(self):
        from Bio.PDB import PDBParser

        result = self.post("/api/pdb", {"sequence": "ALKD"})
        structure = PDBParser(QUIET=True).get_structure("test", io.StringIO(result["pdb"]))
        self.assertEqual([residue.resname for residue in structure.get_residues()], ["ALA", "LEU", "LYS", "ASP"])
        for atom in structure.get_atoms():
            self.assertTrue(all(math.isfinite(float(x)) for x in atom.coord))

    def test_request_limits(self):
        specificity = {"target_environment": self.env_a, "off_target_environments": [self.env_b]}
        invalid = [
            ("/api/score", {"sequence": ""}),
            ("/api/score", {"sequence": "ALXD"}),
            ("/api/score", {"sequence": "ALKD", "residues": ["A", "L", "K", "D", "B"]}),
            ("/api/score", {"sequence": "ALKD", "environment": {"preset": "interfacial_unknown"}}),
            ("/api/score", {"sequence": "ALKD", "environment": {"membrane_charge": "negative"}}),
            ("/api/score", {"sequence": "ALKD", "environment": {"wheel_halfwidth_deg": 1000}}),
            ("/api/score", {"sequence": "ALKD", "weights": {"lambda_env": -1}}),
            ("/api/score", {"sequence": "ALKD", "seed": -1}),
            ("/api/design", {"length": 41}),
            ("/api/design", {"residues": ["A"]}),
            ("/api/optimal-penetration", {"sequence": "ALKD", "environment": self.env_b}),
            # Cross-field checks must run even when the optional seed is omitted.
            ("/api/optimal-penetration", {"sequence": "ALKD", "environment": self.env_a, "percent_step": 20}),
            ("/api/specificity-design", {**specificity, "off_target_environments": [self.env_a, self.env_b]}),
            ("/api/specificity-design", {**specificity, "off_target_environments": [
                {"preset": "interfacial_neg", "wheel_halfwidth_deg": 90.0}]}),
            ("/api/specificity-design", {**specificity, "off_target_environments": [
                {"preset": "homogeneous_polar", "wheel_halfwidth_deg": float(angle)} for angle in range(5)
            ]}),
            ("/api/specificity-design", {**specificity, "num_sequences": 5, "restarts": 4}),
            # Same Hamiltonian under another preset name.
            ("/api/specificity-design", {**specificity, "off_target_environments": [
                {"preset": "interfacial_pos", "membrane_charge": "neg"}]}),
            ("/api/score", {"sequence": "ALKD", "n_decoy": 100}),
        ]
        for route, payload in invalid:
            with self.subTest(route=route, payload=payload):
                self.assertEqual(self.client.post(route, json=payload).status_code, 422)

    def test_non_finite_numbers_are_rejected(self):
        bodies = [
            ("/api/score", '{"sequence": "ALKD", "weights": {"lambda_env": NaN}}'),
            ("/api/score", '{"sequence": "ALKD", "environment": {"wheel_halfwidth_deg": Infinity}}'),
            ("/api/cross-design", '{"environment_a": {}, "environment_b": {}, "lambda_gap": NaN}'),
        ]
        for route, body in bodies:
            with self.subTest(route=route, body=body):
                response = self.client.post(route, content=body, headers={"Content-Type": "application/json"})
                self.assertEqual(response.status_code, 422, response.text)
                self.assertTrue(response.json()["detail"])

    def test_request_size_and_security_headers(self):
        response = self.client.post("/api/score", content=b"{" + b" " * 70000 + b"}",
                                    headers={"Content-Type": "application/json"})
        self.assertEqual(response.status_code, 413)
        chunked = (part for part in [b'{"sequence": "ALKD", "pad": "'] + [b"x" * 4096] * 20 + [b'"}'])
        response = self.client.post("/api/score", content=chunked, headers={"Content-Type": "application/json"})
        self.assertEqual(response.status_code, 413)
        page = self.client.get("/")
        self.assertEqual(page.headers["x-content-type-options"], "nosniff")
        self.assertIn("frame-ancestors 'none'", page.headers["content-security-policy"])

    def test_busy_server_and_time_limit(self):
        from app import main

        slots = [main._job_slots.acquire(blocking=False) for _ in range(main.MAX_CONCURRENT_JOBS)]
        try:
            self.assertTrue(all(slots))
            self.assertEqual(self.client.post("/api/design", json=self.design).status_code, 503)
            with self.client.stream("POST", "/api/design-family-stream", json={**self.design, "family_size": 2}) as response:
                events = [json.loads(line) for line in response.iter_lines() if line]
            self.assertEqual([event["type"] for event in events], ["error"])
        finally:
            for _ in slots:
                main._job_slots.release()

        previous = main.MAX_JOB_SECONDS
        main.MAX_JOB_SECONDS = 0.0
        try:
            response = self.client.post("/api/design", json=self.design)
            self.assertEqual(response.status_code, 400)
            self.assertIn("time limit", response.json()["detail"])
            with self.client.stream("POST", "/api/design-family-stream", json={**self.design, "family_size": 2}) as response:
                events = [json.loads(line) for line in response.iter_lines() if line]
            self.assertEqual(events[-1]["type"], "error")
            self.assertIn("time limit", events[-1]["detail"])
        finally:
            main.MAX_JOB_SECONDS = previous
        self.assertEqual(self.client.post("/api/design", json=self.design).status_code, 200)

    def test_results_include_provenance(self):
        from app import main

        request = {"sequence": "ALKD", "residues": ["A", "L", "K", "D"], "n_decoys": 100, "n_random": 100, "seed": 7}
        result = self.post("/api/score", request)
        provenance = result["provenance"]
        self.assertEqual(provenance["version"], main.APP_VERSION)
        self.assertEqual(provenance["parameters_sha256"], main.PARAMETERS_SHA256)
        self.assertEqual({key: provenance["request"][key] for key in request}, request)
        self.assertEqual(provenance["request"]["environment"]["preset"], "interfacial_neg")
        self.assertIn("seed_protocol", provenance)
        self.assertEqual(result["calibration"]["n_decoys"], 100)
        self.assertEqual(result["calibration"]["seed_entropy"], 7)
        self.assertEqual(result["random_reference"]["seed_entropy"], [7, 1])
        family = self.stream("/api/design-family-stream", {**self.design, "family_size": 2, "oversample_factor": 1})
        self.assertEqual(family["provenance"]["request"]["family_size"], 2)

    def test_stream_candidates_include_provenance(self):
        from app import main

        requests = {
            "/api/design-family-stream": {
                **self.design, "family_size": 2, "oversample_factor": 1,
            },
            "/api/specificity-design-stream": {
                **self.design, "target_environment": self.env_a,
                "off_target_environments": [self.env_b], "num_sequences": 1,
            },
        }
        for route, request in requests.items():
            with self.subTest(route=route):
                with self.client.stream("POST", route, json=request) as response:
                    self.assertEqual(response.status_code, 200)
                    events = [json.loads(line) for line in response.iter_lines() if line]
                self.assertEqual(events[0]["type"], "meta")
                self.assertEqual(events[-1]["type"], "final")
                provenance = events[0]["payload"]["provenance"]
                self.assertEqual(provenance["version"], main.APP_VERSION)
                self.assertEqual(provenance["parameters_sha256"], main.PARAMETERS_SHA256)
                request_model = (main.FamilyDesignRequest if route == "/api/design-family-stream"
                                 else main.SpecificityDesignRequest)
                self.assertEqual(provenance["request"], request_model(**request).model_dump(mode="json"))
                candidates = [event["payload"]["member"] for event in events if event["type"] == "candidate"]
                self.assertTrue(candidates)
                for member in candidates:
                    self.assertEqual(member["provenance"], provenance)
                self.assertEqual(events[-1]["payload"]["provenance"], provenance)

    def test_calibration_honours_cancellation(self):
        import threading

        cancelled = threading.Event()
        cancelled.set()
        env = self.engine.resolve_environment(self.env_a)
        with tempfile.TemporaryDirectory(prefix="helix-cache-") as cache_dir:
            with self.engine.calculation_limits(self.engine.CalculationLimits(cancelled=cancelled)):
                with self.assertRaises(self.engine.CalculationCancelled):
                    self.engine.build_builder(12, list(self.engine.DEFAULT_ALPHABET_16), env,
                                              n_decoys=5000, stats_cache_dir=cache_dir)

    def test_stream_disconnect_releases_calculation_slot(self):
        """A real server: closing a stream must stop the calculation and free its slot."""
        import socket
        import subprocess
        import sys
        import time
        import httpx

        with socket.socket() as sock:
            sock.bind(("127.0.0.1", 0))
            port = sock.getsockname()[1]
        env = {**os.environ, "HELIX_MAX_CONCURRENT_JOBS": "1",
               "HELIX_CACHE_DIR": str(Path(self.temporary.name) / "cache-disconnect")}
        server = subprocess.Popen([sys.executable, "-m", "uvicorn", "app.main:app", "--port", str(port)],
                                  cwd=Path(__file__).resolve().parents[1], env=env,
                                  stdout=subprocess.DEVNULL, stderr=subprocess.DEVNULL)
        self.addCleanup(server.wait, 10)
        self.addCleanup(server.terminate)
        base = f"http://127.0.0.1:{port}"
        for _ in range(100):
            try:
                httpx.get(f"{base}/api/health", timeout=1)
                break
            except httpx.TransportError:
                time.sleep(0.1)
        heavy = {**self.design, "length": 30, "steps": 20000, "restarts": 10, "family_size": 4}
        with httpx.stream("POST", f"{base}/api/design-family-stream", json=heavy, timeout=30) as response:
            self.assertEqual(json.loads(next(response.iter_lines()))["type"], "meta")
        deadline = time.monotonic() + 10
        status = None
        while time.monotonic() < deadline:
            status = httpx.post(f"{base}/api/design", json=self.design, timeout=30).status_code
            if status == 200:
                break
            time.sleep(0.2)
        self.assertEqual(status, 200, "the calculation slot was not released after the client disconnected")

    def test_usage_logging_enabled(self):
        """Logging is configured at import time, so it is exercised in a fresh interpreter."""
        import subprocess
        import sys

        script = """
import json, pathlib
from fastapi.testclient import TestClient
from app import main
client = TestClient(main.app)
headers = {"X-Anonymous-Id": "visitor-1"}
assert client.post("/api/visit", headers=headers).json()["total_visits"] == 1
assert client.post("/api/visit", headers={"X-Anonymous-Id": "visitor-2"}).json()["unique_visitors"] == 2
assert client.post("/api/visit", headers={"Referer": "https://[invalid"}).status_code == 200
payload = {"sequence": "ALKD", "residues": ["A", "L", "K", "D"], "n_decoys": 100, "n_random": 100}
assert client.post("/api/score", json=payload, headers={"Referer": "https://[invalid"}).status_code == 200
assert client.post("/api/score", json={**payload, "sequence": "ALKE"}).status_code == 422
main.MAX_JOB_SECONDS = 0.0
assert client.post("/api/score", json=payload).status_code == 400
main.MAX_JOB_SECONDS = 1800.0
assert client.get("/api/usage-metrics").json() == {"total_visits": 3, "unique_visitors": 2, "total_runs": 1}
assert client.get("/api/health").status_code == 200, "client errors must not degrade health"
main.score_sequence = lambda **kwargs: 1 / 0
assert client.post("/api/score", json=payload).status_code == 500
health = client.get("/api/health")
assert health.status_code == 503 and health.json()["recent"]["failing_endpoints"] == {"score": 1}
log = pathlib.Path(main.USAGE_LOG_PATH).read_text()
assert "ALKD" not in log and "testclient" not in log
assert all(json.loads(line)["timestamp_utc"] for line in log.splitlines())
"""
        env = {**os.environ, "HELIX_USAGE_LOGGING": "1",
               "HELIX_VAR_DIR": str(Path(self.temporary.name) / "var-enabled")}
        completed = subprocess.run([sys.executable, "-c", script], cwd=Path(__file__).resolve().parents[1],
                                   env=env, capture_output=True, text=True, timeout=300)
        self.assertEqual(completed.returncode, 0, completed.stderr)

    def test_usage_logging_disabled(self):
        from app import main

        self.assertFalse(main.USAGE_LOGGING_ENABLED)
        self.assertEqual(self.client.post("/api/visit", headers={"X-Anonymous-Id": "visitor"}).status_code, 200)
        self.assertEqual(self.client.get("/api/usage-metrics").json()["total_visits"], 0)
        self.assertFalse(main.USAGE_LOG_PATH.exists())
        self.assertFalse((Path(self.temporary.name) / "var" / "usage_events.jsonl").exists())

    def test_raw_terms_do_not_depend_on_alphabet_order(self):
        """The same chemical sequence has the same raw terms for any alphabet order."""
        alphabet = list(self.engine.DEFAULT_ALPHABET_16)
        orders = [alphabet, alphabet[::-1], alphabet[5:] + alphabet[:5], sorted(alphabet)]
        env = self.engine.resolve_environment(self.env_a)
        terms = ("_raw_helix_local", "_raw_env_pol", "_raw_env_chg", "_raw_pw_int",
                 "_raw_electrostatic", "_raw_helix_neigh")
        for sequence in ("PPPP", "AEKLWKEAGH", "DKTSQEMFYVAL"):
            values = []
            for residues in orders:
                builder = self.engine.build_builder(len(sequence), residues, env, n_decoys=100)
                codes = self.engine.seq_to_codes(sequence, residues)
                values.append([getattr(builder, term)(codes) for term in terms])
            for other in values[1:]:
                with self.subTest(sequence=sequence):
                    for expected, observed in zip(values[0], other):
                        self.assertAlmostEqual(expected, observed, places=12)

    def test_neighbor_tables_are_indexed_by_residue_label(self):
        """Entries are looked up by residue label (row: residue i, column: residue i+k)."""
        residues = ["K", "P", "E", "A", "C", "L"]
        builder = self.engine.build_builder(6, residues, self.engine.resolve_environment(self.env_a), n_decoys=100)
        index = {aa: i for i, aa in enumerate(residues)}
        # Stored values reproduce the published tables; the scoring convention
        # subtracts M1 and adds M3/M4 after averaging each separation.
        self.assertAlmostEqual(builder.M1[index["A"], index["A"]], 2.734)
        self.assertAlmostEqual(builder.M1[index["K"], index["E"]], 2.264)
        self.assertAlmostEqual(builder.M3[index["E"], index["K"]], -0.150)
        self.assertAlmostEqual(builder.M4[index["E"], index["K"]], -0.420)
        self.assertAlmostEqual(builder.M4[index["K"], index["E"]], 0.100)
        self.assertAlmostEqual(builder.mj_matrix[index["L"], index["K"]], builder.mj_matrix[index["K"], index["L"]])
        for matrix in (builder.M3, builder.M4):
            self.assertFalse(matrix[index["C"]].any())
            self.assertFalse(matrix[:, index["P"]].any())

    def test_invalid_cached_sigma_is_recomputed(self):
        env = self.engine.resolve_environment(self.env_a)
        residues = list(self.engine.DEFAULT_ALPHABET_16)
        with tempfile.TemporaryDirectory(prefix="helix-cache-") as cache_dir:
            fresh = self.engine.build_builder(5, residues, env, n_decoys=100, stats_cache_dir=cache_dir)
            cache_path = fresh._stats_cache_path(100)
            for bad_sigma in (0.0, -1.0):
                with open(cache_path, encoding="utf-8") as handle:
                    cached = json.load(handle)
                cached["stats"]["env_pol"]["sigma"] = bad_sigma
                with open(cache_path, "w", encoding="utf-8") as handle:
                    json.dump(cached, handle)
                reloaded = self.engine.build_builder(5, residues, env, n_decoys=100, stats_cache_dir=cache_dir)
                with self.subTest(sigma=bad_sigma):
                    self.assertEqual(reloaded.stats, fresh.stats)

    def test_corrupt_calibration_cache_is_recomputed(self):
        env = self.engine.resolve_environment(self.env_a)
        residues = list(self.engine.DEFAULT_ALPHABET_16)
        with tempfile.TemporaryDirectory(prefix="helix-cache-") as cache_dir:
            first = self.engine.build_builder(5, residues, env, n_decoys=100, stats_cache_dir=cache_dir)
            cache_path = first._stats_cache_path(100)
            with open(cache_path, "w", encoding="utf-8") as handle:
                handle.write('{"stats": {"helix_local": {"mu"')
            second = self.engine.build_builder(5, residues, env, n_decoys=100, stats_cache_dir=cache_dir)
            self.assertEqual(first.stats, second.stats)
            with open(cache_path, encoding="utf-8") as handle:
                self.assertEqual(json.load(handle)["metadata"]["n_decoys"], 100)
            self.assertEqual([name for name in os.listdir(cache_dir) if name.endswith(".tmp")], [])

    def test_literature_residue_scales(self):
        """Fauchère–Pliska (1983) hydrophobicities and Pace–Scholtz (1998) helix propensities."""
        from core import hamiltonian_builder as model

        fauchere_pliska = {
            "A": 0.31, "R": -1.01, "N": -0.60, "D": -0.77, "C": 1.54, "Q": -0.22, "E": -0.64,
            "G": 0.00, "H": 0.13, "I": 1.80, "L": 1.70, "K": -0.99, "M": 1.23, "F": 1.79,
            "P": 0.72, "S": -0.04, "T": 0.26, "W": 2.25, "Y": 0.96, "V": 1.22,
        }
        pace_scholtz = {
            "A": 0.00, "L": 0.21, "R": 0.21, "M": 0.24, "K": 0.26, "Q": 0.39, "E": 0.40,
            "I": 0.41, "W": 0.49, "S": 0.50, "Y": 0.53, "F": 0.54, "V": 0.61, "H": 0.61,
            "N": 0.65, "T": 0.66, "C": 0.68, "D": 0.69, "G": 1.00, "P": 3.16,
        }
        self.assertEqual(model.FAUCHERE_PLISKA_HYDROPHOBICITY, fauchere_pliska)
        self.assertEqual(model.PACE_SCHOLTZ_HELIX_PROPENSITY, pace_scholtz)
        self.assertIs(self.engine.HYDROPHOBICITY_FP, model.FAUCHERE_PLISKA_HYDROPHOBICITY)
        script = (Path(__file__).resolve().parents[1] / "app" / "static" / "app.js").read_text(encoding="utf-8")
        block = re.search(r"const HYDRO_SCALE = \{(.*?)\};", script, re.S).group(1)
        browser_scale = {aa: float(value) for aa, value in re.findall(r"([A-Z]):\s*(-?[0-9.]+)", block)}
        self.assertEqual(browser_scale, fauchere_pliska)

    def test_helix_neighbor_term_uses_negative_m1_and_source_energy_signs(self):
        """Subtract mean M1 and add mean M3/M4, retaining the published raw tables."""
        env = self.engine.resolve_environment(self.env_b)
        residues = ["A", "E", "G", "L", "P"]
        builder = self.engine.build_builder(10, residues, env, n_decoys=100)
        codes = self.engine.seq_to_codes("AEGLPAEGLP", residues)
        expected = 0.0
        for k, matrix, sign in ((1, builder.M1, -1), (3, builder.M3, 1), (4, builder.M4, 1)):
            pairs = [matrix[codes[i], codes[i + k]] for i in range(len(codes) - k)]
            expected += sign * sum(pairs) / (len(codes) - k)
        self.assertAlmostEqual(builder._raw_helix_neigh(codes), expected, places=12)
        # M1 holds the raw propensity values from the file, not a logarithm of them.
        self.assertAlmostEqual(builder.M1[residues.index("A"), residues.index("A")], 2.734)

    def test_random_sequences_are_not_favored_by_phase_alignment(self):
        """Query, calibration decoys and reference sequences share the same orientation rule."""
        import numpy as np

        residues = list(self.engine.DEFAULT_ALPHABET_16)
        rng = np.random.default_rng(7)
        percentiles = []
        for _ in range(40):
            sequence = "".join(rng.choice(residues, size=12))
            result = self.engine.score_sequence(
                sequence, self.env_a, residues, n_decoys=200, n_random=200, seed=42
            )
            percentiles.append(result["percentile_lower_is_better"])
        self.assertGreater(sum(percentiles) / len(percentiles), 35)
        self.assertLess(sum(percentiles) / len(percentiles), 65)

    def test_search_and_score_share_oriented_calibration_and_reference(self):
        """The search is calibrated with independently aligned decoys, not fixed-phase decoys."""
        import numpy as np
        from unittest.mock import patch

        residues = ["A", "K", "G", "L"]
        env = self.engine.resolve_environment({**self.env_a, "wheel_phase_deg": 37.0})
        weights = self.engine.normalize_weights(None)
        with tempfile.TemporaryDirectory() as cache_dir, patch.dict(os.environ, {"HELIX_CACHE_DIR": cache_dir}):
            design, scoring = self.engine.report_models(
                6, residues, env, weights, n_decoys=100, seed=7, n_random=100,
            )
            self.assertIs(design, scoring)
            self.assertTrue(design.builder.align_phase)
            self.assertEqual(len(list(Path(cache_dir).glob("zscore_stats_*.json"))), 1)
            # Reconstruct the environmental calibration directly with an independent
            # moment-angle calculation, preserving the published decoy sampling seed.
            rng = np.random.default_rng(7)
            raw_environment = []
            for _ in range(100):
                codes = rng.integers(0, len(residues), size=6).tolist()
                sequence = self.engine.codes_to_seq(codes, residues)
                angles = np.deg2rad(np.arange(6) * 100.0)
                hydro = np.array([self.engine.HYDROPHOBICITY_FP[aa] for aa in sequence])
                mx, my = np.sum(hydro * np.cos(angles)), np.sum(hydro * np.sin(angles))
                phase = 37.0 if math.hypot(mx, my) < 1e-10 else -math.degrees(math.atan2(my, mx)) % 360
                mask = np.abs((np.arange(6) * 100.0 + phase + 180.0) % 360.0 - 180.0) <= 90.0
                raw_environment.append(float(np.mean(np.where(mask, -hydro, hydro))))
            mu, sigma = design.builder.stats["env_pol"]
            self.assertAlmostEqual(mu, float(np.mean(raw_environment)), places=12)
            self.assertAlmostEqual(sigma, float(np.std(raw_environment)), places=12)
            rescored = self.engine.score_sequence(
                "AKGLKA", env, residues, weights, n_decoys=100, n_random=100, seed=7,
            )
            self.assertEqual(rescored["calibration"]["terms"],
                             self.engine.score_with_model("AKGLKA", design, residues, weights)["calibration"]["terms"])
            self.assertEqual(rescored["random_reference"]["mean"], design.reference["mean"])
            self.assertEqual(len(list(Path(cache_dir).glob("zscore_stats_*.json"))), 1)

    def test_every_search_proposal_uses_sequence_dependent_alignment(self):
        """Observe the actual objective evaluations, not only rescoring of final candidates."""
        from unittest.mock import patch

        original = self.engine.compute_breakdown
        evaluations = []

        def observe(builder, codes, weights):
            if builder.kwargs["wheel_halfwidth_deg"] == 90.0:
                self.assertTrue(builder.align_phase)
                mx = sum(self.engine.HYDROPHOBICITY_FP[builder.amino_acids[code]] *
                         math.cos(math.radians(index * 100.0)) for index, code in enumerate(codes))
                my = sum(self.engine.HYDROPHOBICITY_FP[builder.amino_acids[code]] *
                         math.sin(math.radians(index * 100.0)) for index, code in enumerate(codes))
                phase = (builder.kwargs["wheel_phase_deg"] if math.hypot(mx, my) < 1e-10
                         else -math.degrees(math.atan2(my, mx)) % 360.0)
                difference = (builder.phase_for(codes) - phase + 180.0) % 360.0 - 180.0
                self.assertAlmostEqual(difference, 0.0, places=10)
                evaluations.append(tuple(codes))
            return original(builder, codes, weights)

        settings = dict(length=6, residues=["A", "K", "L", "G"], n_decoys=100, steps=20, restarts=1)
        workflows = {
            "Design": lambda: self.engine.design_sequence(environment=self.env_a, **settings),
            "Family": lambda: self.engine.design_family(environment=self.env_a, family_size=2,
                                                       oversample_factor=1, **settings),
            "Cross-Design": lambda: self.engine.cross_environment_design(
                environment_a=self.env_a, environment_b=self.env_b, lambda_gap=0.5, **settings),
            "Specificity": lambda: self.engine.design_specificity(
                target_environment=self.env_a, off_target_environments=[self.env_b],
                num_sequences=1, **settings),
        }
        # Small internal reference populations distinguish the many objective
        # evaluations from the few final-score/reference evaluations in this test.
        with patch.object(self.engine, "compute_breakdown", side_effect=observe), \
                patch.object(self.engine, "DESIGN_N_RANDOM", 5), \
                patch.object(self.engine, "SPECIFICITY_N_RANDOM", 5):
            for name, run in workflows.items():
                with self.subTest(workflow=name):
                    evaluations.clear()
                    run()
                    self.assertGreater(len(evaluations), 20)
                    self.assertGreater(len(set(evaluations)), 5)

    def test_aligned_search_handles_vanishing_moment_and_homogeneous_media(self):
        from unittest.mock import patch

        env = self.engine.resolve_environment({**self.env_a, "wheel_phase_deg": 37.0})
        with patch.object(self.engine, "simulated_annealing", return_value={"sequence": "GGGG"}):
            designed = self.engine.design_sequence(4, env, ["G", "A"], n_decoys=100, steps=1, restarts=1)
        self.assertEqual(designed["environment"]["wheel_phase_deg"], 37.0)
        self.assertFalse(designed["environment"]["phase_aligned_to_hydrophobic_moment"])
        self.assertEqual(designed["design_geometry"]["wheel_phase_deg"], 37.0)
        self.assertEqual(designed["design_geometry"]["orientation"],
                         "preset wheel phase (vanishing hydrophobic moment)")
        self.assertEqual(designed["design_geometry"]["energy"], designed["energy"])
        for preset in ("homogeneous_polar", "homogeneous_apolar"):
            with self.subTest(preset=preset):
                first = self.engine.design_sequence(4, {"preset": preset, "wheel_phase_deg": 37.0},
                                                   ["G", "A"], n_decoys=100, steps=20, restarts=1)
                second = self.engine.design_sequence(4, {"preset": preset, "wheel_phase_deg": 151.0},
                                                    ["G", "A"], n_decoys=100, steps=20, restarts=1)
                self.assertEqual(first["sequence"], second["sequence"])
                self.assertEqual(first["energy"], second["energy"])
                self.assertEqual(first["z_score"], second["z_score"])
                self.assertEqual(first["design_geometry"]["orientation"], "fixed wheel phase")

    def test_specificity_keys_include_orientation_rule_for_partial_sectors(self):
        """An explicitly overridden homogeneous preset remains fixed-phase, unlike an interface."""
        aligned = {"preset": "interfacial_neu", "wheel_halfwidth_deg": 90.0}
        fixed = {"preset": "homogeneous_polar", "wheel_halfwidth_deg": 90.0}
        self.assertNotEqual(self.engine.physical_environment_key(aligned), self.engine.physical_environment_key(fixed))
        result = self.post("/api/specificity-design", {
            **self.design, "target_environment": aligned, "off_target_environments": [fixed], "num_sequences": 1,
        })
        self.assertEqual(len(result["off_target_environments"]), 1)

    def test_pairwise_term_counts_contacts_up_to_i_plus_4(self):
        residues = list(self.engine.DEFAULT_ALPHABET_16)
        builder = self.engine.build_builder(10, residues, self.engine.resolve_environment(self.env_b), n_decoys=100)
        self.assertEqual(builder.kwargs["max_interaction_dist"], 4)
        codes = self.engine.seq_to_codes("AEKLWKEAGH", residues)
        expected = sum(builder.mj_matrix[codes[i], codes[j]] for i in range(10) for j in range(i + 1, min(i + 5, 10)))
        self.assertAlmostEqual(builder._raw_pw_int(codes), expected / (10 * 4))

    def test_workflows_report_consistent_energies(self):
        """All four design workflows optimize exactly the scores displayed and exported."""
        def assert_matches_score(result, n_random=500):
            score_request = {
                **self.common, "sequence": result["sequence"],
                "environment": {"preset": result["environment"]["preset"]}, "n_random": n_random,
            }
            # Specificity uses a 3000-sequence internal reference; the web Score
            # request limit is 2000, so compare its complete reference directly.
            rescored = (self.engine.score_sequence(**score_request) if n_random == 3000
                        else self.post("/api/score", score_request))
            for key in ("energy", "z_score", "breakdown", "random_reference", "calibration"):
                self.assertEqual(result[key], rescored[key], key)
            alias = result["design_geometry"]
            for key in ("energy", "z_score", "breakdown", "percentile_lower_is_better"):
                self.assertEqual(alias[key], result[key], key)
            self.assertEqual(alias["wheel_phase_deg"], result["environment"]["wheel_phase_deg"])
            if self.engine.is_interfacial(result["environment"]):
                self.assertEqual(alias["orientation"], "own hydrophobic moment")
                phase_sum = (result["environment"]["wheel_phase_deg"] +
                             result["observables"]["mu_h_phase_deg"]) % 360
                self.assertAlmostEqual(min(phase_sum, 360.0 - phase_sum), 0.0, places=9)

        designed = self.post("/api/design", self.design)
        assert_matches_score(designed)
        family = self.post("/api/design-family", {**self.design, "family_size": 2, "oversample_factor": 1})
        for member in family["members"]:
            assert_matches_score(member)
        self.assertEqual([member["energy"] for member in family["members"]],
                         sorted(member["energy"] for member in family["members"]))

        environments = {"environment_a": self.env_a, "environment_b": self.env_b}
        cross = self.post("/api/cross-design", {**self.design, **environments, "lambda_gap": 0.7})
        a, b = cross["environment_a"], cross["environment_b"]
        assert_matches_score(a)
        assert_matches_score(b)
        self.assertEqual(cross["cross_design"]["energy_a"], a["energy"])
        self.assertEqual(cross["cross_design"]["energy_b"], b["energy"])
        self.assertAlmostEqual(cross["cross_design"]["objective"],
                               a["energy"] + b["energy"] + 0.7 * abs(a["energy"] - b["energy"]), places=12)

        specificity = self.post("/api/specificity-design", {
            **self.design, "target_environment": self.env_a,
            "off_target_environments": [self.env_b, {"preset": "interfacial_pos"}],
            "lambda_balance": 0.35, "num_sequences": 1,
        })
        for candidate in specificity["candidates"]:
            target = candidate["target_environment"]
            off = candidate["off_target_environments"]
            for scored in [target, *off]:
                assert_matches_score(scored, n_random=3000)
            metrics = candidate["specificity_design"]
            worst = min(item["energy"] for item in off)
            self.assertAlmostEqual(metrics["objective"], target["energy"] - 0.65 * worst, places=12)
            self.assertEqual(metrics["energy_margin"], worst - target["energy"])
            self.assertEqual(metrics["z_score_margin"], min(item["z_score"] for item in off) - target["z_score"])

    def test_invalid_parameter_tables_are_rejected(self):
        import numpy as np
        from data_loaders.energy_matrix_loader import _validate_labeled_matrix

        standard = list("ACDEFGHIKLMNPQRSTVWY")
        tables = {
            "non-finite": (np.full((20, 20), np.nan), standard),
            "1x1": (np.zeros((1, 1)), ["A"]),
            "non-standard label": (np.zeros((20, 20)), standard[:-1] + ["B"]),
        }
        for name, (matrix, symbols) in tables.items():
            with self.subTest(table=name), self.assertRaises(ValueError):
                _validate_labeled_matrix(matrix, symbols, name)

    def test_parameter_tables_have_unique_residue_labels(self):
        from data_loaders import energy_matrix_loader as loader

        canonical = set("ACDEFGHIKLMNPQRSTVWY")
        for load in (loader._load_energy_matrix_file, loader._load_first_neighbors_matrix_file,
                     loader._load_third_neighbors_matrix_file, loader._load_fourth_neighbors_matrix_file):
            with self.subTest(table=load.__name__):
                matrix, symbols = load()
                self.assertEqual(len(symbols), 20)
                self.assertEqual(set(symbols), canonical)
                self.assertEqual(matrix.shape, (20, 20))


if __name__ == "__main__":
    unittest.main()
```
