## Additional file 1. Evaluated software source and dependencies for "Helix Design Studio: designing antimicrobial peptide candidates for membrane environments": test_specificity_progress.py

```
"""Keep streamed specificity progress consistent with its displayed candidate."""

from types import SimpleNamespace
import unittest
from unittest.mock import Mock, patch

import numpy as np

from app import engine


class SpecificityProgressTests(unittest.TestCase):
    def test_progress_objective_belongs_to_selected_penalized_best(self):
        target = SimpleNamespace(L=4)
        off_target = SimpleNamespace(L=4)

        def target_score(codes):
            return -4.0 * max(0, sum(codes) - 2)

        def breakdown(builder, codes, weights):
            return {"total": target_score(codes) if builder is target else 0.0}

        # Restart 1 retains XXXX, objective 0 and no diversity penalty.
        # Restart 2 retains AAAX, raw objective -4 but penalized objective 2,
        # because it lies one mutation from the previously accepted AAAA.
        # Progress must therefore show XXXX together with its objective 0.
        rng = Mock()
        rng.integers.side_effect = [
            np.array([0, 0, 0, 0]), 0, 0,
            np.array([1, 1, 1, 0]), 3, 0,
        ]
        rng.random.return_value = 0.99
        with patch.object(engine, "compute_breakdown", side_effect=breakdown), patch.object(
            engine.np.random, "default_rng", return_value=rng
        ):
            updates = list(engine._iter_specificity_search(
                target_builder=target,
                off_builders=[off_target],
                amino_acids=["X", "A"],
                weights={},
                lambda_balance=1.0,
                steps=1,
                restarts=2,
                seed=42,
                accepted_codes=[[1, 1, 1, 1]],
                min_distance=2,
                penalty_weight=6.0,
            ))

        progress = [event["payload"] for event in updates if event["type"] == "progress"]
        for event in progress:
            codes = [int(aa == "A") for aa in event["best_sequence"]]
            self.assertEqual(event["best_objective"], target_score(codes))
        self.assertEqual(progress[-1]["best_sequence"], "XXXX")
        self.assertEqual(progress[-1]["best_objective"], 0.0)
        final = updates[-1]["payload"]
        self.assertEqual(updates[-1]["type"], "result")
        self.assertEqual(final["sequence"], "XXXX")
        self.assertEqual(final["objective"], 0.0)
        self.assertEqual(final["penalized_objective"], 0.0)


if __name__ == "__main__":
    unittest.main()
```
