## Additional file 1. Evaluated software source and dependencies for "Helix Design Studio: designing antimicrobial peptide candidates for membrane environments": test_table1_conventions.py

```
"""Scoring controls must preserve their declared formula and restore the model."""

from pathlib import Path
import sys
import unittest
from unittest.mock import patch

import numpy as np

ROOT = Path(__file__).resolve().parents[1]
sys.path.insert(0, str(ROOT / "Code"))
from core import hamiltonian_builder as model  # noqa: E402
from tools import reproduce_article_table1 as comparison  # noqa: E402


class TableOneConventionTests(unittest.TestCase):
    def builder(self):
        instance = model.HamiltonianBuilder.__new__(model.HamiltonianBuilder)
        instance.L = 2
        instance.amino_acids = ["A", "G"]
        instance._load_matrices_from_files()
        return instance

    def test_current_and_positive_control_are_explicit_and_restore_state(self):
        original_weights = model.NEIGHBOR_MATRIX_WEIGHTS
        original_convention = model.NEIGHBOR_SCORING_CONVENTION
        original_fingerprint = model.parameter_set_sha256()
        with comparison.m1_convention(model, "current"):
            self.assertAlmostEqual(self.builder()._raw_helix_neigh([0, 0]), -2.734)
            metadata = comparison.convention_metadata(model, "current")
            self.assertFalse(metadata["control_override"])
            self.assertTrue(metadata["neighbor_formula"].startswith("-1*mean(M1)"))
        with comparison.m1_convention(model, "positive_m1"):
            self.assertAlmostEqual(self.builder()._raw_helix_neigh([0, 0]), 2.734)
            metadata = comparison.convention_metadata(model, "positive_m1")
            self.assertTrue(metadata["control_override"])
            self.assertTrue(metadata["neighbor_formula"].startswith("+1*mean(M1)"))
            self.assertNotEqual(metadata["parameters_sha256"], original_fingerprint)
        self.assertIs(model.NEIGHBOR_MATRIX_WEIGHTS, original_weights)
        self.assertEqual(model.NEIGHBOR_SCORING_CONVENTION, original_convention)
        self.assertEqual(model.parameter_set_sha256(), original_fingerprint)

    def test_negative_log_source_zeros_are_rejected_without_side_effects(self):
        original_loader = model.HamiltonianBuilder._load_matrices_from_files
        original_fingerprint = model.parameter_set_sha256()
        with self.assertRaisesRegex(ValueError, "published table contains zeros.*No pseudocount"):
            with comparison.m1_convention(model, "negative_log"):
                self.fail("The zero-containing source table must be rejected before scoring")
        self.assertIs(model.HamiltonianBuilder._load_matrices_from_files, original_loader)
        self.assertEqual(model.parameter_set_sha256(), original_fingerprint)

    def test_negative_log_control_is_not_double_negated(self):
        _, symbols = model._load_first_neighbors_matrix_file()
        positive_source = np.full((len(symbols), len(symbols)), 2.0)
        with patch.object(model, "_load_first_neighbors_matrix_file", return_value=(positive_source, symbols)):
            with comparison.m1_convention(model, "negative_log"):
                self.assertAlmostEqual(self.builder()._raw_helix_neigh([0, 0]), -np.log(2.0))
                metadata = comparison.convention_metadata(model, "negative_log")
                self.assertTrue(metadata["neighbor_formula"].startswith("+1*mean(-ln(M1))"))
                self.assertEqual(metadata["m1_transformation"], "negative_log")

    def test_control_restores_state_after_an_interrupted_calculation(self):
        original_weights = model.NEIGHBOR_MATRIX_WEIGHTS
        original_convention = model.NEIGHBOR_SCORING_CONVENTION
        with self.assertRaisesRegex(RuntimeError, "interrupted"):
            with comparison.m1_convention(model, "positive_m1"):
                raise RuntimeError("interrupted")
        self.assertIs(model.NEIGHBOR_MATRIX_WEIGHTS, original_weights)
        self.assertEqual(model.NEIGHBOR_SCORING_CONVENTION, original_convention)


if __name__ == "__main__":
    unittest.main()
```
