## Additional file 1. Evaluated software source and dependencies for "Helix Design Studio: designing antimicrobial peptide candidates for membrane environments": THIRD_PARTY_NOTICES.md

# Third-party notices and scientific sources

The root MIT license applies to original project code and documentation. It does not replace notices or licenses on included third-party material.

## Included software

- **3Dmol.js 2.4.2**, copyright University of Pittsburgh and contributors, is distributed under BSD-3-Clause. Its license includes notices for GLmol, Three.js and jQuery. The complete upstream text is preserved in [LICENSES/3Dmol.txt](LICENSES/3Dmol.txt), with the companion bundle notice in `app/static/3Dmol-min.js.LICENSE.txt`. The bundle also contains pako (MIT and zlib), UPNG.js (MIT), iobuffer (MIT) and netcdfjs (MIT); their notices are reproduced in [LICENSES/3Dmol-bundled-dependencies.txt](LICENSES/3Dmol-bundled-dependencies.txt).
- `Code/data_loaders/energy_matrix_loader.py` derives from the IBM Qiskit protein-folding data loaders and carries the original **IBM 2021, 2022 / Apache-2.0** notice together with a modification notice. `Code/core/resources/mj_matrix.txt` is the Miyazawa–Jernigan table distributed with Qiskit under the same license. `Code/data_loaders/__init__.py` and `Code/core/resources/__init__.py` retain their original IBM notices. The Apache license is included in [LICENSES/Apache-2.0.txt](LICENSES/Apache-2.0.txt).
- Python dependencies are installed separately from the package index and retain their respective upstream licenses; they are not vendored in this repository.

## Parameter provenance

All tables are looked up by residue label; their headers are validated when loaded. The independent source comparison is reproducible with `python tools/verify_parameter_sources.py`. Published table transcriptions, source PDF checksums, axis definitions and transformation metadata are included in `tools/data/parameter_sources/`. The comparison requires exact equality at the printed precision for all 400 M1 values and the 324 published values in each of M3/M4. Cys/Pro defaults are checked separately.

- `mj_matrix.txt`: Miyazawa S, Jernigan RL. *Residue–residue potentials with a favorable contact pair term and an unfavorable high packing density term, for simulation and threading*. J Mol Biol **256**, 623–644 (1996), Table 3. [doi:10.1006/jmbi.1996.0114](https://doi.org/10.1006/jmbi.1996.0114).
- `helix_pairs_prop.txt` (M1): Nacar C. *Propensities of Amino Acid Pairings in Secondary Structure of Globular Proteins*. Protein J **39**, 21–32 (2020), Figure 3, p. 25. [doi:10.1007/s10930-020-09880-6](https://doi.org/10.1007/s10930-020-09880-6), [primary supplement](https://static-content.springer.com/esm/art%3A10.1007%2Fs10930-020-09880-6/MediaObjects/10930_2020_9880_MOESM1_ESM.pdf). The source matrix reports dimensionless relative propensities for hydrogen-bonded n,n+4 pairs; Figure 3 columns identify the earlier carbonyl acceptor i, and rows the later amide donor j. The application transposes this table to index rows by the earlier residue and columns by the later residue. It reuses the values at i,i+1 as an **empirical model adaptation**; the source does not measure adjacent-residue propensities. All source values, including the zero donor-Pro row (the runtime Pro column), are retained at their printed three-decimal precision. In Methods §2.4 the source defines `Po = FH[j,i]/306352` and `Pr = AH[j]*AH[i]/(1350293*(1350293−1))`; reconstruction from the supplement rounds to all 400 printed Figure 3 values. The [erratum](https://doi.org/10.1007/s10930-020-09881-5) corrects a section cross-reference from §10 to §3 and changes no matrix values.
- `helix_sd_i_3.txt` and `helix_sd_i_4.txt` (M3/M4): Muñoz V, Serrano L. *Elucidating the folding problem of helical peptides using empirical parameters*. Nat Struct Biol **1**, 399–409 (1994), Table 2, p. 403. [doi:10.1038/nsb0694-399](https://doi.org/10.1038/nsb0694-399). The first and second entry for each pair correspond to i,i+3 and i,i+4, respectively, in kcal/mol. Rows identify residue i and columns residue i+k. The printed rows place F before Y, whereas the columns place Y before F; each axis is mapped by its own residue labels. These files reproduce all 324 published entries per separation. The source omits Cys and Pro; pairs containing either residue receive zero contribution in the application. This 76-cell extension per matrix is a **model convention, not experimentally established zero interaction energy**.
- Helix propensities in the scoring code: Pace CN, Scholtz JM. *A helix propensity scale based on experimental studies of peptides and proteins*. Biophys J **75**, 422–427 (1998).
- Hydrophobicity values in the scoring code: Fauchère JL, Pliska V. *Hydrophobic parameters II of amino acid side-chains from the partitioning of N-acetyl-amino acid amides*. Eur J Med Chem **18** (1983).

The neighbor-scoring convention is `negative_m1_plus_m3_plus_m4`: `−mean(M1) + mean(M3) + mean(M4)`, with each mean taken over the L−k pairs at its sequence separation. The source M1 table remains unchanged. Subtracting its odds ratios makes larger helical enrichments favorable in the minimized score; a zero source entry contributes zero to the M1 sum. M3/M4 retain their source energy signs, with negative values favorable. These three means have fixed internal coefficients −1, +1 and +1 and are standardized together against calibration sequences. Combining numerical values from different source units and structural contexts defines an empirical descriptor with fixed internal scaling, not an absolute thermodynamic energy. The convention and internal coefficients are included in calibration metadata and in the parameter-set checksum. See [model scope](README.md#model-scope-and-limitations).

Named peptide examples in the interface are demonstration inputs.

## Branding

The SIMBIOS and Universidade de Santiago de Compostela names and logos identify the project and institution. They are excluded from the MIT grant; this distribution grants no trademark rights or endorsement. Replace these assets and institutional presentation when deploying an unrelated service.
