## Additional file 1. Evaluated software source and dependencies for "Helix Design Studio: designing antimicrobial peptide candidates for membrane environments": measure_changes.py

```
"""Measure the numerical effect of each model change between 0.1.0 and the current candidate.

Each variant restores one 0.1.0 behaviour on top of the current code with the other settings
held fixed. With --baseline, the complete 0.1.0 source tree is
also run for an end-to-end comparison of the scoring benchmark.

    python tools/measure_changes.py --baseline /path/to/helix-design-studio-web-0.1.0

Benchmark: the reference peptides of the interface scored in the five environment presets
with the full alphabet, the mean percentile of random sequences in an interfacial
environment, and designs with fixed seeds. Sample sizes are set below; results are printed
as Markdown tables.
"""

from __future__ import annotations

import argparse
import contextlib
import hashlib
import io
import json
import os
import platform
import subprocess
import sys
import tempfile
from pathlib import Path

import numpy as np

REPO = Path(__file__).resolve().parents[1]
sys.path.insert(0, str(REPO))

N_DECOYS = 1000
N_RANDOM = 500
SEED = 42
ALPHABET = list("ARNDCQEGHILKMFPSTWYV")
PRESETS = ["homogeneous_polar", "homogeneous_apolar", "interfacial_neg", "interfacial_neu", "interfacial_pos"]
PEPTIDES = {
    "(EAAAK)3": "EAAAKEAAAKEAAAK",
    "(AAKAA)3": "AAKAAAAKAAAAKAA",
    "LL-37": "LLGDFFRKSKEKIGKEFKRIVQRIKDFLRNLVPRTES",
    "Melittin": "GIGAVLKVLTTGLPALISWIKRKRQQ",
    "Magainin 2": "GIGKFLHSAKKFGKAFVGEIMNS",
    "Cecropin A": "KWKLFKKIEKVGQNIRDGIIKAGPAVAVVGQATQIAK",
    "PGLa": "GMASKAGAIAGKIAKVALKAL",
    "Piscidin 1": "FFHHIFRGIVHVGKTIHRLVTG",
    "BP100": "KKLFKKILKYL",
    "Poly-Ala 15": "AAAAAAAAAAAAAAA",
    "KALP17": "GKKLALALALALALKKA",
    "WALP23": "GWWLALALALALALALALALWWA",
}
RANDOM_LENGTH = 18
RANDOM_COUNT = 150
DESIGN = {"length": 18, "steps": 4000, "restarts": 2, "seeds": (1, 2, 3), "presets": ("interfacial_neg", "homogeneous_polar")}


def quiet():
    return contextlib.redirect_stdout(io.StringIO())


def load_current():
    os.environ["HELIX_CACHE_DIR"] = tempfile.mkdtemp(prefix="helix-measure-")
    with quiet():
        from app import engine
        from core import hamiltonian_builder as model
    return engine, model


engine, model = load_current()


# ----------------------------------------------------------------------------- 0.1.0 behaviours
@contextlib.contextmanager
def positional_tables():
    """0.1.0 indexed M1/M3/M4 by position in the requested alphabet (file order ignored)."""
    from data_loaders import energy_matrix_loader as loader

    original = model.HamiltonianBuilder._load_matrices_from_files

    def patched(self):
        original(self)
        n = self.n_aa
        self.M1 = np.array(loader._load_first_neighbors_matrix_file()[0])[:n, :n]
        self.M3 = np.array(loader._load_third_neighbors_matrix_file()[0])[:n, :n]
        self.M4 = np.array(loader._load_fourth_neighbors_matrix_file()[0])[:n, :n]

    model.HamiltonianBuilder._load_matrices_from_files = patched
    try:
        yield
    finally:
        model.HamiltonianBuilder._load_matrices_from_files = original


@contextlib.contextmanager
def patched_values(table, values):
    saved = {key: table[key] for key in values}
    table.update(values)
    try:
        yield
    finally:
        table.update(saved)


@contextlib.contextmanager
def patched_attribute(obj, name, value):
    saved = getattr(obj, name)
    setattr(obj, name, value)
    try:
        yield
    finally:
        setattr(obj, name, saved)


def query_only_orientation_score(sequence, environment, residues, weights=None, n_decoys=2000, n_random=500, seed=42):
    """0.1.0 scoring: the query was oriented by its hydrophobic moment; calibration decoys and
    random references were evaluated at that fixed phase, without orienting them."""
    env = engine.resolve_environment(environment)
    if engine.is_interfacial(env):
        phase = model.aligned_phase_deg(sequence)
        if phase is not None:
            env["wheel_phase_deg"] = phase
    w = engine.normalize_weights(weights)
    fixed = engine.scoring_model(len(sequence), list(residues), env, w, n_decoys=n_decoys, seed=seed,
                                 n_random=n_random, oriented=False)
    return engine.score_with_model(sequence, fixed, list(residues), w)


VARIANTS = {
    "Table indexing by alphabet position": positional_tables,
    "Ser/Thr hydrophobicities swapped": lambda: patched_values(model.FAUCHERE_PLISKA_HYDROPHOBICITY, {"S": 0.26, "T": -0.04}),
    "Pro helix propensity 3.15": lambda: patched_values(model.PACE_SCHOLTZ_HELIX_PROPENSITY, {"P": 3.15}),
    "MJ contacts up to i+1 (d_max = 1)": lambda: patched_attribute(engine, "MJ_MAX_DISTANCE", 1),
    "Only the query oriented in scoring": lambda: patched_attribute(engine, "score_sequence", query_only_orientation_score),
}
SCORING_ONLY = {"Only the query oriented in scoring"}


# ----------------------------------------------------------------------------- benchmarks
def score_benchmark():
    rows = {}
    for name, sequence in PEPTIDES.items():
        for preset in PRESETS:
            with quiet():
                result = engine.score_sequence(sequence, {"preset": preset}, ALPHABET,
                                               n_decoys=N_DECOYS, n_random=N_RANDOM, seed=SEED)
            rows[f"{name} | {preset}"] = result["z_score"]
    return rows


def random_percentile():
    rng = np.random.default_rng(7)
    values = []
    for _ in range(RANDOM_COUNT):
        sequence = "".join(rng.choice(ALPHABET, size=RANDOM_LENGTH))
        with quiet():
            result = engine.score_sequence(sequence, {"preset": "interfacial_neg"}, ALPHABET,
                                           n_decoys=N_DECOYS, n_random=N_RANDOM, seed=SEED)
        values.append(result["percentile_lower_is_better"])
    return float(np.mean(values))


@contextlib.contextmanager
def search_seed_offset(offset):
    """Change only the search stream; retain calibration and reference seeds."""
    original = engine.rng_seed

    def adjusted(seed, stream, *index):
        return original(seed + offset if stream == engine.SEARCH_STREAM else seed, stream, *index)

    with patched_attribute(engine, "rng_seed", adjusted):
        yield


def designs(search_offset=0):
    out = {}
    with search_seed_offset(search_offset):
        for preset in DESIGN["presets"]:
            for seed in DESIGN["seeds"]:
                with quiet():
                    result = engine.design_sequence(DESIGN["length"], {"preset": preset}, ALPHABET, n_decoys=N_DECOYS,
                                                    steps=DESIGN["steps"], restarts=DESIGN["restarts"], seed=seed)
                out[(preset, seed)] = result["sequence"]
    return out


def spearman(a, b):
    ranks = lambda x: np.argsort(np.argsort(x))
    return float(np.corrcoef(ranks(np.array(a)), ranks(np.array(b)))[0, 1])


def compare_scores(reference, other):
    keys = list(reference)
    delta = np.array([other[k] - reference[k] for k in keys])
    worst = keys[int(np.argmax(np.abs(delta)))]
    return {
        "median_abs_dz": float(np.median(np.abs(delta))),
        "max_abs_dz": float(np.max(np.abs(delta))),
        "max_case": worst,
        "spearman": spearman([reference[k] for k in keys], [other[k] for k in keys]),
    }


def changed_positions(reference, other):
    fractions = [sum(x != y for x, y in zip(reference[key], other[key])) / len(reference[key]) for key in reference]
    return float(np.mean(fractions))


def baseline_scores(path):
    """Run the scoring benchmark with a complete 0.1.0 source tree in a separate interpreter."""
    code = f"""
import contextlib, io, json, sys, numpy as np
sys.path.insert(0, {str(path)!r})
with contextlib.redirect_stdout(io.StringIO()):
    from app.engine import score_sequence
peptides, presets, alphabet = {json.dumps(PEPTIDES)}, {json.dumps(PRESETS)}, {json.dumps(ALPHABET)}
out = {{}}
with contextlib.redirect_stdout(io.StringIO()):
    for name, seq in peptides.items():
        for preset in presets:
            out[name + " | " + preset] = score_sequence(seq, {{"preset": preset}}, alphabet, n_decoys={N_DECOYS},
                                                        n_random={N_RANDOM}, seed={SEED})["z_score"]
    rng = np.random.default_rng(7)
    pct = [score_sequence("".join(rng.choice(alphabet, size={RANDOM_LENGTH})), {{"preset": "interfacial_neg"}},
                          alphabet, n_decoys={N_DECOYS}, n_random={N_RANDOM}, seed={SEED})["percentile_lower_is_better"]
           for _ in range({RANDOM_COUNT})]
print(json.dumps({{"scores": out, "random_percentile": float(np.mean(pct))}}))
"""
    env = {**os.environ, "HELIX_CACHE_DIR": tempfile.mkdtemp(prefix="helix-baseline-")}
    completed = subprocess.run([sys.executable, "-c", code], cwd=path, env=env, capture_output=True, text=True, check=True)
    return json.loads(completed.stdout.strip().splitlines()[-1])


def main():
    parser = argparse.ArgumentParser(description=__doc__.split("\n\n")[0])
    parser.add_argument("--baseline", type=Path, help="checkout of the 0.1.0 source tree for an end-to-end comparison")
    parser.add_argument("--output", type=Path, help="save protocol, scores and summary statistics as JSON")
    args = parser.parse_args()

    current_scores = score_benchmark()
    current_percentile = random_percentile()
    current_designs = designs()
    alternate_designs = designs(search_offset=100)
    search_control = changed_positions(current_designs, alternate_designs)
    design_rows = lambda values: [{"preset": preset, "seed": seed, "sequence": sequence}
                                  for (preset, seed), sequence in values.items()]
    report = {
        "protocol": {"alphabet": ALPHABET, "presets": PRESETS, "peptides": PEPTIDES,
                     "n_decoys": N_DECOYS, "n_random": N_RANDOM, "score_seed": SEED,
                     "weights": engine.DEFAULT_WEIGHTS,
                     "random_sequences": {"count": RANDOM_COUNT, "length": RANDOM_LENGTH, "seed": 7},
                     "design": DESIGN, "search_control_offset": 100,
                     "search_control": "Only SEARCH_STREAM changes from [seed,2] to [seed+100,2]; calibration and reference seeds remain fixed.",
                     "seed_protocol": engine.SEED_PROTOCOL},
        "runtime": {"python": platform.python_version(), "numpy": np.__version__},
        "source_files_sha256": {name: hashlib.sha256((REPO / name).read_bytes()).hexdigest()
                                for name in ("app/engine.py", "Code/core/hamiltonian_builder.py", "Code/data_loaders/energy_matrix_loader.py")},
        "parameters_sha256": model.parameter_set_sha256(),
        "current_scores": current_scores,
        "current_random_percentile": current_percentile,
        "current_designs": design_rows(current_designs),
        "alternate_search_designs": design_rows(alternate_designs),
        "search_control_changed_fraction": search_control,
        "variants": {},
    }

    print(f"Benchmark: {len(PEPTIDES)} reference peptides x {len(PRESETS)} presets, full alphabet, "
          f"n_decoys={N_DECOYS}, n_random={N_RANDOM}, seed={SEED}; random sequences: {RANDOM_COUNT} x L={RANDOM_LENGTH} "
          f"(interfacial_neg); designs: L={DESIGN['length']}, {DESIGN['steps']} steps x {DESIGN['restarts']} restarts, "
          f"seeds {DESIGN['seeds']}, presets {DESIGN['presets']}.\n")
    print(f"Current version: mean percentile of random sequences {current_percentile:.1f} (unbiased: 50). "
          f"Search-stream control: with calibration and random references fixed, designs using search entropy "
          f"[seed+100,2] instead of [seed,2] differ in {100 * search_control:.0f}% of positions. "
          "This descriptive control does not attribute model-change effects to search randomness.\n")
    print("| Restored 0.1.0 behaviour | median abs dZ | max abs dZ (case) | Spearman of Z | random percentile | design positions changed |")
    print("|---|---|---|---|---|---|")
    for label, variant in VARIANTS.items():
        with variant():
            scores = score_benchmark()
            percentile = random_percentile()
            variant_designs = None if label in SCORING_ONLY else designs()
            fraction = None if variant_designs is None else changed_positions(current_designs, variant_designs)
            changed = "n/a (scoring only)" if fraction is None else f"{100 * fraction:.0f}%"
        stats = compare_scores(current_scores, scores)
        report["variants"][label] = {**stats, "scores": scores, "random_percentile": percentile,
                                      "design_changed_fraction": fraction,
                                      "designs": None if variant_designs is None else design_rows(variant_designs)}
        print(f"| {label} | {stats['median_abs_dz']:.3f} | {stats['max_abs_dz']:.3f} ({stats['max_case']}) | "
              f"{stats['spearman']:.3f} | {percentile:.1f} | {changed} |")
    if args.baseline:
        baseline = baseline_scores(args.baseline.resolve())
        stats = compare_scores(current_scores, baseline["scores"])
        report["baseline"] = {**baseline, **stats}
        print(f"| Complete 0.1.0 source tree | {stats['median_abs_dz']:.3f} | {stats['max_abs_dz']:.3f} ({stats['max_case']}) | "
              f"{stats['spearman']:.3f} | {baseline['random_percentile']:.1f} | not compared (different random streams) |")
    if args.output:
        args.output.parent.mkdir(parents=True, exist_ok=True)
        args.output.write_text(json.dumps(report, indent=2, ensure_ascii=False) + "\n", encoding="utf-8")


if __name__ == "__main__":
    main()
```
