## Additional file 1. Evaluated software source and dependencies for "Helix Design Studio: designing antimicrobial peptide candidates for membrane environments": measure_parameter_corrections.py

```
"""Measure M1/M4 restoration while holding the scoring/search implementation fixed.

Extract tag v1.0.0rc1 into a separate directory, then run:
    python tools/measure_parameter_corrections.py --baseline /path/to/1.0.0rc1 --output comparison.json
"""
from __future__ import annotations

import argparse
import contextlib
import hashlib
import json
import os
from pathlib import Path
import platform
import tempfile

import numpy as np
import measure_changes as benchmark


def sha(path):
    return hashlib.sha256(path.read_bytes()).hexdigest()


def read_matrix(path):
    lines = [line.split() for line in path.read_text().splitlines() if line.strip()]
    labels = lines[0]
    matrix = np.asarray(lines[1:], dtype=float)
    assert matrix.shape == (20, 20) and len(set(labels)) == 20
    return matrix, labels


@contextlib.contextmanager
def parameter_case(overrides):
    """Override only numerical arrays; each builder fingerprints its effective arrays."""
    original = benchmark.model.HamiltonianBuilder._load_matrices_from_files
    def load(self):
        original(self)
        for name, (matrix, labels) in overrides.items():
            setattr(self, name, self._matrix_for_alphabet(matrix, labels))
    benchmark.model.HamiltonianBuilder._load_matrices_from_files = load
    previous = os.environ.get('HELIX_CACHE_DIR')
    try:
        with tempfile.TemporaryDirectory(prefix='helix-parameter-case-') as cache:
            os.environ['HELIX_CACHE_DIR'] = cache
            yield
    finally:
        benchmark.model.HamiltonianBuilder._load_matrices_from_files = original
        if previous is None:
            os.environ.pop('HELIX_CACHE_DIR', None)
        else:
            os.environ['HELIX_CACHE_DIR'] = previous


def ranks(values):
    values = np.asarray(values)
    return np.array([np.flatnonzero(np.sort(values) == v).mean() for v in values])


def summarize(reference, current):
    keys = list(reference['scores'])
    delta = np.array([current['scores'][k]-reference['scores'][k] for k in keys])
    old_design = {(r['preset'], r['seed']): r['sequence'] for r in reference['designs']}
    new_design = {(r['preset'], r['seed']): r['sequence'] for r in current['designs']}
    differences = [sum(a != b for a,b in zip(old_design[k], new_design[k])) for k in old_design]
    return {'score_count': len(delta), 'median_absolute_delta_z': float(np.median(abs(delta))),
            'maximum_absolute_delta_z': float(max(abs(delta))),
            'maximum_delta_case': keys[int(np.argmax(abs(delta)))],
            'spearman_z': float(np.corrcoef(ranks(list(reference['scores'].values())),
                                          ranks(list(current['scores'].values())))[0,1]),
            'design_count': len(differences), 'design_positions_changed': int(sum(differences)),
            'design_positions_evaluated': sum(map(len, old_design.values())),
            'design_positions_changed_fraction': float(np.mean([d/len(old_design[k]) for d,k in zip(differences,old_design)]))}


def main():
    parser = argparse.ArgumentParser(description=__doc__.split('\n\n')[0])
    parser.add_argument('--baseline', type=Path, required=True)
    parser.add_argument('--output', type=Path, required=True)
    args = parser.parse_args()
    root = benchmark.REPO
    old_paths = {name:args.baseline/'Code/core/resources'/file for name,file in
                 [('M1','helix_pairs_prop.txt'), ('M4','helix_sd_i_4.txt')]}
    expected = {'M1':'92208ffa4172357f928950f674163d42bd3786838a551255f04ae4670b5c7359',
                'M4':'44c54eb5ec3ea972fd029da96ba51db8a78dddd8e1b5d0a593d48c3e49872045'}
    assert {name:sha(path) for name,path in old_paths.items()} == expected, 'Expected v1.0.0rc1 parameter files'
    old = {name:read_matrix(path) for name,path in old_paths.items()}
    files = ['app/engine.py','Code/core/hamiltonian_builder.py','Code/data_loaders/energy_matrix_loader.py',
             'Code/core/resources/helix_pairs_prop.txt','Code/core/resources/helix_sd_i_3.txt',
             'Code/core/resources/helix_sd_i_4.txt']
    fingerprints = {name:sha(root/name) for name in files}
    report = {'comparison': 'Paired parameter restoration; current engine in every case',
              'baseline_parameter_version':'1.0.0rc1', 'restored_parameter_version':'1.0.0rc2',
              'baseline_parameter_sha256':expected, 'current_source_sha256':fingerprints,
              'runtime':{'python':platform.python_version(), 'numpy':np.__version__},
              'protocol':{'alphabet':benchmark.ALPHABET, 'peptides':benchmark.PEPTIDES,
                          'presets':benchmark.PRESETS, 'weights':benchmark.engine.DEFAULT_WEIGHTS,
                          'n_decoys':benchmark.N_DECOYS, 'n_random':benchmark.N_RANDOM,
                          'score_seed':benchmark.SEED, 'design':benchmark.DESIGN,
                          'random_sequence_count':benchmark.RANDOM_COUNT, 'random_sequence_length':benchmark.RANDOM_LENGTH,
                          'random_sequence_seed':7, 'seed_protocol':benchmark.engine.SEED_PROTOCOL,
                          'calibration':'Recomputed for each parameter case with the same sampled sequences; empty case-specific caches',
                          'interpretation':'Design differences are outcomes of paired searches under changed objectives, not experimental effects'},
              'cases':{}, 'effects':{}}
    cases = [('previous_parameters',old), ('M1_restored',{'M4':old['M4']}),
             ('M4_restored',{'M1':old['M1']}), ('both_restored',{})]
    for name,overrides in cases:
        with parameter_case(overrides):
            design = benchmark.designs()
            report['cases'][name] = {'scores':benchmark.score_benchmark(),
                                    'random_sequence_mean_percentile':benchmark.random_percentile(),
                                    'designs':[{'preset':preset,'seed':seed,'sequence':seq}
                                               for (preset,seed),seq in design.items()]}
        print(f'Completed {name}', flush=True)
    for name in ('M1_restored','M4_restored','both_restored'):
        report['effects'][name] = summarize(report['cases']['previous_parameters'],report['cases'][name])
    assert fingerprints == {name:sha(root/name) for name in files}, 'Runtime files changed during measurement'
    args.output.parent.mkdir(parents=True, exist_ok=True)
    args.output.write_text(json.dumps(report, indent=2, ensure_ascii=False, allow_nan=False)+'\n')
    print(json.dumps(report['effects'], indent=2))


if __name__ == '__main__':
    main()
```
