## Additional file 1. Evaluated software source and dependencies for "Helix Design Studio: designing antimicrobial peptide candidates for membrane environments": reproduce_article_table1.py

```
"""Rescore the exact sequence strings and Z-scores of the method article's Table 1.

This is a specified rerun with current software, not a reconstruction of the historical
calculation. Optional scoring controls are identified separately from the current model.
The bundled input records its manuscript source and transcription checksums.

    python tools/reproduce_article_table1.py --output article_table1_comparison.json
"""
from __future__ import annotations

import argparse
import contextlib
import hashlib
import json
import os
from pathlib import Path
import platform
import re
import sys
import tempfile

import numpy as np

ROOT = Path(__file__).resolve().parents[1]
sys.path.insert(0, str(ROOT))
DEFAULT_INPUT = ROOT / "tools" / "data" / "article_table1.json"
ALPHABET = list("ARNDCQEGHILKMFPSTWYV")
PRESETS = ("homogeneous_polar", "homogeneous_apolar")
WEIGHTS = {"lambda_env": 6.0, "lambda_charge": 3.0, "lambda_local": 1.5,
           "lambda_pairwise": 0.1, "lambda_helix_pairs": 1.5, "lambda_electrostatic": 4.0}


def file_sha256(path):
    return hashlib.sha256(path.read_bytes()).hexdigest()


def source_fingerprints():
    files = ("app/engine.py", "Code/core/hamiltonian_builder.py", "Code/data_loaders/energy_matrix_loader.py")
    return {name: file_sha256(ROOT / name) for name in files}


@contextlib.contextmanager
def m1_convention(model, convention):
    """Apply an explicit, isolated scoring control and restore the current model."""
    validate_m1_convention(model, convention)
    original_loader = model.HamiltonianBuilder._load_matrices_from_files
    original_weights = model.NEIGHBOR_MATRIX_WEIGHTS
    original_convention = model.NEIGHBOR_SCORING_CONVENTION
    try:
        if convention != "current":
            model.NEIGHBOR_MATRIX_WEIGHTS = {**original_weights, 1: 1.0}
            model.NEIGHBOR_SCORING_CONVENTION = (
                "positive_m1_plus_m3_plus_m4" if convention == "positive_m1"
                else "negative_log_m1_plus_m3_plus_m4"
            )
        if convention == "negative_log":
            def load(self):
                original_loader(self)
                self.M1 = -np.log(self.M1)
            # Add -ln(M1) with coefficient +1; subtracting it would invert the control.
            model.HamiltonianBuilder._load_matrices_from_files = load
        yield
    finally:
        model.HamiltonianBuilder._load_matrices_from_files = original_loader
        model.NEIGHBOR_MATRIX_WEIGHTS = original_weights
        model.NEIGHBOR_SCORING_CONVENTION = original_convention


def validate_m1_convention(model, convention):
    if convention not in {"current", "positive_m1", "negative_log"}:
        raise ValueError(f"Unknown scoring convention: {convention}")
    if convention == "negative_log":
        source, _ = model._load_first_neighbors_matrix_file()
        if not np.all(np.isfinite(source) & (source > 0)):
            raise ValueError(
                "The negative-log control requires finite, strictly positive M1 source entries; "
                "the published table contains zeros. No pseudocount or smoothing is applied."
            )


def convention_metadata(model, convention):
    coefficients = {f"M{k}": value for k, value in model.NEIGHBOR_MATRIX_WEIGHTS.items()}
    terms = [
        f"{value:+g}*mean(-ln(M1))" if k == 1 and convention == "negative_log"
        else f"{value:+g}*mean(M{k})"
        for k, value in model.NEIGHBOR_MATRIX_WEIGHTS.items()
    ]
    return {
        "neighbor_score_convention": model.NEIGHBOR_SCORING_CONVENTION,
        "neighbor_matrix_weights": coefficients,
        "m1_transformation": "negative_log" if convention == "negative_log" else "identity",
        "neighbor_formula": " ".join(terms),
        "formula_operands": "M1, M3 and M4 in the formula refer to the stored source coefficients",
        "control_override": convention != "current",
        "parameters_sha256": model.parameter_set_sha256(),
    }


def main():
    parser = argparse.ArgumentParser(description=__doc__.split("\n\n")[0])
    parser.add_argument("--input", type=Path, default=DEFAULT_INPUT)
    parser.add_argument("--output", type=Path, required=True)
    parser.add_argument("--n-decoys", type=int, default=2000)
    parser.add_argument("--n-random", type=int, default=10000)
    parser.add_argument("--seed", type=int, default=42)
    parser.add_argument("--include-negative-log", action="store_true",
                        help="Also add mean(-ln(M1)) as a control; rejects zeros without smoothing")
    parser.add_argument("--include-positive-m1", action="store_true",
                        help="Also add mean(M1), the earlier scoring convention, as a controlled comparison")
    args = parser.parse_args()
    if args.n_decoys < 2 or args.n_random < 2 or not 0 <= args.seed <= 2**32 - 1:
        parser.error("Sample sizes must be at least 2 and seed must be between 0 and 2**32-1")
    inputs = json.loads(args.input.read_text(encoding="utf-8"))
    source_rows = "\n".join(inputs["source"]["source_rows"]) + "\n"
    if hashlib.sha256(source_rows.encode("utf-8")).hexdigest() != inputs["source"]["source_rows_utf8_sha256"]:
        raise ValueError("Source-row transcription checksum mismatch")
    row_pattern = re.compile(r"\\texttt\{([^}]+)\} & ([^&]+) & (\d+) & \\texttt\{([A-Z]+)\} & ([-\d.]+) & ([-\d.]+)")
    transcribed = {}
    for row in inputs["source"]["source_rows"]:
        match = row_pattern.search(row)
        if match is None:
            raise ValueError("Could not parse a transcribed table row")
        transcribed[match[1]] = (int(match[3]), match[4], float(match[5]), float(match[6]))
    if set(transcribed) != {peptide["name"] for peptide in inputs["peptides"]}:
        raise ValueError("Sequence names differ from the source-row transcription")
    for peptide in inputs["peptides"]:
        if len(peptide["sequence"]) != peptide["evaluated_length"]:
            raise ValueError("Evaluated sequence length does not match the input record")
        values = (peptide["printed_length"], peptide["sequence"], peptide["published_z"][PRESETS[0]], peptide["published_z"][PRESETS[1]])
        if transcribed[peptide["name"]] != values:
            raise ValueError("An input sequence or reference score differs from its source row")
    sources = source_fingerprints()
    with tempfile.TemporaryDirectory(prefix="helix-table1-") as cache:
        os.environ["HELIX_CACHE_DIR"] = cache
        from app import engine
        from app.main import APP_VERSION
        from core import hamiltonian_builder as model
        conventions = ["current"]
        if args.include_positive_m1:
            conventions.append("positive_m1")
        if args.include_negative_log:
            conventions.append("negative_log")
        # Reject undefined optional transforms before any scoring runs are performed.
        for convention in conventions:
            validate_m1_convention(model, convention)
        report = {
            "comparison": "Current software using exact method-article Table 1 sequence strings",
            "software_version": APP_VERSION,
            "input_sha256": file_sha256(args.input),
            "source": inputs["source"],
            "notes": inputs["notes"],
            "source_files_sha256": sources,
            "parameters_sha256": model.parameter_set_sha256(),
            "current_scoring_convention": convention_metadata(model, "current"),
            "runtime": {"python": platform.python_version(), "numpy": np.__version__},
            "protocol": {"alphabet": ALPHABET, "environments": PRESETS, "weights": WEIGHTS,
                         "n_decoys": args.n_decoys, "n_random": args.n_random, "seed": args.seed,
                         "calibration_seed_entropy": [args.seed], "reference_seed_entropy": engine.rng_seed(args.seed, engine.REFERENCE_STREAM),
                         "seed_protocol": engine.SEED_PROTOCOL, "mj_max_distance": engine.MJ_MAX_DISTANCE,
                         "neighbor_normalization": "Average each separation over L-k pairs, apply its declared internal coefficient, then standardize the combined helix_neigh term",
                         "comparison_scope": ("Only the declared M1 scoring convention differs between controls; calibration is recomputed using the same random sequences. These are current-engine controls, not complete reconstructions of historical software."
                                              if len(conventions) > 1 else
                                              "Current-model rerun; deviations from published scores do not reconstruct their historical calculation protocol")},
            "variants": {},
        }
        for convention in conventions:
            records = []
            with m1_convention(model, convention):
                metadata = convention_metadata(model, convention)
                for peptide in inputs["peptides"]:
                    for preset in PRESETS:
                        result = engine.score_sequence(peptide["sequence"], {"preset": preset}, ALPHABET,
                                                       weights=WEIGHTS, n_decoys=args.n_decoys,
                                                       n_random=args.n_random, seed=args.seed)
                        expected = peptide["published_z"][preset]
                        records.append({"name": peptide["name"], "sequence": peptide["sequence"],
                                        "printed_length": peptide["printed_length"], "evaluated_length": len(peptide["sequence"]),
                                        "environment": preset, "published_z": expected,
                                        "computed_z": result["z_score"], "delta_z": result["z_score"] - expected})
            differences = np.array([row["delta_z"] for row in records])
            report["variants"][convention] = {**metadata, "software_version": APP_VERSION,
                                               "rows": records, "mean_absolute_delta_z": float(np.abs(differences).mean()),
                                               "max_absolute_delta_z": float(np.abs(differences).max()),
                                               "root_mean_square_delta_z": float(np.sqrt(np.mean(differences**2)))}
        if source_fingerprints() != sources or model.parameter_set_sha256() != report["parameters_sha256"]:
            raise RuntimeError("Source or parameter files changed during the calculation; repeat on a fixed version")
    args.output.parent.mkdir(parents=True, exist_ok=True)
    args.output.write_text(json.dumps(report, indent=2, ensure_ascii=False, allow_nan=False) + "\n", encoding="utf-8")
    for convention, result in report["variants"].items():
        print(f"{convention}: mean |ΔZ| = {result['mean_absolute_delta_z']:.6f}; max |ΔZ| = {result['max_absolute_delta_z']:.6f}")


if __name__ == "__main__":
    main()
```
