## Additional file 1. Evaluated software source and dependencies for "Helix Design Studio: designing antimicrobial peptide candidates for membrane environments": verify_parameter_sources.py

```
#!/usr/bin/env python3
"""Compare runtime neighbor matrices with independently transcribed source tables.

Uses only the Python standard library. Source CSVs retain their published axis
orders, including the unequal F/Y row and column order in Muñoz–Serrano Table 2.
The comparison distinguishes published cells from the C/P zero extension.
"""

import argparse
import csv
from decimal import Decimal
import hashlib
import json
from pathlib import Path


REPOSITORY = Path(__file__).resolve().parents[1]
SOURCE_DIRECTORY = REPOSITORY / "tools" / "data" / "parameter_sources"
RESOURCES_DIRECTORY = REPOSITORY / "Code" / "core" / "resources"
STANDARD_RESIDUES = set("ARNDCQEGHILKMFPSTWYV")


def _labels(values):
    if len(values) != len(set(values)) or not set(values) <= STANDARD_RESIDUES:
        raise ValueError("Matrix labels must be unique standard residue codes")
    return values


def _value(token):
    value = Decimal(token)
    if not value.is_finite():
        raise ValueError("Matrix entries must be finite")
    return value


def load_source_csv(path):
    """Return values keyed by each independently labelled source axis."""
    with Path(path).open(newline="", encoding="utf-8") as handle:
        records = list(csv.reader(handle))
    columns = _labels(records[0][1:])
    rows = _labels([record[0] for record in records[1:]])
    values = {}
    for row, record in zip(rows, records[1:]):
        if len(record) != len(columns) + 1:
            raise ValueError(f"Wrong number of source columns for {row}")
        for column, token in zip(columns, record[1:]):
            values[row, column] = _value(token)
    return rows, columns, values


def load_runtime(path):
    """Runtime files use their one header for both row and column labels."""
    lines = Path(path).read_text(encoding="utf-8").splitlines()
    tokens = lines[0].split()
    labels = _labels(list(tokens[0]) if len(tokens) == 1 else tokens)
    if len(lines) != len(labels) + 1:
        raise ValueError("Wrong number of runtime rows")
    values = {}
    for row, line in zip(labels, lines[1:]):
        tokens = line.split()
        if len(tokens) != len(labels):
            raise ValueError(f"Wrong number of runtime columns for {row}")
        for column, token in zip(labels, tokens):
            values[row, column] = _value(token)
    if set(labels) != STANDARD_RESIDUES:
        raise ValueError("Runtime neighbor matrices must cover all 20 residues")
    return labels, values


def verify_matrix(name, description, resources_directory, source_directory):
    source_path = Path(source_directory) / description["source_csv"]
    source_hash = hashlib.sha256(source_path.read_bytes()).hexdigest()
    if source_hash != description["source_csv_sha256"]:
        raise ValueError(f"{name}: source CSV checksum differs from metadata")
    rows, columns, published = load_source_csv(source_path)
    if rows != list(description["source_row_order"]):
        raise ValueError(f"{name}: source row order differs from metadata")
    if columns != list(description["source_column_order"]):
        raise ValueError(f"{name}: source column order differs from metadata")
    runtime_path = Path(resources_directory) / description["runtime_file"]
    labels, runtime = load_runtime(runtime_path)
    mapping = description["mapping"]
    if mapping == "transpose":
        expected = {(column, row): value for (row, column), value in published.items()}
    elif mapping == "by_labels":
        expected = published
    else:
        raise ValueError(f"{name}: unknown source mapping {mapping!r}")
    differences = []
    for (row, column), value in expected.items():
        actual = runtime[row, column]
        if actual != value:
            differences.append({"residue_i": row, "residue_i_plus_k": column,
                                "source": str(value), "runtime": str(actual)})
    missing = set(description["missing_residues"])
    if set(rows) != STANDARD_RESIDUES - missing or set(columns) != STANDARD_RESIDUES - missing:
        raise ValueError(f"{name}: source residue coverage differs from metadata")
    extensions = [(row, column) for row in labels for column in labels
                  if (row, column) not in expected]
    extension_errors = [{"residue_i": row, "residue_i_plus_k": column,
                         "runtime": str(runtime[row, column])}
                        for row, column in extensions if runtime[row, column] != 0]
    return {
        "source_doi": description["reference"]["doi"],
        "source_location": description["reference"]["location"],
        "source_mapping": mapping,
        "source_separation": description["source_separation"],
        "runtime_separation": description["runtime_separation"],
        "source_csv_sha256": source_hash,
        "runtime_sha256": hashlib.sha256(runtime_path.read_bytes()).hexdigest(),
        "published_cells": len(expected),
        "published_cells_matching": len(expected) - len(differences),
        "published_cell_differences": differences,
        "model_zero_extension_cells": len(extensions),
        "model_zero_extension_errors": extension_errors,
        "verified": not differences and not extension_errors,
    }


def verify(resources_directory=RESOURCES_DIRECTORY, source_directory=SOURCE_DIRECTORY):
    metadata = json.loads((Path(source_directory) / "metadata.json").read_text(encoding="utf-8"))
    matrices = {name: verify_matrix(name, description, resources_directory, source_directory)
                for name, description in metadata["matrices"].items()}
    return {"comparison_policy": metadata["comparison_policy"], "matrices": matrices,
            "verified": all(result["verified"] for result in matrices.values())}


def main():
    parser = argparse.ArgumentParser(description=__doc__)
    parser.add_argument("--resources-dir", type=Path, default=RESOURCES_DIRECTORY)
    parser.add_argument("--output", type=Path, help="Save the complete JSON comparison")
    args = parser.parse_args()
    result = verify(args.resources_dir)
    serialized = json.dumps(result, indent=2, ensure_ascii=False) + "\n"
    if args.output:
        args.output.write_text(serialized, encoding="utf-8")
    print(serialized, end="")
    return 0 if result["verified"] else 1


if __name__ == "__main__":
    raise SystemExit(main())
```
