## Additional file 1. Evaluated software source and dependencies for "Helix Design Studio: designing antimicrobial peptide candidates for membrane environments": 3Dmol-bundled-dependencies.txt

Third-party libraries bundled inside app/static/3Dmol-min.js (3Dmol.js 2.4.2).
Runtime dependency notices are reproduced from the published npm packages
(pako 2.1.0, upng-js 2.1.0, iobuffer 5.3.2, netcdfjs 3.0.0).
Notices for embedded MMTF/MessagePack and Mol*/CIFTools.js code follow them.

================================================================================
pako
================================================================================

(The MIT License)

Copyright (C) 2014-2017 by Vitaly Puzrin and Andrei Tuputcyn

Permission is hereby granted, free of charge, to any person obtaining a copy
of this software and associated documentation files (the "Software"), to deal
in the Software without restriction, including without limitation the rights
to use, copy, modify, merge, publish, distribute, sublicense, and/or sell
copies of the Software, and to permit persons to whom the Software is
furnished to do so, subject to the following conditions:

The above copyright notice and this permission notice shall be included in
all copies or substantial portions of the Software.

THE SOFTWARE IS PROVIDED "AS IS", WITHOUT WARRANTY OF ANY KIND, EXPRESS OR
IMPLIED, INCLUDING BUT NOT LIMITED TO THE WARRANTIES OF MERCHANTABILITY,
FITNESS FOR A PARTICULAR PURPOSE AND NONINFRINGEMENT. IN NO EVENT SHALL THE
AUTHORS OR COPYRIGHT HOLDERS BE LIABLE FOR ANY CLAIM, DAMAGES OR OTHER
LIABILITY, WHETHER IN AN ACTION OF CONTRACT, TORT OR OTHERWISE, ARISING FROM,
OUT OF OR IN CONNECTION WITH THE SOFTWARE OR THE USE OR OTHER DEALINGS IN
THE SOFTWARE.

Portions of pako (lib/zlib) are distributed under the zlib license:

(C) 1995-2013 Jean-loup Gailly and Mark Adler
(C) 2014-2017 Vitaly Puzrin and Andrey Tupitsin

This software is provided 'as-is', without any express or implied
warranty. In no event will the authors be held liable for any damages
arising from the use of this software.

Permission is granted to anyone to use this software for any purpose,
including commercial applications, and to alter it and redistribute it
freely, subject to the following restrictions:

1. The origin of this software must not be misrepresented; you must not
  claim that you wrote the original software. If you use this software
  in a product, an acknowledgment in the product documentation would be
  appreciated but is not required.
2. Altered source versions must be plainly marked as such, and must not be
  misrepresented as being the original software.
3. This notice may not be removed or altered from any source distribution.

================================================================================
upng-js
================================================================================

MIT License

Copyright (c) 2017 Photopea

Permission is hereby granted, free of charge, to any person obtaining a copy
of this software and associated documentation files (the "Software"), to deal
in the Software without restriction, including without limitation the rights
to use, copy, modify, merge, publish, distribute, sublicense, and/or sell
copies of the Software, and to permit persons to whom the Software is
furnished to do so, subject to the following conditions:

The above copyright notice and this permission notice shall be included in all
copies or substantial portions of the Software.

THE SOFTWARE IS PROVIDED "AS IS", WITHOUT WARRANTY OF ANY KIND, EXPRESS OR
IMPLIED, INCLUDING BUT NOT LIMITED TO THE WARRANTIES OF MERCHANTABILITY,
FITNESS FOR A PARTICULAR PURPOSE AND NONINFRINGEMENT. IN NO EVENT SHALL THE
AUTHORS OR COPYRIGHT HOLDERS BE LIABLE FOR ANY CLAIM, DAMAGES OR OTHER
LIABILITY, WHETHER IN AN ACTION OF CONTRACT, TORT OR OTHERWISE, ARISING FROM,
OUT OF OR IN CONNECTION WITH THE SOFTWARE OR THE USE OR OTHER DEALINGS IN THE
SOFTWARE.

================================================================================
iobuffer
================================================================================

The MIT License (MIT)

Copyright (c) 2015 Michaël Zasso

Permission is hereby granted, free of charge, to any person obtaining a copy
of this software and associated documentation files (the "Software"), to deal
in the Software without restriction, including without limitation the rights
to use, copy, modify, merge, publish, distribute, sublicense, and/or sell
copies of the Software, and to permit persons to whom the Software is
furnished to do so, subject to the following conditions:

The above copyright notice and this permission notice shall be included in all
copies or substantial portions of the Software.

THE SOFTWARE IS PROVIDED "AS IS", WITHOUT WARRANTY OF ANY KIND, EXPRESS OR
IMPLIED, INCLUDING BUT NOT LIMITED TO THE WARRANTIES OF MERCHANTABILITY,
FITNESS FOR A PARTICULAR PURPOSE AND NONINFRINGEMENT. IN NO EVENT SHALL THE
AUTHORS OR COPYRIGHT HOLDERS BE LIABLE FOR ANY CLAIM, DAMAGES OR OTHER
LIABILITY, WHETHER IN AN ACTION OF CONTRACT, TORT OR OTHERWISE, ARISING FROM,
OUT OF OR IN CONNECTION WITH THE SOFTWARE OR THE USE OR OTHER DEALINGS IN THE
SOFTWARE.


================================================================================
netcdfjs
================================================================================

The MIT License (MIT)

Copyright (c) 2016 cheminfo

Permission is hereby granted, free of charge, to any person obtaining a copy
of this software and associated documentation files (the "Software"), to deal
in the Software without restriction, including without limitation the rights
to use, copy, modify, merge, publish, distribute, sublicense, and/or sell
copies of the Software, and to permit persons to whom the Software is
furnished to do so, subject to the following conditions:

The above copyright notice and this permission notice shall be included in all
copies or substantial portions of the Software.

THE SOFTWARE IS PROVIDED "AS IS", WITHOUT WARRANTY OF ANY KIND, EXPRESS OR
IMPLIED, INCLUDING BUT NOT LIMITED TO THE WARRANTIES OF MERCHANTABILITY,
FITNESS FOR A PARTICULAR PURPOSE AND NONINFRINGEMENT. IN NO EVENT SHALL THE
AUTHORS OR COPYRIGHT HOLDERS BE LIABLE FOR ANY CLAIM, DAMAGES OR OTHER
LIABILITY, WHETHER IN AN ACTION OF CONTRACT, TORT OR OTHERWISE, ARISING FROM,
OUT OF OR IN CONNECTION WITH THE SOFTWARE OR THE USE OR OTHER DEALINGS IN THE
SOFTWARE.

================================================================================
MMTF v1.0.1, including its MessagePack codec
================================================================================

Bundled source: https://github.com/3dmol/3Dmol.js/blob/2.4.2/src/vendor/mmtf.js
This vendor file is byte-identical to the MMTF v1.0.1 distribution:
https://github.com/rcsb/mmtf-javascript/blob/v1.0.1/dist/mmtf.js
SHA-256 (both files): fee8488f97c36f9bb2e3e750cfde394886eb361c7ea5b7a496455b2353721b39
License source: https://github.com/rcsb/mmtf-javascript/blob/v1.0.1/LICENSE

The MIT licensed applies to all files unless otherwise noted.

//

The MIT License

Copyright (c) 2015-2016, Alexander S. Rose <>

Permission is hereby granted, free of charge, to any person obtaining a copy
of this software and associated documentation files (the "Software"), to deal
in the Software without restriction, including without limitation the rights
to use, copy, modify, merge, publish, distribute, sublicense, and/or sell
copies of the Software, and to permit persons to whom the Software is
furnished to do so, subject to the following conditions:

The above copyright notice and this permission notice shall be included in
all copies or substantial portions of the Software.

THE SOFTWARE IS PROVIDED "AS IS", WITHOUT WARRANTY OF ANY KIND, EXPRESS OR
IMPLIED, INCLUDING BUT NOT LIMITED TO THE WARRANTIES OF MERCHANTABILITY,
FITNESS FOR A PARTICULAR PURPOSE AND NONINFRINGEMENT. IN NO EVENT SHALL THE
AUTHORS OR COPYRIGHT HOLDERS BE LIABLE FOR ANY CLAIM, DAMAGES OR OTHER
LIABILITY, WHETHER IN AN ACTION OF CONTRACT, TORT OR OTHERWISE, ARISING FROM,
OUT OF OR IN CONNECTION WITH THE SOFTWARE OR THE USE OR OTHER DEALINGS IN
THE SOFTWARE.

================================================================================
MessagePack codec contributions by Tim Caswell
================================================================================

The MMTF v1.0.1 source retains the following attribution for its encoder and decoder:
Copyright (c) 2013 Tim Caswell <>, MIT License.
https://github.com/rcsb/mmtf-javascript/blob/v1.0.1/src/msgpack-encode.js
https://github.com/rcsb/mmtf-javascript/blob/v1.0.1/src/msgpack-decode.js
License source:
https://github.com/creationix/msgpack-js/blob/52ca83d0903472591fe6c1991dcaa4133f68f06e/LICENSE

The MIT License (MIT)

Copyright (c) 2013 Tim Caswell <>

Permission is hereby granted, free of charge, to any person obtaining a copy
of this software and associated documentation files (the "Software"), to deal
in the Software without restriction, including without limitation the rights
to use, copy, modify, merge, publish, distribute, sublicense, and/or sell
copies of the Software, and to permit persons to whom the Software is
furnished to do so, subject to the following conditions:

The above copyright notice and this permission notice shall be included in
all copies or substantial portions of the Software.

THE SOFTWARE IS PROVIDED "AS IS", WITHOUT WARRANTY OF ANY KIND, EXPRESS OR
IMPLIED, INCLUDING BUT NOT LIMITED TO THE WARRANTIES OF MERCHANTABILITY,
FITNESS FOR A PARTICULAR PURPOSE AND NONINFRINGEMENT. IN NO EVENT SHALL THE
AUTHORS OR COPYRIGHT HOLDERS BE LIABLE FOR ANY CLAIM, DAMAGES OR OTHER
LIABILITY, WHETHER IN AN ACTION OF CONTRACT, TORT OR OTHERWISE, ARISING FROM,
OUT OF OR IN CONNECTION WITH THE SOFTWARE OR THE USE OR OTHER DEALINGS IN
THE SOFTWARE.

================================================================================
Mol* / CIFTools.js BinaryCIF decoding code
================================================================================

The 3Dmol.js 2.4.2 sources retain these copyright notices:
Copyright (c) 2017-2024 mol* contributors, licensed under MIT.
Copyright (c) 2017 mol* contributors, licensed under MIT.
From CIFTools.js; author David Sehnal <>.
Fixed point, delta, RLE and integer packing adaptations credit Alexander Rose,
MIT License, Copyright (c) 2016; the MMTF MIT license is reproduced above.
Sources:
https://github.com/3dmol/3Dmol.js/blob/2.4.2/src/parsers/cifutils/decoder.ts
https://github.com/3dmol/3Dmol.js/blob/2.4.2/src/parsers/cifutils/encoding.ts
https://github.com/3dmol/3Dmol.js/blob/2.4.2/src/parsers/cifutils/category.ts
The MIT terms below are reproduced from the Mol* license:
https://github.com/molstar/molstar/blob/v4.8.0/LICENSE

The MIT License

    Copyright (c) 2017 - now, Mol* contributors

Permission is hereby granted, free of charge, to any person obtaining a copy
of this software and associated documentation files (the "Software"), to deal
in the Software without restriction, including without limitation the rights
to use, copy, modify, merge, publish, distribute, sublicense, and/or sell
copies of the Software, and to permit persons to whom the Software is
furnished to do so, subject to the following conditions:

The above copyright notice and this permission notice shall be included in
all copies or substantial portions of the Software.

THE SOFTWARE IS PROVIDED "AS IS", WITHOUT WARRANTY OF ANY KIND, EXPRESS OR
IMPLIED, INCLUDING BUT NOT LIMITED TO THE WARRANTIES OF MERCHANTABILITY,
FITNESS FOR A PARTICULAR PURPOSE AND NONINFRINGEMENT. IN NO EVENT SHALL THE
AUTHORS OR COPYRIGHT HOLDERS BE LIABLE FOR ANY CLAIM, DAMAGES OR OTHER
LIABILITY, WHETHER IN AN ACTION OF CONTRACT, TORT OR OTHERWISE, ARISING FROM,
OUT OF OR IN CONNECTION WITH THE SOFTWARE OR THE USE OR OTHER DEALINGS IN
THE SOFTWARE.
