## Additional file 1. Evaluated software source and dependencies for "Helix Design Studio: designing antimicrobial peptide candidates for membrane environments": README.txt

Supplemental third-party notices for the evaluated source snapshot.

The helix-design-studio-web/ directory preserves the exact tracked files of d6ccec1920f256a393b15215df63fd2dead61cd5. The supplemental license text applies to the same unchanged 3Dmol.js 2.4.2 bundle. No runtime source, numerical parameter or result is modified.
