## Supplementary figures and images for "Helix Design Studio: designing antimicrobial peptide candidates for membrane environments"

### figure_1_concept.png

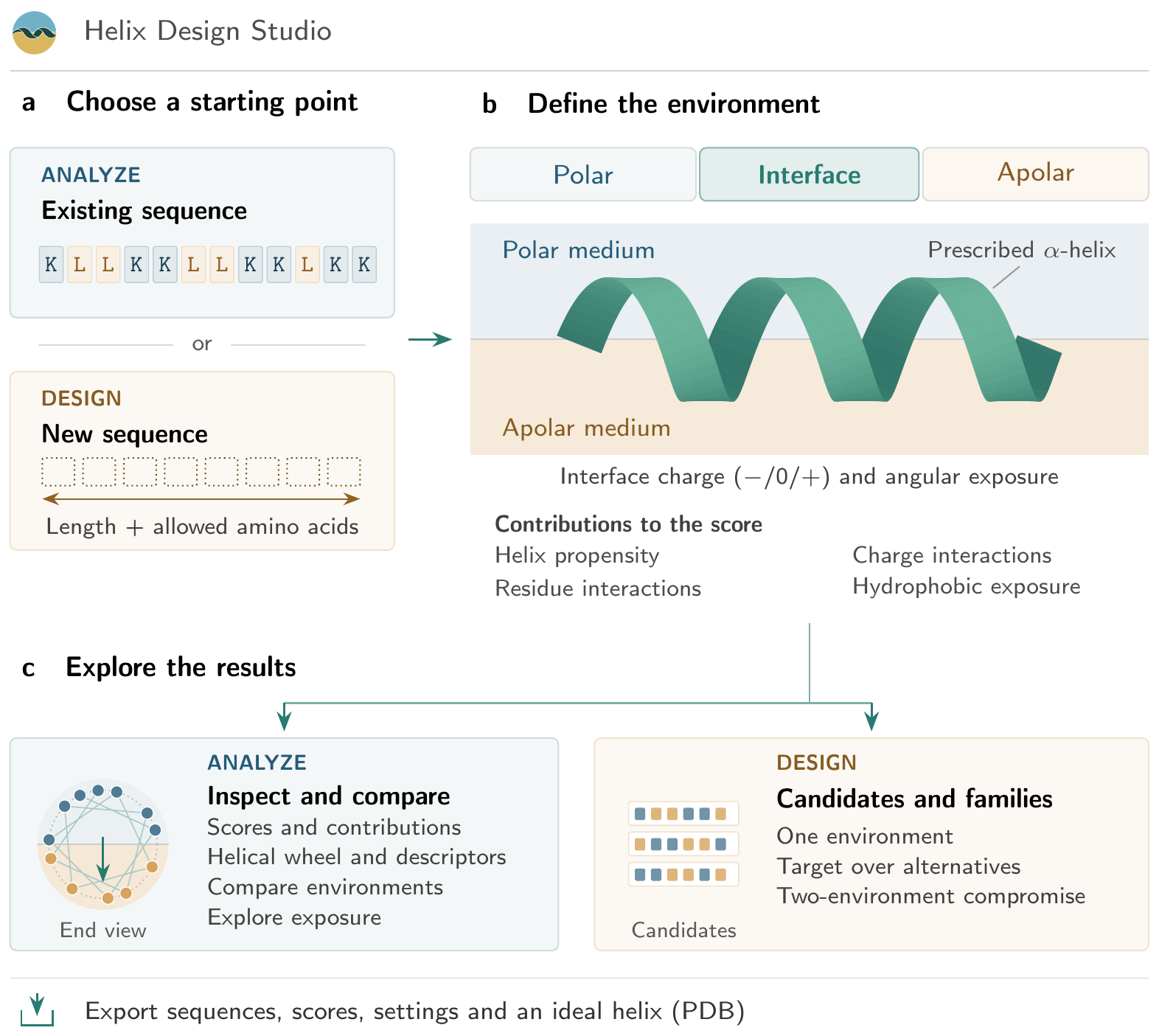

### figure_2_architecture.pdf

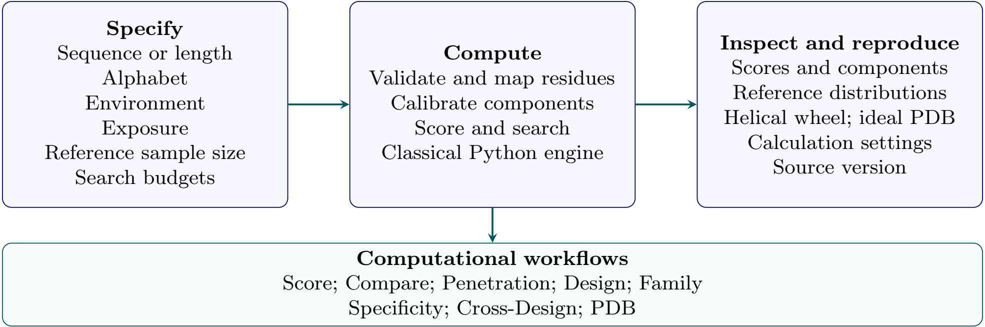

### figure_2_architecture.png

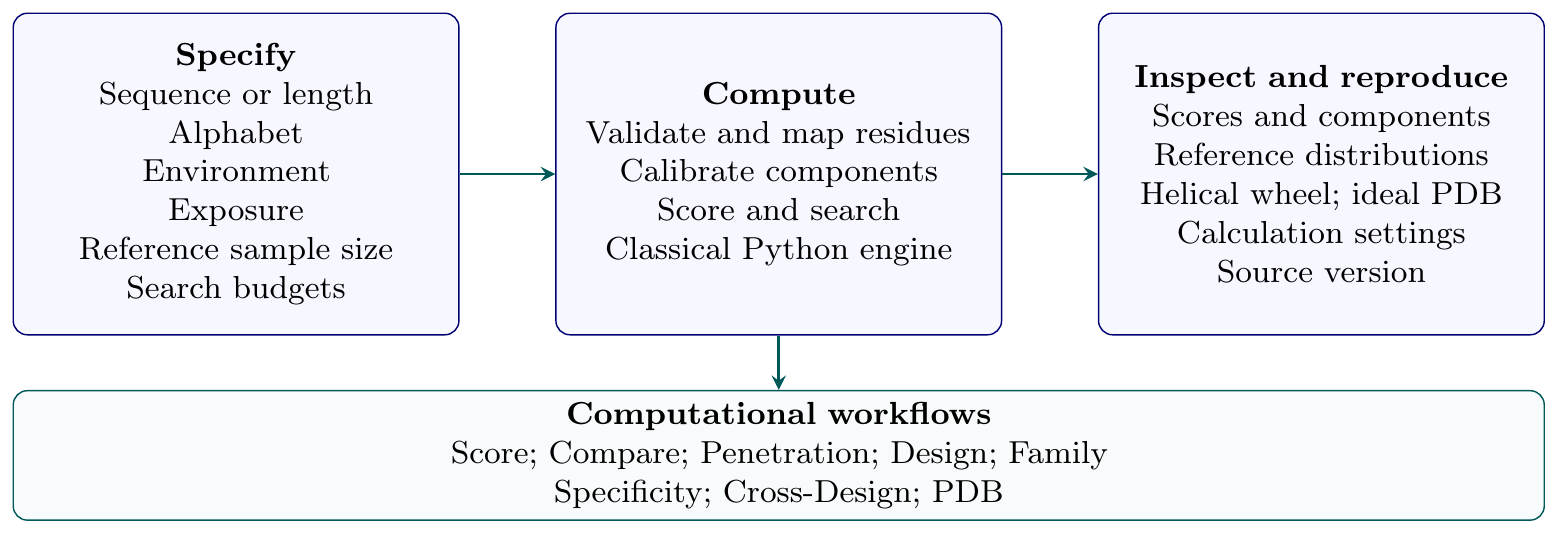

### figure_3_profiles.png

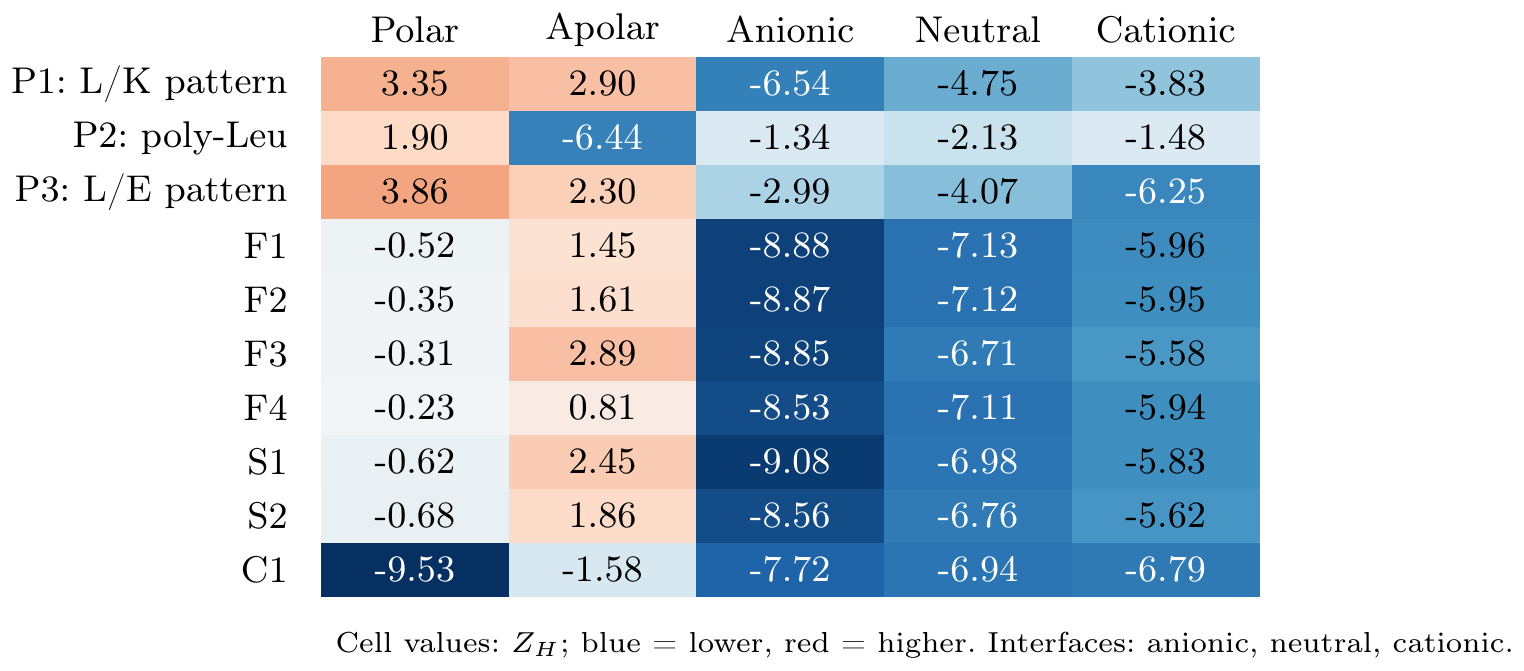

### figure_4_collections.pdf

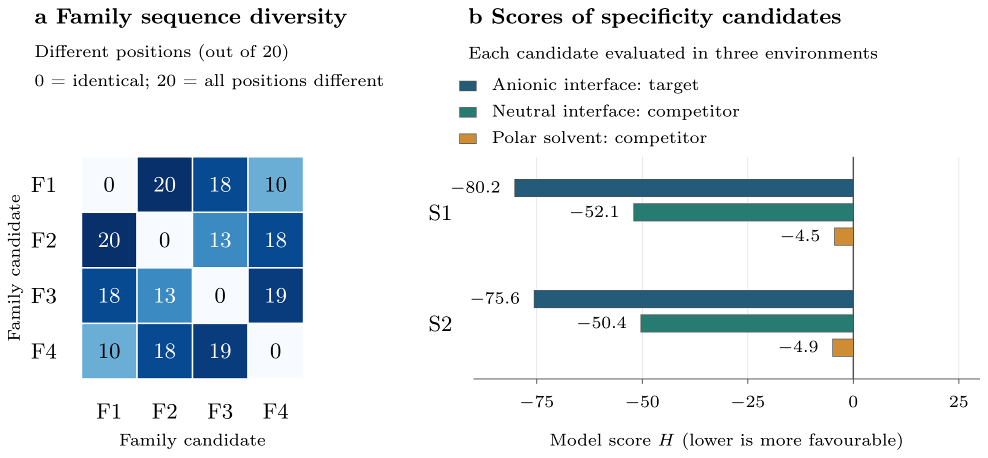

### figure_4_collections.png

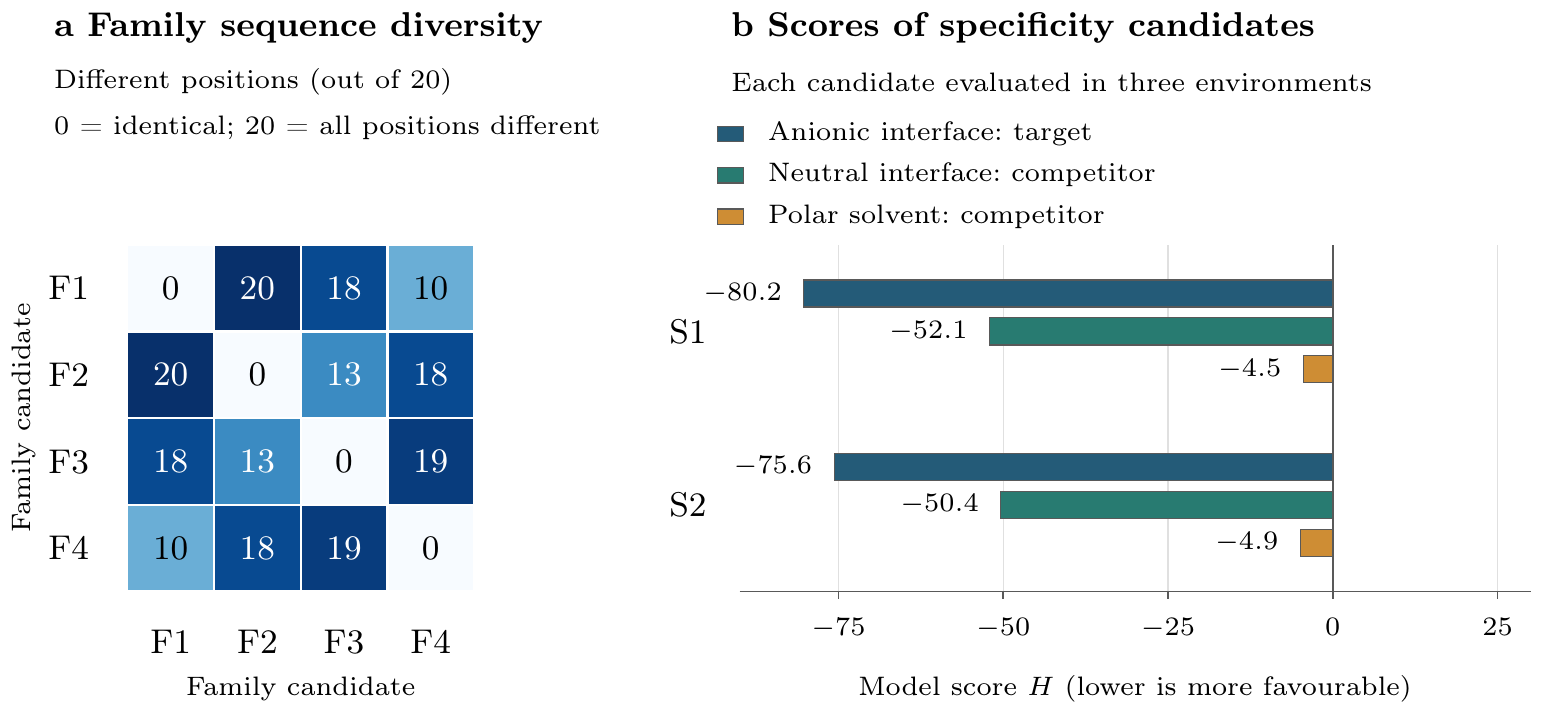

### figure_5_exposure.pdf

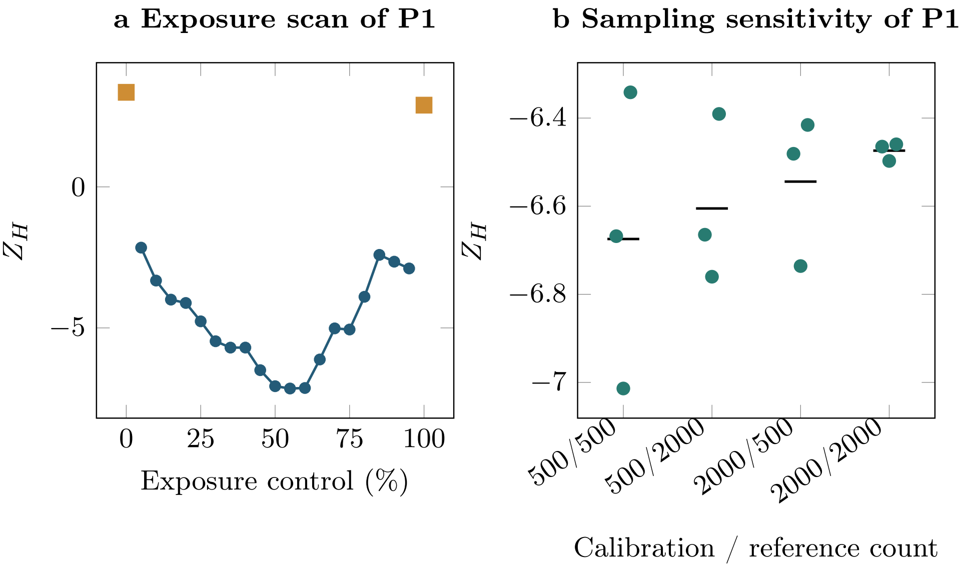

### figure_5_exposure.png

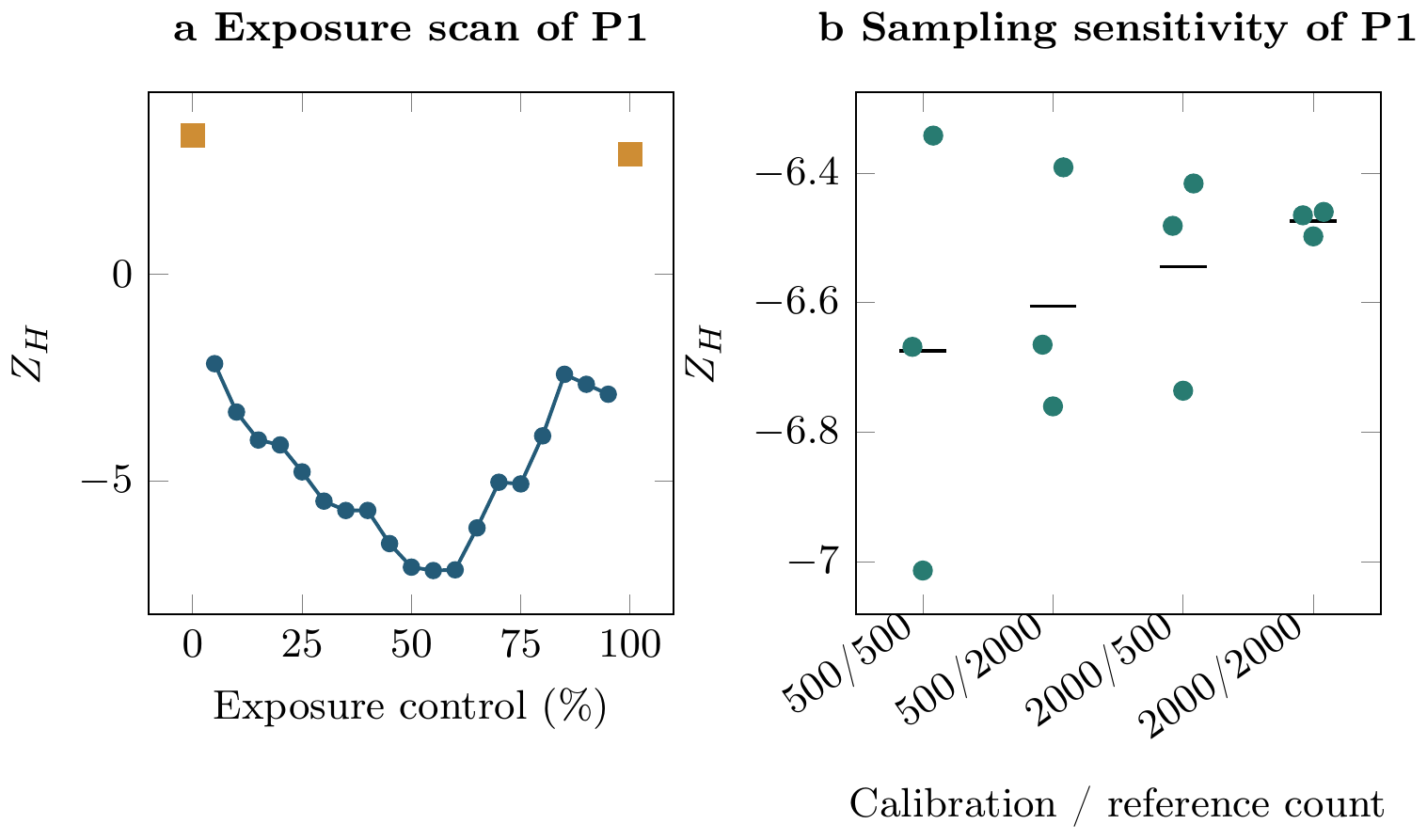

### figure_6_timings.pdf

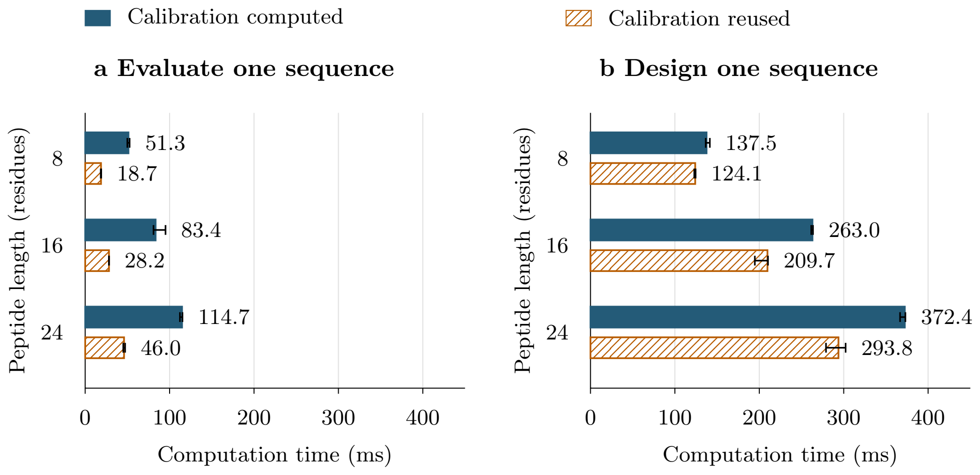

### figure_6_timings.png

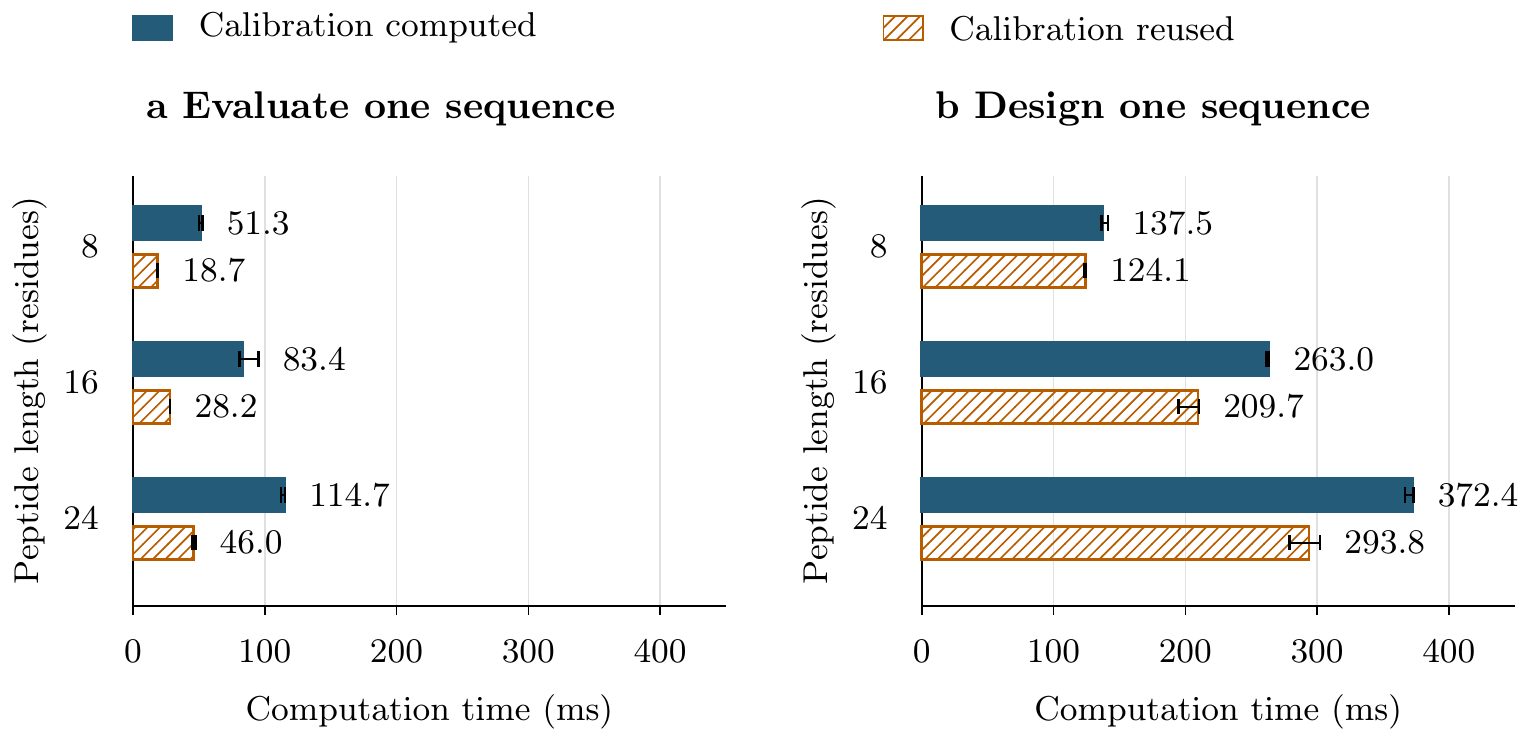

### figure_7_alignment.pdf

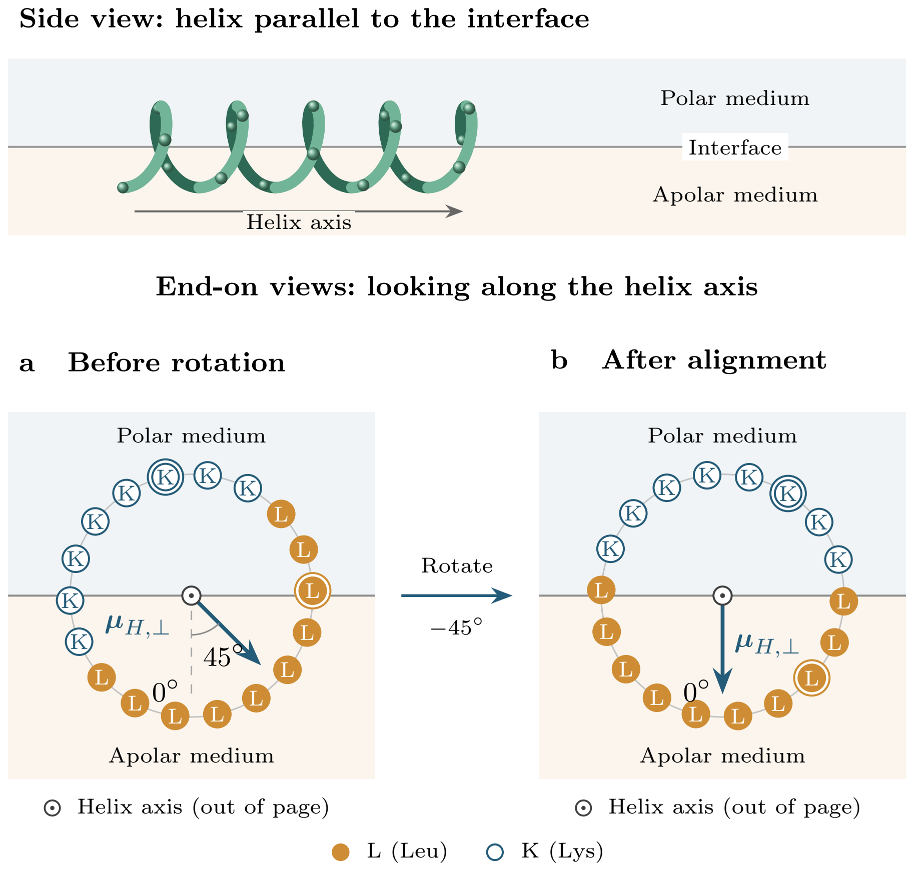

### figure_7_alignment.png

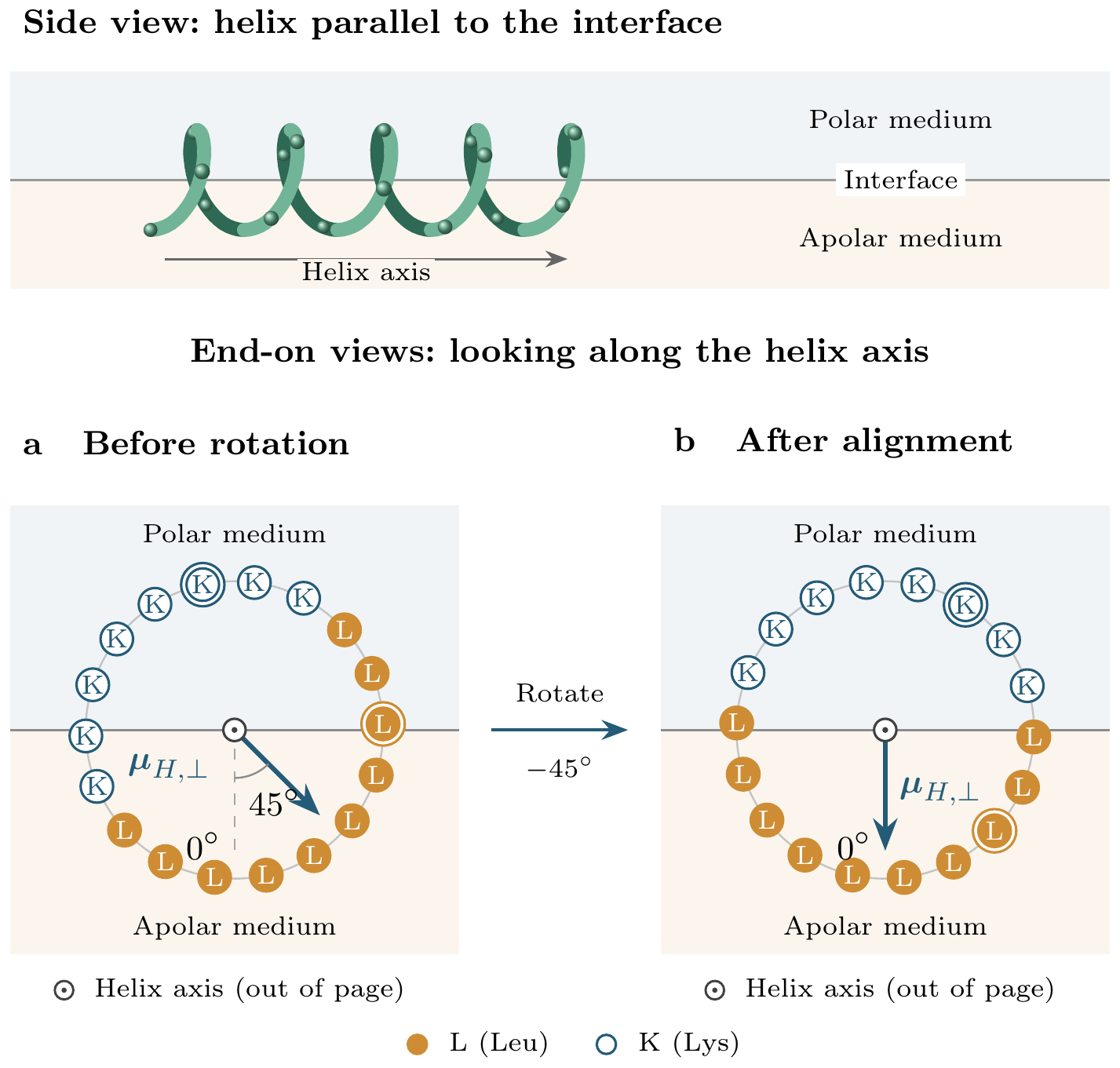

### helix-environment-mark.png

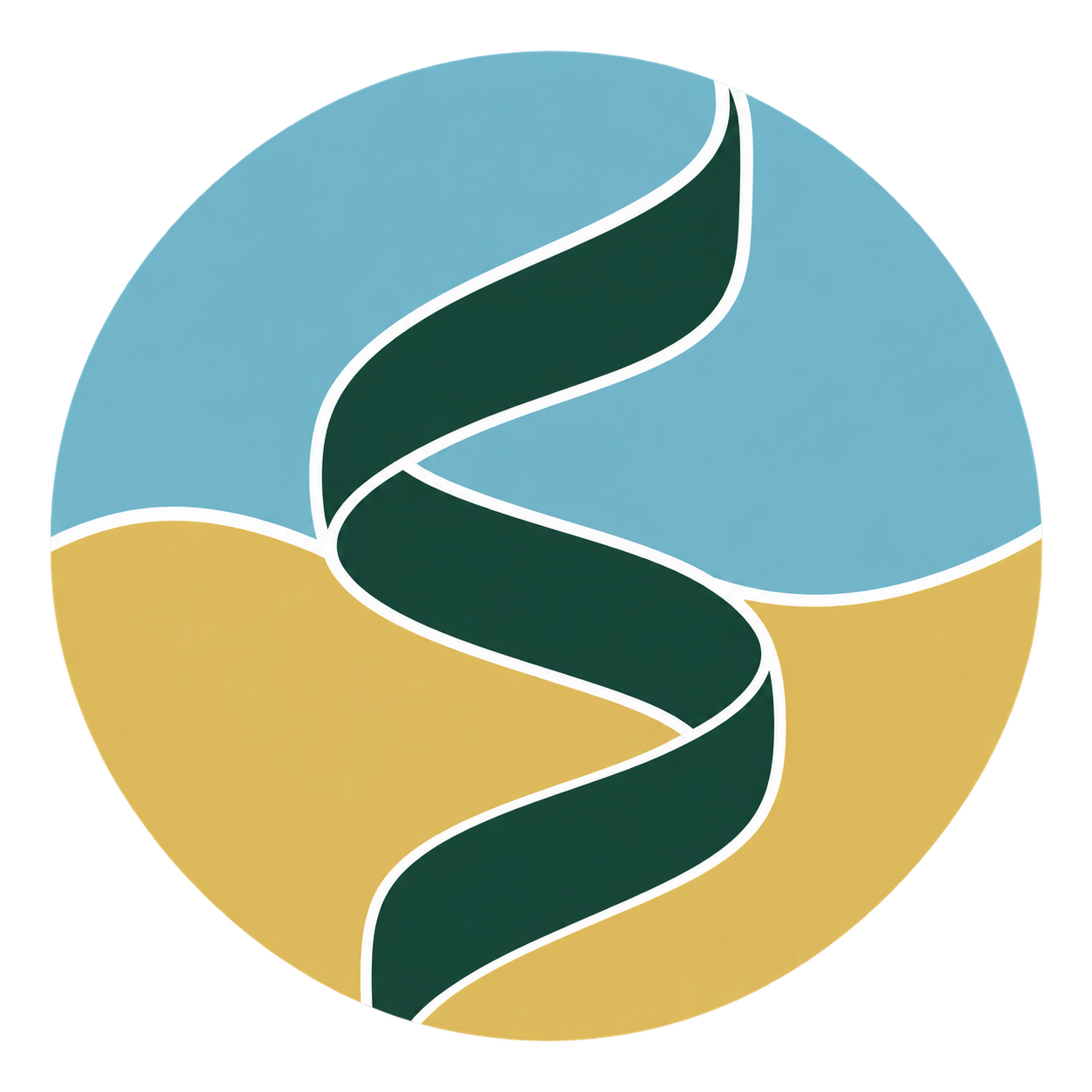

### helix-membrane-horizontal.png

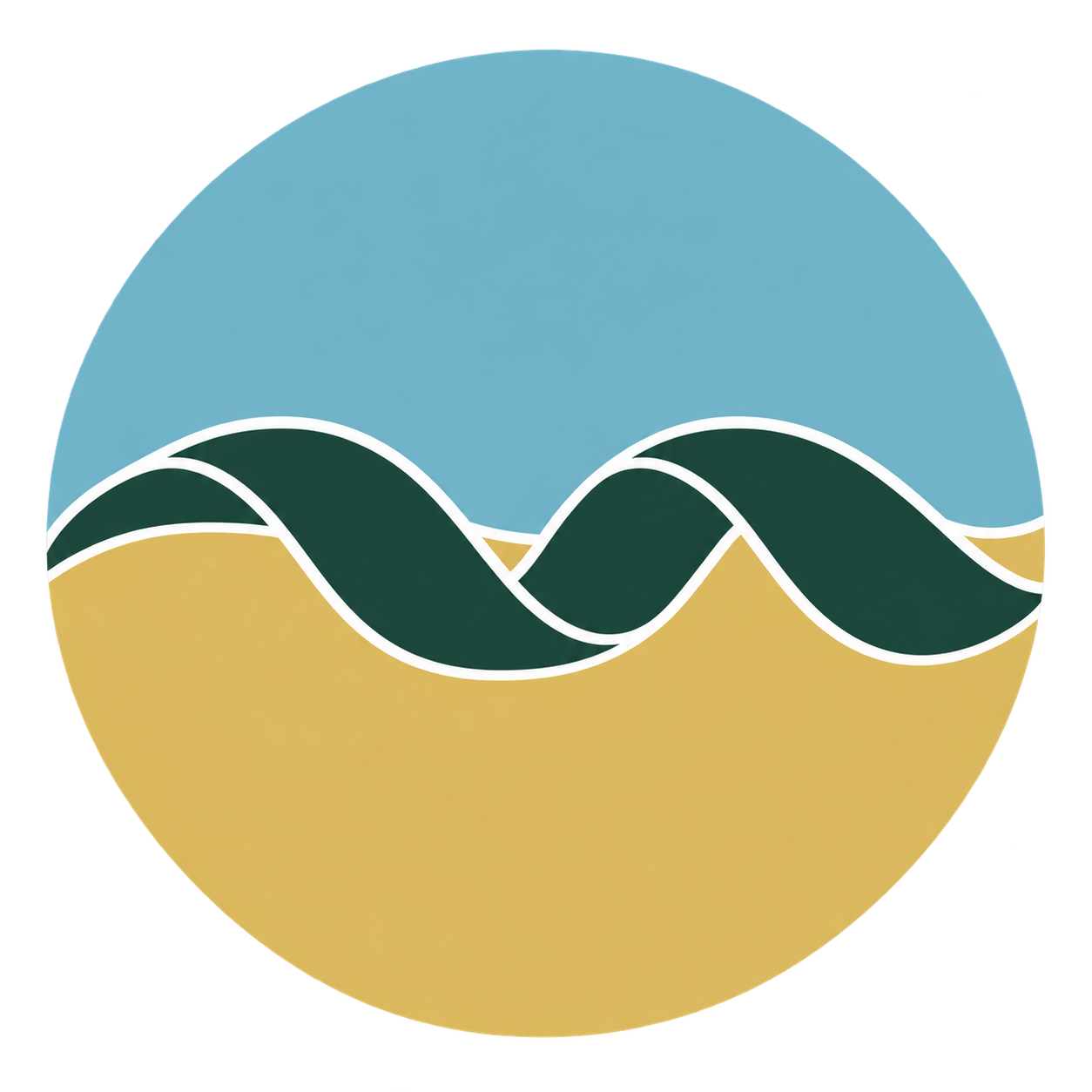

### logo_USC.gif

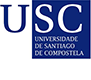

### simbios-logo.png

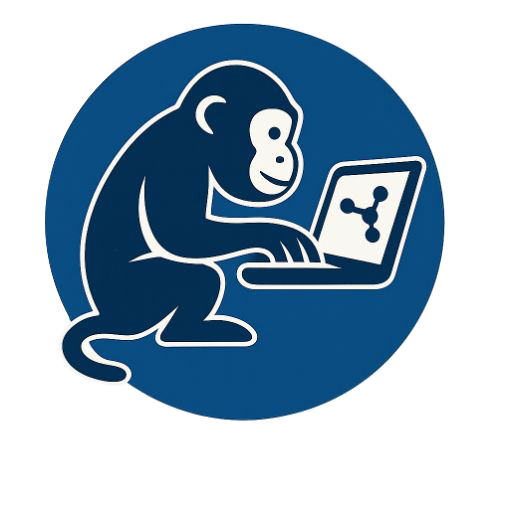
