## Additional file 2. Computational evaluation, inputs and results for "Helix Design Studio: designing antimicrobial peptide candidates for membrane environments": audit_scoring_direction.py

```
"""Bounded audit of score directions, formulas, calibration and minimization.

Uses the real parameter loaders and raw methods without constructing caches.
The existing exact-search result is inspected, not recomputed. No runtime files
are changed. Run with the application dependencies installed.
"""
from __future__ import annotations

import argparse
import ast
from datetime import datetime, timezone
import hashlib
import inspect
import json
from pathlib import Path
import subprocess
import sys

import numpy as np


def sha(path):
    return hashlib.sha256(path.read_bytes()).hexdigest()


def main():
    parser = argparse.ArgumentParser(description=__doc__)
    base = Path(__file__).resolve().parents[1]
    parser.add_argument('--repo', type=Path, default=base.parent / 'helix-design-studio-web')
    parser.add_argument('--results', type=Path, default=base / 'results')
    args = parser.parse_args()
    repo, results = args.repo.resolve(), args.results.resolve()
    sys.path.insert(0, str(repo))
    sys.path.insert(0, str(repo / 'Code'))
    from core import hamiltonian_builder as model
    from app import engine

    seed = 0
    alphabet = list('ARNDCQEGHILKMFPSTWYV')

    def bare(sequence, halfwidth=-1, charge='neu'):
        b = model.HamiltonianBuilder.__new__(model.HamiltonianBuilder)
        b.L, b.amino_acids, b.n_aa = len(sequence), alphabet, len(alphabet)
        b.kwargs = {'wheel_halfwidth_deg': halfwidth, 'wheel_phase_deg': 0,
                    'membrane_charge': charge, 'zscore_seed': seed,
                    'max_interaction_dist': 4, 'align_phase': False}
        b.align_phase, b.stats_cache_version = False, 5
        b._init_properties()
        b._load_matrices_from_files()
        return b, [alphabet.index(a) for a in sequence]

    def raw(method, sequence, halfwidth=-1, charge='neu'):
        b, codes = bare(sequence, halfwidth, charge)
        return float(getattr(b, method)(codes))

    probes = {}
    probes['exposure'] = {f'{sequence}_{medium}': raw('_raw_env_pol', sequence, width)
                          for sequence in ('LLLL', 'KKKK')
                          for medium, width in [('polar', -1), ('apolar', 180)]}
    e = probes['exposure']
    assert e['LLLL_apolar'] < e['KKKK_apolar'] and e['KKKK_polar'] < e['LLLL_polar']
    probes['interface_charge'] = {f'{sequence}_{charge}': raw('_raw_env_chg', sequence, -1, charge)
                                  for sequence in ('KKKK', 'DDDD', 'AAAA')
                                  for charge in ('neg', 'neu', 'pos')}
    c = probes['interface_charge']
    assert c['KKKK_neg'] == c['DDDD_pos'] == -1
    assert c['DDDD_neg'] == c['KKKK_pos'] == c['AAAA_neg'] == c['AAAA_pos'] == 0
    assert raw('_raw_env_chg', 'KKKK', 180, 'neg') == 0
    probes['local_helix'] = {s: raw('_raw_helix_local', s) for s in ('AA', 'GG', 'PP')}
    assert probes['local_helix']['AA'] < probes['local_helix']['GG'] < probes['local_helix']['PP']
    probes['mj_contacts'] = {s: raw('_raw_pw_int', s) for s in ('AA', 'WW')}
    assert probes['mj_contacts']['WW'] < probes['mj_contacts']['AA'] < 0
    probes['neighbors'] = {s: raw('_raw_helix_neigh', s) for s in ('AA', 'GG', 'AP')}
    assert np.isclose(probes['neighbors']['AA'], -2.734)
    assert probes['neighbors']['AA'] < probes['neighbors']['GG'] < probes['neighbors']['AP'] == 0
    b, codes = bare('IAAF'); b.M1.fill(0)
    probes['neighbors']['M3_isolated_IAAF'] = float(b._raw_helix_neigh(codes))
    b, codes = bare('WAAAH'); b.M1.fill(0); b.M3.fill(0)
    probes['neighbors']['M4_isolated_WAAAH'] = float(b._raw_helix_neigh(codes))
    assert np.isclose(probes['neighbors']['M3_isolated_IAAF'], -.45)
    assert np.isclose(probes['neighbors']['M4_isolated_WAAAH'], -.9)
    probes['electrostatics'] = {f'{s}_{medium}': raw('_raw_electrostatic', s, width)
                                for s in ('KK', 'KD')
                                for medium, width in [('polar', -1), ('mixed', 90), ('apolar', 180)]}
    for medium in ('polar', 'mixed', 'apolar'):
        assert probes['electrostatics'][f'KD_{medium}'] < 0 < probes['electrostatics'][f'KK_{medium}']
    assert raw('_raw_electrostatic', 'KAAAAAAAK') > 0
    assert raw('_raw_electrostatic', 'KAAAAAAAAK') == 0

    b, _ = bare('AKLDGWER')
    stats = b._calibrate_z_scores(100)
    rng = np.random.default_rng(seed)
    independent = []
    for _ in range(100):
        codes = rng.integers(0, len(alphabet), size=b.L).tolist()
        means = [float(np.mean([m[codes[i], codes[i+k]] for i in range(b.L-k)]))
                 for k, m in ((1, b.M1), (3, b.M3), (4, b.M4))]
        independent.append(-means[0] + means[1] + means[2])
    expected = float(np.mean(independent)), float(np.std(independent))
    assert np.allclose(stats['helix_neigh'], expected, atol=1e-14, rtol=0)
    assert all(w > 0 for w in engine.DEFAULT_WEIGHTS.values())

    source = (repo / 'app/engine.py').read_text()
    functions = {n.name: n for n in ast.parse(source).body if isinstance(n, ast.FunctionDef)}
    optimizer_checks = {}
    for name in ('simulated_annealing', '_iter_specificity_search', 'cross_environment_design'):
        fn = functions[name]
        deltas = [n for n in ast.walk(fn) if isinstance(n, ast.Assign)
                  and any(isinstance(t, ast.Name) and t.id == 'delta' for t in n.targets)]
        assert len(deltas) == 1
        delta = ast.unparse(deltas[0].value)
        expected_delta = 'prop_obj - current_obj' if name == 'cross_environment_design' else 'proposal_objective - current_objective'
        assert delta == expected_delta
        accepts_downhill = any(isinstance(n, ast.Compare) and isinstance(n.left, ast.Name)
                               and n.left.id == 'delta' and isinstance(n.ops[0], ast.LtE)
                               and isinstance(n.comparators[0], ast.Constant) and n.comparators[0].value == 0
                               for n in ast.walk(fn))
        assert accepts_downhill
        function_source = ast.get_source_segment(source, fn)
        optimizer_checks[name] = {'status': 'passed', 'delta': delta, 'accepts_downhill': True,
                                 'source_sha256': hashlib.sha256(function_source.encode()).hexdigest()}
    family = inspect.getsource(engine.design_family_stream)
    assert 'candidate = simulated_annealing(' in family
    assert 'ranked = sorted(unique_results.values(), key=lambda item: item["design_geometry"]["energy"])' in family
    optimizer_checks['family'] = {'status': 'passed', 'search': 'simulated_annealing plus nonnegative diversity penalty',
                                  'ranking': 'ascending reported model score'}
    exact_path = results / 'exact_search_summary.json'
    exact = json.loads(exact_path.read_text())
    assert exact['candidate_count'] == 4096
    trials = exact['trials']
    assert len(trials) == 5 and all(t['recovered_exact_minimum'] for t in trials)
    commit = subprocess.check_output(['git', '-C', str(repo), 'rev-parse', 'HEAD'], text=True).strip()
    dirty = subprocess.check_output(['git', '-C', str(repo), 'status', '--porcelain'], text=True).strip()
    assert not dirty, 'Runtime must be an immutable clean revision.'
    study_metadata = json.loads((results/'metadata.json').read_text())
    study_source = study_metadata.get('final_source', study_metadata.get('source', {}))
    assert study_source['commit'] == commit, 'Existing exact-search evidence must match the audited revision.'
    main_ast = ast.parse((repo/'app/main.py').read_text())
    version = next(ast.literal_eval(n.value) for n in main_ast.body if isinstance(n, ast.Assign)
                   and any(isinstance(t, ast.Name) and t.id == 'APP_VERSION' for t in n.targets))
    result = {
        'title': 'Directional audit of all six raw score components and minimization conventions',
        'status': 'passed', 'verified_utc': datetime.now(timezone.utc).isoformat(),
        'version': version, 'source_commit': commit, 'git_status': dirty,
        'parameters_sha256': model.parameter_set_sha256(), 'seed': seed,
        'method': 'Actual raw methods and parameter loaders; bare objects bypass caches. One 100-sequence calibration; no new optimization searches.',
        'formulas': {
            'exposure': 'mean((1-2*b_i)*FP(s_i)); b_i=1 in apolar sector',
            'interface_charge': 'sum_polar(min(surface_charge*q_i,0))/L',
            'local_helix': 'mean(Pace-Scholtz penalty(s_i))',
            'mj_contacts': 'sum_{i<j,j-i<=4} MJ(s_i,s_j)/(4*L)',
            'neighbors': '-mean(M1_i,i+1)+mean(M3_i,i+3)+mean(M4_i,i+4); omit unavailable separations',
            'electrostatics': '2/[L*(L-1)] * sum_{i<j,j-i<=8} q_i*q_j/[(1+j-i)*epsilon_ij]',
            'combined_score': 'sum_a positive_weight_a*(raw_a-mean_a)/positive_population_std_a',
        },
        'small_raw_probes': probes,
        'joint_neighbor_calibration': {'Ncal': 100, 'seed': seed, 'length': 8, 'ordered_alphabet': alphabet,
                                       'actual_mu_sigma': list(stats['helix_neigh']), 'independent_mu_sigma': list(expected)},
        'optimizer_direction_checks': optimizer_checks,
        'existing_exact_search_evidence': {'file': exact_path.name, 'sha256': sha(exact_path),
                                           'source_commit': study_source['commit'],
                                           'candidate_count': exact['candidate_count'], 'exact_minimum': exact['exact_minimum'],
                                           'exact_minimizers': exact['exact_minimizers'], 'successful_trials': len(trials),
                                           'note': 'Inspected existing evaluation output; enumeration and searches were not rerun.'},
        'limitations': [
            'Directional checks establish implementation behavior, not biological validity or thermodynamic accuracy.',
            'M1 at separation1 and its linear combination with M3/M4 are empirical model choices.',
            'Interface coupling rewards opposite charges; same-sign and neutral residues both contribute zero raw value.',
            'Zero raw surface coupling can become positive after centering; this is not explicit repulsion.',
            'Electrostatics uses sequence separation, a cutoff8 and effective epsilon values .1/.3/1, not atomistic distances or salt-dependent Debye screening.',
            'Uniform positive rescaling cancels on standardization, including the homogeneous polar/apolar dielectric multiplier.',
            'Hydrophobic-moment alignment prescribes wheel phase; it does not minimize the total score with respect to angle.',
            'Finite exact-search evidence establishes optimality only within its enumerated sequence space.',
        ],
        'source_sha256': {str(p.relative_to(repo)): sha(p) for p in [repo/'Code/core/hamiltonian_builder.py',repo/'app/engine.py',repo/'app/main.py',*[repo/'Code/core/resources'/n for n in model.RESOURCE_FILES]]},
        'audit_script_sha256': sha(Path(__file__)),
    }
    output = results/'scoring_direction_audit.json'
    output.write_text(json.dumps(result, indent=2, ensure_ascii=False, allow_nan=False)+'\n')
    print(json.dumps({'status': result['status'], 'output': str(output), 'source_commit': commit,
                      'component_count': len(probes), 'optimizer_checks': len(optimizer_checks)}))


if __name__ == '__main__':
    main()
```
