## Additional file 2. Computational evaluation, inputs and results for "Helix Design Studio: designing antimicrobial peptide candidates for membrane environments": compare_descriptors.py

```
"""Actual modlAMP/HDS descriptor comparison; HELIQUEST is a documentary reference only."""
import collections
import datetime
import hashlib
import importlib.metadata
import inspect
import json
import math
import pathlib
import platform
import subprocess
import sys

root = pathlib.Path(sys.argv[1]).resolve()
output = pathlib.Path(sys.argv[2]).resolve()
sys.path.insert(0, str(root))
from app import engine
from modlamp.descriptors import PeptideDescriptor, GlobalDescriptor
import modlamp.descriptors

fp = {'A':0.31,'R':-1.01,'N':-0.60,'D':-0.77,'C':1.54,'Q':-0.22,'E':-0.64,'G':0.,'H':0.13,'I':1.80,'L':1.70,'K':-0.99,'M':1.23,'F':1.79,'P':0.72,'S':-0.04,'T':0.26,'W':2.25,'Y':0.96,'V':1.22}
probes = {'probe_amphiphilic':'LKKLLKLLKKLLKLLKKLLK','probe_hydrophobic':'LLLLLLLLLLLLLLLLLLLL','probe_acidic':'LEELLELLEELLELLEELLE'}
controls = {'control_ST_alternating':'STSTSTSTSTSTSTSTSTST','control_S_homopolymer':'S'*20,'control_T_homopolymer':'T'*20}
inputs = dict(probes, **controls)
seqs = list(inputs.values())

def make_descriptor():
    descriptor = PeptideDescriptor(seqs, 'eisenberg')
    descriptor.scale = {aa:[value] for aa,value in fp.items()}
    descriptor.scalename = 'Fauchere-Pliska supplied explicitly from HELIQUEST Help & Methods'
    return descriptor

means = make_descriptor()
means.calculate_global(window=20, modality='max')
moments = make_descriptor()
moments.calculate_moment(window=20, angle=100, modality='max')
charges = GlobalDescriptor(seqs)
charges.calculate_charge(ph=7.4, amide=False)

rows = []
for i,(identifier,seq) in enumerate(inputs.items()):
    n = len(seq)
    mu = engine.hydrophobic_moment_descriptor(seq)
    # Independently implement the equation documented by HELIQUEST; no server execution.
    cosine = sum(fp[aa]*math.cos(math.radians(j*100.0)) for j,aa in enumerate(seq))
    sine = sum(fp[aa]*math.sin(math.radians(j*100.0)) for j,aa in enumerate(seq))
    reference_mean = sum(fp[aa] for aa in seq)/n
    reference_moment = math.hypot(cosine,sine)/n
    reference_charge = seq.count('K')+seq.count('R')-seq.count('D')-seq.count('E')
    engine_mean = sum(engine.HYDROPHOBICITY_FP[aa] for aa in seq)/n
    package_mean = float(means.descriptor[i,0])
    package_moment = float(moments.descriptor[i,0])
    rows.append({
        'id':identifier,'role':'primary_probe' if identifier in probes else 'scale_regression_control',
        'sequence':seq,'length':n,'composition':dict(sorted(collections.Counter(seq).items())),
        'helix_design_studio_executed':{
            'hydrophobic_moment_descriptor_raw':mu,
            'mean_hydrophobicity_computed_from_engine_scale':engine_mean,
            'model_net_charge':engine.model_net_charge(seq)},
        'modlamp_executed':{'mean_hydrophobicity':package_mean,
            'hydrophobic_moment_per_residue':package_moment,
            'charge_ph_7_4_unamidated_package_rounds_to_3_decimals':float(charges.descriptor[i,0])},
        'heliquest_documented_formula_independent_local_calculation':{
            'mean_hydrophobicity':reference_mean,'hydrophobic_moment_per_residue':reference_moment,
            'integer_sidechain_charge':reference_charge},
        'absolute_difference_hds_vs_modlamp':{
            'mean_hydrophobicity':abs(engine_mean-package_mean),
            'hydrophobic_moment_per_residue':abs(mu['magnitude_per_residue']-package_moment)}
    })

source_file = pathlib.Path(__file__)
engine_file = pathlib.Path(engine.__file__)
package_file = pathlib.Path(modlamp.descriptors.__file__)

def git_value(*arguments):
    """Source archives without .git remain executable; file hashes identify their code."""
    try:
        process = subprocess.run(['git', *arguments], cwd=root, text=True,
                                 capture_output=True, check=False)
    except FileNotFoundError:
        return None
    return process.stdout.strip() if process.returncode == 0 else None

scale_mismatches = {aa:{'hds':engine.HYDROPHOBICITY_FP.get(aa),'documented_FP':value} for aa,value in fp.items() if engine.HYDROPHOBICITY_FP.get(aa)!=value}
result = {
    'title':'Actual modlAMP 4.3.2 versus Helix Design Studio descriptor comparison, with HELIQUEST documentary formula reference',
    'execution_timestamp_utc':datetime.datetime.now(datetime.timezone.utc).isoformat(),
    'documentation_access_date':'2026-09-29',
    'heliquest_live_execution':{'executed':False,'reason':'HELIQUEST is used as a documentary reference for the scale and formula.', 'raw_visible_server_result_fields':None},
    'executed_comparators':['Helix Design Studio app.engine hydrophobic_moment_descriptor(), model_net_charge()','modlAMP PeptideDescriptor.calculate_global(), calculate_moment(), GlobalDescriptor.calculate_charge()'],
    'protocol':{
        'inputs':'Three hand-specified synthetic 20-residue probes plus three S/T scale regression controls.',
        'window_length':20,'angle_deg':100,'moment_normalization':'Magnitude divided by sequence length N.',
        'scale_name':'Fauchere-Pliska 1983, exact values documented by HELIQUEST',
        'scale_values':fp,
        'modlamp_scale_configuration':'PeptideDescriptor instantiated with Eisenberg then public scale dictionary replaced by the explicitly supplied FP mapping {residue:[value]}; scalename label changed. The executed algorithm is unmodified modlAMP. These are not default-scale modlAMP results.',
        'hds_mean_note':'HDS moment and charge functions were executed directly; H mean was computed from the actual engine HYDROPHOBICITY_FP dictionary, not returned by an HDS mean-H API.',
        'heliquest_formula_note':'Independent local formula check based on HELIQUEST documentation; never a live-server result.',
        'charge_conventions':'HDS and documented HELIQUEST use K+R-D-E with neutral histidine and uncharged termini. modlAMP charge uses pKa-based fractional ionization at pH 7.4 with free termini (amide=False); its results round to three decimals. Their different charges are expected and are not agreement failures.',
        'scope_limit':'Descriptor correctness check on three synthetic probes, not a biological, runtime, optimization-quality, usability, or broad software benchmark.'},
    'provenance':{
        'hds_repository':'https://github.com/SIMBIOS-USC/helix-design-studio-web',
        'hds_git_head':git_value('rev-parse','HEAD'),
        'hds_git_status':git_value('status','--short'),
        'engine_path':engine_file.resolve().relative_to(root).as_posix(),
        'engine_sha256':hashlib.sha256(engine_file.read_bytes()).hexdigest(),
        'engine_function_source':inspect.getsource(engine.hydrophobic_moment_descriptor),
        'modlamp_descriptors_sha256':hashlib.sha256(package_file.read_bytes()).hexdigest(),
        'python_version':platform.python_version(),'platform':platform.platform(),
        'packages':{name:importlib.metadata.version(name) for name in ['modlamp','numpy','scipy','joblib']},
        'environment':'Isolated comparator virtual environment; production dependencies were not changed.',
        'rerun_command':'python experiments/compare_descriptors.py ../helix-design-studio-web results/heliquest_comparison.json',
        'installation_commands':['python -m venv .venv-comparison', '.venv-comparison/bin/python -m pip install modlamp==4.3.2'],
        'reproduction_script_path':'experiments/compare_descriptors.py',
        'reproduction_script_sha256':hashlib.sha256(source_file.read_bytes()).hexdigest(),
        'reproduction_script':source_file.read_text()},
    'sources':[
        {'url':'https://heliquest.ipmc.cnrs.fr/HelpProcedure.htm','kind':'Official HELIQUEST methods and numerical FP table','facts_checked':{'Ser':-0.040,'Thr':0.260,'Leu':1.700,'Lys':-0.990,'Glu':-0.640,'angle_deg':100,'charge_pH_nominal':7.4,'charge_convention':'K,R +1; D,E -1; His and termini neutral'}},
        {'url':'https://modlamp.org/modlamp.html','kind':'Official modlAMP descriptor documentation (website labels v4.3.0; executed package 4.3.2)'},
        {'url':'https://doi.org/10.1093/bioinformatics/btx285','kind':'modlAMP original paper'},
        {'url':'https://doi.org/10.1093/bioinformatics/btn392','kind':'HELIQUEST original paper'}],
    'scale_audit':{
        'all_20_hds_scale_values_equal_documented_FP':not bool(scale_mismatches),'mismatches':scale_mismatches,
        'control_rationale':'Balanced alternating ST can hide an S/T swap in mean and moment magnitude; S and T homopolymers and a full 20-residue table comparison also check scale identity.'},
    'rows':rows,
    'summary':{
        'primary_probe_count':3,'regression_control_count':3,
        'max_abs_difference_mean_H_all_six':max(r['absolute_difference_hds_vs_modlamp']['mean_hydrophobicity'] for r in rows),
        'max_abs_difference_moment_all_six':max(r['absolute_difference_hds_vs_modlamp']['hydrophobic_moment_per_residue'] for r in rows)}
}
assert not scale_mismatches, scale_mismatches
assert result['summary']['max_abs_difference_moment_all_six'] < 1e-12
output.parent.mkdir(parents=True,exist_ok=True)
output.write_text(json.dumps(result,indent=2,ensure_ascii=False,allow_nan=False)+'\n')
print(json.dumps({'output':str(output),'summary':result['summary'],'rows':rows},indent=2))
```
