## Additional file 2. Computational evaluation, inputs and results for "Helix Design Studio: designing antimicrobial peptide candidates for membrane environments": compare_m1_scoring.py

```
#!/usr/bin/env python3
"""Isolate the effect of subtracting M1 while retaining source-verified tables.

Both conditions use the same current engine, aligned geometry, component weights,
random samples and search budgets. The positive-M1 condition is an experimental
control created in memory, not a run of a historical software release.

    python experiments/compare_m1_scoring.py --repo ../helix-design-studio-web \
        --output results/m1_scoring_comparison.json
"""
from __future__ import annotations

import argparse
import ast
from contextlib import contextmanager
from datetime import datetime, timezone
import hashlib
import json
import math
import os
from pathlib import Path
import platform
import sys
import tempfile
import time

import numpy as np

from run_examples import application_version, dump, source_metadata


def sha256(path):
    return hashlib.sha256(path.read_bytes()).hexdigest()


def benchmark_settings(repo):
    """Read the existing benchmark's literal inputs without importing its runtime."""
    path = repo / "tools/measure_changes.py"
    names = {"N_DECOYS", "N_RANDOM", "SEED", "ALPHABET", "PRESETS", "PEPTIDES", "DESIGN"}
    values = {}
    for node in ast.parse(path.read_text(encoding="utf-8")).body:
        if isinstance(node, ast.Assign):
            for target in node.targets:
                if isinstance(target, ast.Name) and target.id in names:
                    value = node.value
                    # ALPHABET is declared as list("...") in the benchmark.
                    if target.id == "ALPHABET" and isinstance(value, ast.Call):
                        if not (isinstance(value.func, ast.Name) and value.func.id == "list"
                                and len(value.args) == 1 and not value.keywords):
                            raise ValueError("Unexpected benchmark alphabet declaration")
                        values[target.id] = list(ast.literal_eval(value.args[0]))
                    else:
                        values[target.id] = ast.literal_eval(value)
    if set(values) != names:
        raise ValueError(f"Missing benchmark inputs: {sorted(names - set(values))}")
    if len(values["PEPTIDES"]) * len(values["PRESETS"]) != 60:
        raise ValueError("The expected benchmark comprises 12 sequences in five environments")
    design = values["DESIGN"]
    if len(design["seeds"]) * len(design["presets"]) != 6:
        raise ValueError("The expected design benchmark comprises six paired searches")
    return values, {"path": str(path.relative_to(repo)), "sha256": sha256(path)}


def require_unchanged(before, repo):
    after = source_metadata(repo)
    if before != after:
        raise RuntimeError("Runtime source or Git state changed during the experiment; discard its outputs")
    return after


def positive_m1_neighbour(self, sequence):
    """Pre-correction sign, applied to the same current numerical matrices."""
    total = 0.0
    for separation, matrix in ((1, self.M1), (3, self.M3), (4, self.M4)):
        if self.L <= separation:
            continue
        scale = 1.0 / (self.L - separation)
        for index in range(self.L - separation):
            total += matrix[sequence[index], sequence[index + separation]] * scale
    return total


@contextmanager
def convention_case(model, positive, cache):
    """Change only neighbour convention in memory; keep isolated empty caches."""
    original_method = model.HamiltonianBuilder._raw_helix_neigh
    original_name = model.NEIGHBOR_SCORING_CONVENTION
    original_weights = model.NEIGHBOR_MATRIX_WEIGHTS
    previous_cache = os.environ.get("HELIX_CACHE_DIR")
    if positive:
        model.HamiltonianBuilder._raw_helix_neigh = positive_m1_neighbour
        model.NEIGHBOR_SCORING_CONVENTION = "positive_m1_plus_m3_plus_m4_experimental_control"
        model.NEIGHBOR_MATRIX_WEIGHTS = {1: 1.0, 3: 1.0, 4: 1.0}
    os.environ["HELIX_CACHE_DIR"] = str(cache)
    try:
        yield
    finally:
        model.HamiltonianBuilder._raw_helix_neigh = original_method
        model.NEIGHBOR_SCORING_CONVENTION = original_name
        model.NEIGHBOR_MATRIX_WEIGHTS = original_weights
        if previous_cache is None:
            os.environ.pop("HELIX_CACHE_DIR", None)
        else:
            os.environ["HELIX_CACHE_DIR"] = previous_cache


def sequence_sample_hash(length, alphabet_size, count, entropy):
    """Identify the shared uniform random sample by its ordered residue indices."""
    rng = np.random.default_rng(entropy)
    digest = hashlib.sha256()
    for _ in range(count):
        digest.update(rng.integers(0, alphabet_size, size=length).astype(np.uint8).tobytes())
    return digest.hexdigest()


def convention_checks(model):
    """Check direction, linearity and zero handling without calibration or caches."""
    builder = object.__new__(model.HamiltonianBuilder)
    builder.L = 5
    builder.M1 = np.array([[0.0, 0.7], [1.3, 0.0]])
    builder.M3 = np.zeros((2, 2))
    builder.M4 = np.zeros((2, 2))
    sequence = [0, 1, 0, 1, 0]
    original = float(builder._raw_helix_neigh(sequence))
    builder.M1[0, 1] += 0.5
    changed = float(builder._raw_helix_neigh(sequence))
    if not (math.isclose(original, -1.0, abs_tol=1e-12)
            and math.isclose(changed - original, -0.25, abs_tol=1e-12)):
        raise RuntimeError("The current neighbour formula does not linearly favour greater M1")
    builder.M1[:] = 0.0
    zeros = float(builder._raw_helix_neigh(sequence))
    if not math.isfinite(zeros) or zeros != 0.0:
        raise RuntimeError("Zero M1 propensities must remain finite zero contributions")
    return {"increasing_m1_lowers_raw_neighbour_score": True,
            "tested_m1_increment": 0.5, "affected_pairs": 2, "pair_count": 4,
            "raw_neighbour_change": changed - original,
            "zero_m1_raw_contribution": zeros,
            "transformation": "Signed linear coefficients; no logarithmic transform"}


def evaluate_case(engine, model, settings, positive, cache):
    started = time.perf_counter()
    alphabet = settings["ALPHABET"]
    n_decoys, n_random, seed = (settings[key] for key in ("N_DECOYS", "N_RANDOM", "SEED"))
    weights = dict(engine.DEFAULT_WEIGHTS)
    scores, designs = [], []
    with convention_case(model, positive, cache):
        info = {"neighbor_score_convention": model.NEIGHBOR_SCORING_CONVENTION,
                "neighbor_matrix_weights": dict(model.NEIGHBOR_MATRIX_WEIGHTS),
                "parameter_set_sha256": model.parameter_set_sha256(),
                "experimental_control": bool(positive)}
        for name, sequence in settings["PEPTIDES"].items():
            for preset in settings["PRESETS"]:
                result = engine.score_sequence(sequence, {"preset": preset}, alphabet, weights,
                                               n_decoys=n_decoys, n_random=n_random, seed=seed)
                scores.append({"sequence_name": name, "sequence": sequence, "environment": preset,
                               "energy": result["energy"], "z_score": result["z_score"],
                               "calibration": result["calibration"],
                               "random_reference": result["random_reference"],
                               "wheel_phase_deg": result["environment"]["wheel_phase_deg"]})
        design = settings["DESIGN"]
        for preset in design["presets"]:
            for design_seed in design["seeds"]:
                result = engine.design_sequence(design["length"], {"preset": preset}, alphabet, weights,
                                                n_decoys=n_decoys, steps=design["steps"],
                                                restarts=design["restarts"], seed=design_seed)
                designs.append({"environment": preset, "seed": design_seed, "sequence": result["sequence"],
                                "energy": result["energy"], "z_score": result["z_score"],
                                "calibration": result["calibration"],
                                "random_reference": result["random_reference"],
                                "wheel_phase_deg": result["environment"]["wheel_phase_deg"]})
    return {**info, "scores": scores, "designs": designs, "wall_seconds": time.perf_counter() - started}


def tied_ranks(values):
    ordered = np.sort(np.asarray(values, dtype=float))
    return np.array([float(np.flatnonzero(ordered == value).mean()) for value in values])


def spearman(left, right):
    a, b = tied_ranks(left), tied_ranks(right)
    if np.std(a) == 0 or np.std(b) == 0:
        return None
    return float(np.corrcoef(a, b)[0, 1])


def compare_cases(positive, negative):
    score_pairs = []
    for old, new in zip(positive["scores"], negative["scores"], strict=True):
        keys = ("sequence_name", "sequence", "environment")
        if any(old[key] != new[key] for key in keys):
            raise RuntimeError("Score rows do not describe matched inputs")
        score_pairs.append({**{key: old[key] for key in keys},
                            "positive_m1_energy": old["energy"], "negative_m1_energy": new["energy"],
                            "delta_energy": new["energy"] - old["energy"],
                            "positive_m1_z_score": old["z_score"], "negative_m1_z_score": new["z_score"],
                            "delta_z_score": new["z_score"] - old["z_score"]})
    design_pairs = []
    for old, new in zip(positive["designs"], negative["designs"], strict=True):
        if (old["environment"], old["seed"]) != (new["environment"], new["seed"]):
            raise RuntimeError("Design rows do not describe matched searches")
        distance = sum(a != b for a, b in zip(old["sequence"], new["sequence"], strict=True))
        design_pairs.append({"environment": old["environment"], "seed": old["seed"],
                             "positive_m1_sequence": old["sequence"], "negative_m1_sequence": new["sequence"],
                             "exact_match": old["sequence"] == new["sequence"], "hamming_distance": distance,
                             "length": len(old["sequence"])})
    summary = {"score_count": len(score_pairs), "paired_design_count": len(design_pairs),
               "design_record_count": len(positive["designs"]) + len(negative["designs"]),
               "exact_design_matches": sum(row["exact_match"] for row in design_pairs),
               "changed_design_positions": sum(row["hamming_distance"] for row in design_pairs),
               "design_positions_evaluated": sum(row["length"] for row in design_pairs),
               "minimum_design_hamming_distance": min(row["hamming_distance"] for row in design_pairs),
               "maximum_design_hamming_distance": max(row["hamming_distance"] for row in design_pairs)}
    summary["changed_design_position_fraction"] = summary["changed_design_positions"] / summary["design_positions_evaluated"]
    for name, field in (("energy", "energy"), ("z_score", "z_score")):
        delta = np.array([row[f"delta_{field}"] for row in score_pairs])
        summary[f"median_absolute_delta_{name}"] = float(np.median(np.abs(delta)))
        summary[f"maximum_absolute_delta_{name}"] = float(np.max(np.abs(delta)))
        summary[f"spearman_{name}"] = spearman([row[field] for row in positive["scores"]],
                                               [row[field] for row in negative["scores"]])
    return summary, score_pairs, design_pairs


def main():
    parser = argparse.ArgumentParser(description=__doc__.split("\n\n")[0])
    parser.add_argument("--repo", required=True, type=Path)
    parser.add_argument("--output", required=True, type=Path)
    args = parser.parse_args()
    repo, output = args.repo.resolve(), args.output.resolve()
    if output.exists():
        raise SystemExit("Refusing to replace an existing result; choose a new --output file")
    before = source_metadata(repo)
    if not before["commit"] or before["git_status"]:
        raise SystemExit("An identified, clean runtime revision is required")
    settings, benchmark_source = benchmark_settings(repo)
    os.environ.update(HELIX_CODE_DIR=str(repo / "Code"), QFOLD_CODE_DIR=str(repo / "Code"), HELIX_USAGE_LOGGING="0")
    sys.path.insert(0, str(repo))
    from app import engine
    from core import hamiltonian_builder as model
    if Path(engine.__file__).resolve() != repo / "app/engine.py":
        raise RuntimeError("Imported an engine from another repository")
    if model.NEIGHBOR_MATRIX_WEIGHTS != {1: -1.0, 3: 1.0, 4: 1.0}:
        raise RuntimeError("Expected the current negative-M1 scoring convention")
    checks = convention_checks(model)
    started = datetime.now(timezone.utc).isoformat()
    cases = {}
    for positive, name in ((True, "positive_m1_control"), (False, "negative_m1_current")):
        with tempfile.TemporaryDirectory(prefix=f"helix-m1-{name}-") as cache:
            cases[name] = evaluate_case(engine, model, settings, positive, Path(cache))
        require_unchanged(before, repo)
        print(f"Completed {name}", flush=True)
    summary, score_pairs, design_pairs = compare_cases(cases["positive_m1_control"], cases["negative_m1_current"])
    ensembles = []
    lengths = sorted({len(sequence) for sequence in settings["PEPTIDES"].values()})
    samples = {(length, settings["SEED"]) for length in lengths}
    samples.update((settings["DESIGN"]["length"], seed) for seed in settings["DESIGN"]["seeds"])
    for length, seed in sorted(samples):
        ensembles.append({"length": length, "seed": seed,
                          "calibration_sample_sha256": sequence_sample_hash(length, len(settings["ALPHABET"]), settings["N_DECOYS"], seed),
                          "reference_sample_sha256": sequence_sample_hash(length, len(settings["ALPHABET"]), settings["N_RANDOM"], engine.rng_seed(seed, engine.REFERENCE_STREAM))})
    report = {"comparison": "Isolated change from +M1 to -M1 with the same source-verified numerical tables",
              "started_utc": started, "finished_utc": datetime.now(timezone.utc).isoformat(),
              "source": before, "final_source": require_unchanged(before, repo), "source_unchanged": True,
              "software_version": application_version(repo),
              "experiment_script_sha256": sha256(Path(__file__).resolve()),
              "benchmark_source": benchmark_source,
              "runtime": {"python": sys.version, "numpy": np.__version__, "platform": platform.platform()},
              "parameter_tables_sha256": {name: sha256(repo / "Code/core/resources" / name) for name in model.RESOURCE_FILES},
              "protocol": {"alphabet": settings["ALPHABET"], "peptides": settings["PEPTIDES"],
                           "presets": settings["PRESETS"], "weights": dict(engine.DEFAULT_WEIGHTS),
                           "n_decoys": settings["N_DECOYS"], "n_random": settings["N_RANDOM"],
                           "score_seed": settings["SEED"], "design": settings["DESIGN"],
                           "seed_protocol": engine.SEED_PROTOCOL,
                           "orientation": "Each query, proposal, calibration sequence and random reference uses its own hydrophobic-moment alignment at interfaces",
                           "environment_settings": {preset: engine.resolve_environment({"preset": preset}) for preset in settings["PRESETS"]},
                           "calibration": "Recomputed separately in fresh condition-specific caches, using identical sampled sequences",
                           "cache_policy": "One newly created temporary directory per convention, removed after that condition",
                           "shared_random_samples": ensembles,
                           "control_scope": "Positive-M1 control is a process-local intervention in the identified current engine, not a historical software release",
                           "delta_definition": "negative-M1 current minus positive-M1 experimental control",
                           "interpretation": "Paired model calculations and stochastic searches; native scores use condition-specific calibration and reference statistics"},
              "formula_checks": checks, "cases": cases, "score_pairs": score_pairs,
              "design_pairs": design_pairs, "summary": summary}
    dump(output, report)
    print(json.dumps(summary, indent=2, allow_nan=False))


if __name__ == "__main__":
    main()
```
