## Additional file 2. Computational evaluation, inputs and results for "Helix Design Studio: designing antimicrobial peptide candidates for membrane environments": compare_search_orientation.py

```
#!/usr/bin/env python3
"""Compare fixed and hydrophobic-moment-aligned searches on one immutable runtime.

Only the search-model orientation is changed. Each mode/seed is executed in a
fresh subprocess and private cache. All recovered candidates are then evaluated
in a separate process using one common aligned calibration/reference protocol.
No requests are sent to a deployed service and no runtime source is modified.
"""
from __future__ import annotations

import argparse
import ast
from contextlib import redirect_stdout
from datetime import datetime, timezone
import hashlib
import itertools
import json
import math
import os
from pathlib import Path
import platform
import statistics
import subprocess
import sys
import time

from run_examples import ALPHABET, PRESETS, WEIGHTS, application_version, csv_write, dump, finite, source_metadata


BASELINE = "18adbab6f4076580a0481e559d84aebe724814f1"
SEEDS = (42, 43, 44)
COMMON = {"length": 20, "residues": ALPHABET, "weights": WEIGHTS, "n_decoys": 2000, "steps": 3000}
REQUESTS = {
    "design": {**COMMON, "environment": {"preset": "interfacial_neg"}, "restarts": 2},
    "family": {**COMMON, "environment": {"preset": "interfacial_neg"}, "restarts": 2,
               "family_size": 4, "oversample_factor": 2},
    "specificity": {**COMMON, "target_environment": {"preset": "interfacial_neg"},
                    "off_target_environments": [{"preset": "interfacial_neu"}, {"preset": "homogeneous_polar"}],
                    "restarts": 4, "num_sequences": 2, "lambda_balance": 0.5},
    "cross_design": {**COMMON, "environment_a": {"preset": "interfacial_neg"},
                     "environment_b": {"preset": "homogeneous_polar"}, "restarts": 2, "lambda_gap": 0.5},
}


def sha256(path):
    return hashlib.sha256(path.read_bytes()).hexdigest()


def ensure_same_source(before, after):
    if before != after:
        raise RuntimeError("Runtime source or Git state changed during the comparison; results must be discarded.")


def import_engine(repo, output):
    cache = output / "calibration_cache"
    cache.mkdir(parents=True, exist_ok=True, mode=0o700)
    os.environ.update(HELIX_CACHE_DIR=str(cache), HELIX_VAR_DIR=str(output / "var"),
                      HELIX_USAGE_LOGGING="0", HELIX_CODE_DIR=str(repo / "Code"), QFOLD_CODE_DIR=str(repo / "Code"))
    sys.path.insert(0, str(repo))
    from app import engine
    from core.hamiltonian_builder import parameter_set_sha256
    return engine, parameter_set_sha256()


def native_objective(workflow, candidate):
    if workflow in ("design", "family"):
        return candidate["design_geometry"]["energy"]
    if workflow == "specificity":
        target = candidate["target_environment"]["design_geometry"]["energy"]
        off = [score["design_geometry"]["energy"] for score in candidate["off_target_environments"]]
        reconstructed = target - 0.5 * min(off)
        stored = candidate["specificity_design"]["objective"]
    else:
        a, b = [candidate[name]["design_geometry"]["energy"] for name in ("environment_a", "environment_b")]
        reconstructed = a + b + 0.5 * abs(a - b)
        stored = candidate["cross_design"]["objective"]
    if not math.isclose(stored, reconstructed, rel_tol=0, abs_tol=1e-8):
        raise RuntimeError(f"Native {workflow} objective cannot be reconstructed from returned search scores")
    return stored


def search_worker(args):
    repo, output = args.repo.resolve(), args.output.resolve()
    output.mkdir(parents=True, exist_ok=True)
    before = source_metadata(repo)
    engine, parameter_sha = import_engine(repo, output)
    original_report_models = engine.report_models

    def selected_models(length, amino_acids, environment, weights, *, n_decoys, seed, n_random):
        aligned = engine.scoring_model(length, amino_acids, environment, weights,
                                       n_decoys=n_decoys, seed=seed, n_random=n_random, oriented=True)
        if args.worker == "aligned" or not engine.is_interfacial(environment):
            return aligned, aligned
        fixed = engine.scoring_model(length, amino_acids, environment, weights,
                                     n_decoys=n_decoys, seed=seed, n_random=n_random, oriented=False)
        return fixed, aligned

    # This process-local replacement is the sole experimental intervention.
    engine.report_models = selected_models
    functions = {"design": engine.design_sequence, "family": engine.design_family,
                 "specificity": engine.design_specificity, "cross_design": engine.cross_environment_design}
    candidates, calls = [], []
    started = time.perf_counter()
    with (output / "runtime.log").open("w", encoding="utf-8") as log:
        for workflow, template in REQUESTS.items():
            request = {**template, "seed": args.seed}
            call_start = time.perf_counter()
            with redirect_stdout(log):
                result = functions[workflow](**request)
            elapsed = time.perf_counter() - call_start
            if not finite(result):
                raise RuntimeError(f"Nonfinite {workflow} result")
            members = (result["members"] if workflow == "family" else
                       result["candidates"] if workflow == "specificity" else [result])
            returned_ranks = {candidate["sequence"]: rank for rank, candidate in enumerate(members, 1)}
            # Preserve the response verbatim, but classify each collection by
            # its actual search objective. A standard-report sort would mix
            # orientation rules in the fixed-search counterfactual.
            if workflow == "family":
                members = sorted(members, key=lambda item: item["design_geometry"]["energy"])
            elif workflow == "specificity":
                members = sorted(members, key=lambda item: (
                    round(float(item["specificity_design"]["objective"]), 8),
                    -float(item["specificity_design"]["z_score_margin"]),
                    float(item["target_environment"]["design_geometry"]["z_score"]), item["sequence"]))
            records = []
            for rank, candidate in enumerate(members, 1):
                record = {"mode": args.worker, "seed": args.seed, "workflow": workflow,
                          "native_rank": rank, "returned_rank": returned_ranks[candidate["sequence"]],
                          "sequence": candidate["sequence"],
                          "native_objective": native_objective(workflow, candidate)}
                records.append(record)
            candidates.extend(records)
            call = {"workflow": workflow, "mode": args.worker, "seed": args.seed,
                    "request": request, "wall_seconds": elapsed, "returned_candidates": len(records)}
            calls.append(call)
            dump(output / f"{workflow}.json", {**call, "response": result})
            print(f"{args.worker} seed {args.seed} {workflow}: {elapsed:.3f}s", flush=True)
    engine.report_models = original_report_models
    after = source_metadata(repo)
    ensure_same_source(before, after)
    dump(output / "candidates.json", candidates)
    dump(output / "metadata.json", {"source": before, "final_source": after, "parameter_set_sha256": parameter_sha,
                                    "mode": args.worker, "seed": args.seed, "calls": calls,
                                    "total_wall_seconds": time.perf_counter() - started,
                                    "intervention": "process-local replacement of engine.report_models; search oriented=" + str(args.worker == "aligned"),
                                    "native_ranking": "family: search energy; specificity: rounded search objective, decreasing search Z margin, target search Z, sequence; returned_rank preserves runtime reporting order",
                                    "reporting": "both modes retain aligned standard reports; design_geometry records the actual search rule",
                                    "resolved_environment_keys": {preset: list(engine.physical_environment_key(engine.resolve_environment({"preset": preset}))) for preset in PRESETS}})


def common_objective(workflow, scores):
    target = scores["interfacial_neg"]["energy"]
    if workflow in ("design", "family"):
        return target
    if workflow == "specificity":
        return target - 0.5 * min(scores[preset]["energy"] for preset in ("interfacial_neu", "homogeneous_polar"))
    a, b = target, scores["homogeneous_polar"]["energy"]
    return a + b + 0.5 * abs(a - b)


def rescore_worker(args):
    repo, output = args.repo.resolve(), args.output.resolve()
    output.mkdir(parents=True, exist_ok=True)
    before = source_metadata(repo)
    engine, parameter_sha = import_engine(repo, output)
    all_candidates = json.loads((output.parent / "candidates.json").read_text())
    sequences = sorted({item["sequence"] for item in all_candidates})
    scores = {sequence: {} for sequence in sequences}
    references = {}
    with (output / "runtime.log").open("w", encoding="utf-8") as log, redirect_stdout(log):
        for preset in PRESETS:
            environment = engine.resolve_environment({"preset": preset})
            model = engine.scoring_model(20, ALPHABET, environment, WEIGHTS,
                                         n_decoys=2000, seed=42, n_random=1000, oriented=True)
            references[preset] = {"environment": environment, "reference": model.reference,
                                  "calibration": model.builder.stats, "align_phase": bool(model.builder.align_phase)}
            for sequence in sequences:
                scores[sequence][preset] = engine.score_with_model(sequence, model, ALPHABET, WEIGHTS)
    dump(output / "scores.json", scores)
    dump(output / "backgrounds.json", references)
    rows = []
    for item in all_candidates:
        evaluated = scores[item["sequence"]]
        row = {**item, "common_aligned_objective": common_objective(item["workflow"], evaluated)}
        if item["workflow"] == "specificity":
            off = [evaluated[preset] for preset in ("interfacial_neu", "homogeneous_polar")]
            row["common_energy_margin"] = min(score["energy"] for score in off) - evaluated["interfacial_neg"]["energy"]
            row["common_z_score_margin"] = min(score["z_score"] for score in off) - evaluated["interfacial_neg"]["z_score"]
        else:
            row["common_energy_margin"], row["common_z_score_margin"] = None, None
        rows.append(row)
    for row in rows:
        peers = [other for other in rows if (other["mode"], other["seed"], other["workflow"]) == (row["mode"], row["seed"], row["workflow"])]
        row["common_rank_within_collection"] = 1 + sum(other["common_aligned_objective"] < row["common_aligned_objective"] - 1e-9 for other in peers)
        pooled = [other for other in rows if (other["seed"], other["workflow"]) == (row["seed"], row["workflow"])]
        row["common_rank_within_matched_seed"] = 1 + sum(other["common_aligned_objective"] < row["common_aligned_objective"] - 1e-9 for other in pooled)
    csv_write(output / "candidate_scores.csv", rows)
    dump(output / "candidate_scores.json", rows)
    after = source_metadata(repo)
    ensure_same_source(before, after)
    dump(output / "metadata.json", {"source": before, "final_source": after, "parameter_set_sha256": parameter_sha,
                                    "protocol": {"alphabet": ALPHABET, "weights": WEIGHTS, "n_decoys": 2000,
                                                 "n_random": 1000, "seed": 42, "orientation": "own hydrophobic moment at interfaces"},
                                    "unique_sequences": len(sequences)})


def parameter_identity(repo, baseline):
    records = {}
    for path in sorted((repo / "Code/core/resources").glob("*.txt")):
        relative = str(path.relative_to(repo))
        original = subprocess.run(["git", "-C", str(repo), "show", f"{baseline}:{relative}"], check=True, capture_output=True).stdout
        records[relative] = {"baseline_sha256": hashlib.sha256(original).hexdigest(), "current_sha256": sha256(path)}
    if not all(record["baseline_sha256"] == record["current_sha256"] for record in records.values()):
        raise RuntimeError("Parameter files differ from the specified baseline; this would confound the comparison")
    # The parameter checksum also includes the per-residue scales, so check
    # those explicitly without importing the baseline runtime into this process.
    relative = "Code/core/hamiltonian_builder.py"
    original = subprocess.run(["git", "-C", str(repo), "show", f"{baseline}:{relative}"], check=True, capture_output=True, text=True).stdout
    names = {"PACE_SCHOLTZ_HELIX_PROPENSITY", "FAUCHERE_PLISKA_HYDROPHOBICITY"}
    def scales(source):
        return {target.id: ast.literal_eval(node.value)
                for node in ast.parse(source).body if isinstance(node, ast.Assign)
                for target in node.targets if isinstance(target, ast.Name) and target.id in names}
    old_scales, new_scales = scales(original), scales((repo / relative).read_text())
    if set(old_scales) != names or old_scales != new_scales:
        raise RuntimeError("Per-residue parameter scales differ from the baseline")
    records["per_residue_scales"] = {"baseline_sha256": hashlib.sha256(json.dumps(old_scales, sort_keys=True).encode()).hexdigest(),
                                     "current_sha256": hashlib.sha256(json.dumps(new_scales, sort_keys=True).encode()).hexdigest()}
    return records


def hamming_distances(rows):
    sequences = [row["sequence"] for row in rows]
    values = [sum(a != b for a, b in zip(left, right)) for left, right in itertools.combinations(sequences, 2)]
    return {"unique_sequences": len(set(sequences)), "pair_count": len(values),
            "minimum_hamming_distance": min(values) if values else None,
            "mean_hamming_distance": statistics.fmean(values) if values else None}


def comparison_master(args):
    repo, output = args.repo.resolve(), args.output.resolve()
    if output.exists() and any(output.iterdir()):
        raise SystemExit("Refusing to overwrite an existing comparison output; choose an empty --output directory")
    output.mkdir(parents=True, exist_ok=True)
    before = source_metadata(repo)
    if not before["commit"] or before["git_status"]:
        raise SystemExit("Use an identified, clean runtime checkout for the comparison")
    baseline_commit = subprocess.run(["git", "-C", str(repo), "rev-parse", args.baseline_ref], check=True, capture_output=True, text=True).stdout.strip()
    protocol = {"source": before, "version": application_version(repo), "baseline_commit": baseline_commit,
                "parameter_file_identity": parameter_identity(repo, baseline_commit),
                "python": sys.version, "platform": platform.platform(), "seeds": list(SEEDS), "request_templates": REQUESTS,
                "comparison_script_sha256": sha256(Path(__file__)),
                "started_utc": datetime.now(timezone.utc).isoformat(),
                "intervention": "One runtime snapshot; only the search orientation flag differs between modes. Each mode uses calibration and reference sequences under its own search rule; standard reporting is aligned in both modes.",
                "baseline_interpretation": "The fixed mode is a counterfactual on the identified current runtime, not an execution of the historical baseline commit. The baseline identifies the unchanged numerical parameter set. Family/Specificity ranking is reconstructed under the actual search objective, with returned order retained separately; reordering does not change the collected sequence set.",
                "common_evaluation": "All returned candidates rescored together in a separate process, aligned, n_decoys=2000, n_random=1000, seed=42. Objective energy is independent of reference-ensemble size.",
                "runtime_interpretation": "Sequential local engine wall times include search-model calibration, reference calculation, search and aligned reporting. They are not pure optimizer timings, deployment timings or capacity measurements; unrelated machine processes are not suspended.",
                "interpretation": "Paired finite-budget computational comparison only. Lower common objective is preferred by this scoring model; no biological performance or global optimum is implied."}
    dump(output / "protocol.json", protocol)
    started, candidates, calls = time.perf_counter(), [], []
    for index, seed in enumerate(SEEDS):
        # Alternate order to reduce a consistent first-mode timing advantage.
        for mode in (("fixed", "aligned") if index % 2 == 0 else ("aligned", "fixed")):
            child = output / f"{mode}_seed{seed}"
            subprocess.run([sys.executable, str(Path(__file__).resolve()), "--repo", str(repo), "--output", str(child), "--worker", mode, "--seed", str(seed)], check=True)
            candidates.extend(json.loads((child / "candidates.json").read_text()))
            calls.extend(json.loads((child / "metadata.json").read_text())["calls"])
            ensure_same_source(before, source_metadata(repo))
    dump(output / "candidates.json", candidates)
    subprocess.run([sys.executable, str(Path(__file__).resolve()), "--repo", str(repo), "--output", str(output / "common_aligned"), "--worker", "rescore"], check=True)
    rows = json.loads((output / "common_aligned/candidate_scores.json").read_text())
    paired, diversity = [], []
    for seed in SEEDS:
        for workflow in REQUESTS:
            groups = {mode: [row for row in rows if (row["seed"], row["workflow"], row["mode"]) == (seed, workflow, mode)] for mode in ("fixed", "aligned")}
            best = {mode: min(group, key=lambda row: row["common_aligned_objective"]) for mode, group in groups.items()}
            difference = best["aligned"]["common_aligned_objective"] - best["fixed"]["common_aligned_objective"]
            pair = {"seed": seed, "workflow": workflow,
                    "fixed_best_common_objective": best["fixed"]["common_aligned_objective"],
                    "aligned_best_common_objective": best["aligned"]["common_aligned_objective"],
                    "aligned_minus_fixed_common_objective": difference,
                    "preferred_mode_under_common_objective": "tie" if abs(difference) <= 1e-9 else "aligned" if difference < 0 else "fixed",
                    "fixed_best_sequence": best["fixed"]["sequence"], "aligned_best_sequence": best["aligned"]["sequence"],
                    "shared_sequences": len({row["sequence"] for row in groups["fixed"]} & {row["sequence"] for row in groups["aligned"]})}
            for mode, group in groups.items():
                pair[f"{mode}_rank_inversions_under_common_rule"] = sum((left["native_rank"] - right["native_rank"]) * (left["common_rank_within_collection"] - right["common_rank_within_collection"]) < 0 for left, right in itertools.combinations(group, 2))
                pair[f"{mode}_wall_seconds"] = next(call["wall_seconds"] for call in calls if (call["seed"], call["workflow"], call["mode"]) == (seed, workflow, mode))
                diversity.append({"seed": seed, "workflow": workflow, "mode": mode, **hamming_distances(group)})
            paired.append(pair)
    csv_write(output / "paired_objectives.csv", paired)
    csv_write(output / "collection_diversity.csv", diversity)
    csv_write(output / "calls.csv", calls)
    after = source_metadata(repo)
    ensure_same_source(before, after)
    summary = {"paired_results": paired, "collection_diversity": diversity, "candidate_count": len(rows),
               "workflow_summary": {workflow: {"aligned_preferred": sum(row["workflow"] == workflow and row["preferred_mode_under_common_objective"] == "aligned" for row in paired),
                                               "fixed_preferred": sum(row["workflow"] == workflow and row["preferred_mode_under_common_objective"] == "fixed" for row in paired),
                                               "ties": sum(row["workflow"] == workflow and row["preferred_mode_under_common_objective"] == "tie" for row in paired)} for workflow in REQUESTS},
               "source_unchanged": True, "final_source": after,
               "finished_utc": datetime.now(timezone.utc).isoformat(), "total_wall_seconds": time.perf_counter() - started,
               "exact_optimizer_check": "Run the companion run_examples.py, which enumerates all 4096 aligned length-six A/E/K/L sequences and compares five annealing trials. It is intentionally not duplicated here.",
               "interpretation": protocol["interpretation"]}
    dump(output / "summary.json", summary)
    retained = [path for path in output.rglob("*") if path.is_file() and path.suffix != ".log" and "calibration_cache" not in path.parts]
    dump(output / "file_manifest.json", {str(path.relative_to(output)): sha256(path) for path in sorted(retained)})
    print(f"Completed {len(paired)} paired comparisons, {len(rows)} candidates, in {summary['total_wall_seconds']:.1f}s", flush=True)


def main():
    parser = argparse.ArgumentParser(description=__doc__)
    parser.add_argument("--repo", type=Path, required=True)
    parser.add_argument("--output", type=Path, required=True)
    parser.add_argument("--baseline-ref", default=BASELINE)
    parser.add_argument("--worker", choices=("fixed", "aligned", "rescore"), help=argparse.SUPPRESS)
    parser.add_argument("--seed", type=int, choices=SEEDS, help=argparse.SUPPRESS)
    args = parser.parse_args()
    if args.worker == "rescore":
        rescore_worker(args)
    elif args.worker:
        if args.seed is None:
            parser.error("Search workers require --seed")
        search_worker(args)
    else:
        comparison_master(args)


if __name__ == "__main__":
    main()
```
