## Additional file 2. Computational evaluation, inputs and results for "Helix Design Studio: designing antimicrobial peptide candidates for membrane environments": export_parameter_tables.py

```
#!/usr/bin/env python3
"""Verify the primary-source mappings and export the complete runtime tables.

ReportLab is a document-generation dependency, not an application dependency.
Source transcriptions and their independent axis labels are distributed with
the application in tools/data/parameter_sources/.
"""
from __future__ import annotations

import argparse
import ast
from datetime import datetime, timezone
from decimal import Decimal
import hashlib
import importlib.metadata
import json
from pathlib import Path
import platform
import runpy
import subprocess
import sys

from reportlab.lib import colors
from reportlab.lib.pagesizes import A4, landscape
from reportlab.pdfgen import canvas
from reportlab.platypus import Table, TableStyle


ROOT = Path(__file__).resolve().parents[1]
STANDARD = set("ARNDCQEGHILKMFPSTWYV")


def sha256(path):
    return hashlib.sha256(Path(path).read_bytes()).hexdigest()


def version(repo):
    for node in ast.parse((repo / "app/main.py").read_text(encoding="utf-8")).body:
        if isinstance(node, ast.Assign) and any(isinstance(target, ast.Name) and target.id == "APP_VERSION" for target in node.targets):
            value = ast.literal_eval(node.value)
            if isinstance(value, str) and value:
                return value
    raise ValueError("Missing application version")


def resource_table(path):
    lines = [line for line in path.read_text(encoding="utf-8").splitlines() if line.strip()]
    symbols = list("".join(lines[0].split()))
    tokens = [line.split() for line in lines[1:]]
    if len(symbols) != len(set(symbols)) or set(symbols) != STANDARD:
        raise ValueError(f"Invalid residue labels: {path.name}")
    if len(tokens) != 20 or any(len(row) != 20 for row in tokens):
        raise ValueError(f"Wrong runtime matrix dimensions: {path.name}")
    if any(not Decimal(value).is_finite() for row in tokens for value in row):
        raise ValueError(f"Nonfinite matrix value: {path.name}")
    return symbols, tokens


def make_pdf(path, matrices, commit, software_version):
    width, height = landscape(A4)
    margin = 34
    navy = colors.HexColor("#15324A")
    ink = colors.HexColor("#263744")
    grey = colors.HexColor("#56636E")
    grid = colors.HexColor("#C9D7DF")
    stripe = colors.HexColor("#F2F6F8")
    supplemental = colors.HexColor("#E4E8EB")
    pdf = canvas.Canvas(str(path), pagesize=(width, height), pageCompression=1, invariant=1)
    pdf.setTitle("Helix Design Studio: verified runtime parameter tables")
    pdf.setAuthor("Daniel Conde-Torres; Rebeca Garcia-Fandino; Angel Pineiro")
    pdf.setSubject("Primary-source values, indexing and documented model adaptations")
    for page, (name, matrix) in enumerate(matrices.items(), 1):
        k = matrix["sequence_separation"]
        symbols = matrix["row_residues"]
        pdf.setFillColor(navy)
        pdf.setFont("Helvetica-Bold", 10)
        pdf.drawString(margin, height - 29, f"HELIX DESIGN STUDIO  {software_version}")
        pdf.setFillColor(grey)
        pdf.setFont("Helvetica", 9)
        pdf.drawRightString(width - margin, height - 29, "VERIFIED RUNTIME PARAMETERS")
        pdf.setStrokeColor(grid)
        pdf.line(margin, height - 39, width - margin, height - 39)
        pdf.setFillColor(navy)
        pdf.setFont("Helvetica-Bold", 22)
        pdf.drawString(margin, height - 69, f"{name}: sequence separation k = {k}")
        pdf.setFillColor(ink)
        pdf.setFont("Helvetica", 10)
        pdf.drawString(margin, height - 88, "Rows: residue at i. Columns: residue at i+k. Each table contains 400 runtime values.")
        source = matrix["primary_reference"]
        if name == "M1":
            note = "Nacar (2020), Fig. 3, p. 25; source layout transposed to preserve sequence direction."
            adaptation = "Source pairs are n,n+4; the model reuses these dimensionless propensities at i,i+1."
        else:
            note = f"Munoz and Serrano (1994), Table 2, p. 403; i,i+{k} energies in kcal/mol, mapped by labels."
            adaptation = "C/P pairs (grey cells) are a 76-cell zero-contribution extension; they are absent from the source."
        pdf.drawString(margin, height - 105, note)
        pdf.drawString(margin, height - 120, adaptation)
        pdf.setFont("Helvetica", 8.5)
        pdf.drawString(margin, height - 135, "Raw source values. Score: -mean(M1) + mean(M3) + mean(M4), followed by joint standardization and weighting.")

        formatted = [[f"{Decimal(value):.3f}" for value in row] for row in matrix["numeric_tokens"]]
        data = [["i / i+k", *symbols]] + [[aa, *row] for aa, row in zip(symbols, formatted)]
        label_width = 29
        column_width = (width - 2 * margin - label_width) / 20
        table = Table(data, colWidths=[label_width] + [column_width] * 20, rowHeights=[23] + [17.5] * 20)
        style = [
            ("BACKGROUND", (0, 0), (-1, 0), navy),
            ("TEXTCOLOR", (0, 0), (-1, 0), colors.white),
            ("FONTNAME", (0, 0), (-1, 0), "Helvetica-Bold"),
            ("FONTSIZE", (0, 0), (0, 0), 7),
            ("FONTSIZE", (1, 0), (-1, 0), 9),
            ("FONTNAME", (0, 1), (0, -1), "Helvetica-Bold"),
            ("FONTSIZE", (0, 1), (0, -1), 9),
            ("TEXTCOLOR", (0, 1), (0, -1), navy),
            ("FONTNAME", (1, 1), (-1, -1), "Helvetica"),
            ("FONTSIZE", (1, 1), (-1, -1), 8.8),
            ("TEXTCOLOR", (1, 1), (-1, -1), ink),
            ("ALIGN", (0, 0), (-1, 0), "CENTER"),
            ("ALIGN", (0, 1), (0, -1), "CENTER"),
            ("ALIGN", (1, 1), (-1, -1), "RIGHT"),
            ("VALIGN", (0, 0), (-1, -1), "MIDDLE"),
            ("TOPPADDING", (0, 0), (-1, -1), 0),
            ("BOTTOMPADDING", (0, 0), (-1, -1), 0),
            ("LEFTPADDING", (0, 0), (-1, -1), 2),
            ("RIGHTPADDING", (0, 0), (-1, -1), 2),
            ("GRID", (0, 1), (-1, -1), 0.25, grid),
            ("LINEAFTER", (0, 0), (0, -1), 0.6, navy),
        ]
        for row in range(1, 21):
            style.append(("BACKGROUND", (0, row), (-1, row), stripe if row % 2 == 0 else colors.white))
        if name != "M1":
            for index, residue in enumerate(symbols, 1):
                if residue in {"C", "P"}:
                    style.append(("BACKGROUND", (1, index), (-1, index), supplemental))
                    style.append(("BACKGROUND", (index, 1), (index, -1), supplemental))
        table.setStyle(TableStyle(style))
        _, table_height = table.wrap(width - 2 * margin, height)
        bottom = height - 147 - table_height
        if bottom < 72:
            raise ValueError("Parameter table overlaps the citation/footer")
        table.drawOn(pdf, margin, bottom)

        pdf.setFillColor(grey)
        pdf.setFont("Helvetica", 8.4)
        citation = f"Source DOI: {source['doi']}"
        pdf.drawString(margin, 61, citation)
        pdf.linkURL(f"https://doi.org/{source['doi']}", (margin, 59, margin + 255, 71), relative=0, thickness=0)
        pdf.setFont("Helvetica", 7.7)
        resource_label = matrix["resource"]
        pdf.drawString(margin, 47, resource_label)
        pdf.linkURL(f"https://github.com/SIMBIOS-USC/helix-design-studio-web/blob/{commit}/{resource_label}",
                    (margin, 45, margin + 400, 55), relative=0, thickness=0)
        pdf.setFont("Helvetica", 6.9)
        pdf.drawString(margin, 34, f"Commit: {commit}")
        pdf.drawString(margin, 22, f"SHA-256 of resource: {matrix['sha256']}")
        pdf.setFont("Helvetica", 8)
        pdf.drawRightString(width - margin, 22, f"{page} / 3")
        pdf.showPage()
    pdf.save()


def main():
    parser = argparse.ArgumentParser(description=__doc__)
    parser.add_argument("--repo", type=Path, default=ROOT.parent / "helix-design-studio-web")
    parser.add_argument("--results", type=Path, default=ROOT / "results")
    args = parser.parse_args()
    repo, results = args.repo.resolve(), args.results.resolve()
    commit = subprocess.check_output(["git", "rev-parse", "HEAD"], cwd=repo, text=True).strip()
    status = subprocess.check_output(["git", "status", "--porcelain"], cwd=repo, text=True).strip()
    if status:
        raise SystemExit("Parameter exports require an identified clean source revision")
    software_version = version(repo)
    # Load definitions without constructing a builder or creating calibration caches.
    sys.path.insert(0, str(repo / "Code"))
    model = runpy.run_path(str(repo / "Code/core/hamiltonian_builder.py"))
    convention = model["NEIGHBOR_SCORING_CONVENTION"]
    neighbor_weights = model["NEIGHBOR_MATRIX_WEIGHTS"]
    if convention != "negative_m1_plus_m3_plus_m4" or neighbor_weights != {1: -1.0, 3: 1.0, 4: 1.0}:
        raise SystemExit("This table export requires the identified negative-M1 scoring convention")
    parameter_fingerprint = model["parameter_set_sha256"]()
    tool_path = repo / "tools/verify_parameter_sources.py"
    source_dir = repo / "tools/data/parameter_sources"
    verification = runpy.run_path(str(tool_path))["verify"](repo / "Code/core/resources", source_dir)
    if not verification["verified"]:
        raise SystemExit("Parameter tables differ from the independently transcribed sources")
    descriptions = json.loads((source_dir / "metadata.json").read_text(encoding="utf-8"))
    matrices = {}
    for name, description in descriptions["matrices"].items():
        path = repo / "Code/core/resources" / description["runtime_file"]
        symbols, tokens = resource_table(path)
        matrices[name] = {
            "resource": path.relative_to(repo).as_posix(), "sha256": sha256(path),
            "shape": [20, 20], "sequence_separation": description["runtime_separation"],
            "source_sequence_separation": description["source_separation"],
            "row_residues": symbols, "column_residues": symbols,
            "indexing": "row = residue at i; column = residue at i+k",
            "numeric_tokens": tokens, "units_of_source_values": description["units"],
            "internal_scoring_coefficient": neighbor_weights[description["runtime_separation"]],
            "primary_reference": description["reference"],
            "source_mapping": description["mapping"], "source_axes": description["source_axes"],
            "model_adaptation": description.get("model_adaptation"),
            "missing_value_policy": description.get("missing_value_policy"),
            "verification": verification["matrices"][name],
        }
    results.mkdir(parents=True, exist_ok=True)
    pdf_path = results / "parameter_tables.pdf"
    make_pdf(pdf_path, matrices, commit, software_version)
    record = {
        "status": "primary_source_values_verified_with_documented_model_adaptations",
        "status_date": datetime.now(timezone.utc).date().isoformat(),
        "software_version": software_version, "software_commit": commit,
        "parameters_sha256": parameter_fingerprint,
        "parameter_fingerprint_scope": "Source matrices, per-residue scales and the neighbor-scoring convention with its internal coefficients",
        "neighbor_score_convention": convention,
        "neighbor_matrix_weights": {f"M{k}": value for k, value in neighbor_weights.items()},
        "neighbor_score_formula": "-mean(M1) + mean(M3) + mean(M4)",
        "bibliographic_reference": {
            "status": "primary_sources_identified_and_numerically_verified",
            "independent_bibliographic_verification_claimed": True,
            "verification_scope": "Exact printed coefficients mapped by independently labelled source axes; model adaptations are recorded separately.",
        },
        "application": "Raw source coefficients are preserved. Average each matrix over its L-k pairs, subtract the M1 mean and add the M3 and M4 means, then standardize the combined empirical neighbour component before applying its configured component weight. The internal coefficients -1,+1,+1 fix the relative numerical scaling of heterogeneous source quantities; the result is a comparative descriptor. Nacar's hydrogen-bonded n,n+4 enrichments are reused empirically at k=1, with greater enrichment lowering the minimized score. A zero M1 source value contributes zero to its unstandardized sum.",
        "matrices": matrices,
        "verification_tool": {"file": tool_path.relative_to(repo).as_posix(), "sha256": sha256(tool_path)},
        "source_metadata_sha256": sha256(source_dir / "metadata.json"),
        "export_runtime": {"python": platform.python_version(), "reportlab": importlib.metadata.version("reportlab")},
        "table_export": {"file": f"{results.name}/parameter_tables.pdf", "sha256": sha256(pdf_path), "numeric_tokens": 1200,
                         "published_coefficients": sum(item["published_cells"] for item in verification["matrices"].values()),
                         "zero_extension_coefficients": sum(item["model_zero_extension_cells"] for item in verification["matrices"].values())},
        "computation_status": "Parameter records identify this corrected software revision. Reported numerical outputs must be obtained from a separate study of the same commit and table hashes.",
    }
    output = results / "parameter_source_checks.json"
    output.write_text(json.dumps(record, indent=2, ensure_ascii=False, allow_nan=False) + "\n", encoding="utf-8")
    print(json.dumps({"software_version": software_version, "software_commit": commit,
                      "parameters_sha256": parameter_fingerprint, "neighbor_score_convention": convention,
                      "pdf": str(pdf_path), "json": str(output), "verified_published_coefficients": record["table_export"]["published_coefficients"],
                      "documented_zero_extension_coefficients": record["table_export"]["zero_extension_coefficients"]}, indent=2))


if __name__ == "__main__":
    main()
```
