## Additional file 2. Computational evaluation, inputs and results for "Helix Design Studio: designing antimicrobial peptide candidates for membrane environments": README.md

# Reproducible software evaluation

This study evaluates **Helix Design Studio 1.0.0rc6**, commit `d6ccec1920f256a393b15215df63fd2dead61cd5`, on **17 October 2026**. The neighbour tables were checked against their primary sources: Nacar (2020), Figure 3, for M1, and Muñoz and Serrano (1994), Table 2, for M3/M4. Source transcriptions retain independent row and column labels in the source repository. M1 is transposed to preserve sequence direction; its source observations concern hydrogen-bonded n,n+4 pairs, and the model explicitly reuses these coefficients at i,i+1. The 18×18 M3/M4 tables are extended to 20 residues with 76 zero-contribution cells each for pairs containing C or P. These extensions are model conventions. `results/parameter_source_checks.json` records the exact source comparison, all 1,200 runtime entries, labels, adaptations and checksums; `results/parameter_tables.pdf` reproduces the complete tables. Numerical source agreement does not validate the model adaptations or biological activity.

## Run and preserve the study

Install the source repository's pinned development requirements, then run from the submission directory:

```bash
python experiments/run_examples.py --repo ../helix-design-studio-web --output results
python experiments/export_parameter_tables.py --repo ../helix-design-studio-web --results results
python experiments/summarize_results.py --results results
```

Use a new `--output` directory when `results/` already contains a completed study. `--plan-only` writes explicit request templates without computing results; the completed-output guard also applies in this mode. All requests run locally through FastAPI TestClient. The runner records the source commit, Git state, source/table hashes and package versions, and checks that the source did not change during calculation. Missing optional hardware or Git commands produce unavailable metadata. Use the exact Git checkout for a new manuscript build, whose source-identity checks require an identified revision.

The parameter exporter requires ReportLab and a clean identified source revision, verifies the runtime files against the distributed source CSVs, and produces the matrix PDF and comparison record. The standard-library summarizer validates completed outputs and regenerates `summary.json`, `experiments/RESULTS_NOTES.md` and `reproduction_manifest.json`. Run it after final source/document edits. The manifest includes the available evaluation and manuscript scripts; diagnostic `runtime.log` and bytecode caches are excluded from the distribution.

## Sequences and search protocols

Three hand-specified 20-residue probes demonstrate contrasting compositions and residue patterns. No experimental activity or preferred conformation is asserted for them. The study requests four family members, two specificity candidates and one compromise design, retains every returned sequence, and scores the ten illustrated sequences under all five environment presets.

All main search examples use **2000 component-calibration sequences**, including Specificity, seed 42, the ordered alphabet `A,R,N,D,C,Q,E,G,H,I,L,K,M,F,P,S,T,W,Y,V`, and interfacial apolar-sector half-width **90°**. These are explicit experimental settings. The browser's **Advanced settings** expose calibration sample size, random seed and an exact apolar-sector half-width override. Specificity's default calibration count is 10000; set it to 2000 to reproduce this example. The standard 50% exposure slider setting maps to 95°, so use the exact-angle field for 90°. Specificity shares that override across its target and interfacial off-targets; Compare/Cross-Design have separate fields for their two environments.

The family request uses the anionic interface, 3,000 steps per restart, two restarts per candidate attempt, four members and `oversample_factor=2`. The attempt budget is at least four times the requested family size; collection stops once enough unique members are found. Diversity is encouraged by a soft penalty, not guaranteed by a hard distance constraint. Reported Hamming distances describe the actual collection.

Specificity uses length 20, 3,000 steps, four **total** restarts shared across attempts, two requested candidates, λ = 0.5, an anionic target and neutral-interface/homogeneous-polar off-targets. Unsuccessful attempts consume the restart budget. Compromise design uses length 20, 3,000 steps, two restarts and γ = 0.5 for the anionic-interface and homogeneous-polar environments.

All searches rotate each proposed interfacial sequence about its helical axis so that its transverse hydrophobic moment points toward the apolar sector. `native_design_scores.csv` extracts the aligned design scores. Specificity and compromise objectives are reconstructed from those same aligned model scores. The compatibility field `design_geometry` duplicates the parent-level score values in this release.

## Standard scores, backgrounds and random streams

Every row in `scores.csv` uses the same ordered 20-residue alphabet, six default weights, 2,000 calibration decoys, 1,000 reference sequences and seed 42. At interfaces, the standard score aligns **each** query, calibration sequence and reference sequence to its own hydrophobic moment. Design responses use these same alignment conventions. A single calibrated model and reference ensemble is reused for search and result reporting; `design_geometry` is a compatibility copy of the reported score.

Native Design, Family, Cross-Design and Compare outputs use 500 references; Specificity uses 3,000. Common rescoring uses 1,000. Differences in reference size can therefore change a Z-score even when the orientation rule is the same. The model score H is dimensionless.

The seed protocol uses separate NumPy SeedSequence entropy inputs:

- Calibration decoys: `[seed]`.
- Reference ensemble: `[seed, 1]`.
- Single search: `[seed, 2]`.
- Family/Specificity search run i: `[seed, 2, i]`.

The full request, software version, parameter checksum and seed protocol are retained in response provenance. Environments are evaluated with the same declared base seed; their different Hamiltonians still produce different distributions. The neighbour component is **−mean(M1) + mean(M3) + mean(M4)**, with each matrix averaged over the available pairs at its own separation, then the combined component standardized. The 400 published M1 coefficients remain unchanged, including zeros. Greater M1 propensity lowers this raw component through the signed linear combination. The scoring-convention identifier and matrix weights are included in parameter and calibration-cache fingerprints. Miyazawa–Jernigan contacts cover sequence separations one through four.

## Exposure and sampling sensitivity

The exposure scan evaluates 21 settings from 0 to 100 in increments of five. All points use the same supplied seed and the same separate calibration/reference streams. Homogeneous polar and apolar presets are used at the endpoints; intermediate values alter the interfacial angular mask. The reported minimum is conditional on this model geometry and finite background samples. The percentage is not a physical insertion depth. The full profile and native response are retained. Idealized PDB output represents prescribed alpha-helical geometry, not a folding prediction.

The sampling-sensitivity check scores the same amphiphilic probe at the anionic interface with seeds 42–44 and the four combinations of 500/2,000 calibration decoys and 500/2,000 reference sequences. All 12 outputs are retained. Changing the seed changes both sampled backgrounds; this fixed-sequence experiment contains no search noise and does not estimate general model uncertainty.

## Timing interpretation

Timing requests cover lengths 8, 16 and 24 and seeds 42, 43 and 44. Each cold request starts with an empty dedicated calibration cache and is immediately repeated with identical payload and populated cache. These are calibration-cache measurements, not operating-system cold starts. Python, FastAPI and the application are already initialized, requests run sequentially, and unrelated machine processes are not suspended.

Score and Design each build one aligned calibration model. Warm requests reuse these calibration statistics but still calculate reference scores and, for Design, run the search. The timing comparison includes search and reporting work; it is not a measurement of the optimizer alone.

Wall time includes in-process request handling and JSON serialization. It excludes HTTP-server startup, network transport, deployment queues and browser rendering. Report medians and observed ranges across the three seeds, not confidence intervals or service-capacity claims. Score uses 1,000 references; Design uses its native 500-reference protocol, 3,000 steps and two restarts.

The recorded system is **Apple M3 Max, 16 logical CPUs, 48 GiB RAM, macOS 26.6.2, Python 3.12.8**. `metadata.json` preserves the measured hardware and package versions. At length 24, median Score times were 114.7 ms with calibration computed and 46.0 ms with calibration reused; Design times were 372.4 and 293.8 ms. These timings describe this sequential benchmark.

## Exact optimization check and evidence files

The exact benchmark enumerates all 4,096 sequences of length six over A/E/K/L under the default anionic-interface environment with per-sequence hydrophobic-moment alignment, default weights, 2,000 calibration decoys and calibration seed 42. Five annealing searches use this same objective, seeds 42–46, 1,000 steps and two restarts. `exact_search_summary.json` records the minimizers, each score gap and recovery count within absolute tolerance 1e-9. The result checks search behavior for that finite objective; it establishes neither the physical validity of the score nor global optimality for longer designs.

The unique minimum is **LLKKLL**, H = **−24.555206117239166**, recovered in **5/5** searches. The evaluated revision also passed **48 Python tests and 29 frontend tests**; the exact commands and output hashes are recorded in `software_verification.json`.

`responses/` preserves requests and complete responses. `calls.json` records timing and calibration-cache identities. `calibration_statistics.json` records calibration metadata and component statistics. `metadata.json` captures hardware, package versions, source hashes, seeds, timestamps and repeated-request checks. CSV files supply the figures and tables. `software_verification.json` links the separate Python/frontend test results to the evaluated revision. The independent modlAMP comparison has its own environment and provenance record; HELIQUEST is a documentary reference, not an executed live-server comparator.

## Primary parameter sources

- Nacar C. Propensities of Amino Acid Pairings in Secondary Structure of Globular Proteins. Protein J 39:21–32 (2020), Figure 3, p.25. [doi:10.1007/s10930-020-09880-6](https://doi.org/10.1007/s10930-020-09880-6). Figure columns identify the earlier residue and rows the later residue. The published proline-donor zeros are retained in the corresponding runtime column.
- Muñoz V, Serrano L. Elucidating the folding problem of helical peptides using empirical parameters. Nat Struct Biol 1:399–409 (1994), Table 2, p.403. [doi:10.1038/nsb0694-399](https://doi.org/10.1038/nsb0694-399). The source has different F/Y row and column orders; both axes are mapped independently. Source energies retain their kcal/mol numerical values before joint neighbour-term standardization.

`parameter_correction_comparison.json` evaluates four numerical-table conditions with this same rc6 engine and scoring convention: initial tables, updated M1 only, updated M4 only, and both updated. The report identifies the initial table hashes and the current engine hashes separately. It is a table intervention, not a run of an older complete release. Across 60 paired scores, updating both tables gives median/max |ΔZH| **0.108390/0.244802** and Spearman correlation **0.995276**; the six paired searches differ at **60/108** positions.

## Paired M1 scoring-convention comparison

`compare_m1_scoring.py` changes only the sign with which M1 enters the neighbour component, keeping all source-verified matrix values and the current engine fixed. The positive-M1 condition is a process-local experimental control; the current condition uses −M1+M3+M4. Both conditions use their own freshly computed calibration, with identical calibration and reference sequences, identified by hashes. Independent temporary cache directories and distinct convention fingerprints prevent statistics from being shared between conditions.

```bash
python experiments/compare_m1_scoring.py --repo ../helix-design-studio-web --output results/m1_scoring_comparison.json
```

Inputs are read from the existing `tools/measure_changes.py` benchmark: 12 reference sequences, five environments, 1000 calibration sequences, 500 references and scoring seed 42; six paired length-18 designs use seeds 1, 2 and 3 in anionic-interface and homogeneous-polar environments, 4000 steps and two restarts. The source-verified tables and per-sequence alignment are common to both conditions. The report retains all 60 score pairs, 12 native design records, six paired sequence comparisons, complete calibration/reference statistics, exact software version and source hashes. Changes are defined as **negative-M1 current minus positive-M1 control**; condition-specific native scores retain their respective background normalizations.

The resulting median/max |ΔH| are **2.709000/23.327558** and median/max |ΔZH| are **0.362826/3.438784**. Spearman correlations are **0.816004** for H and **0.817394** for ZH. All six designed sequences differ, with **76/108 positions changed (70.37%)** and paired Hamming distances 8–18. The largest score change occurs for poly-Ala15 in homogeneous polar medium: ZH changes from +0.694854 to −2.743931. These describe effects within the model, rather than experimental improvements in peptides.

## Paired orientation comparison

`compare_search_orientation.py` runs fixed and aligned search variants on the same clean runtime with the same parameter set, environments, budgets and three seeds. The fixed variant is a controlled counterfactual; it is not an execution of an older complete software release. Six subprocesses preserve all outputs and a separate process reevaluates every candidate with one aligned protocol (2,000 calibration sequences, 1,000 references and seed 42). Search acceptance can cause random streams to diverge, so matching seeds does not imply identical mutation proposals. Runtime records include calibration and reporting and are not pure optimizer timings.

```bash
python experiments/compare_search_orientation.py --repo ../helix-design-studio-web --output results/search_orientation_comparison
python experiments/validate_search_alignment.py --repo ../helix-design-studio-web --output results/independent_alignment_audit.json
```

The paired comparison retains three seeds per workflow and every returned candidate. Outcomes are reported by workflow, without treating shared candidates or workflows as independent statistical replicates. The independent alignment audit calculates phases and environmental exposure for 81 sequences directly and instruments all four design workflows. Neither check validates biological activity or equilibrium peptide orientation.

The rc6 comparison contains 48 candidate records, 21 distinct sequences per orientation rule, with no sequence shared between rules across the tested workflows and seeds. Best candidates from matched searches differ at 6–20 of 20 positions. Aligned search gives the lower common best objective in **3/3 seeds for each of Design, Family, Specificity and Cross-Design**. The independent audit passed, including 8717 observed aligned evaluations and exact reconstruction of Cross-Design and Specificity objectives from their displayed scores.

The fixed-orientation Specificity counterfactual preserves the `margins_geometry` text emitted by the current runtime. That inherited text is not used in the comparison: the explicit experimental mode, each nested `design_geometry.orientation`, and the recorded search energies identify the fixed condition. Numerical margins and objectives are reconstructed from those energies; the original responses are retained unchanged.

## Manuscript figures and export provenance

The manuscript contains **seven main figures**: concept, architecture, environmental profiles, candidate diversity and scores, exposure and sampling sensitivity, computation times, and transverse hydrophobic-moment alignment. The conceptual Figure 1 and alignment Figure 7 are retained as manuscript artwork; the numerical panels use the preserved evaluation records.

Build the standalone LaTeX before exporting its artwork:

```bash
python source/build_manuscript.py
python source/make_figures.py
python experiments/summarize_results.py
```

The second command runs `source/export_manuscript_figures.py` and then `source/make_graphical_abstract.py`. To regenerate only the seven main figures from an existing final LaTeX, run `python source/export_manuscript_figures.py` without arguments. It compiles the exact inline drawings with the original manuscript preamble, crops the vector PDFs and renders PNGs at **300 dpi**. `figures/FIGURE_CAPTIONS.md` retains each caption verbatim. The main-figure inventory is **seven PDFs plus seven PNGs**; the graphical abstract is a separate 920 × 300 pixel schematic.

`results/figure_export_manifest.json` records the manuscript SHA-256, figure order and labels, artwork/caption hashes, output hashes and dimensions. It is included in the reproduction manifest. Package creation rejects stale exports or an inventory other than seven matched PDF/PNG pairs. The active manuscript sources include `manuscript_references.tex`, `transverse_alignment_figure.tex`, the builder and both exporters; the seven figures are exported from the complete manuscript rather than separately reconstructed illustrations.

Export requires `pdflatex` with TikZ/PGFPlots, `pdfcrop` with Ghostscript, Poppler's `pdftoppm`, and `pypdf`. The builder and separate graphical abstract use Matplotlib. See `REPRODUCE.md` for the installation and regeneration order.
