## Additional file 2. Computational evaluation, inputs and results for "Helix Design Studio: designing antimicrobial peptide candidates for membrane environments": RESULTS_NOTES.md

# Measured results for the release-candidate draft

All values below were produced by the completed run of `run_examples.py`, using commit `d6ccec1920f256a393b15215df63fd2dead61cd5`. Initial Git status: `clean`; final Git status: `clean`. Runtime source hashes were identical before and after the run. The numerical parameter dataset and its checksums are recorded in `results/parameter_source_checks.json`; the neighbour matrices are also reproduced in `results/parameter_tables.pdf`. These calculations characterize the implemented model; they do not establish biological activity.

The run performed 104 successful local API requests in 13.443 s. It retained 50 common-protocol scores, 12 native aligned design scores, 39 distinct calibration records, 21 profile points, 12 reference-sampling sensitivity measurements, and 4,096 exhaustive-reference energies. All 18 identical payload repeats returned identical JSON with populated calibration caches. The run created no usage log.

## Example observations

For `LKKLLKLLKKLLKLLKKLLK`, common-protocol reference Z scores were -6.536 (anionic interface), -4.752 (neutral interface), -3.829 (cationic interface), 3.352 (homogeneous polar), 2.897 (homogeneous apolar). For the acidic analogue `LEELLELLEELLELLEELLE`, the anionic, neutral, and cationic interface values were -2.986, -4.073, -6.251. Poly-leucine had its lowest value in the homogeneous apolar preset (-6.443). These are illustrative responses of the implemented objective, not experimental validation.

The family returned 4 members, with aligned model scores from -78.453 to -75.347 and pairwise Hamming distances from 10 to 20. The lowest-native-score sequence was `RERRWWKRWWKEWRKWEKKW`. The W/K/R-rich outputs illustrate that this unconstrained objective can push composition toward extreme residues; no synthesis, toxicity, activity, aggregation, or developability filter is implied. Common rescoring in `scores.csv` uses the same alignment rule; its reference count can differ from that of the native design result.

Specificity candidates had native target-versus-lowest-off-target score margins of 28.191, 25.275, and native objectives of -54.217, -50.453. Both specificity objectives and the cross-design objective (-136.205) were reconstructed from their returned native component scores. These margins are normalized model-score differences, not physical transfer free energies or experimental selectivity measurements.

The scan minimum occurred at the displayed 55% setting (Z=-7.148, score=-63.466), with interfacial half-width 100.5 degrees. For interior percentage settings p, half-width = 40 + 1.1p degrees; endpoints 0 and 100 are replaced by homogeneous polar/apolar presets. The percentage is a geometric index, not measured membrane insertion. The same seed protocol is used across scan points; this minimum has no experimental confidence interval.

## Exact reference and finite sampling

At length 6 and alphabet A/E/K/L, exhaustive enumeration found 1 minimum sequence(s): LLKKLL, with score -24.555206117239166. 5/5 annealing searches recovered it within absolute tolerance 1e-9. This verifies one finite optimization instance of the same objective; it does not establish biological validity or global optimality for larger examples.

Observed reference Z-score ranges across the three sensitivity seeds were:

- 500 calibration decoys / 500 references: -7.014 to -6.342.
- 500 calibration decoys / 2,000 references: -6.760 to -6.391.
- 2,000 calibration decoys / 500 references: -6.736 to -6.416.
- 2,000 calibration decoys / 2,000 references: -6.497 to -6.460.

These are sampling ranges for one sequence, not biological prediction uncertainty or inferential confidence intervals.

## Timing measurements

Recorded hardware: Apple M3 Max; 16 logical CPUs; 48 GiB RAM. In-process API wall-time medians over three seeds:

| Length | Score, empty calibration cache | Score, populated cache | Design, empty calibration cache | Design, populated cache |
|---:|---:|---:|---:|---:|
| 8 | 0.051 s | 0.019 s | 0.138 s | 0.124 s |
| 16 | 0.083 s | 0.028 s | 0.263 s | 0.210 s |
| 24 | 0.115 s | 0.046 s | 0.372 s | 0.294 s |

Score requests used 2,000 calibration decoys and 1,000 reference sequences; design requests used 3,000 steps and two restarts with the native 500-reference output. Imports had already completed. Values exclude network transport, queueing, server startup, and browser rendering. Observed ranges are in `timing_summary.csv`.

Single examples took 0.981 s (family), 1.625 s (specificity), 0.506 s (cross-design), and 1.949 s (21-point scan). These are one measurement each, unlike the three-seed size benchmark.

The manifest hashes all result files except itself and `runtime.log`, and available experiment/manuscript sources. Regenerate it with `python experiments/summarize_results.py` after the final source edits.
