## Additional file 2. Computational evaluation, inputs and results for "Helix Design Studio: designing antimicrobial peptide candidates for membrane environments": run_examples.py

```
#!/usr/bin/env python3
"""Reproduce manuscript examples through the public FastAPI request models.

No requests are sent to the deployed service. See README.md for interpretation.
Requires the application requirements and httpx (requirements-dev.txt).
"""
from __future__ import annotations

import argparse
import ast
from contextlib import redirect_stdout
import csv
from datetime import datetime, timezone
import hashlib
import importlib.metadata
import io
import itertools
import json
import math
import os
from pathlib import Path
import platform
import statistics
import subprocess
import sys
import tempfile
import time


ALPHABET = list("ARNDCQEGHILKMFPSTWYV")
PRESETS = ["homogeneous_polar", "homogeneous_apolar", "interfacial_neg", "interfacial_neu", "interfacial_pos"]
WEIGHTS = {"lambda_env": 6.0, "lambda_charge": 3.0, "lambda_local": 1.5,
           "lambda_pairwise": 0.1, "lambda_helix_pairs": 1.5, "lambda_electrostatic": 4.0}
# Hand-specified demonstration inputs, not experimentally characterized peptides.
PROBES = {
    "probe_amphiphilic": "LKKLLKLLKKLLKLLKKLLK",
    "probe_hydrophobic": "LLLLLLLLLLLLLLLLLLLL",
    "probe_acidic": "LEELLELLEELLELLEELLE",
}
assert all(len(sequence) == 20 for sequence in PROBES.values())
COMMON = {"residues": ALPHABET, "weights": WEIGHTS, "n_decoys": 2000, "seed": 42}


def dump(path: Path, value):
    path.parent.mkdir(parents=True, exist_ok=True)
    path.write_text(json.dumps(value, indent=2, sort_keys=True, allow_nan=False) + "\n", encoding="utf-8")


def csv_write(path: Path, rows):
    if not rows:
        return
    with path.open("w", newline="", encoding="utf-8") as handle:
        writer = csv.DictWriter(handle, fieldnames=list(rows[0]))
        writer.writeheader()
        writer.writerows(rows)


def command(*args):
    try:
        result = subprocess.run(args, capture_output=True, text=True, check=False)
    except OSError:
        # Optional metadata tools (for example sysctl or Git) may be absent.
        # FileNotFoundError is an OSError subclass.
        return None
    return result.stdout.strip() if result.returncode == 0 else None


def source_metadata(repo: Path):
    paths = [*repo.glob("app/*.py"), *repo.glob("Code/**/*.py"), *repo.glob("Code/**/*.txt"), repo / "requirements.txt"]
    hashes = {str(path.relative_to(repo)): hashlib.sha256(path.read_bytes()).hexdigest()
              for path in sorted(paths) if path.is_file()}
    return {
        "repository": "https://github.com/SIMBIOS-USC/helix-design-studio-web",
        "commit": command("git", "-C", str(repo), "rev-parse", "HEAD"),
        "git_status": command("git", "-C", str(repo), "status", "--porcelain"),
        "runtime_source_sha256": hashes,
        "runtime_source_manifest_sha256": hashlib.sha256(json.dumps(hashes, sort_keys=True).encode()).hexdigest(),
    }


def application_version(repo: Path) -> str:
    """Read the declared application version without importing runtime state."""
    for node in ast.parse((repo / "app/main.py").read_text(encoding="utf-8")).body:
        if isinstance(node, ast.Assign) and any(isinstance(target, ast.Name) and target.id == "APP_VERSION" for target in node.targets):
            value = ast.literal_eval(node.value)
            if isinstance(value, str) and value:
                return value
    raise ValueError("Application APP_VERSION must be a nonempty string literal")


def finite(value):
    if isinstance(value, dict):
        return all(finite(child) for child in value.values())
    if isinstance(value, list):
        return all(finite(child) for child in value)
    if isinstance(value, (int, float)):
        return math.isfinite(value)
    return True


def main():
    parser = argparse.ArgumentParser(description=__doc__)
    parser.add_argument("--repo", type=Path, default=Path(__file__).resolve().parents[2] / "helix-design-studio-web")
    parser.add_argument("--output", type=Path, default=Path(__file__).resolve().parents[1] / "results")
    parser.add_argument("--plan-only", action="store_true", help="Write explicit request templates without running calculations.")
    args = parser.parse_args()
    repo, output = args.repo.resolve(), args.output.resolve()
    if (output / "metadata.json").exists():
        raise SystemExit("Refusing to overwrite an existing completed experiment. Choose a new --output directory.")
    output.mkdir(parents=True, exist_ok=True)
    templates = {
        "score": {**COMMON, "sequence": PROBES["probe_amphiphilic"], "environment": {"preset": "interfacial_neg"}, "n_random": 1000},
        "family": {**COMMON, "length": 20, "environment": {"preset": "interfacial_neg"}, "steps": 3000,
                   "restarts": 2, "family_size": 4, "oversample_factor": 2},
        "specificity": {**COMMON, "length": 20, "target_environment": {"preset": "interfacial_neg"},
                        "off_target_environments": [{"preset": "interfacial_neu"}, {"preset": "homogeneous_polar"}],
                        "steps": 3000, "restarts": 4, "num_sequences": 2, "lambda_balance": 0.5},
        "compare": {**COMMON, "sequence": PROBES["probe_amphiphilic"],
                    "environment_a": {"preset": "interfacial_neg"}, "environment_b": {"preset": "homogeneous_polar"}},
        "cross_design": {**COMMON, "length": 20, "steps": 3000, "restarts": 2, "lambda_gap": 0.5,
                         "environment_a": {"preset": "interfacial_neg"}, "environment_b": {"preset": "homogeneous_polar"}},
        "penetration": {**COMMON, "sequence": PROBES["probe_amphiphilic"], "environment": {"preset": "interfacial_neg"},
                        "n_random": 1000, "percent_min": 0, "percent_max": 100, "percent_step": 5},
    }
    dump(output / "request_templates.json", templates)
    dump(output / "probe_sequences.json", {"description": "Hand-specified synthetic demonstration inputs; no biological activity is asserted.", "sequences": PROBES})
    if args.plan_only:
        print("Request templates written; no scientific calculation performed.")
        return
    start = time.perf_counter()
    metadata = {
        "started_utc": datetime.now(timezone.utc).isoformat(),
        "parameter_mapping_status": "resource_labels_define_the_evaluated_parameterization",
        "publication_status": "Reproducible evaluation of the identified software revision.",
        "parameter_mapping_limitation": "The run identifies parameter files by labels and checksums; independent bibliographic verification is not performed by this runner.",
        "source": source_metadata(repo),
        "python": sys.version,
        "platform": platform.platform(),
        "architecture": platform.machine(),
        "cpu": command("sysctl", "-n", "machdep.cpu.brand_string") or platform.processor(),
        "logical_cpus": os.cpu_count(),
        "memory_bytes": command("sysctl", "-n", "hw.memsize"),
        "packages": {dist.metadata["Name"]: dist.version for dist in importlib.metadata.distributions()},
        "protocol": {"alphabet": ALPHABET, "weights": WEIGHTS, "n_decoys": 2000,
                     "version": application_version(repo), "search_orientation": "own hydrophobic moment in interfaces, applied to proposals, calibration decoys and references",
                     "standard_score_orientation": "own hydrophobic moment in interfaces, applied to decoys and references too",
                     "mj_max_interaction_distance": 4,
                     "common_rescoring_n_random": 1000, "common_rescoring_seed": 42,
                     "benchmark_seeds": [42, 43, 44], "benchmark_lengths": [8, 16, 24],
                     "benchmark_steps": 3000, "benchmark_restarts": 2,
                     "measurement": "In-process FastAPI TestClient wall time; excludes network/browser; application and imports already initialized.",
                     "cache_states": "Each cold request starts with an empty dedicated calibration cache. Warm repeats the identical request immediately, without clearing that cache.",
                     "concurrency_control": "Sequential requests, no deliberate competing benchmark workload. Other machine processes were not suspended."},
    }
    calls = []
    score_rows = []
    calibration_records = {}
    runtime_rows = []
    with tempfile.TemporaryDirectory(prefix="helix-manuscript-") as temporary:
        cache = Path(temporary) / "cache"
        cache.mkdir()
        os.environ["HELIX_CACHE_DIR"] = str(cache)
        os.environ["HELIX_VAR_DIR"] = str(Path(temporary) / "var")
        os.environ["HELIX_USAGE_LOGGING"] = "0"
        os.environ["QFOLD_CODE_DIR"] = str(repo / "Code")
        sys.path.insert(0, str(repo))
        from fastapi.testclient import TestClient
        from app.main import app
        from app import engine

        # A publication run must not proceed with the known label-indexing defect.
        sequence = "APWYEDLKRQNASVITMHCG"
        raw_values = []
        with redirect_stdout(io.StringIO()):
            for alphabet in (ALPHABET, list(reversed(ALPHABET))):
                builder = engine.build_builder(len(sequence), alphabet, engine.resolve_environment({"preset": "interfacial_neg"}), n_decoys=100)
                raw_values.append(builder._raw_helix_neigh(engine.seq_to_codes(sequence, alphabet)))
        if not math.isclose(*raw_values, rel_tol=1e-12, abs_tol=1e-12):
            raise RuntimeError(f"Matrix alphabet-order invariance failed: {raw_values}. Correct the implementation before publication experiments.")
        for path in cache.glob("zscore_stats_*.json"):
            path.unlink()
        metadata["matrix_order_guard"] = {"sequence": sequence, "raw_neighbor_scores": raw_values, "passed": True}

        with TestClient(app) as client, (output / "runtime.log").open("w", encoding="utf-8") as log:
            def capture_calibration():
                for path in cache.glob("zscore_stats_*.json"):
                    calibration_records[path.stem] = json.loads(path.read_text())

            def post(name, route, payload):
                before = sorted(path.name for path in cache.glob("zscore_stats_*.json"))
                call_start = time.perf_counter()
                with redirect_stdout(log):
                    response = client.post(route, json=payload)
                elapsed = time.perf_counter() - call_start
                if response.status_code != 200:
                    raise RuntimeError(f"{name}: HTTP {response.status_code}: {response.text}")
                result = response.json()
                if not finite(result):
                    raise RuntimeError(f"{name}: nonfinite result")
                after = sorted(path.name for path in cache.glob("zscore_stats_*.json"))
                record = {"name": name, "route": route, "request": payload, "wall_seconds": elapsed,
                          "calibration_files_before": before, "calibration_files_after": after,
                          "new_calibration_files": sorted(set(after) - set(before))}
                dump(output / "responses" / f"{name}.json", {**record, "response": result})
                calls.append(record)
                capture_calibration()
                print(f"{name}: {elapsed:.3f} s", flush=True)
                return result, elapsed

            def add_score_rows(label, origin, result, protocol, design_geometry=False):
                score = result["design_geometry"] if design_geometry else result
                if design_geometry:
                    # The compatibility alias must describe the objective actually
                    # optimized, which now has the same orientation as Score.
                    for key in ("energy", "z_score", "percentile_lower_is_better"):
                        if not math.isclose(score[key], result[key], rel_tol=0, abs_tol=1e-12):
                            raise RuntimeError(f"Search/Score mismatch for {label}: {key}")
                    expected_orientation = result["calibration"]["orientation"]
                    if (expected_orientation == "own hydrophobic moment" and
                            not result["environment"].get("phase_aligned_to_hydrophobic_moment", True)):
                        expected_orientation = "preset wheel phase (vanishing hydrophobic moment)"
                    if score["orientation"] != expected_orientation:
                        raise RuntimeError(f"Search/Score orientation mismatch for {label}")
                phase = score["wheel_phase_deg"] if design_geometry else result["environment"]["wheel_phase_deg"]
                orientation = score["orientation"] if design_geometry else result["calibration"]["orientation"]
                row = {"id": label, "origin": origin, "sequence": result["sequence"], "length": result["length"],
                       "environment": result["environment"]["preset"], "energy": score["energy"], "z_score": score["z_score"],
                       "percentile_lower_is_better": score["percentile_lower_is_better"],
                       "model_net_charge": result["model_net_charge"],
                       "hydrophobic_moment_per_residue": result["observables"]["abs_mu_h_per_residue"],
                       "phase_deg": phase,
                       "n_random": result["random_reference"]["n"],
                       "reference_seed": json.dumps(result["random_reference"]["seed_entropy"]),
                       "n_decoys": result["calibration"]["n_decoys"],
                       "calibration_seed": result["calibration"]["seed_entropy"],
                       "orientation_mode": orientation, "protocol": protocol}
                row.update({f"term_{key}": value for key, value in score["breakdown"].items()})
                score_rows.append(row)

            # All plotted environment profiles use one explicit evaluation protocol.
            all_sequences = [(name, "hand_specified", sequence) for name, sequence in PROBES.items()]
            family, _ = post("family", "/api/design-family", templates["family"])
            specificity, _ = post("specificity", "/api/specificity-design", templates["specificity"])
            cross, _ = post("cross_design", "/api/cross-design", templates["cross_design"])
            for i, member in enumerate(family["members"], 1):
                all_sequences.append((f"family_{i}", "family_design", member["sequence"]))
                add_score_rows(f"family_{i}", "family_design", member, "native_aligned_design_2000_500", True)
            for i, member in enumerate(specificity["candidates"], 1):
                all_sequences.append((f"specificity_{i}", "specificity_design", member["sequence"]))
                for score in [member["target_environment"], *member["off_target_environments"]]:
                    add_score_rows(f"specificity_{i}", "specificity_design", score, "native_aligned_design_2000_3000", True)
                data = member["specificity_design"]
                reconstructed = member["target_environment"]["design_geometry"]["energy"] + (data["lambda_balance"] - 1) * min(score["design_geometry"]["energy"] for score in member["off_target_environments"])
                if not math.isclose(data["objective"], reconstructed, rel_tol=0, abs_tol=1e-8):
                    raise RuntimeError("Native specificity objective is inconsistent with returned search scores")
            all_sequences.append(("cross_1", "cross_design", cross["sequence"]))
            for score in [cross["environment_a"], cross["environment_b"]]:
                add_score_rows("cross_1", "cross_design", score, "native_aligned_design_2000_500", True)
            a, b = cross["environment_a"]["design_geometry"]["energy"], cross["environment_b"]["design_geometry"]["energy"]
            if not math.isclose(cross["cross_design"]["objective"], a + b + 0.5 * abs(a - b), rel_tol=0, abs_tol=1e-8):
                raise RuntimeError("Native cross-design objective is inconsistent with returned search scores")
            csv_write(output / "native_design_scores.csv", score_rows)
            score_rows.clear()
            for label, origin, sequence in all_sequences:
                for preset in PRESETS:
                    scored, _ = post(f"score_{label}_{preset}", "/api/score", {
                        **COMMON, "sequence": sequence, "environment": {"preset": preset}, "n_random": 1000})
                    add_score_rows(label, origin, scored, "common_2000_1000_seed42_autoaligned")
            csv_write(output / "scores.csv", score_rows)
            dump(output / "sequences.json", [{"id": label, "origin": origin, "sequence": sequence} for label, origin, sequence in all_sequences])
            (output / "examples.fasta").write_text("".join(f">{label} origin={origin}\n{sequence}\n" for label, origin, sequence in all_sequences), encoding="utf-8")
            distances = [{"member_a": i + 1, "member_b": j + 1,
                          "hamming_distance": sum(a != b for a, b in zip(left["sequence"], right["sequence"]))}
                         for i, left in enumerate(family["members"]) for j, right in enumerate(family["members"]) if j > i]
            csv_write(output / "family_hamming_distances.csv", distances)
            post("comparison_native_protocol", "/api/compare", templates["compare"])
            profile, _ = post("penetration_native_protocol", "/api/optimal-penetration", templates["penetration"])
            csv_write(output / "penetration.csv", profile["profile"])
            pdb, _ = post("idealized_helix", "/api/pdb", {"sequence": family["members"][0]["sequence"]})
            (output / "family_1_idealized.pdb").write_text(pdb["pdb"], encoding="utf-8")

            sensitivity_rows = []
            for n_decoys in (500, 2000):
                for n_random in (500, 2000):
                    for seed in (42, 43, 44):
                        result, elapsed = post(f"sensitivity_d{n_decoys}_r{n_random}_seed{seed}", "/api/score", {
                            **templates["score"], "n_decoys": n_decoys, "n_random": n_random, "seed": seed})
                        sensitivity_rows.append({"n_decoys": n_decoys, "n_random": n_random, "seed": seed,
                                                 "energy": result["energy"], "z_score": result["z_score"],
                                                 "percentile_lower_is_better": result["percentile_lower_is_better"],
                                                 "wall_seconds": elapsed})
            csv_write(output / "reference_sampling_sensitivity.csv", sensitivity_rows)

            # Exhaustive reference for one tractable aligned objective. This tests
            # optimizer behavior, not the physical accuracy of the score.
            exact_alphabet = list("AEKL")
            exact_environment = engine.resolve_environment({"preset": "interfacial_neg"})
            with redirect_stdout(log):
                exact_builder = engine.build_builder(6, exact_alphabet, exact_environment, n_decoys=2000, zscore_seed=42, oriented=True)
            capture_calibration()
            exact_start = time.perf_counter()
            exact_rows = []
            for codes in itertools.product(range(len(exact_alphabet)), repeat=6):
                exact_rows.append({"sequence": engine.codes_to_seq(list(codes), exact_alphabet),
                                   "energy": engine.compute_breakdown(exact_builder, list(codes), WEIGHTS)["total"]})
            exact_elapsed = time.perf_counter() - exact_start
            exact_minimum = min(row["energy"] for row in exact_rows)
            exact_minimizers = [row["sequence"] for row in exact_rows if math.isclose(row["energy"], exact_minimum, rel_tol=0, abs_tol=1e-9)]
            csv_write(output / "exact_search_all.csv", exact_rows)
            exact_trials = []
            for seed in range(42, 47):
                trial_start = time.perf_counter()
                with redirect_stdout(log):
                    result = engine.simulated_annealing(exact_builder, exact_alphabet, WEIGHTS, steps=1000, restarts=2, seed=seed)
                elapsed = time.perf_counter() - trial_start
                gap = result["energy"] - exact_minimum
                if gap < -1e-9:
                    raise RuntimeError("Annealing result below exhaustively enumerated minimum")
                exact_trials.append({"seed": seed, "sequence": result["sequence"], "energy": result["energy"],
                                     "exact_minimum": exact_minimum, "absolute_optimality_gap": gap,
                                     "recovered_exact_minimum": abs(gap) <= 1e-9, "wall_seconds": elapsed})
            csv_write(output / "exact_search_annealing.csv", exact_trials)
            dump(output / "exact_search_summary.json", {
                "length": 6, "alphabet": exact_alphabet, "candidate_count": len(exact_rows),
                "environment": exact_environment, "weights": WEIGHTS, "n_decoys": 2000,
                "calibration_seed": 42, "search_seeds": list(range(42, 47)), "steps": 1000, "restarts": 2,
                "orientation": "each candidate, calibration decoy and reference sequence uses its own hydrophobic-moment alignment", "absolute_tolerance": 1e-9,
                "exact_minimum": exact_minimum, "exact_minimizers": exact_minimizers,
                "enumeration_wall_seconds": exact_elapsed,
                "successful_trials": sum(row["recovered_exact_minimum"] for row in exact_trials), "trials": exact_trials,
                "interpretation": "Optimization check on this finite objective only; does not establish biological validity or optimality at other lengths/alphabets."})
            print(f"exact search: {len(exact_rows)} sequences, {sum(row['recovered_exact_minimum'] for row in exact_trials)}/5 annealing successes", flush=True)

            # Three fixed seeds per size; each pair has identical request content.
            for length in (8, 16, 24):
                for seed in (42, 43, 44):
                    for workflow in ("score", "design"):
                        for path in cache.glob("zscore_stats_*.json"):
                            path.unlink()
                        payload = {**COMMON, "seed": seed, "environment": {"preset": "interfacial_neg"}}
                        if workflow == "score":
                            payload.update(sequence=(PROBES["probe_amphiphilic"] * 2)[:length], n_random=1000)
                        else:
                            payload.update(length=length, steps=3000, restarts=2)
                        previous = None
                        for state in ("cold", "warm"):
                            name = f"timing_{workflow}_L{length}_seed{seed}_{state}"
                            result, elapsed = post(name, f"/api/{workflow}", payload)
                            if state == "warm" and result != previous:
                                raise RuntimeError(f"Identical request yielded different results for {name}")
                            previous = result
                            if state == "cold" and len(calls[-1]["new_calibration_files"]) != 1:
                                raise RuntimeError(f"Expected one calibration model for {name}")
                            runtime_rows.append({"workflow": workflow, "length": length, "seed": seed,
                                                 "calibration_cache": state, "wall_seconds": elapsed,
                                                 "sequence": result["sequence"], "energy": result["energy"], "z_score": result["z_score"],
                                                 "new_calibration_count": len(calls[-1]["new_calibration_files"])})
                            csv_write(output / "timings.csv", runtime_rows)
            metadata["deterministic_warm_repeats"] = 18
            metadata["usage_log_created"] = (Path(temporary) / "var" / "usage_events.jsonl").exists()

    timing_summary = []
    for workflow in ("score", "design"):
        for length in (8, 16, 24):
            for state in ("cold", "warm"):
                values = [row["wall_seconds"] for row in runtime_rows if row["workflow"] == workflow and row["length"] == length and row["calibration_cache"] == state]
                timing_summary.append({"workflow": workflow, "length": length, "calibration_cache": state,
                                       "n": len(values), "median_seconds": statistics.median(values),
                                       "minimum_seconds": min(values), "maximum_seconds": max(values)})
    csv_write(output / "timing_summary.csv", timing_summary)
    dump(output / "calibration_statistics.json", calibration_records)
    dump(output / "calls.json", calls)
    metadata.update(finished_utc=datetime.now(timezone.utc).isoformat(), total_wall_seconds=time.perf_counter() - start,
                    request_count=len(calls), distinct_calibration_records=len(calibration_records),
                    final_source=source_metadata(repo))
    if metadata["source"]["runtime_source_sha256"] != metadata["final_source"]["runtime_source_sha256"]:
        raise RuntimeError("Source changed during the experiment; discard these results and rerun from an immutable source snapshot.")
    dump(output / "metadata.json", metadata)
    print(f"Completed {len(calls)} requests in {metadata['total_wall_seconds']:.1f} s.", flush=True)


if __name__ == "__main__":
    main()
```
