## Additional file 2. Computational evaluation, inputs and results for "Helix Design Studio: designing antimicrobial peptide candidates for membrane environments": summarize_results.py

```
#!/usr/bin/env python3
"""Validate completed outputs and regenerate summaries and their SHA256 manifest.

Uses only the Python standard library and performs no new scientific calculations.
Run this again after the manuscript sources have received their final edits.
"""
from __future__ import annotations

import argparse
import csv
import hashlib
import json
import math
from pathlib import Path
import statistics


def read_json(path):
    return json.loads(path.read_text(encoding="utf-8"))


def write_json(path, value):
    path.write_text(json.dumps(value, indent=2, sort_keys=True, allow_nan=False) + "\n", encoding="utf-8")


def read_csv(path):
    with path.open(newline="", encoding="utf-8") as handle:
        return list(csv.DictReader(handle))


def main():
    parser = argparse.ArgumentParser(description=__doc__)
    parser.add_argument("--results", type=Path, default=Path(__file__).resolve().parents[1] / "results")
    args = parser.parse_args()
    results = args.results.resolve()
    submission = results.parent
    metadata = read_json(results / "metadata.json")
    score_rows = read_csv(results / "scores.csv")
    timing_rows = read_csv(results / "timings.csv")
    native_rows = read_csv(results / "native_design_scores.csv")
    calls = read_json(results / "calls.json")
    response_paths = sorted((results / "responses").glob("*.json"))
    if len(calls) != metadata["request_count"] or len(response_paths) != len(calls):
        raise ValueError("Incomplete API request/response provenance")
    if metadata["source"]["runtime_source_sha256"] != metadata["final_source"]["runtime_source_sha256"]:
        raise ValueError("Runtime source changed during computation")
    if metadata["usage_log_created"]:
        raise ValueError("Unexpected usage log in the offline experiment")
    for row in score_rows:
        if int(row["n_random"]) != 1000 or int(row["n_decoys"]) != 2000 or int(row["calibration_seed"]) != 42:
            raise ValueError("Inconsistent common evaluation protocol")
        if not math.isclose(float(row["energy"]), float(row["term_total"]), rel_tol=0, abs_tol=1e-10):
            raise ValueError("Score and breakdown total differ")
    for row in timing_rows:
        expected_calibrations = 1 if row["calibration_cache"] == "cold" else 0
        if int(row["new_calibration_count"]) != expected_calibrations:
            raise ValueError("Unexpected calibration cache state")
    response = lambda name: read_json(results / "responses" / f"{name}.json")["response"]
    family = response("family")
    specificity = response("specificity")
    cross = response("cross_design")
    depth = response("penetration_native_protocol")
    exact = read_json(results / "exact_search_summary.json")
    distances = [int(row["hamming_distance"]) for row in read_csv(results / "family_hamming_distances.csv")]
    family_energies = [member["design_geometry"]["energy"] for member in family["members"]]
    parameter_record = read_json(results / "parameter_source_checks.json") if (results / "parameter_source_checks.json").exists() else None
    if parameter_record:
        if parameter_record.get("software_commit") != metadata["source"]["commit"] or parameter_record.get("software_version") != metadata["protocol"]["version"]:
            raise ValueError("Parameter-source record identifies a different software revision")
        for name, matrix in parameter_record["matrices"].items():
            if matrix["sha256"] != metadata["source"]["runtime_source_sha256"][matrix["resource"]]:
                raise ValueError(f"Parameter-source record differs from the evaluated {name} resource")
            if not matrix["verification"]["verified"]:
                raise ValueError(f"{name} failed primary-source verification")
        if hashlib.sha256((results / "parameter_tables.pdf").read_bytes()).hexdigest() != parameter_record["table_export"]["sha256"]:
            raise ValueError("Parameter-table PDF differs from the recorded export")
    parameter_status = parameter_record["status"] if parameter_record else metadata["parameter_mapping_status"]
    common_energy = {(row["id"], row["environment"]): float(row["energy"]) for row in score_rows}
    for row in native_rows:
        if not math.isclose(float(row["energy"]), common_energy[row["id"], row["environment"]], rel_tol=0, abs_tol=1e-10):
            raise ValueError("Aligned native and common-profile energies differ under the same calibration")
    for candidate in [*family["members"],
                      *(score for member in specificity["candidates"] for score in [member["target_environment"], *member["off_target_environments"]]),
                      cross["environment_a"], cross["environment_b"]]:
        for key in ("energy", "z_score", "percentile_lower_is_better", "breakdown"):
            if candidate[key] != candidate["design_geometry"][key]:
                raise ValueError(f"Search score and reported {key} differ")
    summary = {
        "status": parameter_status,
        "commit": metadata["source"]["commit"],
        "total_requests": len(calls),
        "common_profile_rows": len(score_rows),
        "native_score_rows": len(native_rows),
        "family": {
            "returned": family["family_size_returned"],
            "native_energy_range": [min(family_energies), max(family_energies)],
            "hamming_range": [min(distances), max(distances)],
        },
        "specificity": [{"sequence": member["sequence"], **member["specificity_design"]}
                        for member in specificity["candidates"]],
        "depth": {
            "points": len(depth["profile"]),
            "minimum_percent": depth["best_penetration_percent"],
            "minimum_z_score": depth["best_score"]["z_score"],
            "halfwidth_degrees": depth["best_score"]["environment"]["wheel_halfwidth_deg"],
        },
        "exact_search": exact,
        "timing_summary": read_csv(results / "timing_summary.csv"),
        "reference_sensitivity": [],
    }
    sensitivities = read_csv(results / "reference_sampling_sensitivity.csv")
    for n_decoys in (500, 2000):
        for n_random in (500, 2000):
            selected = [row for row in sensitivities if int(row["n_decoys"]) == n_decoys and int(row["n_random"]) == n_random]
            values = [float(row["z_score"]) for row in selected]
            summary["reference_sensitivity"].append({
                "n_decoys": n_decoys, "n_random": n_random,
                "seeds": [int(row["seed"]) for row in selected], "z_values": values,
                "mean_z": statistics.mean(values), "sample_sd_z": statistics.stdev(values),
                "min_z": min(values), "max_z": max(values),
            })
    write_json(results / "summary.json", summary)

    # Keep interpretation notes synchronized with the measured run, including timing.
    score = {(row["id"], row["environment"]): float(row["z_score"]) for row in score_rows}
    timings = {(row["workflow"], int(row["length"]), row["calibration_cache"]): float(row["median_seconds"])
               for row in summary["timing_summary"]}
    runtime = {call["name"]: call["wall_seconds"] for call in calls}
    cpu = metadata.get("cpu") or "unavailable CPU model"
    memory = metadata.get("memory_bytes")
    memory_label = f"{int(memory) / 2**30:g} GiB RAM" if memory else "unavailable RAM metadata"
    cpu_count = metadata.get("logical_cpus")
    core_label = f"{cpu_count} logical CPUs" if cpu_count is not None else "unavailable logical-CPU count"
    status_label = lambda value: "unavailable" if value is None else value or "clean"
    commit_label = metadata["source"].get("commit") or "unavailable (no Git metadata)"
    notes = [
        "# Measured results for the release-candidate draft\n",
        f"All values below were produced by the completed run of `run_examples.py`, using commit `{commit_label}`. "
        f"Initial Git status: `{status_label(metadata['source']['git_status'])}`; final Git status: `{status_label(metadata['final_source']['git_status'])}`. "
        "Runtime source hashes were identical before and after the run. "
        "The numerical parameter dataset and its checksums are recorded in `results/parameter_source_checks.json`; "
        "the neighbour matrices are also reproduced in `results/parameter_tables.pdf`. "
        "These calculations characterize the implemented model; they do not establish biological activity.\n",
        f"The run performed {len(calls)} successful local API requests in {metadata['total_wall_seconds']:.3f} s. "
        f"It retained {len(score_rows)} common-protocol scores, {len(native_rows)} native aligned design scores, "
        f"{metadata['distinct_calibration_records']} distinct calibration records, {len(depth['profile'])} profile points, "
        f"{len(sensitivities)} reference-sampling sensitivity measurements, and {exact['candidate_count']:,} exhaustive-reference energies. "
        f"All {metadata['deterministic_warm_repeats']} identical payload repeats returned identical JSON with populated calibration caches. "
        "The run created no usage log.\n",
        "## Example observations\n",
        "For `LKKLLKLLKKLLKLLKKLLK`, common-protocol reference Z scores were "
        + ", ".join(f"{score['probe_amphiphilic', env]:.3f} ({label})" for env, label in [
            ("interfacial_neg", "anionic interface"), ("interfacial_neu", "neutral interface"),
            ("interfacial_pos", "cationic interface"), ("homogeneous_polar", "homogeneous polar"),
            ("homogeneous_apolar", "homogeneous apolar")])
        + ". For the acidic analogue `LEELLELLEELLELLEELLE`, the anionic, neutral, and cationic interface values were "
        + ", ".join(f"{score['probe_acidic', env]:.3f}" for env in ["interfacial_neg", "interfacial_neu", "interfacial_pos"])
        + f". Poly-leucine had its lowest value in the homogeneous apolar preset ({score['probe_hydrophobic', 'homogeneous_apolar']:.3f}). "
        "These are illustrative responses of the implemented objective, not experimental validation.\n",
        f"The family returned {family['family_size_returned']} members, with aligned model scores from {min(family_energies):.3f} "
        f"to {max(family_energies):.3f} and pairwise Hamming distances from {min(distances)} to {max(distances)}. "
        f"The lowest-native-score sequence was `{family['members'][0]['sequence']}`. "
        "The W/K/R-rich outputs illustrate that this unconstrained objective can push composition toward extreme residues; "
        "no synthesis, toxicity, activity, aggregation, or developability filter is implied. "
        "Common rescoring in `scores.csv` uses the same alignment rule; its reference count can differ from that of the native design result.\n",
        "Specificity candidates had native target-versus-lowest-off-target score margins of "
        + ", ".join(f"{member['specificity_design']['energy_margin']:.3f}" for member in specificity['candidates'])
        + ", and native objectives of "
        + ", ".join(f"{member['specificity_design']['objective']:.3f}" for member in specificity['candidates'])
        + f". Both specificity objectives and the cross-design objective ({cross['cross_design']['objective']:.3f}) "
        "were reconstructed from their returned native component scores. These margins are normalized model-score differences, "
        "not physical transfer free energies or experimental selectivity measurements.\n",
        f"The scan minimum occurred at the displayed {depth['best_penetration_percent']}% setting "
        f"(Z={depth['best_score']['z_score']:.3f}, score={depth['best_score']['energy']:.3f}), "
        f"with interfacial half-width {depth['best_score']['environment']['wheel_halfwidth_deg']:g} degrees. "
        "For interior percentage settings p, half-width = 40 + 1.1p degrees; endpoints 0 and 100 are replaced "
        "by homogeneous polar/apolar presets. The percentage is a geometric index, not measured membrane insertion. "
        "The same seed protocol is used across scan points; this minimum has no experimental confidence interval.\n",
        "## Exact reference and finite sampling\n",
        f"At length {exact['length']} and alphabet {'/'.join(exact['alphabet'])}, exhaustive enumeration found "
        f"{len(exact['exact_minimizers'])} minimum sequence(s): {', '.join(exact['exact_minimizers'])}, "
        f"with score {exact['exact_minimum']:.15f}. {exact['successful_trials']}/{len(exact['trials'])} annealing searches "
        "recovered it within absolute tolerance 1e-9. This verifies one finite optimization instance of the same objective; "
        "it does not establish biological validity or global optimality for larger examples.\n",
        "Observed reference Z-score ranges across the three sensitivity seeds were:\n",
    ]
    notes.extend(f"- {row['n_decoys']:,} calibration decoys / {row['n_random']:,} references: "
                 f"{row['min_z']:.3f} to {row['max_z']:.3f}." for row in summary["reference_sensitivity"])
    notes.extend([
        "\nThese are sampling ranges for one sequence, not biological prediction uncertainty or inferential confidence intervals.\n",
        "## Timing measurements\n",
        f"Recorded hardware: {cpu}; {core_label}; {memory_label}. In-process API wall-time medians over three seeds:\n",
        "| Length | Score, empty calibration cache | Score, populated cache | Design, empty calibration cache | Design, populated cache |",
        "|---:|---:|---:|---:|---:|",
    ])
    notes.extend(f"| {length} | {timings['score', length, 'cold']:.3f} s | {timings['score', length, 'warm']:.3f} s | "
                 f"{timings['design', length, 'cold']:.3f} s | {timings['design', length, 'warm']:.3f} s |"
                 for length in (8, 16, 24))
    notes.extend([
        "\nScore requests used 2,000 calibration decoys and 1,000 reference sequences; design requests used 3,000 steps "
        "and two restarts with the native 500-reference output. Imports had already completed. Values exclude network "
        "transport, queueing, server startup, and browser rendering. Observed ranges are in `timing_summary.csv`.\n",
        f"Single examples took {runtime['family']:.3f} s (family), {runtime['specificity']:.3f} s (specificity), "
        f"{runtime['cross_design']:.3f} s (cross-design), and {runtime['penetration_native_protocol']:.3f} s (21-point scan). "
        "These are one measurement each, unlike the three-seed size benchmark.\n",
        "The manifest hashes all result files except itself and `runtime.log`, and available experiment/manuscript "
        "sources. Regenerate it with `python experiments/summarize_results.py` after the final source edits.\n",
    ])
    (submission / "experiments" / "RESULTS_NOTES.md").write_text("\n".join(notes), encoding="utf-8")

    manifest = {
        path.relative_to(results).as_posix(): hashlib.sha256(path.read_bytes()).hexdigest()
        for path in sorted(results.rglob("*"))
        if path.is_file() and path.name not in {"reproduction_manifest.json", "runtime.log"}
        and "__pycache__" not in path.parts and path.suffix != ".pyc"
    }
    sources = [
        "experiments/run_examples.py", "experiments/compare_descriptors.py", "experiments/summarize_results.py",
        "experiments/export_parameter_tables.py", "experiments/compare_search_orientation.py",
        "experiments/validate_search_alignment.py", "experiments/compare_m1_scoring.py",
        "experiments/audit_scoring_direction.py",
        "experiments/README.md", "experiments/RESULTS_NOTES.md", "experiments/runner_checks.json",
        "source/build_manuscript.py", "source/make_figures.py", "source/export_manuscript_figures.py",
        "source/make_graphical_abstract.py", "source/manuscript_body.tex", "source/manuscript_preamble.tex",
        "source/manuscript_references.tex", "source/transverse_alignment_figure.tex", "source/sn-jnl.cls",
        "figures/FIGURE_CAPTIONS.md", "REPRODUCE.md",
    ]
    for relative in sources:
        path = submission / relative
        if path.is_file():
            manifest[f"../{relative}"] = hashlib.sha256(path.read_bytes()).hexdigest()
    write_json(results / "reproduction_manifest.json", manifest)
    print(f"Validated {len(calls)} responses and {len(score_rows)} common scores; regenerated summary, notes, and {len(manifest)} SHA256 entries.")


if __name__ == "__main__":
    main()
```
