## Additional file 2. Computational evaluation, inputs and results for "Helix Design Studio: designing antimicrobial peptide candidates for membrane environments": validate_search_alignment.py

```
#!/usr/bin/env python3
"""Independently check hydrophobic-moment alignment in a selected source checkout.

Run with the checkout's Python dependencies installed, for example:
    python experiments/validate_search_alignment.py --repo ../helix-design-studio-web \
        --output results/independent_alignment_audit.json

This is an implementation consistency check, not biological model validation or a
sampling-convergence benchmark. The temporary calibration cache is discarded.
"""

from __future__ import annotations

import argparse
import ast
from datetime import datetime, timezone
import hashlib
import importlib
import itertools
import json
import math
import os
from pathlib import Path
import platform
import subprocess
import sys
import tempfile
from unittest.mock import patch


def require(condition: bool, message: str) -> None:
    """Keep checks active even when Python is invoked with optimization enabled."""
    if not condition:
        raise AssertionError(message)


def git_output(repo: Path, *arguments: str) -> str:
    return subprocess.check_output(
        ["git", "-C", str(repo), *arguments], text=True, stderr=subprocess.PIPE,
    ).strip()


def sha256_file(path: Path) -> str:
    return hashlib.sha256(path.read_bytes()).hexdigest()


def source_version(repo: Path) -> str:
    tree = ast.parse((repo / "app/main.py").read_text())
    for node in tree.body:
        if isinstance(node, ast.Assign) and any(
            isinstance(target, ast.Name) and target.id == "APP_VERSION"
            for target in node.targets
        ):
            return str(ast.literal_eval(node.value))
    raise ValueError("APP_VERSION was not found in app/main.py")


def validate(repo: Path) -> dict:
    repo = repo.expanduser().resolve(strict=True)
    commit = git_output(repo, "rev-parse", "HEAD")
    tree = git_output(repo, "rev-parse", "HEAD^{tree}")
    dirty = bool(git_output(repo, "status", "--porcelain"))
    version = source_version(repo)
    sys.path.insert(0, str(repo))
    with patch.dict(os.environ, {"HELIX_CODE_DIR": str(repo / "Code")}):
        engine = importlib.import_module("app.engine")
    require(Path(engine.__file__).resolve() == repo / "app/engine.py",
            "The imported engine does not belong to the requested checkout")
    core = importlib.import_module(engine.HamiltonianBuilder.__module__)
    require(Path(core.__file__).resolve() == repo / "Code/core/hamiltonian_builder.py",
            "The imported Hamiltonian does not belong to the requested checkout")
    numpy = importlib.import_module("numpy")
    source_paths = [
        "app/engine.py", "app/main.py", "Code/core/hamiltonian_builder.py",
        "Code/data_loaders/energy_matrix_loader.py",
        *[f"Code/core/resources/{name}" for name in (
            "mj_matrix.txt", "helix_pairs_prop.txt", "helix_sd_i_3.txt", "helix_sd_i_4.txt",
        )],
    ]
    report = {
        "status": "passed",
        "created_utc": datetime.now(timezone.utc).isoformat(),
        "provenance": {
            "repo_argument_resolved": str(repo),
            "commit": commit,
            "commit_tree": tree,
            "working_tree_dirty": dirty,
            "software_version": version,
            "python": platform.python_version(),
            "numpy": numpy.__version__,
            "seed_protocol": engine.SEED_PROTOCOL,
            "parameters_sha256": core.parameter_set_sha256(),
            "source_sha256": {name: sha256_file(repo / name) for name in source_paths},
            "validation_script_sha256": sha256_file(Path(__file__).resolve()),
        },
        "protocol": {
            "length": 4,
            "residues": ["A", "K", "L"],
            "seed": 23,
            "n_decoys": 100,
            "orientation_oracle_reference_size": 50,
            "steps": 100,
            "restarts": 1,
            "family_size": 2,
            "oversample_factor": 1,
            "num_specificity_sequences": 1,
            "cross_lambda_gap": 0.7,
            "specificity_lambda_balance": 0.35,
            "environment_a": {"preset": "interfacial_neg", "wheel_halfwidth_deg": 63.0},
            "environment_b": {"preset": "interfacial_pos"},
            "design_reference_size": engine.DESIGN_N_RANDOM,
            "specificity_reference_size": engine.SPECIFICITY_N_RANDOM,
            "sampling_convergence_tested": False,
            "biological_validity_tested": False,
        },
        "independent_checks": {},
    }
    with tempfile.TemporaryDirectory(prefix="helix-alignment-audit-") as cache_dir, patch.dict(
        os.environ, {"HELIX_CACHE_DIR": cache_dir},
    ):
        alphabet = report["protocol"]["residues"]
        environment = engine.resolve_environment(report["protocol"]["environment_a"])
        weights = engine.normalize_weights(None)
        search, displayed = engine.report_models(
            4, alphabet, environment, weights, n_decoys=100, seed=23, n_random=50,
        )
        require(search is displayed, "Search and displayed scores use different models")
        count = 0
        phases = set()
        maximum_phase_error = 0.0
        maximum_exposure_error = 0.0
        for residues in itertools.product(alphabet, repeat=4):
            sequence = "".join(residues)
            codes = engine.seq_to_codes(sequence, alphabet)
            hydro = [engine.HYDROPHOBICITY_FP[aa] for aa in sequence]
            mx = sum(h * math.cos(math.radians(100 * i)) for i, h in enumerate(hydro))
            my = sum(h * math.sin(math.radians(100 * i)) for i, h in enumerate(hydro))
            phase = -math.degrees(math.atan2(my, mx)) % 360.0
            phase_error = abs(search.builder.phase_for(codes) - phase)
            require(phase_error <= 1e-10, f"Incorrect aligned phase for {sequence}")
            mask = [abs((100 * i + phase + 180) % 360 - 180) <= 63.0 for i in range(4)]
            exposure = sum(-h if inside else h for h, inside in zip(hydro, mask)) / 4
            exposure_error = abs(search.builder.raw_terms(codes)["env_pol"] - exposure)
            require(exposure_error <= 1e-12, f"Incorrect exposure term for {sequence}")
            maximum_phase_error = max(maximum_phase_error, phase_error)
            maximum_exposure_error = max(maximum_exposure_error, exposure_error)
            phases.add(round(phase, 8))
            count += 1
        report["independent_checks"]["exhaustive_4_residue_AKL"] = {
            "sequences": count,
            "distinct_aligned_phases": len(phases),
            "manual_phase_and_environment_term_match": True,
            "maximum_phase_error_degrees": maximum_phase_error,
            "maximum_raw_exposure_error": maximum_exposure_error,
        }
        report["independent_checks"]["search_and_report_share_same_model_object"] = True
        captured = []
        original = engine.compute_breakdown

        def tracked(builder, codes, supplied_weights):
            captured.append((builder.align_phase, tuple(codes), builder.phase_for(codes)))
            return original(builder, codes, supplied_weights)

        common = dict(length=4, residues=alphabet, n_decoys=100, steps=100, restarts=1, seed=23)
        with patch.object(engine, "compute_breakdown", side_effect=tracked):
            design = engine.design_sequence(environment=environment, **common)
            family = engine.design_family(
                environment=environment, family_size=2, oversample_factor=1, **common,
            )
            cross = engine.cross_environment_design(
                environment_a=environment, environment_b={"preset": "interfacial_pos"},
                lambda_gap=0.7, **common,
            )
            specificity = engine.design_specificity(
                target_environment=environment, off_target_environments=[{"preset": "interfacial_pos"}],
                lambda_balance=0.35, num_sequences=1, **common,
            )
        require(bool(captured), "No score computations were observed")
        require(all(aligned for aligned, _, _ in captured), "A workflow used a fixed-phase builder")
        for scored in [design, *family["members"], cross["environment_a"], cross["environment_b"],
                       specificity["target_environment"], *specificity["off_target_environments"]]:
            for key in ("energy", "z_score", "breakdown", "percentile_lower_is_better"):
                require(scored[key] == scored["design_geometry"][key], f"Inconsistent alias field {key}")
            require(scored["environment"]["wheel_phase_deg"] == scored["design_geometry"]["wheel_phase_deg"],
                    "The reported geometry phase differs from the scoring phase")
        a, b = cross["environment_a"]["energy"], cross["environment_b"]["energy"]
        cross_error = abs(cross["cross_design"]["objective"] - (a + b + 0.7 * abs(a - b)))
        require(cross_error <= 1e-12, "Cross-design objective differs from displayed energies")
        target = specificity["target_environment"]["energy"]
        off = min(item["energy"] for item in specificity["off_target_environments"])
        specificity_error = abs(specificity["specificity_design"]["objective"] - (target - 0.65 * off))
        require(specificity_error <= 1e-12, "Specificity objective differs from displayed energies")
        report["independent_checks"]["four_workflow_evaluations_are_aligned"] = {
            "calls_observed": len(captured),
            "distinct_phases": len({round(phase, 8) for _, _, phase in captured}),
        }
        report["independent_checks"]["export_aliases_match_containing_scores"] = True
        report["independent_checks"]["cross_and_specificity_objectives_match_displayed_energies"] = {
            "passed": True,
            "absolute_cross_objective_error": cross_error,
            "absolute_specificity_objective_error": specificity_error,
        }
    return report


def main() -> None:
    parser = argparse.ArgumentParser(description=__doc__)
    parser.add_argument("--repo", type=Path,
                        default=Path(__file__).resolve().parents[2] / "helix-design-studio-web",
                        help="Source checkout to validate (its commit and source hashes are recorded)")
    parser.add_argument("--output", type=Path, help="Optional JSON output path")
    args = parser.parse_args()
    report = validate(args.repo)
    encoded = json.dumps(report, indent=2, ensure_ascii=False) + "\n"
    if args.output is not None:
        args.output.parent.mkdir(parents=True, exist_ok=True)
        args.output.write_text(encoded)
    print(encoded, end="")


if __name__ == "__main__":
    main()
```
