## Additional file 2. Computational evaluation, inputs and results for "Helix Design Studio: designing antimicrobial peptide candidates for membrane environments": parameter_tables.pdf

### M1: sequence separation k = 1

Rows: residue at i. Columns: residue at i+k. Each table contains 400 runtime values.

Nacar (2020), Fig. 3, p. 25; source layout transposed to preserve sequence direction.

Source pairs are n,n+4; the model reuses these dimensionless propensities at i,i+1.

Raw source values. Score: -mean(M1) + mean(M3) + mean(M4), followed by joint standardization and weighting.

| i / i+k | A | R | N | D | C | Q | E | G | H | I | L | K | M | F | P | S | T | W | Y | V |
|---|---|---|---|---|---|---|---|---|---|---|---|---|---|---|---|---|---|---|---|---|
| A | 2.734 | 1.724 | 0.817 | 0.721 | 1.221 | 1.524 | 1.326 | 0.764 | 1.114 | 1.251 | 1.782 | 1.316 | 1.637 | 1.301 | 0.000 | 1.036 | 0.799 | 1.218 | 1.329 | 1.107 |
| R | 1.648 | 1.466 | 0.886 | 1.197 | 0.697 | 1.593 | 2.216 | 0.517 | 0.902 | 0.983 | 1.404 | 1.005 | 1.248 | 0.870 | 0.000 | 0.799 | 0.771 | 1.003 | 0.876 | 0.728 |
| N | 1.224 | 1.013 | 0.882 | 0.669 | 0.873 | 1.329 | 1.066 | 0.338 | 0.663 | 0.935 | 1.075 | 1.076 | 1.138 | 0.822 | 0.000 | 0.741 | 0.689 | 0.981 | 0.841 | 0.699 |
| D | 1.769 | 1.804 | 0.976 | 0.729 | 0.775 | 1.557 | 1.292 | 0.389 | 0.820 | 0.927 | 1.201 | 1.615 | 1.325 | 0.988 | 0.000 | 0.859 | 0.815 | 1.127 | 0.882 | 0.756 |
| C | 1.448 | 0.893 | 0.644 | 0.445 | 1.800 | 0.724 | 0.696 | 0.486 | 0.855 | 1.051 | 1.480 | 0.716 | 1.575 | 1.057 | 0.000 | 0.688 | 0.558 | 1.276 | 0.998 | 0.814 |
| Q | 1.777 | 1.483 | 1.433 | 1.954 | 0.822 | 1.943 | 2.046 | 0.526 | 0.980 | 1.045 | 1.429 | 1.428 | 1.462 | 1.087 | 0.000 | 1.012 | 0.906 | 1.382 | 0.974 | 0.704 |
| E | 1.900 | 2.523 | 1.460 | 0.873 | 0.794 | 2.144 | 1.806 | 0.501 | 1.146 | 1.063 | 1.451 | 2.490 | 1.315 | 1.015 | 0.000 | 1.074 | 0.914 | 1.141 | 0.967 | 0.763 |
| G | 1.155 | 0.770 | 0.370 | 0.386 | 0.606 | 0.602 | 0.512 | 0.370 | 0.453 | 0.550 | 0.825 | 0.524 | 0.819 | 0.590 | 0.000 | 0.507 | 0.457 | 0.577 | 0.551 | 0.514 |
| H | 1.238 | 1.107 | 0.627 | 0.570 | 1.088 | 1.148 | 0.900 | 0.365 | 0.913 | 1.017 | 1.264 | 0.881 | 1.403 | 0.749 | 0.000 | 0.676 | 0.736 | 0.885 | 0.701 | 0.994 |
| I | 1.275 | 1.117 | 0.908 | 0.423 | 1.640 | 1.109 | 0.856 | 0.432 | 0.947 | 1.690 | 1.946 | 1.228 | 1.731 | 1.311 | 0.000 | 0.619 | 0.753 | 0.988 | 1.186 | 1.167 |
| L | 1.575 | 1.374 | 0.761 | 0.519 | 1.705 | 1.250 | 1.008 | 0.495 | 1.201 | 1.690 | 2.268 | 1.200 | 1.909 | 1.496 | 0.000 | 0.766 | 0.778 | 1.175 | 1.501 | 1.238 |
| K | 1.297 | 1.016 | 0.957 | 1.551 | 0.566 | 1.545 | 2.264 | 0.381 | 0.724 | 0.928 | 1.077 | 1.465 | 1.061 | 0.817 | 0.000 | 0.787 | 0.692 | 0.751 | 0.865 | 0.553 |
| M | 1.580 | 1.244 | 0.996 | 0.542 | 1.795 | 1.237 | 0.994 | 0.569 | 1.206 | 1.719 | 2.031 | 1.191 | 2.878 | 1.571 | 0.000 | 0.756 | 0.752 | 1.281 | 1.429 | 1.197 |
| F | 1.143 | 0.918 | 0.688 | 0.335 | 1.884 | 0.977 | 0.644 | 0.343 | 1.089 | 1.165 | 1.637 | 0.860 | 1.844 | 1.328 | 0.000 | 0.625 | 0.682 | 1.132 | 1.391 | 1.283 |
| P | 1.276 | 1.000 | 0.421 | 0.410 | 0.405 | 0.986 | 0.855 | 0.238 | 0.492 | 0.531 | 0.726 | 0.898 | 0.673 | 0.516 | 0.000 | 0.442 | 0.398 | 0.559 | 0.486 | 0.471 |
| S | 1.484 | 1.244 | 0.839 | 0.632 | 0.888 | 1.303 | 0.945 | 0.439 | 0.845 | 1.010 | 1.211 | 1.216 | 1.324 | 1.019 | 0.000 | 0.804 | 0.716 | 0.960 | 0.876 | 0.712 |
| T | 1.297 | 1.192 | 0.865 | 0.536 | 1.036 | 1.153 | 0.863 | 0.384 | 0.821 | 1.332 | 1.598 | 0.989 | 1.513 | 0.999 | 0.000 | 0.579 | 0.653 | 0.983 | 0.875 | 0.995 |
| W | 1.342 | 1.244 | 1.046 | 0.419 | 1.803 | 1.496 | 0.977 | 0.447 | 1.246 | 1.181 | 1.434 | 1.133 | 1.704 | 1.309 | 0.000 | 0.777 | 0.819 | 1.328 | 1.197 | 1.053 |
| Y | 1.069 | 1.206 | 0.794 | 0.338 | 1.393 | 1.135 | 0.730 | 0.361 | 1.196 | 1.162 | 1.526 | 1.362 | 1.603 | 1.024 | 0.000 | 0.636 | 0.671 | 0.899 | 1.074 | 1.072 |
| V | 1.197 | 1.063 | 0.699 | 0.356 | 1.322 | 0.924 | 0.783 | 0.338 | 0.820 | 1.386 | 1.793 | 1.054 | 1.615 | 1.004 | 0.000 | 0.563 | 0.692 | 0.874 | 0.952 | 1.062 |

Source DOI: 10.1007/s10930-020-09880-6

Code/core/resources/helix_pairs_prop.txt

Commit: d6ccec1920f256a393b15215df63fd2dead61cd5

SHA-256 of resource: 266410b1335e1de1341e39e031ecd23054e174d3017322d51be1bb20c4e19684

### M3: sequence separation k = 3

Rows: residue at i. Columns: residue at i+k. Each table contains 400 runtime values.

Munoz and Serrano (1994), Table 2, p. 403; i,i+3 energies in kcal/mol, mapped by labels.

C/P pairs (grey cells) are a 76-cell zero-contribution extension; they are absent from the source.

Raw source values. Score: -mean(M1) + mean(M3) + mean(M4), followed by joint standardization and weighting.

| i / i+k | A | G | S | T | N | D | Q | E | H | K | R | Y | F | W | V | I | M | L | C | P |
|---|---|---|---|---|---|---|---|---|---|---|---|---|---|---|---|---|---|---|---|---|
| A | 0.000 | 0.000 | 0.050 | -0.050 | -0.050 | -0.050 | -0.050 | -0.150 | -0.150 | -0.150 | -0.150 | -0.130 | -0.130 | -0.130 | -0.050 | -0.150 | -0.150 | -0.250 | 0.000 | 0.000 |
| G | 0.200 | 0.400 | 0.600 | 0.600 | 0.600 | 0.600 | 0.000 | 0.000 | 0.020 | 0.000 | 0.000 | 0.050 | 0.050 | 0.050 | 0.000 | 0.000 | 0.000 | 0.000 | 0.000 | 0.000 |
| S | 0.150 | 0.400 | 0.400 | 0.400 | 0.600 | 0.600 | 0.000 | 0.000 | 0.020 | 0.000 | 0.000 | 0.050 | 0.050 | 0.050 | 0.000 | 0.000 | 0.000 | 0.000 | 0.000 | 0.000 |
| T | 0.150 | 0.400 | 0.400 | 0.400 | 0.600 | 0.600 | 0.000 | 0.000 | 0.020 | 0.000 | 0.000 | 0.050 | 0.050 | 0.050 | 0.000 | 0.000 | 0.000 | 0.000 | 0.000 | 0.000 |
| N | 0.150 | 0.400 | 0.600 | 0.600 | 0.600 | 0.600 | 0.000 | 0.000 | 0.020 | 0.000 | 0.000 | 0.050 | 0.050 | 0.050 | 0.000 | 0.000 | 0.000 | 0.000 | 0.000 | 0.000 |
| D | 0.150 | 0.400 | 0.600 | 0.600 | 0.600 | 0.300 | 0.000 | 0.300 | -0.100 | -0.200 | -0.200 | 0.050 | 0.050 | 0.050 | 0.000 | 0.000 | 0.000 | 0.000 | 0.000 | 0.000 |
| Q | -0.000 | -0.050 | 0.050 | -0.050 | -0.050 | -0.050 | 0.000 | -0.100 | -0.050 | -0.100 | -0.100 | -0.200 | -0.200 | -0.200 | -0.150 | -0.150 | -0.150 | -0.150 | 0.000 | 0.000 |
| E | -0.000 | -0.050 | 0.050 | -0.050 | -0.050 | 0.300 | -0.140 | 0.300 | -0.100 | -0.150 | -0.150 | -0.150 | -0.150 | -0.150 | -0.100 | -0.100 | -0.100 | -0.100 | 0.000 | 0.000 |
| H | 0.000 | 0.020 | 0.000 | 0.000 | 0.000 | 0.000 | 0.000 | -0.200 | 0.100 | 0.150 | 0.150 | -0.250 | -0.200 | -0.300 | -0.110 | -0.110 | -0.110 | -0.110 | 0.000 | 0.000 |
| K | 0.000 | 0.000 | 0.000 | 0.000 | 0.000 | 0.200 | 0.000 | 0.200 | 0.200 | 0.300 | 0.300 | 0.000 | 0.000 | 0.000 | 0.000 | 0.000 | 0.000 | 0.000 | 0.000 | 0.000 |
| R | 0.000 | 0.000 | 0.000 | 0.000 | 0.000 | 0.200 | 0.000 | 0.200 | 0.200 | 0.300 | 0.300 | 0.000 | 0.000 | 0.000 | 0.000 | 0.000 | 0.000 | 0.000 | 0.000 | 0.000 |
| Y | 0.000 | 0.130 | 0.130 | 0.130 | 0.130 | 0.130 | 0.000 | -0.310 | -0.250 | -0.310 | -0.310 | -0.200 | -0.200 | -0.200 | -0.250 | -0.250 | -0.250 | -0.250 | 0.000 | 0.000 |
| F | 0.000 | 0.130 | 0.130 | 0.130 | 0.130 | 0.130 | 0.000 | -0.310 | -0.250 | -0.310 | -0.310 | -0.200 | -0.200 | -0.200 | -0.250 | -0.250 | -0.250 | -0.250 | 0.000 | 0.000 |
| W | 0.000 | 0.130 | 0.130 | 0.130 | 0.130 | 0.130 | 0.000 | -0.310 | -0.250 | -0.310 | -0.310 | -0.200 | -0.200 | -0.200 | -0.250 | -0.250 | -0.250 | -0.250 | 0.000 | 0.000 |
| V | 0.100 | 0.000 | -0.000 | 0.000 | 0.000 | 0.000 | -0.150 | -0.150 | -0.350 | -0.150 | -0.150 | -0.450 | -0.450 | -0.450 | -0.350 | -0.350 | -0.350 | -0.350 | 0.000 | 0.000 |
| I | 0.100 | 0.000 | -0.000 | 0.000 | 0.000 | 0.000 | -0.150 | -0.150 | -0.350 | -0.150 | -0.150 | -0.450 | -0.450 | -0.450 | -0.350 | -0.350 | -0.350 | -0.350 | 0.000 | 0.000 |
| M | 0.100 | 0.000 | -0.000 | 0.000 | 0.000 | 0.000 | -0.150 | -0.150 | -0.300 | -0.150 | -0.150 | -0.450 | -0.450 | -0.450 | -0.350 | -0.350 | -0.350 | -0.350 | 0.000 | 0.000 |
| L | 0.100 | 0.000 | -0.000 | 0.000 | 0.000 | 0.000 | -0.150 | -0.150 | -0.350 | -0.150 | -0.150 | -0.450 | -0.450 | -0.450 | -0.350 | -0.350 | -0.350 | -0.350 | 0.000 | 0.000 |
| C | 0.000 | 0.000 | 0.000 | 0.000 | 0.000 | 0.000 | 0.000 | 0.000 | 0.000 | 0.000 | 0.000 | 0.000 | 0.000 | 0.000 | 0.000 | 0.000 | 0.000 | 0.000 | 0.000 | 0.000 |
| P | 0.000 | 0.000 | 0.000 | 0.000 | 0.000 | 0.000 | 0.000 | 0.000 | 0.000 | 0.000 | 0.000 | 0.000 | 0.000 | 0.000 | 0.000 | 0.000 | 0.000 | 0.000 | 0.000 | 0.000 |

Source DOI: 10.1038/nsb0694-399

Code/core/resources/helix_sd_i_3.txt

Commit: d6ccec1920f256a393b15215df63fd2dead61cd5

SHA-256 of resource: ac0fcc7c17ffeaeea6664906b5473c1c9ddd2f2145d0d75626e674b7eb0505c5

### M4: sequence separation k = 4

Rows: residue at i. Columns: residue at i+k. Each table contains 400 runtime values.

Munoz and Serrano (1994), Table 2, p. 403; i,i+4 energies in kcal/mol, mapped by labels.

C/P pairs (grey cells) are a 76-cell zero-contribution extension; they are absent from the source.

Raw source values. Score: -mean(M1) + mean(M3) + mean(M4), followed by joint standardization and weighting.

| i / i+k | A | G | S | T | N | D | Q | E | H | K | R | Y | F | W | V | I | M | L | C | P |
|---|---|---|---|---|---|---|---|---|---|---|---|---|---|---|---|---|---|---|---|---|
| **A** | 0.000 | 0.000 | 0.210 | 0.210 | 0.210 | 0.210 | 0.000 | -0.100 | -0.050 | -0.100 | -0.100 | -0.100 | -0.100 | -0.100 | -0.050 | -0.100 | -0.100 | -0.200 | 0.000 | 0.000 |
| **G** | 0.300 | 0.600 | 0.600 | 0.600 | 0.600 | 0.600 | -0.070 | -0.070 | 0.000 | 0.100 | 0.100 | 0.170 | 0.170 | 0.170 | 0.400 | 0.400 | 0.400 | 0.400 | 0.000 | 0.000 |
| **S** | 0.100 | 0.600 | 0.600 | 0.600 | 0.600 | 0.600 | -0.100 | -0.100 | -0.050 | -0.100 | -0.100 | 0.170 | 0.170 | 0.170 | 0.400 | 0.400 | 0.400 | 0.400 | 0.000 | 0.000 |
| **T** | 0.100 | 0.600 | 0.600 | 0.600 | 0.600 | 0.600 | -0.100 | -0.100 | -0.050 | -0.100 | -0.100 | 0.170 | 0.170 | 0.170 | 0.400 | 0.400 | 0.400 | 0.400 | 0.000 | 0.000 |
| **N** | 0.100 | 0.600 | 0.600 | 0.600 | 0.600 | 0.600 | -0.100 | -0.100 | -0.050 | -0.100 | -0.100 | 0.170 | 0.170 | 0.170 | 0.400 | 0.400 | 0.400 | 0.400 | 0.000 | 0.000 |
| **D** | 0.100 | 0.600 | 0.600 | 0.600 | 0.600 | 0.600 | 0.000 | 0.000 | -0.200 | -0.320 | -0.400 | 0.170 | 0.170 | 0.170 | 0.400 | 0.400 | 0.400 | 0.400 | 0.000 | 0.000 |
| **Q** | -0.000 | 0.000 | 0.000 | 0.000 | 0.000 | -0.350 | 0.000 | -0.100 | -0.150 | -0.250 | -0.300 | -0.260 | -0.260 | -0.260 | -0.050 | -0.050 | -0.050 | -0.050 | 0.000 | 0.000 |
| **E** | -0.000 | 0.020 | 0.000 | 0.000 | 0.000 | 0.000 | 0.000 | 0.000 | -0.300 | -0.420 | -0.500 | -0.100 | -0.100 | -0.100 | 0.000 | 0.000 | 0.000 | 0.000 | 0.000 | 0.000 |
| **H** | 0.000 | 0.000 | 0.000 | 0.000 | 0.000 | 0.000 | 0.000 | 0.100 | 0.100 | 0.150 | 0.150 | -0.250 | -0.200 | -0.300 | -0.107 | -0.107 | -0.107 | -0.107 | 0.000 | 0.000 |
| **K** | 0.000 | 0.020 | 0.000 | 0.000 | 0.000 | 0.150 | 0.000 | 0.100 | 0.200 | 0.300 | 0.300 | -0.200 | -0.200 | -0.200 | -0.150 | -0.150 | -0.150 | -0.150 | 0.000 | 0.000 |
| **R** | 0.000 | 0.020 | 0.000 | 0.000 | 0.000 | 0.150 | 0.000 | 0.100 | 0.200 | 0.300 | 0.300 | -0.200 | -0.200 | -0.200 | -0.150 | -0.150 | -0.150 | -0.150 | 0.000 | 0.000 |
| **Y** | 0.000 | 0.320 | 0.320 | 0.320 | 0.320 | 0.320 | 0.000 | 0.000 | -0.850 | 0.000 | 0.000 | -0.500 | -0.500 | -0.500 | -0.300 | -0.300 | -0.300 | -0.300 | 0.000 | 0.000 |
| **F** | 0.000 | 0.320 | 0.320 | 0.320 | 0.320 | 0.320 | 0.000 | 0.000 | -0.800 | 0.000 | 0.000 | -0.500 | -0.500 | -0.500 | -0.300 | -0.300 | -0.300 | -0.300 | 0.000 | 0.000 |
| **W** | 0.000 | 0.320 | 0.320 | 0.320 | 0.320 | 0.320 | 0.000 | 0.000 | -0.900 | 0.000 | 0.000 | -0.500 | -0.500 | -0.500 | -0.300 | -0.300 | -0.300 | -0.300 | 0.000 | 0.000 |
| **V** | 0.100 | 0.000 | 1.100 | 1.100 | 1.100 | 1.100 | -0.080 | -0.080 | -0.100 | -0.080 | -0.080 | 0.000 | 0.000 | 0.000 | -0.450 | -0.450 | -0.450 | -0.450 | 0.000 | 0.000 |
| **I** | 0.100 | 0.000 | 1.100 | 1.100 | 1.100 | 1.100 | -0.080 | -0.080 | -0.100 | -0.080 | -0.080 | 0.000 | 0.000 | 0.000 | -0.450 | -0.450 | -0.450 | -0.450 | 0.000 | 0.000 |
| **M** | 0.100 | 0.000 | 1.100 | 1.100 | 1.100 | 1.100 | -0.080 | -0.080 | -0.100 | -0.080 | -0.080 | 0.000 | 0.000 | 0.000 | -0.450 | -0.450 | -0.450 | -0.450 | 0.000 | 0.000 |
| **L** | 0.100 | 0.000 | 1.100 | 1.100 | 1.100 | 1.100 | -0.080 | -0.080 | -0.100 | -0.080 | -0.080 | 0.000 | 0.000 | 0.000 | -0.450 | -0.450 | -0.450 | -0.450 | 0.000 | 0.000 |
| **C** | 0.000 | 0.000 | 0.000 | 0.000 | 0.000 | 0.000 | 0.000 | 0.000 | 0.000 | 0.000 | 0.000 | 0.000 | 0.000 | 0.000 | 0.000 | 0.000 | 0.000 | 0.000 | 0.000 | 0.000 |
| **P** | 0.000 | 0.000 | 0.000 | 0.000 | 0.000 | 0.000 | 0.000 | 0.000 | 0.000 | 0.000 | 0.000 | 0.000 | 0.000 | 0.000 | 0.000 | 0.000 | 0.000 | 0.000 | 0.000 | 0.000 |

Source DOI: 10.1038/nsb0694-399

Code/core/resources/helix_sd_i_4.txt

Commit: d6ccec1920f256a393b15215df63fd2dead61cd5

SHA-256 of resource: 492bb545cc304bd51a417ad7ffa4e9cac6babf8cf4b256a828406551ccab4897
