## Additional file 3. Figures and captions for "Helix Design Studio: designing antimicrobial peptide candidates for membrane environments": figure_1_concept.pdf

### a Choose a starting point

**ANALYZE**

**Existing sequence**

K L L K K L L K K L K K

or

**DESIGN**

**New sequence**

Length + allowed amino acids

### b Define the environment

Polar | **Interface** | Apolar

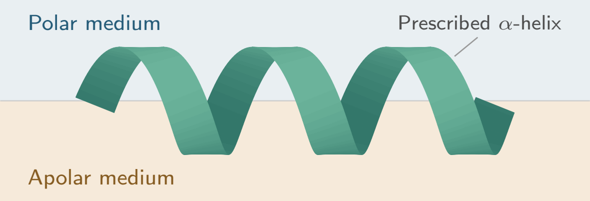


Interface charge (−/0/+) and angular exposure

**Contributions to the score**

- Helix propensity
- Residue interactions
- Charge interactions
- Hydrophobic exposure

### c Explore the results

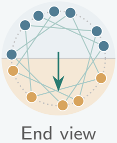


**ANALYZE**

**Inspect and compare**

- Scores and contributions
- Helical wheel and descriptors
- Compare environments
- Explore exposure

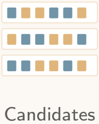


**DESIGN**

**Candidates and families**

- One environment
- Target over alternatives
- Two-environment compromise

Export sequences, scores, settings and an ideal helix (PDB)
