## Additional file 3. Figures and captions for "Helix Design Studio: designing antimicrobial peptide candidates for membrane environments": figure_3_profiles.pdf

| | Polar | Apolar | Anionic | Neutral | Cationic |
|---|---|---|---|---|---|
| P1: L/K pattern | 3.35 | 2.90 | -6.54 | -4.75 | -3.83 |
| P2: poly-Leu | 1.90 | -6.44 | -1.34 | -2.13 | -1.48 |
| P3: L/E pattern | 3.86 | 2.30 | -2.99 | -4.07 | -6.25 |
| F1 | -0.52 | 1.45 | -8.88 | -7.13 | -5.96 |
| F2 | -0.35 | 1.61 | -8.87 | -7.12 | -5.95 |
| F3 | -0.31 | 2.89 | -8.85 | -6.71 | -5.58 |
| F4 | -0.23 | 0.81 | -8.53 | -7.11 | -5.94 |
| S1 | -0.62 | 2.45 | -9.08 | -6.98 | -5.83 |
| S2 | -0.68 | 1.86 | -8.56 | -6.76 | -5.62 |
| C1 | -9.53 | -1.58 | -7.72 | -6.94 | -6.79 |

Cell values: $Z_H$; blue = lower, red = higher. Interfaces: anionic, neutral, cationic.
